# ‘Targeting’ the Search: An Upgraded Structural and Functional Repository of Antimicrobial Peptides for Biofilm Studies (B-AMP v2.0) with a Focus on Biofilm Protein Targets

**DOI:** 10.1101/2022.08.11.503698

**Authors:** Shashank Ravichandran, Sai Supriya Avatapalli, Yatindrapravanan Narasimhan, Karishma S Kaushik, Ragothaman M Yennamalli

## Abstract

Bacterial biofilms, often as multispecies communities, are recalcitrant to conventional antibiotics, making the treatment of biofilm infections a challenge. There is a push towards developing novel anti-biofilm approaches, such as antimicrobial peptides (AMPs), with activity against specific biofilm targets. In previous work, we developed Biofilm-AMP, a structural and functional repository of AMPs for biofilm studies (B-AMP v1.0) with more than 5000 structural models of AMPs and a vast library of AMP annotations to existing biofilm literature. In this study, we present an upgraded version of B-AMP, with a focus on existing and novel bacterial biofilm targets. B-AMP v2.0 hosts a curated collection of 2502 biofilm protein targets across 473 bacterial species, with structural protein models and functional annotations from PDB, UniProt, and PubMed databases. The biofilm targets can be searched for using the name of the source organism, and function and type of protein, and results include designated Target IDs (unique to B-AMP v2.0), UniProt IDs, 3D predicted protein structures, PDBQT files, pre-defined protein function, and relevant scientific literature. To present an example of the combined applicability of both, the AMP and biofilm target libraries in the repository, we present two case studies. In the first case study, we expand an *in silico* pipeline to evaluate AMPs against a single biofilm target in the multidrug resistant, bacterial pathogen *Corynebacterium striatum*, using 3D protein-peptide docking models from previous work and Molecular Dynamics simulations (∼1.18 µs). In the second case study, we build an *in silico* pipeline to identify candidate AMPs (using AMPs with both anti-Gram positive and anti-Gram negative activity) against two biofilm targets with a common functional classification in *Pseudomonas aeruginosa* and *Staphylococcus aureus*, widely-encountered bacterial co-pathogens. With its enhanced structural and functional capabilities, B-AMP v2.0 serves as a comprehensive resource for AMP investigations related to biofilm studies. B-AMP v2.0 is freely available at https://b-amp.karishmakaushiklab.com and will be regularly updated with structural models of AMPs and biofilm targets, as well as 3D protein-peptide interaction models for key biofilm-forming pathogens.

## Introduction

Biofilms are multicellular aggregates or communities of microorganisms, such as bacteria, encased in an extracellular matrix (Penesyan et al., 2021). In clinical settings, bacterial biofilms, often as multispecies states, are associated with persistent infection states, and exhibit increased tolerance to conventional antibiotics (Høiby et al., 2010a, 2011; Orazi and O’Toole, 2020). This often results in prolonged and repeated antibiotic use, which in turn, fuels the emergence and spread of antibiotic resistance (Olivares et al., 2020; Dostert et al., 2021). Given this, there is a concerted focus on developing alternate and adjunct approaches to treat and prevent biofilm infections (Römling et al., 2014; Wu et al., 2014; Hughes and Webber, 2017).

Antimicrobial peptides (AMPs) are being increasingly recognized for their potential as novel anti-biofilm approaches (Yasir et al., 2018; Galdiero et al., 2019; di Somma et al., 2020; Hancock et al., 2021). A diverse class of naturally-occurring and synthetic peptides, AMPs display activity against specific bacterial targets, and can thereby inhibit or interfere with processes in biofilm development. This is particularly relevant given that biofilm formation occurs via a series of sequential stages, where each event, as well as transitions across events, are coordinated by unique structural and signaling factors (Donlan, 2001). Biofilm formation starts with the initial attachment of bacteria, either as single cells of small aggregates, to biotic or abiotic surfaces. Following this initial transient attachment, bacteria release extracellular matrix factors that lead to irreversible surface attachment. Subsequently, the attached bacteria proliferate to form mature, three-dimensional communities. Mature biofilms can disperse to seed new sites via the release of bacteria as small or large clumps of cells, or single cells. Across different biofilm-forming bacterial species, these stages of biofilm formation are mediated by a range of bacterial proteins, including adhesins, structural proteins, signaling molecules and regulatory elements (Latasa et al., 2006; Fong and Yildiz, 2015; Hobley et al., 2015; Wolska et al., 2015). Therefore, bacterial proteins involved in biofilm formation can serve as ‘biofilm targets’, and AMPs that interfere with, or inhibit the function of these specific targets, could lead to hitherto unexplored AMP-biofilm target combinations.

In our previous work, we developed Biofilm-AMP (B-AMP v1.0), a curated structural and functional repository of AMPs for biofilm studies (Antimicrobial Peptide Repository for Biofilms | B-AMP; Mhade et al., 2021). B-AMP hosts structural models of 5766 AMPs (AMP sequences were obtained from DRAMP v3.0), including filtered-lists of AMPs with known anti-Gram positive and anti-Gram negative activity, and a vast library of 11,865 AMP annotations to existing biofilm literature. The AMP library in B-AMP v1.0 is easily accessible using search-enabled filters, and includes FASTA sequences, 3D predicted structures and relevant literature. As a case study to evaluate the application of B-AMP v1.0 for the *in silico* screening and identification of AMPs with anti-biofilm potential, select 3D structural models of AMPs with known anti-Gram positive activity, were docked with the catalytic site residues of the sortase C protein of the in the multidrug resistant pathogen *Corynebacterium striatum*, known to be important for biofilm formation in the multidrug resistant pathogen (Antimicrobial Peptide Repository for Biofilms | B-AMP; Mhade et al., 2021). Based on interacting residues and docking scores, a preference score was proposed to categorize AMPs for future *in silico* and laboratory evaluation against *C. striatum* biofilms. Taken together, B-AMP v1.0 provides a vast structural and functional repository of AMPs, along with a proposed *in silico* pipeline, that can be used to evaluate candidate AMP interactions with biofilm targets. However, in addition to a library of AMPs, identifying candidate AMP-biofilm target combinations would require a comprehensive curation of potential biofilm targets across a wide-range of pathogens.

While B-AMP v1.0 largely focused on building a library of AMPs for biofilm studies, in this study, we present an upgraded version of B-AMP (B-AMP v2.0), with a focus on existing and novel biofilm targets. B-AMP v2.0 hosts a curated collection of 2502 unique biofilm targets, representing 470 biofilm-forming bacterial species. This includes annotated structural and functional information, with source data obtained from three primary databases, namely PDB, UniProt, and PubMed. In B-AMP v2.0, biofilm targets can be searched for using the function or type of protein, and name of the source organism, with collated results that include Target IDs (unique to B-AMP v2.0), UniProt IDs, 3D structure of the target proteins, PDBQT files, pre-defined protein functions, and relevant scientific literature. To present an example of the combined applicability of both, the AMP and biofilm target libraries in the repository, we present two case studies. In the first case study, we focus on the *in silico* evaluation of AMPs against a single biofilm target in a multidrug resistant bacterial pathogen, using previously built 3D protein-peptide docking models of AMPs with the sortase C protein of *C. striatum*. For this, we use MD simulations (∼1.18 µs) to validate our previously proposed preference scores for AMPs with the sortase C protein (based on docking scores and interacting residues). In the second case study, we focus on two biofilm targets with common functional annotations across the widely-encountered bacterial co-pathogens, *P. aeruginosa* and *S. aureus*. For this, we build an *in silico* pipeline to identify candidate AMPs (using AMPs with both anti-Gram positive and anti-Gram negative activity) against two biofilm targets from both pathogens involved in cell adhesion. Given the comprehensive resources and approaches, B-AMP v2.0 lends itself well for AMP studies relevant to biofilms for a range of bacterial species and biofilm targets, which includes high-throughput and large-scale *in silico* identification and evaluation of candidate AMP-biofilm target combinations.

## Methods

### Updates and analytics related to the AMP library of B-AMP v1.0

As part of updates to B-AMP v1.0, additional AMPs were identified from the DRAMP database v3.0, using previously published methods, and modeled using PEPFOLD3 (Lamiable et al., 2016; Mhade et al., 2021; Shi et al., 2022). The AMP library was updated with new tiles for each additional AMP containing Pep IDs (unique to B-AMP), FASTA sequences, PDB structures and PDBQT files. Further, for each AMP (including newly-added AMPs), relevant biofilm literature sourced from PubMed, was hyperlinked to each tile. The reference sheet for the AMP library, and the PubMed biofilm literature and counts sheets were updated. Analytics on the usage of B-AMP v1.0, including web page views, visits and downloads, location and language of website access was retrieved from GoatCounter (from December 16, 2021 till June 28, 2022) (GoatCounter – open source web analytics). Metrics related to the scientific publication associated with B-AMP v1.0 was downloaded from the journal website (from December 16, 2021 till June 28, 2022) (Mhade et al., 2021).

### Data retrieval and query construction for biofilm targets for B-AMP v2.0

To identify and retrieve data for biofilm protein targets across a range of bacterial species, three primary databases were used, namely, UniProt (for sequence and literature data), PDB (for structural data), and PubMed (for biofilm related literature data). For each database, a query was constructed according to the query syntax of the database using boolean operators. In the construction of the query syntaxes, ‘biofilm formation’ was designated as a mandatory term. Other terms included keywords such as ‘adhesion’, ‘maturation’, ‘dispersal’,‘quorumsensing’, ‘MSCRAMM’,‘matrix’,and ‘autoinducer’, which were used with the OR boolean operator. Using specific query builders in the UniProt, PDB, and PubMed databases, custom queries were used to retrieve the data. Therefore, ‘(Biofilm AND formation) AND ((adhesion OR maturation OR dispersion OR autoinducer OR MSCRAMM OR target OR matrix) OR (quorum AND sensing)) AND bacteria’ was the query used across the databases, modified according to the specific syntax. *In-house* Python API scripts were used to automate the data retrieval from the databases (https://github.com/KarishmaKaushikLab/BAMP-v2-scripts), and data was stored in a Javascript Object Notation (JSON) format.

### Biofilm target data integration for B-AMP v2.0

Data retrieved from all three primary sources were screened for completeness and redundancy by considering the intersection of data across the databases. Taking into account the dogma of the protein world, where sequence dictates function and function, in turn, dictates structure, the UniProt ID for each biofilm target was considered as the primary data. Further, for a given biofilm protein target there can be more than one PDB structure and more than one associated PubMed ID. Given this, we searched for UniProt entries that satisfied the criteria of having at least one entry in PDB or at least one entry in PubMed. Each UniProt ID was mapped onto PDB data and PubMed data, and vice versa. Using this approach, the data retrieved from the three databases were mapped to obtain a complete and nonredundant dataset of biofilm protein targets. The list of UniProt IDs was matched with the UniProt metadata in the PDB structures, which enabled the removal of false-positive entries from PDB. Similarly, the primary citation for each PDB ID was mapped to the list of PubMed metadata. This multi-directional mapping across the three datasets helped identify UniProt IDs without experimentally validated PDB structures and PubMed IDs not mapped in UniProt and PDB. The non-redundant data from the three sources was used to create a masterlist with the following information for each biofilm target: Target ID (unique to B-AMP v2.0), bacterial protein name, organism name and strain, UniProt ID, PDB ID (if available), PubMed ID, classification as per UniProt’s GO (Gene Ontology - Molecular Function) term and classification of function as per PDB.

### Modeling of biofilm protein targets of *P. aeruginosa* and *S. aureus* using RoseTTAFold

For 90 biofilm targets of *P. aeruginosa* and *S. aureus* (that did not have experimentally determined structures), we used RoseTTAFold, to predict the tertiary structure of the protein target (Baek et al., 2021). For this, RoseTTAFold was installed locally on a workstation with 16 Intel(R) Xeon(R) Silver 4208 processors by cloning the repository from GitHub (https://github.com/RosettaCommons/RoseTTAFold). For a given protein sequence, a multiple sequence alignment was constructed, which was followed by residue-residue interaction mapping using Graph-transformer (Yun et al., 2019). Finally, a coordinate space was used to derive multiple conformations, which were refined in an iterative manner to provide highly ranked models as output. Computational models of these 90 biofilm targets are available in B-AMP v2.0, along with additional target information such as bacterial protein name, organism name and strain, UniProt ID, PubMed ID, classification as per UniProt GO term and classification of function as per PDB.

### Validation of 3D protein-peptide docking models of select AMPs with the sortase C protein of *C. striatum* using MD simulations

To validate the molecular docking results of 60 select 3D protein-peptide docking models of AMPs (and the LPMTG pilin motif) with the sortase C protein of *C. striatum*, MD simulations were performed using GROMACS v2020.3 (Berendsen et al., 1995; Abraham et al., 2015; Bauer et al., 2022). For this, the sortase C-AMP complex structure was placed at the center of a cubic box, at a distance of 1 nm from the edge of the box. Water molecules were added in the box using the SPC6 water model (Jorgensen et al., 1998). The generalized all atom OPLS force field was implemented, and each system was neutralized with the appropriate number of Na^+^ and Cl^-^ ions (Jorgensen et al., 1996). The protein-peptide complex along with water and ions was energy minimized using the steepest descent algorithm for 50000 steps or converging at an earlier step (Berendsen et al., 1981). Further, each system was equilibrated in the canonical NVT phase at a constant temperature of 300K for 50 picoseconds (ps) and in the canonical NPT phase with a constant pressure of 1 bar for 50 ps. Finally, the production MD simulation was set to 20 nanoseconds (ns), where a frame or snapshot of velocities and coordinates was stored for every 10 ps. The trajectory obtained was processed by removing the periodic boundary condition effect and jumps, and centered by rotating and translating the complex. The processed trajectory was visualized, and the interactions between each AMP and sortase C was mapped using DIMPLOT (Laskowski and Swindells, 2011).

### *In silico* molecular docking of select AMPs with both anti-Gram positive and anti-Gram negative activity against cell adhesion biofilm targets of *P. aeruginosa* and *S. aureus*

For *P. aeruginosa*, the crystal structure for Target 37 (PDB ID: 5CYL) or the ‘fimbrial subunit CupB6’, was analyzed along with the primary citation, and the polyproline region was identified as the active site. The biologically functional unit is a monomer and a single chain was processed by adding non-polar hydrogens and Kollman charges, and saved as a PDBQT file. Grid maps were generated for a grid box of dimensions 47.25 Å× 47.25 Å× 47.25 Å. For *S. aureus*, the crystal structure for Target 1 (PDB ID: 7C7U) or the ‘biofilm-associated surface protein’, was analyzed along with its primary citation (Ma et al., 2021). The N-terminal lobe of the protein was identified to have a significant role in biofilm formation, and active sites of the protein were visualized to select Gln728 as the center of the gridbox (Ma et al., 2021). The biologically functional unit was then processed by adding non-polar hydrogens and Kollman charges, and saved as a PDBQT file. Grid maps were generated for a grid box of dimensions 47.25 Å X 47.25 ÅX 47.25 Å. A filtered list of AMPs with both anti-Gram positive and anti-Gram negative activity was prepared using the B-AMP library (Mhade et al., 2021). For protein-peptide *in silico* molecular docking, Autodock 4.2 was used, with each AMP processed by adding hydrogens and Gasteiger charges, and setting the torsions to a maximum of 32 with most atoms (saved as PDBQT files). For both Target ID 1 from *S. aureus* and Target ID 37 from *P. aeruginosa*, each AMP was docked to the protein target using a Lamarckian genetic algorithm. Following this, the protein-peptide docked complexes were analyzed using *in house* python scripts. Based on binding energy scores, candidate AMPs were identified for their ability to interact with both targets.

### Building the user interface of B-AMP v2.0 to host biofilm targets and 3D protein-peptide docking models

In the upgraded version of B-AMP, a new section ‘Biofilm targets’ was created to house the structural and functional library of biofilm targets. For each biofilm target, the assigned Target ID was considered as the primary identifier. Relevant information related to each Target ID was extracted as individual JSON files. Using these, JavaScript codes were executed to create tiles for each Target ID, that can be searched for using a range of queries such as biofilm protein target name, name of the source organism, and function of the protein target based on PDB or UniProt GO annotations. For a given query, filters in the form of two additional drop-down menus were designed with organism name and classification of function as per PDB. Further, each biofilm target tile was color coded using a color legend based on functional classification of the target function as per PDB. Each tile displays the Target ID, protein name, organism name and strain, and classification of function as per PDB. The target protein structure, if available, is displayed as a static image, and the PDB coordinates are available for download. RoseTTAFold models are also available for download as hyperlinks in the tile. Each tile also contains hyperlinks to UniProt and PubMed databases. For the additional 3D protein-peptide docking models, a new section ‘AMPs docked to dual biofilm targets’ was created, and *in silico* predicted models for candidate AMPs with the cell adhesion biofilm targets of *P. aeruginosa* and *S. aureus* are displayed. Each model tile displays PDBQT IN/OUT files, the downloadable 3D model, and bond information.

### Availability of scripts used to upgrade B-AMP v2.0

The in-house scripts used to build the structural and functional repository of biofilm targets have been added to B-AMP (https://b-amp.karishmakaushiklab.com/code.html).

## Results

### B-AMP v1.0 as a resource for AMP studies relevant to biofilms

In previous work, we published Biofilm-AMP (B-AMP v1.0), a curated structural and functional repository of AMPs for biofilm studies (Antimicrobial Peptide Repository for Biofilms | B-AMP; Mhade et al., 2021). In the original version, B-AMP hosted 5544 3D predicted structural models of AMPs (AMP sequences from the DRAMP v3.0 database), with 613 AMP annotated to 11,611 biofilm literature references. For each AMP, the results include downloadable FASTA files, PDB files, images of predicted and chosen 3D models, and PubMed links to relevant literature. In addition, for specific-user case scenarios, filtered lists of 2534 AMPs with known anti-Gram positive activity and 2389 AMPs with known anti-Gram negative activity are also available. To demonstrate the applicability of the AMP structural library in B-AMP, 100 select AMPs with known anti-Gram positive activity were subject to *in silico* molecular docking (using AutoDock Vina) with the semi-open lid conformation of the sortase C protein of *C. striatum*. This includes 88 AMPs with a length of 2-8 amino acids and 12 representative AMPs each ranging from 9-20 amino acids in length. The 3D predicted protein-peptide interaction models, including an *in house* built preference score based on interacting residues and docking scores, are also made available in B-AMP v1.0.

From December 15, 2021 (B-AMP v1.0 was published on December 16, 2021) to June 28, 2022, B-AMP v1.0 received 1328 page views and 1106 visits, which resulted in ∼185 page views per month (website analytics retrieved from GoatCounter) (GoatCounter – open source web analytics). The highest number of page views were on the home page (482/1328), followed by the AMP library (219/1328), AMPs docked to biofilm targets (105/1328), code (87/1328), Anti-Gram positive AMP library (79/1328), Anti-Gram negative AMP library (69/1328), and references (36/1328) **(Figure 1A)**. The remaining B-AMP v1.0 engagements (321/1328) were with the AMP and 3D protein-peptide docking models, and other features such as GitHub pages and API scripts. Based on the data retrieved, 66% of visits accessed B-AMP v1.0 from India (729/1106), 17% from the USA (188/1106), 14% from China (153/1106), with the remaining 3% from Mexico, Germany and Malaysia, Hong Kong, Canada, Ecuador, France, Switzerland, Italy, Kazakhstan, Philippines, United Kingdom (36/1106) **(Figure 1B)**. Further, 85% (939/1106) of visits were in English, followed by 14% (154/1106) in Chinese, and a smaller percentage (∼1%) in Portuguese, Kazakh and French (13/1106) **(Figure 1C;** webpage translation via Google). In addition to the B-AMP v1.0 website, the related published work has received a total of 2074 views (from December 16, 2021 to June 15, 2022) **(Figure 1D**, month wise data from Frontiers from December, 2021 to June, 2022, accessed on June 28, 2022).

**Figure 1:**
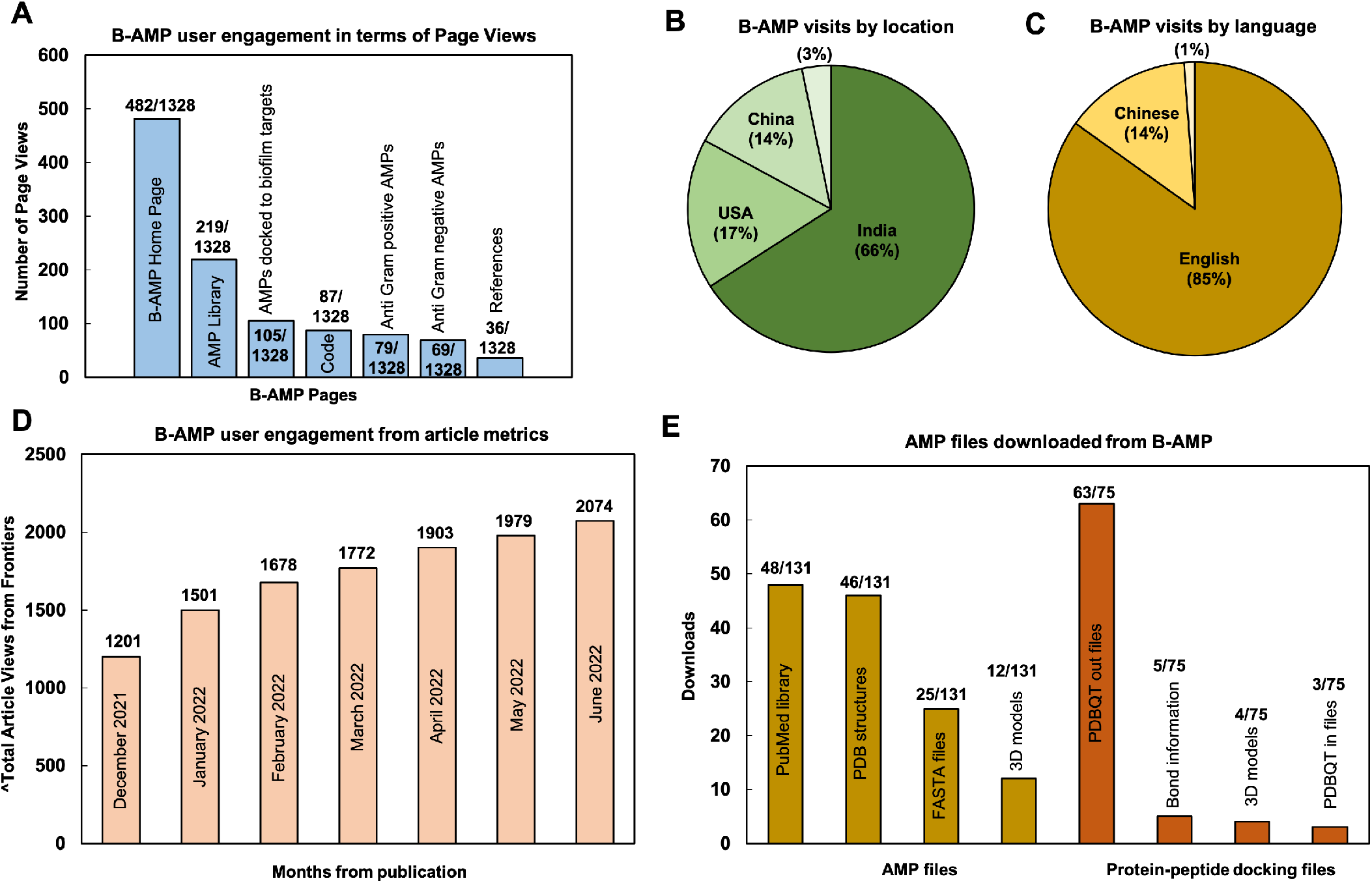
User metrics of B-AMP v1.0 as a resource for AMP studies relevant to biofilms. Data reflects user engagement from December 16, 2021 till June 28, 2022; in this period B-AMP received 1328 page views and 1106 visits **(A)** Since publication of the repository, B-AMP v1.0 has received a total of 1328 user visits, of which the highest page views were the home page (482/1328), followed by the AMP library (219/1328) and AMPs docked to biofilm targets (105/1328). The remaining B-AMP v1.0 engagements were with the code, Anti-Gram positive and Anti-Gram negative AMPs, and other features such as GitHub pages and API scripts. **(B)** Based on location, 66% (729/1106) of B-AMP visits were from India, followed by 17% (188/1106) from the USA and 14% (153/1106) from China. **(C)** Based on language, 85% (939/1106) of B-AMP visits were in English, followed by 14% (154/1106) in Chinese, and smaller percentage (∼1%) in Portuguese, Kazakh and French (13/1106) (webpage translation via Google). **(D)** B-AMP v1.0 Frontiers article views from December 16, 2021 to June 28, 2022, showing a steady increase in total article views from publication (month wise data from Frontiers analytics). **(E)** The 148 different AMPs and protein-peptide models with user engagement included 78 AMPs and 70 protein-peptide docking models, with 131 and 75 user downloads respectively. For the 78 AMPs, highest downloads were for the PubMed links (48/131), followed by PDB structures (46/131), FASTA files (25/131) and 3D models (12/131). For the 70 protein-peptide docking models, highest downloads were for the PDBQT out files (63/75), followed by bond information (5/75), 3D models (4/75) and PDBQT in files (3/75). Data shows the highest downloads for AMPs and protein-peptide docking models.

Additionally, the manuscript has received 2 citations, and the pre-print has received 1532 abstract views, 147 full-text HTML views and 401 PDF downloads (from August 18, 2021 to June 28, 2022) (Antunes et al., 2022; El-Omar et al., 2022). Among the user engagements with the AMP libraries on B-AMP v1.0, for 148 different AMPs (78 AMPs and 70 protein-peptide docking models) there were 206 downloads (131 for AMPs and 75 for protein-peptide docking models) for related information, including the provided FASTA files, PDB structures, 3D models, PDBQT files or PubMed links. Of this, for 78 AMPs, highest downloads were for the PubMed links, followed by PDB structures, FASTA files and 3D models **(Figure 1E)**. For the 70 protein-peptide docking models, highest downloads were for the PDBQT out files, followed by bond information, 3D models and PDBQT IN files **(Figure 1E)**.

Since the first version, we have updated B-AMP with 220 additional AMP structural models, based on updated sequences in the DRAMP v3.0 database. These additional AMPs are reflected in the master list of AMPs, as well as in the biofilm literature annotations (https://b-amp.karishmakaushiklab.com/all.html). Given this, the AMP library in B-AMP now hosts a total of 5766 AMP structural models with 11,865 annotations (from 622 AMPs) to current biofilm literature. Taken together, B-AMP v1.0 presents a comprehensive community resource of AMPs for biofilm studies, with structural and functional resources, that can be leveraged to identify and evaluate AMP interactions with potential biofilm targets.

### Curation of biofilm protein targets for B-AMP v2.0

In addition to a library of AMPs, identifying candidate AMP-biofilm target combinations would require a large curation of potential biofilm targets across a wide-range of pathogens. Therefore, while B-AMP v1.0 largely focused on building a library of AMPs for biofilm studies, in this study, we focused on building an upgraded version of B-AMP (B-AMP v2.0) with existing and novel biofilm protein targets. To build an upgraded version of B-AMP with a biofilm target repository, data on protein targets relevant to biofilm formation was retrieved from three major databases, namely UniProt (for sequence and literature data), PDB (for structural data), and PubMed (for biofilm related literature data) using specific queries **(Figure 2A)**. The retrieved data revealed 3095 targets from PDB, 10066 from UniProt and 12726 from PubMed. Between the UniProt and PDB data, there were 59 UniProt IDs that are shared between the two datasets. In other words, from the 3095 targets obtained from PDB database for the customized query, 5146 UniProt IDs were retrieved. A higher number indicates presence of multiple chains or oligomeric proteins in PDB data. Next, we mapped the list of 5146 PDB IDs (2063 Uniprot IDs) to the 10066 targets obtained directly from the UniProt database, followed by similar mapping of the overlap between the other datasets. The table does not reflect PubMed’s overlap between UniProt and PDB, since the data from PubMed is unidirectional, PubMed IDs can be derived from UniProt and/or PDB databases, but not vice-versa. Using the protein world dogma of sequence dictating function and function dictating the structure, UniProt IDs were used as the primary set of data. Using UniProt IDs as the starting point, we searched for UniProt entries that satisfy the criteria of having at least one entry in PDB or at least one entry in PubMed. This resulted in a final number of 2502 of targets with two important criteria, the biofilm target must have a UniProt ID and must have at least one PDB ID OR at least one PubMed entry (**Table 1**).

**Table 1:** Mapping of biofilm targets across UniProt (for sequence and literature data), PDB (for structural data), and PubMed (for biofilm related literature data) databases.

| Database | Total number of hits | Non-redundant data in each dataset | Filtering Criteria |  |
| --- | --- | --- | --- | --- |
| Uniprot | 10066 | 2063 | Having at least one PDB entry | Having at least one PubMed entry |
|  |  |  | 102 | 139 |
| PDB | 3095 | 102 | Having at least one UniProt ID | Having a PubMed ID |
|  |  |  | 5146 | 2978 |
| PubMed | 12726 | 1437 | - | - |

**Figure 2:**
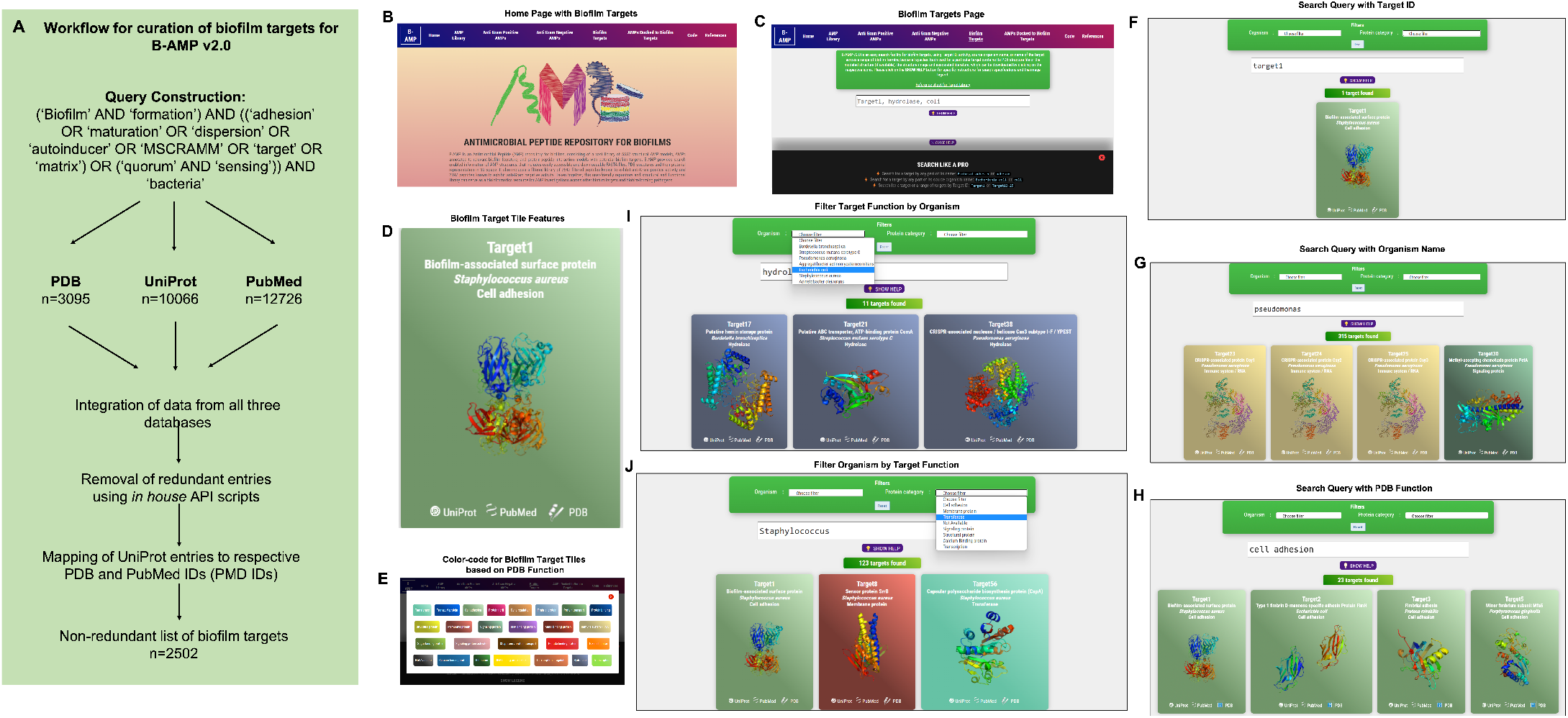
Workflow and features of B-AMP v2.0. **(A)** A specific query, using mandatory and optional keywords relevant to biofilm targets, was constructed and applied across three primary databases, UniProt, PDB and PubMed, following which retrieved data was compiled and mapped to produce a non-redundant list of 2502 protein targets relevant to biofilm formation. **(B)** The B-AMP home page shows a new tab for biofilm targets. **(C)** The newly added biofilm targets page shows a search bar and instructions to search for biofilm targets. **(D)** The search query populates results on biofilm targets in color-coded tiles, based on their PDB function. Each tile contains information such as Target IDs, name of source organism, FASTA files, PDB or UniProt GO functions, PDBQT files, and PubMed links to relevant literature. Further, if the 3D predicted structure of the protein is available on PDB, the role also contains the downloadable PDB structure. If the protein target was modeled using RoseTTAFold, the tile also contains the 3D predicted protein structure. **(E)** The ‘legend’ tab below the search bar provides the legend for the color-coded tiles. **(F-H)** Biofilm targets can be searched for using Target IDs (for example, Target19), name of the source organism (for example, *Pseudomonas aeruginosa*) or function of the protein target based on PDB annotation (for example, cell adhesion). **(I-J)** For search queries related to organism name and PDB target function, the results can be further filtered using the ‘filter by organism’ or ‘filter by function’ feature in the drop-down menu.

The 2502 non-redundant biofilm protein targets are compiled in a master list **(Suppl File 1)**, with the following information for each biofilm target: Target ID (unique to B-AMP v2.0), bacterial protein name, organism name, strain, UniProt ID, PDB ID, PubMed ID, classification as per UniProt GO (Molecular Function) term, and classification of function as per PDB.

### Features of B-AMP v2.0 as a structural and functional repository of biofilm protein targets

In the upgraded version, B-AMP v2.0 (https://b-amp.karishmakaushiklab.com/) features a new tab ‘Biofilm targets’ that houses the structural and functional library of biofilm protein targets **(Figure 2B)**. In this section, biofilm targets can be searched for using Target IDs (e.g: Target19), or name of the source organism (e.g: *Pseudomonas aeruginosa*), or function of the protein target in biofilm formation based on PDB annotation (e.g: cell adhesion). The ‘show help’ pop-up tab below the search bar provides information on the supported search format and queries **(Figure 2C)**. For all search formats, the search query populates results on biofilm targets in color-coded tiles, based on their PDB function. Each tile contains information such as Target IDs, name of source organism, FASTA files, PDB or UniProt GO functions, PDBQT files, and PubMed links to relevant literature. Further, if the 3D predicted structure of the protein is available on PDB, the tile also contains the downloadable PDB structure. If the protein target was modeled using RoseTTAFold, as for 90 target proteins from *P. aeruginosa* and *S. aureus*, the tile contains the 3D modeled structure **(Figure 2D)**. Finally, if the biofilm target structure is available, the thumbnail image of the structure is provided on the face of the tile. The ‘legend’ tab below the search bar provides the legend for the color-coded tiles **(Figure 2E)**. Using the Target ID, a search query can be used to obtain results containing the protein name with the PDB ID, UniProt ID, GO function and relevant literature. For example, typing Target ID 1 (Target1) in the search bar **(Figure 2F)**, displays a single tile corresponding to the unique biofilm target, with corresponding information such as biofilm-associated surface protein (target name), *S. aureus* (organism name), cell adhesion (target function), along with links to UniProt, PubMed and PDB, and the 3D protein structure on the face of the tile. Along these lines, specific to the biofilm-forming bacterial species under investigation, the organism name (genus or species) can be used as a search query **(Figure 2G)**. For example, the search query ‘*Pseudomonas*’ populates target tiles belonging to the specific genus, which includes target tiles from across different *Pseudomonas* species. Further, biofilm targets can also be searched for using function, based on the PDB annotation of protein function **(Figure 2H)**. For example, using ‘cell adhesion’ as a search query provides target tiles from a range of different organisms, with information on Target ID, organism name, protein name and 3D protein structure (if available). For search queries related to organism name and PDB function, the results can be further filtered using the ‘filter by organism’ or ‘filter by function’ feature in the drop-down menu. For example, the search query ‘hydrolase’ can be further filtered by organisms to find ‘hydrolases’ specific for ‘*Escherichia coli*’ **(Figure 2I)**. Similarly, the search query ‘*Staphylococcus aureus*’ can be filtered for biofilm targets with specified functions such as ‘transferases’ (**Figure 2J)**. If the 3D protein structure for a particular target is not available (from PDB or among the RoseTTAFold models), the face of the tile reads ‘structure not available’.

### Diversity and distribution of biofilm targets in B-AMP v2.0

The biofilm protein targets in B-AMP v2.0 show diverse distribution in terms of source organism, target function and 3D protein structural models.

### Biofilm target distribution based on source organism

Based on the source organism, the 2502 biofilm targets belong to 227 different bacterial class, family, genera or descriptions (as per NCBI Taxonomy), of which the ten most represented genera are *Pseudomonas* (315 biofilm targets), *Escherichia* (259 biofilm targets), *Salmonella* (216 biofilm targets), *Acinetobacter* (160 biofilm targets), *Staphylococcus* (123 biofilm targets), *Stenotrophomonas* (79 biofilm targets), *Klebsiella* (70 biofilm targets), *Yersinia* (69 biofilm targets), *Xanthomonas* (49 biofilm targets) and *Enterobacter* (48 biofilm targets) **(Figure 3A, Suppl File 2)** (Taxonomy | UniProt help | UniProt). The bacterial genera include 473 different bacterial species or species descriptions (as per NCBI Taxonomy), of which the highest number of biofilm targets are from *Escherichia coli*, with a total of 249 biofilm targets, followed by *Salmonella enterica* with 118 biofilm targets and *S. aureus* with 81 biofilm targets **(Figure 3B, Suppl File 3)** (Taxonomy | UniProt help | UniProt). This is followed by *Acinetobacter baumanii* (72 biofilm targets), *Klebsiella pneumoniae* (64 biofilm targets), *Salmonella typhimurium* (58 biofilm targets), *Pseudomonas fluorescens* (51 biofilm targets), *Pseudomonas aeruginosa* (45 biofilm targets), *Yersinia enterocolitica* (38 biofilm targets) and *Stenotrophomonas maltophilia* (31 biofilm targets). Of the 473 bacterial species with biofilm targets in B-AMP v2.0, 368 are Gram-negative bacterial species and 86 are Gram-positive bacterial species; the remaining 19 species are either Gram-variable or with an uncharacterized Gram stain reaction **(Figure 3C)**. Importantly, this includes bacterial pathogens widely associated with biofilm infections in humans such as *P. aeruginosa, S. aureus, S. epidermidis, E. coli, B. cenocepacia, S. pneumoniae, A. baumanii, V. cholerae*, as well as multidrug resistant bacterial species, such as *Enterobacter* spp, *Citrobacter* spp, *Klebsiella* spp, *Eikenella* spp, *Serratia* spp, and *Stenotrophomonas* spp (Hall-Stoodley and Stoodley, 2005; van Duin and Paterson, 2016; Pérez-Rodríguez and Mercanoglu Taban, 2019; Vestby et al., 2020; Schulze et al., 2021).

**Figure 3:**
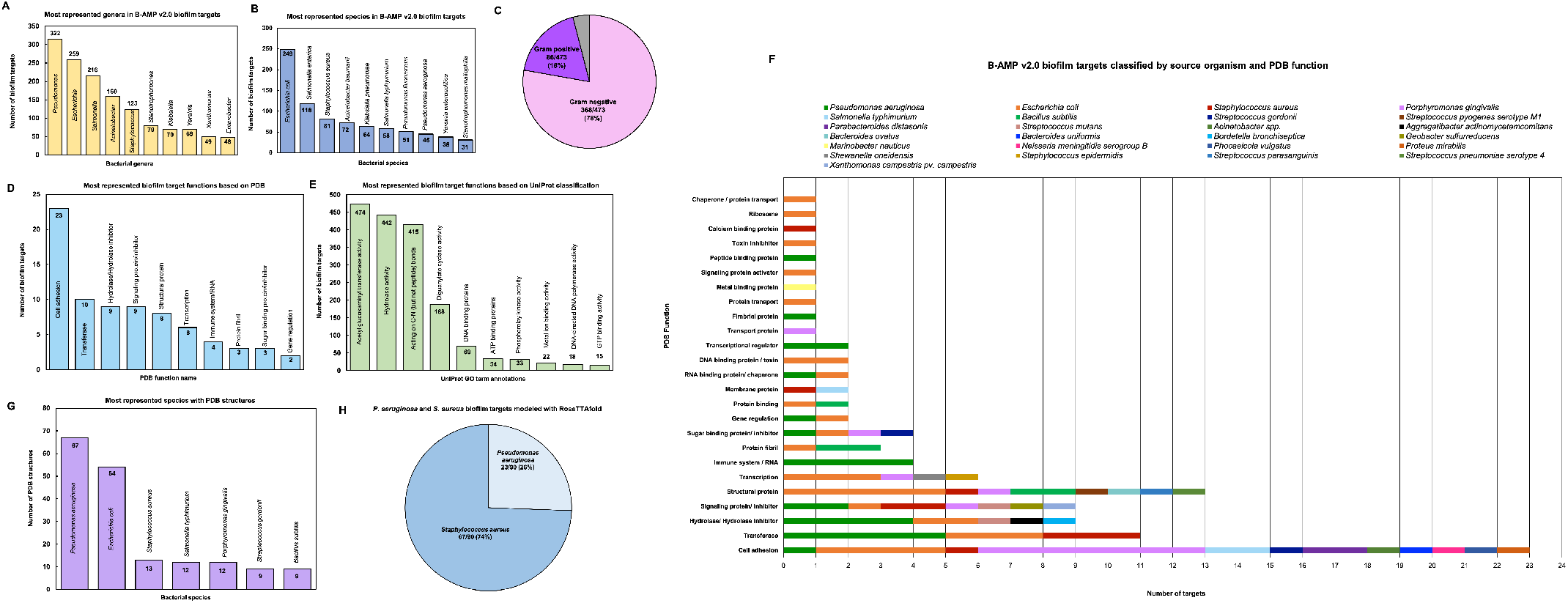
Diversity and distribution of biofilm targets in B-AMP v2.0. **(A)** In the 2502 non-redundant biofilm targets in B-AMP v2.0, the ten most represented genera are *Pseudomonas* (315 biofilm targets), *Escherichia* (259 biofilm targets), *Salmonella* (216 biofilm targets), *Acinetobacter* (160 biofilm targets), *Staphylococcus* (123 biofilm targets), *Stenotrophomonas* (79 biofilm targets), *Klebsiella* (70 biofilm targets), *Yersinia* (69 biofilm targets), *Xanthomonas* (49 biofilm targets) and *Enterobacter* (48 biofilm targets) **(B)** Of the total biofilm targets, the ten most represented species include *Escherichia coli* (249 biofilm targets), *Salmonella enterica* (118 biofilm targets), *S. aureus* (81 biofilm targets), *Acinetobacter baumanii* (72 biofilm targets), *Klebsiella pneumoniae* (64 biofilm targets), *Salmonella typhimurium* (58 biofilm targets), *Pseudomonas fluorescens* (51 biofilm targets), *Pseudomonas aeruginosa* (45 biofilm targets), *Yersinia enterocolitica* (38 biofilm targets) and *Stenotrophomonas maltophilia* (31 biofilm targets). **(C)** The bacterial species represented in the biofilm targets in B-AMP v2.0 include 368 Gram-negative bacterial species and 86 Gram-positive bacterial species, with 19 species that are either Gram-variable or with an uncharacterized Gram stain reaction. **(D)** Of the 98 biofilm targets in PDB with defined function, the largest group of targets belong to the ‘cell adhesion’ category (23 biofilm targets), which represents proteins involved in adhesion of the bacterial cell to host or abiotic surfaces. This is followed by the ‘transferase’ category (10 biofilm targets), ‘hydrolase/ hydrolase inhibitor’ (9 biofilm targets), ‘signaling protein/inhibitor’ (9 biofilm targets), ‘structural protein’ (8 biofilm targets), ‘transcription’ category (6 biofilm targets), ‘immune system/RNA’ (4 biofilm targets), ‘protein fibril’ (3 biofilm targets), ‘sugar binding protein/ inhibitor’ (3 biofilm targets) and ‘gene regulation’ (2 biofilm targets). **(E)** Of the 1321 biofilm targets with annotation of function in UniProt, 474 targets are annotated with ‘acetyl glucosaminyl transferase activity’, 442 targets with ‘hydrolase activity’, 415 targets described as ‘acting on carbon-nitrogen (but not peptide) bonds’, 188 targets with ‘diguanylate cyclase’ activity, 69 targets described as ‘DNA binding’ proteins, 34 targets described as ‘ATP binding’ proteins, 33 targets with ‘phosphorelay kinase activity’, 22 targets with ‘metal ion binding’ activity, 18 targets with ‘DNA-directed DNA polymerase’ activity, and 15 targets with ‘GTP binding’ activity. **(F)** Of the total biofilm targets in B-AMP v2.0, the predicted 3D coordinates for 102 targets (Target IDs 1-102) are available in PDB, with the highest number of PDB structures from *P. aeruginosa* (67 PDB structures), followed by *E. coli* (54 PDB structures), *S. aureus* (13 PDB structures), *S. typhimurium* and *P. gingivalis* (12 PDB structures each), and *S. gordonii* and *B. subtilis* (9 PDB structures each). **(G)** In addition to the biofilm target structures available in PDB, 90 biofilm target structures from *P. aeruginosa* and *S. aureus* were modeled using RoseTTAFold, which includes 23 models for *P. aeruginosa* and 67 models for *S. aureus*. **(H)** For the 98 biofilm targets with PDB annotation, classification by source organism reveals a range of bacterial species represented across the targets, notably for the biofilm targets related to cell adhesion, transferase activity, hydrolase/hydrolase inhibitor activity, signaling proteins/inhibitor activity, structural proteins, transcription proteins and sugar binding proteins/inhibitor activity.

### Biofilm target distribution based on protein function (from PDB and UniProt)

For each of the 2502 biofilm targets, the functional annotation was obtained from PDB and UniProt (RCSB PDB: Homepage; UniProt). In PDB, the classification by protein function is based on the type of the protein, and can be obtained from the ‘struct_keywords’ data category in the PDBx/mmCIF file (Data Category struct_keywords). Based on annotation of function in PDB, out of 2502 biofilm targets, 2404 are designated with ‘no GO function’. Of the 98 biofilm targets with defined function, the largest group of targets belong to the ‘cell adhesion’ category (23 biofilm targets), which represents proteins involved in adhesion of the bacterial cell to biotic or abiotic surfaces. This is followed by the ‘transferase’ category (10 biofilm targets), ‘hydrolase/ hydrolase inhibitor’ (9 biofilm targets), ‘signaling protein/inhibitor’ (9 biofilm targets), ‘structural protein’ (8 biofilm targets), ‘transcription’ category (6 biofilm targets), ‘immune system/RNA’ (4 biofilm targets), ‘protein fibril’ (3 biofilm targets), ‘sugar binding protein/ inhibitor’ (3 biofilm targets) and ‘gene regulation’ (2 biofilm targets) **(Figure 3D)**. The full list of PDB target functions is provided in **Suppl File 4**.

In UniProt, the protein target function is classified based on GO term annotations. Given this, there can be more than one term to describe a protein function, with a single target having multiple descriptions of function. Based on annotation of function in UniProt, out of 2502 biofilm targets, 1181 are designated with ‘no GO function’ (UniProt). The remaining 1321 targets were described across 131 GO terms, with 474 targets annotated with ‘acetyl glucosaminyl transferase activity’, 442 targets with annotated ‘hydrolase activity’, 415 targets annotated as ‘acting on carbon-nitrogen (but not peptide) bonds’, 188 targets annotated with ‘diguanylate cyclase’ activity, 69 targets annotated as ‘DNA binding’ proteins, 34 targets annotated as ‘ATP binding’ proteins, 33 targets annotated with ‘phosphorelay kinase activity’, 22 targets annotated with ‘metal ion binding’ activity, 18 targets annotated with ‘DNA-directed DNA polymerase’ activity, and 15 targets annotated with ‘GTP binding’ activity. **(Figure 3E)** The full list of UniProt target functions is provided in **Suppl File 5**.

When the 98 biofilm targets in B-AMP v2.0 with PDB annotations are classified by source organism, the data reveals a range of bacterial species represented representing several target functions **(Figure 3F)**. For the most represented target function ‘cell adhesion’, biofilm targets belong to 10 different bacterial species, including common biofilm-forming pathogens such as *E. coli, P. gingivalis, S. typhimurium, S. gordonii, Acinetobacter* spp, and *P. mirabilis*, as well as widely-encountered biofilm-forming co-pathogens *P. aeruginosa* and *S. aureus*. A large segment of the biofilm targets annotated with the function ‘cell adhesion’ include bacterial surface proteins, such as fimbrial subunit proteins and adhesins. Similarly, for the other target functions, biofilm targets represent a wide range of pathogens and bacterial proteins. The full list of PDB functions for the biofilm targets with source organisms and bacterial proteins is listed in **Suppl File 6**.

### Biofilm target distribution based on 3D structural models

In B-AMP v2.0, for 102 biofilm targets (out of 2502, for Target IDs 1-102) the experimentally determined 3D coordinates are available in the PDB. Considering the possibility of multiple structures for the same sequence and oligomeric nature of the structure (in some cases), this results in 198 PDB IDs across 25 bacterial species. The highest number of PDB structures are from *P. aeruginosa* (67 PDB structures), followed by *E. coli* (54 PDB structures), *S. aureus* (13 PDB structures), *S. typhimurium* and *P. gingivalis* (12 PDB structures each), and *S. gordonii* and *B. subtilis* (9 PDB structures each) **(Figure 3G)**. The full list of bacterial species with targets with PDB structures is provided in **Suppl File 7**. Further, the PDB IDs for each of these biofilm targets have been provided in the master list of targets **(Suppl File 1)** and in the B-AMP v2.0 user-interface, the tiles corresponding to these 102 targets have been linked to the respective PDB structures.

In addition to the biofilm target structures available in PDB, 90 biofilm target structures from *P. aeruginosa* and *S. aureus* were modeled using RoseTTAFold, which includes 23 models for *P. aeruginosa*, and 67 models for *S. aureus* **(Figure 3H)**. *P. aeruginosa* and *S. aureus* are widely-encountered biofilm-forming pathogens, and both species are often observed together (as co-pathogens) in several clinical infection states (Alves et al., 2018; Yung et al., 2021). Given this, and the increasing trend in resistance to conventional antibiotics demonstrated by these two pathogens, identifying AMP-biofilm target combinations for *P. aeruginosa* and *S. aureus* infections holds significance (Lister and Horswill, 2014; Bhattacharya et al., 2015; Ciofu and Tolker-Nielsen, 2019; Sindeldecker and Stoodley, 2021). The full list of *P. aeruginosa* and *S. aureus* biofilm targets modeled using RoseTTAFold with the Target IDs, UniProt IDs and functional details are provided in **Suppl Files 8 and 9**. Further, in the B-AMP v2.0 user-interface, the tiles corresponding to these 90 targets have links to the respective RoseTTAFold predicted structures.

### A case study of using B-AMP for the *in silico* evaluation of AMPs against a single biofilm target in a multidrug resistant bacterial pathogen

*C. striatum* is a multidrug resistant, biofilm-forming, bacterial pathogen, increasingly associated with a range of wound, eye, skin and ear infections (de Souza et al., 2015; McMullen et al., 2017; Datta et al., 2021; Mhade et al., 2021). In *C. striatum*, the sortase-pilin machinery encodes the pilus-specific sortase C enzyme, known to be important for biofilm formation. Therefore, interfering with the function of the sortase C protein could prevent, disrupt or retard biofilm development in *C. striatum*, and therefore serve as a target for candidate AMPs. In B-AMP v1.0, we evaluated the interactions of 100 select predicted 3D AMP models with the catalytic site residues of the semi-open lid conformation of the *C. striatum* sortase C protein (using AutoDock Vina) (Mhade et al., 2021). This filtered subset included 88 AMPs ranging from 2 to 8 amino acid residues in length and 12 AMPs ranging from 9 to 20 residues in length. In addition, as a positive control or standard, the LPMTG motif of the pilin subunit of *C. striatum* was docked to the sortase C protein. Based on binding energy scores and interacting residues, we proposed a preference scale (from 0-10, with 0 being the lowest score and 10 being the highest score) using which candidate AMPs could be taken up for further evaluations (Mhade et al., 2021). An important next step to validate this scoring system includes *in silico* approaches such as MD simulations, which enable the investigation of AMP-protein interactions in explicit solvent environments (Geng et al., 2019; Wang et al., 2019). Based on the previously proposed preference score, we selected 58 AMP-sortase C docking models for 20 nanoseconds (ns) MD simulations, distributed across preference scores 0, 1, 8, 9 and 10 (a combination of low and high scores), in addition to the standard LPMTG motif. Taken together, this resulted in a total of ∼1.18 µs of trajectory data from the 59 AMP-sortase C complexes (including the LPMTG motif) **(Table 2)**.

**Table 2:** *In silico* molecular docking models of select AMP-sortase C interactions analyzed using MD simulations for a cumulative period of ∼1.18 µs.

| **AMP-sortase C docking models** | **Number of docking models analyzed in MD simulation** | **Number of docking models where the AMP shows longevity of interactions** | **Number of docking models where the AMP drifts away** |
| --- | --- | --- | --- |
| LPMTG motif of the pilin subunit of *C. striatum* | 1 | 1 | None |
| Preference Score 10 | 6 | 5 | 1<br>(Pep4785) |
| Preference Score 9 | 6 | 6 | None |
| Preference Score 8 | 42 | 41 | 1<br>(Pep5487) |
| Preference Score 1 | 3 | 2 | 1<br>(Pep3309) |
| Preference Score 0 | 1 | 1<br>(Pep993) | None |

For each simulation setup, the AMP-sortase C complex was equilibrated with solvent, followed by neutralization with ions and further equilibration. The total number of atoms in each system ranged from 49440 to 56333. After processing, the trajectories were analyzed visually, and the interactions mapped using DIMPLOT were correlated for the initial and final snapshots. The standard LPMTG motif of the *C. striatum* pilin subunit (Pep1), known to interact with the catalytic site residues of the sortase C protein during pilus assembly, showed longevity of interactions with the residues His168, Gly170, Ile235, and Asn236, of the sortase C protein, where Cys230-His168-Arg239 constitutes the active site catalytic triad (**Figure 4A**) (Ton-That and Schneewind, 2004). At the end of the 20 ns simulation, the majority of the AMP-sortase C models from the high preference scores (52/54, preference scores 8, 9 and 10), remained bound to the semi-open lid conformation of the sortase C protein **(Figures 4B and C)**. The remaining 2 AMPs from the high preference scores (Pep4785 in preference score 10 and Pep5487 in preference score 8), drifted away from the active site region **(Figure 4D)**. For the 4 AMP-sortase C complexes in the lower preference scores (preference scores 0 and 1), 3 of the AMPs were seen to remain complexed with the sortase C protein, with 1 AMP (Pep3309) drifting away (**Figures 4E and F**). Taken together, this indicates that the proposed preference score based on *in silico* molecular docking provides a candidate list of AMPs for further evaluation. This is notably observed for AMP-sortase C models with high preference scores (scores 8, 9 and 10), where MD simulations showed the majority of the candidate AMPs demonstrating a high specificity of interactions with the sortase C protein. Among the low preference scores (scores 0 and 1), 3 out of the 4 classified AMPs demonstrated longevity of interactions with the sortase C protein. This indicates that AMPs with low preference scores based on *in silico* molecular docking should be evaluated with additional approaches, which could prevent their erroneous exclusion as potential anti-biofilm candidates. Taken together, 55 out of 58 AMP-sortase C complexes across all preference scores were validated based on longevity of interactions with or drifting away from, the sortase C protein, underscoring the value of the proposed *in silico* approach to identify and filter candidate AMPs for further *in vitro* and *in vivo* testing for anti-biofilm potential against *C. striatum*. The MD simulation trajectory movies, snapshots, and interactions of the entire AMP-sortase C complexes is provided in **Suppl File 10**.

**Figure 4:**
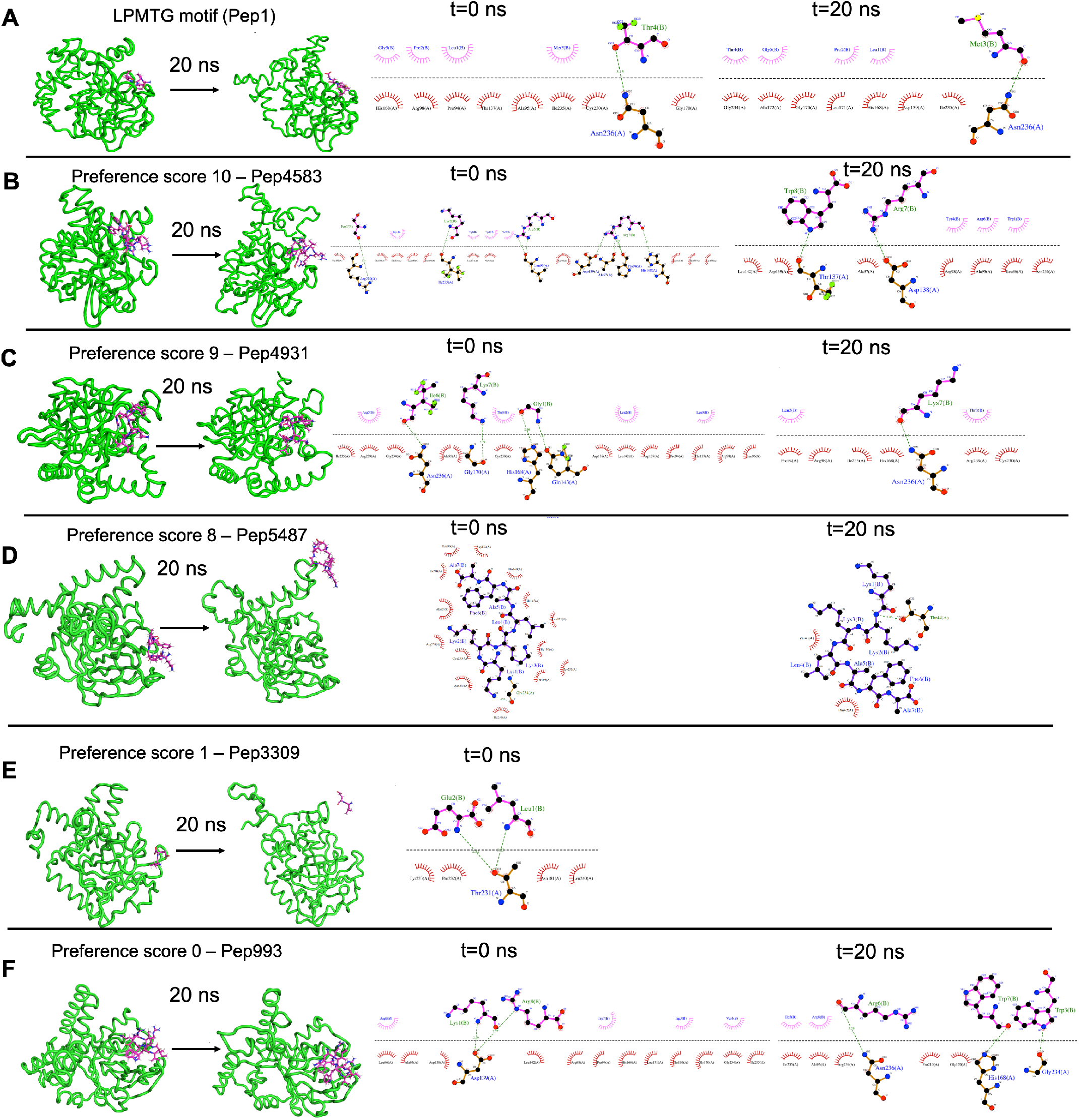
Representative results from Molecular Dynamics (MD) simulations to validate select AMP-biofilm target interactions with the catalytic site of the *C. striatum* sortase C protein. **(A)** The LPMTG motif (Pep1) of the *C. striatum* pilin subunit (magenta) remains in complex with the 3D predicted structure of the semi-open lid conformation of the *C. striatum* sortase C (green) at the end of 20 ns trajectory. The adjacent interaction maps at t=0 ns and t=20 ns indicate the longevity of the interactions. **(B)** Pep4583, categorized in preference score 10 (highest score, based on *in silico* molecular docking) showed longevity of interactions with the *C. striatum* sortase C protein at the end of 20 ns trajectory. **(C)** Among the AMPs in preference score 9, all AMPs, including Pep4931, showed longevity of interactions at the end of the 20ns simulation. **(D)** For AMPs in preference score 8, all AMPs showed longevity of interactions with the *C. striatum* sortase C protein, except Pep5487 which drifted away. **(E)** Pep3309, categorized in preference score 1 (lower score, based on *in silico* molecular docking), was observed to drift away from the sortase C protein at the end of 20 ns trajectory. **(F)** Pep993, categorized in preference score 0 (lowest score, based on *in silico* molecular docking), continued to be in complex with the sortase C protein at the end of the 20 ns simulation.

### A case study of using B-AMP for the *in silico* evaluation of AMPs against two biofilm targets in widely-encountered bacterial co-pathogens

The Gram-negative pathogen *P. aeruginosa* and Gram-positive pathogen *S. aureus* are widely-encountered bacterial co-pathogens, present together in a range of infection states, including wounds and disease-affected lungs (Alves et al., 2018; Yung et al., 2021). In biofilm infections, the two pathogens are found in close association with each other, and their concomitant presence is believed to worsen the outcome of the infection (DeLeon et al., 2014; Briaud et al., 2020). Therefore, AMPs with the potential to target both pathogens, would not only provide a composite treatment approach, but would also reduce the use and need of antibiotic therapy. Given that initial attachment is the first step in biofilm formation, and the well-established role of cell adhesion in mediating initial attachment of *P. aeruginosa* and *S. aureus* to biotic and abiotic surfaces, we selected cell adhesion as the common biofilm target function across the two co-pathogens (Donlan, 2001; Dunne, 2002; Vallet et al., 2004; Garrett et al., 2008; Paharik and Horswill, 2016). The search query ‘*Pseudomonas aeruginosa*’ and ‘*Staphylococcus aureus*’ in the biofilm target repository in B-AMP v2.0, with filtering using the PDB function ‘cell adhesion’, identified one target each for *P. aeruginosa* and *S. aureus*; for *P. aeruginosa* this was Target ID 37 or the ‘fimbrial subunit CupB6’ and for *S. aureus* this was Target ID 1 or ‘biofilm-associated surface protein’. Part of the chaperone-usher system in *P. aeruginosa*, known to assemble pili on the bacterial surface, the fimbrial subunit CupB6 (Target 37) of *P. aeruginosa* is the tip adhesion subunit that is attached to the main pilin shaft. The surface-exposed polyproline helix of CupB6 is part of the adhesion domain, and is believed to mediate a range of protein-protein interactions (Rasheed et al., 2016). In *S. aureus*, Target ID 37 or ‘biofilm-associated surface protein’, mediates intercellular adhesion and biofilm formation, with the N-terminal lobe of the protein identified to have a significant role (Ma et al., 2021). A filtered list of 2035 AMPs known to have both anti-Gram positive and anti-Gram negative activity (from B-AMP v1.0) **(Suppl File 11)**, were docked with the identified active site regions of the biofilm targets using AutoDock v4.2. Based on the docking results, we filtered AMPs with negative energy binding scores and binding energy differences in the range of ± 1 kcal/mol for both Target ID 1 and Target ID 37. This resulted in a filtered list of 25 candidate AMPs for both biofilm targets, which bind to the identified active sites of the biofilm targets **(Table 3)**. Based on the filtering criteria applied, the predicted complexes and interacting residues (hydrogen and hydrophobic bonds) are displayed for the highest five candidate AMPs for both biofilm targets (**Figure 5**) in B-AMP v2.0 (https://b-amp.karishmakaushiklab.com/docked_dual.html). Based on the proposed candidate list, AMPs can be taken up for further *in silico* evaluation such as AMP-target dynamic interactions and MD simulations, as well as *in vitro* and *in vivo* evaluation against mixed-species *P. aeruginosa* and *S. aureus* biofilms. It is interesting to note that of the 25 candidate AMPs, 10 AMPs (Pep5241, Pep2533, Pep4707, Pep167, Pep3239, Pep4709, Pep3292, Pep3052, Pep3240 and Pep5488) were previously evaluated using *in silico* molecular docking with the sortase C protein of *striatum* (Mhade et al., 2021). Of these 10 AMPs, 6 AMPs (Pep5241, Pep4707, Pep3239, Pep4709, Pep3292, Pep3240), were categorized into high preference scores (scores 8, 9 and 10) based on interacting residues with the predicted model of sortase C. While this could be due promiscuity in interactions displayed by select AMPs, it underscores the fact that the proposed *in silico* pipelines can serve to identify potential candidate AMPs from a vast library, which can then be taken up for further evaluation.

**Table 3:** *In silico* molecular docking of 25 candidate AMPs with binding energy differences of ± 1 kcal/mol) for both biofilm targets, Target 1 or ‘biofilm-associated surface protein’ of *S. aureus* and Target 37 or ‘fimbrial subunit CupB6’ of *P. aeruginosa*.

| **AMP PepID**<br>**(from B-AMP)** | **Binding Energy**<br>**(kcal/mol) with**<br>**Target ID 1** | **Binding Energy**<br>**(kcal/mol) with**<br>**Target ID 37** | **Difference in**<br>**Binding Energies**<br>**(kcal/mol)** |
| --- | --- | --- | --- |
| Pep4975 | -0.05 | -0.04 | 0.01 |
| Pep5241 | -2.43 | -2.38 | 0.05 |
| Pep4710 | -3.96 | -4.03 | 0.07 |
| Pep1217 | -0.77 | -0.93 | 0.16 |
| Pep5037 | -0.17 | -0.36 | 0.19 |
| Pep5193 | -5.06 | -4.84 | 0.22 |
| Pep169 | -1.44 | -1.17 | 0.27 |
| Pep2533 | -3.04 | -3.41 | 0.37 |
| Pep3961 | -1.13 | -0.74 | 0.39 |
| Pep4707 | -6.01 | -5.58 | 0.43 |
| Pep783 | -2.23 | -1.78 | 0.45 |
| Pep167 | -1.03 | -0.57 | 0.46 |
| Pep2896 | -0.59 | -1.06 | 0.47 |
| Pep662 | -2.16 | -2.64 | 0.48 |
| Pep3239 | -1.77 | -2.25 | 0.48 |
| Pep4709 | -4.79 | -5.39 | 0.6 |
| Pep4842 | -1.99 | -1.38 | 0.61 |
| Pep3292 | -3.59 | -4.31 | 0.72 |
| Pep3052 | -3.14 | -2.4 | 0.74 |
| Pep407 | -0.32 | -1.07 | 0.75 |
| Pep3240 | -1.32 | -2.08 | 0.76 |
| Pep69 | -0.54 | -1.33 | 0.79 |
| Pep2530 | -1.62 | -2.44 | 0.82 |
| Pep5057 | -1.72 | -0.89 | 0.83 |
| Pep5488 | -3.38 | -2.38 | 1 |

**Figure 5:**
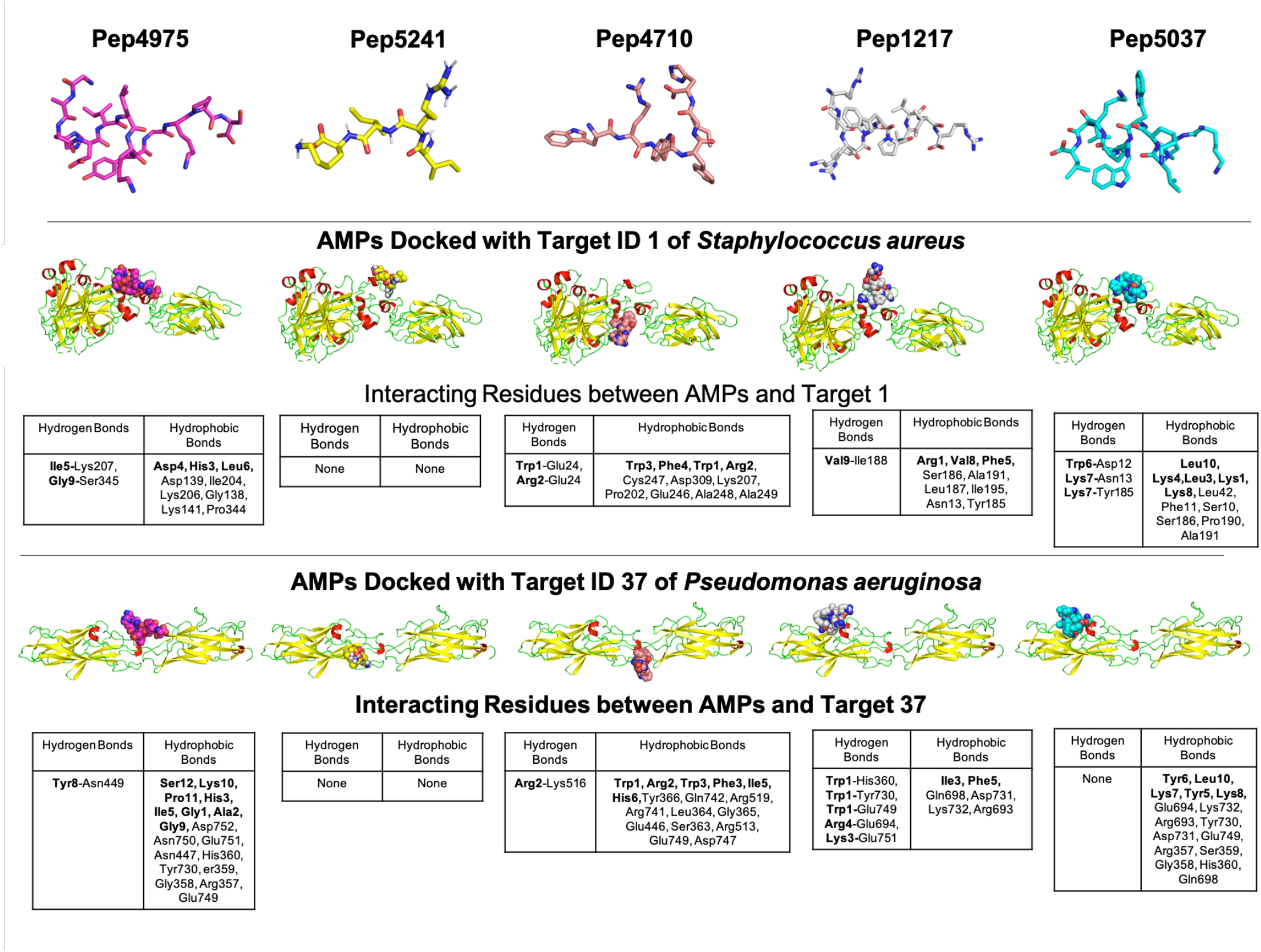
Representative results from *in silico* molecular docking for the filtered list of anti-Gram positive and anti-Gram negative AMPs with Target ID 37 or the ‘fimbrial subunit CupB6’ of *P. aeruginosa* and Target ID 1 or the ‘biofilm-associated surface protein’ of *S. aureus*. **(A)** Based on negative binding energy scores and binding energy differences in the range of ± 1 kcal/mol for both Target ID 1 and Target ID 37, the highest ranked AMPs were Pep4975, Pep5241, Pep4710, Pep1217 and Pep5037. **(B)** For the highest ranked AMPs, *in silico* docking models with Target ID 37 or the ‘fimbrial subunit CupB6’ of *P. aeruginosa* reveal a range of hydrogen and hydrophobic bonds for all AMPs, except for Pep5241. **(C)** For the highest ranked AMPs, *in silico* docking models with Target ID 1 or the ‘biofilm-associated surface protein’ of *S. aureus* reveal a range of hydrogen and hydrophobic bonds, for all AMPs, except for Pep5241. For Pep5241, there were no hydrogen or hydrophobic bonds detected, possibly owing to the short length of the AMP (4 amino acid residues).

**Figure 6:**
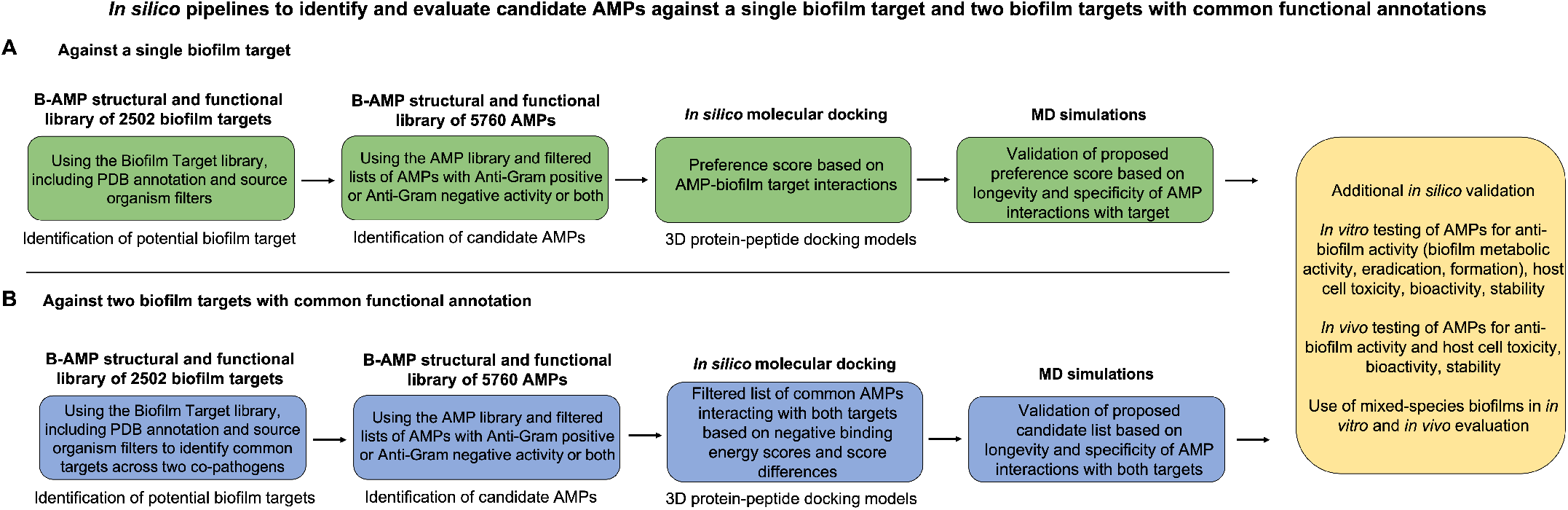
Summary of the proposed *in silico* pipelines to identify and evaluate candidate AMPs against a single biofilm target and two biofilm targets with common functional annotations. To present an example of the combined applicability of both the AMP and biofilm target libraries in the B-AMP v2.0 repository, we present two *in silico* pipelines to identify and evaluate AMP-biofilm target interactions. **(A)** For evaluating AMPs against a single biofilm target, the biofilm target library can be used to identify a suitable protein target based on the bacterial species under study and the role of the target in biofilm formation. Next, filtered lists of AMPs with anti-Gram positive or anti-Gram negative activity or both can be subject to *in silico* molecular docking with the biofilm target, followed by development of a preference score of candidate AMPs based on docking scores and interacting residues. The predicted 3D protein-peptide docking models can be validated using MD simulations, followed by possible *in vitro* and *in vivo* testing. **(B)** For evaluating AMPs against two biofilm targets, the biofilm target library can be used to identify suitable protein targets, possibly with a common functional classification, across two biofilm-forming bacterial co-pathogens. Next, filtered lists of AMPs with anti-Gram positive or anti-Gram negative activity or both can be subject to *in silico* molecular docking with the biofilm targets, followed by development of a candidate list of AMPs based on negative binding energy scores and binding energy differences across both targets. The predicted 3D protein-peptide docking models can be validated using MD simulations, followed by possible *in vitro* and *in vivo* testing with mixed-species biofilms.

## Discussion

### AMPs as potential anti-biofilm approaches

In clinical infection states, biofilms are observed as multicellular aggregates, often consisting of more than one bacterial species. Under these conditions, biofilms display increased tolerance to antibiotic treatments, resulting in prolonged and repeated antibiotic usage (Høiby et al., 2010). Given this, identifying and evaluating novel anti-biofilm approaches, such as AMPs, is both necessary and important. For several reasons, AMPs are well-suited to be developed into anti-biofilm approaches (Batoni et al., 2011; Pompilio et al., 2012; Dawgul et al., 2014; Chung and Khanum, 2017; Jorge et al., 2017; Yasir et al., 2018; Galdiero et al., 2019; di Somma et al., 2020; Huan et al., 2020; Hancock et al., 2021). To start with, a vast repertoire of natural and synthetic AMPs exists, and they lend themselves well for *in silico, in vitro* and *in vivo* evaluation against biofilm-forming pathogens. Further, AMPs have been shown to act on bacteria with slow growth rates and low metabolic activity, including multidrug resistant strains, and the emergence of bacterial resistance to AMPs is rare (Batoni et al., 2011; Dawgul et al., 2014; Yasir et al., 2018; el Shazely et al., 2020). Finally, in addition to their membrane disrupting features, AMPs also have the ability to interfere with or act against a range of specific bacterial targets and processes. This provides them the ability to target the different stages of biofilm formation, as well as more than one bacterial pathogen, which is particularly relevant in the context of mixed-species biofilm infections.

### Relevance of biofilm protein targets in B-AMP v2.0

Biofilm formation is a complex and coordinated process, involving several different bacterial proteins with a wide range of functions. Given this, bacterial proteins involved in biofilm formation and development are potential biofilm targets, and AMPs that either interfere with or inhibit these targets can reduce, prevent, and possibly disrupt biofilms (Jorge et al., 2012; Zapotoczna et al., 2017; Yasir et al., 2018; Kalsy et al., 2020; Portelinha and Angeles-Boza, 2021). While most AMP studies focus on their membrane and cell wall disrupting properties, AMPs also exert their mechanism of action via several intracellular bacterial systems, including interference with cell adhesion proteins, interruption of quorum sensing systems, degradation of extracellular matrix proteins, blockade of the alarmone signaling system, destruction of nucleic acids, downregulation of transport binding proteins and inhibition of fundamental cell process such as transcription, translation and energy metabolism (de la Fuente-Núñez et al., 2014; Yasir et al., 2018; Luo and Song, 2021). Based on their functional classification and designated nomenclature, the biofilm targets in B-AMP v2.0 represent proteins that are involved in (based on PDB functional annotations) or could be involved in (based on bacterial protein names) several of these bacterial processes and systems **(Suppl File 1)**, and can therefore serve as candidate targets for AMPs with anti-biofilm potential.

### Potential applications of resources in B-AMP for biofilm studies

Taken together, B-AMP v2.0 is a structural and functional repository consisting of (i) 5766 AMPs from natural and synthetic sources, (ii) 2502 bacterial protein targets from a range of bacterial species, and (iii) AMP-biofilm target models for widely-recognized biofilm-forming pathogens. In addition, updated codes and in-house python scripts used to compile and build the repository, and the references of relevance have been listed. Given this, the curated resources in B-AMP v2.0 can be leveraged for a range of AMP studies of relevance to biofilms, including *in silico* screening, and *in vitro* and *in vivo* experimental studies. Additionally, the repository also hosts *in silico* pipelines to present examples of the utilization of the AMP and biofilm target libraries. In B-AMP v1.0 a subset of the filtered list of AMPs with anti-Gram positive activity were docked to the sortase C biofilm target of *C. striatum*. In B-AMP v2.0, this work was extended to validate select 3D protein-peptide docked complexes using MD simulations. In addition to a pipeline to evaluate AMPs against a single biofilm target, B-AMP v2.0 presents a case study to evaluate AMPs against dual biofilm targets with a common functional classification, using biofilm-forming co-pathogens *P. aeruginosa* and *S. aureus*. For this, a filtered list of AMPs with both anti-Gram positive and anti-Gram negative activity were subject to *in silico* molecular docking with cell adhesion targets of these two pathogens. Based on these results, a categorization approach for AMPs into preference scores for a single biofilm target, and dual biofilm targets is also presented. Taken together, as the two case studies exemplify, the collective resources in B-AMP, consisting of a structural and functional library of AMPs and biofilm targets, and 3D AMP-biofilm target docking models, complement each other for studies related to identifying potential AMP-biofilm target interactions, and provide an accessible set of *in silico* resources to extend the studies to a range of biofilm-forming pathogens and mixed-species biofilm states.

### Limitations of B-AMP v2.0

In the upgraded version, B-AMP v2.0 includes biofilm protein targets, with data collected and compiled from PDB and UniProt databases. Based on the limitations of B-AMP v1.0, where we discussed the explicit need to include biofilm targets in the repository, B-AMP v2.0 is a more comprehensive resource for AMP studies relevant to biofilms. However, given that the sources of data are the PDB and UniProt databases, biofilm targets in B-AMP v2.0 are exclusively protein in nature. This is a limitation, given that AMPs are known to target bacterial components such as surface lipids (AMP-lipid interactions are involved in membrane disruption) and polysaccharides, that are also known to play a role in biofilm formation (Hollmann et al., 2018; Benfield and Henriques, 2020; Lin et al., 2020; Zhang et al., 2021). Future expansions of B-AMP v2.0 can include bacterial lipid and lipoprotein targets from databases such as LIPID MAPS Consortium and DOLOP, and bacterial carbohydrates using Polysac DB, Bacterial CSDB, and CAZy (www.cazy.org/) (LIPID MAPS® Lipidomics Gateway; Babu et al., 2006; Onyango et al., 2021; Leggieri et al., 2022; Liew et al., 2022). An additional limitation in B-AMP v2.0 is that the targets represent only bacterial targets (no fungal targets), and 3D structural models are not available for all biofilm targets. While expanding the biofilm target library in B-AMP v2.0 would be useful, the current resources in B-AMP v2.0 serve as an excellent starting point for *in silico* AMP-biofilm target investigations for bacterial biofilm pathogens.

## Conclusions

There has been a concerted push to build dedicated AMP resources for biofilm studies, with a focus on enabling *in silico* investigations as high-throughput screening and predictive tools to identify and evaluate AMPs with anti-biofilm potential. In this upgraded version, B-AMP v2.0 is a comprehensive repository of AMPs, bacterial protein targets and AMP-biofilm target interactions for relevant bacterial pathogens, with pre-determined structural models and annotations to existing scientific literature. B-AMP v2.0 continues to be freely available to the community at https://b-amp.karishmakaushiklab.com, and will be regularly updated with AMP structural models, biofilm targets and 3D protein-peptide interaction models for a range of biofilm-forming pathogens.

## Acknowledgements

We thank Shreeya Mhade, Stutee Panse, Gandhar Tendulkar and Rohit Awate for updating B-AMP v1.0 with additional 3D structural AMP models. We thank Pulkit Anupam Srivastava for initial discussions on data retrieval and database development. The computational infrastructure and support provided by the Bioinformatics Research and Applications Facility (BRAF) funded by the National Supercomputing Mission, Government of India at the Centre For Development of Advanced Computing, Pune are gratefully acknowledged. YN, SR, and RMY thank SASTRA Deemed to be University for infrastructural support and research facilities.

## Supplementary Files

**Suppl File 1:** Non-redundant list of 2502 biofilm protein targets in B-AMP v2.0

**Suppl Table 2:** Distribution of biofilm targets in B-AMP v2.0 based on bacterial class, family, genera or description as per NCBI Taxonomy

**Suppl Table 3:** Distribution of biofilm targets in B-AMP v2.0 based on bacterial species or description as per NCBI taxonomy

**Suppl Table 4:** List of PDB target functions of biofilm targets in B-AMP v2.0

**Suppl Table 5:** List of UniProt target functions of biofilm targets in B-AMP v2.0

**Suppl File 6:** Table of PDB target functions with source organisms and names of biofilm protein targets

**Suppl Table 7:** List of bacterial species with PDB structures in B-AMP v2.0

**Suppl Table 8:** List of *P. aeruginosa* biofilm targets modeled using RoseTTAFold in B-AMP v2.0

**Suppl Table 9:** List of *S. aureus* biofilm targets modeled using RoseTTAFold in B-AMP v2.0

**Suppl Table 10:** Full list of AMP-sortase C docking models for *C. striatum* analyzed using MD simulations (preference scores 10, 9, 8, 1 and 0)

**Suppl File 11:** Filtered list of 2035 AMPs with both anti-Gram positive and anti-Gram negative activity used for *in silico* molecular docking against Target 37 of *P. aeruginosa* and Target 1 of *S. aureus*.

