## Supplementary material for "‘Targeting’ the Search: An Upgraded Structural and Functional Repository of Antimicrobial Peptides for Biofilm Studies (B-AMP v2.0) with a Focus on Biofilm Protein Targets": Suppl File 1

| Target ID | Bacterial Protein Name | Organism Name | Strain | UniProt ID | PDB ID | PubMed ID | Classification as per UniProt GO Molecular Function | Classification of Function as per PDB |
|---|---|---|---|---|---|---|---|---|
| 1 | Biofilm-associated surface protein | *Staphylococcus aureus* | | Q79LN3 | 7C7U | 11292810; 34046916 | Not Available | Cell adhesion |
| 2 | Type 1 fimbrin D-mannose specific adhesin (Protein FimH) | *Escherichia coli* | Strain K12 | P08191 | 6YHW | 2890081; 7905476; 7610040; 9278503; 16738553; 2900235; 1971261; 12010488 | Mannose-binding | Cell adhesion |
| 3 | Fimbrial adhesin | *Proteus mirabilis* | Strain HI4320 | B4EUK6 | 6Y4E | 18375554; 32780778 | Not Available | Cell adhesion |
| 4 | Putative large adhesion protein (Lap) involved in biofilm formation | *Marinobacter nauticus* | Strain ATCC 49840 / DSM 8798 / CIP 103578 / SP17 | H8W6K8 | 6XI3 | 22689231; 31335890; 32984811 | Calcium ion binding | Metal binding protein |
| 5 | Minor fimbrium subunit Mfa5 | *Porphyromonas gingivalis* | Strain ATCC 33277 / DSM 20709 / CIP 103683 / JCM 12257 / NCTC 11834 / 2561 | B2RHG5 | 6TNJ | 18524787; 19589838; 24118823; 26001707; 26437277 | Not Available | Cell adhesion |
| 6 | Fimbrin-like protein FimI | *Escherichia coli* | Strain K12 | P39264 | 6SWH | 7610040; 9278503; 16738553; 6147250 | Not Available | Structural protein |
| 7 | TasA anchoring / assembly protein (Biofilm assembly accessory protein TapA) | *Bacillus subtilis* | Strain 168 | P40949 | 6QAY | 8969508; 9384377; 2507524; 10464223; 10559173; 16430695; 16430696; 21477127; 23646920; 24488317 | Not Available | Protein fibril |
| 8 | Sensor protein SrrB (EC 2.7.13.3) (Staphylococcal respiratory response protein B) | *Staphylococcus aureus* | | Q9L523 | 6PAJ, 7JL6 | 11157922; 15060046, 32354997 | ATP binding, phosphorelay sensor kinase activity | Membrane protein |
| 9 | Uncharacterized fimbrial-like protein YdeQ | *Escherichia coli* | Strain K12 | P77588 | 6MAP | 9097039; 9278503; 16738553 | Not Available | Sugar binding protein |
| 10 | Transcriptional regulatory protein LytR | *Staphylococcus aureus* | Strain NCTC 8325 / PS 47 | P60611 | 6M8O | 8550490; 8824633; 19502411; 23733465; 25491472 | DNA-binding Transcription activator activity, phosphorelay response regulator activity, Transcription cis-regulatory region binding | Signaling protein |
| 11 | Curli production assembly / transport component CsgG | *Escherichia coli* | Strain K12 | P0AEA2 | 6L7C | 8817489; 8905232; 9278503; 16738553 | Identical protein binding | Protein transport |
| 12 | Major fimbrium subunit FimA type-1 (Fimbrillin) (Fimbrilin) (Major fimbrial subunit protein type I) | *Porphyromonas gingivalis* | Strain ATCC 33277 / DSM 20709 / CIP 103683 / JCM 12257 / NCTC 11834 / 2561 | B2RH54 | 6JZK | 7902712; 18524787; 8778568; 9786913; 10377095; 12593606; 15165251; 17675496; 17526848; 20530728; 23809984; 27062925 | Structural molecule | Cell adhesion |
| 13 | RNA polymerase sigma-H factor (Sigma-30) | *Porphyromonas gingivalis* | Strain ATCC 15692 / DSM 22644 / CIP 104116 / JCM 14847 / LMG 12228 / 1C / PRS 101 / PAO1 | Q06198 | 6IN7 | 8432708; 8378309; 7961421; 10984043; 7737518 | DNA-binding Transcription activator activity, sigma factor activity, Transcription cis-regulatory region binding | Transcription |
| 14 | Major curlin subunit | *Escherichia coli* | Strain K12 | P28307 | 6G8C | 8459772; 8817489; 8905232; 9278503; 16738553; 1357528; 1677357; 17010156 | Identical protein binding | Protein fibril |
| 15 | Methyl-accepting chemotaxis protein TlpQ (TlpQ chemoreceptor) | *Porphyromonas gingivalis* | Strain ATCC 15692 / DSM 22644 / CIP 104116 / JCM 14847 / LMG 12228 / 1C / PRS 101 / PAO1 | Q9I0I4 | 6FU4 | 10984043; 33097715; 30425146 | Not Available | Signaling protein |
| 16 | Streptococcal hemagglutinin (Hs antigen) (Sialic acid-binding adhesin) | *Streptococcus gordonii* | Strain ChalliS / ATCC 35105 / BCRC 15272 / CH1 / DL1 / V288 | A8AWU7 | 6EFC, 6EFD, 6X3K | 11854202; 17720781; 8366515; 15213130; 19884334; 21071690; 27616700 | Not Available | Cell adhesion |
| 17 | Putative hemin storage protein | *Bordetella bronchiseptica* | Strain ATCC BAA-588 / NCTC 13252 / RB50 | A0A0H3LKK6 | 6AU1 | 12910271; 29684093 | Hydrolase activity, acting on carbon-nitrogen (but not peptide) bonds | Hydrolase |
| 18 | Autoinducer-2 kinase (AI-2 kinase) (EC 2.7.1.189) | *Escherichia coli* | Strain K12 | P77432 | 5YA1 | 9097039; 9278503; 16738553; 8649811; 15601708; 17557827; 20025244 | Autoinducer-2 kinase activity, phosphoTransferase activity, alcohol group as acceptor | Structural protein |
| 19 | Putative outer Membrane protein | *Salmonella typhimurium* | Strain LT2 / SGSC1412 / ATCC 700720 | Q8ZRK4 | 5Y9H | 11677609; 16793551; 18448124; 29125121 | Not Available | Cell adhesion |
| 20 | Putative fimbriae subunit | *Salmonella typhimurium* | Strain LT2 / SGSC1412 / ATCC 700720 | Q7CR56 | 5Y9H | 11677609; 29125121 | Not Available | Cell adhesion |
| 21 | Putative ABC transporter, ATP-binding protein ComA | *Streptococcus mutans serotype C* | Strain ATCC 700610 / UA159 | Q8DW05 | 5XE9 | 12397186; 20100826; 23534432; 28642545 | ABC-type bacteriocin transporter activity, ATP binding, cysteine-type peptidase activity | Hydrolase |

| Target ID | Bacterial Protein Name | Organism Name | Strain | UniProt ID | PDB ID | PubMed ID | Classification as per UniProt GO Molecular Function | Classification of Function as per PDB |
|---|---|---|---|---|---|---|---|---|
| 22 | HTH-type Transcriptional regulator RcdA (Regulator of csgD) | *Escherichia coli* | Strain K12 | P75811 | 5X5I | 8905232; 9278503; 16738553; 23233451 | DNA binding transcription factor activity | Transcription |
| 23 | CRISPR-associated protein Csy1 | *Pseudomonas aeruginosa* | Strain UCBPP-PA14 | Q02ML9 | 5UZ9, 6B44, 6B45, 6B47, 6B48, 6NE0 | 17038190; 21398535; 21081758; 21536913; 22522703; 28340349; 28985564; 28985564; 28985564; 28985564; 30872121 | Not Available | Immune system / RNA |
| 24 | CRISPR-associated protein Csy2 | *Pseudomonas aeruginosa* | Strain UCBPP-PA14 | Q02MM0 | 5UZ9, 6B44, 6B45, 6B47, 6B48, 6NE0 | 17038190; 21398535; 21081758; 21536913; 22522703; 28340349;28985564;28985564;28985564; 28985564; 30872121 | Not Available | Immune system / RNA |
| 25 | CRISPR-associated protein Csy3 | *Pseudomonas aeruginosa* | Strain UCBPP-PA14 | Q02MM1 | 5UZ9, 5XLO, 5XLP, 6B44, 6B45, 6B46, 6B47, 6B48, 6NE0 | 17038190; 21398535; 21081758; 21536913; 22522703; 28340349; 28574055; 28574055; 28985564; 28985564; 28985564; 28985564; 28985564; 30872121 | Not Available | Immune system / RNA |
| 26 | Major biofilm matrix component (Spore coat-associated protein N) (Translocation-dependent antimicrobial spore component) | *Bacillus subtilis* | Strain 168 | P54507 | 5OF1 | 8969508; 9384377; 3096962; 10368135; 10658653; 10049401; 10464223; 12429099; 16430695; 16430696; 20080671; 23646920; 29531041 | Identical protein binding | Protein fibril |
| 27 | Minor fimbrium anchoring subunit Mfa2 (Minor fimbrial antigen 2) | *Porphyromonas gingivalis* | Strain ATCC 33277 / DSM 20709 / CIP 103683 / JCM 12257 / NCTC 11834 / 2561 | B2RHG2 | 5NFI | 18524787; 19589838; 26437277; 27062925; 29379120 | Not Available | Cell adhesion |
| 28 | Minor fimbrium tip subunit Mfa3 | *Porphyromonas gingivalis* | Strain ATCC 33277 / DSM 20709 / CIP 103683 / JCM 12257 / NCTC 11834 / 2561 | B2RHG3 | 5NF4 | 18524787; 19589838; 24118823; 26001707; 26437277; 29379120 | Not Available | Cell adhesion |
| 29 | Minor fimbrium subunit Mfa1 (Pg-II fim a) | *Porphyromonas gingivalis* | Strain ATCC 33277 / DSM 20709 / CIP 103683 / JCM 12257 / NCTC 11834 / 2561 | B2RHG1 | 5NF2, 5NF3 | 18524787; 9786913; 11083792; 12593606; 15972485; 19589838; 23809984; 24118823; 26001707; 26437277; 29379120; 29379120 | Not Available | Cell adhesion |
| 30 | Methyl-accepting chemotaxis protein PctA | *Pseudomonas aeruginosa* | Strain ATCC 15692 / DSM 22644 / CIP 104116 / JCM 14847 / LMG 12228 / 1C / PRS 101 / PAO1 | G3XD24 | 5LTX, 5T65, 5T7M | 8522505; 9353923; 10984043; 16233808; 23650915; 27031335; 30425146; 33097715; 31964737 | Amino acid binding, transmembrane signaling receptor activity | Signaling protein |
| 31 | Regulatory protein RsaL (RsaL protein) (Virulence gene repressor RsaL) | *Pseudomonas aeruginosa* | | Q9X7H4 | 5J2Y | 1902216; 10094696 | DNA binding | Gene regulation |
| 32 | Sensor protein kinase WalK (EC 2.7.13.3) | *Staphylococcus aureus* | Strain NCTC 8325 / PS 47 | Q2G2U4 | 5IS1 | 10368139; 14762013; 17827301; 22825451; 27389096 | ATP binding, metal ion binding, phosphorelay response regulator activity, phosphorelay sensor kinase activity, protein kinase activator activity | Transferase |
| 33 | Protein FimG | *Escherichia coli* | Strain K12 | P08190 | 5IQO | 2890081; 7610040; 9278503; 16738553 | Not Available | Cell adhesion |
| 34 | 2-heptyl-4(1H)-quinolone synthase subunit PqsB | *Pseudomonas aeruginosa* | Strain ATCC 15692 / DSM 22644 / CIP 104116 / JCM 14847 / LMG 12228 / 1C / PRS 101 / PAO1 | Q9I4X2 | 5DWZ | 10984043; 24239007; 26811339 | AcylTransferase activity | Transferase |
| 35 | 2-heptyl-4(1H)-quinolone synthase subunit PqsC (EC 2.3.1.230) | *Pseudomonas aeruginosa* | Strain ATCC 15692 / DSM 22644 / CIP 104116 / JCM 14847 / LMG 12228 / 1C / PRS 101 / PAO1 | Q9I4X1 | 5DWZ | 10984043; 24239007; 26811339 | 3-oxoacyl-[acyl-carrier-protein] synthase activity, beta-ketoacyl-acyl-carrier-protein synthase III activity | Transferase |

| Target ID | Bacterial Protein Name | Organism Name | Strain | UniProt ID | PDB ID | PubMed ID | Classification as per UniProt GO Molecular Function | Classification of Function as per PDB |
|---|---|---|---|---|---|---|---|---|
| 36 | Minor fimbrium tip subunit MfA4 (Immunoreactive 32 kDa antigen) | *Porphyromonas gingivalis* | Strain ATCC 33277 / DSM 20709 / CIP 103683 / JCM 12257 / NCTC 11834 / 2561 | B2RHG4 | 5DHM | 18524787; 19589838; 24118823; 26001707; 26437277; 26972441 | Not Available | Cell adhesion |
| 37 | Fimbrial subunit CupB6 | *Pseudomonas aeruginosa* | Strain ATCC 15692 / DSM 22644 / CIP 104116 / JCM 14847 / LMG 12228 / 1C / PRS 101 / PAO1 | Q9HWU7 | 5CYL | 10984043; 27481165 | Not Available | Cell adhesion |
| 38 | CRISPR-associated nuclease / helicase Cas3 subtype I-F / YPEST (EC 3.1.-.-) (EC 3.6.4.-) | *Pseudomonas aeruginosa* | Strain UCBPP-PA14 | Q02ML8 | 5B7I, 5GQH | 17038190; 18952788; 21398535; 21081758; 27455460; 27585537 | ATP binding, helicase activity, Hydrolase activity, Metal ion binding | Hydrolase |
| 39 | Diguanylate cyclase DosC (DGC) (EC 2.7.7.65) (Direct oxygen-sensing cyclase) | *Escherichia coli* | Strain K12 | P0AA89 | 4ZVA | 9097039; 9278503; 16738553; 16418169; 17010156; 19764732; 19332833; 21067162; 20303158; 20553324; 20576684; 22005448; 26148715 | Carbon monoxide binding, Diguanylate cyclase activity, GTP binding, heme binding, Metal ion binding, oxygen binding | Signaling Protein |
| 40 | PslG | *Pseudomonas aeruginosa* | Strain ATCC 15692 / DSM 22644 / CIP 104116 / JCM 14847 / LMG 12228 / 1C / PRS 101 / PAO1 | Q9I1N2 | 4ZN2 | 10984043; 26611635; 26424791 | Hydrolase activity, hydrolyzing O-glycosyl compounds | Hydrolase |
| 41 | GGDEF domain-containing protein | *Pseudomonas aeruginosa* | Strain ATCC 15692 / DSM 22644 / CIP 104116 / JCM 14847 / LMG 12228 / 1C / PRS 101 / PAO1 | Q9HW69 | 4WXO | 10984043 | Diguanylate cyclase activity, Metal ion binding | Transferase |
| 42 | Transcriptional regulator FleQ | *Pseudomonas aeruginosa* | Strain ATCC 15692 / DSM 22644 / CIP 104116 / JCM 14847 / LMG 12228 / 1C / PRS 101 / PAO1 | G3XCV0 | 4WXM | 10984043; 9287015; 11673434; 22581773; 26362077; 26712005 | ATP binding, cis-regulatory region Sequence-specific DNA binding, cyclic-di-GMP binding, DNA-binding Transcription activator activity, DNA-binding Transcription repressor activity, Transcription cis-regulatory region binding | Transcriptional regulator |
| 43 | Surface protein G | *Staphylococcus aureus* | Strain NCTC 8325 / PS 47 | Q2G2B2 | 4WVE | 11830639; 12634333; 14523109; 17660408 | Not Available | Structural protein |
| 44 | Fimbrial protein (Pilin) | *Neisseria meningitidis serogroup B* | Strain MC58 | P05431 | 4V1J | 3141743; 10710307; 413571; 8645220; 7934852; 8993351; 9515697; 17121595; 27698424 | Not Available | Cell adhesion |
| 45 | Transcriptional regulatory protein CpxR | *Escherichia coli* | Strain K12 | P0AE88 | 4UHJ, 4UHK, 4UHS, 4UHT | 8294007; 8346018; 9278503; 16738553; 3058985; 7883164; 9351822; 9401031; 9473036; 10972835; 11830644; 16166523; 17259177; 23416055; 25207645; 28552574; 28552574; 30086390; 30086390 | DNA-binding Transcription activator activity, DNA-binding Transcription factor activity, phosphorelay response regulator activity, Transcription cis-regulatory region binding | Transcription |
| 46 | PgaB | *Aggregatibacter actinomycetemcomitans* | Actinobacillus actinomycetemcomitans, Haemophilus actinomycetemcomitans | A5HJW8 | 4U10 | 17851029 | Hydrolase activity, acting on carbon-nitrogen (but not peptide) bonds, Metal ion binding | Hydrolase |
| 47 | Outer membrane porin F | *Pseudomonas aeruginosa* | Strain ATCC 15692 / DSM 22644 / CIP 104116 / JCM 14847 / LMG 12228 / 1C / PRS 101 / PAO1 | P13794 | 4RLC, 5U1H | 2447060; 10984043; 25846137; 25846137 | Calcium ion binding, complement component C3b binding, porin activity | Peptide binding protein |
| 48 | Minor fimbrium tip subunit MfA4 (Immunoreactive 32 kDa antigen PG49) | *Porphyromonas gingivalis* | Strain ATCC BAA-308 / W83 | Q7MXK0 | 4RDB | 12949112; 27062925; 27062925 | Not Available | Structural protein |
| 49 | Fimbrium subunit Fim1C | *Phocaeicola vulgatus* | Strain ATCC 8482 / DSM 1447 / JCM 5826 / CCUG 4940 / NBRC 14291 / NCTC 11154 BacteroideS vulgatuS | A6L3B5 | 4QB7 | 17579514; 27062925; 27062925 | Not Available | Cell adhesion |
| 50 | Major fimbrium subunit FimA type-4 (Fimbrillin) (Fimbrilin) (Major fimbrial subunit protein type-4) (FimA4) | *Porphyromonas gingivalis* | Strain ATCC BAA-308 / W83 | P59914 | 4Q98 | 12949112; 23809984; 27062925; 27062925 | Structural molecule | Cell adhesion |

| Target ID | Bacterial Protein Name | Organism Name | Strain | UniProt ID | PDB ID | PubMed ID | Classification as per UniProt GO Molecular Function | Classification of Function as per PDB |
|---|---|---|---|---|---|---|---|---|
| 51 | Adhesin / invasin TibA autotransporter [Cleaved into: Adhesin / invasin TibA (Glycoprotein TibA); Adhesin / invasin TibA translocator] | *Escherichia coli O78:H11* | Strain H10407 / ETEC | Q9XD84 | 4Q1Q | 10417177; 20802035; 8039917; 11119488; 15784535; 25211077 | Not Available | Cell adhesion |
| 52 | Phosphate-binding protein PstS | *Porphyromonas gingivalis* | Strain ATCC 15692 / DSM 22644 / CIP 104116 / JCM 14847 / LMG 12228 / 1C / PRS 101 / PAO1 | G3XDA8 | 4OMB | 16232467; 10984043; 12596240; 18282104; 6436026; 16232935; 24023943; 25005086 | Phosphate ion binding | Transport protein |
| 53 | AIDA-I autotransporter (AIDA) [Cleaved into: Adhesin AIDA-I; AIDA-I translocator (AIDAc)] | *Escherichia coli* | | Q03155 | 4MEE | 1625582; 15950405; 8899706; 10559167; 15547278 | Not Available | Protein binding |
| 54 | Serine-rich adhesin for platelets (Adhesin SraP) (Staphylococcus aureus surface protein A) (SasA) | *Staphylococcus aureus* | Strain NCTC 8325 / PS 47 | Q2FUW1 | 4M00, 4M01, 4M02, 4M03 | 12634333; 15784571; 11830639; 18800056; 18621893; 19202081; 20472795; 20714350; 24901708; 24901708; 24901708; 24901708; 24901708 | Calcium ion binding | Calcium binding protein |
| 55 | Putative fimbrium subunit Fim1C | *Bacteroides uniformis* | Strain ATCC 8492 / DSM 6597 / CCUG 4942 / CIP 103695 / JCM 5828 / KCTC 5204 / NCTC 13054 / VPI 0061 | A7UZ95 | 4K4K | 2.71E+15 | Not Available | Cell adhesion |
| 56 | CapA (Capsular polysaccharide biosynthesis protein CapA) (Capsular polysaccharide type 8 biosynthesis protein cap8A) (Capsule biosynthesis protein CapA) (Chain length regulator (Capsular polysaccharide biosynthesis)) (Membrane protein CapB2) | *Staphylococcus aureus* | | Q7BSA3 | 4JMP | 10496925; 16565080; 24146800; 26048971; 3305292524146800 | Not Available | Transferase |
| 57 | Putative fimbrium tip subunit Fim1C | *Parabacteroides distasonis* | Strain ATCC 8503 / DSM 20701 / CIP 104284 / JCM 5825 / NCTC 11152 | A6LHQ6 | 4JG5 | 17579514; 26972441; 27062925; 27062925 | Not Available | Cell adhesion |
| 58 | Sensor protein VraS (EC 2.7.13.3) | *Staphylococcus aureus* | Strain Mu50 / ATCC 700699 | Q7A2Q0 | 4GT8 | 11418146; 10708580 | ATP binding, phosphorelay sensor kinase activity, protein dimerization activity | Transferase |
| 59 | Exodeoxyribonuclease III (EC 3.1.11.2) | *Pseudomonas aeruginosa* | | Q51380 | 4F1R | 8824606; 23281037; 23717639; 30758287 | Exodeoxyribonuclease III activity | Transcriptional regulator |
| 60 | Putative fimbrium tip subunit Fim1C | *Bacteroides ovatus* | Strain ATCC 8483 / DSM 1896 / JCM 5824 / BCRC 10623 / CCUG 4943 / NCTC 11153 | A7M4E1 | 4EPS | 27062925; | Not Available | Structural protein |
| 61 | Sensor histidine kinase CpxA (EC 2.7.13.3) | *Escherichia coli* | Strain K12 | P0AE82 | 4BIU | 3007473; 3058985; 8346018; 9278503; 16738553; 2185221; 8432716; 7883164; 9351822; 9401031; 9473036; 10972835; 11830644; 15919996; 16166523; 17259177; 20005847; 21317318; 21239493; 25207645; 24492262 | ATP binding, Identical protein binding, phosphoprotein phosphatase activity, phosphorelay sensor kinase activity | Transferase |
| 62 | Biofilm-surface layer protein A (ORF-1) | *Bacillus subtilis* | Strain 168 | P71014 | 4BHU | 9384377; 8752328; 18978066; 21742882; 22571672; 28701036; 23904481; 26324938 | Not Available | Structural protein |
| 63 | signaltransduction protein TRAP (Target of RNAIII-activating protein) | *Staphylococcus aureus* | Strain NCTC 8325 / PS 47 | Q2G2F3 | 4AE5 | 11160124; 14726534; 16177293 | Not Available | Signaling protein |
| 64 | Pneumococcal serine-rich repeat protein (PsrP) (Adhesin PsrP) (Serine-rich repeat protein PsrP) | *Streptococcus pneumoniae serotype 4* | Strain ATCC BAA-334 / TIGR4 | A0A0H2URK1 | 3ZGH | 11463916; 16861665; 18507531; 19202081; 19627498; 20714350; 24936067; 28456649; 28246170; 24430336; 27582320 | DNA binding | Structural protein |

| Target ID | Bacterial Protein Name | Organism Name | Strain | UniProt ID | PDB ID | PubMed ID | Classification as per UniProt GO Molecular Function | Classification of Function as per PDB |
|---|---|---|---|---|---|---|---|---|
| 65 | Trimeric autotransporter adhesin AtaA (TAA AtaA) (Type 5 secretion system autotransporter AtaA) | *Acinetobacter spp.* | Strain Tol 5 | K7ZP88 | 3WPA | 23155410; 17090933; 23893702; 27074146; 27305955; 28720107; 31653547; 31972092; 26698633 | Not Available | Cell adhesion |
| 66 | Poly-beta-1, 6-N-acetyl-D-glucosamine N-deacetylase (PGA N-deacetylase) (Poly-beta-1, 6-GlcNAc N-deacetylase) (EC 3.5.1.-) | *Escherichia coli* | Strain K12 | P75906 | 3VUS | 8905232; 9278503; 16738553; 15090514; 18359807 | Hydrolase activity, acting on carbon-nitrogen (but not peptide) bonds, Hydrolase activity, hydrolyzing O-glycosyl compounds | Hydrolase |
| 67 | Diguanylate cyclase DgcZ (DGC) (EC 2.7.7.65) (Zinc-sensory diguanylate cyclase) | *Escherichia coli* | Strain K12 | P31129 | 3TVK | 8383113; 9097039; 9278503; 16738553; 18713317; 19332833; 19460094; 20582742; 26148715; 23769666 | Diguanylate cyclase activity, GTP binding, Identical protein binding, zinc ion binding | Transferase |
| 68 | Acyl-homoserine lactone acylase PvdQ (AHL acylase PvdQ) (Acyl-HSL acylase PvdQ) (EC 3.5.1.97) [Cleaved into: Acyl-homoserine lactone acylase PvdQ subunit alpha (Acyl-HSL acylase PvdQ subunit alpha); Acyl-homoserine lactone acylase PvdQ subunit beta (Acyl-HSL acylase PvdQ subunit beta)] | *Pseudomonas aeruginosa* | Strain ATCC 15692 / DSM 22644 / CIP 104116 / JCM 14847 / LMG 12228 / 1C / PRS 101 / PAO1 | Q9I194 | 3SRA | 10984043; 16495538; 12207696; 14532048; 12686626 | ADP-dependent short-chain-acyl-CoA Hydrolase activity, Hydrolase activity, acting on carbon-nitrogen (but not peptide) bonds, in linear amides, Identical protein binding | Hydrolase |
| 69 | Fap1 adhesin (Adhesin Fap1) (Fimbriae-associated protein Fap1) (Serine-rich repeat protein Fap1) | *Streptococcus parasanguinis* | | A1C3L3 | 3RGU | 10594831; 16997950; 9632253; 20164186; 20584910 | Not Available | Structural protein |
| 70 | Putative fimbrium tip subunit Fim1F | *Parabacteroides distasonis* | Strain ATCC 8503 / DSM 20701 / CIP 104284 / JCM 5825 / NCTC 11152 | A6LHQ9 | 3R4R | 17579514; 26972441; 27062925; 27062925 | Not Available | Cell adhesion |
| 71 | Response regulator | *Xanthomonas campestris pv. campestris* | Strain ATCC 33913 / DSM 3586 / NCPPB 528 / LMG 568 / P 25 | Q8P559 | 3QYY | 12024217; 22120736 | Diguanylate cyclase activity, nucleotide binding | Signaling protein |
| 72 | Platelet binding protein GspB (Adhesin GspB) (Serine-rich adhesin for platelets) (Serine-rich repeat protein GspB) | *Streptococcus gordonii* | | Q939N5 | 3QC5, 3QC6, 5IUC, 6EF7, 6EF9, 6EFA | 12010500; 8366515; 14729688; 15049820; 15489421; 19202081; 20714350; 21531800; 26884191; 21765814; 21765814; 21765814; 21765814 | Not Available | Sugar binding protein |
| 73 | Diguanylate cyclase with PAS sensory domain (EC 2.7.7.65) | *Shewanella oneidensis* | Strain MR-1 | Q8EGB0 | 3MFX | 12368813 | Diguanylate cyclase activity | Transcription |
| 74 | Probable two-component response regulator | *Pseudomonas aeruginosa* | Strain ATCC 15692 / DSM 22644 / CIP 104116 / JCM 14847 / LMG 12228 / 1C / PRS 101 / PAO1 | Q9HXT9 | 3I5C | 10984043; 18366254; 19695263 | Diguanylate cyclase activity, Identical protein binding, nucleotide binding | Signaling protein |
| 75 | mRNA interferase toxin MqsR (EC 3.1.-.-) (Endoribonuclease MqsR) (Motility quorum-sensing regulator MqsR) (Toxin MqsR) | *Escherichia coli* | Strain K12 | Q46865 | 3HI2 | 9278503; 16738553; 16768798; 16352847; 19690171; 19909729; 20105222; 19943910; 21516113; 21788497; 29531044; 23432955; 23289863; 23172222; 25534751; 20041169 | Endoribonuclease activity | DNA binding protein / toxin |
| 76 | Antitoxin MqsA | *Escherichia coli* | Strain K12 | Q46864 | 3HI2 | 9278503; 16738553; 16768798; 19690171; 20105222; 19943910; 21516113; 23432955; 23172222; 24212724; 25534751; 20823526; 20041169; 21068382; 22789559 | Metal ion binding, Sequence-specific DNA binding | DNA binding protein / toxin |
| 77 | Anthraniloyl-CoA anthraniloylTransferase (EC 2.3.1.262) (2-heptyl-4(1H)-quinolone synthase PqsD) (PqsD) | *Pseudomonas aeruginosa* | Strain ATCC 15692 / DSM 22644 / CIP 104116 / JCM 14847 / LMG 12228 / 1C / PRS 101 / PAO1 | P20582 | 3H76, 3H77, 3H78 | 10984043; 2153661; 18728009; 21425231; 22992202; 24239007; 19694421; 19694421; 19694421;19694421 | 3-oxoacyl-[acyl-carrier-protein] synthase activity, acylTransferase activity | Transferase |
| 78 | Diguanylate cyclase | *Geobacter sulfurreducens* | Strain ATCC 51573 / DSM 12127 / PCA | Q74FR4 | 3EZU | 14671304 | Diguanylate cyclase activity | Signaling protein |

| Target ID | Bacterial Protein Name | Organism Name | Strain | UniProt ID | PDB ID | PubMed ID | Classification as per UniProt GO Molecular Function | Classification of Function as per PDB |
|---|---|---|---|---|---|---|---|---|
| 79 | Fucose-binding lectin PA-IIL | *Pseudomonas aeruginosa* | Strain ATCC 15692 / DSM 22644 / CIP 104116 / JCM 14847 / LMG 12228 / 1C / PRS 101 / PAO1 | Q9HYN5 | 3DCQ | 10984043; 12415289; 12909014; 15790314; 15573375; 16438968; 17540045; 17623286; 19101469; 23719508; 27308201; 26898721; 29896342; 29147502; 29163899; 30830745; 31146126 | Carbohydrate binding, Metal ion binding | Sugar binding protein |
| 80 | Pilin (Lancefield T antigen) (Pilus backbone Structural protein) | *Streptococcus pyogenes serotype M1* | | Q9A1S2 | 3B2M, 3GLD, 3GLE | 11296296; 16088826; 17501921; 18063798; 18063798 | Not Available | Structural protein |
| 81 | Biofilm operon icaADBC HTH-type negative Transcriptional regulator IcaR (Intercellular adhesion protein R) | *Staphylococcus epidermidis* | Strain ATCC 35984 / RP62A | Q5HKQ1 | 2ZCM | 8809760; 15774886; 12142410; 18208836 | DNA binding | Transcription |
| 82 | Lipoprotein NlpE (Copper homeostasis protein CutF) | *Escherichia coli* | Strain K12 | P40710 | 2Z4H, 2Z4I | 7635808; 7635807; 9278503; 16738553; 11830644; 15252048; 17698001; 17698001; 17698001 | Identical protein binding | Signaling protein activator |
| 83 | CRISPR-associated endonuclease Cas6 / Csy4 (EC 3.1.-.-) | *Pseudomonas aeruginosa* | Strain UCBPP-PA14 | Q02MM2 | 2XLI, 2XLJ, 2XLK, 4AL5, 4AL6, 4AL7, 5UZ9, 6B44, 6B45, 6B46, 6B47, 6B48, 6NE0 | 17038190; 21398535; 21081758; 21536913; 20829488; 22522703; 20829488; 20829488; 20829488; 22522703; 22522703; 22522703; 28340349; 28985564; 28985564; 28985564; 28985564; 28985564; 30872121 | Endonuclease activity, RNA binding | Immune system / RNA |
| 84 | Autotransporter adhesin SadA (Salmonella adhesin A) (Type 5 secretion system autotransporter SadA) | *Salmonella typhimurium* | Strain LT2 / SGSC1412 / ATCC 700720 | Q8ZL64 | 2WPQ, 2WPR, 2WPS, 2YNY, 2YNZ, 2YO0, 2YO1, 2YO2, 2YO3, 3ZMF | 11677609; 21859856; 24369174; 18093992; 19805097; 23213248; 19805097; 19805097;19805097; 23213248; 23213248; 23213248; 23213248; 23213248; 23213248;18093992 | Not Available | Membrane protein |
| 85 | 2-aminobenzoylacetyl-CoA thioesterase (EC 3.1.2.32) | *Pseudomonas aeruginosa* | Strain ATCC 15692 / DSM 22644 / CIP 104116 / JCM 14847 / LMG 12228 / 1C / PRS 101 / PAO1 | P20581 | 2Q0I, 2Q0J, 2VW8, 3DH8, 5HIO, 5HIP, 5HIQ, 5HIS, 7KGW, 7KGX | 2153661; 10984043; 12426334; 18776012; 25960261; 27851827; 19788310; 20419351; 27082157; 19788310; 19788310; 20419351; 19788310; 27082157; 27082157; 27082157; 27082157; 33793200; 33793200 | Beta-lactamase activity, Metal ion binding | Hydrolase |
| 86 | Probable two-component-system connector protein AriR | *Escherichia coli* | Strain K12 | P75993 | 2OXL | 9278503; 16738553; 15470704; 17511876; 18174134; 19172264; 19240136; 17765265; 17765265 | DNA binding | Gene regulation |

| Target ID | Bacterial Protein Name | Organism Name | Strain | UniProt ID | PDB ID | PubMed ID | Classification as per UniProt GO Molecular Function | Classification of Function as per PDB |
|---|---|---|---|---|---|---|---|---|
| 87 | Transcription antitermination protein RfaH | *Escherichia coli* | Strain K12 | P0AFW0 | 2OUG, 5OND, 6C6S, 6C6T | 1584020; 1379743; 9278503; 16738553; 8606157; 8951819; 9171395; 9426123; 10660066; 12007406; 11983161; 16452414; 17434131; 17434131; 29741479; 29887376; 29887376 | Bacterial-type RNA polymerase core enzyme binding, DNA binding, Transcription antitermination factor activity, DNA binding, translation activator activity | Transcription |
| 88 | High-affinity zinc uptake system binding-protein ZnuA | *Bacillus subtilis* | Strain 168 | O34966 | 2O1E | 9274031; 9384377; 9811636; 12426338; 14730579; 21813502; 22882210 | Metal ion binding | Structural protein |
| 89 | Curli production assembly / transport component CsgE | *Escherichia coli* | Strain K12 | P0AE95 | 2NA4 | 8817489; 8905232; 9278503; 16738553 | Identical protein binding | Chaperone / protein transport |
| 90 | Type-1 fimbrial protein, A chain (Type-1A pilin) | *Escherichia coli* | Strain K12 | P04128 | 2M5G | 2858471; 6147250; 7610040; 9278503; 16738553 | Identical protein binding | Structural protein |
| 91 | Sensor protein QseC (EC 2.7.13.3) | *Escherichia coli* | Strain K12 | P40719 | 2KSE, 3JZ3 | 9278503; 16738553; 11929534; 15919996; 16352847; 20498088; 20594156 | ATP binding, phosphorelay sensor kinase activity | Transferase |
| 92 | Protein FimF | *Escherichia coli* | Strain K12 | P08189 | 2JMR | 2890081; 7610040; 9278503; 16738553; 1970114 | Not Available | Cell adhesion |
| 93 | Competence stimulating peptide (Competence stimulating protein) | *Streptococcus mutans* | | Q99QI5 | 2I2J | 11208787; 16936029 | Pheromone activity | Signaling protein |
| 94 | Toxin YoeB (EC 3.1.-.-) (Putative endoribonuclease YoeB) (Putative mRNA interferase Yoeb) | *Escherichia coli* | Strain K12 | P69348 | 2A6Q, 2A6R, 2A6S, 4V8X, 6NY6 | 9278503; 16738553; 14672926; 15009896; 15980067; 16768798; 17170003; 18854355; 19028895; 19124462; 19400780; 21788497; 29531044; 16109374; 16109374; 16109374; 16109374; 23945936; 31501867 | Endoribonuclease activity, endoribonuclease activity, producing 3'-phosphomonoesters, ribosomal small subunit binding, RNA binding | Ribosome |
| 95 | Antitoxin YefM | *Escherichia coli* | Strain K12 | P69346 | 2A6Q | 9097040; 9278503; 16738553; 6170941; 7567469; 14672926; 15009896; 15980067; 16768798; 17170003; 19028895; 19400780; 16109374; 16109374 | Sequence-specific DNA binding, toxic substance binding | Toxin inhhibitor |
| 96 | Global transcriptional regulator Spx (Redox-responsive Transcription factor Spx) (Suppressor of clpP and clpX) (Spx) | *Bacillus subtilis* | Strain 168 | O31602 | 1Z3E, 3GFK, 3IHQ, 6GHB, 6GHO | 9384377; 11544224; 11703662; 12028382; 12057962; 12642660; 14597697; 15659166; 16885442; 17158660; 17434969; 17908206; 18662407; 18687074; 19074380; 22307755; 23894131; 24417481; 24942655; 29271514; 30001325; 30718304; 30830258; 31655740; 16249335; 19580872; 20084284; 30982633; 16249335; 19580872; 20084284; 30982633; 30982633 | Not Available | Protein binding |

| Target ID | Bacterial Protein Name | Organism Name | Strain | UniProt ID | PDB ID | PubMed ID | Classification as per UniProt GO Molecular Function | Classification of Function as per PDB |
|---|---|---|---|---|---|---|---|---|
| 97 | Carbon storage regulator (Translational dual regulator CsrA) | *Escherichia coli* | Strain K12 | P69913 | 1Y00, 5Z38 | 8393005; 8604133; 9205837; 9278503; 16738553; 7751274; 7493933; 9211896; 11298291; 11741870; 12193630; 12067347; 12694612; 16352847; 16923806; 18223068; 18713317; 19103924; 19460094; 21488981; 21696456; 28851853; 28924029; 25833324; 15866937; 15866937; 29567971 | mRNA 5'-UTR binding | RNA binding protein |
| 98 | Translational regulator CsrA (Carbon storage regulator) (Global Translational regulatory protein RsmA) | *Pseudomonas aeruginosa* | Strain ATCC 15692 / DSM 22644 / CIP 104116 / JCM 14847 / LMG 12228 / 1C / PRS 101 / PAO1 | O69078 | 1VPZ | 11673439; 10984043; 15126453; 16359708; 16104018 | mRNA 5'-UTR binding | RNA binding protein |
| 99 | Endoribonuclease toxin MazF (EC 3.1.27.-) (Toxin MazF) (mRNA interferase MazF) | *Escherichia coli* | Strain K12 | P0AE70 | 1UB4, 3NFC, 5CK9, 5CKB, 5CKD, 5CKE, 5CKF, 5CKH, 5CO7, 5CQX, 5CQY, 5CR2 | 2844820; 8226627; 9278503; 16738553; 8650219; 11071896; 11222603; 12123459; 12972253; 15150257; 15576778; 15316771; 15537630; 15837428; 16390452; 17962566; 18310334; 18854355; 19251848; 19707553; 21944167; 21419338; 21788497; 29531044; 20005847; 22412352; 23432955; 23416055; 23280569; 24375411; 25564525; 19215780; 26395283; 12718874; 16413577; 12718874; 27026704; 27026704; 27026704; 27026704; 27026704; 27026704; 27026704; 27026704; 27026704; 27026704 | DNA binding, endoribonuclease activity, protein homodimerization activity, protein-containing complex binding, RNA binding | Hydrolase |
| 100 | O-acetyl-ADP-ribose deacetylase (EC 3.1.1.106) (Regulator of RNAse III activity) | *Escherichia coli* | Strain K12 | P0A8D6 | 1SPV | 8905232; 9278503; 16738553; 19141481; 21257746; 24267348; 25546632; 28034758; 28582517; 26481419 | Deacetylase activity, endoribonuclease inhibitor activity, enzyme binding, O-acetyl-ADP-ribose deacetylase activity, purine nucleoside binding | Structural protein |
| 101 | Surface composition regulator | *Escherichia coli* | Strain K12 | P26649 | 1RRZ | 1324388; 9205837; 9278503; 16738553; 23537328; 9208918; 15161493 | Not Available | Structural protein |
| 102 | Type IV major pilin protein PilA (Pilin) | *Pseudomonas aeruginosa* | Strain ATCC 15692 / DSM 22644 / CIP 104116 / JCM 14847 / LMG 12228 / 1C / PRS 101 / PAO1 | P04739 | 1PAO | 2997119; 10984043; 8330261; 8845350; 1429457; 9282737; 23266901; 26041805; 27162347; 31431558 | Not Available | Fimbrial protein |
| 103 | AI-2 transport protein TqsA (Transport of quorum-sensing signal protein) | *Escherichia coli* | Strain K12 | P0AFS5 | -- | 9097039; 9278503; 16738553; 15919996; 16385049 | Efflux transmembrane transporter activity | Not Available |
| 104 | HTH-type quorum sensing-dependent Transcriptional regulator VjbR | *Brucella melitensis biotype 1* | Strain 16M / ATCC 23456 / NCTC 10094 | Q8YAY5 | -- | 11756688; 16008582; 17056750; 17557825; 20529360 | DNA binding | Not Available |
| 105 | Sensory / regulatory protein RpfC (EC 2.7.13.3) | *Xanthomonas campestris pv. campestris* | Strain 8004 | P0C0F7 | -- | 11123673; 15899963; 1645442; 12960398; 16611728; 28784774; 16940295; 20231439; 28369120 | ATP binding, phosphorelay sensor kinase activity | Not Available |
| 106 | Exopolyphosphatase (ExopolyPase) (EC 3.6.1.11) (Polyphosphate:ADP phosphoTransferase) (PolyP:ADP phosphoTransferase) (EC 2.7.4.1) | *Pseudomonas aeruginosa* | Strain ATCC 15692 / DSM 22644 / CIP 104116 / JCM 14847 / LMG 12228 / 1C / PRS 101 / PAO1 | Q9ZN70 | -- | 10382967; 10984043; 26576296 | Exopolyphosphatase activity, polyphosphate kinase activity, pyrophosphatase activity | Not Available |

| Target ID | Bacterial Protein Name | Organism Name | Strain | UniProt ID | PDB ID | PubMed ID | Classification as per UniProt GO Molecular Function | Classification of Function as per PDB |
|---|---|---|---|---|---|---|---|---|
| 107 | Two-component response regulator PprB | *Pseudomonas aeruginosa* | Strain ATCC 15692 / DSM 22644 / CIP 104116 / JCM 14847 / LMG 12228 / 1C / PRS 101 / PAO1 | Q9HWA4 | -- | 10984043; 12499175; 15882421; 18684246; 19151143; 21091863; 23209420; 31492668 | DNA binding, phosphorelay response regulator activity | Not Available |
| 108 | Platelet adherence protein A (Adhesin PadA) | *Streptococcus gordonii* | Strain Challis / ATCC 35105 / BCRC 15272 / CH1 / DL1 / V288 | A8AZP4 | -- | 17720781; 8366515; 19884334; 21071690; 24136582; 27616700 | Not Available | Not Available |
| 109 | S-ribosylhomocysteine lyase | *Listeria monocytogenes serovar 1 / 2a* | Strain ATCC BAA-679 / EGD-e | Q8Y7I9 | -- | 12039883; 11679669 | Iron ion binding, S-ribosylhomocysteine lyase activity | Not Available |
| 110 | Adhesin Ata autotransporter (Acinetobacter trimeric autotransporter) (Type 5 secretion system autotransporter Ata) | *Acinetobacter baumannii* | Strain ATCC 17978 / CIP 53.77 / LMG 1025 / NCDC KC755 / 5377 | A3M3H0 | -- | 17344419; 22825448; 22609912 | Not Available | Not Available |
| 111 | Uncharacterized protein | *Desulfitobacterium hafniense* | Strain Y51 | Q24NU6 | -- | 16513756 | Not Available | Not Available |
| 112 | Poly-beta-1, 6-N-acetyl-D-glucosamine N-deacetylase PgaB | *Pseudomonas spp. GM25* | | J2X9J6 | -- | 23045501 | Hydrolase activity, acting on carbon-nitrogen (but not peptide) bonds | Not Available |
| 113 | Uncharacterized protein | *Limosilactobacillus mucosae LM1* | | A0A0D4CKI7 | -- | 22887668 | Not Available | Not Available |
| 114 | Poly-beta-1, 6-N-acetyl-D-glucosamine synthase (Poly-beta-1, 6-Glcc synthase) (EC 2.4.1.-) | *Bordetella pertussis* | Strain Tohama I / ATCC BAA-589 / NCTC 13251 | Q7VX65 | -- | 12910271 | Acetyl glucosaminyl transferase activity | Not Available |
| 115 | Uncharacterized protein | *Bordetella pertussis* | Strain Tohama I / ATCC BAA-589 / NCTC 13251 | Q7VY41 | -- | 12910271 | Not Available | Not Available |
| 116 | Motility quorum-sensing regulator MqsR | *Pseudomonas spp. S35* | | A0A6I6VNC7 | -- | 25398872 | Not Available | Not Available |
| 117 | Type II toxin-antitoxin system MqsR family toxin | *Pseudomonas caspiana* | | A0A502JPJ4 | -- | 31206918 | Not Available | Not Available |
| 118 | Putative hemin storage protein | *Bordetella pertussis* | Strain Tohama I / ATCC BAA-589 / NCTC 13251 | Q7VX64 | -- | 12910271 | Hydrolase activity, acting on carbon-nitrogen (but not peptide) bonds | Not Available |
| 119 | Surface composition regulator | *Klebsiella oxytoca* | Strain ATCC 8724 / DSM 4798 / JCM 20051 / NBRC 3318 / NRRL B-199 / KCTC 1686 | A0A0H3H1S4 | -- | 22493189 | Not Available | Not Available |
| 120 | Uncharacterized protein | *Serpentinomona mccroryi* | | A0A060NNA1 | -- | 24845058 | Not Available | Not Available |
| 121 | Poly-beta-1, 6-N-acetyl-D-glucosamine synthase (Poly-beta-1, 6-GlcNAc synthase) (EC 2.4.1.-) | *Cupriavidus gilardii CR3* | | A0A0M4KYJ3 | -- | 26301592 | Acetyl glucosaminyl transferase activity | Not Available |
| 122 | Poly-beta-1, 6-N-acetyl-D-glucosamine biosynthesis protein PgaD | *Pseudomonas viciae* | | A0A4P7PA43 | -- | 30941113 | Not Available | Not Available |
| 123 | Poly-beta-1, 6-N-acetyl-D-glucosamine biosynthesis protein PgaD | *Sedimenticola spp.* | | A0A7C1YJ10 | -- | 31911466 | Not Available | Not Available |
| 124 | Poly-beta-1, 6-N-acetyl-D-glucosamine synthase (Poly-beta-1, 6-GlcNAc synthase) (EC 2.4.1.-) | *Pseudomonas viciae* | | A0A4P7PA38 | -- | 30941113 | Acetyl glucosaminyl transferase activity | Not Available |
| 125 | Type II toxin-antitoxin system MqsR family toxin | *Pseudomonas gessardii* | | A0A7Y1QPP7 | -- | 32587583 | Not Available | Not Available |
| 126 | Poly-beta-1, 6-N-acetyl-D-glucosamine biosynthesis protein PgaD | *Pseudomonas gessardii* | | A0A7Y1MQN0 | -- | 32587583 | Not Available | Not Available |
| 127 | Poly-beta-1, 6-N-acetyl-D-glucosamine synthase (Poly-beta-1, 6-GlcNAc synthase) (EC 2.4.1.-) | *Sedimenticola spp.* | | A0A7C1YCS2 | -- | 31911466 | Acetyl glucosaminyl transferase activity | Not Available |
| 128 | Poly-beta-1, 6-N-acetyl-D-glucosamine N-deacetylase PgaB (EC 3.5.1.-) | *Pseudomonas gessardii* | | A0A7Y1MR48 | -- | 32587583 | Hydrolase activity, acting on carbon-nitrogen (but not peptide) bonds | Not Available |
| 129 | Uncharacterized protein | *Cupriavidus gilardii CR3* | | A0A0M3SW47 | -- | 26301592 | Not Available | Not Available |
| 130 | Poly-beta-1, 6-N-acetyl-D-glucosamine N-deacetylase PgaB | *Sedimenticola spp.* | | A0A7C1YIT8 | -- | 31911466 | Hydrolase activity, acting on carbon-nitrogen (but not peptide) bonds | Not Available |
| 131 | Poly-beta-1, 6-N-acetyl-D-glucosamine biosynthesis protein PgaD | *Candidatus Desulfofervidus auxilii* | | A0A7C0Y839 | -- | 31911466 | Not Available | Not Available |
| 132 | Putative membrane protein | *Bordetella bronchiseptica 253* | | A0A0C6P5Z7 | -- | 23051057 | Not Available | Not Available |
| 133 | Surface composition regulator | *Klebsiella pneumoniae subspp. pneumoniae* | Strain HS11286 | A0A0H3GYK7 | -- | 22408243 | Not Available | Not Available |
| 134 | Hemin storage system HmsS protein | *Klebsiella pneumoniae subspp. pneumoniae* | Strain HS11286 | A0A0H3GQE1 | -- | 22408243 | Not Available | Not Available |
| 135 | Poly-beta-1, 6-N-acetyl-D-glucosamine synthase (Poly-beta-1, 6-GlcNAc synthase) (EC 2.4.1.-) | *Klebsiella pneumoniae subspp. pneumoniae* | Strain HS11286 | A0A0H3GH44 | -- | 22408243 | Acetyl glucosaminyl transferase activity | Not Available |
| 136 | Uncharacterized protein | *Collinsella aerofaciens* | | A0A6N9JJB5 | -- | 31477907 | Not Available | Not Available |
| 137 | Motility quorum-sensing regulator | *Escherichia coli O17:K52:H18* | Strain UMN026 / ExPEC | B7ND13 | -- | 19165319 | Not Available | Not Available |
| 138 | Poly-beta-1, 6-N-acetyl-D-glucosamine synthase (Poly-beta-1, 6-GlcNAc synthase) (EC 2.4.1.-) | *Pseudomonas spp. CFBP13528* | | A0A4U3IFB2 | -- | 31040301 | Acetyl glucosaminyl transferase activity | Not Available |
| 139 | Outer membrane N-deacetylase | *Pseudomonas spp.* | Strain M1 | W5IKV6 | -- | 10388678; 23405299 | Hydrolase activity, acting on carbon-nitrogen (but not peptide) bonds | Not Available |
| 140 | Polysaccharide deacetylase | *Burkholderia spp. TJI49* | | F0FYV7 | -- | 23653265 | Hydrolase activity, acting on carbon-nitrogen (but not peptide) bonds | Not Available |
| 141 | mRNA interferase MqsR | *Pseudomonas spp.* | Strain M1 | W5IYY9 | -- | 10388678; 23405299 | Not Available | Not Available |
| 142 | Poly-beta-1, 6-N-acetyl-D-glucosamine N-deacetylase PgaB | *Stenotrophomonas spp. ZAC14D1_naIMI4_6* | | A0A3Q8FCT2 | -- | 29765358 | Hydrolase activity, acting on carbon-nitrogen (but not peptide) bonds | Not Available |
| 143 | Poly-beta-1, 6-N-acetyl-D-glucosamine synthase (Poly-beta-1, 6-GlcNAc synthase) (EC 2.4.1.-) | *Stenotrophomonas spp. ZAC14D1_naIMI4_6* | | A0A3Q8F836 | -- | 29765358 | Acetyl glucosaminyl transferase activity | Not Available |
| 144 | Predicted inner membrane protein | *Escherichia coli* | Strain ATCC 9637 / CCM 2024 / DSM 1116 / LMG 11080 / NBRC 13500 / NCIMB 8666 / NRRL B-766 / W | E0IZ23 | -- | 21208457 | Not Available | Not Available |
| 145 | Poly-beta-1, 6-N-acetyl-D-glucosamine synthase (Poly-beta-1, 6-GlcNAc synthase) (EC 2.4.1.-) | *Burkholderia spp. TJI49* | | F0FYV6 | -- | 23653265 | Acetyl glucosaminyl transferase activity | Not Available |
| 146 | Poly-beta-1, 6-N-acetyl-D-glucosamine biosynthesis protein PgaD | *Pseudomonas spp. CFBP13528* | | A0A4U3IG94 | -- | 31040301 | Not Available | Not Available |
| 147 | Uncharacterized protein | *Burkholderia spp. TJI49* | | F0FZG2 | -- | 23653265 | Not Available | Not Available |
| 148 | Uncharacterized protein | *Gordonibacter urolithinfaciens* | | A0A7K0IDQ7 | -- | 31477907 | Not Available | Not Available |
| 149 | Poly-beta-1, 6-N-acetyl-D-glucosamine N-deacetylase PgaB | *Pseudomonas spp. CFBP13528* | | A0A4U3IGQ5 | -- | 31040301 | Hydrolase activity, acting on carbon-nitrogen (but not peptide) bonds | Not Available |

| Target ID | Bacterial Protein Name | Organism Name | Strain | UniProt ID | PDB ID | PubMed ID | Classification as per UniProt GO Molecular Function | Classification of Function as per PDB |
|---|---|---|---|---|---|---|---|---|
| 150 | Uncharacterized protein | *Candidatus thiomargarita nelsonii* | | A0A0A6P4A5 | -- | 27199933 | Not Available | Not Available |
| 151 | Poly-beta-1, 6-N-acetyl-D-glucosamine N-deacetylase | *Pseudomonas spp. S35* | | A0A6I6VSZ2 | -- | 25398872 | Hydrolase activity, acting on carbon-nitrogen (but not peptide) bonds | Not Available |
| 152 | Lipoprotein for biofilm formation | *Bacillus pumilus* | Strain SAFR-032 | A8FF48 | -- | 17895969 | Not Available | Not Available |
| 153 | Poly-beta-1, 6-N-acetyl-D-glucosamine biosynthesis protein PgaD | *Pseudomonas spp. S35* | | A0A6I6VN30 | -- | 25398872 | Not Available | Not Available |
| 154 | Uncharacterized protein | *Candidatus thiomargarita nelsonii* | | A0A0A6P6A5 | -- | 27199933 | Not Available | Not Available |
| 155 | Uncharacterized protein | *Candidatus thiomargarita nelsonii* | | A0A0A6S368 | -- | 27199933 | Not Available | Not Available |
| 156 | Poly-beta-1, 6-N-acetyl-D-glucosamine synthase (Poly-beta-1, 6-GlcNAc synthase) (EC 2.4.1.-) | *Pseudomonas spp. S35* | | A0A6I6VPE8 | -- | 25398872 | Acetyl glucosaminyl transferase activity | Not Available |
| 157 | Uncharacterized protein | *Gamma proteobacterium HTCC5015* | | B5JTY1 | -- | 20472792 | Not Available | Not Available |
| 158 | Uncharacterized protein | *Bacteroidetes bacterium* | | A0A7C5HKG5 | -- | 31911466 | Not Available | Not Available |
| 159 | YcdR | *Gamma proteobacterium HTCC5015* | | B5JTX9 | -- | 20472792 | Hydrolase activity, acting on carbon-nitrogen (but not peptide) bonds | Not Available |
| 160 | Poly-beta-1, 6-N-acetyl-D-glucosamine synthase (Poly-beta-1, 6-GlcNAc synthase) (EC 2.4.1.-) | *Gamma proteobacterium HTCC5015* | | B5JTY0 | -- | 20472792 | Acetyl glucosaminyl transferase activity | Not Available |
| 161 | Poly-beta-1, 6-N-acetyl-D-glucosamine synthase (Poly-beta-1, 6-GlcNAc synthase) (EC 2.4.1.-) | *Pseudomonas gessardii* | | A0A7Y1MQR9 | -- | 32587583 | Acetyl glucosaminyl transferase activity | Not Available |
| 162 | Outer membrane N-deacetylase | *Klebsiella aerogenes* | Strain ATCC 13048 / DSM 30053 / JCM 1235 / KCTC 2190 / NBRC 13534 / NCIMB 10102 / NCTC 10006 | A0A0H3FMM8 | -- | 22493190 | Hydrolase activity, acting on carbon-nitrogen (but not peptide) bonds | Not Available |
| 163 | Toxin-antitoxin system, toxin component | *Nitrosomonadales bacterium* | | A0A658BNF9 | -- | 29662147 | Not Available | Not Available |
| 164 | Type II toxin-antitoxin system MqsR family toxin | *Pseudomonas viciae* | | A0A4P7PLV8 | -- | 30941113 | Not Available | Not Available |
| 165 | Outer membrane N-deacetylase | *Cupriavidus gilardii CR3* | | A0A0M5KLW7 | -- | 26301592 | Hydrolase activity, acting on carbon-nitrogen (but not peptide) bonds | Not Available |
| 166 | Poly-beta-1, 6-N-acetyl-D-glucosamine synthase (Poly-beta-1, 6-GlcNAc synthase) (EC 2.4.1.-) | *Rhodocyclaceae bacterium* | | A0A7C6VQ62 | -- | 32123542 | Acetyl glucosaminyl transferase activity | Not Available |
| 167 | GGDEF family protein | *Vibrio cholerae serotype O1* | Strain ATCC 39315 / El Tor Inaba N16961 | Q9KTK2 | -- | 10952301 | Diguanylate cyclase activity | Not Available |
| 168 | Poly-beta-1, 6-N-acetyl-D-glucosamine synthase (Poly-beta-1, 6-GlcNAc synthase) (EC 2.4.1.-) | *Xenorhabdus stockiae* | | A0A2D0KWC7 | -- | 28993611 | Acetyl glucosaminyl transferase activity | Not Available |
| 169 | Uncharacterized protein | *Lachnospiraceae bacterium* | | A0A3C1WTK7 | -- | 30148503 | Not Available | Not Available |
| 170 | Histidinekinase (EC 2.7.13.3) | *Vibrio cholerae serotype O1* | Strain ATCC 39315 / El Tor Inaba N16961 | Q9KLP6 | -- | 10952301 | Diguanylate cyclase activity, protein histidine kinase activity | Not Available |
| 171 | GGDEF family protein | *Vibrio cholerae serotype O1* | Strain ATCC 39315 / El Tor Inaba N16961 | Q9KM29 | -- | 10952301 | Diguanylate cyclase activity | Not Available |
| 172 | GGDEF family protein | *Vibrio cholerae serotype O1* | Strain ATCC 39315 / El Tor Inaba N16961 | Q9KSA6 | -- | 10952301 | Diguanylate cyclase activity | Not Available |
| 173 | GGDEF family protein | *Vibrio cholerae serotype O1* | Strain ATCC 39315 / El Tor Inaba N16961 | Q9KT76 | -- | 10952301 | Diguanylate cyclase activity | Not Available |
| 174 | GGDEF family protein | *Vibrio cholerae serotype O1* | Strain ATCC 39315 / El Tor Inaba N16961 | Q9KMV1 | -- | 10952301 | Diguanylate cyclase activity | Not Available |
| 175 | GGDEF family protein | *Vibrio cholerae serotype O1* | Strain ATCC 39315 / El Tor Inaba N16961 | Q9KRN7 | -- | 10952301 | Diguanylate cyclase activity | Not Available |
| 176 | GGDEF family protein | *Vibrio cholerae serotype O1* | Strain ATCC 39315 / El Tor Inaba N16961 | Q9KSP0 | -- | 10952301 | Diguanylate cyclase activity, Metal ion binding | Not Available |
| 177 | GGDEF family protein | *Vibrio cholerae serotype O1* | Strain ATCC 39315 / El Tor Inaba N16961 | Q9KS87 | -- | 10952301 | Diguanylate cyclase activity | Not Available |
| 178 | Poly-beta-1, 6-N-acetyl-D-glucosamine N-deacetylase PgaB | *Enterobacter kobei* | | A0A2J0PG21 | -- | 29088362 | Hydrolase activity, acting on carbon-nitrogen (but not peptide) bonds | Not Available |
| 179 | Sensory box / GGDEF family protein | *Vibrio cholerae serotype O1* | Strain ATCC 39315 / El Tor Inaba N16961 | Q9KL11 | -- | 10952301 | Diguanylate cyclase activity | Not Available |
| 180 | Sensory box / GGDEF family protein | *Vibrio cholerae serotype O1* | Strain ATCC 39315 / El Tor Inaba N16961 | Q9KPJ7 | -- | 10952301 | Diguanylate cyclase activity | Not Available |
| 181 | Poly-beta-1, 6-N-acetyl-D-glucosamine biosynthesis protein PgaD | *Xenorhabdus stockiae* | | A0A2D0KWD2 | -- | 28993611 | Not Available | Not Available |
| 182 | GGDEF family protein | *Vibrio cholerae serotype O1* | Strain ATCC 39315 / El Tor Inaba N16961 | Q9KSS1 | -- | 10952301 | Diguanylate cyclase activity | Not Available |
| 183 | GGDEF family protein | *Vibrio cholerae serotype O1* | Strain ATCC 39315 / El Tor Inaba N16961 | Q9KL98 | -- | 10952301 | Diguanylate cyclase activity | Not Available |
| 184 | GGDEF family protein | *Vibrio cholerae serotype O1* | Strain ATCC 39315 / El Tor Inaba N16961 | Q9KT01 | -- | 10952301 | Diguanylate cyclase activity | Not Available |
| 185 | Poly-beta-1, 6-N-acetyl-D-glucosamine biosynthesis protein PgaD | *Lachnospiraceae bacterium* | | A0A3D0DPQ1 | -- | 30148503 | Not Available | Not Available |
| 186 | GGDEF family protein | *Vibrio cholerae serotype O1* | Strain ATCC 39315 / El Tor Inaba N16961 | Q9KNN4 | -- | 10952301 | Diguanylate cyclase activity | Not Available |
| 187 | GGDEF family protein | *Vibrio cholerae serotype O1* | Strain ATCC 39315 / El Tor Inaba N16961 | Q9KM32 | -- | 10952301 | Diguanylate cyclase activity | Not Available |
| 188 | Poly-beta-1, 6-N-acetyl-D-glucosamine synthase (Poly-beta-1, 6-GlcNAc synthase) (EC 2.4.1.-) | *Pseudomonas spp.* | | A0A3D4NT76 | -- | 30148503 | Acetyl glucosaminyl transferase activity | Not Available |
| 189 | mRNA interferase MqsR | *Pseudomonas spp.* | | A0A3D0JRX8 | -- | 30148503 | Not Available | Not Available |
| 190 | Surface composition regulator | *Salmonella agona* | Strain SL483 | B5F692 | -- | 21602358 | Not Available | Not Available |
| 191 | mRNA interferase MqsR | *Pseudomonas spp.* | | A0A355RA73 | -- | 30148503 | Not Available | Not Available |
| 192 | Type II toxin-antitoxin system MqsR family toxin | *Pseudomonas spp.* | | A0A3D1VUK6 | -- | 30148503 | Not Available | Not Available |
| 193 | Poly-beta-1, 6-N-acetyl-D-glucosamine biosynthesis protein PgaD | *Pseudomonas spp.* | | A0A3D4NT61 | -- | 30148503 | Not Available | Not Available |
| 194 | Poly-beta-1, 6-N-acetyl-D-glucosamine synthase (Poly-beta-1, 6-GlcNAc synthase) (EC 2.4.1.-) | *Xenorhabdus budapestensis* | | A0A2D0J1F4 | -- | 28993611 | Acetyl glucosaminyl transferase activity | Not Available |
| 195 | Poly-beta-1, 6-N-acetyl-D-glucosamine biosynthesis protein PgaD | *Priestia megaterium* | Bacillus megaterium | A0A2T7JNY8 | -- | 31228823 | Not Available | Not Available |
| 196 | Hypothetical lipoprotein ycdR | *Escherichia coli O6:H1* | Strain CFT073 / ATCC 700928 / UPEC | A0A0H2V5Y9 | -- | 12471157 | Hydrolase activity, acting on carbon-nitrogen (but not peptide) bonds | Not Available |
| 197 | Putative polysaccharide deacetylase associated with biofilm formation putative lipoprotein (PgaB) | *Ralstonia solanacearum CMR15* | | D8NG92 | -- | 20550686 | Hydrolase activity, acting on carbon-nitrogen (but not peptide) bonds | Not Available |
| 198 | Type II toxin-antitoxin system MqsR family toxin | *Desulfovibrio spp.* | | A0A3D4GJY3 | -- | 30148503 | Not Available | Not Available |
| 199 | Uncharacterized protein | *Serpentinomonas raichei* | | A0A060NIR8 | -- | 24845058 | Not Available | Not Available |
| 200 | Putative transmembrane protein | *Ralstonia solanacearum CMR15* | | D8NG94 | -- | 20550686 | Not Available | Not Available |
| 201 | Poly-beta-1, 6-N-acetyl-D-glucosamine synthase (Poly-beta-1, 6-GlcNAc synthase) (EC 2.4.1.-) | *Ralstonia solanacearum CMR15* | | D8NG93 | -- | 20550686 | Acetyl glucosaminyl transferase activity | Not Available |
| 202 | Poly-beta-1, 6-N-acetyl-D-glucosamine synthase (Poly-beta-1, 6-GlcNAc synthase) (EC 2.4.1.-) | *Escherichia coli O6:H1* | Strain CFT073 / ATCC 700928 / UPEC | A0A0H2V816 | -- | 12471157 | Acetyl glucosaminyl transferase activity | Not Available |

| Target ID | Bacterial Protein Name | Organism Name | Strain | UniProt ID | PDB ID | PubMed ID | Classification as per UniProt GO Molecular Function | Classification of Function as per PDB |
|---|---|---|---|---|---|---|---|---|
| 203 | Putative polysaccharide deacetylase | *Klebsiella pneumoniae subspp. pneumoniae* | Strain ATCC 700721 / MGH 78578 | A6TH34 | -- | 11677609 | Hydrolase activity, acting on carbon-nitrogen (but not peptide) bonds | Not Available |
| 204 | Poly-beta-1, 6-N-acetyl-D-glucosamine synthase (Poly-beta-1, 6-GlcNAc synthase) (EC 2.4.1.-) | *Klebsiella pneumoniae subspp. pneumoniae* | Strain ATCC 700721 / MGH 78578 | A6TH33 | -- | 11677609 | Acetyl glucosaminyl transferase activity | Not Available |
| 205 | Poly-beta-1, 6-N-acetyl-D-glucosamine synthase (Poly-beta-1, 6-GlcNAc synthase) (EC 2.4.1.-) | *Tetragenococcus halophilus* | Pediococcus halophilus | A0A3G5FLQ7 | -- | 21357458 | Acetyl glucosaminyl transferase activity | Not Available |
| 206 | Uncharacterized protein | *Hydrogenophaga spp.* | | A0A350SRA6 | -- | 30148503 | Not Available | Not Available |
| 207 | Uncharacterized protein | *Escherichia coli O6:H1* | Strain CFT073 / ATCC 700928 / UPEC | A0A0H2V5Z5 | -- | 12471157 | Not Available | Not Available |
| 208 | Surface composition regulator | *Edwardsiella ictaluri* | Strain 93-146 | C5B9Z2 | -- | 22247535 | Not Available | Not Available |
| 209 | Poly-beta-1, 6-N-acetyl-D-glucosamine N-deacetylase | *Tatumella spp. OPLPL6* | | A0A2G8BYL8 | -- | 29067021 | Hydrolase activity, acting on carbon-nitrogen (but not peptide) bonds | Not Available |
| 210 | GGDEF domain-containing protein | *Rhodopirellula baltica* | Strain DSM 10527 / NCIMB 13988 / SH1 | Q7UJ25 | -- | 12835416 | Diguanylate cyclase activity | Not Available |
| 211 | Poly-beta-1, 6-N-acetyl-D-glucosamine synthase (Poly-beta-1, 6-GlcNAc synthase) (EC 2.4.1.-) | *Tatumella spp. OPLPL6* | | A0A2G8BYM4 | -- | 29067021 | Acetyl glucosaminyl transferase activity | Not Available |
| 212 | Probable two-component response regulator | *Rhodopirellula baltica* | Strain DSM 10527 / NCIMB 13988 / SH1 | Q7URT2 | -- | 12835416 | Diguanylate cyclase activity | Not Available |
| 213 | GGDEF family protein | *Rhodopirellula baltica* | Strain DSM 10527 / NCIMB 13988 / SH1 | Q7UIF2 | -- | 12835416 | Diguanylate cyclase activity | Not Available |
| 214 | Uncharacterized protein | *Rhodopirellula baltica* | Strain DSM 10527 / NCIMB 13988 / SH1 | Q7UWZ9 | -- | 12835416 | Diguanylate cyclase activity | Not Available |
| 215 | Poly-beta-1, 6-N-acetyl-D-glucosamine N-deacetylase PgaB | *Acinetobacter seifertii* | | N8SCM4 | -- | 25563912 | Hydrolase activity, acting on carbon-nitrogen (but not peptide) bonds | Not Available |
| 216 | Surface composition regulator | *Salmonella enterica subspp. enterica serovar* | Urbana Str. R8-2977 | G5S153 | -- | 21859443 | Not Available | Not Available |
| 217 | Poly-beta-1, 6-N-acetyl-D-glucosamine synthase (Poly-beta-1, 6-GlcNAc synthase) (EC 2.4.1.-) | *Mammaliicoccus fleurettii* | | A0A3A0I0Q2 | -- | 28066335 | Acetyl glucosaminyl transferase activity | Not Available |
| 218 | Poly-beta-1, 6-N-acetyl-D-glucosamine synthase (Poly-beta-1, 6-GlcNAc synthase) (EC 2.4.1.-) | *Aerococcus urinaehominis* | | A0A109RGK8 | -- | 27103727 | Acetyl glucosaminyl transferase activity | Not Available |
| 219 | Poly-beta-1, 6-N-acetyl-D-glucosamine synthase (Poly-beta-1, 6-GlcNAc synthase) (EC 2.4.1.-) | *Stenotrophomonas geniculata N1* | | A0A0L8AEC2 | -- | 22689240 | Acetyl glucosaminyl transferase activity | Not Available |
| 220 | Uncharacterized protein | *Candidatus Fermentibacter daniensis* | | A0A163WHS5 | -- | 27058503 | Not Available | Not Available |
| 221 | Hemin storage protein | *Stenotrophomonas geniculata N1* | | A0A0L8AES1 | -- | 22689240 | Not Available | Not Available |
| 222 | Uncharacterized protein | *Novosphingobium nitrogenifigens DSM 19370* | | F1ZCU9 | -- | 22156397 | Not Available | Not Available |
| 223 | Surface composition regulator | *Salmonella dublin* | | A0A3T3IH62 | -- | 30286803 | Not Available | Not Available |
| 224 | Poly-beta-1, 6-N-acetyl-D-glucosamine N-deacetylase PgaB | *Stenotrophomonas spp. YAU14A_MKIMI4_1* | | A0A7U5WKB3 | -- | 29765358 | Hydrolase activity, acting on carbon-nitrogen (but not peptide) bonds | Not Available |
| 225 | Poly-beta-1, 6-N-acetyl-D-glucosamine synthase (Poly-beta-1, 6-GlcNAc synthase) (EC 2.4.1.-) | *Pseudomonas proteolytica* | | A0A7Y1GX08 | -- | 32587583 | Acetyl glucosaminyl transferase activity | Not Available |
| 226 | mRNA interferase MqsR | *Lautropia spp. SCN 69-89* | | A0A1D2SFM5 | -- | 26031303 | Not Available | Not Available |
| 227 | Poly-beta-1, 6-N-acetyl-D-glucosamine biosynthesis protein PgaD | *Lautropia spp. SCN 69-89* | | A0A1D2SJT2 | -- | 26031303 | Not Available | Not Available |
| 228 | Poly-beta-1, 6-N-acetyl-D-glucosamine synthase (Poly-beta-1, 6-GlcNAc synthase) (EC 2.4.1.-) | *Pseudomonas proteolytica* | | A0A7Y1E338 | -- | 32587583 | Acetyl glucosaminyl transferase activity | Not Available |
| 229 | Poly-beta-1, 6-N-acetyl-D-glucosamine biosynthesis protein PgaD | *Stenotrophomonas spp. YAU14A_MKIMI4_1* | | A0A7U6AE53 | -- | 29765358 | Not Available | Not Available |
| 230 | Poly-beta-1, 6-N-acetyl-D-glucosamine biosynthesis protein PgaD | *Candidatus Accumulibacter spp.* | | A0A3D3NIE8 | -- | 30148503 | Not Available | Not Available |
| 231 | Surface composition regulator | *Pectobacterium peruviense* | | A0A0J5XXP7 | -- | 26251497 | DNA binding, DNA-directed DNA polymerase activity | Not Available |
| 232 | CDA1 protein | *Mannheimia succiniciproducens* | Strain MBEL55E | Q65TJ0 | -- | 15378067 | Hydrolase activity, acting on carbon-nitrogen (but not peptide) bonds | Not Available |
| 233 | Poly-beta-1, 6-N-acetyl-D-glucosamine synthase (Poly-beta-1, 6-GlcNAc synthase) (EC 2.4.1.-) | *Pectobacterium peruviense* | | A0A0J5XL48 | -- | 26251497 | Acetyl glucosaminyl transferase activity | Not Available |
| 234 | mRNA interferase MqsR | *Candidatus Accumulibacter spp.* | | A0A3D6DTJ4 | -- | 30148503 | Not Available | Not Available |
| 235 | Poly-beta-1, 6-N-acetyl-D-glucosamine synthase (Poly-beta-1, 6-GlcNAc synthase) (EC 2.4.1.-) | *Burkholderia multivorans CGD2* | | B9BJ73 | -- | 23105085 | Acetyl glucosaminyl transferase activity | Not Available |
| 236 | Hemin storage protein | *Pectobacterium peruviense* | | A0A0J5XP71 | -- | 26251497 | Not Available | Not Available |
| 237 | Uncharacterized protein | *Lautropia spp. SCN 69-89* | | A0A1D2SI36 | -- | 26031303 | Not Available | Not Available |
| 238 | Poly-beta-1, 6-N-acetyl-D-glucosamine N-deacetylase (EC 3.5.1.-) | *Aeromona salmonicida subspp. pectinolytica 34mel* | | T0PMK3 | -- | 29304740 | Hydrolase activity, acting on carbon-nitrogen (but not peptide) bonds | Not Available |
| 239 | Type II toxin-antitoxin system MqsR family toxin | *Nitrospirae bacterium* | | A0A7C0YTH1 | -- | 31911466 | Not Available | Not Available |
| 240 | Poly-beta-1, 6-N-acetyl-D-glucosamine synthase (Poly-beta-1, 6-GlcNAc synthase) (EC 2.4.1.-) | *Pseudomonas spp. GM30* | | W6VHT4 | -- | 23045501 | Acetyl glucosaminyl transferase activity | Not Available |
| 241 | Poly-beta-1, 6-N-acetyl-D-glucosamine synthase (Poly-beta-1, 6-GlcNAc synthase) (EC 2.4.1.-) | *Aeromona salmonicida subspp. pectinolytica 34mel* | | T0PR46 | -- | 29304740 | Acetyl glucosaminyl transferase activity | Not Available |
| 242 | Poly-beta-1, 6-N-acetyl-D-glucosamine biosynthesis protein PgaD | *Burkholderiales bacterium* | | A0A3A0ELI9 | -- | 30199629 | Not Available | Not Available |
| 243 | Poly-beta-1, 6-N-acetyl-D-glucosamine N-deacetylase PgaB | *Acinetobacter baumannii* | | A0A5P1UIB3 | -- | 25136007 | Hydrolase activity, acting on carbon-nitrogen (but not peptide) bonds | Not Available |
| 244 | Poly-beta-1, 6-N-acetyl-D-glucosamine synthase (Poly-beta-1, 6-GlcNAc synthase) (EC 2.4.1.-) | *Burkholderiales bacterium* | | A0A3A0EKV6 | -- | 30199629 | Acetyl glucosaminyl transferase activity | Not Available |
| 245 | Poly-beta-1, 6-N-acetyl-D-glucosamine N-deacetylase PgaB | *Burkholderiales bacterium* | | A0A3A0ES14 | -- | 30199629 | Hydrolase activity, acting on carbon-nitrogen (but not peptide) bonds | Not Available |
| 246 | Poly-beta-1, 6-N-acetyl-D-glucosamine N-deacetylase PgaB | *Acinetobacter baumannii* | | A0A7Z2AB22 | -- | 27917170 | Hydrolase activity, acting on carbon-nitrogen (but not peptide) bonds | Not Available |
| 247 | Poly-beta-1, 6-N-acetyl-D-glucosamine synthase (Poly-beta-1, 6-GlcNAc synthase) (EC 2.4.1.-) | *Xanthomonadaceae bacterium SCN 69-320* | | A0A1E4K406 | -- | 26031303 | Acetyl glucosaminyl transferase activity | Not Available |
| 248 | Poly-beta-1, 6-N-acetyl-D-glucosamine biosynthesis protein PgaD | *Acinetobacter baumannii* | | A0A7H1PNT2 | -- | 27917170 | Not Available | Not Available |
| 249 | Poly-beta-1, 6-N-acetyl-D-glucosamine N-deacetylase PgaB | *Xanthomonadaceae bacterium SCN 69-320* | | A0A1E4K488 | -- | 26031303 | Hydrolase activity, acting on carbon-nitrogen (but not peptide) bonds | Not Available |
| 250 | Poly-beta-1, 6-N-acetyl-D-glucosamine biosynthesis protein PgaD | *Xanthomonadaceae bacterium SCN 69-320* | | A0A1E4K403 | -- | 26031303 | Not Available | Not Available |
| 251 | Surface composition regulator | *Salmonella virchow* | | A0A3Y8PE59 | -- | 30286803 | Not Available | Not Available |
| 252 | S-ribosylhomocysteine lyase (EC 4.4.1.21) (AI-2 synthesis protein) (Autoinducer-2 production protein LuxS) | *Streptococcus anginosus* | | Q4PLF8 | -- | 16897563 | Iron ion binding, S-ribosylhomocysteine lyase activity | Not Available |
| 253 | Surface composition regulator | *Salmonella enterica subspp. enterica serovar* | London | A0A5J1SKH1 | -- | 30286803 | Not Available | Not Available |

| Target ID | Bacterial Protein Name | Organism Name | Strain | UniProt ID | PDB ID | PubMed ID | Classification as per UniProt GO Molecular Function | Classification of Function as per PDB |
|---|---|---|---|---|---|---|---|---|
| 254 | DUF1190 domain-containing protein (Lipoprotein) | *Xanthomonas campestris* | | A0A6N3X0V8 | -- | 32966572 | Not Available | Not Available |
| 255 | Surface composition regulator | *Salmonella enterica subspp. enterica serovar* | Stanley | A0A486WXB0 | -- | 30286803; 31099590 | Not Available | Not Available |
| 256 | Surface composition regulator | *Serratia marcescens* | | A0A0G3STT8 | -- | 28636609 | Not Available | Not Available |
| 257 | Uncharacterized protein | *Pseudomonas spp.* | Strain GM41 2012 | W6VMB4 | -- | 23045501 | Not Available | Not Available |
| 258 | Poly-beta-1, 6-N-acetyl-D-glucosamine N-deacetylase PgaB (EC 3.5.1.-) | *Pseudomonas proteolytica* | | A0A7Y1H7P0 | -- | 32587583 | Hydrolase activity, acting on carbon-nitrogen (but not peptide) bonds | Not Available |
| 259 | Poly-beta-1, 6-N-acetyl-D-glucosamine N-deacetylase PgaB (EC 3.5.1.-) | *Pseudomonas proteolytica* | | A0A7Y1DEZ8 | -- | 32587583 | Hydrolase activity, acting on carbon-nitrogen (but not peptide) bonds | Not Available |
| 260 | Poly-beta-1, 6-N-acetyl-D-glucosamine synthase (Poly-beta-1, 6-GlcNAc synthase) (EC 2.4.1.-) | *Shigella spp.* | | A0A351N4J1 | -- | 30148503 | Acetyl glucosaminyl transferase activity | Not Available |
| 261 | Type II toxin-antitoxin system MqsR family toxin | *Pseudomonas spp. WS 5011* | | A0A7Y1BLX1 | -- | 32587583 | Not Available | Not Available |
| 262 | Poly-beta-1, 6-N-acetyl-D-glucosamine synthase (Poly-beta-1, 6-GlcNAc synthase) (EC 2.4.1.-) | *Klebsiella pneumoniae* | | A0A2L0KKC0 | -- | 29437920 | Acetyl glucosaminyl transferase activity | Not Available |
| 263 | mRNA interferase MqsR of the MqsA-MqsR antitoxin / toxin complex and DNA-binding Transcriptionalrepressor (EC 3.1.-.-) | *Pseudomonas putida* | Strain ATCC 47054 / DSM 6125 / CFBP 8728 / NCIMB 11950 / KT2440 | Q88F93 | -- | 12534463 | DNA binding, Hydrolase activity | Not Available |
| 264 | Hemin storage protein | *Stenotrophomonas maltophilia MF89* | | T5KKA0 | -- | 24092793 | Not Available | Not Available |
| 265 | Poly-beta-1, 6-N-acetyl-D-glucosamine biosynthesis protein PgaD | *Pseudomonas orientalis* | | A0A2L0RPX3 | -- | 29479340 | Not Available | Not Available |
| 266 | Hemin storage system protein | *Yersinia entomophaga* | | A0A3S6EX35 | -- | 27187466 | Not Available | Not Available |
| 267 | Uncharacterized protein | *Lentilactobacillus diolivorans DSM 14421* | | A0A0R1SPG5 | -- | 26415554 | Not Available | Not Available |
| 268 | Uncharacterized protein | *Gammaproteobacteria bacterium RBG_16_51_14* | | A0A1G0FW62 | -- | 27774985 | Not Available | Not Available |
| 269 | Poly-beta-1, 6-N-acetyl-D-glucosamine synthase (Poly-beta-1, 6-GlcNAc synthase) (EC 2.4.1.-) | *Burkholderia multivorans CGD1* | | B9B6W1 | -- | 23105085 | Acetyl glucosaminyl transferase activity | Not Available |
| 270 | Type II toxin-antitoxin system MqsR family toxin | *Companilactobacillus salsicarnum* | | A0A646JPD0 | -- | 31668878 | Not Available | Not Available |
| 271 | Outer membrane N-deacetylase | *Yersinia entomophaga* | | A0A3S6EYN4 | -- | 27187466 | Hydrolase activity, acting on carbon-nitrogen (but not peptide) bonds | Not Available |
| 272 | Biofilm PGA synthesis lipoprotein PgaB | *Xanthomonas oryzae pv. oryzicola* | Strain BLS256 | G7TD27 | -- | 21784931 | Hydrolase activity, acting on carbon-nitrogen (but not peptide) bonds | Not Available |
| 273 | Biofilm PGA synthesis lipoprotein PgaB (EC 3.-.-.-) | *Burkholderia multivorans CGD1* | | B9B6W2 | -- | 23105085 | Hydrolase activity, acting on carbon-nitrogen (but not peptide) bonds | Not Available |
| 274 | Uncharacterized protein | *Pelagibacterium spp. SCN 64-44* | | A0A1E4G1S5 | -- | 26031303 | Not Available | Not Available |
| 275 | Biofilm PGA synthesis protein PgaD | *Enterobacter cloacae subspp. cloacae* | Strain ATCC 13047 / DSM 30054 / NBRC 13535 / NCTC 10005 / WDCM 00083 / NCDC 279-56 | A0A0H3CRL8 | -- | 20207761 | Not Available | Not Available |
| 276 | Putative membrane protein | *Streptomyces coelicolor* | Strain ATCC BAA-471 / A3 2 / M145 | Q9AD78 | -- | 12000953 | Diguanylate cyclase activity | Not Available |
| 277 | Poly-beta-1, 6-N-acetyl-D-glucosamine N-deacetylase | *Pseudomonas orientalis* | | A0A2L0RPV9 | -- | 29479340 | Hydrolase activity, acting on carbon-nitrogen (but not peptide) bonds | Not Available |
| 278 | Uncharacterized protein | *Burkholderia multivorans CGD1* | | B9B078 | -- | 23105085 | Not Available | Not Available |
| 279 | Uncharacterized protein | *Levilactobacillus koreensis JCM 16448* | | A0A0R1KV26 | -- | 26415554 | Not Available | Not Available |
| 280 | Poly-beta-1, 6-N-acetyl-D-glucosamine synthase (Poly-beta-1, 6-GlcNAc synthase) (EC 2.4.1.-) | *Stenotrophomonas maltophilia* | Pseudomonas maltophilia Xanthomonas maltophilia | A0A2J0SNY9 | -- | 29170656 | Acetyl glucosaminyl transferase activity | Not Available |
| 281 | Biofilm PGA synthesis lipoprotein pgaB | *Acinetobacter spp. HA* | | I4ZWY1 | -- | 22933775 | Hydrolase activity, acting on carbon-nitrogen (but not peptide) bonds | Not Available |
| 282 | mRNA interferase MqsR | *Photorhabdus spp. S9-53* | | A0A329WFL1 | -- | 29877789 | Not Available | Not Available |
| 283 | Poly-beta-1, 6-N-acetyl-D-glucosamine N-deacetylase PgaB | *Candidatus Manganitrophus noduliformans* | | A0A7X6DPV0 | -- | 32669693 | Hydrolase activity, acting on carbon-nitrogen (but not peptide) bonds | Not Available |
| 284 | Poly-beta-1, 6-N-acetyl-D-glucosamine synthase (Poly-beta-1, 6-GlcNAc synthase) (EC 2.4.1.-) | *Schinkia azotoformans LMG 9581* | | K6DNV1 | -- | 23087684 | Acetyl glucosaminyl transferase activity | Not Available |
| 285 | Poly-beta-1, 6-N-acetyl-D-glucosamine synthase (Poly-beta-1, 6-GlcNAc synthase) (EC 2.4.1.-) | *Oceanospirillales bacterium* | | A0A7V2SRN9 | -- | 31911466 | Acetyl glucosaminyl transferase activity | Not Available |
| 286 | Poly-beta-1, 6-N-acetyl-D-glucosamine synthase (Poly-beta-1, 6-GlcNAc synthase) (EC 2.4.1.-) | *Candidatus Manganitrophus noduliformans* | | A0A7X6DPT7 | -- | 32669693 | Acetyl glucosaminyl transferase activity | Not Available |
| 287 | Surface composition regulator | *Salmonella typhimurium* | Strain D23580 | A0A6C7IEY4 | -- | 19901036 | Not Available | Not Available |
| 288 | Uncharacterized protein | *Acinetobacter baumannii TYTH-1* | | A0A7U4BP68 | -- | 23209228 | Not Available | Not Available |
| 289 | Poly-beta-1, 6-N-acetyl-D-glucosamine N-deacetylase PgaB | *Enterobacter chengduensis* | | A0A5C1C483 | -- | 30302649 | Hydrolase activity, acting on carbon-nitrogen (but not peptide) bonds | Not Available |
| 290 | Poly-beta-1, 6-N-acetyl-D-glucosamine synthase (Poly-beta-1, 6-GlcNAc synthase) (EC 2.4.1.-) | *Priestia megaterium* | Strain DSM 319 / IMG 1521 Bacillus megaterium | D5DKD8 | -- | 21705586 | Acetyl glucosaminyl transferase activity | Not Available |
| 291 | Poly-beta-1, 6-N-acetyl-D-glucosamine synthase (Poly-beta-1, 6-GlcNAc synthase) (EC 2.4.1.-) | *Acinetobacter baumannii TYTH-1* | | A0A7U4BPK8 | -- | 23209228 | Acetyl glucosaminyl transferase activity | Not Available |
| 292 | Xylanase / chitin deacetylase | *Acinetobacter baumannii TYTH-1* | | A0A7U4F894 | -- | 23209228 | Hydrolase activity, acting on carbon-nitrogen (but not peptide) bonds, Hydrolase activity, acting on glycosyl bonds | Not Available |
| 293 | Lipoprotein HmsF | *Acinetobacter baumannii TYTH-1* | | A0A7U4F6K9 | -- | 23209228 | Hydrolase activity, acting on carbon-nitrogen (but not peptide) bonds | Not Available |
| 294 | Uncharacterized protein | *Priestia megaterium* | Strain DSM 319 / IMG 1521 Bacillus megaterium | D5DKD7 | -- | 21705586 | Not Available | Not Available |
| 295 | Poly-beta-1, 6-N-acetyl-D-glucosamine biosynthesis protein PgaD | *Acinetobacter baumannii TYTH-1* | | A0A7U4BLE2 | -- | 23209228 | Not Available | Not Available |
| 296 | Poly-beta-1, 6-N-acetyl-D-glucosamine synthase (Poly-beta-1, 6-GlcNAc synthase) (EC 2.4.1.-) | *Acinetobacter baumannii TYTH-1* | | A0A7U4BLC4 | -- | 23209228 | Acetyl glucosaminyl transferase activity | Not Available |
| 297 | Poly-beta-1, 6-N-acetyl-D-glucosamine biosynthesis protein PgaD (Fragment) | *Guyparkeria spp. SCN-R1* | | A0A426QDS2 | -- | 24673239 | Not Available | Not Available |
| 298 | Poly-beta-1, 6-N-acetyl-D-glucosamine synthase (Poly-beta-1, 6-GlcNAc synthase) (EC 2.4.1.-) | *Lentisphaerae bacterium GWF2_52_8* | | A0A1G0ZGS6 | -- | 27774985 | Acetyl glucosaminyl transferase activity | Not Available |

| Target ID | Bacterial Protein Name | Organism Name | Strain | UniProt ID | PDB ID | PubMed ID | Classification as per UniProt GO Molecular Function | Classification of Function as per PDB |
|---|---|---|---|---|---|---|---|---|
| 299 | Predicted enzyme associated with biofilm formation | *Escherichia coli* | Strain ATCC 9637 / CCM 2024 / DSM 1116 / LMG 11080 / NBRC 13500 / NCIMB 8666 / NRRL B-766 / W | E0IZ21 | -- | 21208457 | Hydrolase activity, acting on carbon-nitrogen (but not peptide) bonds | Not Available |
| 300 | Surface composition regulator | *Escherichia coli* | Strain ATCC 9637 / CCM 2024 / DSM 1116 / LMG 11080 / NBRC 13500 / NCIMB 8666 / NRRL B-766 / W | E0IVV8 | -- | 21208457 | Not Available | Not Available |
| 301 | Uncharacterized protein | *Collinsella aerofaciens* | | A0A6N9JPD9 | -- | 31477907 | Not Available | Not Available |
| 302 | Poly-beta-1, 6-N-acetyl-D-glucosamine synthase (Poly-beta-1, 6-GlcNAc synthase) (EC 2.4.1.-) | *Pseudomonas spp.* | Strain M1 | W5IMC6 | -- | 10388678; 23405299 | Acetyl glucosaminyl transferase activity | Not Available |
| 303 | Poly-beta-1, 6-N-acetyl-D-glucosamine synthase (Poly-beta-1, 6-GlcNAc synthase) (EC 2.4.1.-) | *Escherichia coli* | Strain ATCC 9637 / CCM 2024 / DSM 1116 / LMG 11080 / NBRC 13500 / NCIMB 8666 / NRRL B-766 / W | E0IZ22 | -- | 21208457 | Acetyl glucosaminyl transferase activity | Not Available |
| 304 | Poly-beta-1, 6-N-acetyl-D-glucosamine biosynthesis protein PgaD | *Stenotrophomonas spp. ZAC14D1_naIMI4_6* | | A0A3Q8FAH5 | -- | 29765358 | Not Available | Not Available |
| 305 | Poly-beta-1, 6-N-acetyl-D-glucosamine biosynthesis protein PgaD | *Stenotrophomonas spp. ZAC14D1_naIMI4_6* | | A0A3Q8F8L9 | -- | 29765358 | Not Available | Not Available |
| 306 | Poly-beta-1, 6-N-acetyl-D-glucosamine synthase (Poly-beta-1, 6-GlcNAc synthase) (EC 2.4.1.-) | *Stenotrophomonas spp. ZAC14D1_naIMI4_6* | | A0A3Q8FBD5 | -- | 29765358 | Acetyl glucosaminyl transferase activity | Not Available |
| 307 | Type II toxin-antitoxin system MqsR family toxin | *Pseudomonas spp. WS 5414* | | A0A7Y1CDF1 | -- | 32587583 | Not Available | Not Available |
| 308 | Uncharacterized protein | *Zetaproteobacteria bacterium CG2_30_46_52* | | A0A1J5IY78 | -- | 27112493 | Not Available | Not Available |
| 309 | Poly-beta-1, 6-N-acetyl-D-glucosamine N-deacetylase | *Leclercia spp. LSNIH1* | | A0A2I8NTT5 | -- | 29437920 | Hydrolase activity, acting on carbon-nitrogen (but not peptide) bonds | Not Available |
| 310 | Biofilm PGA synthesis lipoprotein PgaB (EC 3.-.-.-) | *Pseudomonas fluorescens* | Strain ATCC BAA-477 / NRRL B-23932 / Pf-5 | Q4KKC4 | -- | 15980861 | Hydrolase activity, acting on carbon-nitrogen (but not peptide) bonds | Not Available |
| 311 | Poly-beta-1, 6-N-acetyl-D-glucosamine synthase (Poly-beta-1, 6-GlcNAc synthase) (EC 2.4.1.-) | *Pseudomonas fluorescens* | Strain ATCC BAA-477 / NRRL B-23932 / Pf-5 | Q4KKC3 | -- | 15980861 | Acetyl glucosaminyl transferase activity | Not Available |
| 312 | Uncharacterized protein | *Yersinia pseudotuberculosis serotype I* | Strain IP32953 | Q666X0 | -- | 15358858 | Not Available | Not Available |
| 313 | Uncharacterized protein | *Thiotrichales bacterium SG8_50* | | A0A0S8EKY3 | -- | 25922666 | Not Available | Not Available |
| 314 | Motility quorum-sensing regulator MqsR | *Pseudomonas fluorescens* | Strain ATCC BAA-477 / NRRL B-23932 / Pf-5 | Q4KHU9 | -- | 15980861 | Not Available | Not Available |
| 315 | Type II toxin-antitoxin system MqsR family toxin | *Methylosinus sporium* | | A0A2U1STG6 | -- | 29725720 | Not Available | Not Available |
| 316 | Poly-beta-1, 6-N-acetyl-D-glucosamine N-deacetylase PgaB (Fragment) | *Planctomycetaceae bacterium* | | A0A3B8ZFN5 | -- | 30148503 | Hydrolase activity, acting on carbon-nitrogen (but not peptide) bonds | Not Available |
| 317 | Poly-beta-1, 6-N-acetyl-D-glucosamine synthase (Poly-beta-1, 6-GlcNAc synthase) (EC 2.4.1.-) | *Yersinia pseudotuberculosis serotype I* | Strain IP32953 | Q66B29 | -- | 15358858 | Acetyl glucosaminyl transferase activity | Not Available |
| 318 | Poly-beta-1, 6-N-acetyl-D-glucosamine synthase (Poly-beta-1, 6-GlcNAc synthase) (EC 2.4.1.-) (Fragment) | *Planctomycetaceae bacterium* | | A0A3B8ZEJ0 | -- | 30148503 | Acetyl glucosaminyl transferase activity | Not Available |
| 319 | Poly-beta-1, 6-N-acetyl-D-glucosamine synthase (Poly-beta-1, 6-GlcNAc synthase) (EC 2.4.1.-) | *Acinetobacter baumannii* | | A0A2I8CSE1 | -- | 29437920 | Acetyl glucosaminyl transferase activity | Not Available |
| 320 | Uncharacterized protein | *Xanthomonas oryzae pv. oryzae* | Strain KACC10331 / KXO85 | Q5GVV4 | -- | 15673718 | Not Available | Not Available |
| 321 | Poly-beta-1, 6-N-acetyl-D-glucosamine biosynthesis protein PgaD | *Acinetobacter baumannii* | | A0A7Z1WPV2 | -- | 27917170 | Not Available | Not Available |
| 322 | Poly-beta-1, 6-N-acetyl-D-glucosamine synthase (Poly-beta-1, 6-GlcNAc synthase) (EC 2.4.1.-) | *Acinetobacter baumannii* | | A0A1E3M1V1 | -- | 25136007 | Acetyl glucosaminyl transferase activity | Not Available |
| 323 | Uncharacterized protein | *Novosphingobium nitrogenifigens DSM 19370* | | F1ZCX2 | -- | 22156397 | Not Available | Not Available |
| 324 | HmsF protein | *Xanthomonas oryzae pv. oryzae* | Strain KACC10331 / KXO85 | Q5GVV6 | -- | 15673718 | Hydrolase activity, acting on carbon-nitrogen (but not peptide) bonds | Not Available |
| 325 | Poly-beta-1, 6-N-acetyl-D-glucosamine synthase (Poly-beta-1, 6-GlcNAc synthase) (EC 2.4.1.-) | *Xanthomonas oryzae pv. oryzae* | Strain KACC10331 / KXO85 | Q5GVV5 | -- | 15673718 | Acetyl glucosaminyl transferase activity | Not Available |
| 326 | Poly-beta-1, 6-N-acetyl-D-glucosamine N-deacetylase PgaB | *Acinetobacter baumannii* | | A0A2I8CS57 | -- | 29437920 | Hydrolase activity, acting on carbon-nitrogen (but not peptide) bonds | Not Available |
| 327 | Poly-beta-1, 6-N-acetyl-D-glucosamine synthase (Poly-beta-1, 6-GlcNAc synthase) (EC 2.4.1.-) | *Acinetobacter baumannii* | | A0A0D5YG83 | -- | 25845845 | Acetyl glucosaminyl transferase activity | Not Available |
| 328 | Putative inner Membrane protein associated with biofilm formation | *Escherichia coli O45:K1* | Strain S88 / ExPEC | B7MIG7 | -- | 19165319 | Not Available | Not Available |
| 329 | Putative polysaccharide deacetylase associated with biofilm formation putative lipoprotein | *Escherichia coli O45:K1* | Strain S88 / ExPEC | B7MIG9 | -- | 19165319 | Hydrolase activity, acting on carbon-nitrogen (but not peptide) bonds | Not Available |
| 330 | Uncharacterized protein | *Deltaproteobacteria bacterium RIFOXYD12_FULL_57_12* | | A0A1F9LU99 | -- | 27774985 | Not Available | Not Available |
| 331 | Uncharacterized protein | *Serratia spp. M24T3* | | I0QPB7 | -- | 22740681 | Not Available | Not Available |
| 332 | Biofilm PGA synthesis lipoprotein PgaB | *Actinobacillus pleuropneumoniae serotype 5b* | Strain L20 | A3N3L6 | -- | 18065534 | Hydrolase activity, acting on carbon-nitrogen (but not peptide) bonds | Not Available |
| 333 | Uncharacterized protein | *Halomonas spp. TD01* | | F7SRR8 | -- | 22040376 | Not Available | Not Available |
| 334 | GGDEF family protein | *Deinococcus radiodurans* | Strain ATCC 13939 / DSM 20539 / JCM 16871 / LMG 4051 / NBRC 15346 / NCIMB 9279 / R1 / VKM B-1422 | Q9RZN0 | -- | 10567266 | Diguanylate cyclase activity | Not Available |
| 335 | GGDEF family protein | *Deinococcus radiodurans* | Strain ATCC 13939 / DSM 20539 / JCM 16871 / LMG 4051 / NBRC 15346 / NCIMB 9279 / R1 / VKM B-1422 | Q9RVP4 | -- | 10567266 | Diguanylate cyclase activity | Not Available |
| 336 | GGDEF family protein | *Deinococcus radiodurans* | Strain ATCC 13939 / DSM 20539 / JCM 16871 / LMG 4051 / NBRC 15346 / NCIMB 9279 / R1 / VKM B-1422 | Q9RTW9 | -- | 10567266 | Diguanylate cyclase activity | Not Available |

| Target ID | Bacterial Protein Name | Organism Name | Strain | UniProt ID | PDB ID | PubMed ID | Classification as per UniProt GO Molecular Function | Classification of Function as per PDB |
|---|---|---|---|---|---|---|---|---|
| 337 | GGDEF family protein | *Deinococcus radiodurans* | Strain ATCC 13939 / DSM 20539 / JCM 16871 / LMG 4051 / NBRC 15346 / NCIMB 9279 / R1 / VKM B-1422 | Q9RRJ2 | -- | 10567266 | Diguanylate cyclase activity | Not Available |
| 338 | GGDEF family protein | *Deinococcus radiodurans* | Strain ATCC 13939 / DSM 20539 / JCM 16871 / LMG 4051 / NBRC 15346 / NCIMB 9279 / R1 / VKM B-1422 | Q9RZS9 | -- | 10567266 | Diguanylate cyclase activity | Not Available |
| 339 | Poly-beta-1, 6-N-acetyl-D-glucosamine synthase (Poly-beta-1, 6-GlcNAc synthase) (EC 2.4.1.-) | *Rubrivivax benzoatilyticus JA2 = ATCC BAA-35* | | F3LKS7 | -- | 21478355 | Acetyl glucosaminyl transferase activity | Not Available |
| 340 | Poly-beta-1, 6-N-acetyl-D-glucosamine synthase (Poly-beta-1, 6-GlcNAc synthase) (EC 2.4.1.-) | *Paraburkholderia azotifigens* | | A0A5C6VQD9 | -- | 29185955 | Acetyl glucosaminyl transferase activity | Not Available |
| 341 | Poly-beta-1, 6-N-acetyl-D-glucosamine synthase (Poly-beta-1, 6-GlcNAc synthase) (EC 2.4.1.-) | *Acidiferrobacteraceae bacterium* | | A0A7V2NLQ8 | -- | 31911466 | Acetyl glucosaminyl transferase activity | Not Available |
| 342 | Poly-beta-1, 6-N-acetyl-D-glucosamine N-deacetylase PgaB (EC 3.5.1.-) | *Pseudomonas spp. WS 5010* | | A0A7Y0ZL61 | -- | 32587583 | Hydrolase activity, acting on carbon-nitrogen (but not peptide) bonds | Not Available |
| 343 | Uncharacterized protein | *Lacticaseibacillus fabifermentans T30PCM01* | | W6T8S0 | -- | 24558238 | Not Available | Not Available |
| 344 | Uncharacterized protein | *Xanthomonas oryzae pv. oryzicola* | Strain BLS256 | G7TD25 | -- | 21784931 | Not Available | Not Available |
| 345 | NodB homology domain-containing protein | *Chromobacterium violaceum* | Strain ATCC 12472 / DSM 30191 / JCM 1249 / NBRC 12614 / NCIMB 9131 / NCTC 9757 | Q7NQ63 | -- | 14500782 | Hydrolase activity, acting on carbon-nitrogen (but not peptide) bonds | Not Available |
| 346 | Type II toxin-antitoxin system MqsR family toxin | *Gemmataceae bacterium* | | A0A7V3VXR8 | -- | 31911466 | Not Available | Not Available |
| 347 | Uncharacterized protein | *Lacticaseibacillus fabifermentans T30PCM01* | | W6T5A4 | -- | 24558238 | Not Available | Not Available |
| 348 | Poly-beta-1, 6-N-acetyl-D-glucosamine synthase (Poly-beta-1, 6-GlcNAc synthase) (EC 2.4.1.-) | *Chromobacterium violaceum* | Strain ATCC 12472 / DSM 30191 / JCM 1249 / NBRC 12614 / NCIMB 9131 / NCTC 9757 | Q7NTW2 | -- | 14500782 | Acetyl glucosaminyl transferase activity | Not Available |
| 349 | Hemin storage signal peptide protein | *Chromobacterium violaceum* | Strain ATCC 12472 / DSM 30191 / JCM 1249 / NBRC 12614 / NCIMB 9131 / NCTC 9757 | Q7NTW1 | -- | 14500782 | Hydrolase activity, acting on carbon-nitrogen (but not peptide) bonds | Not Available |
| 350 | Uncharacterized protein | *Methylomicrobium album BG8* | | H8GLP1 | -- | 23580712 | Not Available | Not Available |
| 351 | Uncharacterized protein | *Chromobacterium violaceum* | Strain ATCC 12472 / DSM 30191 / JCM 1249 / NBRC 12614 / NCIMB 9131 / NCTC 9757 | Q7NTW3 | -- | 14500782 | Not Available | Not Available |
| 352 | Poly-beta-1, 6-N-acetyl-D-glucosamine synthase (Poly-beta-1, 6-GlcNAc synthase) (EC 2.4.1.-) | *Pseudomonas orientalis* | | A0A2L0RQV7 | -- | 29479340 | Acetyl glucosaminyl transferase activity | Not Available |
| 353 | Poly-beta-1, 6-N-acetyl-D-glucosamine synthase (Poly-beta-1, 6-GlcNAc synthase) (EC 2.4.1.-) | *Xanthomonas oryzae pv. oryzicola* | Strain BLS256 | G7TD26 | -- | 21784931 | Acetyl glucosaminyl transferase activity | Not Available |
| 354 | Poly-beta-1, 6-N-acetyl-D-glucosamine N-deacetylase PgaB | *Acinetobacter spp. ACNIH4* | | A0A2S4TAE7 | -- | 29437920 | Hydrolase activity, acting on carbon-nitrogen (but not peptide) bonds | Not Available |
| 355 | Poly-beta-1, 6-N-acetyl-D-glucosamine N-deacetylase PgaB | *Acinetobacter spp. ACNIH4* | | A0A2S4TAJ4 | -- | 29437920 | Hydrolase activity, acting on carbon-nitrogen (but not peptide) bonds | Not Available |
| 356 | GGDEF domain-containing protein | *Thermotoga maritima* | Strain ATCC 43589 / DSM 3109 / JCM 10099 / NBRC 100826 / MSB8 | Q9X067 | -- | 10360571 | Diguanylate cyclase activity | Not Available |
| 357 | Response regulator | *Thermotoga maritima* | Strain ATCC 43589 / DSM 3109 / JCM 10099 / NBRC 100826 / MSB8 | Q9WXZ0 | -- | 10360571 | Diguanylate cyclase activity | Not Available |
| 358 | GGDEF domain-containing protein | *Thermotoga maritima* | Strain ATCC 43589 / DSM 3109 / JCM 10099 / NBRC 100826 / MSB8 | Q9X1T0 | -- | 10360571 | Diguanylate cyclase activity | Not Available |
| 359 | Poly-beta-1, 6-N-acetyl-D-glucosamine synthase (Poly-beta-1, 6-GlcNAc synthase) (EC 2.4.1.-) | *Staphylococcus simiae CCM 7213 = CCUG 51256* | | G5JK07 | -- | 22272658 | Acetyl glucosaminyl transferase activity | Not Available |
| 360 | GGDEF domain-containing protein | *Thermotoga maritima* | Strain ATCC 43589 / DSM 3109 / JCM 10099 / NBRC 100826 / MSB8 | Q9X1S4 | -- | 10360571 | Diguanylate cyclase activity | Not Available |
| 361 | Poly-beta-1, 6-N-acetyl-D-glucosamine synthase (Poly-beta-1, 6-GlcNAc synthase) (EC 2.4.1.-) | *Escherichia coli O45:K1* | Strain S88 / ExPEC | B7MIG8 | -- | 19165319 | Acetyl glucosaminyl transferase activity | Not Available |
| 362 | Poly-beta-1, 6-N-acetyl-D-glucosamine synthase (Poly-beta-1, 6-GlcNAc synthase) (EC 2.4.1.-) | *Pseudomonas chlororaphis* | | A0A0D5Y7R2 | -- | 25679537 | Acetyl glucosaminyl transferase activity | Not Available |
| 363 | Hemin storage protein | *Stenotrophomonas geniculata N1* | | A0A0L8AEA0 | -- | 22689240 | Hydrolase activity, acting on carbon-nitrogen (but not peptide) bonds | Not Available |
| 364 | Uncharacterized protein | *Actinobacteria bacterium* | | A0A7V4MZE2 | -- | 31911466 | Not Available | Not Available |
| 365 | Poly-beta-1, 6-N-acetyl-D-glucosamine synthase (Poly-beta-1, 6-GlcNAc synthase) (EC 2.4.1.-) | *Pseudomonas spp. WS 5413* | | A0A7Y0Z1V7 | -- | 32587583 | Acetyl glucosaminyl transferase activity | Not Available |
| 366 | Uncharacterized protein | *Hyphomonaceae bacterium* | | A0A3D2W3N1 | -- | 30148503 | Not Available | Not Available |
| 367 | Putative PGA biosynthesis protein | *Escherichia coli O83:H1* | Strain NRG 857C / AIEC | A0A0H3EHC0 | -- | 21108814 | Not Available | Not Available |
| 368 | Poly-beta-1, 6-N-acetyl-D-glucosamine biosynthesis protein PgaD | *Pseudomonas spp. WS 5413* | | A0A7Y0Z1X4 | -- | 32587583 | Not Available | Not Available |
| 369 | Poly-beta-1, 6-N-acetyl-D-glucosamine N-deacetylase PgaB (EC 3.5.1.-) | *Pseudomonas spp. WS 5413* | | A0A7Y0Z1Z3 | -- | 32587583 | Hydrolase activity, acting on carbon-nitrogen (but not peptide) bonds | Not Available |
| 370 | Poly-beta-1, 6-N-acetyl-D-glucosamine N-deacetylase PgaB | *Methylophaga spp.* | | A0A7C5BNS2 | -- | 31911466 | Hydrolase activity, acting on carbon-nitrogen (but not peptide) bonds | Not Available |
| 371 | Uncharacterized protein | *Stenotrophomonas spp.* | | A0A3D0GBU3 | -- | 30148503 | Not Available | Not Available |
| 372 | mRNA interferase MqsR | *Hyphomonaceae bacterium* | | A0A352CC32 | -- | 30148503 | Not Available | Not Available |
| 373 | Poly-beta-1, 6-N-acetyl-D-glucosamine synthase (Poly-beta-1, 6-GlcNAc synthase) (EC 2.4.1.-) | *Stenotrophomonas spp.* | | A0A3C0X5Z0 | -- | 30148503 | Acetyl glucosaminyl transferase activity | Not Available |
| 374 | Poly-beta-1, 6-N-acetyl-D-glucosamine biosynthesis protein PgaD | *Stenotrophomonas spp.* | | A0A357N0H1 | -- | 30148503 | Not Available | Not Available |
| 375 | Poly-beta-1, 6-N-acetyl-D-glucosamine synthase (Poly-beta-1, 6-GlcNAc synthase) (EC 2.4.1.-) | *Stenotrophomonas spp.* | | A0A349Y4K4 | -- | 30148503 | Acetyl glucosaminyl transferase activity | Not Available |

| Target ID | Bacterial Protein Name | Organism Name | Strain | UniProt ID | PDB ID | PubMed ID | Classification as per UniProt GO Molecular Function | Classification of Function as per PDB |
|---|---|---|---|---|---|---|---|---|
| 376 | Poly-beta-1, 6-N-acetyl-D-glucosamine synthase (Poly-beta-1, 6-GlcNAc synthase) (EC 2.4.1.-) | *Escherichia coli O83:H1* | Strain NRG 857C / AIEC | A0A0H3EI51 | -- | 21108814 | Acetyl glucosaminyl transferase activity | Not Available |
| 377 | Surface composition regulator | *Escherichia coli O83:H1* | Strain NRG 857C / AIEC | A0A0H3EM10 | -- | 21108814 | Not Available | Not Available |
| 378 | Poly-beta-1, 6-N-acetyl-D-glucosamine N-deacetylase PgaB | *Deltaproteobacteria bacterium RBG_13_60_28* | | A0A1F9BIL4 | -- | 27774985 | Hydrolase activity, acting on carbon-nitrogen (but not peptide) bonds | Not Available |
| 379 | Poly-beta-1, 6-N-acetyl-D-glucosamine synthase (Poly-beta-1, 6-GlcNAc synthase) (EC 2.4.1.-) | *Deltaproteobacteria bacterium RBG_13_60_28* | | A0A1F9BKU9 | -- | 27774985 | Acetyl glucosaminyl transferase activity | Not Available |
| 380 | Poly-beta-1, 6-N-acetyl-D-glucosamine biosynthesis protein PgaD | *Stenotrophomonas spp.* | | A0A3D0GC17 | -- | 30148503 | Not Available | Not Available |
| 381 | Poly-beta-1, 6-N-acetyl-D-glucosamine N-deacetylase PgaB | *Stenotrophomonas spp.* | | A0A3C0X5X6 | -- | 30148503 | Hydrolase activity, acting on carbon-nitrogen (but not peptide) bonds | Not Available |
| 382 | Poly-beta-1, 6-N-acetyl-D-glucosamine synthase (Poly-beta-1, 6-GlcNAc synthase) (EC 2.4.1.-) | *Stenotrophomonas spp.* | | A0A356JD40 | -- | 30148503 | Acetyl glucosaminyl transferase activity | Not Available |
| 383 | Poly-beta-1, 6-N-acetyl-D-glucosamine biosynthesis protein PgaD (Fragment) | *Stenotrophomonas spp.* | | A0A3C0X5Y8 | -- | 30148503 | Not Available | Not Available |
| 384 | Poly-beta-1, 6-N-acetyl-D-glucosamine synthase (Poly-beta-1, 6-GlcNAc synthase) (EC 2.4.1.-) | *Stenotrophomonas spp.* | | A0A3D4THW1 | -- | 30148503 | Acetyl glucosaminyl transferase activity | Not Available |
| 385 | Poly-beta-1, 6-N-acetyl-D-glucosamine N-deacetylase PgaB | *Stenotrophomonas spp.* | | A0A349Y4K5 | -- | 30148503 | Hydrolase activity, acting on carbon-nitrogen (but not peptide) bonds | Not Available |
| 386 | Poly-beta-1, 6-N-acetyl-D-glucosamine synthase (Poly-beta-1, 6-GlcNAc synthase) (EC 2.4.1.-) | *Methylococcaceae bacterium* | | A0A351V597 | -- | 30148503 | Acetyl glucosaminyl transferase activity | Not Available |
| 387 | Poly-beta-1, 6-N-acetyl-D-glucosamine synthase (Poly-beta-1, 6-GlcNAc synthase) (EC 2.4.1.-) | *Paraburkholderia hospita BS001* | | I5CU28 | -- | 22843604 | Acetyl glucosaminyl transferase activity | Not Available |
| 388 | Type IV pilin PilA | *Vibrio vulnificus* | | Q8VVV3 | -- | 15731039 | Not Available | Not Available |
| 389 | Uncharacterized protein | *Lactococcus lactis subspp. cremoris IBB477* | | A0A1E7G187 | -- | 27687992 | Not Available | Not Available |
| 390 | Putative transmembrane protein | *blood disease bacterium R229* | | G2ZXS3 | -- | 21931687 | Not Available | Not Available |
| 391 | Surface composition regulator | *Salmonella enterica subspp. enterica serovar* | Hvittingfoss | A0A5I0UXC3 | -- | 30286803 | Not Available | Not Available |
| 392 | IcaA (Fragment) | *Staphylococcus simulans* | | Q83ZH8 | -- | 12957956 | Not Available | Not Available |
| 393 | Surface composition regulator | *Salmonella enterica subspp. enterica serovar* | Java | A0A741RTZ8 | -- | 30286803 | Not Available | Not Available |
| 394 | PsoA | *Pseudomonas putida* | Arthrobacter Siderocapsulatus | A8MN36 | -- | 16585751; 18599835 | Catalytic activity, phosphopantetheine binding | Not Available |
| 395 | Putative transmembrane protein | *Ralstonia syzygii R24* | | G3A8J7 | -- | 21931687 | Not Available | Not Available |
| 396 | Poly-beta-1, 6-N-acetyl-D-glucosamine synthase (Poly-beta-1, 6-GlcNAc synthase) (EC 2.4.1.-) | *Staphylococcus aureus* | | A0A1D0C161 | -- | 20876371 | Acetyl glucosaminyl transferase activity | Not Available |
| 397 | Hemin storage signal peptide protein | *Paraburkholderia hospita BS001* | | I5CU27 | -- | 22843604 | Hydrolase activity, acting on carbon-nitrogen (but not peptide) bonds | Not Available |
| 398 | Surface composition regulator | *Salmonella enterica subspp. indica serovar* | 45:a:e, n, x | A0A737BKL7 | -- | 30286803 | Not Available | Not Available |
| 399 | Poly-beta-1, 6-N-acetyl-D-glucosamine synthase (Poly-beta-1, 6-GlcNAc synthase) (EC 2.4.1.-) | *blood disease bacterium R229* | | G2ZXS2 | -- | 21931687 | Acetyl glucosaminyl transferase activity | Not Available |
| 400 | Putative polysaccharide deacetylase associated with biofilm formation putative lipoprotein (PgaB) | *blood disease bacterium R229* | | G2ZXS1 | -- | 21931687 | Hydrolase activity, acting on carbon-nitrogen (but not peptide) bonds | Not Available |
| 401 | Surface composition regulator | *Salmonella enterica subspp. houtenae serovar* | 16:z4, z32:- | A0A735L3N9 | -- | 30286803 | Not Available | Not Available |
| 402 | IcaA (Fragment) | *Staphylococcus aureus* | | Q83ZH9 | -- | 12957956 | Not Available | Not Available |
| 403 | Poly-beta-1, 6-N-acetyl-D-glucosamine synthase (Poly-beta-1, 6-GlcNAc synthase) (EC 2.4.1.-) | *Staphylococcus aureus* | | A0A6A9GS78 | -- | 31682959 | Acetyl glucosaminyl transferase activity | Not Available |
| 404 | Glycerol esther hydrolase (Fragment) | *Staphylococcus aureus* | | Q9RQP5 | -- | 10496925 | Hydrolase activity | Not Available |
| 405 | Surface composition regulator | *Salmonella enterica subspp. indica serovar* | 45:a:e, n, x | A0A737BK14 | -- | 30286803 | Not Available | Not Available |
| 406 | Conjugative transfer protein Pilx7 (PilX7) (Pilx7 protein) (Putative PilX7) | *Escherichia coli* | | B0ZDY1 | -- | 15328093; 18440636; 19717626; 21831988; 24862095 | Not Available | Not Available |
| 407 | Poly-beta-1, 6-N-acetyl-D-glucosamine N-deacetylase PgaB (Polysaccharide deacetylase) | *Burkholderia ambifaria* | Strain ATCC BAA-244 / AMMD Burkholderia cepacia Strain AMMD | Q0B9V5 | -- | 25931592 | Hydrolase activity, acting on carbon-nitrogen (but not peptide) bonds | Not Available |
| 408 | Poly-beta-1, 6-N-acetyl-D-glucosamine synthase (Poly-beta-1, 6-GlcNAc synthase) (EC 2.4.1.-) | *Stenotrophomonas maltophilia* | Pseudomonas maltophilia Xanthomonas maltophilia | A0A0K2IYC8 | -- | 26324280; 29170656 | Acetyl glucosaminyl transferase activity | Not Available |
| 409 | Poly-beta-1, 6-N-acetyl-D-glucosamine synthase (Poly-beta-1, 6-GlcNAc synthase) (EC 2.4.1.-) | *Enterobacter kobei* | | A0A2J0PJF5 | -- | 29088362 | Acetyl glucosaminyl transferase activity | Not Available |
| 410 | Type II toxin-antitoxin system MqsR family toxin | *Deltaproteobacteria bacterium* | | A0A7C8AFX8 | -- | 32737307 | Not Available | Not Available |
| 411 | Poly-beta-1, 6-N-acetyl-D-glucosamine biosynthesis protein PgaD | *Escherichia coli* | | A0A0A1A417 | -- | 30286803; 30947268 | Not Available | Not Available |
| 412 | Poly-beta-1, 6-N-acetyl-D-glucosamine N-deacetylase PgaB (EC 3.5.1.-) | *Escherichia coli* | | A0A795PNG2 | -- | 30286803 | Hydrolase activity, acting on carbon-nitrogen (but not peptide) bonds | Not Available |
| 413 | GCU-specific mRNA interferase toxin of the MqsR-MqsA toxin-antitoxin system (Type II toxin-antitoxin system toxin MqsR) (EC 3.1.26.-) (mRNA interferase MqsR) (EC 3.1.-.-) | *Escherichia coli* | | A0A0H0HXD8 | -- | 30286803; 31477907 | Hydrolase activity | Not Available |
| 414 | Lipoprotein YcdR (EC 3.5.1.-) (Poly-beta-1, 6-N-acetyl-D-glucosamine N-deacetylase PgaB) | *Escherichia coli* | | A0A2T3TFG9 | -- | 31477907 | Hydrolase activity, acting on carbon-nitrogen (but not peptide) bonds | Not Available |
| 415 | PGA biosynthesis protein (Poly-beta-1, 6-N-acetyl-D-glucosamine biosynthesis protein PgaD) | *Escherichia coli* | | A0A0D8VWU3 | -- | 30286803; 31477907; 32021174 | Not Available | Not Available |
| 416 | Poly-beta-1, 6-N-acetyl-D-glucosamine synthase (Poly-beta-1, 6-GlcNAc synthase) (EC 2.4.1.-) | *Escherichia coli* | | A0A1U9U1S6 | -- | 30286803; 31477907 | Acetyl glucosaminyl transferase activity | Not Available |
| 417 | Poly-beta-1, 6-N-acetyl-D-glucosamine synthase (Poly-beta-1, 6-GlcNAc synthase) (EC 2.4.1.-) | *Escherichia coli* | | A0A3P1J9P9 | -- | 30286803 | Acetyl glucosaminyl transferase activity | Not Available |
| 418 | Poly-beta-1, 6-N-acetyl-D-glucosamine biosynthesis protein PgaD | *Enterobacter kobei* | | A0A2J0PIC7 | -- | 29088362 | Not Available | Not Available |
| 419 | Surface composition regulator | *Salmonella enterica subspp. enterica serovar* | Give | A0A5C5HDB9 | -- | 26139723; 30286803; 30902850 | Not Available | Not Available |
| 420 | Poly-beta-1, 6-N-acetyl-D-glucosamine synthase (Poly-beta-1, 6-GlcNAc synthase) (EC 2.4.1.-) | *Escherichia coli* | | C3TEV3 | -- | 19439656 | Acetyl glucosaminyl transferase activity | Not Available |
| 421 | Surface composition regulator | *Salmonella typhimurium* | Strain SL1344 | A0A0H3NI14 | -- | 22538806; 30286803 | Not Available | Not Available |
| 422 | Poly-beta-1, 6-N-acetyl-D-glucosamine N-deacetylase PgaB | *Escherichia coli* | | A0A3K0QE29 | -- | 30286803 | Hydrolase activity, acting on carbon-nitrogen (but not peptide) bonds | Not Available |

| Target ID | Bacterial Protein Name | Organism Name | Strain | UniProt ID | PDB ID | PubMed ID | Classification as per UniProt GO Molecular Function | Classification of Function as per PDB |
|---|---|---|---|---|---|---|---|---|
| 423 | Uncharacterized protein | *Deltaproteobacteria bacterium* | | A0A661K4W2 | -- | 31911466 | Not Available | Not Available |
| 424 | LuxR regulator (SmcR-like protein VvpR) | *Vibrio scophthalmi LMG 19158* | | F9RJF4 | -- | 23198796; 21930677 | DNA binding | Not Available |
| 425 | Poly-beta-1, 6-N-acetyl-D-glucosamine biosynthesis protein PgaD | *Escherichia coli* | | A0A2A6Q924 | -- | 28221970; 30924410 | Not Available | Not Available |
| 426 | Poly-beta-1, 6-N-acetyl-D-glucosamine biosynthesis protein PgaD (Transporter) | *Acinetobacter pittii* | Acinetobacter genomospp. 3 | A0A242STX0 | -- | 30036696 | Not Available | Not Available |
| 427 | Poly-beta-1, 6-N-acetyl-D-glucosamine N-deacetylase PgaB | *Escherichia coli* | | A0A2A6Q8V5 | -- | 28221970; 30924410 | Hydrolase activity, acting on carbon-nitrogen (but not peptide) bonds | Not Available |
| 428 | Poly-beta-1, 6-N-acetyl-D-glucosamine N-deacetylase PgaB (EC 3.5.1.-) | *Escherichia coli* | | A0A233ZEF3 | -- | 30286803; 32021174 | Hydrolase activity, acting on carbon-nitrogen (but not peptide) bonds | Not Available |
| 429 | Surface composition regulator | *Escherichia coli* | | A0A1M0PIU6 | -- | 30286803; 31477907 | Not Available | Not Available |
| 430 | Poly-beta-1, 6-N-acetyl-D-glucosamine synthase (Poly-beta-1, 6-GlcNAc synthase) (EC 2.4.1.-) | *Escherichia coli* | | A0A2A6Q915 | -- | 28221970; 30924410 | Acetyl glucosaminyl transferase activity | Not Available |
| 431 | Molecular chaperone (MrkB) (Putative bacterial pili assembly chaperone) (Putative fimbrial chaperone protein) (Type 3 fimbria chaperone MrkB) | *Escherichia coli* | | B0LHG6 | -- | 18055599; 19717626; 22479389; 26441869; 30286803 | Not Available | Not Available |
| 432 | Plasmid stability protein StbD (Putative antitoxin) (Plasmid stability protein, antitoxin StbD) (Plasmid stabilization protein) (Putative antitoxin of toxin-antitoxin system) (RelB / StbD replicon stabilization protein (Antitoxin to RelE / StbE)) (Stability protein) (Stability protein StbD) (StbD) (StbD stability protein) (Toxin-antitoxin system, antitoxin component) | *Escherichia coli* | | B0EYQ2 | -- | 15328093; 17936903; 18440636; 19717626; 21831988; 22470007; 22479389; 25403661; 25421479; 26048440; 30286803; 30924410; 31642700; 31652858 | Not Available | Not Available |
| 433 | Poly-beta-1, 6-N-acetyl-D-glucosamine synthase (Poly-beta-1, 6-GlcNAc synthase) (EC 2.4.1.-) | *Escherichia coli* | | A0A1Y5BGC5 | -- | 30286803 | Acetyl glucosaminyl transferase activity | Not Available |
| 434 | Biofilm PGA synthesis protein PgaD (PGA biosynthesis protein) (Poly-beta-1, 6-N-acetyl-D-glucosamine biosynthesis) (Poly-beta-1, 6-N-acetyl-D-glucosamine biosynthesis protein PgaD) | *Escherichia coli* | | C3TEV8 | -- | 19439656; 23275093; 26441869; 30286803; 29437920; 30924410; 31642700; 31652858; 31477907; 30948755; 32739475; 28953937 | Not Available | Not Available |
| 435 | Poly-beta-1, 6-N-acetyl-D-glucosamine synthase (Poly-beta-1, 6-GlcNAc synthase) (EC 2.4.1.-) | *Escherichia coli* | | J7QDP1 | -- | 23275093; 30286803; 29437920; 30924410; 31642700; 31652858; 31477907; 30948755; 28953937 | Acetyl glucosaminyl transferase activity | Not Available |
| 436 | Surface composition regulator | *Escherichia coli* | | J7QHM3 | -- | 23275093; 26441869; 26227606; 28686503; 30286803; 29437920; 30924410; 31642700; 31652858; 31477907; 30947268; 30948755; 32739475; 32599678; 28953937 | Not Available | Not Available |
| 437 | Poly-beta-1, 6-N-acetyl-D-glucosamine N-deacetylase PgaB (EC 3.5.1.-) | *Escherichia coli* | | A0A192EMZ6 | -- | 30286803 | Hydrolase activity, acting on carbon-nitrogen (but not peptide) bonds | Not Available |
| 438 | Poly-beta-1, 6-N-acetyl-D-glucosamine synthase (Poly-beta-1, 6-GlcNAc synthase) (EC 2.4.1.-) | *Escherichia coli* | | W8TQ48 | -- | 26441869 | Acetyl glucosaminyl transferase activity | Not Available |
| 439 | Surface composition regulator | *Salmonella newport* | | A0A0R9NRZ2 | -- | 30286803 | Not Available | Not Available |
| 440 | Poly-beta-1, 6-N-acetyl-D-glucosamine synthase (Poly-beta-1, 6-GlcNAc synthase) (EC 2.4.1.-) | *Escherichia coli* | | A0A1D7PYQ0 | -- | 30286803; 30947268 | Acetyl glucosaminyl transferase activity | Not Available |
| 441 | Surface composition regulator | *Escherichia coli* | | E2QE75 | -- | 28221970; 28686503; 30286803; 30924410; 31477907 | Not Available | Not Available |
| 442 | Surface composition regulator | *Salmonella enterica subspp. enterica serovar* | Weltevreden | A0A3V7I4R8 | -- | 31099590 | Not Available | Not Available |
| 443 | Poly-beta-1, 6-N-acetyl-D-glucosamine N-deacetylase PgaB (EC 3.5.1.-) | *Escherichia coli* | | A0A4Y8EDQ8 | -- | 32739475 | Hydrolase activity, acting on carbon-nitrogen (but not peptide) bonds | Not Available |
| 444 | Poly-beta-1, 6-N-acetyl-D-glucosamine synthase (Poly-beta-1, 6-GlcNAc synthase) (EC 2.4.1.-) | *Acinetobacter pittii* | Acinetobacter genomospp. 3 | A0A242SD62 | -- | 30036696 | Acetyl glucosaminyl transferase activity | Not Available |
| 445 | Poly-beta-1, 6-N-acetyl-D-glucosamine synthase (Poly-beta-1, 6-GlcNAc synthase) (EC 2.4.1.-) | *Acinetobacter pittii* | Acinetobacter genomospp. 3 | A0A0N9ALG4 | -- | 30036696 | Acetyl glucosaminyl transferase activity | Not Available |
| 446 | ActX (ActX protein) (Transcription termiNot Availabletion factor NusG) (Transcriptionalactivator) | *Escherichia coli* | | B0ZDY9 | -- | 15328093; 18440636; 19717626; 21831988; 30286803; 31652858 | Not Available | Not Available |
| 447 | Surface composition regulator | *Salmonella enterica subspp. enterica serovar* | Sandiego | A0A5J1IID0 | -- | 30286803 | Not Available | Not Available |

| Target ID | Bacterial Protein Name | Organism Name | Strain | UniProt ID | PDB ID | PubMed ID | Classification as per UniProt GO Molecular Function | Classification of Function as per PDB |
|---|---|---|---|---|---|---|---|---|
| 448 | Chromosome partitioning protein ParA (ParA family protein) (ParF) (Partitioning protein ParA) (Plasmid partition protein) (Plasmid partition protein ParF) (Plasmid partitioning protein ParF) (Putative crown gall tumor protein VirC1) | *Escherichia coli* | | B0EYM3 | -- | 17936903; 19717626; 22470007; 22479389; 31652858 | Not Available | Not Available |
| 449 | Surface composition regulator | *Pluralibacter gergoviae* | Enterobacter gergoviae | A0A1S2BI09 | -- | 25502672 | Not Available | Not Available |
| 450 | Poly-beta-1, 6-N-acetyl-D-glucosamine biosynthesis protein PgaD | *Escherichia coli* | | A0A4Y8EDW4 | -- | 32739475 | Not Available | Not Available |
| 451 | Poly-beta-1, 6-N-acetyl-D-glucosamine N-deacetylase (EC 3.5.1.-) (Poly-beta-1, 6-N-acetyl-D-glucosamine N-deacetylase PgaB) | *Escherichia coli* | | A0A2X6F457 | -- | 30286803 | Hydrolase activity, acting on carbon-nitrogen (but not peptide) bonds | Not Available |
| 452 | Poly-beta-1, 6-N-acetyl-D-glucosamine synthase (Poly-beta-1, 6-GlcNAc synthase) (EC 2.4.1.-) | *Escherichia coli* | | A0A1Y2ZN44 | -- | 30924410 | Acetyl glucosaminyl transferase activity | Not Available |
| 453 | Lipoprotein YcdR (Poly-beta-1, 6-N-acetyl-D-glucosamine N-deacetylase) (Poly-beta-1, 6-N-acetyl-D-glucosamine N-deacetylase PgaB) (EC 3.5.1.-) | *Escherichia coli* | | A0A0L6Y2B4 | -- | 30286803 | Hydrolase activity, acting on carbon-nitrogen (but not peptide) bonds | Not Available |
| 454 | PGA biosynthesis protein (Poly-beta-1, 6-N-acetyl-D-glucosamine biosynthesis protein PgaD) | *Escherichia coli* | | A0A178T084 | -- | 30286803 | Not Available | Not Available |
| 455 | Poly-beta-1, 6-N-acetyl-D-glucosamine N-deacetylase PgaB (EC 3.5.1.-) | *Escherichia coli* | | A0A0J2BYQ8 | -- | 28686503; 30286803 | Hydrolase activity, acting on carbon-nitrogen (but not peptide) bonds | Not Available |
| 456 | Poly-beta-1, 6-N-acetyl-D-glucosamine N-deacetylase (Poly-beta-1, 6-N-acetyl-D-glucosamine N-deacetylase PgaB) (EC 3.5.1.-) | *Escherichia coli* | | A0A178T0C2 | -- | 30286803 | Hydrolase activity, acting on carbon-nitrogen (but not peptide) bonds | Not Available |
| 457 | Surface composition regulator | *Salmonella enterica subspp. enterica serovar* | Bredeney | A0A3V9PUT8 | -- | 30286803 | Not Available | Not Available |
| 458 | S-ribosylhomocysteine lyase (EC 4.4.1.21) (AI-2 synthesis protein) (Autoinducer-2 production protein LuxS) | *Streptococcus salivarius K12* | | J7TQ69 | -- | 23045482 | Iron ion binding, S-ribosylhomocysteine lyase activity | Not Available |
| 459 | Poly-beta-1, 6-N-acetyl-D-glucosamine N-deacetylase PgaB (EC 3.5.1.-) | *Escherichia coli* | | A0A0A1A2S7 | -- | 30286803; 30947268 | Hydrolase activity, acting on carbon-nitrogen (but not peptide) bonds | Not Available |
| 460 | MrkA fimbrial protein (MrkA fimbrial protein major fimbrial subunit) (Type 1 fimbrial protein) (Type 3 fimbria major subunit MrkA) | *Escherichia coli* | | B0ZDW4 | -- | 15328093; 18440636; 30286803; 31652858 | Not Available | Not Available |
| 461 | Conjugal transfer protein (Conjugal transfer protein Pilx9) (PilX9) (Pilx9 protein) (Pilx9 protein, putative conjugal transfer protein, putative exported protein) (TrbG / VirB9 family P-type conjugative transfer protein) | *Escherichia coli* | | B0ZDX9 | -- | 15328093; 18440636; 19717626; 22479389; 31652858 | Not Available | Not Available |
| 462 | Poly-beta-1, 6-N-acetyl-D-glucosamine synthase (Poly-beta-1, 6-GlcNAc synthase) (EC 2.4.1.-) | *Staphylococcus aureus* | Strain Newman | A0A0H3KAW6 | -- | 17951380 | Acetyl glucosaminyl transferase activity | Not Available |
| 463 | Surface composition regulator | *Salmonella enterica subspp. enterica serovar* | Schwarzengrund | A0A3T2ZVE9 | -- | 30286803 | Not Available | Not Available |
| 464 | Poly-beta-1, 6-N-acetyl-D-glucosamine biosynthesis protein PgaD | *Acinetobacter cumulans* | | A0A498CYW7 | -- | 30808586 | Not Available | Not Available |
| 465 | Motility quorum-sensing regulator / GCU-specific mRNA interferase toxin (Type II toxin-antitoxin system MqsR family toxin) | *Pseudomonas chengduensis* | | A0A1G6UUI8 | -- | 32587583 | Not Available | Not Available |
| 466 | Bacteriophage like protein (Putative toxin-antitoxin system, toxin component) | *Lactobacillus johnsonii ATCC 33200* | | C2E3I3 | -- | 26415554 | Not Available | Not Available |
| 467 | Major facilitator superfamily permease | *Acinetobacter oleivorans* | Strain JCM 16667 / KCTC 23045 / DR1 | D3JTG4 | -- | 20629796; 20639327 | Transmembrane transporter activity | Not Available |
| 468 | Acyl-CoA dehydrogenase | *Acinetobacter oleivorans* | Strain JCM 16667 / KCTC 23045 / DR1 | D3JTG9 | -- | 20629796; 20639327 | Flavin adenine dinucleotide binding, oxidoreductase activity, acting on the CH-CH group of donors | Not Available |
| 469 | AqsR (Transcriptionalactivator protein) | *Acinetobacter oleivorans* | Strain JCM 16667 / KCTC 23045 / DR1 | D3JTG7 | -- | 20629796; 20639327 | DNA binding | Not Available |
| 470 | Poly-beta-1, 6-N-acetyl-D-glucosamine synthase (Poly-beta-1, 6-GlcNAc synthase) (EC 2.4.1.-) | *Acinetobacter schindleri* | | A0A2S1FF20 | -- | 29954893 | Acetyl glucosaminyl transferase activity | Not Available |
| 471 | Surface composition regulator | *Salmonella diarizonae* | | A0A2I5HLE4 | -- | 30286803 | Not Available | Not Available |
| 472 | Poly-beta-1, 6-N-acetyl-D-glucosamine N-deacetylase PgaB | *Acinetobacter schindleri* | | A0A2S1FF79 | -- | 29954893 | Hydrolase activity, acting on carbon-nitrogen (but not peptide) bonds | Not Available |
| 473 | Poly-beta-1, 6-N-acetyl-D-glucosamine N-deacetylase PgaB | *Acinetobacter schindleri* | | A0A2S1FF30 | -- | 29954893 | Hydrolase activity, acting on carbon-nitrogen (but not peptide) bonds | Not Available |
| 474 | Acyl-homoserine-lactone synthase (EC 2.3.1.184) (Autoinducer synthesis protein) | *Acinetobacter oleivorans* | Strain JCM 16667 / KCTC 23045 / DR1 | D3JTG5 | -- | 20629796; 20639327 | N-acyl homoserine lactone synthase activity | Not Available |
| 475 | Poly-beta-1, 6-N-acetyl-D-glucosamine biosynthesis protein PgaD | *Piscirickettsiaceae bacterium* | | A0A2A4PM66 | -- | 29099490 | Not Available | Not Available |
| 476 | Acyl-CoA synthetase (AMP-forming) / AMP-acid ligase II (Long chain fatty acid-CoA ligase) | *Acinetobacter oleivorans* | Strain JCM 16667 / KCTC 23045 / DR1 | D3JTG8 | -- | 20629796; 20639327 | Ligase activity | Not Available |
| 477 | Poly-beta-1, 6-N-acetyl-D-glucosamine N-deacetylase PgaB (Polysaccharide deacetylase) (Xylanae / chitin deacetylase) (EC 3.5.1.-) | *Acinetobacter baumannii* | | A0A0D5YHR1 | -- | 25845845 | Hydrolase activity, acting on carbon-nitrogen (but not peptide) bonds, Hydrolase activity, acting on glycosyl bonds | Not Available |
| 478 | Putative inner Membrane protein (Uncharacterized protein yigG) | *Salmonella typhimurium* | Strain LT2 / SGSC1412 / ATCC 700720 | Q7CPD6 | -- | 11677609 | Not Available | Not Available |
| 479 | Surface composition regulator | *Salmonella enterica subspp. enterica serovar* | Java | A0A426WQB0 | -- | 30286803; 30902850; 31099590 | Not Available | Not Available |
| 480 | Surface composition regulator | *Citrobacter pasteurii* | | A0A7X5FWW 8 | -- | 30087402 | Not Available | Not Available |
| 481 | Surface composition regulator | *Salmonella enterica subspp. enterica serovar* | Tennessee | A0A3G3DVF2 | -- | 30286803; 30902850 | Not Available | Not Available |
| 482 | Biofilm PGA synthesis protein pgaD (Biofilm synthesis auxiliary protein) (Poly-beta-1, 6-N-acetyl-D-glucosamine biosynthesis protein PgaD) (Transporter) | *Acinetobacter baumannii* | | A0A059ZP60 | -- | 25136007; 26170289; 29137647; 30124845 | Not Available | Not Available |
| 483 | Biofilm PGA synthesis lipoprotein pgaB (Lipoprotein HmsF) (EC 3.5.1.-) (Poly-beta-1, 6-N-acetyl-D-glucosamine N-deacetylase PgaB) (Polysaccharide deacetylase) | *Acinetobacter baumannii* | | A0A090BDC4 | -- | 25845845; 27917170; 29137647 | Hydrolase activity, acting on carbon-nitrogen (but not peptide) bonds | Not Available |
| 484 | Csu fimbrial tip adhesin CsuE (CsuE) | *Acinetobacter baumannii* | | Q6XBY2 | -- | 14663080 | Not Available | Not Available |
| 485 | Poly-beta-1, 6-N-acetyl-D-glucosamine biosynthesis protein PgaD | *Citrobacter pasteurii* | | A0A7X5JM17 | -- | 30087402 | Not Available | Not Available |
| 486 | Poly-beta-1, 6-N-acetyl-D-glucosamine biosynthesis protein PgaD | *Acinetobacter baumannii* | | A0A090B5L8 | -- | 27917170; 29137647; 30124845; 29437920 | Not Available | Not Available |

| Target ID | Bacterial Protein Name | Organism Name | Strain | UniProt ID | PDB ID | PubMed ID | Classification as per UniProt GO Molecular Function | Classification of Function as per PDB |
|---|---|---|---|---|---|---|---|---|
| 487 | Poly-beta-1, 6-N-acetyl-D-glucosamine N-deacetylase PgaB (Polysaccharide deacetylase) | *Acinetobacter baumannii* | | A0A6I1NXE1 | -- | 26170289 | Hydrolase activity, acting on carbon-nitrogen (but not peptide) bonds | Not Available |
| 488 | Hemin storage protein (Poly-beta-1, 6-N-acetyl-D-glucosamine N-deacetylase PgaB) | *Acinetobacter baumannii* | | A0A090C165 | -- | 25136007; 26170289 | Hydrolase activity, acting on carbon-nitrogen (but not peptide) bonds | Not Available |
| 489 | Surface composition regulator | *Salmonella enterica I* | | A0A3U0IR89 | -- | 30286803 | Not Available | Not Available |
| 490 | Poly-beta-1, 6-N-acetyl-D-glucosamine synthase (Poly-beta-1, 6-GlcNAc synthase) (EC 2.4.1.-) | *Pseudomonas baetica* | | A0A2N0DQC6 | -- | 29579234 | Acetyl glucosaminyl transferase activity | Not Available |
| 491 | Biofilm PGA synthesis protein PgaD (Poly-beta-1, 6-N-acetyl-D-glucosamine biosynthesis protein PgaD) | *Pseudomonas baetica* | | A0A2N0DQB9 | -- | 29579234 | Not Available | Not Available |
| 492 | Poly-beta-1, 6-N-acetyl-D-glucosamine biosynthesis protein PgaD | *Xanthomonas cucurbitae* | | A0A2S7DKA1 | -- | 33734871 | Not Available | Not Available |
| 493 | Poly-beta-1, 6-N-acetyl-D-glucosamine biosynthesis protein PgaD | *Pseudomonas proteolytica* | | A0A7Y1H7A3 | -- | 32587583 | Not Available | Not Available |
| 494 | Motility quorum-sensing regulator / GCU-specific mRNA interferase toxin (Type II toxin-antitoxin system MqsR family toxin) | *Pseudomonas baetica* | | A0A2N0DMT7 | -- | 29579234 | Not Available | Not Available |
| 495 | Surface composition regulator | *Salmonella enterica subspp. enterica serovar* | Mikawasima | A0A5J0RCF5 | -- | 30286803 | Not Available | Not Available |
| 496 | Surface composition regulator | *Salmonella enterica subspp. diarizonae serovar* | 48:i:z | A0A735VU22 | -- | 30286803 | Not Available | Not Available |
| 497 | Probable regulator protein RsaL (RsaL) (RsaL protein) | *Pseudomonas putida* | Arthrobacter Siderocapsulatus | Q8GEL9 | -- | 12450862; 15345437; 16585751 | Not Available | Not Available |
| 498 | Uncharacterized protein | *Desulfuromona spp.* | | A0A357YHN4 | -- | 30148503 | Not Available | Not Available |
| 499 | Hemin storage protein (Poly-beta-1, 6-N-acetyl-D-glucosamine biosynthesis protein PgaD) | *Stenotrophomonas maltophilia* | Pseudomonas maltophilia Xanthomonas maltophilia | A0A0K2IM64 | -- | 26324280 | Not Available | Not Available |
| 500 | Surface composition regulator | *Salmonella enterica subspp. enterica serovar* | 4, [5], 12:b:- | A0A741FNZ1 | -- | 30286803 | Not Available | Not Available |
| 501 | Surface composition regulator | *Salmonella enterica subspp. houtenae serovar* | 45:g, z51:- | A0A701UF81 | -- | 30286803 | Not Available | Not Available |
| 502 | Putative cytoplasmic protein (Transposase) | *Escherichia coli* | | B0LHH0 | -- | 18055599; 19717626 | Sequence-specific DNA binding | Not Available |
| 503 | Poly-beta-1, 6-N-acetyl-D-glucosamine N-deacetylase PgaB | *Pseudomonas spp. GM79* | | J2X5H7 | -- | 23045501 | Hydrolase activity, acting on carbon-nitrogen (but not peptide) bonds | Not Available |
| 504 | mRNA interferase MqsR | *Photorhabdus spp. S10-54* | | A0A329WE01 | -- | 29877789 | Not Available | Not Available |
| 505 | Poly-beta-1, 6-N-acetyl-D-glucosamine synthase (Poly-beta-1, 6-GlcNAc synthase) (EC 2.4.1.-) | *Pseudomonas spp. GM79* | | J3IHV5 | -- | 23045501 | Acetyl glucosaminyl transferase activity | Not Available |
| 506 | Uncharacterized protein | *Gammaproteobacteria bacterium* | | A0A3C0I1D7 | -- | 30148503 | Not Available | Not Available |
| 507 | Poly-beta-1, 6-N-acetyl-D-glucosamine synthase (Poly-beta-1, 6-GlcNAc synthase) (EC 2.4.1.-) | *Gammaproteobacteria bacterium* | | A0A7C5JBS2 | -- | 31911466 | Acetyl glucosaminyl transferase activity | Not Available |
| 508 | Uncharacterized protein | *Gammaproteobacteria bacterium* | | A0A7C1MU14 | -- | 31911466 | Not Available | Not Available |
| 509 | Poly-beta-1, 6-N-acetyl-D-glucosamine N-deacetylase PgaB (Fragment) | *Gammaproteobacteria bacterium* | | A0A3D1S1W2 | -- | 30148503 | Hydrolase activity, acting on carbon-nitrogen (but not peptide) bonds | Not Available |
| 510 | Poly-beta-1, 6-N-acetyl-D-glucosamine N-deacetylase PgaB | *Gammaproteobacteria bacterium* | | A0A7C1XX91 | -- | 31911466 | Hydrolase activity, acting on carbon-nitrogen (but not peptide) bonds | Not Available |
| 511 | Poly-beta-1, 6-N-acetyl-D-glucosamine synthase (Poly-beta-1, 6-GlcNAc synthase) (EC 2.4.1.-) | *Gammaproteobacteria bacterium* | | A0A7C1TYP8 | -- | 31911466 | Acetyl glucosaminyl transferase activity | Not Available |
| 512 | Uncharacterized protein | *Treponema denticola* | Strain ATCC 35405 / DSM 14222 / CIP 103919 / JCM 8153 / KCTC 15104 | Q73JV3 | -- | 15064399 | Not Available | Not Available |
| 513 | Poly-beta-1, 6-N-acetyl-D-glucosamine biosynthesis protein PgaD | *Gammaproteobacteria bacterium* | | A0A7C5JCE7 | -- | 31911466 | Not Available | Not Available |
| 514 | Poly-beta-1, 6-N-acetyl-D-glucosamine synthase (Poly-beta-1, 6-GlcNAc synthase) (EC 2.4.1.-) | *Metakosakonia spp. MRY16-398* | | A0A348DF59 | -- | 30524415 | Acetyl glucosaminyl transferase activity | Not Available |
| 515 | Poly-beta-1, 6-N-acetyl-D-glucosamine N-deacetylase PgaB (Fragment) | *Gammaproteobacteria bacterium* | | A0A7C6AWG6 | -- | 31911466 | Hydrolase activity, acting on carbon-nitrogen (but not peptide) bonds | Not Available |
| 516 | Poly-beta-1, 6-N-acetyl-D-glucosamine N-deacetylase PgaB | *Gammaproteobacteria bacterium* | | A0A7C3CT95 | -- | 31911466 | Hydrolase activity, acting on carbon-nitrogen (but not peptide) bonds | Not Available |
| 517 | Poly-beta-1, 6-N-acetyl-D-glucosamine synthase (Poly-beta-1, 6-GlcNAc synthase) (EC 2.4.1.-) | *Gammaproteobacteria bacterium* | | A0A3D5AXG0 | -- | 30148503 | Acetyl glucosaminyl transferase activity | Not Available |
| 518 | Poly-beta-1, 6-N-acetyl-D-glucosamine biosynthesis protein PgaD | *Gammaproteobacteria bacterium* | | A0A3D5AWF3 | -- | 30148503 | Not Available | Not Available |
| 519 | Poly-beta-1, 6-N-acetyl-D-glucosamine biosynthesis protein PgaD | *Gammaproteobacteria bacterium* | | A0A7C1XYQ8 | -- | 31911466 | Not Available | Not Available |
| 520 | Fimbrial subunit CupA1 | *Pseudomonas aeruginosa* | Strain ATCC 15692 / DSM 22644 / CIP 104116 / JCM 14847 / LMG 12228 / 1C / PRS 101 / PAO1 | Q9I1Y7 | -- | 10984043 | Not Available | Not Available |
| 521 | Poly-beta-1, 6-N-acetyl-D-glucosamine biosynthesis protein PgaD | *Stenotrophomonas spp.* | | A0A349Y4K3 | -- | 30148503 | Not Available | Not Available |
| 522 | Poly-beta-1, 6-N-acetyl-D-glucosamine biosynthesis protein PgaD | *Stenotrophomonas spp.* | | A0A356JDL1 | -- | 30148503 | Not Available | Not Available |
| 523 | Poly-beta-1, 6-N-acetyl-D-glucosamine biosynthesis protein PgaD | *Deltaproteobacteria bacterium RBG_13_60_28* | | A0A1F9BIK0 | -- | 27774985 | Not Available | Not Available |
| 524 | Surface composition regulator | *Shigella flexneri serotype X* | Strain 2002017 | D2A7N5 | -- | 19955273 | Not Available | Not Available |
| 525 | GGDEF domain-containing protein | *Pseudomonas aeruginosa* | Strain ATCC 15692 / DSM 22644 / CIP 104116 / JCM 14847 / LMG 12228 / 1C / PRS 101 / PAO1 | Q9I6W3 | -- | 10984043 | Diguanylate cyclase activity | Not Available |
| 526 | GGDEF domain-containing protein | *Pseudomonas aeruginosa* | Strain ATCC 15692 / DSM 22644 / CIP 104116 / JCM 14847 / LMG 12228 / 1C / PRS 101 / PAO1 | Q9HYQ2 | -- | 10984043 | Diguanylate cyclase activity | Not Available |
| 527 | GGDEF domain-containing protein | *Pseudomonas aeruginosa* | Strain ATCC 15692 / DSM 22644 / CIP 104116 / JCM 14847 / LMG 12228 / 1C / PRS 101 / PAO1 | Q9I4M8 | -- | 10984043 | Diguanylate cyclase activity | Not Available |

| Target ID | Bacterial Protein Name | Organism Name | Strain | UniProt ID | PDB ID | PubMed ID | Classification as per UniProt GO Molecular Function | Classification of Function as per PDB |
|---|---|---|---|---|---|---|---|---|
| 528 | Exodeoxyribonuclease III (EC 3.1.11.2) | *Pseudomonas aeruginosa* | Strain ATCC 15692 / DSM 22644 / CIP 104116 / JCM 14847 / LMG 12228 / 1C / PRS 101 / PAO1 | G3XDB0 | -- | 10984043 | DNA-(apurinic or apyrimidinic site) endonuclease activity, double-stranded DNA 3'-5' exodeoxyribonuclease activity, exodeoxyribonuclease III activity, phosphoric diester Hydrolase activity | Not Available |
| 529 | Fimbrial domain-containing protein | *Pseudomonas aeruginosa* | Strain ATCC 15692 / DSM 22644 / CIP 104116 / JCM 14847 / LMG 12228 / 1C / PRS 101 / PAO1 | Q9HTS0 | -- | 10984043 | Not Available | Not Available |
| 530 | Probable fimbrial subunit CupB1 | *Pseudomonas aeruginosa* | Strain ATCC 15692 / DSM 22644 / CIP 104116 / JCM 14847 / LMG 12228 / 1C / PRS 101 / PAO1 | Q9HWU2 | -- | 10984043 | Not Available | Not Available |
| 531 | Fimbrial subunit CupC1 | *Pseudomonas aeruginosa* | Strain ATCC 15692 / DSM 22644 / CIP 104116 / JCM 14847 / LMG 12228 / 1C / PRS 101 / PAO1 | Q9I4X7 | -- | 10984043 | Not Available | Not Available |
| 532 | GGDEF domain-containing protein | *Pseudomonas aeruginosa* | Strain ATCC 15692 / DSM 22644 / CIP 104116 / JCM 14847 / LMG 12228 / 1C / PRS 101 / PAO1 | Q9HZ57 | -- | 10984043 | Diguanylate cyclase activity | Not Available |
| 533 | 50S ribosomal subunit assembly factor BipA (EC 3.6.5.-) (GTP-binding protein BipA) | *Pseudomonas aeruginosa* | Strain ATCC 15692 / DSM 22644 / CIP 104116 / JCM 14847 / LMG 12228 / 1C / PRS 101 / PAO1 | Q9HU67 | -- | 10984043 | GTP binding, GTPase activity, ribosome binding, rRNA binding, tRNA binding | Not Available |
| 534 | Uncharacterized protein | *Pseudomonas aeruginosa* | Strain ATCC 15692 / DSM 22644 / CIP 104116 / JCM 14847 / LMG 12228 / 1C / PRS 101 / PAO1 | Q9I6K0 | -- | 10984043 | Diguanylate cyclase activity | Not Available |
| 535 | Regulatory protein RsaL | *Pseudomonas aeruginosa* | Strain ATCC 15692 / DSM 22644 / CIP 104116 / JCM 14847 / LMG 12228 / 1C / PRS 101 / PAO1 | G3XD78 | -- | 10984043 | DNA binding, DNA-binding Transcription repressor activity, Transcription cis-regulatory region binding | Not Available |
| 536 | Uncharacterized protein | *Xylella fastidiosa* | Strain M23 | B2I812 | -- | 20601474 | Not Available | Not Available |
| 537 | Uncharacterized protein | *Pseudomonas spp. GM79* | | J2WV44 | -- | 23045501 | Not Available | Not Available |
| 538 | Poly-beta-1, 6-N-acetyl-D-glucosamine biosynthesis protein PgaD | *Pseudomonas spp. GM79* | | J3DHL5 | -- | 23045501 | Not Available | Not Available |
| 539 | Biofilm PGA synthesis deacetylase PgaB | *Acinetobacter baumannii NCGM 237* | | A0A0E1PPJ3 | -- | 24550340 | Hydrolase activity, acting on carbon-nitrogen (but not peptide) bonds | Not Available |
| 540 | Poly-beta-1, 6-N-acetyl-D-glucosamine synthase (Poly-beta-1, 6-GlcNAc synthase) (EC 2.4.1.-) | *Acinetobacter baumannii NCGM 237* | | A0A0E1PT59 | -- | 24550340 | Acetyl glucosaminyl transferase activity | Not Available |
| 541 | Poly-beta-1, 6-N-acetyl-D-glucosamine N-deacetylase PgaB | *Acinetobacter baumannii NCGM 237* | | A0A0E1PVR4 | -- | 24550340 | Hydrolase activity, acting on carbon-nitrogen (but not peptide) bonds | Not Available |
| 542 | Poly-beta-1, 6-N-acetyl-D-glucosamine auxiliary protein PgaD | *Acinetobacter baumannii NCGM 237* | | A0A0E1PU81 | -- | 24550340 | Not Available | Not Available |
| 543 | PgaB | *Pseudomonas ogarae* | Strain DSM 112162 / CECT 30235 / F113 | G8PY99 | -- | 22328765 | Hydrolase activity, acting on carbon-nitrogen (but not peptide) bonds | Not Available |
| 544 | PgaD | *Pseudomonas ogarae* | Strain DSM 112162 / CECT 30235 / F113 | G8PYA1 | -- | 22328765 | Not Available | Not Available |
| 545 | Poly-beta-1, 6-N-acetyl-D-glucosamine synthase (Poly-beta-1, 6-GlcNAc synthase) (EC 2.4.1.-) | *Pseudomonas ogarae* | Strain DSM 112162 / CECT 30235 / F113 | G8PYA0 | -- | 22328765 | Acetyl glucosaminyl transferase activity | Not Available |
| 546 | Poly-beta-1, 6-N-acetyl-D-glucosamine synthase (Poly-beta-1, 6-GlcNAc synthase) (EC 2.4.1.-) | *Acinetobacter baumannii NCGM 237* | | A0A0E1PS38 | -- | 24550340 | Acetyl glucosaminyl transferase activity | Not Available |
| 547 | Toxin-antitoxin system | *Pseudomonas ogarae* | Strain DSM 112162 / CECT 30235 / F113 | G8Q9Z1 | -- | 22328765 | Not Available | Not Available |
| 548 | Poly-beta-1, 6-N-acetyl-D-glucosamine synthase (Poly-beta-1, 6-GlcNAc synthase) (EC 2.4.1.-) | *Candidatus Competibacteraceae bacterium* | | A0A433I7B0 | -- | 30578268 | Acetyl glucosaminyl transferase activity | Not Available |
| 549 | Poly-beta-1, 6-N-acetyl-D-glucosamine biosynthesis protein PgaD | *Candidatus Competibacteraceae bacterium* | | A0A3S0WCV8 | -- | 30578268 | Not Available | Not Available |
| 550 | Poly-beta-1, 6-N-acetyl-D-glucosamine biosynthesis protein PgaD | *Pseudomonas spp. GM24* | | J2QNC2 | -- | 23045501 | Not Available | Not Available |
| 551 | Poly-beta-1, 6-N-acetyl-D-glucosamine N-deacetylase PgaB | *Candidatus Competibacteraceae bacterium* | | A0A3S0W208 | -- | 30578268 | Hydrolase activity, acting on carbon-nitrogen (but not peptide) bonds | Not Available |
| 552 | Poly-beta-1, 6-N-acetyl-D-glucosamine synthase (Poly-beta-1, 6-GlcNAc synthase) (EC 2.4.1.-) | *Pseudomonas spp. GM24* | | J3FT73 | -- | 23045501 | Acetyl glucosaminyl transferase activity | Not Available |
| 553 | Biofilm PGA synthesis auxiliary protein PgaD | *Acinetobacter baumannii NCGM 237* | | A0A0E1PVN7 | -- | 24550340 | Not Available | Not Available |
| 554 | Poly-beta-1, 6-N-acetyl-D-glucosamine N-deacetylase PgaB | *Pseudomonas spp. GM24* | | J2R4Q4 | -- | 23045501 | Hydrolase activity, acting on carbon-nitrogen (but not peptide) bonds | Not Available |
| 555 | Uncharacterized protein | *Kangiella koreensis* | Strain DSM 16069 / KCTC 12182 / SW-125 | C7RBF2 | -- | 21304661 | Not Available | Not Available |
| 556 | Uncharacterized protein | *Pseudomonas spp. TCU-HL1* | | A0A1B3E8H3 | -- | 27542933 | Not Available | Not Available |
| 557 | Uncharacterized protein | *Candidatus Lambdaproteobacteria bacterium RIFOXYD2_FULL_56_26* | | A0A1F6H0I8 | -- | 27774985 | Not Available | Not Available |
| 558 | Poly-beta-1, 6-N-acetyl-D-glucosamine synthase (Poly-beta-1, 6-GlcNAc synthase) (EC 2.4.1.-) | *Acinetobacter spp. ACNIH2* | | A0A2L0HWN6 | -- | 29437920 | Acetyl glucosaminyl transferase activity | Not Available |
| 559 | Poly-beta-1, 6-N-acetyl-D-glucosamine synthase (Poly-beta-1, 6-GlcNAc synthase) (EC 2.4.1.-) | *Carnobacterium divergens* | Lactobacillus divergens | A0A7I0FF87 | -- | 30081251 | Acetyl glucosaminyl transferase activity | Not Available |
| 560 | Poly-beta-1, 6-N-acetyl-D-glucosamine synthase (Poly-beta-1, 6-GlcNAc synthase) (EC 2.4.1.-) | *Lysobacter enzymogenes* | | A0A1J1EEU3 | -- | 28065880 | Acetyl glucosaminyl transferase activity | Not Available |

| Target ID | Bacterial Protein Name | Organism Name | Strain | UniProt ID | PDB ID | PubMed ID | Classification as per UniProt GO Molecular Function | Classification of Function as per PDB |
|---|---|---|---|---|---|---|---|---|
| 561 | HmsF protein | *Lysobacter enzymogenes* | | A0A1J1EKH2 | -- | 28065880 | Hydrolase activity, acting on carbon-nitrogen (but not peptide) bonds | Not Available |
| 562 | Motility quorum-sensing regulator MqsR | *Pseudomonas moraviensis R28-S* | | V8R8K1 | -- | 24558233 | Not Available | Not Available |
| 563 | Surface composition regulator | *Salmonella enterica subspp. enterica serovar* | Senftenberg Str. A4-543 | G5R5Q1 | -- | 21859443 | Not Available | Not Available |
| 564 | Polysaccharide deacetylase | *Kangiella koreensis* | Strain DSM 16069 / KCTC 12182 / SW-125 | C7RBF0 | -- | 21304661 | Hydrolase activity, acting on carbon-nitrogen (but not peptide) bonds | Not Available |
| 565 | Uncharacterized protein | *Pseudomonas spp. TCU-HL1* | | A0A1B3E8H9 | -- | 27542933 | Not Available | Not Available |
| 566 | Poly-beta-1, 6-N-acetyl-D-glucosamine biosynthesis protein PgaD | *Acinetobacter spp. ACNIH2* | | A0A2L0HWJ0 | -- | 29437920 | Not Available | Not Available |
| 567 | Uncharacterized protein | *Achromobacter xylosoxidans* | Strain A8 | E3HKX9 | -- | 21097610 | Not Available | Not Available |
| 568 | Uncharacterized protein | *Thioalkalivibrio sulfidiphilus* | Strain HL-EbGR7 | B8GTH7 | -- | 21475584 | Not Available | Not Available |
| 569 | Poly-beta-1, 6-N-acetyl-D-glucosamine biosynthesis protein PgaD | *Variovorax spp. CF313* | | J2JIW7 | -- | 23045501 | Not Available | Not Available |
| 570 | Poly-beta-1, 6-N-acetyl-D-glucosamine synthase (Poly-beta-1, 6-GlcNAc synthase) (EC 2.4.1.-) | *Ralstonia solanacearum* | Strain GMI1000 Pseudomonas solanacearum | Q8XT27 | -- | 11823852 | Acetyl glucosaminyl transferase activity | Not Available |
| 571 | Poly-beta-1, 6-N-acetyl-D-glucosamine biosynthesis protein PgaD | *Serratia marcescens* | | A0A1L6QPC1 | -- | 28536292 | Not Available | Not Available |
| 572 | Poly-beta-1, 6-N-acetyl-D-glucosamine N-deacetylase PgaB | *Variovorax spp. CF313* | | J2SQM8 | -- | 23045501 | Hydrolase activity, acting on carbon-nitrogen (but not peptide) bonds | Not Available |
| 573 | GGDEF family protein | *Leptospira interrogans serogroup Icterohaemorrhagiae serovar Lai* | Strain 56601 | Q8F244 | -- | 12712204 | Diguanylate cyclase activity | Not Available |
| 574 | Response regulator with GGDEF domain | *Leptospira interrogans serogroup Icterohaemorrhagiae serovar Lai* | Strain 56601 | Q8EZC6 | -- | 12712204 | Diguanylate cyclase activity | Not Available |
| 575 | Poly-beta-1, 6-N-acetyl-D-glucosamine biosynthesis protein PgaD | *Enterobacteriaceae bacterium* | | A0A3C0H2G8 | -- | 30148503 | Not Available | Not Available |
| 576 | Uncharacterized protein | *Enterobacteriaceae bacterium* | | A0A3C2BJ06 | -- | 30148503 | Not Available | Not Available |
| 577 | Poly-beta-1, 6-N-acetyl-D-glucosamine biosynthesis protein PgaD | *Xenorhabdus kozodoii* | | A0A2D0LH81 | -- | 28993611 | Not Available | Not Available |
| 578 | GGDEF family protein | *Leptospira interrogans serogroup Icterohaemorrhagiae serovar Lai* | Strain 56601 | Q8F251 | -- | 12712204 | Diguanylate cyclase activity | Not Available |
| 579 | Poly-beta-1, 6-N-acetyl-D-glucosamine synthase (Poly-beta-1, 6-GlcNAc synthase) (EC 2.4.1.-) | *Xenorhabdus kozodoii* | | A0A2D0LIF9 | -- | 28993611 | Acetyl glucosaminyl transferase activity | Not Available |
| 580 | Toxin-antitoxin system, toxin component | *Enterobacteriaceae bacterium* | | A0A3C0H4J1 | -- | 30148503 | Not Available | Not Available |
| 581 | Poly-beta-1, 6-N-acetyl-D-glucosamine N-deacetylase PgaB | *Enterobacteriaceae bacterium* | | A0A3C2BHH0 | -- | 30148503 | Hydrolase activity, acting on carbon-nitrogen (but not peptide) bonds | Not Available |
| 582 | GGDEF family protein | *Leptospira interrogans serogroup Icterohaemorrhagiae serovar Lai* | Strain 56601 | Q8F248 | -- | 12712204 | Diguanylate cyclase activity | Not Available |
| 583 | GGDEF family protein | *Leptospira interrogans serogroup Icterohaemorrhagiae serovar Lai* | Strain 56601 | Q8F245 | -- | 12712204 | Diguanylate cyclase activity | Not Available |
| 584 | Uncharacterized protein | *Xylella fastidiosa* | Strain Temecula1 / ATCC 700964 | Q87EE2 | -- | 12533478 | Not Available | Not Available |
| 585 | Poly-beta-1, 6-N-acetyl-D-glucosamine biosynthesis protein PgaD | *Enterobacteriaceae bacterium* | | A0A3C2BGH1 | -- | 30148503 | Not Available | Not Available |
| 586 | GGDEF domain receiver component of a two component response regulator | *Leptospira interrogans serogroup Icterohaemorrhagiae serovar Lai* | Strain 56601 | Q8F383 | -- | 12712204 | Diguanylate cyclase activity | Not Available |
| 587 | GGDEF family protein | *Leptospira interrogans serogroup Icterohaemorrhagiae serovar Lai* | Strain 56601 | Q8EXH2 | -- | 12712204 | Diguanylate cyclase activity | Not Available |
| 588 | Surface composition regulator | *Cronobacter turicensis* | Strain DSM 18703 / CCUG 55852 / LMG 23827 / z3032 | C9Y2V2 | -- | 21037008 | Not Available | Not Available |
| 589 | GGDEF family protein | *Leptospira interrogans serogroup Icterohaemorrhagiae serovar Lai* | Strain 56601 | Q8F246 | -- | 12712204 | Diguanylate cyclase activity | Not Available |
| 590 | GGDEF family protein | *Leptospira interrogans serogroup Icterohaemorrhagiae serovar Lai* | Strain 56601 | Q8F247 | -- | 12712204 | Diguanylate cyclase activity | Not Available |
| 591 | Poly-beta-1, 6-N-acetyl-D-glucosamine N-deacetylase PgaB | *Enterobacteriaceae bacterium* | | A0A3C0H2V4 | -- | 30148503 | Hydrolase activity, acting on carbon-nitrogen (but not peptide) bonds | Not Available |
| 592 | Poly-beta-1, 6-N-acetyl-D-glucosamine biosynthesis protein PgaD | *Enterobacteriaceae bacterium* | | A0A351NN88 | -- | 30148503 | Not Available | Not Available |
| 593 | Poly-beta-1, 6-N-acetyl-D-glucosamine N-deacetylase PgaB | *Enterobacteriaceae bacterium* | | A0A351NN90 | -- | 30148503 | Hydrolase activity, acting on carbon-nitrogen (but not peptide) bonds | Not Available |
| 594 | Polysaccharide deacetylase | *Xenorhabdus kozodoii* | | A0A2D0LH91 | -- | 28993611 | Hydrolase activity, acting on carbon-nitrogen (but not peptide) bonds | Not Available |
| 595 | Sensory box / GGDEF family protein | *Bacillus cereus* | Strain ATCC 14579 / DSM 31 / CCUG 7414 / JCM 2152 / NBRC 15305 / NCIMB 9373 / NCTC 2599 / NRRL B-3711 | Q819E1 | -- | 12721630 | Diguanylate cyclase activity | Not Available |
| 596 | Poly-beta-1, 6-N-acetyl-D-glucosamine biosynthesis protein PgaD | *Enterobacter cloacae complex spp. ECNIH11* | | A0A2S4SKF2 | -- | 29437920 | Not Available | Not Available |
| 597 | Poly-beta-1, 6-N-acetyl-D-glucosamine synthase (Poly-beta-1, 6-GlcNAc synthase) (EC 2.4.1.-) | *Enterobacteriaceae bacterium* | | A0A351NN89 | -- | 30148503 | Acetyl glucosaminyl transferase activity | Not Available |
| 598 | Biofilm PGA synthesis deacetylase PgaB | *Paraburkholderia caribensis MBA4* | | A0A0P0RAM0 | -- | 24558235 | Hydrolase activity, acting on carbon-nitrogen (but not peptide) bonds | Not Available |

| Target ID | Bacterial Protein Name | Organism Name | Strain | UniProt ID | PDB ID | PubMed ID | Classification as per UniProt GO Molecular Function | Classification of Function as per PDB |
|---|---|---|---|---|---|---|---|---|
| 599 | GGDEF family protein | *Leptospira interrogans serogroup Icterohaemorrhagiae serovar Lai* | Strain 56601 | Q8EXH5 | -- | 12712204 | Diguanylate cyclase activity | Not Available |
| 600 | GGDEF family protein | *Leptospira interrogans serogroup Icterohaemorrhagiae serovar Lai* | Strain 56601 | Q8F249 | -- | 12712204 | Diguanylate cyclase activity | Not Available |
| 601 | Poly-beta-1, 6-N-acetyl-D-glucosamine synthase (Poly-beta-1, 6-GlcNAc synthase) (EC 2.4.1.-) | *Aquitalea magnusonii* | | A0A3G9GKV2 | -- | 28293292 | Acetyl glucosaminyl transferase activity | Not Available |
| 602 | Poly-beta-1, 6-N-acetyl-D-glucosamine biosynthesis protein PgaD | *Xanthomonas vasicola pv. zeae* | | A0A7C7F7Z7 | -- | 32040377 | Not Available | Not Available |
| 603 | Biofilm PGA synthesis deacetylase PgaB | *Aquitalea magnusonii* | | A0A3G9GKG4 | -- | 28293292 | Hydrolase activity, acting on carbon-nitrogen (but not peptide) bonds | Not Available |
| 604 | Biofilm PGA synthesis protein PgaD | *Luteimona cucumeris* | | A0A562L7U6 | -- | 26203337 | Not Available | Not Available |
| 605 | Poly-beta-1, 6-N-acetyl-D-glucosamine N-deacetylase PgaB | *Xanthomonas vasicola pv. zeae* | | A0A7C7B1H5 | -- | 32040377 | Hydrolase activity, acting on carbon-nitrogen (but not peptide) bonds | Not Available |
| 606 | Poly-beta-1, 6-N-acetyl-D-glucosamine N-deacetylase PgaB (EC 3.5.1.-) | *Pseudomonas spp. WS 5503* | | A0A7Y0WES9 | -- | 32587583 | Hydrolase activity, acting on carbon-nitrogen (but not peptide) bonds | Not Available |
| 607 | Poly-beta-1, 6-N-acetyl-D-glucosamine biosynthesis protein PgaD | *Pseudomonas spp. WS 5503* | | A0A7Y0WEE3 | -- | 32587583 | Not Available | Not Available |
| 608 | HmsF protein | *Xanthomonas axonopodis pv. citri* | Strain 306 | Q8PLI6 | -- | 12024217 | Hydrolase activity, acting on carbon-nitrogen (but not peptide) bonds | Not Available |
| 609 | GGDEF domain-containing protein | *Rhodopirellula baltica* | Strain DSM 10527 / NCIMB 13988 / SH1 | Q7UXF6 | -- | 12835416 | Diguanylate cyclase activity | Not Available |
| 610 | Probable response regulatory protein | *Rhodopirellula baltica* | Strain DSM 10527 / NCIMB 13988 / SH1 | Q7UF69 | -- | 12835416 | Diguanylate cyclase activity | Not Available |
| 611 | Uncharacterized protein | *Xanthomonas axonopodis pv. citri* | Strain 306 | Q8PLI8 | -- | 12024217 | Not Available | Not Available |
| 612 | Probable response regulatory protein | *Rhodopirellula baltica* | Strain DSM 10527 / NCIMB 13988 / SH1 | Q7UR09 | -- | 12835416 | Diguanylate cyclase activity | Not Available |
| 613 | Poly-beta-1, 6-N-acetyl-D-glucosamine biosynthesis protein PgaD | *Tatumella spp. OPLPL6* | | A0A2G8BYZ7 | -- | 29067021 | Not Available | Not Available |
| 614 | Poly-beta-1, 6-N-acetyl-D-glucosamine synthase (Poly-beta-1, 6-GlcNAc synthase) (EC 2.4.1.-) | *Enterobacter cloacae complex spp. ECNIH11* | | A0A2S4SKE8 | -- | 29437920 | Acetyl glucosaminyl transferase activity | Not Available |
| 615 | Poly-beta-1, 6-N-acetyl-D-glucosamine N-deacetylase PgaB | *Enterobacter cloacae complex spp. ECNIH11* | | A0A2S4SKH4 | -- | 29437920 | Hydrolase activity, acting on carbon-nitrogen (but not peptide) bonds | Not Available |
| 616 | Conserved hypothetical Membrane protein | *Pseudomonas fluorescens* | Strain SBW25 | C3KAG0 | -- | 19432983 | Not Available | Not Available |
| 617 | Putative lipoprotein | *Pseudomonas fluorescens* | Strain SBW25 | C3KAF8 | -- | 19432983 | Hydrolase activity, acting on carbon-nitrogen (but not peptide) bonds | Not Available |
| 618 | Poly-beta-1, 6-N-acetyl-D-glucosamine synthase (Poly-beta-1, 6-GlcNAc synthase) (EC 2.4.1.-) (Fragment) | *Lentisphaerae bacterium GWF2_44_16* | | A0A1G0YRV0 | -- | 27774985 | Acetyl glucosaminyl transferase activity | Not Available |
| 619 | Surface composition regulator | *Enterobacter cloacae complex spp. ECNIH11* | | A0A2S4SUC2 | -- | 29437920 | Not Available | Not Available |
| 620 | Haemin storage system, HmsF protein | *Yersinia pseudotuberculosis serotype I* | Strain IP32953 | Q66B30 | -- | 15358858 | Hydrolase activity, acting on carbon-nitrogen (but not peptide) bonds | Not Available |
| 621 | Poly-beta-1, 6-N-acetyl-D-glucosamine synthase (Poly-beta-1, 6-GlcNAc synthase) (EC 2.4.1.-) | *Luteibacter yeojuensis* | | A0A7X5QS05 | -- | 16957102 | Acetyl glucosaminyl transferase activity | Not Available |
| 622 | Poly-beta-1, 6-N-acetyl-D-glucosamine N-deacetylase PgaB | *Luteibacter yeojuensis* | | A0A7X5QS01 | -- | 16957102 | Hydrolase activity, acting on carbon-nitrogen (but not peptide) bonds | Not Available |
| 623 | Surface composition regulator | *Serratia plymuthica S13* | | S4YI83 | -- | 23929484 | Not Available | Not Available |
| 624 | Biofilm PGA synthesis lipoprotein PgaB | *Luteimona cucumeris* | | A0A562L7K0 | -- | 26203337 | Hydrolase activity, acting on carbon-nitrogen (but not peptide) bonds | Not Available |
| 625 | PgaB protein | *Escherichia coli* | Strain 55989 / EAEC | B7LFD3 | -- | 19165319 | Hydrolase activity, acting on carbon-nitrogen (but not peptide) bonds | Not Available |
| 626 | Poly-beta-1, 6-N-acetyl-D-glucosamine biosynthesis protein PgaD | *Luteibacter yeojuensis* | | A0A7X5QS23 | -- | 16957102 | Not Available | Not Available |
| 627 | Poly-beta-1, 6-N-acetyl-D-glucosamine synthase (Poly-beta-1, 6-GlcNAc synthase) (EC 2.4.1.-) | *Escherichia coli O6:K15:H31* | Strain 536 / UPEC | A0A454A2Y6 | -- | 16879640 | Acetyl glucosaminyl transferase activity | Not Available |
| 628 | Haemin storage system, HmsS protein | *Yersinia pseudotuberculosis serotype I* | Strain IP32953 | Q66B28 | -- | 15358858 | Not Available | Not Available |
| 629 | Biofilm PGA synthesis lipoprotein PgaB (EC 3.-.-.-) | *Klebsiella pneumoniae* | Strain 342 | B5Y382 | -- | 18654632 | Hydrolase activity, acting on carbon-nitrogen (but not peptide) bonds | Not Available |
| 630 | Poly-beta-1, 6-N-acetyl-D-glucosamine synthase (Poly-beta-1, 6-GlcNAc synthase) (EC 2.4.1.-) | *Escherichia coli* | Strain 55989 / EAEC | B7LFD2 | -- | 19165319 | Acetyl glucosaminyl transferase activity | Not Available |
| 631 | Surface composition regulator | *Klebsiella pneumoniae* | Strain 342 | B5XU41 | -- | 18654632 | Not Available | Not Available |
| 632 | Poly-beta-1, 6-N-acetyl-D-glucosamine N-deacetylase PgaB (EC 3.5.1.-) | *Pseudomonas spp. WS 5027* | | A0A7Y1B5A9 | -- | 32587583 | Hydrolase activity, acting on carbon-nitrogen (but not peptide) bonds | Not Available |
| 633 | Putative biofilm PGA synthesis protein PgaD | *Klebsiella pneumoniae* | Strain 342 | B5Y384 | -- | 18654632 | Not Available | Not Available |
| 634 | Poly-beta-1, 6-N-acetyl-D-glucosamine synthase (Poly-beta-1, 6-GlcNAc synthase) (EC 2.4.1.-) | *Luteimona cucumeris* | | A0A562L7D7 | -- | 26203337 | Acetyl glucosaminyl transferase activity | Not Available |
| 635 | Poly-beta-1, 6-N-acetyl-D-glucosamine synthase (Poly-beta-1, 6-GlcNAc synthase) (EC 2.4.1.-) | *Xanthomonas vasicola pv. zeae* | | A0A7C7B0Q2 | -- | 32040377 | Acetyl glucosaminyl transferase activity | Not Available |
| 636 | Poly-beta-1, 6-N-acetyl-D-glucosamine synthase (Poly-beta-1, 6-GlcNAc synthase) (EC 2.4.1.-) | *Klebsiella pneumoniae* | Strain 342 | B5Y383 | -- | 18654632 | Acetyl glucosaminyl transferase activity | Not Available |
| 637 | Poly-beta-1, 6-N-acetyl-D-glucosamine N-deacetylase PgaB | *Acinetobacter cumulans* | | A0A4P8DU29 | -- | 30808586 | Hydrolase activity, acting on carbon-nitrogen (but not peptide) bonds | Not Available |
| 638 | Poly-beta-1, 6-N-acetyl-D-glucosamine biosynthesis protein PgaD | *Candidatus Muproteobacteria bacterium RBG_19FT_COMBO_61_10* | | A0A1F6UP26 | -- | 27774985 | Not Available | Not Available |
| 639 | Surface composition regulator | *Cronobacter malonaticus* | | A0A423XSW2 | -- | 30416493 | Not Available | Not Available |
| 640 | Poly-beta-1, 6-N-acetyl-D-glucosamine synthase (Poly-beta-1, 6-GlcNAc synthase) (EC 2.4.1.-) | *Acinetobacter baumannii* | Strain AB0057 | A0A7U4DG52 | -- | 18931120 | Acetyl glucosaminyl transferase activity | Not Available |
| 641 | Surface composition regulator | *Cronobacter malonaticus* | | V5U106 | -- | 24435860 | Not Available | Not Available |
| 642 | Uncharacterized protein | *Syntrophobacter spp. DG_60* | | A0A0S7XQF1 | -- | 25922666 | Not Available | Not Available |
| 643 | mRNA interferase MqsR | *Paraburkholderia madseniana* | | A0A6N6W8U5 | -- | 32027304 | Not Available | Not Available |
| 644 | Poly-beta-1, 6-N-acetyl-D-glucosamine synthase (Poly-beta-1, 6-GlcNAc synthase) (EC 2.4.1.-) | *Sulfurimona spp. RIFOXYD2_FULL_34_21* | | A0A1G3UF66 | -- | 27774985 | Acetyl glucosaminyl transferase activity | Not Available |
| 645 | Fimbrial protein | *Yersinia pestis* | | A0A384KL23 | -- | 11586360 | Not Available | Not Available |
| 646 | Poly-beta-1, 6-N-acetyl-D-glucosamine synthase (Poly-beta-1, 6-GlcNAc synthase) (EC 2.4.1.-) | *Citrobacter freundii complex spp. CFNIH4* | | A0A2L0TFK3 | -- | 29437920 | Acetyl glucosaminyl transferase activity | Not Available |

| Target ID | Bacterial Protein Name | Organism Name | Strain | UniProt ID | PDB ID | PubMed ID | Classification as per UniProt GO Molecular Function | Classification of Function as per PDB |
|---|---|---|---|---|---|---|---|---|
| 647 | Hypothetical lipoprotein YcdR | *Escherichia coli* | Strain UTI89 / UPEC | Q1RDJ4 | -- | 16585510 | Hydrolase activity, acting on carbon-nitrogen (but not peptide) bonds | Not Available |
| 648 | Poly-beta-1, 6-N-acetyl-D-glucosamine biosynthesis protein PgaD | *Citrobacter freundii complex spp. CFNIH4* | | A0A2L0TFH9 | -- | 29437920 | Not Available | Not Available |
| 649 | Poly-beta-1, 6-N-acetyl-D-glucosamine N-deacetylase PgaB | *Sulfurimona spp. RIFOXYD2_FULL_34_21* | | A0A1G3UF37 | -- | 27774985 | Hydrolase activity, acting on carbon-nitrogen (but not peptide) bonds | Not Available |
| 650 | Surface composition regulator | *Citrobacter freundii complex spp. CFNIH4* | | A0A2L0TH79 | -- | 29437920 | Not Available | Not Available |
| 651 | Poly-beta-1, 6-N-acetyl-D-glucosamine synthase (Poly-beta-1, 6-GlcNAc synthase) (EC 2.4.1.-) | *Yersinia pestis* | | Q0WFJ6 | -- | 11586360 | Acetyl glucosaminyl transferase activity | Not Available |
| 652 | Uncharacterized protein | *Escherichia coli* | Strain UTI89 / UPEC | Q1RDJ6 | -- | 16585510 | Not Available | Not Available |
| 653 | Poly-beta-1, 6-N-acetyl-D-glucosamine N-deacetylase | *Citrobacter freundii complex spp. CFNIH4* | | A0A2L0TG22 | -- | 29437920 | Hydrolase activity, acting on carbon-nitrogen (but not peptide) bonds | Not Available |
| 654 | Putative motility quorum-sensing regulator | *Magnetospirillum gryphiswaldense* | Strain DSM 6361 / JCM 21280 / NBRC 15271 / MSR-1 | V6F2G1 | -- | 24625872 | Not Available | Not Available |
| 655 | Poly-beta-1, 6-N-acetyl-D-glucosamine biosynthesis protein PgaD | *Sulfurimona spp. RIFOXYD2_FULL_34_21* | | A0A1G3UG59 | -- | 27774985 | Not Available | Not Available |
| 656 | Hypothetical cytosolic protein | *Syntrophus aciditrophicus* | Strain SB | Q2LYD0 | -- | 17442750 | Not Available | Not Available |
| 657 | Poly-beta-1, 6-N-acetyl-D-glucosamine synthase (Poly-beta-1, 6-GlcNAc synthase) (EC 2.4.1.-) | *Alcaligenaceae bacterium* | | A0A7C6NLD6 | -- | 32123542 | Acetyl glucosaminyl transferase activity | Not Available |
| 658 | Poly-beta-1, 6-N-acetyl-D-glucosamine biosynthesis protein PgaD | *Alcaligenaceae bacterium* | | A0A7C6NL24 | -- | 32123542 | Not Available | Not Available |
| 659 | Putative membrane protein | *Xanthomonas campestris pv. campestris* | Strain B100 | A0A1X7QFS0 | -- | 18304669 | Not Available | Not Available |
| 660 | Uncharacterized protein | *Rhodanobacter spp. 115* | | I4VP81 | -- | 22843592 | Not Available | Not Available |
| 661 | Poly-beta-1, 6-N-acetyl-D-glucosamine synthase (Poly-beta-1, 6-GlcNAc synthase) (EC 2.4.1.-) | *Pasteurellaceae bacterium* | | A0A356E647 | -- | 30148503 | Acetyl glucosaminyl transferase activity | Not Available |
| 662 | Poly-beta-1, 6-N-acetyl-D-glucosamine biosynthesis protein PgaD | *Alcaligenaceae bacterium* | | A0A2D8B7A5 | -- | 29337314 | Not Available | Not Available |
| 663 | Poly-beta-1, 6-N-acetyl-D-glucosamine N-deacetylase PgaB (Fragment) | *Pasteurellaceae bacterium* | | A0A356EBH6 | -- | 30148503 | Hydrolase activity, acting on carbon-nitrogen (but not peptide) bonds | Not Available |
| 664 | Poly-beta-1, 6-N-acetyl-D-glucosamine synthase (Poly-beta-1, 6-GlcNAc synthase) (EC 2.4.1.-) | *Alcaligenaceae bacterium* | | A0A2D8B791 | -- | 29337314 | Acetyl glucosaminyl transferase activity | Not Available |
| 665 | Poly-beta-1, 6-N-acetyl-D-glucosamine N-deacetylase PgaB (EC 3.5.1.-) | *Alcaligenaceae bacterium* | | A0A7C6NPE6 | -- | 32123542 | Hydrolase activity, acting on carbon-nitrogen (but not peptide) bonds | Not Available |
| 666 | Poly-beta-1, 6-N-acetyl-D-glucosamine synthase (Poly-beta-1, 6-GlcNAc synthase) (EC 2.4.1.-) | *Alcaligenes spp. EGD-AK7* | | U1YHE7 | -- | 24407646 | Acetyl glucosaminyl transferase activity | Not Available |
| 667 | Poly-beta-1, 6-N-acetyl-D-glucosamine synthase (Poly-beta-1, 6-GlcNAc synthase) (EC 2.4.1.-) | *Pseudomonas trivialis* | | A0A0H5A7V6 | -- | 26337878 | Acetyl glucosaminyl transferase activity | Not Available |
| 668 | Outer membrane N-deacetylase | *Pseudomonas trivialis* | | A0A0H5ABT7 | -- | 26337878 | Hydrolase activity, acting on carbon-nitrogen (but not peptide) bonds | Not Available |
| 669 | Motility quorum-sensing regulator MqsR | *Pseudomonas trivialis* | | A0A0H5A8B6 | -- | 26337878 | Not Available | Not Available |
| 670 | Uncharacterized protein | *Sulfurimona denitrificans* | Strain ATCC 33889 / DSM 1251 Thiomicrospira denitrificans Strain ATCC 33889 / DSM 1251 | Q30SJ9 | -- | 18065616 | Not Available | Not Available |
| 671 | Uncharacterized protein | *Alcaligenes spp. EGD-AK7* | | U1XZ69 | -- | 24407646 | Not Available | Not Available |
| 672 | Poly-beta-1, 6-N-acetyl-D-glucosamine synthase (Poly-beta-1, 6-GlcNAc synthase) (EC 2.4.1.-) | *Sulfurimona denitrificans* | Strain ATCC 33889 / DSM 1251 Thiomicrospira denitrificans Strain ATCC 33889 / DSM 1251 | Q30SK0 | -- | 18065616 | Acetyl glucosaminyl transferase activity | Not Available |
| 673 | Polysaccharide deacetylase | *Sulfurimona denitrificans* | Strain ATCC 33889 / DSM 1251 Thiomicrospira denitrificans Strain ATCC 33889 / DSM 1251 | Q30SK1 | -- | 18065616 | Hydrolase activity, acting on carbon-nitrogen (but not peptide) bonds | Not Available |
| 674 | Biofilm PGA synthesis protein PgaD | *Pseudomonas trivialis* | | A0A0H5AAQ8 | -- | 26337878 | Not Available | Not Available |
| 675 | Outer membrane N-deacetylase | *Alcaligenes spp. EGD-AK7* | | U1ZPF8 | -- | 24407646 | Hydrolase activity, acting on carbon-nitrogen (but not peptide) bonds | Not Available |
| 676 | Uncharacterized protein | *Pediococcus damnosus* | | A0A0R2HGU2 | -- | 26415554 | Not Available | Not Available |
| 677 | Biofilm PGA synthesis lipoprotein PgaB | *Xanthomonas oryzae pv. oryzae* | Strain PXO99A | A0A0K0GHF3 | -- | 18452608 | Hydrolase activity, acting on carbon-nitrogen (but not peptide) bonds | Not Available |
| 678 | Poly-beta-1, 6-N-acetyl-D-glucosamine biosynthesis protein PgaD | *Pseudomonas spp. S04* | | A0A6I6SR34 | -- | 25398872 | Not Available | Not Available |
| 679 | Putative toxin-antitoxin system, toxin component | *Bordetella pertussis STO1-CHOC-0019* | | A0A6N3RY39 | -- | 24356839 | Not Available | Not Available |
| 680 | Poly-beta-1, 6-N-acetyl-D-glucosamine synthase (Poly-beta-1, 6-GlcNAc synthase) (EC 2.4.1.-) | *Pseudomonas spp. S04* | | A0A6I6SST5 | -- | 25398872 | Acetyl glucosaminyl transferase activity | Not Available |
| 681 | Surface composition regulator | *Yersinia pestis bv. Antiqua* | Strain Antiqua | A0A0E1NXP9 | -- | 16740952 | DNA binding, DNA-directed DNA polymerase activity | Not Available |
| 682 | Poly-beta-1, 6-N-acetyl-D-glucosamine synthase (Poly-beta-1, 6-GlcNAc synthase) (EC 2.4.1.-) | *Methylomona methanica* | Strain MC09 | G0A376 | -- | 22123758 | Acetyl glucosaminyl transferase activity | Not Available |
| 683 | Poly-beta-1, 6-N-acetyl-D-glucosamine N-deacetylase PgaB (EC 3.5.1.-) | *Bordetella pertussis STO1-CHOC-0019* | | A0A6N3RXJ4 | -- | 24356839 | Hydrolase activity, acting on carbon-nitrogen (but not peptide) bonds | Not Available |
| 684 | Uncharacterized protein | *Methylomona methanica* | Strain MC09 | G0A375 | -- | 22123758 | Not Available | Not Available |
| 685 | Poly-beta-1, 6-N-acetyl-D-glucosamine synthase (Poly-beta-1, 6-GlcNAc synthase) (EC 2.4.1.-) | *Lysobacter gummosus* | | A0A0S2G4Y6 | -- | 26597042 | Acetyl glucosaminyl transferase activity | Not Available |
| 686 | Haemin storage system, HmsF protein | *Yersinia pestis bv. Antiqua* | Strain Antiqua | A0A0E1NRK0 | -- | 16740952 | Hydrolase activity, acting on carbon-nitrogen (but not peptide) bonds | Not Available |
| 687 | GGDEF family protein | *Coxiella burnetii* | Strain RSA 493 / Nine Mile phaSe I | Q83A76 | -- | 12704232; 19047403 | Diguanylate cyclase activity | Not Available |
| 688 | Poly-beta-1, 6-N-acetyl-D-glucosamine N-deacetylase | *Pseudomonas spp. S04* | | A0A6I6SX07 | -- | 25398872 | Hydrolase activity, acting on carbon-nitrogen (but not peptide) bonds | Not Available |
| 689 | Poly-beta-1, 6-N-acetyl-D-glucosamine synthase (Poly-beta-1, 6-GlcNAc synthase) (EC 2.4.1.-) | *Anaeromyxobacter spp. PSR-1* | | A0A0D6QFS3 | -- | 23709511 | Acetyl glucosaminyl transferase activity | Not Available |
| 690 | Biofilm PGA synthesis lipoprotein PgaB | *Actinobacillus pleuropneumoniae serotype 3* | Strain JL03 | B0BTK3 | -- | 18197260 | Hydrolase activity, acting on carbon-nitrogen (but not peptide) bonds | Not Available |
| 691 | Poly-beta-1, 6-N-acetyl-D-glucosamine synthase (Poly-beta-1, 6-GlcNAc synthase) (EC 2.4.1.-) | *Pseudomonas kairouanensis* | | A0A4Z0ANC1 | -- | 30910423 | Acetyl glucosaminyl transferase activity | Not Available |

| Target ID | Bacterial Protein Name | Organism Name | Strain | UniProt ID | PDB ID | PubMed ID | Classification as per UniProt GO Molecular Function | Classification of Function as per PDB |
|---|---|---|---|---|---|---|---|---|
| 692 | Poly-beta-1, 6-N-acetyl-D-glucosamine N-deacetylase PgaB | *Pseudomonas kairouanensis* | | A0A4Z0API9 | -- | 30910423 | Hydrolase activity, acting on carbon-nitrogen (but not peptide) bonds | Not Available |
| 693 | Uncharacterized protein | *Macrococcus goetzii* | | A0A395G7Z9 | -- | 29951040 | Not Available | Not Available |
| 694 | Type II toxin-antitoxin system MqsR family toxin | *Pseudomonas kairouanensis* | | A0A4Z0AZZ6 | -- | 30910423 | Not Available | Not Available |
| 695 | Type II toxin-antitoxin system MqsR family toxin | *Thermoanaerobacter spp.* | | A0A7C7E8X0 | -- | 32123542 | Not Available | Not Available |
| 696 | Uncharacterized protein | *Rhodopila globiformis* | RhodoPseudomonas globiformis | A0A2S6NP98 | -- | 29423563 | Not Available | Not Available |
| 697 | Poly-beta-1, 6-N-acetyl-D-glucosamine biosynthesis protein PgaD | *Pseudomonas kairouanensis* | | A0A4Z0ANQ1 | -- | 30910423 | Not Available | Not Available |
| 698 | Toxin-antitoxin system, toxin component | *Burkholderiales bacterium RIFOXYC12_FULL_60_6* | | A0A1F4P0B8 | -- | 27774985 | Not Available | Not Available |
| 699 | GGDEF domain-containing protein | *Pseudomonas aeruginosa* | Strain ATCC 15692 / DSM 22644 / CIP 104116 / JCM 14847 / LMG 12228 / 1C / PRS 101 / PAO1 | Q9I2P4 | -- | 10984043 | Diguanylate cyclase activity | Not Available |
| 700 | Surface composition regulator | *Salmonella paratyphi C* | Strain RKS4594 | C0PYX0 | -- | 19229335 | Not Available | Not Available |
| 701 | Poly-beta-1, 6-N-acetyl-D-glucosamine synthase (Poly-beta-1, 6-GlcNAc synthase) (EC 2.4.1.-) | *Macrococcus goetzii* | | A0A395G880 | -- | 29951040 | Acetyl glucosaminyl transferase activity | Not Available |
| 702 | Poly-beta-1, 6-N-acetyl-D-glucosamine synthase (Poly-beta-1, 6-GlcNAc synthase) (EC 2.4.1.-) | *Exiguobacterium chiriqhucha RW-2* | | U1LZK0 | -- | 23929485 | Acetyl glucosaminyl transferase activity | Not Available |
| 703 | Surface composition regulator | *Enterobacter lignolyticus* | Strain SCF1 | E3GBM1 | -- | 22180812 | Not Available | Not Available |
| 704 | Uncharacterized protein | *Acinetobacter baumannii* | Strain AYE | B0V954 | -- | 18350144 | Not Available | Not Available |
| 705 | NodB homology domain-containing protein | *Acinetobacter baumannii* | Strain AYE | B0V7F4 | -- | 18350144 | Hydrolase activity, acting on carbon-nitrogen (but not peptide) bonds | Not Available |
| 706 | Poly-beta-1, 6-N-acetyl-D-glucosamine synthase (Poly-beta-1, 6-GlcNAc synthase) (EC 2.4.1.-) | *Halomonas spp. 54_146* | | A0A101CZX4 | -- | 26787827 | Acetyl glucosaminyl transferase activity | Not Available |
| 707 | NodB homology domain-containing protein | *Halomonas spp. 54_146* | | A0A101CZW4 | -- | 26787827 | Hydrolase activity, acting on carbon-nitrogen (but not peptide) bonds | Not Available |
| 708 | Poly-beta-1, 6-N-acetyl-D-glucosamine synthase (Poly-beta-1, 6-GlcNAc synthase) (EC 2.4.1.-) | *Acinetobacter baumannii* | Strain AYE | B0V7F5 | -- | 18350144 | Acetyl glucosaminyl transferase activity | Not Available |
| 709 | GGDEF domain-containing protein | *Pseudomonas aeruginosa* | Strain ATCC 15692 / DSM 22644 / CIP 104116 / JCM 14847 / LMG 12228 / 1C / PRS 101 / PAO1 | Q9HZX6 | -- | 10984043 | Diguanylate cyclase activity | Not Available |
| 710 | Uncharacterized protein | *Acinetobacter baumannii* | Strain AYE | B0V7F6 | -- | 18350144 | Not Available | Not Available |
| 711 | Probable two-component response regulator | *Pseudomonas aeruginosa* | Strain ATCC 15692 / DSM 22644 / CIP 104116 / JCM 14847 / LMG 12228 / 1C / PRS 101 / PAO1 | Q9HUW7 | -- | 10984043 | Diguanylate cyclase activity | Not Available |
| 712 | Putative lipoprotein (Haemin storage system) (HmsF) | *Acinetobacter baumannii* | Strain AYE | B0V956 | -- | 18350144 | Hydrolase activity, acting on carbon-nitrogen (but not peptide) bonds | Not Available |
| 713 | Poly-beta-1, 6-N-acetyl-D-glucosamine N-deacetylase PgaB | *Aeromona salmonicida* | | A0A428VR46 | -- | 30502493 | Hydrolase activity, acting on carbon-nitrogen (but not peptide) bonds | Not Available |
| 714 | Poly-beta-1, 6-N-acetyl-D-glucosamine synthase (Poly-beta-1, 6-GlcNAc synthase) (EC 2.4.1.-) | *Acinetobacter baumannii* | Strain AYE | B0V955 | -- | 18350144 | Acetyl glucosaminyl transferase activity | Not Available |
| 715 | Uncharacterized protein | *Caldanaerobacter subterraneus subspp. yonseiensis KB-1* | | U5CHV1 | -- | 24201201 | Not Available | Not Available |
| 716 | Poly-beta-1, 6-N-acetyl-D-glucosamine synthase (Poly-beta-1, 6-GlcNAc synthase) (EC 2.4.1.-) | *Acinetobacter ursingii* | | A0A3D2SJG2 | -- | 30148503 | Acetyl glucosaminyl transferase activity | Not Available |
| 717 | Poly-beta-1, 6-N-acetyl-D-glucosamine N-deacetylase PgaB | *Enterobacter hormaechei* | | A0A6G3F3N1 | -- | 31477907 | Hydrolase activity, acting on carbon-nitrogen (but not peptide) bonds | Not Available |
| 718 | Biofilm PGA synthesis auxiliary protein PgaD | *Acinetobacter ursingii* | | A0A3G9G0U0 | -- | 28293292 | Not Available | Not Available |
| 719 | Poly-beta-1, 6-N-acetyl-D-glucosamine N-deacetylase PgaB (Fragment) | *Acinetobacter ursingii* | | A0A3D2SPM8 | -- | 30148503 | Hydrolase activity, acting on carbon-nitrogen (but not peptide) bonds | Not Available |
| 720 | Surface composition regulator | *Enterobacter hormaechei* | | A0A2G4ZVG2 | -- | 29088362 | Not Available | Not Available |
| 721 | Poly-beta-1, 6-N-acetyl-D-glucosamine synthase (Poly-beta-1, 6-GlcNAc synthase) (EC 2.4.1.-) | *Enterobacter hormaechei* | | A0A2J0PY96 | -- | 29088362 | Acetyl glucosaminyl transferase activity | Not Available |
| 722 | Surface composition regulator | *Enterobacter hormaechei* | | A0A6G3F999 | -- | 31477907 | Not Available | Not Available |
| 723 | Poly-beta-1, 6-N-acetyl-D-glucosamine synthase (Poly-beta-1, 6-GlcNAc synthase) (EC 2.4.1.-) | *Achromobacter marplatensis* | | J4QR07 | -- | 23105084 | Acetyl glucosaminyl transferase activity | Not Available |
| 724 | Uncharacterized protein | *Achromobacter marplatensis* | | J4J7C4 | -- | 23105084 | Not Available | Not Available |
| 725 | Poly-beta-1, 6-N-acetyl-D-glucosamine N-deacetylase PgaB | *Methylomona denitrificans* | | A0A140E5K7 | -- | 25580993 | Hydrolase activity, acting on carbon-nitrogen (but not peptide) bonds | Not Available |
| 726 | Poly-beta-1, 6-N-acetyl-D-glucosamine biosynthesis protein PgaD | *Enterobacter hormaechei* | | A0A6G3F3X4 | -- | 31477907 | Not Available | Not Available |
| 727 | Uncharacterized protein | *Achromobacter marplatensis* | | J4J6D3 | -- | 23105084 | Not Available | Not Available |
| 728 | Hemin storage protein | *Achromobacter marplatensis* | | J4YMC5 | -- | 23105084 | Hydrolase activity, acting on carbon-nitrogen (but not peptide) bonds | Not Available |
| 729 | Uncharacterized protein | *Methylomona denitrificans* | | A0A140E5K9 | -- | 25580993 | Not Available | Not Available |
| 730 | Poly-beta-1, 6-N-acetyl-D-glucosamine synthase (Poly-beta-1, 6-GlcNAc synthase) (EC 2.4.1.-) | *Achromobacter marplatensis* | | J4Y513 | -- | 23105084 | Acetyl glucosaminyl transferase activity | Not Available |
| 731 | Biofilm PGA synthesis auxiliary protein PgaD | *Acinetobacter ursingii* | | A0A3G9FU89 | -- | 28293292 | Not Available | Not Available |
| 732 | Poly-beta-1, 6-N-acetyl-D-glucosamine synthase (Poly-beta-1, 6-GlcNAc synthase) (EC 2.4.1.-) | *Methylomona denitrificans* | | A0A140E5K8 | -- | 25580993 | Acetyl glucosaminyl transferase activity | Not Available |
| 733 | Poly-beta-1, 6-N-acetyl-D-glucosamine synthase (Poly-beta-1, 6-GlcNAc synthase) (EC 2.4.1.-) | *Acinetobacter ursingii* | | A0A3G9FTN3 | -- | 28293292 | Acetyl glucosaminyl transferase activity | Not Available |
| 734 | Poly-beta-1, 6-N-acetyl-D-glucosamine N-deacetylase PgaB | *Enterobacter hormaechei* | | A0A2J0PYU0 | -- | 29088362 | Hydrolase activity, acting on carbon-nitrogen (but not peptide) bonds | Not Available |
| 735 | Polysaccharide deacetylase | *Escherichia coli O78:H11* | Strain H10407 / ETEC | E3PJJ0 | -- | 20802035 | Hydrolase activity, acting on carbon-nitrogen (but not peptide) bonds | Not Available |
| 736 | Surface composition regulator | *Escherichia coli O78:H11* | Strain H10407 / ETEC | E3PJW4 | -- | 20802035 | Not Available | Not Available |
| 737 | Poly-beta-1, 6-N-acetyl-D-glucosamine synthase (Poly-beta-1, 6-GlcNAc synthase) (EC 2.4.1.-) | *Escherichia coli O78:H11* | Strain H10407 / ETEC | E3PJI9 | -- | 20802035 | Acetyl glucosaminyl transferase activity | Not Available |
| 738 | Poly-beta-1, 6-N-acetyl-D-glucosamine synthase (Poly-beta-1, 6-GlcNAc synthase) (EC 2.4.1.-) | *Carnobacterium divergens DSM 20623* | | A0A0R2HW17 | -- | 26415554 | Acetyl glucosaminyl transferase activity | Not Available |
| 739 | Uncharacterized protein | *Staphylococcus aureus subspp. aureus 71193* | | A0A0E0VNH0 | -- | 22375071 | Not Available | Not Available |

| Target ID | Bacterial Protein Name | Organism Name | Strain | UniProt ID | PDB ID | PubMed ID | Classification as per UniProt GO Molecular Function | Classification of Function as per PDB |
|---|---|---|---|---|---|---|---|---|
| 740 | Poly-beta-1, 6-N-acetyl-D-glucosamine synthase (Poly-beta-1, 6-GlcNAc synthase) (EC 2.4.1.-) | *Candidatus Muproteobacteria bacterium RBG_19FT_COMBO_61_10* | | A0A1F6UP22 | -- | 27774985 | Acetyl glucosaminyl transferase activity | Not Available |
| 741 | Poly-beta-1, 6-N-acetyl-D-glucosamine synthase (Poly-beta-1, 6-GlcNAc synthase) (EC 2.4.1.-) | *Acidobacteria bacterium* | | A0A2V7YF32 | -- | 29899444 | Acetyl glucosaminyl transferase activity | Not Available |
| 742 | Poly-beta-1, 6-N-acetyl-D-glucosamine synthase (Poly-beta-1, 6-GlcNAc synthase) (EC 2.4.1.-) | *Aggregatibacter aphrophilus* | Haemophilus aphrophilus | A0A3M6PFA6 | -- | 30598730 | Acetyl glucosaminyl transferase activity | Not Available |
| 743 | Poly-beta-1, 6-N-acetyl-D-glucosamine N-deacetylase PgaB | *Gammaproteobacteria bacterium HGW-Gammaproteobacteria-10* | | A0A2N1ZS52 | -- | 28350393 | Hydrolase activity, acting on carbon-nitrogen (but not peptide) bonds | Not Available |
| 744 | Poly-beta-1, 6-N-acetyl-D-glucosamine synthase (Poly-beta-1, 6-GlcNAc synthase) (EC 2.4.1.-) | *Alicyclobacillus acidocaldarius* | Strain Tc-4-1 Bacills acidocaldarius | F8IJL3 | -- | 21914900 | Acetyl glucosaminyl transferase activity | Not Available |
| 745 | Poly-beta-1, 6-N-acetyl-D-glucosamine synthase (Poly-beta-1, 6-GlcNAc synthase) (EC 2.4.1.-) | *Lautropia spp. SCN 70-15* | | A0A1D2TKP6 | -- | 26031303 | Acetyl glucosaminyl transferase activity | Not Available |
| 746 | mRNA interferase MqsR | *Janthinobacterium spp.* | | A0A349L6R0 | -- | 30148503 | Not Available | Not Available |
| 747 | Poly-beta-1, 6-N-acetyl-D-glucosamine synthase (Poly-beta-1, 6-GlcNAc synthase) (EC 2.4.1.-) | *Gammaproteobacteria bacterium HGW-Gammaproteobacteria-10* | | A0A2N1ZS36 | -- | 28350393 | Acetyl glucosaminyl transferase activity | Not Available |
| 748 | Uncharacterized protein | *Alicyclobacillus acidocaldarius* | Strain Tc-4-1 Bacills acidocaldarius | F8IJL4 | -- | 21914900 | Not Available | Not Available |
| 749 | Poly-beta-1, 6-N-acetyl-D-glucosamine biosynthesis protein PgaD | *Gammaproteobacteria bacterium HGW-Gammaproteobacteria-10* | | A0A2N1ZS44 | -- | 28350393 | Not Available | Not Available |
| 750 | Biofilm PGA synthesis protein PgaD | *Escherichia coli O139:H28* | Strain E24377A / ETEC | A7ZKC6 | -- | 18676672 | Not Available | Not Available |
| 751 | Uncharacterized protein | *Pseudomonas putida H8234* | | R9VGH1 | -- | 23868128 | Not Available | Not Available |
| 752 | Motility quorum-sensing regulator MqsR | *Deltaproteobacteria bacterium RIFOXYD12_FULL_56_24* | | A0A1F9L5Q3 | -- | 27774985 | Not Available | Not Available |
| 753 | Poly-beta-1, 6-N-acetyl-D-glucosamine N-deacetylase PgaB | *Lautropia spp. SCN 70-15* | | A0A1D2TKR0 | -- | 26031303 | Hydrolase activity, acting on carbon-nitrogen (but not peptide) bonds | Not Available |
| 754 | Poly-beta-1, 6-N-acetyl-D-glucosamine biosynthesis protein PgaD | *Lautropia spp. SCN 70-15* | | A0A1D2TKI8 | -- | 26031303 | Not Available | Not Available |
| 755 | Poly-beta-1, 6-N-acetyl-D-glucosamine N-deacetylase PgaB | *Aggregatibacter aphrophilus* | Haemophilus aphrophilus | A0A3M6PFB5 | -- | 30598730 | Hydrolase activity, acting on carbon-nitrogen (but not peptide) bonds | Not Available |
| 756 | Uncharacterized protein | *Deltaproteobacteria bacterium RIFOXYD12_FULL_56_24* | | A0A1F9L1K1 | -- | 27774985 | Not Available | Not Available |
| 757 | Toxin-antitoxin system, toxin component | *Hafnia alvei FB1* | | A0A097R795 | -- | 25075225 | Not Available | Not Available |
| 758 | Poly-beta-1, 6-N-acetyl-D-glucosamine synthase (Poly-beta-1, 6-GlcNAc synthase) (EC 2.4.1.-) | *Staphylococcus aureus subspp. aureus CN1* | | T1YDR1 | -- | 23977354 | Acetyl glucosaminyl transferase activity | Not Available |
| 759 | Polysaccharide deacetylase | *Xenorhabdus ehlersii* | | A0A2D0ILC7 | -- | 28993611 | Hydrolase activity, acting on carbon-nitrogen (but not peptide) bonds | Not Available |
| 760 | Poly-beta-1, 6-N-acetyl-D-glucosamine synthase (Poly-beta-1, 6-GlcNAc synthase) (EC 2.4.1.-) | *Escherichia coli O139:H28* | Strain E24377A / ETEC | A7ZKC7 | -- | 18676672 | Acetyl glucosaminyl transferase activity | Not Available |
| 761 | Biofilm PGA synthesis lipoprotein pgaB (EC 3.-.-.-) | *Escherichia coli O139:H28* | Strain E24377A / ETEC | A7ZKC8 | -- | 18676672 | Hydrolase activity, acting on carbon-nitrogen (but not peptide) bonds | Not Available |
| 762 | Hemin storage protein | *Hafnia alvei FB1* | | A0A097R0G2 | -- | 25075225 | Hydrolase activity, acting on carbon-nitrogen (but not peptide) bonds | Not Available |
| 763 | Poly-beta-1, 6-N-acetyl-D-glucosamine synthase (Poly-beta-1, 6-GlcNAc synthase) (EC 2.4.1.-) | *Xenorhabdus ehlersii* | | A0A2D0ILF0 | -- | 28993611 | Acetyl glucosaminyl transferase activity | Not Available |
| 764 | Putative intercellular adhesion protein A (Biofilm formation) | *Vibrio parahaemolyticus serotype O3:K6* | Strain RIMD 2210633 | Q87PP2 | -- | 12620739 | Not Available | Not Available |
| 765 | Surface composition regulator | *Hafnia alvei FB1* | | A0A097R276 | -- | 25075225 | Not Available | Not Available |
| 766 | Uncharacterized protein | *Legionella drancourtii LLAP12* | | G9ERV4 | -- | 22047552 | Not Available | Not Available |
| 767 | Poly-beta-1, 6-N-acetyl-D-glucosamine biosynthesis protein PgaD | *Xenorhabdus ehlersii* | | A0A2D0ILC2 | -- | 28993611 | Not Available | Not Available |
| 768 | Poly-beta-1, 6-N-acetyl-D-glucosamine synthase (Poly-beta-1, 6-GlcNAc synthase) (EC 2.4.1.-) | *Hafnia alvei FB1* | | A0A097R0I0 | -- | 25075225 | Acetyl glucosaminyl transferase activity | Not Available |
| 769 | Surface composition regulator | *Edwardsiella tarda* | Strain FL6-60 | A0A0H3DQQ5 | -- | 22003892 | Not Available | Not Available |
| 770 | Surface composition regulator | *Salmonella paratyphi A* | Strain ATCC 9150 / SARB42 | A0A735JW74 | -- | 30286803 | Not Available | Not Available |
| 771 | Poly-beta-1, 6-N-acetyl-D-glucosamine synthase (Poly-beta-1, 6-GlcNAc synthase) (EC 2.4.1.-) | *Yersinia enterocolitica* | type O:2 Str. YE3094 / 96 | A0A7U7IYU5 | -- | 24753568 | Acetyl glucosaminyl transferase activity | Not Available |
| 772 | Surface composition regulator | *Yersinia enterocolitica* | type O:2 Str. YE3094 / 96 | A0A7U7FZ92 | -- | 24753568 | DNA binding, DNA-directed DNA polymerase activity | Not Available |
| 773 | Uncharacterized protein | *Xanthomonas hortorum pv. carotae* | Str. M081 | V7ZEF0 | -- | 21722296 | Not Available | Not Available |
| 774 | Surface composition regulator | *Salmonella enterica subspp. enterica serovar* | 6, 7:c:1, 5 | A0A740VH22 | -- | 30286803 | Not Available | Not Available |
| 775 | Surface composition regulator | *Yersinia enterocolitica* | type O:5, 27 Str. YE149 / 02 | A0A7U7IQG9 | -- | 24753568 | DNA binding, DNA-directed DNA polymerase activity | Not Available |
| 776 | Haemin storage system, HmsS protein | *Yersinia enterocolitica* | type O:5, 27 Str. YE149 / 02 | A0A7U7FMU7 | -- | 24753568 | Not Available | Not Available |
| 777 | IcaA (Fragment) | *Staphylococcus condimenti* | | Q83ZI0 | -- | 12957956 | Not Available | Not Available |
| 778 | Uncharacterized protein | *Yersinia enterocolitica* | type O:2 Str. YE3094 / 96 | A0A7U7G2K0 | -- | 24753568 | Not Available | Not Available |
| 779 | Surface composition regulator | *Salmonella enterica subspp. houtenae serovar* | 44:z4, z24:- | A0A737NPA5 | -- | 30286803 | Not Available | Not Available |
| 780 | Surface composition regulator | *Salmonella enterica subspp. enterica serovar* | Miami | A0A5I2DLX0 | -- | 30286803 | Not Available | Not Available |
| 781 | Haemin storage system, HmsS protein | *Yersinia enterocolitica* | type O:2 Str. YE3094 / 96 | A0A7U7IYG1 | -- | 24753568 | Not Available | Not Available |
| 782 | Surface composition regulator | *Salmonella enterica subspp. houtenae serovar 48:z4, z32:-* | | A0A729FR63 | -- | 30286803 | Not Available | Not Available |
| 783 | Surface composition regulator | *Salmonella enterica subspp. enterica serovar Typhi* | Str. CT18 | A0A718C8V5 | -- | 30286803 | Not Available | Not Available |
| 784 | Surface composition regulator | *Salmonella enterica subspp. arizonae serovar* | 41:z4, z23:- | A0A741N0E9 | -- | 30286803 | Not Available | Not Available |
| 785 | S-ribosylhomocysteine lyase (EC 4.4.1.21) (AI-2 synthesis protein) (Autoinducer-2 production protein LuxS) | *Aeromona hydrophila* | | B1PWE7 | -- | 18930130 | Iron ion binding, S-ribosylhomocysteine lyase activity | Not Available |
| 786 | PsoR | *Pseudomonas putida* | Arthrobacter siderocapsulatus | A8MN34 | -- | 16585751; 18599835 | DNA binding | Not Available |
| 787 | IcaA (Fragment) | *Staphylococcus condimenti* | | Q83ZH5 | -- | 12957956 | Not Available | Not Available |
| 788 | Biofilm PGA synthesis lipoprotein pgaB | *Yersinia enterocolitica W22703* | | F4N0N6 | -- | 21453472 | Hydrolase activity, acting on carbon-nitrogen (but not peptide) bonds | Not Available |
| 789 | Uncharacterized protein (Fragment) | *Klebsiella pneumoniae* | | Q1G0U9 | -- | 16861646 | Not Available | Not Available |

| Target ID | Bacterial Protein Name | Organism Name | Strain | UniProt ID | PDB ID | PubMed ID | Classification as per UniProt GO Molecular Function | Classification of Function as per PDB |
|---|---|---|---|---|---|---|---|---|
| 790 | Uncharacterized protein | *Klebsiella pneumoniae* | | Q1G0V5 | -- | 16861646 | Not Available | Not Available |
| 791 | Surface composition regulator | *Salmonella enterica subspp. enterica serovar Kouka* | | A0A729KY13 | -- | 30286803 | Not Available | Not Available |
| 792 | LysR_substrate domain-containing protein (Fragment) | *Klebsiella pneumoniae* | | Q1G0W0 | -- | 16861646 | Not Available | Not Available |
| 793 | Uncharacterized protein | *Klebsiella pneumoniae* | | Q1G0V8 | -- | 16861646 | Not Available | Not Available |
| 794 | Surface composition regulator | *Salmonella enterica subspp. houtenae serovar* | 1, 40:z4, z32:- | A0A730WRJ9 | -- | 30286803 | Not Available | Not Available |
| 795 | Uncharacterized protein | *Klebsiella pneumoniae* | | Q1G0U0 | -- | 16861646 | Not Available | Not Available |
| 796 | Aldedh domain-containing protein | *Klebsiella pneumoniae* | | Q1G0V4 | -- | 16861646 | Oxidoreductase activity, acting on the aldehyde or oxo group of donors, NAD or NADP as acceptor | Not Available |
| 797 | Uncharacterized protein (Fragment) | *Klebsiella pneumoniae* | | Q1G0T9 | -- | 16861646 | Not Available | Not Available |
| 798 | Uncharacterized protein (Fragment) | *Klebsiella pneumoniae* | | Q1G0U6 | -- | 16861646 | Not Available | Not Available |
| 799 | Uncharacterized protein | *uncultured Desulfobacterium spp.* | | E1YIC6 | -- | 21176053 | Not Available | Not Available |
| 800 | Poly-beta-1, 6-N-acetyl-D-glucosamine synthase (Poly-beta-1, 6-GlcNAc synthase) (EC 2.4.1.-) | *Acinetobacter baumannii* | | C8YYH7 | -- | 19633088 | Acetyl glucosaminyl transferase activity | Not Available |
| 801 | Autoinducer-2 ABC transporter permease protein | *Klebsiella pneumoniae* | | C8BNA6 | -- | 20600864 | Transmembrane transporter activity | Not Available |
| 802 | Uncharacterized protein | *Klebsiella pneumoniae* | | Q1G0U5 | -- | 16861646 | Not Available | Not Available |
| 803 | Urease accessory protein UreF | *Staphylococcus spp. T-02* | | B9A8F9 | -- | 20208338 | Nickel cation binding | Not Available |
| 804 | Surface composition regulator | *Yersinia enterocolitica W22703* | | F4N5Y4 | -- | 21453472 | DNA binding, DNA-directed DNA polymerase activity | Not Available |
| 805 | Poly-beta-1, 6-N-acetyl-D-glucosamine synthase (Poly-beta-1, 6-GlcNAc synthase) (EC 2.4.1.-) (Fragment) | *Yersinia enterocolitica W22703* | | F4MUX8 | -- | 21453472 | Acetyl glucosaminyl transferase activity | Not Available |
| 806 | Poly-beta-1, 6-N-acetyl-D-glucosamine synthase (Poly-beta-1, 6-GlcNAc synthase) (EC 2.4.1.-) | *Staphylococcus capitis* | | Q6YI89 | -- | 12957956 | Acetyl glucosaminyl transferase activity | Not Available |
| 807 | Surface composition regulator | *Salmonella typhimurium* | | A0A701JYH4 | -- | 30286803 | Not Available | Not Available |
| 808 | Surface composition regulator | *Salmonella enterica subspp. enterica serovar* | Orion | A0A3T2WB67 | -- | 30286803 | Not Available | Not Available |
| 809 | Surface composition regulator | *Salmonella gallinarum* | | A0A3V9NPH3 | -- | 30286803 | Not Available | Not Available |
| 810 | IcaR (Fragment) | *Staphylococcus capitis* | | Q6YI86 | -- | 12957956 | DNA binding | Not Available |
| 811 | Uncharacterized protein | *Klebsiella pneumoniae* | | Q1G0U2 | -- | 16861646 | Not Available | Not Available |
| 812 | IcaA (Fragment) | *Staphylococcus capitis* | | Q83ZH3 | -- | 12957956 | Not Available | Not Available |
| 813 | PTS EIIC type-3 domain-containing protein (Fragment) | *Klebsiella pneumoniae* | | Q1G0U4 | -- | 16861646 | Protein-N(PI)-phosphohistidine-sugar phosphoTransferase activity | Not Available |
| 814 | Biofilm polysaccharide intercellular adhesin deacetylase (Intercellular adhesion protein B) (Poly-beta-1, 6-N-acetyl-D-glucosamine N-deacetylase) (Fragment) | *Staphylococcus capitis* | | Q6YI87 | -- | 12957956 | Hydrolase activity, acting on carbon-nitrogen (but not peptide) bonds, in linear amides | Not Available |
| 815 | Surface composition regulator | *Salmonella enterica subspp. enterica serovar* | Ohio | A0A3V5VUX0 | -- | 30286803 | Not Available | Not Available |
| 816 | Surface composition regulator | *Salmonella enterica subspp. houtenae serovar* | 43:z4, z23:- | A0A730K0K8 | -- | 30286803 | Not Available | Not Available |
| 817 | Uncharacterized protein | *Yersinia enterocolitica W22703* | | F4MUX9 | -- | 21453472 | Not Available | Not Available |
| 818 | Surface composition regulator | *Salmonella enterica subspp. enterica serovar Paratyphi B* | | A0A3V7VTQ9 | -- | 30286803 | Not Available | Not Available |
| 819 | Surface composition regulator | *Salmonella enterica subspp. enterica serovar* | Rubislaw Str. ATCC 10717 | A0A6W0NR32 | -- | 30286803 | Not Available | Not Available |
| 820 | Biofilm polysaccharide intercellular adhesin synthesis protein IcaD (Intercellular adhesion protein D) (Poly-beta-1, 6-N-acetyl-D-glucosamine synthesis protein IcaD) (Fragment) | *Staphylococcus capitis* | | Q6YI88 | -- | 12957956 | Not Available | Not Available |
| 821 | S-adenosylmethionine decarboxylase proenzyme (AdoMetDC) (SAMDC) (EC 4.1.1.50) [Cleaved into: S-adenosylmethionine decarboxylase beta chain; S-adenosylmethionine decarboxylase alpha chain] | *Bacillus cereus* | | A0A063CLJ3 | -- | 25301660; 30637927 | Adenosylmethionine decarboxylase activity | Not Available |
| 822 | Ribonuclease J (RNAse J) (EC 3.1.-.-) | *Bacillus cereus* | | A0A068NAJ3 | -- | 25301660 | 5'-3' exoribonuclease activity, endoribonuclease activity, RNA binding, zinc ion binding | Not Available |
| 823 | Poly-beta-1, 6-N-acetyl-D-glucosamine synthase (Poly-beta-1, 6-GlcNAc synthase) (EC 2.4.1.-) | *Staphylococcus aureus* | | A0A2S6DQT8 | -- | 29479347 | Acetyl glucosaminyl transferase activity | Not Available |
| 824 | Global transcriptional regulator Spx | *Bacillus cereus* | | A0A2A8ED49 | -- | 25301660; 31228823; 30637927; 33353020 | Not Available | Not Available |
| 825 | Surface composition regulator | *Salmonella hadar* | | A0A3V2W5E6 | -- | 30286803; 30902850 | Not Available | Not Available |
| 826 | Ribonuclease PH (RNAse PH) (EC 2.7.7.56) (tRNA nucleotidyl transferase) | *Bacillus cereus* | | A0A080TUN6 | -- | 31228823; 30637927 | 3'-5'-exoribonuclease activity, tRNA binding, tRNA nucleotidylTransferase activity | Not Available |
| 827 | Ribonuclease J (RNAse J) (EC 3.1.-.-) | *Bacillus cereus* | | A0A063CPQ3 | -- | 25301660 | 5'-3' exoribonuclease activity, endoribonuclease activity, RNA binding, zinc ion binding | Not Available |
| 828 | Surface composition regulator | *Salmonella enterica subspp. enterica serovar* | Braenderup | A0A3T3G645 | -- | 30286803 | Not Available | Not Available |
| 829 | Type II toxin-antitoxin system MqsR family toxin | *Yersinia pestis* | | A0A3N4BIT9 | -- | 11586360 | Not Available | Not Available |
| 830 | Surface composition regulator | *Salmonella enterica subspp. enterica serovar* | Javiana | A0A3V7PI12 | -- | 30286803 | Not Available | Not Available |
| 831 | Surface composition regulator | *Salmonella anatum* | | A0A2T8MEV5 | -- | 30286803; 30902850 | Not Available | Not Available |
| 832 | Surface composition regulator | *Salmonella potsdam* | | A0A5I6RXB1 | -- | 30286803 | Not Available | Not Available |
| 833 | Surface composition regulator | *Salmonella enterica subspp. enterica serovar* | Coeln | A0A5I3IDD6 | -- | 30286803 | Not Available | Not Available |
| 834 | Surface composition regulator | *Salmonella enterica subspp. enterica serovar* | Napoli | A0A5H6JAD5 | -- | 30286803 | Not Available | Not Available |
| 835 | Poly-beta-1, 6-N-acetyl-D-glucosamine synthase (Poly-beta-1, 6-GlcNAc synthase) (EC 2.4.1.-) | *Staphylococcus lugdunensis* | | A8DMY8 | -- | 17635864 | Acetyl glucosaminyl transferase activity | Not Available |
| 836 | Surface composition regulator | *Salmonella enterica subspp. enterica serovar* | | A0A720D3V5 | -- | 30286803 | Not Available | Not Available |
| 837 | AHL-dependent Transcriptional regulator | *Edwardsiella tarda* | | Q60HM0 | -- | 16233705; 19283379 | DNA binding | Not Available |
| 838 | Surface composition regulator | *Salmonella enteritidis PT4* | Strain P125109 | A0A724LRF7 | -- | 30286803 | Not Available | Not Available |
| 839 | Surface composition regulator | *Salmonella enterica subspp. enterica serovar* | Havana | A0A3V3TZ71 | -- | 30286803 | Not Available | Not Available |

| Target ID | Bacterial Protein Name | Organism Name | Strain | UniProt ID | PDB ID | PubMed ID | Classification as per UniProt GO Molecular Function | Classification of Function as per PDB |
|---|---|---|---|---|---|---|---|---|
| 840 | Surface composition regulator | *Salmonella enterica subspp. enterica serovar* | Eastbourne | A0A3V5UUL8 | -- | 30286803 | Not Available | Not Available |
| 841 | Surface composition regulator | *Salmonella enterica subspp. enterica serovar* | Haifa | A0A3V5ZEC4 | -- | 30902850 | Not Available | Not Available |
| 842 | Surface composition regulator | *Edwardsiella piscicida* | | A0A034SUT9 | -- | 24926054 | Not Available | Not Available |
| 843 | Adenosylhomocysteinae (EC 3.3.1.1) (S-adenosyl-L-homocysteine Hydrolase) (AdoHcyase) | *Pseudomonas aeruginosa* | | K7QUT8 | -- | 22942290 | Adenosylhomocysteinae activity | Not Available |
| 844 | Poly-beta-1, 6-N-acetyl-D-glucosamine N-deacetylase (Poly-beta-1, 6-N-acetyl-D-glucosamine N-deacetylase PgaB) (EC 3.5.1.-) | *Pseudomonas lactis* | | A0A219A6N9 | -- | 32587583 | Hydrolase activity, acting on carbon-nitrogen (but not peptide) bonds | Not Available |
| 845 | Surface composition regulator | *Salmonella enterica subspp. enterica serovar* | Bareilly | A0A4Z8Z843 | -- | 30286803; 30902850 | Not Available | Not Available |
| 846 | Poly-beta-1, 6-N-acetyl-D-glucosamine N-deacetylase | *Pseudomonas prosekii* | | A0A3L8CDD7 | -- | 29867824 | Hydrolase activity, acting on carbon-nitrogen (but not peptide) bonds | Not Available |
| 847 | Biofilm PGA synthesis lipoprotein PgaB (EC 3.-.-.-) (Poly-beta-1, 6-N-acetyl-D-glucosamine N-deacetylase PgaB) (EC 3.5.1.-) | *Pseudomonas lactis* | | I4KF65 | -- | 22792073; 32587583 | Hydrolase activity, acting on carbon-nitrogen (but not peptide) bonds | Not Available |
| 848 | DNA polymerase III subunit delta (EC 2.7.7.7) (HolA) | *Staphylococcus lugdunensis* | | A0A0N7BA97 | -- | 26416910 | DNA binding, DNA-directed DNA polymerase activity | Not Available |
| 849 | 3-hydroxylisobutyryl-CoA Hydrolase (Putative enoyl-CoA hydratase / isomerase family protein) | *Acinetobacter oleivorans* | Strain JCM 16667 / KCTC 23045 / DR1 | D3JTG3 | -- | 20629796; 20639327 | 3-hydroxyisobutyryl-CoA Hydrolase activity, isomerase activity | Not Available |
| 850 | Surface composition regulator | *Salmonella enterica subspp. enterica serovar* | Concord | A0A2R4D758 | -- | 30286803 | Not Available | Not Available |
| 851 | Poly-beta-1, 6-N-acetyl-D-glucosamine biosynthesis protein PgaD | *Pseudomonas prosekii* | | A0A3L8CDF8 | -- | 29867824 | Not Available | Not Available |
| 852 | Poly-beta-1, 6-N-acetyl-D-glucosamine biosynthesis protein PgaD | *Acinetobacter schindleri* | | A0A2S1FF05 | -- | 29954893 | Not Available | Not Available |
| 853 | Poly-beta-1, 6-N-acetyl-D-glucosamine biosynthesis protein PgaD (Putative biofilm PGA synthesis protein PgaD) | *Pseudomonas lactis* | | A0A0R3BJY0 | -- | 22792073; 32587583 | Not Available | Not Available |
| 854 | Poly-beta-1, 6-N-acetyl-D-glucosamine synthase (Poly-beta-1, 6-GlcNAc synthase) (EC 2.4.1.-) | *Acinetobacter cumulans* | | A0A3A8G879 | -- | 30808586 | Acetyl glucosaminyl transferase activity | Not Available |
| 855 | Poly-beta-1, 6-N-acetyl-D-glucosamine synthase (Poly-beta-1, 6-GlcNAc synthase) (EC 2.4.1.-) | *Pseudomonas prosekii* | | A0A3L8CCK9 | -- | 29867824 | Acetyl glucosaminyl transferase activity | Not Available |
| 856 | ComE operon protein 2 (ComEB) | *Staphylococcus lugdunensis* | | A0A0N7B1F5 | -- | 26416910 | dCMP deaminae activity, zinc ion binding | Not Available |
| 857 | Poly-beta-1, 6-N-acetyl-D-glucosamine synthase (Poly-beta-1, 6-GlcNAc synthase) (EC 2.4.1.-) | *Pseudomonas lactis* | | I4KC34 | -- | 22792073; 32587583 | Acetyl glucosaminyl transferase activity | Not Available |
| 858 | Surface composition regulator | *Klebsiella oxytoca* | | A0A0G3S4R7 | -- | 29437920 | Not Available | Not Available |
| 859 | Motility quorum-sensing regulator MqsR (Type II toxin-antitoxin system MqsR family toxin) | *Pseudomonas lactis* | | I4KAK6 | -- | 22792073; 32587583 | Not Available | Not Available |
| 860 | Poly-beta-1, 6-N-acetyl-D-glucosamine synthase (Poly-beta-1, 6-GlcNAc synthase) (EC 2.4.1.-) | *Yersinia pestis* | | A0A1U8QLL4 | -- | 12142430; 15368893 | Acetyl glucosaminyl transferase activity | Not Available |
| 861 | HTH-type Transcriptional regulator LuxR (LuxR regulator) | *Vibrio scophthalmi* | | H6URH3 | -- | 23198796 | DNA binding | Not Available |
| 862 | Acyl-homoserine-lactone synthase (EC 2.3.1.184) (Autoinducer synthesis protein) | *Aeromona hydrophila* | | Q27IV3 | -- | 19729404 | N-acyl homoserine lactone synthase activity | Not Available |
| 863 | Surface composition regulator | *Yersinia pestis* | | Q0WDU2 | -- | 11586360 | DNA binding, DNA-directed DNA polymerase activity | Not Available |
| 864 | Fimbrial protein (Similar to pilin F17d-A) (Type 1 fimbrial protein) | *Yersinia pestis* | | Q9ZC32 | -- | 11586360 | Not Available | Not Available |
| 865 | Poly-beta-1, 6-N-acetyl-D-glucosamine N-deacetylase PgaB | *Acinetobacter wuhouensis* | | A0A385C6B4 | -- | 30117799 | Hydrolase activity, acting on carbon-nitrogen (but not peptide) bonds | Not Available |
| 866 | AhyR (LuxR family Transcriptional regulator) (LuxR-type Transcriptional regulator) | *Aeromona hydrophila* | | Q27IV4 | -- | 19729404 | DNA binding | Not Available |
| 867 | Surface composition regulator | *Salmonella enterica subspp. enterica serovar* | Lattenkamp | A0A5W2LQM8 | -- | 30286803 | Not Available | Not Available |
| 868 | HmsT (HmsT protein) | *Yersinia pestis* | | Q9X5X6 | -- | 9249062; 10438760; 10463178; 11586360 | Diguanylate cyclase activity | Not Available |
| 869 | Haemin storage system, HmsF protein (HmsF protein) (Poly-beta-1, 6-N-acetyl-D-glucosamine N-deacetylase PgaB) (EC 3.5.1.-) | *Yersinia pestis* | | Q9R7V4 | -- | 11586360 | Hydrolase activity, acting on carbon-nitrogen (but not peptide) bonds | Not Available |
| 870 | Surface composition regulator | *Salmonella enterica subspp. enterica serovar* | Mbandaka | A0A418ZAK8 | -- | 30286803; 31099590 | Not Available | Not Available |
| 871 | Frimbrial protein (Putative frimbrial protein) (Type 1 fimbrial protein) | *Yersinia pestis* | | Q306M1 | -- | 11586360 | Not Available | Not Available |
| 872 | Haemin storage system, HmsS protein (HmsS) (HmsS protein) (Poly-beta-1, 6-N-acetyl-D-glucosamine biosynthesis protein PgaD) | *Yersinia pestis* | | P74989 | -- | 9249062; 11586360 | Not Available | Not Available |
| 873 | Poly-beta-1, 6-N-acetyl-D-glucosamine biosynthesis protein PgaD | *Acinetobacter wuhouensis* | | A0A385C8R0 | -- | 30117799 | Not Available | Not Available |
| 874 | Poly-beta-1, 6-N-acetyl-D-glucosamine biosynthesis protein PgaD | *Bacillus megaterium* | Strain ATCC 14581 / DSM 32 / JCM 2506 / NBRC 15308 / NCIMB 9376 / NCTC 10342 / NRRL B-14308 / VKM B-512 | A0A0B6AKG3 | -- | 25931591 | Not Available | Not Available |
| 875 | Amyloid fiber anchoring / assembly protein TapA (Biofilm formation protein) (Lipoprotein for biofilm formation) | *Bacillus glycinifermentans* | | A0A0T6BTZ5 | -- | 26297378 | Not Available | Not Available |
| 876 | Surface composition regulator | *Citrobacter pasteurii* | | A0A6N6K9P3 | -- | 30087402 | Not Available | Not Available |
| 877 | Surface composition regulator | *Salmonella enterica I* | | A0A3Z0GZU8 | -- | 30286803 | Not Available | Not Available |
| 878 | Poly-beta-1, 6-N-acetyl-D-glucosamine synthase (Poly-beta-1, 6-GlcNAc synthase) (EC 2.4.1.-) | *Acinetobacter baumannii* | | A0A0D8GQW0 | -- | 26170289 | Acetyl glucosaminyl transferase activity | Not Available |
| 879 | Anti-FlhC(2)FlhD(4) factor YdiV | *Salmonella typhimurium* | Strain LT2 / SGSC1412 / ATCC 700720 | Q8ZLB9 | -- | 11260463; 11677609; 11929533 | Not Available | Not Available |
| 880 | Biofilm synthesis auxiliary protein PgaD (Poly-beta-1, 6-N-acetyl-D-glucosamine auxiliary protein PgaD) (Poly-beta-1, 6-N-acetyl-D-glucosamine biosynthesis protein PgaD) | *Acinetobacter baumannii* | | A0A059ZRK6 | -- | 25136007; 25845845; 26170289 | Not Available | Not Available |
| 881 | Biofilm PGA synthesis lipoprotein pgaB (Poly-beta-1, 6-N-acetyl-D-glucosamine N-deacetylase PgaB) | *Acinetobacter baumannii* | | A0A6L8MEI5 | -- | 27917170; 29137647 | Hydrolase activity, acting on carbon-nitrogen (but not peptide) bonds | Not Available |
| 882 | Type II toxin-antitoxin system MqsR family toxin | *Pseudomonas spp. WS 5010* | | A0A7Y0ZGX5 | -- | 32587583 | Not Available | Not Available |
| 883 | mRNA interferase MqsR | *Gallionellales bacterium RIFCSPHIGHO2_02_FULL_57_16* | | A0A1G0EFX2 | -- | 27774985 | Not Available | Not Available |
| 884 | Poly-beta-1, 6-N-acetyl-D-glucosamine synthase (Poly-beta-1, 6-GlcNAc synthase) (EC 2.4.1.-) | *Pseudomonas spp. WS 5010* | | A0A7Y0ZL97 | -- | 32587583 | Acetyl glucosaminyl transferase activity | Not Available |

| Target ID | Bacterial Protein Name | Organism Name | Strain | UniProt ID | PDB ID | PubMed ID | Classification as per UniProt GO Molecular Function | Classification of Function as per PDB |
|---|---|---|---|---|---|---|---|---|
| 885 | Poly-beta-1, 6-N-acetyl-D-glucosamine biosynthesis protein PgaD | *Pseudomonas spp. WS 5010* | | A0A7Y0ZL43 | -- | 32587583 | Not Available | Not Available |
| 886 | Uncharacterized protein | *Burkholderiales bacterium GWF1_66_17* | | A0A1F4H8D6 | -- | 27774985 | Not Available | Not Available |
| 887 | Poly-beta-1, 6-N-acetyl-D-glucosamine biosynthesis protein PgaD | *Rhodocyclaceae bacterium* | | A0A7C6VPP3 | -- | 32123542 | Not Available | Not Available |
| 888 | Outer membrane N-deacetylase | *Achromobacter arsenitoxydans SY8* | | H0F5P5 | -- | 22328747 | Hydrolase activity, acting on carbon-nitrogen (but not peptide) bonds | Not Available |
| 889 | Poly-beta-1, 6-N-acetyl-D-glucosamine biosynthesis protein PgaD | *Pseudomonas spp. GM25* | | J2P592 | -- | 23045501 | Not Available | Not Available |
| 890 | Putative polysaccharide deacetylase | *Klebsiella pneumoniae subspp. pneumoniae* | Strain HS11286 | A0A0H3GL72 | -- | 22408243 | Hydrolase activity, acting on carbon-nitrogen (but not peptide) bonds | Not Available |
| 891 | Uncharacterized protein | *Rhodocyclales bacterium RIFCSPLOWO2_02_FULL_63_24* | | A0A1G3HT42 | -- | 27774985 | Not Available | Not Available |
| 892 | Uncharacterized protein | *Pseudomonas spp. GM25* | | J2XBE6 | -- | 23045501 | Not Available | Not Available |
| 893 | Biofilm PGA synthesis protein PgaD | *Pseudomonas spp.* | Strain M1 | W5INN3 | -- | 10388678; 23405299 | Not Available | Not Available |
| 894 | Uncharacterized protein | *Rhodocyclales bacterium RIFCSPLOWO2_02_FULL_63_24* | | A0A1G3HJF3 | -- | 27774985 | Not Available | Not Available |
| 895 | Poly-beta-1, 6-N-acetyl-D-glucosamine N-deacetylase PgaB (Fragment) | *Candidatus Desulfofervidus auxilii* | | A0A7C0Y3M0 | -- | 31911466 | Hydrolase activity, acting on carbon-nitrogen (but not peptide) bonds | Not Available |
| 896 | Putative hemin storage protein | *Bordetella bronchiseptica 253* | | A0A0C6P1R7 | -- | 23051057 | Hydrolase activity, acting on carbon-nitrogen (but not peptide) bonds | Not Available |
| 897 | Poly-beta-1, 6-N-acetyl-D-glucosamine synthase (Poly-beta-1, 6-GlcNAc synthase) (EC 2.4.1.-) | *Candidatus Desulfofervidus auxilii* | | A0A7C0Y6F6 | -- | 31911466 | Acetyl glucosaminyl transferase activity | Not Available |
| 898 | Surface composition regulator | *Kosakonia radicincitans DSM 16656* | | A0A1V0LHN3 | -- | 22965092 | Not Available | Not Available |
| 899 | Poly-beta-1, 6-N-acetyl-D-glucosamine synthase (Poly-beta-1, 6-GlcNAc synthase) (EC 2.4.1.-) | *Pseudomonas spp. GM25* | | J3F4T7 | -- | 23045501 | Acetyl glucosaminyl transferase activity | Not Available |
| 900 | Uncharacterized protein | *Bordetella bronchiseptica 253* | | A0A0C6P5T2 | -- | 23051057 | Not Available | Not Available |
| 901 | Poly-beta-1, 6-N-acetyl-D-glucosamine synthase (Poly-beta-1, 6-GlcNAc synthase) (EC 2.4.1.-) | *Bordetella bronchiseptica 253* | | A0A0C6P4Z6 | -- | 23051057 | Acetyl glucosaminyl transferase activity | Not Available |
| 902 | Motility quorum-sensing regulator MqsR | *Pseudomonas fluorescens* | | A0A166MX06 | -- | 29769716 | Not Available | Not Available |
| 903 | Poly-beta-1, 6-N-acetyl-D-glucosamine synthase (Poly-beta-1, 6-GlcNAc synthase) (EC 2.4.1.-) | *Stenotrophomonas spp. DDT-1* | | A0A149QNS6 | -- | 26888254 | Acetyl glucosaminyl transferase activity | Not Available |
| 904 | mRNA interferase MqsR | *Betaproteobacteria bacterium HGW-Betaproteobacteria-3* | | A0A2N2RGM4 | -- | 28350393 | Not Available | Not Available |
| 905 | Type II toxin-antitoxin system MqsR family toxin | *Pseudomonas fluorescens* | | A0A7Z8NMX6 | -- | 31040301 | Not Available | Not Available |
| 906 | Poly-beta-1, 6-N-acetyl-D-glucosamine synthase (Poly-beta-1, 6-GlcNAc synthase) (EC 2.4.1.-) | *Pseudomonas fluorescens* | | A0A160A1X0 | -- | 29769716 | Acetyl glucosaminyl transferase activity | Not Available |
| 907 | Hemin storage protein | *Stenotrophomonas spp. DDT-1* | | A0A149QNot Available0 | -- | 26888254 | Not Available | Not Available |
| 908 | Poly-beta-1, 6-N-acetyl-D-glucosamine synthase (Poly-beta-1, 6-GlcNAc synthase) (EC 2.4.1.-) | *Pseudomonas fluorescens* | | A0A161Z6J8 | -- | 29769716 | Acetyl glucosaminyl transferase activity | Not Available |
| 909 | Poly-beta-1, 6-N-acetyl-D-glucosamine biosynthesis protein PgaD | *Pseudomonas fluorescens* | | A0A166NCZ6 | -- | 29769716 | Not Available | Not Available |
| 910 | Type II toxin-antitoxin system MqsR family toxin | *Pseudomonas veronii* | | A0A7Y0ZQP0 | -- | 32587583 | Not Available | Not Available |
| 911 | Motility quorum-sensing regulator MqsR | *Pseudomonas spp. S49* | | A0A7Z2M4T1 | -- | 25398872 | Not Available | Not Available |
| 912 | Poly-beta-1, 6-N-acetyl-D-glucosamine N-deacetylase PgaB (EC 3.5.1.-) | *Pseudomonas veronii* | | A0A7Y0ZZU3 | -- | 32587583 | Hydrolase activity, acting on carbon-nitrogen (but not peptide) bonds | Not Available |
| 913 | Poly-beta-1, 6-N-acetyl-D-glucosamine biosynthesis protein PgaD | *Pseudomonas fluorescens* | | A0A0N9WHA4 | -- | 29769716 | Not Available | Not Available |
| 914 | Uncharacterized protein | *Mycetohabitans rhizoxinica* | Strain DSM 19002 / CIP 109453 / HKI 454 Paraburkholderia rhizoxinica | E5AT95 | -- | 21131495 | Not Available | Not Available |
| 915 | Poly-beta-1, 6-N-acetyl-D-glucosamine N-deacetylase | *Pseudomonas fluorescens* | | A0A160A2W4 | -- | 29769716 | Hydrolase activity, acting on carbon-nitrogen (but not peptide) bonds | Not Available |
| 916 | Poly-beta-1, 6-N-acetyl-D-glucosamine biosynthesis protein PgaD | *Pseudomonas fluorescens* | | A0A160A0I6 | -- | 29769716 | Not Available | Not Available |
| 917 | Poly-beta-1, 6-N-acetyl-D-glucosamine N-deacetylase PgaB | *Pseudomonas fluorescens* | | A0A0N9W7I0 | -- | 29769716 | Hydrolase activity, acting on carbon-nitrogen (but not peptide) bonds | Not Available |
| 918 | Poly-beta-1, 6-N-acetyl-D-glucosamine N-deacetylase PgaB (EC 3.5.1.-) | *Pseudomonas veronii* | | A0A7Y1ABU3 | -- | 32587583 | Hydrolase activity, acting on carbon-nitrogen (but not peptide) bonds | Not Available |
| 919 | Putative biofilm formation, controlled by quorum sensing, formation of cell aggregates | *Pseudoalteromonas translucida* | Strain TAC 125 | Q3IK57 | -- | 16169927 | Not Available | Not Available |
| 920 | Poly-beta-1, 6-N-acetyl-D-glucosamine biosynthesis protein PgaD | *Pseudomonas fluorescens* | | A0A0N9WGY1 | -- | 29769716 | Not Available | Not Available |
| 921 | Poly-beta-1, 6-N-acetyl-D-glucosamine synthase (Poly-beta-1, 6-GlcNAc synthase) (EC 2.4.1.-) | *Pseudomonas fluorescens* | | A0A0N9WHZ0 | -- | 29769716 | Acetyl glucosaminyl transferase activity | Not Available |
| 922 | Hemin storage protein | *Stenotrophomonas spp. DDT-1* | | A0A149QN88 | -- | 26888254 | Hydrolase activity, acting on carbon-nitrogen (but not peptide) bonds | Not Available |
| 923 | Poly-beta-1, 6-N-acetyl-D-glucosamine N-deacetylase PgaB | *Pseudomonas fluorescens* | | A0A0N9VTC1 | -- | 29769716 | Hydrolase activity, acting on carbon-nitrogen (but not peptide) bonds | Not Available |
| 924 | Type II toxin-antitoxin system MqsR family toxin | *Pseudomonas veronii* | | A0A7Y1ABA9 | -- | 32587583 | Not Available | Not Available |
| 925 | Putative membrane protein | *Xanthomonas citri pv. mangiferaeindicae LMG 941* | | H8FLX0 | -- | 22582385 | Not Available | Not Available |
| 926 | Polysaccharide deacetylase | *Xenorhabdus spp. KK7.4* | | A0A2D0K518 | -- | 28993611 | Hydrolase activity, acting on carbon-nitrogen (but not peptide) bonds | Not Available |
| 927 | Poly-beta-1, 6-N-acetyl-D-glucosamine synthase (Poly-beta-1, 6-GlcNAc synthase) (EC 2.4.1.-) | *Staphylococcus delphini* | | A0A2A4GXC5 | -- | 28859126 | Acetyl glucosaminyl transferase activity | Not Available |
| 928 | Poly-beta-1, 6-N-acetyl-D-glucosamine biosynthesis protein PgaD | *Pseudomonas veronii* | | A0A7Y1ABU2 | -- | 32587583 | Not Available | Not Available |
| 929 | Poly-beta-1, 6-N-acetyl-D-glucosamine synthase (Poly-beta-1, 6-GlcNAc synthase) (EC 2.4.1.-) | *Pseudomonas veronii* | | A0A7Y1ABV4 | -- | 32587583 | Acetyl glucosaminyl transferase activity | Not Available |
| 930 | Poly-beta-1, 6-N-acetyl-D-glucosamine N-deacetylase (EC 3.5.1.-) | *Xanthomonas citri pv. mangiferaeindicae LMG 941* | | H8FLX2 | -- | 22582385 | Hydrolase activity, acting on carbon-nitrogen (but not peptide) bonds | Not Available |
| 931 | Poly-beta-1, 6-N-acetyl-D-glucosamine biosynthesis protein PgaD | *Pseudomonas spp. S49* | | A0A7Z2M4F2 | -- | 25398872 | Not Available | Not Available |
| 932 | Poly-beta-1, 6-N-acetyl-D-glucosamine synthase (Poly-beta-1, 6-GlcNAc synthase) (EC 2.4.1.-) | *Pseudomonas spp. S49* | | A0A7Z2M356 | -- | 25398872 | Acetyl glucosaminyl transferase activity | Not Available |

| Target ID | Bacterial Protein Name | Organism Name | Strain | UniProt ID | PDB ID | PubMed ID | Classification as per UniProt GO Molecular Function | Classification of Function as per PDB |
|---|---|---|---|---|---|---|---|---|
| 933 | Poly-beta-1, 6-N-acetyl-D-glucosamine synthase (Poly-beta-1, 6-GlcNAc synthase) (EC 2.4.1.-) | *Staphylococcus delphini* | | A0A2A4HFR7 | -- | 28859126 | Acetyl glucosaminyl transferase activity | Not Available |
| 934 | Poly-beta-1, 6-N-acetyl-D-glucosamine biosynthesis protein PgaD | *Xenorhabdus spp. KK7.4* | | A0A2D0K575 | -- | 28993611 | Not Available | Not Available |
| 935 | Poly-beta-1, 6-N-acetyl-D-glucosamine biosynthesis protein PgaD | *Pseudomonas veronii* | | A0A0R2ZU73 | -- | 32587583 | Not Available | Not Available |
| 936 | Poly-beta-1, 6-N-acetyl-D-glucosamine N-deacetylase | *Pseudomonas spp. S49* | | A0A7Z2M3J1 | -- | 25398872 | Hydrolase activity, acting on carbon-nitrogen (but not peptide) bonds | Not Available |
| 937 | Poly-beta-1, 6-N-acetyl-D-glucosamine synthase (Poly-beta-1, 6-GlcNAc synthase) (EC 2.4.1.-) | *Xanthomonas citri pv. mangiferaeindicae LMG 941* | | H8FLX1 | -- | 22582385 | Acetyl glucosaminyl transferase activity | Not Available |
| 938 | Type II toxin-antitoxin system MqsR family toxin | *Solimona terrae* | | A0A6M2BWG2 | -- | 24421276 | Not Available | Not Available |
| 939 | Uncharacterized protein | *Achromobacter arsenitoxydans SY8* | | H0F5P7 | -- | 22328747 | Not Available | Not Available |
| 940 | Poly-beta-1, 6-N-acetyl-D-glucosamine synthase (Poly-beta-1, 6-GlcNAc synthase) (EC 2.4.1.-) | *Achromobacter arsenitoxydans SY8* | | H0F5P6 | -- | 22328747 | Acetyl glucosaminyl transferase activity | Not Available |
| 941 | Poly-beta-1, 6-N-acetyl-D-glucosamine N-deacetylase PgaB | *Xanthomonadales bacterium RIFOXYA1_FULL_69_10* | | A0A1G4A9M8 | -- | 27774985 | Hydrolase activity, acting on carbon-nitrogen (but not peptide) bonds | Not Available |
| 942 | Poly-beta-1, 6-N-acetyl-D-glucosamine biosynthesis protein PgaD | *Enterobacteriaceae bacterium ENNIH2* | | A0A2I8QSJ0 | -- | 29437920 | Not Available | Not Available |
| 943 | Poly-beta-1, 6-N-acetyl-D-glucosamine synthase (Poly-beta-1, 6-GlcNAc synthase) (EC 2.4.1.-) | *Xanthomonadales bacterium RIFOXYA1_FULL_69_10* | | A0A1G4A9H6 | -- | 27774985 | Acetyl glucosaminyl transferase activity | Not Available |
| 944 | Poly-beta-1, 6-N-acetyl-D-glucosamine biosynthesis protein PgaD | *Enterobacter spp. EC-NT1* | | A0A3D8XP45 | -- | 29665424 | Not Available | Not Available |
| 945 | Poly-beta-1, 6-N-acetyl-D-glucosamine synthase (Poly-beta-1, 6-GlcNAc synthase) (EC 2.4.1.-) | *Enterobacter spp. EC-NT1* | | A0A3D8XNY3 | -- | 29665424 | Acetyl glucosaminyl transferase activity | Not Available |
| 946 | Poly-beta-1, 6-N-acetyl-D-glucosamine N-deacetylase PgaB | *Enterobacter spp. EC-NT1* | | A0A3D8XP54 | -- | 29665424 | Hydrolase activity, acting on carbon-nitrogen (but not peptide) bonds | Not Available |
| 947 | Poly-beta-1, 6-N-acetyl-D-glucosamine synthase (Poly-beta-1, 6-GlcNAc synthase) (EC 2.4.1.-) | *Pseudomonas lactis* | | A0A7Y1LIK1 | -- | 32587583 | Acetyl glucosaminyl transferase activity | Not Available |
| 948 | Motility quorum-sensing regulator MqsR | *Agrobacterium rubi* | | A0A1B9TWB4 | -- | 27547538 | Not Available | Not Available |
| 949 | Poly-beta-1, 6-N-acetyl-D-glucosamine biosynthesis protein PgaD | *Pseudomonas lactis* | | A0A7Y1LIN2 | -- | 32587583 | Not Available | Not Available |
| 950 | Poly-beta-1, 6-N-acetyl-D-glucosamine N-deacetylase PgaB | *Pseudomonas spp. GM30* | | W6VK76 | -- | 23045501 | Hydrolase activity, acting on carbon-nitrogen (but not peptide) bonds | Not Available |
| 951 | Poly-beta-1, 6-N-acetyl-D-glucosamine biosynthesis protein PgaD | *Pseudomonas spp. GM30* | | W6W1H7 | -- | 23045501 | Not Available | Not Available |
| 952 | Surface composition regulator | *Enterobacter spp. EC-NT1* | | A0A3D8XGN6 | -- | 29665424 | Not Available | Not Available |
| 953 | Biofilm PGA synthesis protein PgaD | *Pseudomonas spp. Os17* | | A0A0D6B9Q4 | -- | 25496595 | Not Available | Not Available |
| 954 | Uncharacterized protein | *Pseudomonas fuscovaginae* | | A0A0N0VKM0 | -- | 26422147 | Not Available | Not Available |
| 955 | Uncharacterized protein | *Deltaproteobacteria bacterium RIFOXYD12_FULL_50_9* | | A0A1F9MN67 | -- | 27774985 | Not Available | Not Available |
| 956 | Outer membrane N-deacetylase | *Pseudomonas spp. Os17* | | A0A0D6BA39 | -- | 25496595 | Hydrolase activity, acting on carbon-nitrogen (but not peptide) bonds | Not Available |
| 957 | Poly-beta-1, 6-N-acetyl-D-glucosamine synthase (Poly-beta-1, 6-GlcNAc synthase) (EC 2.4.1.-) | *Pseudomonas spp. Os17* | | A0A0D6B9S5 | -- | 25496595 | Acetyl glucosaminyl transferase activity | Not Available |
| 958 | Poly-beta-1, 6-N-acetyl-D-glucosamine synthase (Poly-beta-1, 6-GlcNAc synthase) (EC 2.4.1.-) | *Leclercia spp. LSNIH1* | | A0A2I8NRS9 | -- | 29437920 | Acetyl glucosaminyl transferase activity | Not Available |
| 959 | Poly-beta-1, 6-N-acetyl-D-glucosamine synthase (Poly-beta-1, 6-GlcNAc synthase) (EC 2.4.1.-) | *Pseudomonas fuscovaginae* | | A0A0M9GJ16 | -- | 26422147 | Acetyl glucosaminyl transferase activity | Not Available |
| 960 | Poly-beta-1, 6-N-acetyl-D-glucosamine N-deacetylase PgaB | *Pseudomonas fuscovaginae* | | A0A0N0VKP0 | -- | 26422147 | Hydrolase activity, acting on carbon-nitrogen (but not peptide) bonds | Not Available |
| 961 | Poly-beta-1, 6-N-acetyl-D-glucosamine biosynthesis protein PgaD | *Leclercia spp. LSNIH1* | | A0A2I8NRR5 | -- | 29437920 | Not Available | Not Available |
| 962 | Surface composition regulator | *Salmonella newport* | Strain SL254 | A0A0H3BM79 | -- | 21602358 | Not Available | Not Available |
| 963 | Poly-beta-1, 6-N-acetyl-D-glucosamine synthase (Poly-beta-1, 6-GlcNAc synthase) (EC 2.4.1.-) | *Burkholderia spp. KJ006* | | I2DW95 | -- | 22843575 | Acetyl glucosaminyl transferase activity | Not Available |
| 964 | Poly-beta-1, 6-N-acetyl-D-glucosamine biosynthesis protein PgaD | *Pseudomonas spp. GM80* | | J3DTA0 | -- | 23045501 | Not Available | Not Available |
| 965 | Biofilm PGA synthesis deacetylase PgaB | *Burkholderia spp. KJ006* | | I2DW94 | -- | 22843575 | Hydrolase activity, acting on carbon-nitrogen (but not peptide) bonds | Not Available |
| 966 | Uncharacterized protein | *Acidithiobacillus ferrivorans SS3* | | G0JSS8 | -- | 21705598 | Not Available | Not Available |
| 967 | Uncharacterized protein | *Pseudomonas protegens* | Strain DSM 19095 / LMG 27888 / CFBP 6595 / CHA0 | A0A2C9EED5 | -- | 24762936 | Not Available | Not Available |
| 968 | Surface composition regulator | *Salmonella heidelberg* | Strain SL476 | A0A6C6ZPJ5 | -- | 21602358 | Not Available | Not Available |
| 969 | Surface composition regulator | *Salmonella enterica subspp. enterica serovar* | Lubbock | A0A6C8LYN6 | -- | 31527278 | Not Available | Not Available |
| 970 | Poly-beta-1, 6-N-acetyl-D-glucosamine biosynthesis protein PgaD | *Xanthomonadales bacterium RIFOXYA1_FULL_69_10* | | A0A1G4A9L8 | -- | 27774985 | Not Available | Not Available |
| 971 | Uncharacterized protein | *Pseudomonas protegens* | Strain DSM 19095 / LMG 27888 / CFBP 6595 / CHA0 | A0A2C9EGU0 | -- | 24762936 | Not Available | Not Available |
| 972 | Poly-beta-1, 6-N-acetyl-D-glucosamine N-deacetylase PgaB | *Enterobacteriaceae bacterium ENNIH2* | | A0A2I8QUW8 | -- | 29437920 | Hydrolase activity, acting on carbon-nitrogen (but not peptide) bonds | Not Available |
| 973 | Poly-beta-1, 6-N-acetyl-D-glucosamine N-deacetylase PgaB (EC 3.5.1.-) | *Pseudomonas protegens* | Strain DSM 19095 / LMG 27888 / CFBP 6595 / CHA0 | A0A2C9EE93 | -- | 24762936 | Hydrolase activity, acting on carbon-nitrogen (but not peptide) bonds | Not Available |
| 974 | Poly-beta-1, 6-N-acetyl-D-glucosamine synthase (Poly-beta-1, 6-GlcNAc synthase) (EC 2.4.1.-) | *Enterobacteriaceae bacterium ENNIH2* | | A0A2I8QSF7 | -- | 29437920 | Acetyl glucosaminyl transferase activity | Not Available |
| 975 | Surface composition regulator | *Salmonella enterica subspp. enterica serovar* | Alachua Str. R6-377 | G5LTU2 | -- | 21859443 | Not Available | Not Available |
| 976 | Poly-beta-1, 6-N-acetyl-D-glucosamine synthase (Poly-beta-1, 6-GlcNAc synthase) (EC 2.4.1.-) | *Pseudomonas silesiensis* | | A0A191YLQ5 | -- | 29153257 | Acetyl glucosaminyl transferase activity | Not Available |
| 977 | mRNA interferase MqsR | *Pseudomonas silesiensis* | | A0A191YXN6 | -- | 29153257 | Not Available | Not Available |
| 978 | Poly-beta-1, 6-N-acetyl-D-glucosamine N-deacetylase PgaB | *Ralstonia pickettii OR214* | | R0CKR4 | -- | 23792748 | Hydrolase activity, acting on carbon-nitrogen (but not peptide) bonds | Not Available |
| 979 | Uncharacterized protein | *Arenimona spp. SCN 70-307* | | A0A1D2RX64 | -- | 26031303 | Not Available | Not Available |
| 980 | Polysaccharide deacetylase (EC 3.-.-.-) | *Sulfurimona gotlandica* | Strain DSM 19862 / JCM 16533 / GD1 | H1FST3 | -- | 22203982 | Hydrolase activity, acting on carbon-nitrogen (but not peptide) bonds | Not Available |
| 981 | mRNA interferase MqsR | *Herbaspirillum aquaticum* | | A0A225STC8 | -- | 19671717 | Not Available | Not Available |
| 982 | Uncharacterized protein | *Sulfurimona gotlandica* | Strain DSM 19862 / JCM 16533 / GD1 | B6BJF3 | -- | 22203982 | Not Available | Not Available |

| Target ID | Bacterial Protein Name | Organism Name | Strain | UniProt ID | PDB ID | PubMed ID | Classification as per UniProt GO Molecular Function | Classification of Function as per PDB |
|---|---|---|---|---|---|---|---|---|
| 983 | Poly-beta-1, 6-N-acetyl-D-glucosamine synthase (Poly-beta-1, 6-GlcNAc synthase) (EC 2.4.1.-) | *Ralstonia pickettii OR214* | | R0CK10 | -- | 23792748 | Acetyl glucosaminyl transferase activity | Not Available |
| 984 | Poly-beta-1, 6-N-acetyl-D-glucosamine synthase (Poly-beta-1, 6-GlcNAc synthase) (EC 2.4.1.-) | *Sulfurimona gotlandica* | Strain DSM 19862 / JCM 16533 / GD1 | B6BJF4 | -- | 22203982 | Acetyl glucosaminyl transferase activity | Not Available |
| 985 | Poly-beta-1, 6-N-acetyl-D-glucosamine biosynthesis protein PgaD | *Pseudomonas silesiensis* | | A0A191YLN0 | -- | 29153257 | Not Available | Not Available |
| 986 | Surface composition regulator | *Salmonella enterica subspp. enterica serovar* | Montevideo Str. S5-403 | G5Q1S1 | -- | 21859443 | Not Available | Not Available |
| 987 | Uncharacterized protein | *Lactobacillus spp. ASF360* | | N1ZVW5 | -- | 24723722 | Not Available | Not Available |
| 988 | Poly-beta-1, 6-N-acetyl-D-glucosamine N-deacetylase PgaB | *Pseudomonas silesiensis* | | A0A191YLE3 | -- | 29153257 | Hydrolase activity, acting on carbon-nitrogen (but not peptide) bonds | Not Available |
| 989 | Poly-beta-1, 6-N-acetyl-D-glucosamine N-deacetylase PgaB | *Aeromona veronii Hm21* | | A0A7Z3YI03 | -- | 24092791 | Hydrolase activity, acting on carbon-nitrogen (but not peptide) bonds | Not Available |
| 990 | Uncharacterized protein | *Ralstonia pickettii OR214* | | R0E6J7 | -- | 23792748 | Not Available | Not Available |
| 991 | Poly-beta-1, 6-N-acetyl-D-glucosamine synthase (Poly-beta-1, 6-GlcNAc synthase) (EC 2.4.1.-) | *Aeromona veronii Hm21* | | A0A7Z3YHZ5 | -- | 24092791 | Acetyl glucosaminyl transferase activity | Not Available |
| 992 | mRNA interferase MqsR | *Advenella kashmirensis W13003* | | V8QRA6 | -- | 24482505 | Not Available | Not Available |
| 993 | Uncharacterized protein | *Sphingobium spp. MI1205* | | A0A126RE24 | -- | 27056230 | Not Available | Not Available |
| 994 | Poly-beta-1, 6-N-acetyl-D-glucosamine synthase (Poly-beta-1, 6-GlcNAc synthase) (EC 2.4.1.-) | *Pseudomonas protegens* | Strain DSM 19095 / LMG 27888 / CFBP 6595 / CHA0 | A0A2C9EE84 | -- | 24762936 | Acetyl glucosaminyl transferase activity | Not Available |
| 995 | Poly-beta-1, 6-N-acetyl-D-glucosamine synthase (Poly-beta-1, 6-GlcNAc synthase) (EC 2.4.1.-) | *Xenorhabdus spp. KK7.4* | | A0A2D0K576 | -- | 28993611 | Acetyl glucosaminyl transferase activity | Not Available |
| 996 | Putative intercellular adhesion protein A (Biofilm formation) | *Vibrio spp. AND4* | | A8T1C7 | -- | 20436956 | Not Available | Not Available |
| 997 | Poly-beta-1, 6-N-acetyl-D-glucosamine synthase (Poly-beta-1, 6-GlcNAc synthase) (EC 2.4.1.-) | *Pseudomonas veronii* | | A0A0R2ZUD4 | -- | 32587583 | Acetyl glucosaminyl transferase activity | Not Available |
| 998 | Lipoprotein for biofilm formation | *Bacillus stratosphericus LAMA 585* | | M5QYB3 | -- | 23640380 | Not Available | Not Available |
| 999 | GGDEF domain-containing protein | *Streptomyces coelicolor* | Strain ATCC BAA-471 / A3 2 / M145 | Q9AD37 | -- | 12000953 | Diguanylate cyclase activity, helicase activity | Not Available |
| 1000 | Poly-beta-1, 6-N-acetyl-D-glucosamine synthase (Poly-beta-1, 6-GlcNAc synthase) (EC 2.4.1.-) | *Yersinia entomophaga* | | A0A3S6EX22 | -- | 27187466 | Acetyl glucosaminyl transferase activity | Not Available |
| 1001 | Biofilm PGA synthesis lipoprotein PgaB | *Enterobacter cloacae subspp. cloacae* | Strain ATCC 13047 / DSM 30054 / NBRC 13535 / NCTC 10005 / WDCM 00083 / NCDC 279-56 | A0A0H3CQE5 | -- | 20207761 | Hydrolase activity, acting on carbon-nitrogen (but not peptide) bonds | Not Available |
| 1002 | Uncharacterized protein | *Klebsiella pneumoniae* | | Q1G0V3 | -- | 16861646 | Not Available | Not Available |
| 1003 | Surface composition regulator | *Salmonella enterica subspp. enterica serovar* | Javiana | A0A736M1L3 | -- | 30286803 | Not Available | Not Available |
| 1004 | Surface composition regulator | *Salmonella enterica subspp. salamae serovar* | 42:z:1, 5 | A0A735PV97 | -- | 30286803 | Not Available | Not Available |
| 1005 | IcaA (Fragment) | *Staphylococcus cohnii* | | Q83ZH4 | -- | 12957956 | Not Available | Not Available |
| 1006 | Uncharacterized protein (Fragment) | *Klebsiella pneumoniae* | | Q1G0U7 | -- | 16861646 | Not Available | Not Available |
| 1007 | Uncharacterized protein (Fragment) | *Klebsiella pneumoniae* | | Q1G0W1 | -- | 16861646 | Not Available | Not Available |
| 1008 | Uncharacterized protein | *Klebsiella pneumoniae* | | Q1G0V7 | -- | 16861646 | Not Available | Not Available |
| 1009 | Uncharacterized protein | *Klebsiella pneumoniae* | | Q1G0V9 | -- | 16861646 | Not Available | Not Available |
| 1010 | Aldedh domain-containing protein (Fragment) | *Klebsiella pneumoniae* | | Q1G0V2 | -- | 16861646 | Oxidoreductase activity, acting on the aldehyde or oxo group of donors, NAD or NADP as acceptor | Not Available |
| 1011 | Motility quorum-sensing regulator mqsR | *Yersinia enterocolitica W22703* | | F4N438 | -- | 21453472 | Not Available | Not Available |
| 1012 | Uncharacterized protein | *Klebsiella pneumoniae* | | Q1G0V6 | -- | 16861646 | Not Available | Not Available |
| 1013 | Surface composition regulator | *Salmonella enterica subspp. enterica serovar* | Panama | A0A751YZQ8 | -- | 30286803 | Not Available | Not Available |
| 1014 | Motility quorum-sensing regulator mqsR | *uncultured Desulfobacterium spp.* | | E1YKP6 | -- | 21176053 | Not Available | Not Available |
| 1015 | Uncharacterized protein (Fragment) | *Klebsiella pneumoniae* | | Q1G0V1 | -- | 16861646 | Not Available | Not Available |
| 1016 | Uncharacterized protein (Fragment) | *Klebsiella pneumoniae* | | Q1G0U3 | -- | 16861646 | Not Available | Not Available |
| 1017 | Uncharacterized protein | *Klebsiella pneumoniae* | | Q1G0U1 | -- | 16861646 | Not Available | Not Available |
| 1018 | PTS EIIB type-1 domain-containing protein (Fragment) | *Klebsiella pneumoniae* | | Q1G0V0 | -- | 16861646 | Protein-N(PI)-phosphohistidine-sugar phosphoTransferase activity | Not Available |
| 1019 | Biofilm PGA synthesis protein PgaD | *Pseudomonas moraviensis R28-S* | | V8REE0 | -- | 24558233 | Not Available | Not Available |
| 1020 | Biofilm PGA synthesis protein PgaD, putative | *Pseudomonas fluorescens Q2-87* | | J2ERK9 | -- | 22792073 | Not Available | Not Available |
| 1021 | Surface composition regulator | *Salmonella enterica subspp. enterica serovar* | Javiana | A0A733VDE6 | -- | 30286803 | Not Available | Not Available |
| 1022 | Poly-beta-1, 6-N-acetyl-D-glucosamine synthase (Poly-beta-1, 6-GlcNAc synthase) (EC 2.4.1.-) | *Pseudomonas fluorescens Q2-87* | | J2XWR0 | -- | 22792073 | Acetyl glucosaminyl transferase activity | Not Available |
| 1023 | Surface composition regulator | *Salmonella enterica subspp. salamae serovar* | 30:1, z28:z6 | A0A737XXT1 | -- | 30286803 | Not Available | Not Available |
| 1024 | Biofilm PGA synthesis lipoprotein PgaB (EC 3.-.-.-) | *Pseudomonas fluorescens Q2-87* | | J2MEN5 | -- | 22792073 | Hydrolase activity, acting on carbon-nitrogen (but not peptide) bonds | Not Available |
| 1025 | Surface composition regulator | *Salmonella enterica subspp. salamae serovar* | 58:l, z13, z28:z6 | A0A734HKS5 | -- | 30286803 | Not Available | Not Available |
| 1026 | Uncharacterized protein | *Shewanella spp. 33B* | | B9W0T4 | -- | 19336243 | Not Available | Not Available |
| 1027 | Motility quorum-sensing regulator MqsR | *Pseudomonas fluorescens Q2-87* | | J2YB16 | -- | 22792073 | Not Available | Not Available |
| 1028 | Surface composition regulator | *Salmonella enterica subspp. enterica serovar* | Typhi Str. CT18 | A0A714V9M2 | -- | 30286803 | Not Available | Not Available |
| 1029 | UPF0441 protein GKC49_07595 | *Enterobacter agglomerans* | Erwinia herbicola Pantoea agglomerans | A0A6I6K1P2 | -- | 32966572 | Not Available | Not Available |
| 1030 | Surface composition regulator | *Salmonella enterica subspp. salamae* | | A0A3U3WQ67 | -- | 30286803 | Not Available | Not Available |
| 1031 | Biofilm formation regulator BssS (Biofilm formation regulatory protein BssS) | *Enterobacter agglomerans* | Erwinia herbicola Pantoea agglomerans | A0A356RXP9 | -- | 32966572 | Not Available | Not Available |
| 1032 | Hha toxicity modulator TomB (Toxin overexpression modulator in biofilms) | *Enterobacter agglomerans* | Erwinia herbicola Pantoea agglomerans | A0A1T4JJK3 | -- | 32966572 | Not Available | Not Available |
| 1033 | Surface composition regulator | *Salmonella enterica subspp. enterica serovar* | Panama | A0A3T2YKC2 | -- | 30286803 | Not Available | Not Available |
| 1034 | Poly-beta-1, 6-N-acetyl-D-glucosamine synthase (Poly-beta-1, 6-GlcNAc synthase) (EC 2.4.1.-) | *Pusillimona spp.* | | A0A2E4R4G5 | -- | 29337314 | Acetyl glucosaminyl transferase activity | Not Available |
| 1035 | Surface composition regulator | *Salmonella enterica subspp. enterica serovar* | Reading | A0A5H8IBP9 | -- | 30286803 | Not Available | Not Available |
| 1036 | Poly-beta-1, 6-N-acetyl-D-glucosamine biosynthesis protein PgaD | *Pusillimona spp.* | | A0A2E4R6T6 | -- | 29337314 | Not Available | Not Available |

| Target ID | Bacterial Protein Name | Organism Name | Strain | UniProt ID | PDB ID | PubMed ID | Classification as per UniProt GO Molecular Function | Classification of Function as per PDB |
|---|---|---|---|---|---|---|---|---|
| 1037 | Biofilm PGA synthesis auxiliary protein PgaD (Poly-beta-1, 6-N-acetyl-D-glucosamine biosynthesis protein PgaD) | *Pseudomonas synxantha* | | A0A0R2YFQ5 | -- | 27231365 | Not Available | Not Available |
| 1038 | Poly-beta-1, 6-N-acetyl-D-glucosamine synthase (Poly-beta-1, 6-GlcNAc synthase) (EC 2.4.1.-) | *Staphylococcus pseudintermedius* | | A0A317YUG5 | -- | 29292005; 30173756 | Acetyl glucosaminyl transferase activity | Not Available |
| 1039 | Surface composition regulator | *Salmonella enterica subspp. enterica serovar* | Litchfield | A0A3W0FDC6 | -- | 30286803 | Not Available | Not Available |
| 1040 | Poly-beta-1, 6-N-acetyl-D-glucosamine synthase (Poly-beta-1, 6-GlcNAc synthase) (EC 2.4.1.-) | *Piscirickettsiaceae bacterium* | | A0A2A4W8B8 | -- | 29099490 | Acetyl glucosaminyl transferase activity | Not Available |
| 1041 | Poly-beta-1, 6-N-acetyl-D-glucosamine synthase (Poly-beta-1, 6-GlcNAc synthase) (EC 2.4.1.-) | *Xanthomonas cucurbitae* | | A0A2S7DKF0 | -- | 33734871 | Acetyl glucosaminyl transferase activity | Not Available |
| 1042 | Surface composition regulator | *Salmonella enterica subspp. enterica serovar* | Bovismorbificans | A0A630XF19 | -- | 30286803 | Not Available | Not Available |
| 1043 | Biofilm PGA synthesis lipoprotein PgaB (Poly-beta-1, 6-N-acetyl-D-glucosamine N-deacetylase) | *Pseudomonas baetica* | | A0A2N0DQC4 | -- | 29579234 | Hydrolase activity, acting on carbon-nitrogen (but not peptide) bonds | Not Available |
| 1044 | Enoyl-CoA hydratase | *Acinetobacter oleivorans* | Strain JCM 16667 / KCTC 23045 / DR1 | D3JTG2 | -- | 20629796; 20639327 | catalytic activity | Not Available |
| 1045 | Motility quorum-sensing regulator / GCU-specific mRNA interferase toxin | *Nitrosospira multiformis* | Strain ATCC 25196 / NCIMB 11849 / C 71 | Q2Y6X0 | -- | 18390676 | Not Available | Not Available |
| 1046 | Outer membrane N-deacetylase (EC 3.5.1.-) (Poly-beta-1, 6-N-acetyl-D-glucosamine N-deacetylase) (Poly-beta-1, 6-N-acetyl-D-glucosamine N-deacetylase PgaB) | *Klebsiella pneumoniae* | | A0A2V3JZQ6 | -- | 30373801; 32429555 | Hydrolase activity, acting on carbon-nitrogen (but not peptide) bonds | Not Available |
| 1047 | Poly-beta-1, 6-N-acetyl-D-glucosamine synthase (Poly-beta-1, 6-GlcNAc synthase) (EC 2.4.1.-) | *Carnobacterium divergens* | Lactobacillus divergens | A0A5F0MNT9 | -- | 30081251 | Acetyl glucosaminyl transferase activity | Not Available |
| 1048 | Poly-beta-1, 6-N-acetyl-D-glucosamine synthase (Poly-beta-1, 6-GlcNAc synthase) (EC 2.4.1.-) | *Enterobacter hormaechei* | | A0A331VAX8 | -- | 30286803 | Acetyl glucosaminyl transferase activity | Not Available |
| 1049 | Biofilm PGA synthesis deacetylase, xylanase / chitin deacetylase (Hemin storage protein) (Poly-beta-1, 6-N-acetyl-D-glucosamine N-deacetylase PgaB) | *Acinetobacter baumannii* | | A0A219CD45 | -- | 30124845 | Hydrolase activity, acting on carbon-nitrogen (but not peptide) bonds, Hydrolase activity, acting on glycosyl bonds | Not Available |
| 1050 | Poly-beta-1, 6-N-acetyl-D-glucosamine synthase (Poly-beta-1, 6-GlcNAc synthase) (EC 2.4.1.-) | *Acinetobacter baumannii* | | A0A505MFJ4 | -- | 29137647 | Acetyl glucosaminyl transferase activity | Not Available |
| 1051 | Poly-beta-1, 6-N-acetyl-D-glucosamine biosynthesis protein PgaD | *Klebsiella pneumoniae* | | A0A0S3FBI0 | -- | 30373801; 32429555 | Not Available | Not Available |
| 1052 | Uncharacterized protein | *Companilactobacillus farciminis* | | A0A0H4LUD2 | -- | 26383668 | Not Available | Not Available |
| 1053 | Poly-beta-1, 6-N-acetyl-D-glucosamine synthase (Poly-beta-1, 6-GlcNAc synthase) (EC 2.4.1.-) | *Carnobacterium divergens* | Lactobacillus divergens | A0A2R8A3Q4 | -- | 30081251 | Acetyl glucosaminyl transferase activity | Not Available |
| 1054 | Poly-beta-1, 6-N-acetyl-D-glucosamine biosynthesis protein PgaD | *Acinetobacter baumannii* | | A0A0J8V218 | -- | 29437920 | Not Available | Not Available |
| 1055 | Surface composition regulator | *Salmonella derby* | | A0A3W0AU60 | -- | 30286803; 30902850 | Not Available | Not Available |
| 1056 | Fimbrial protein (Fimbrial-like protein) (MrkF) (Type 1 fimbrial protein) (Type 3 fimbria minor subunit MrkF) | *Klebsiella pneumoniae* | | A0A0W8ANM7 | -- | 21901098; 30373801; 29437920; 32429555 | Not Available | Not Available |
| 1057 | Surface composition regulator | *Salmonella oranienberg* | | A0A3V8Z2E2 | -- | 30286803 | Not Available | Not Available |
| 1058 | Poly-beta-1, 6-N-acetyl-D-glucosamine synthase (Poly-beta-1, 6-GlcNAc synthase) (EC 2.4.1.-) | *Acinetobacter ursingii* | | A0A3F3LB10 | -- | 28293292 | Acetyl glucosaminyl transferase activity | Not Available |
| 1059 | Biofilm PGA synthesis deacetylase PgaB (Poly-beta-1, 6-N-acetyl-D-glucosamine N-deacetylase PgaB) | *Acinetobacter ursingii* | | A0A3F3L5D6 | -- | 28293292 | Hydrolase activity, acting on carbon-nitrogen (but not peptide) bonds | Not Available |
| 1060 | Surface composition regulator | *Salmonella montevideo* | | A0A3A3J9M4 | -- | 30286803; 31527278 | Not Available | Not Available |
| 1061 | Poly-beta-1, 6-N-acetyl-D-glucosamine synthase (Poly-beta-1, 6-GlcNAc synthase) (EC 2.4.1.-) | *Klebsiella pneumoniae* | | A0A0W8A3S5 | -- | 25341126; 30373801; 32429555 | Acetyl glucosaminyl transferase activity | Not Available |
| 1062 | Poly-beta-1, 6-N-acetyl-D-glucosamine synthase (Poly-beta-1, 6-GlcNAc synthase) (EC 2.4.1.-) | *Acinetobacter baumannii* | | V5VBU0 | -- | 27917170; 29137647; 30124845 | Acetyl glucosaminyl transferase activity | Not Available |
| 1063 | Poly-beta-1, 6-N-acetyl-D-glucosamine synthase (Poly-beta-1, 6-GlcNAc synthase) (EC 2.4.1.-) | *Citrobacter pasteurii* | | A0A6N6JYD6 | -- | 30087402 | Acetyl glucosaminyl transferase activity | Not Available |
| 1064 | Poly-beta-1, 6-N-acetyl-D-glucosamine synthase (Poly-beta-1, 6-GlcNAc synthase) (EC 2.4.1.-) | *Acinetobacter baumannii* | | A0A059ZRE0 | -- | 25136007; 25845845; 26170289; 27917170; 30124845; 29437920 | Acetyl glucosaminyl transferase activity | Not Available |
| 1065 | PgaD (Poly-beta-1, 6-N-acetyl-D-glucosamine biosynthesis protein PgaD) | *Acinetobacter baumannii* | | C8YYH8 | -- | 19633088 | Not Available | Not Available |
| 1066 | Biofilm PGA synthesis protein pgaD (Poly-beta-1, 6-N-acetyl-D-glucosamine biosynthesis protein PgaD) | *Acinetobacter baumannii* | | A0A0E1JKH0 | -- | 25845845; 27917170 | Not Available | Not Available |
| 1067 | Biofilm synthesis deacetylase PgaB (Lipoprotein HmsF) (EC 3.5.1.-) (Poly-beta-1, 6-N-acetyl-D-glucosamine N-deacetylase PgaB) | *Acinetobacter baumannii* | | V5VFW3 | -- | 30124845; 29437920 | Hydrolase activity, acting on carbon-nitrogen (but not peptide) bonds | Not Available |
| 1068 | Poly-beta-1, 6-N-acetyl-D-glucosamine synthase (Poly-beta-1, 6-GlcNAc synthase) (EC 2.4.1.-) (Fragment) | *Citrobacter freundii* | | V5YMI7 | -- | 28636609; 30286803; 30087402; 32766684 | Acetyl glucosaminyl transferase activity | Not Available |
| 1069 | Hemin storage system protein (Poly-beta-1, 6-N-acetyl-D-glucosamine biosynthesis protein PgaD) | *Enterobacter hormaechei* | | A0A156GTQ3 | -- | 29088362 | Not Available | Not Available |
| 1070 | DNA invertase Pin homolog (Recombinae family protein) (Resolvase) | *Escherichia coli* | | B0ZE07 | -- | 15328093; 18440636; 19717626; 31642700 | DNA binding, DNA strand exchange activity | Not Available |
| 1071 | InsB (Insertion element protein) (Transposase) (Transposase (Identified by ISEscan HMM)) | *Escherichia coli* | | B0LHG4 | -- | 18055599; 19717626; 26488485 | Not Available | Not Available |
| 1072 | Conjugal transfer protein (Killer protein) (TrbM-domain protein) | *Escherichia coli* | | B0ZDX4 | -- | 15328093; 18440636; 19717626; 22479389; 31652858 | Not Available | Not Available |
| 1073 | Poly-beta-1, 6-N-acetyl-D-glucosamine synthase (Poly-beta-1, 6-GlcNAc synthase) (EC 2.4.1.-) | *Escherichia coli* | | A0A366YGF8 | -- | 30286803 | Acetyl glucosaminyl transferase activity | Not Available |
| 1074 | MrkD fimbrial protein (Type 3 fimbria adhesin subunit MrkD) | *Escherichia coli* | | B0ZDW1 | -- | 15328093; 18440636 | Not Available | Not Available |
| 1075 | Uncharacterized protein | *Klebsiella pneumoniae* | | Q1G0U8 | -- | 16861646 | Not Available | Not Available |
| 1076 | Poly-beta-1, 6-N-acetyl-D-glucosamine synthase (Poly-beta-1, 6-GlcNAc synthase) (EC 2.4.1.-) | *Escherichia coli* | | A0A7B5TBK4 | -- | 30286803 | Acetyl glucosaminyl transferase activity | Not Available |

| Target ID | Bacterial Protein Name | Organism Name | Strain | UniProt ID | PDB ID | PubMed ID | Classification as per UniProt GO Molecular Function | Classification of Function as per PDB |
|---|---|---|---|---|---|---|---|---|
| 1077 | Fimbriae stability-associated protein (Fimbrial protein) (MrkF) (MrkF fimbrial protein) (MrkF fimbrial protein putative assembly / stability function) (Type 1 fimbrial protein) (Type 3 fimbria minor subunit MrkF) | *Escherichia coli* | | B0LHG9 | -- | 15328093; 18055599; 18440636; 19717626; 22479389; 26441869; 30286803; 31652858 | Not Available | Not Available |
| 1078 | Poly-beta-1, 6-N-acetyl-D-glucosamine synthase (Poly-beta-1, 6-GlcNAc synthase) (EC 2.4.1.-) | *Escherichia coli* | | A0A0D6IFI2 | -- | 28686503; 30286803; 31477907; 32021174 | Acetyl glucosaminyl transferase activity | Not Available |
| 1079 | PGA biosynthesis protein (Poly-beta-1, 6-N-acetyl-D-glucosamine biosynthesis protein PgaD) | *Escherichia coli* | | A0A0J2DWH8 | -- | 28686503; 30286803 | Not Available | Not Available |
| 1080 | Cyanide hydratase (EC 3.1.-.-) (GCU-specific mRNA interferase toxin of the MqsR-MqsA toxin-antitoxin system) (GCU-specific mRNA interferase toxin of the MqsRA toxin-antitoxin system) (Type II toxin-antitoxin system MqsR family toxin) (Type II toxin-antitoxin system toxin MqsR) (EC 3.1.26.-) (mRNA interferase MqsR) | *Escherichia coli* | | J7QHJ7 | -- | 23275093; 28686503; 30286803; 29437920; 31642700; 31652858; 31477907; 30947268; 30948755; 32021174; 32739475; 32599678 | Hydrolase activity | Not Available |
| 1081 | Poly-beta-1, 6-N-acetyl-D-glucosamine biosynthesis protein PgaD | *Chromobacterium haemolyticum* | | A0A2T5P0R2 | -- | 29670144 | Not Available | Not Available |
| 1082 | Poly-beta-1, 6-N-acetyl-D-glucosamine biosynthesis (Poly-beta-1, 6-N-acetyl-D-glucosamine biosynthesis protein PgaD) | *Escherichia coli* | | A0A066SWE3 | -- | 30286803; 31477907 | Not Available | Not Available |
| 1083 | H-HA (Haemolysin expression modulating protein Hha) (Hemolysin activation protein) (Hemolysin expression modulating protein) (Hemolysin expression modulator Hha) (Hha) (Hha expression modulating protein) (Hha expression modulation protein) (Histone-like protein) (Putative hemolysin expression modulating protein) | *Escherichia coli* | | B0ZDX1 | -- | 15328093; 18440636; 19717626; 21831988; 22479389; 23836183; 25403661; 30286803; 31652858; 31477907 | Not Available | Not Available |
| 1084 | Poly-beta-1, 6-N-acetyl-D-glucosamine synthase (Poly-beta-1, 6-GlcNAc synthase) (EC 2.4.1.-) | *Escherichia coli* | | A0A1Y2XXH9 | -- | 30286803 | Acetyl glucosaminyl transferase activity | Not Available |
| 1085 | Surface composition regulator | *Klebsiella pneumoniae* | | A0A080SUH1 | -- | 30373801; 29437920; 32429555 | Not Available | Not Available |
| 1086 | Outer membrane N-deacetylase (Poly-beta-1, 6-N-acetyl-D-glucosamine N-deacetylase PgaB) (EC 3.5.1.-) | *Citrobacter freundii* | | A0A0P8KEL7 | -- | 28636609; 30286803 | Hydrolase activity, acting on carbon-nitrogen (but not peptide) bonds | Not Available |
| 1087 | Poly-beta-1, 6-N-acetyl-D-glucosamine biosynthesis protein PgaD | *Citrobacter freundii* | | A0A0P8I821 | -- | 28636609; 30286803 | Not Available | Not Available |
| 1088 | Fimbria adhesin protein (Fimbrial protein) (MrkD) (MrkD protein) (Type 3 fimbria adhesin subunit MrkD) | *Klebsiella pneumoniae* | | G3FT07 | -- | 21901098; 24123820; 25341126; 30373801; 29437920; 32429555 | Not Available | Not Available |
| 1089 | Hemin storage system protein (Poly-beta-1, 6-N-acetyl-D-glucosamine biosynthesis protein PgaD) | *Klebsiella pneumoniae* | | A0A0C7KL91 | -- | 30373801; 32429555 | Not Available | Not Available |
| 1090 | Surface composition regulator | *Citrobacter freundii* | | A0A133LI55 | -- | 30286803; 30087402; 32766684 | Not Available | Not Available |
| 1091 | Outer membrane N-deacetylase (Poly-beta-1, 6-N-acetyl-D-glucosamine N-deacetylase PgaB) (EC 3.5.1.-) | *Citrobacter freundii* | | A0A0D7M2B6 | -- | 30087402; 32766684 | Hydrolase activity, acting on carbon-nitrogen (but not peptide) bonds | Not Available |
| 1092 | Ferrous iron transporter B (Fimbrial protein) (MrkA) (MrkA protein) (Type 1 fimbrial protein) (Type 3 fimbria major subunit MrkA) | *Klebsiella pneumoniae* | | G3FT04 | -- | 21901098; 25341126; 30373801; 29437920; 32429555 | Not Available | Not Available |
| 1093 | Surface composition regulator | *Salmonella thompson* | | A0A403SPK3 | -- | 30286803 | Not Available | Not Available |
| 1094 | Poly-beta-1, 6-N-acetyl-D-glucosamine biosynthesis protein PgaD | *Acinetobacter ursingii* | | A0A3D2SIV6 | -- | 30148503 | Not Available | Not Available |
| 1095 | Autoinducer 2 import system permease LsrD (Autoinducer-2 ABC transporter permease LsrD) (Autoinducer-2 ABC transporter permease protein) | *Klebsiella pneumoniae* | | C8BNA7 | -- | 20600864; 30373801; 32429555 | Transmembrane transporter activity | Not Available |
| 1096 | Surface composition regulator | *Klebsiella pneumoniae* | | W9B3F6 | -- | 25341126 | Not Available | Not Available |
| 1097 | Surface composition regulator | *Salmonella enterica subspp. enterica serovar* | Kentucky | A0A2T8L9D1 | -- | 30286803; 31715658; 31527278 | Not Available | Not Available |
| 1098 | Surface composition regulator | *Salmonella enterica subspp. enterica serovar* | 4, [5], 12:i:- | A0A315I6K6 | -- | 30286803 | Not Available | Not Available |
| 1099 | AI-2 transport protein TqsA (AI-2E family transporter) (PerM family permease) (Permease) (Putative transport protein) | *Klebsiella pneumoniae* | | C8BNA8 | -- | 20600864; 30373801; 29437920; 32429555 | Not Available | Not Available |
| 1100 | Surface composition regulator | *Citrobacter freundii* | | A0A0A5R3I1 | -- | 28636609 | Not Available | Not Available |
| 1101 | Surface composition regulator | *Salmonella senftenberg* | | A0A3R0HNH3 | -- | 30286803 | Not Available | Not Available |
| 1102 | Biofilm PGA synthesis auxiliary protein PgaD (Biofilm PGA synthesis protein PgaD) (Hemin storage protein) (Poly-beta-1, 6-N-acetyl-D-glucosamine biosynthesis protein PgaD) | *Citrobacter freundii* | | A0A0D7LKL1 | -- | 30286803; 30087402; 32766684 | Not Available | Not Available |
| 1103 | Outer membrane N-deacetylase (Poly-beta-1, 6-N-acetyl-D-glucosamine N-deacetylase) (EC 3.5.1.-) (Poly-beta-1, 6-N-acetyl-D-glucosamine N-deacetylase PgaB) | *Escherichia coli* | | A0A0B0VSG0 | -- | 29437920 | Hydrolase activity, acting on carbon-nitrogen (but not peptide) bonds | Not Available |
| 1104 | Surface composition regulator | *Escherichia coli* | | A0A0K4QSN3 | -- | 30286803 | Not Available | Not Available |
| 1105 | Biofilm PGA synthesis protein PgaD (Poly-beta-1, 6-N-acetyl-D-glucosamine biosynthesis protein PgaD) | *Escherichia coli* | | A0A2X5E7W5 | -- | 30286803 | Not Available | Not Available |

| Target ID | Bacterial Protein Name | Organism Name | Strain | UniProt ID | PDB ID | PubMed ID | Classification as per UniProt GO Molecular Function | Classification of Function as per PDB |
|---|---|---|---|---|---|---|---|---|
| 1106 | Lipoprotein YcdR (Outer membrane N-deacetylase) (Poly-beta-1, 6-N-acetyl-D-glucosamine (PGA) N-deacetylase outer membrane export lipoprotein) (Poly-beta-1, 6-N-acetyl-D-glucosamine N-deacetylase) (Poly-beta-1, 6-N-acetyl-D-glucosamine N-deacetylase PgaB) (EC 3.5.1.-) (Poly-beta-1, 6-N-acetyl-D-glucosamine N-deacetylase outer membrane export lipoprotein) | *Escherichia coli* | | J7QZQ0 | -- | 23275093; 26441869; 30286803; 30924410; 31642700; 31652858; 31477907; 30948755; 32739475; 28953937 | Hydrolase activity, acting on carbon-nitrogen (but not peptide) bonds | Not Available |
| 1107 | Poly-beta-1, 6-N-acetyl-D-glucosamine N-deacetylase PgaB (EC 3.5.1.-) | *Escherichia coli* | | A0A061KYR7 | -- | 30286803; 31477907 | Hydrolase activity, acting on carbon-nitrogen (but not peptide) bonds | Not Available |
| 1108 | Conjugal transfer protein (Hypothetical lipoprotein) (Putative lipoprotein) | *Escherichia coli* | | B0ZDX5 | -- | 15328093; 18440636; 19717626; 22479389 | Not Available | Not Available |
| 1109 | Poly-beta-1, 6-N-acetyl-D-glucosamine N-deacetylase PgaB (EC 3.5.1.-) | *Escherichia coli* | | A0A3L1M483 | -- | 30286803 | Hydrolase activity, acting on carbon-nitrogen (but not peptide) bonds | Not Available |
| 1110 | Lipoprotein YcdR (EC 3.5.1.-) (Poly-beta-1, 6-N-acetyl-D-glucosamine N-deacetylase) (Poly-beta-1, 6-N-acetyl-D-glucosamine N-deacetylase PgaB) | *Escherichia coli* | | C3TEU9 | -- | 19439656 | Hydrolase activity, acting on carbon-nitrogen (but not peptide) bonds | Not Available |
| 1111 | Surface composition regulator | *Escherichia coli* | | A0A2X5FFT5 | -- | 30286803 | Not Available | Not Available |
| 1112 | Putative methyl-accepting chemotaxis protein | *Salmonella typhimurium* | Strain LT2 / SGSC1412 / ATCC 700720 | Q8ZPW4 | -- | 11677609 | Diguanylate cyclase activity | Not Available |
| 1113 | Poly-beta-1, 6-N-acetyl-D-glucosamine synthase (Poly-beta-1, 6-GlcNAc synthase) (EC 2.4.1.-) | *Acinetobacter baumannii* | Strain AB307-0294 | A0A5K6CTW7 | -- | 18931120 | Acetyl glucosaminyl transferase activity | Not Available |
| 1114 | Putative signaling protein | *Clostridium botulinum* | Strain Hall / ATCC 3502 / NCTC 13319 / Type A | A5HXY9 | -- | 17519437 | Diguanylate cyclase activity | Not Available |
| 1115 | Fimbrial subunit | *Salmonella typhimurium* | Strain LT2 / SGSC1412 / ATCC 700720 | Q8ZS05 | -- | 11677609 | Not Available | Not Available |
| 1116 | Putative signaling protein | *Clostridium botulinum* | Strain Hall / ATCC 3502 / NCTC 13319 / Type A | A5HYM7 | -- | 17519437 | Diguanylate cyclase activity | Not Available |
| 1117 | Putative diguanylate cyclase / phosphodiesterase domain 1 | *Salmonella typhimurium* | Strain LT2 / SGSC1412 / ATCC 700720 | H9L4B6 | -- | 11677609 | Diguanylate cyclase activity | Not Available |
| 1118 | Putative signaling protein | *Clostridium botulinum* | Strain Hall / ATCC 3502 / NCTC 13319 / Type A | A5I1K0 | -- | 17519437 | Diguanylate cyclase activity | Not Available |
| 1119 | GGDEF family protein | *Deinococcus radiodurans* | Strain ATCC 13939 / DSM 20539 / JCM 16871 / LMG 4051 / NBRC 15346 / NCIMB 9279 / R1 / VKM B-1422 | Q9RYX3 | -- | 10567266 | Diguanylate cyclase activity, phosphorelay sensor kinase activity | Not Available |
| 1120 | Putative fimbrial chaperone protein | *Salmonella typhimurium* | Strain LT2 / SGSC1412 / ATCC 700720 | Q8ZJT9 | -- | 11677609 | Not Available | Not Available |
| 1121 | Putative outer Membrane protein | *Salmonella typhimurium* | Strain LT2 / SGSC1412 / ATCC 700720 | Q8ZQ03 | -- | 11677609 | Not Available | Not Available |
| 1122 | Putative signaling protein | *Clostridium botulinum* | Strain Hall / ATCC 3502 / NCTC 13319 / Type A | A5I786 | -- | 17519437 | Diguanylate cyclase activity | Not Available |
| 1123 | Putative diguanylate cyclase / phosphodiesterase | *Salmonella typhimurium* | Strain LT2 / SGSC1412 / ATCC 700720 | Q8ZMW4 | -- | 11677609 | Diguanylate cyclase activity | Not Available |
| 1124 | Putative diguanylate cyclase / phosphodiesterase domain 1 containing protein | *Salmonella typhimurium* | Strain LT2 / SGSC1412 / ATCC 700720 | Q8ZJW8 | -- | 11677609 | Diguanylate cyclase activity | Not Available |
| 1125 | Uncharacterized protein | *Acinetobacter baumannii* | Strain AB307-0294 | A0A5K6CUI0 | -- | 18931120 | Not Available | Not Available |
| 1126 | Putative fimbrial subunit | *Salmonella typhimurium* | Strain LT2 / SGSC1412 / ATCC 700720 | H9L482 | -- | 11677609 | Not Available | Not Available |
| 1127 | Fimbrial subunit | *Salmonella typhimurium* | Strain LT2 / SGSC1412 / ATCC 700720 | H9L4A8 | -- | 11677609 | Not Available | Not Available |
| 1128 | Putative signaling protein | *Clostridium botulinum* | Strain Hall / ATCC 3502 / NCTC 13319 / Type A | A5I1D2 | -- | 17519437 | Diguanylate cyclase activity | Not Available |
| 1129 | Biofilm PGA synthesis protein PgaD | *Acinetobacter baumannii* | Strain AB307-0294 | A0A5K6CPI1 | -- | 18931120 | Not Available | Not Available |
| 1130 | Putative fimbrial subunit | *Salmonella typhimurium* | Strain LT2 / SGSC1412 / ATCC 700720 | Q8ZJU2 | -- | 11677609 | Not Available | Not Available |
| 1131 | Poly-beta-1, 6-N-acetyl-D-glucosamine N-deacetylase (EC 3.5.1.-) | *Acinetobacter baumannii* | Strain AB307-0294 | A0A5K6CU40 | -- | 18931120 | Hydrolase activity, acting on carbon-nitrogen (but not peptide) bonds | Not Available |
| 1132 | GGDEF family protein | *Deinococcus radiodurans* | Strain ATCC 13939 / DSM 20539 / JCM 16871 / LMG 4051 / NBRC 15346 / NCIMB 9279 / R1 / VKM B-1422 | Q9RVD9 | -- | 10567266 | Diguanylate cyclase activity, phosphorelay sensor kinase activity | Not Available |
| 1133 | Fimbrial subunit | *Salmonella typhimurium* | Strain LT2 / SGSC1412 / ATCC 700720 | H9L408 | -- | 11677609 | Not Available | Not Available |
| 1134 | Putative minor fimbrial subunit | *Salmonella typhimurium* | Strain LT2 / SGSC1412 / ATCC 700720 | H9L4A9 | -- | 11677609 | Not Available | Not Available |
| 1135 | Putative fimbrial subunit | *Salmonella typhimurium* | Strain LT2 / SGSC1412 / ATCC 700720 | Q8ZRR4 | -- | 11677609 | Not Available | Not Available |
| 1136 | Putative fimbriae | *Salmonella typhimurium* | Strain LT2 / SGSC1412 / ATCC 700720 | Q8ZRR7 | -- | 11677609 | Not Available | Not Available |
| 1137 | Putative signaling protein | *Clostridium botulinum* | Strain Hall / ATCC 3502 / NCTC 13319 / Type A | A5I3D2 | -- | 17519437 | Diguanylate cyclase activity | Not Available |
| 1138 | Poly-beta-1, 6-N-acetyl-D-glucosamine synthase (Poly-beta-1, 6-GlcNAc synthase) (EC 2.4.1.-) | *Photobacterium profundum* | Strain SS9 | Q6LK51 | -- | 15746425 | Acetyl glucosaminyl transferase activity | Not Available |
| 1139 | Putative minor fimbrial subunit | *Salmonella typhimurium* | Strain LT2 / SGSC1412 / ATCC 700720 | Q8ZRP9 | -- | 11677609 | Not Available | Not Available |

| Target ID | Bacterial Protein Name | Organism Name | Strain | UniProt ID | PDB ID | PubMed ID | Classification as per UniProt GO Molecular Function | Classification of Function as per PDB |
|---|---|---|---|---|---|---|---|---|
| 1140 | GGDEF family protein | *Deinococcus radiodurans* | Strain ATCC 13939 / DSM 20539 / JCM 16871 / LMG 4051 / NBRC 15346 / NCIMB 9279 / R1 / VKM B-1422 | Q9RWR1 | -- | 10567266 | Diguanylate cyclase activity | Not Available |
| 1141 | NodB homology domain-containing protein | *Photobacterium profundum* | Strain SS9 | Q6LK50 | -- | 15746425 | Hydrolase activity, acting on carbon-nitrogen (but not peptide) bonds | Not Available |
| 1142 | Polysaccharide deacetylase | *Halomonas spp. TD01* | | F7SRR6 | -- | 22040376 | Hydrolase activity, acting on carbon-nitrogen (but not peptide) bonds | Not Available |
| 1143 | Diguanylate cyclase with PAS / PAC sensor | *Dictyoglomus turgidum* | Strain DSM 6724 / Z-1310 | B8DYG5 | -- | 28066333 | Diguanylate cyclase activity | Not Available |
| 1144 | Type II toxin-antitoxin system MqsR family toxin | *Desulfuromonadales bacterium* | | A0A7C8ARH6 | -- | 32737307 | Not Available | Not Available |
| 1145 | Uncharacterized protein | *Proteobacteria bacterium SG_bin4* | | A0A1W9GZE0 | -- | 28390307 | Not Available | Not Available |
| 1146 | Diguanylate cyclase | *Dictyoglomus turgidum* | Strain DSM 6724 / Z-1310 | B8E0L1 | -- | 28066333 | Diguanylate cyclase activity | Not Available |
| 1147 | Diguanylate cyclase | *Dictyoglomus turgidum* | Strain DSM 6724 / Z-1310 | B8E1P2 | -- | 28066333 | Diguanylate cyclase activity | Not Available |
| 1148 | Poly-beta-1, 6-N-acetyl-D-glucosamine synthase (Poly-beta-1, 6-GlcNAc synthase) (EC 2.4.1.-) | *Klebsiella pneumoniae subspp. rhinoscleromatis* | SB3432 | R4Y4T4 | -- | 21629720 | Acetyl glucosaminyl transferase activity | Not Available |
| 1149 | Diguanylate cyclase | *Dictyoglomus turgidum* | Strain DSM 6724 / Z-1310 | B8DZI2 | -- | 28066333 | Diguanylate cyclase activity | Not Available |
| 1150 | Diguanylate cyclase | *Dictyoglomus turgidum* | Strain DSM 6724 / Z-1310 | B8E392 | -- | 28066333 | Diguanylate cyclase activity | Not Available |
| 1151 | Poly-beta-1, 6-N-acetyl-D-glucosamine synthase (Poly-beta-1, 6-GlcNAc synthase) (EC 2.4.1.-) | *Romboutsia weinsteinii* | | A0A371J6B9 | -- | 28982987 | Acetyl glucosaminyl transferase activity | Not Available |
| 1152 | Uncharacterized protein | *Rhizobium freirei PRF 81* | | N6UY50 | -- | 23270491 | Not Available | Not Available |
| 1153 | Uncharacterized protein | *Novosphingobium barchaimii LL02* | | A0A0J7XQU0 | -- | 25850427 | Not Available | Not Available |
| 1154 | Poly-beta-1, 6-N-acetyl-D-glucosamine biosynthesis protein PgaD | *Elusimicrobia bacterium RIFOXYA2_FULL_50_26* | | A0A1F9WSX0 | -- | 27774985 | Not Available | Not Available |
| 1155 | Uncharacterized protein | *Pseudomonas spp. GM21* | | J3EX79 | -- | 23045501 | Not Available | Not Available |
| 1156 | Poly-beta-1, 6-N-acetyl-D-glucosamine N-deacetylase PgaB | *Escherichia spp.* | | A0A352MKB2 | -- | 30148503 | Hydrolase activity, acting on carbon-nitrogen (but not peptide) bonds | Not Available |
| 1157 | Poly-beta-1, 6-N-acetyl-D-glucosamine synthase (Poly-beta-1, 6-GlcNAc synthase) (EC 2.4.1.-) | *Elusimicrobia bacterium RIFOXYA2_FULL_50_26* | | A0A1F9WSW2 | -- | 27774985 | Acetyl glucosaminyl transferase activity | Not Available |
| 1158 | Poly-beta-1, 6-N-acetyl-D-glucosamine synthase (Poly-beta-1, 6-GlcNAc synthase) (EC 2.4.1.-) | *Escherichia spp.* | | A0A352MKB1 | -- | 30148503 | Acetyl glucosaminyl transferase activity | Not Available |
| 1159 | Poly-beta-1, 6-N-acetyl-D-glucosamine biosynthesis protein PgaD | *Escherichia spp.* | | A0A352MKB0 | -- | 30148503 | Not Available | Not Available |
| 1160 | Uncharacterized protein | *Geobacteraceae bacterium GWC2_58_44* | | A0A1G0MDW5 | -- | 27774985 | Not Available | Not Available |
| 1161 | Lipoprotein for biofilm formation | *Bacillus atrophaeus* | Strain 1942 | A0A0H3E3Y9 | -- | 21464989 | Not Available | Not Available |
| 1162 | Surface composition regulator | *Escherichia coli O111:H11 str. CVM9455* | | K4WSM6 | -- | 23052305 | Not Available | Not Available |
| 1163 | Poly-beta-1, 6-N-acetyl-D-glucosamine N-deacetylase PgaB | *Pectobacterium polonicum* | | A0A4V6ESD3 | -- | 30973319 | Hydrolase activity, acting on carbon-nitrogen (but not peptide) bonds | Not Available |
| 1164 | Type II toxin-antitoxin system MqsR family toxin | *Escherichia coli* | | A0A6M0Q153 | -- | 16943019 | Not Available | Not Available |
| 1165 | Poly-beta-1, 6-N-acetyl-D-glucosamine N-deacetylase PgaB | *Escherichia coli* | | A0A7B6D4F3 | -- | 30286803 | Hydrolase activity, acting on carbon-nitrogen (but not peptide) bonds | Not Available |
| 1166 | Poly-beta-1, 6-N-acetyl-D-glucosamine biosynthesis protein PgaD | *Escherichia coli* | | A0A792HNV3 | -- | 30286803 | Not Available | Not Available |
| 1167 | Uncharacterized protein | *Desulfuromonadales bacterium GWC2_61_20* | | A0A1F9PX82 | -- | 27774985 | Not Available | Not Available |
| 1168 | Predicted enzyme associated with biofilm formation | *Escherichia coli O1:K1 / APEC* | | A0A0H2YXW7 | -- | 17293413 | Hydrolase activity, acting on carbon-nitrogen (but not peptide) bonds | Not Available |
| 1169 | Predicted inner Membrane protein | *Escherichia coli O1:K1 / APEC* | | A0A0H2YXK2 | -- | 17293413 | Not Available | Not Available |
| 1170 | Hemin storage protein | *Stenotrophomonas maltophilia MF89* | | T5K7H5 | -- | 24092793 | Hydrolase activity, acting on carbon-nitrogen (but not peptide) bonds | Not Available |
| 1171 | Poly-beta-1, 6-N-acetyl-D-glucosamine synthase (Poly-beta-1, 6-GlcNAc synthase) (EC 2.4.1.-) | *Stenotrophomonas maltophilia MF89* | | T5KI40 | -- | 24092793 | Acetyl glucosaminyl transferase activity | Not Available |
| 1172 | Type II toxin-antitoxin system MqsR family toxin | *Herbaspirillum frisingense* | | A0A7G6B4A5 | -- | 32414048 | Not Available | Not Available |
| 1173 | Putative hemin storage protein | *Stenotrophomonas maltophilia* | Strain K279a | B2FMD0 | -- | 18419807 | Hydrolase activity, acting on carbon-nitrogen (but not peptide) bonds | Not Available |
| 1174 | Poly-beta-1, 6-N-acetyl-D-glucosamine synthase (Poly-beta-1, 6-GlcNAc synthase) (EC 2.4.1.-) | *Aggregatibacter actinomycetemcomitans RhAA1* | | H0KC41 | -- | 22328766 | Acetyl glucosaminyl transferase activity | Not Available |
| 1175 | Putative toxin-antitoxin system, toxin component | *Burkholderia multivorans CF2* | | J4RFZ0 | -- | 23105085 | Not Available | Not Available |
| 1176 | Poly-beta-1, 6-N-acetyl-D-glucosamine biosynthesis protein PgaD | *Pseudomonas spp. WS 5406* | | A0A7Y0VDH4 | -- | 32587583 | Not Available | Not Available |
| 1177 | Poly-beta-1, 6-N-acetyl-D-glucosamine synthase (Poly-beta-1, 6-GlcNAc synthase) (EC 2.4.1.-) | *Pseudomonas spp. WS 5406* | | A0A7Y0VCZ0 | -- | 32587583 | Acetyl glucosaminyl transferase activity | Not Available |
| 1178 | Poly-beta-1, 6-N-acetyl-D-glucosamine synthase (Poly-beta-1, 6-GlcNAc synthase) (EC 2.4.1.-) | *Acidithiobacillales bacterium SM1_46* | | A0A0S8L3M3 | -- | 25922666 | Acetyl glucosaminyl transferase activity | Not Available |
| 1179 | Poly-beta-1, 6-N-acetyl-D-glucosamine N-deacetylase PgaB (EC 3.5.1.-) | *Burkholderia multivorans CF2* | | J4RKZ9 | -- | 23105085 | Hydrolase activity, acting on carbon-nitrogen (but not peptide) bonds | Not Available |
| 1180 | Putative haemin storage system, HmsS transmembrane protein | *Stenotrophomonas maltophilia* | Strain K279a | B2FMD2 | -- | 18419807 | Not Available | Not Available |
| 1181 | Uncharacterized protein | *Acidithiobacillales bacterium SM1_46* | | A0A0S8L209 | -- | 25922666 | Not Available | Not Available |
| 1182 | Poly-beta-1, 6-N-acetyl-D-glucosamine synthase (Poly-beta-1, 6-GlcNAc synthase) (EC 2.4.1.-) | *Burkholderia multivorans CF2* | | J4RKZ2 | -- | 23105085 | Acetyl glucosaminyl transferase activity | Not Available |
| 1183 | Uncharacterized protein | *Weissella cibaria* | | A0A0D1LW17 | -- | 25678547 | Not Available | Not Available |
| 1184 | Poly-beta-1, 6-N-acetyl-D-glucosamine synthase (Poly-beta-1, 6-GlcNAc synthase) (EC 2.4.1.-) | *Stenotrophomonas maltophilia* | Strain K279a | B2FMD1 | -- | 18419807 | Acetyl glucosaminyl transferase activity | Not Available |
| 1185 | Motility quorum-sensing regulator / GCU-specific mRNA interferase toxin | *Lysobacter ruishenii* | | A0A562LCQ9 | -- | 26203337 | Not Available | Not Available |
| 1186 | Uncharacterized protein | *Castellaniella defragrans 65Phen* | | W8X1U6 | -- | 24952578 | Not Available | Not Available |
| 1187 | Biofilm PGA synthesis deacetylase PgaB | *Castellaniella defragrans 65Phen* | | W8X5T9 | -- | 24952578 | Hydrolase activity, acting on carbon-nitrogen (but not peptide) bonds | Not Available |
| 1188 | Poly-beta-1, 6-N-acetyl-D-glucosamine biosynthesis protein PgaD | *Pseudomonas nabeulensis* | | A0A4Z0ANK2 | -- | 30910423 | Not Available | Not Available |

| Target ID | Bacterial Protein Name | Organism Name | Strain | UniProt ID | PDB ID | PubMed ID | Classification as per UniProt GO Molecular Function | Classification of Function as per PDB |
|---|---|---|---|---|---|---|---|---|
| 1189 | Poly-beta-1, 6-N-acetyl-D-glucosamine synthase (Poly-beta-1, 6-GlcNAc synthase) (EC 2.4.1.-) | *Castellaniella defragrans 65Phen* | | W8X1E8 | -- | 24952578 | Acetyl glucosaminyl transferase activity | Not Available |
| 1190 | Poly-beta-1, 6-N-acetyl-D-glucosamine synthase (Poly-beta-1, 6-GlcNAc synthase) (EC 2.4.1.-) | *Pseudomonas nabeulensis* | | A0A4Z0APQ2 | -- | 30910423 | Acetyl glucosaminyl transferase activity | Not Available |
| 1191 | Uncharacterized protein | *Sulfuricurvum spp. PC08-66* | | A0A0C2ZZX5 | -- | 25620962 | Not Available | Not Available |
| 1192 | Poly-beta-1, 6-N-acetyl-D-glucosamine N-deacetylase PgaB (EC 3.5.1.-) | *Pseudomonas spp. WS 5406* | | A0A7Y0VCY5 | -- | 32587583 | Hydrolase activity, acting on carbon-nitrogen (but not peptide) bonds | Not Available |
| 1193 | Uncharacterized protein | *Furfurilactobacillus rossiae DSM 15814* | | A0A0R1RIK6 | -- | 26415554 | Not Available | Not Available |
| 1194 | Poly-beta-1, 6-N-acetyl-D-glucosamine synthase (Poly-beta-1, 6-GlcNAc synthase) (EC 2.4.1.-) | *Methylotenera versatilis* | Strain 301 | D7DJ80 | -- | 21622745 | Acetyl glucosaminyl transferase activity | Not Available |
| 1195 | GGDEF family protein | *Xanthomonas campestris pv. campestris* | Strain ATCC 33913 / DSM 3586 / NCPPB 528 / LMG 568 / P 25 | Q8P4E9 | -- | 12024217 | Diguanylate cyclase activity | Not Available |
| 1196 | Poly-beta-1, 6-N-acetyl-D-glucosamine synthase (Poly-beta-1, 6-GlcNAc synthase) (EC 2.4.1.-) | *Sulfuricurvum kujiense* | Strain ATCC BAA-921 / DSM 16994 / JCM 11577 / YK-1 | E4U030 | -- | 22675602 | Acetyl glucosaminyl transferase activity | Not Available |
| 1197 | Uncharacterized protein | *Methylotenera versatilis* | Strain 301 | D7DK49 | -- | 21622745 | Not Available | Not Available |
| 1198 | Transcriptional regulator | *Xanthomonas campestris pv. campestris* | Strain ATCC 33913 / DSM 3586 / NCPPB 528 / LMG 568 / P 25 | Q8P893 | -- | 12024217 | Diguanylate cyclase activity | Not Available |
| 1199 | GGDEF domain-containing protein | *Xanthomonas campestris pv. campestris* | Strain ATCC 33913 / DSM 3586 / NCPPB 528 / LMG 568 / P 25 | Q8PAG4 | -- | 12024217 | Diguanylate cyclase activity | Not Available |
| 1200 | Response regulator | *Xanthomonas campestris pv. campestris* | Strain ATCC 33913 / DSM 3586 / NCPPB 528 / LMG 568 / P 25 | Q8P782 | -- | 12024217 | Diguanylate cyclase activity | Not Available |
| 1201 | GGDEF domain-containing protein | *Xanthomonas campestris pv. campestris* | Strain ATCC 33913 / DSM 3586 / NCPPB 528 / LMG 568 / P 25 | Q8P5K7 | -- | 12024217 | Diguanylate cyclase activity | Not Available |
| 1202 | Uncharacterized protein | *Xanthomonas campestris pv. campestris* | Strain ATCC 33913 / DSM 3586 / NCPPB 528 / LMG 568 / P 25 | Q8PDU6 | -- | 12024217 | Diguanylate cyclase activity | Not Available |
| 1203 | Polysaccharide deacetylase | *Methylotenera versatilis* | Strain 301 | D7DK51 | -- | 21622745 | Hydrolase activity, acting on carbon-nitrogen (but not peptide) bonds | Not Available |
| 1204 | GGDEF family protein | *Xanthomonas campestris pv. campestris* | Strain ATCC 33913 / DSM 3586 / NCPPB 528 / LMG 568 / P 25 | Q8P9F3 | -- | 12024217 | Diguanylate cyclase activity | Not Available |
| 1205 | Motility quorum-sensing regulator MqsR | *Hyphomonaceae bacterium* | | A0A3D2W0H9 | -- | 30148503 | Not Available | Not Available |
| 1206 | Poly-beta-1, 6-N-acetyl-D-glucosamine synthase (Poly-beta-1, 6-GlcNAc synthase) (EC 2.4.1.-) | *Stenotrophomonas spp.* | | A0A357N2M6 | -- | 30148503 | Acetyl glucosaminyl transferase activity | Not Available |
| 1207 | Poly-beta-1, 6-N-acetyl-D-glucosamine biosynthesis protein PgaD | *Stenotrophomonas spp.* | | A0A357N2K2 | -- | 30148503 | Not Available | Not Available |
| 1208 | Poly-beta-1, 6-N-acetyl-D-glucosamine N-deacetylase PgaB | *Stenotrophomonas spp.* | | A0A356JDF6 | -- | 30148503 | Hydrolase activity, acting on carbon-nitrogen (but not peptide) bonds | Not Available |
| 1209 | Poly-beta-1, 6-N-acetyl-D-glucosamine N-deacetylase PgaB | *Stenotrophomonas spp.* | | A0A3D0GBI1 | -- | 30148503 | Hydrolase activity, acting on carbon-nitrogen (but not peptide) bonds | Not Available |
| 1210 | Poly-beta-1, 6-N-acetyl-D-glucosamine N-deacetylase PgaB | *Stenotrophomonas spp.* | | A0A357N2H1 | -- | 30148503 | Hydrolase activity, acting on carbon-nitrogen (but not peptide) bonds | Not Available |
| 1211 | Poly-beta-1, 6-N-acetyl-D-glucosamine N-deacetylase PgaB | *Stenotrophomonas spp.* | | A0A3D4THG8 | -- | 30148503 | Hydrolase activity, acting on carbon-nitrogen (but not peptide) bonds | Not Available |
| 1212 | Poly-beta-1, 6-N-acetyl-D-glucosamine synthase (Poly-beta-1, 6-GlcNAc synthase) (EC 2.4.1.-) | *Stenotrophomonas spp.* | | A0A3D0GBN0 | -- | 30148503 | Acetyl glucosaminyl transferase activity | Not Available |
| 1213 | Outer membrane N-deacetylase | *Escherichia coli O83:H1* | Strain NRG 857C / AIEC | A0A0H3EFY1 | -- | 21108814 | Hydrolase activity, acting on carbon-nitrogen (but not peptide) bonds | Not Available |
| 1214 | Poly-beta-1, 6-N-acetyl-D-glucosamine biosynthesis protein PgaD | *Stenotrophomonas spp.* | | A0A3D4TH94 | -- | 30148503 | Not Available | Not Available |
| 1215 | Surface composition regulator | *Klebsiella pneumoniae 30660 / NJST258_1* | | A0A0E1CCY4 | -- | 24639510 | Not Available | Not Available |
| 1216 | Poly-beta-1, 6-N-acetyl-D-glucosamine N-deacetylase PgaB | *Methylococcaceae bacterium* | | A0A351V598 | -- | 30148503 | Hydrolase activity, acting on carbon-nitrogen (but not peptide) bonds | Not Available |
| 1217 | Biofilm PGA synthesis lipoprotein PgaB | *Acinetobacter spp. HA* | | I4ZWX7 | -- | 22933775 | Hydrolase activity, acting on carbon-nitrogen (but not peptide) bonds | Not Available |
| 1218 | Hemin storage protein | *Stenotrophomonas maltophilia* | Pseudomonas maltophilia Xanthomonas maltophilia | A0A0M0P251 | -- | 26324280 | Hydrolase activity, acting on carbon-nitrogen (but not peptide) bonds | Not Available |
| 1219 | Poly-beta-1, 6-N-acetyl-D-glucosamine synthase (Poly-beta-1, 6-GlcNAc synthase) (EC 2.4.1.-) | *Stenotrophomonas maltophilia* | Pseudomonas maltophilia Xanthomonas maltophilia | A0A0M0P2A8 | -- | 26324280 | Acetyl glucosaminyl transferase activity | Not Available |
| 1220 | Poly-beta-1, 6-N-acetyl-D-glucosamine biosynthesis protein PgaD | *Stenotrophomonas maltophilia* | Pseudomonas maltophilia Xanthomonas maltophilia | A0A2J0SNZ8 | -- | 29170656 | Not Available | Not Available |
| 1221 | Uncharacterized protein | *Acinetobacter spp. HA* | | I4ZWY0 | -- | 22933775 | Not Available | Not Available |
| 1222 | Poly-beta-1, 6-N-acetyl-D-glucosamine synthase (Poly-beta-1, 6-GlcNAc synthase) (EC 2.4.1.-) | *Stenotrophomonas maltophilia* | Pseudomonas maltophilia Xanthomonas maltophilia | A0A2Y9U8M9 | -- | 29170656 | Acetyl glucosaminyl transferase activity | Not Available |
| 1223 | Poly-beta-1, 6-N-acetyl-D-glucosamine N-deacetylase PgaB | *Stenotrophomonas maltophilia* | Pseudomonas maltophilia Xanthomonas maltophilia | A0A2U4HSV1 | -- | 29170656 | Hydrolase activity, acting on carbon-nitrogen (but not peptide) bonds | Not Available |
| 1224 | Poly-beta-1, 6-N-acetyl-D-glucosamine synthase (Poly-beta-1, 6-GlcNAc synthase) (EC 2.4.1.-) | *Stenotrophomonas maltophilia* | Pseudomonas maltophilia Xanthomonas maltophilia | A0A2J0UEV7 | -- | 29170656 | Acetyl glucosaminyl transferase activity | Not Available |
| 1225 | Poly-beta-1, 6-N-acetyl-D-glucosamine synthase (Poly-beta-1, 6-GlcNAc synthase) (EC 2.4.1.-) | *Stenotrophomonas maltophilia* | Pseudomonas maltophilia Xanthomonas maltophilia | A0A2J0TWK6 | -- | 29170656 | Acetyl glucosaminyl transferase activity | Not Available |

| Target ID | Bacterial Protein Name | Organism Name | Strain | UniProt ID | PDB ID | PubMed ID | Classification as per UniProt GO Molecular Function | Classification of Function as per PDB |
|---|---|---|---|---|---|---|---|---|
| 1226 | Poly-beta-1, 6-N-acetyl-D-glucosamine biosynthesis protein PgaD | *Stenotrophomonas maltophilia* | Pseudomonas maltophilia Xanthomonas maltophilia | A0A2J0UEW8 | -- | 29170656 | Not Available | Not Available |
| 1227 | Poly-beta-1, 6-N-acetyl-D-glucosamine N-deacetylase PgaB | *Stenotrophomonas maltophilia* | Pseudomonas maltophilia Xanthomonas maltophilia | A0A2W6M1R9 | -- | 29170656 | Hydrolase activity, acting on carbon-nitrogen (but not peptide) bonds | Not Available |
| 1228 | Poly-beta-1, 6-N-acetyl-D-glucosamine N-deacetylase PgaB | *Stenotrophomonas maltophilia* | Pseudomonas maltophilia Xanthomonas maltophilia | A0A2J0SNW7 | -- | 29170656 | Hydrolase activity, acting on carbon-nitrogen (but not peptide) bonds | Not Available |
| 1229 | Hemin storage protein | *Stenotrophomonas maltophilia* | Pseudomonas maltophilia Xanthomonas maltophilia | A0A0M0P257 | -- | 26324280 | Not Available | Not Available |
| 1230 | Surface composition regulator | *Escherichia coli O44:H18* | Strain 042 / EAEC | D3GVV9 | -- | 20098708 | Not Available | Not Available |
| 1231 | Poly-beta-1, 6-N-acetyl-D-glucosamine synthase (Poly-beta-1, 6-GlcNAc synthase) (EC 2.4.1.-) | *Acinetobacter spp. HA* | | I4ZWX9 | -- | 22933775 | Acetyl glucosaminyl transferase activity | Not Available |
| 1232 | Poly-beta-1, 6-N-acetyl-D-glucosamine synthase (Poly-beta-1, 6-GlcNAc synthase) (EC 2.4.1.-) | *Stenotrophomonas maltophilia* | Pseudomonas maltophilia Xanthomonas maltophilia | A0A2J0SHH8 | -- | 29170656 | Acetyl glucosaminyl transferase activity | Not Available |
| 1233 | Poly-beta-1, 6-N-acetyl-D-glucosamine biosynthesis protein PgaD | *Thauera spp.* | | A0A3B9PL39 | -- | 30148503 | Not Available | Not Available |
| 1234 | Poly-beta-1, 6-N-acetyl-D-glucosamine synthase (Poly-beta-1, 6-GlcNAc synthase) (EC 2.4.1.-) | *Pseudomonas spp. R84* | | A0A7Z2KP33 | -- | 25398872 | Acetyl glucosaminyl transferase activity | Not Available |
| 1235 | Poly-beta-1, 6-N-acetyl-D-glucosamine N-deacetylase PgaB | *Desulfobacteraceae bacterium* | | A0A3A4SC85 | -- | 28644444 | Hydrolase activity, acting on carbon-nitrogen (but not peptide) bonds | Not Available |
| 1236 | Poly-beta-1, 6-N-acetyl-D-glucosamine biosynthesis protein PgaD | *Pseudomonas spp. R84* | | A0A7Z2KNW1 | -- | 25398872 | Not Available | Not Available |
| 1237 | Poly-beta-1, 6-N-acetyl-D-glucosamine synthase (Poly-beta-1, 6-GlcNAc synthase) (EC 2.4.1.-) | *Thauera spp.* | | A0A3B9PL11 | -- | 30148503 | Acetyl glucosaminyl transferase activity | Not Available |
| 1238 | mRNA interferase MqsR | *Paraburkholderia phytofirmans OLGA172* | | A0A160FSJ7 | -- | 27063562 | Not Available | Not Available |
| 1239 | Poly-beta-1, 6-N-acetyl-D-glucosamine N-deacetylase (EC 3.5.1.-) | *Acinetobacter venetianus* | | A0A150HS26 | -- | 26902269 | Hydrolase activity, acting on carbon-nitrogen (but not peptide) bonds | Not Available |
| 1240 | Poly-beta-1, 6-N-acetyl-D-glucosamine N-deacetylase (EC 3.5.1.-) | *Acinetobacter venetianus* | | A0A150HP82 | -- | 26902269 | Hydrolase activity, acting on carbon-nitrogen (but not peptide) bonds | Not Available |
| 1241 | Poly-beta-1, 6-N-acetyl-D-glucosamine biosynthesis protein PgaD | *Desulfobacteraceae bacterium* | | A0A3A4RZP0 | -- | 28644444 | Not Available | Not Available |
| 1242 | Motility quorum-sensing regulator MqsR | *Pseudomonas spp. R84* | | A0A7Z2KQT3 | -- | 25398872 | Not Available | Not Available |
| 1243 | Poly-beta-1, 6-N-acetyl-D-glucosamine N-deacetylase PgaB | *Herbaspirillum spp. YR522* | | J3HWP5 | -- | 23045501 | Hydrolase activity, acting on carbon-nitrogen (but not peptide) bonds | Not Available |
| 1244 | Uncharacterized protein | *Herbaspirillum spp. YR522* | | J2W5K5 | -- | 23045501 | Not Available | Not Available |
| 1245 | Uncharacterized protein | *Acinetobacter venetianus* | | A0A150HP16 | -- | 26902269 | Not Available | Not Available |
| 1246 | Poly-beta-1, 6-N-acetyl-D-glucosamine synthase (Poly-beta-1, 6-GlcNAc synthase) (EC 2.4.1.-) | *Marine gamma proteobacterium HTCC2143* | | A0YHE0 | -- | 20601481 | Acetyl glucosaminyl transferase activity | Not Available |
| 1247 | Poly-beta-1, 6-N-acetyl-D-glucosamine synthase (Poly-beta-1, 6-GlcNAc synthase) (EC 2.4.1.-) | *Herbaspirillum spp. YR522* | | J3CKU0 | -- | 23045501 | Acetyl glucosaminyl transferase activity | Not Available |
| 1248 | Poly-beta-1, 6-N-acetyl-D-glucosamine synthase (Poly-beta-1, 6-GlcNAc synthase) (EC 2.4.1.-) | *Paraburkholderia phytofirmans OLGA172* | | A0A160FI12 | -- | 27063562 | Acetyl glucosaminyl transferase activity | Not Available |
| 1249 | Poly-beta-1, 6-N-acetyl-D-glucosamine N-deacetylase | *Pseudomonas spp. R84* | | A0A7Z2KNV3 | -- | 25398872 | Hydrolase activity, acting on carbon-nitrogen (but not peptide) bonds | Not Available |
| 1250 | Poly-beta-1, 6-N-acetyl-D-glucosamine N-deacetylase PgaB | *Moraxellaceae bacterium* | | A0A4Q5VP81 | -- | 30498029 | Hydrolase activity, acting on carbon-nitrogen (but not peptide) bonds | Not Available |
| 1251 | Poly-beta-1, 6-N-acetyl-D-glucosamine N-deacetylase PgaB (Fragment) | *Moraxellaceae bacterium* | | A0A4Q6DWC4 | -- | 30498029 | Hydrolase activity, acting on carbon-nitrogen (but not peptide) bonds | Not Available |
| 1252 | Poly-beta-1, 6-N-acetyl-D-glucosamine synthase (Poly-beta-1, 6-GlcNAc synthase) (EC 2.4.1.-) | *Moraxellaceae bacterium* | | A0A4Q5VP12 | -- | 30498029 | Acetyl glucosaminyl transferase activity | Not Available |
| 1253 | Poly-beta-1, 6-N-acetyl-D-glucosamine biosynthesis protein PgaD | *Moraxellaceae bacterium* | | A0A4V2AC17 | -- | 30498029 | Not Available | Not Available |
| 1254 | Outer membrane N-deacetylase | *Pseudomonas fluorescens BRIP34879* | | L7HN69 | -- | 23661484 | Hydrolase activity, acting on carbon-nitrogen (but not peptide) bonds | Not Available |
| 1255 | Poly-beta-1, 6-N-acetyl-D-glucosamine synthase (Poly-beta-1, 6-GlcNAc synthase) (EC 2.4.1.-) | *Moraxellaceae bacterium* | | A0A4Q5VNZ1 | -- | 30498029 | Acetyl glucosaminyl transferase activity | Not Available |
| 1256 | Xylanae / chitin deacetylase | *Acinetobacter oleivorans* | Strain JCM 16667 / KCTC 23045 / DR1 | D8JGK9 | -- | 20639327 | Hydrolase activity, acting on carbon-nitrogen (but not peptide) bonds, Hydrolase activity, acting on glycosyl bonds | Not Available |
| 1257 | Biofilm PGA synthesis lipoprotein pgaB | *Acinetobacter oleivorans* | Strain JCM 16667 / KCTC 23045 / DR1 | D8JE04 | -- | 20639327 | Hydrolase activity, acting on carbon-nitrogen (but not peptide) bonds | Not Available |
| 1258 | Poly-beta-1, 6-N-acetyl-D-glucosamine synthase (Poly-beta-1, 6-GlcNAc synthase) (EC 2.4.1.-) | *Acinetobacter oleivorans* | Strain JCM 16667 / KCTC 23045 / DR1 | D8JE03 | -- | 20639327 | Acetyl glucosaminyl transferase activity | Not Available |
| 1259 | Biofilm PGA synthesis protein PgaD, putative | *Pseudomonas fluorescens BRIP34879* | | L7HLG7 | -- | 23661484 | Not Available | Not Available |
| 1260 | Poly-beta-1, 6-N-acetyl-D-glucosamine synthase (Poly-beta-1, 6-GlcNAc synthase) (EC 2.4.1.-) | *Acinetobacter oleivorans* | Strain JCM 16667 / KCTC 23045 / DR1 | D8JGL0 | -- | 20639327 | Acetyl glucosaminyl transferase activity | Not Available |
| 1261 | Poly-beta-1, 6-N-acetyl-D-glucosamine synthase (Poly-beta-1, 6-GlcNAc synthase) (EC 2.4.1.-) | *Pseudomonas fluorescens BRIP34879* | | L7HM73 | -- | 23661484 | Acetyl glucosaminyl transferase activity | Not Available |
| 1262 | Uncharacterized protein | *Acinetobacter oleivorans* | Strain JCM 16667 / KCTC 23045 / DR1 | D8JGL1 | -- | 20639327 | Not Available | Not Available |
| 1263 | Poly-beta-1, 6-N-acetyl-D-glucosamine synthase (Poly-beta-1, 6-GlcNAc synthase) (EC 2.4.1.-) | *Moraxellaceae bacterium* | | A0A4Q6EPP1 | -- | 30498029 | Acetyl glucosaminyl transferase activity | Not Available |
| 1264 | Poly-beta-1, 6-N-acetyl-D-glucosamine biosynthesis protein PgaD | *Moraxellaceae bacterium* | | A0A4Q6DYT8 | -- | 30498029 | Not Available | Not Available |
| 1265 | Poly-beta-1, 6-N-acetyl-D-glucosamine biosynthesis protein PgaD | *Moraxellaceae bacterium* | | A0A4Q5VNZ4 | -- | 30498029 | Not Available | Not Available |
| 1266 | Poly-beta-1, 6-N-acetyl-D-glucosamine synthase (Poly-beta-1, 6-GlcNAc synthase) (EC 2.4.1.-) (Fragment) | *Moraxellaceae bacterium* | | A0A4Q6CZ11 | -- | 30498029 | Acetyl glucosaminyl transferase activity | Not Available |
| 1267 | Polysaccharide deacetylase | *Actinobacillus succinogenes* | Strain ATCC 55618 / DSM 22257 / CCUG 43843 / 130Z | A6VMM9 | -- | 21118570 | Hydrolase activity, acting on carbon-nitrogen (but not peptide) bonds | Not Available |
| 1268 | Poly-beta-1, 6-N-acetyl-D-glucosamine synthase (Poly-beta-1, 6-GlcNAc synthase) (EC 2.4.1.-) | *Actinobacillus succinogenes* | Strain ATCC 55618 / DSM 22257 / CCUG 43843 / 130Z | A6VMN0 | -- | 21118570 | Acetyl glucosaminyl transferase activity | Not Available |
| 1269 | Putative transmembrane protein | *Ralstonia solanaarum CFBP2957* | | D8P2G5 | -- | 20550686 | Not Available | Not Available |

| Target ID | Bacterial Protein Name | Organism Name | Strain | UniProt ID | PDB ID | PubMed ID | Classification as per UniProt GO Molecular Function | Classification of Function as per PDB |
|---|---|---|---|---|---|---|---|---|
| 1270 | Putative polysaccharide deacetylase associated with biofilm formation putative lipoprotein | *Escherichia coli O7:K1* | Strain IAI39 / ExPEC | A0A0H3MHK2 | -- | 19165319 | Hydrolase activity, acting on carbon-nitrogen (but not peptide) bonds | Not Available |
| 1271 | Motility quorum-sensing regulator | *Escherichia coli O7:K1* | Strain IAI39 / ExPEC | A0A0H3MKK3 | -- | 19165319 | Not Available | Not Available |
| 1272 | Toxin-antitoxin system, toxin component | *Burkholderia pyrrocinia* | Pseudomonas pyrrocinia | A0A0G3X296 | -- | 26150017 | Not Available | Not Available |
| 1273 | MqsR toxin | *Piscirickettsiaceae bacterium NZ-RLO1* | | A0A177W0R7 | -- | 27365345 | Not Available | Not Available |
| 1274 | Motility quorum-sensing regulator mqsR | *Tistrella mobilis* | Strain KA081020-065 | I3TW37 | -- | 22458477 | Not Available | Not Available |
| 1275 | Toxin-antitoxin system, toxin component | *Burkholderia pyrrocinia* | Pseudomonas pyrrocinia | A0A2Z5N2Q2 | -- | 29899517 | Not Available | Not Available |
| 1276 | Poly-beta-1, 6-N-acetyl-D-glucosamine synthase (Poly-beta-1, 6-GlcNAc synthase) (EC 2.4.1.-) | *Burkholderia pyrrocinia* | Pseudomonas pyrrocinia | A0A0G3X2J4 | -- | 26150017 | Acetyl glucosaminyl transferase activity | Not Available |
| 1277 | Poly-beta-1, 6-N-acetyl-D-glucosamine N-deacetylase PgaB | *Burkholderia pyrrocinia* | Pseudomonas pyrrocinia | A0A2Z5N2I1 | -- | 29899517 | Hydrolase activity, acting on carbon-nitrogen (but not peptide) bonds | Not Available |
| 1278 | Poly-beta-1, 6-N-acetyl-D-glucosamine synthase (Poly-beta-1, 6-GlcNAc synthase) (EC 2.4.1.-) | *Burkholderia pyrrocinia* | Pseudomonas pyrrocinia | A0A2Z5N4C5 | -- | 29899517 | Acetyl glucosaminyl transferase activity | Not Available |
| 1279 | Poly-beta-1, 6-N-acetyl-D-glucosamine synthase (Poly-beta-1, 6-GlcNAc synthase) (EC 2.4.1.-) | *Staphylococcus aureus subspp. aureus* | ST228 | A0A7U7F068 | -- | 22720005 | Acetyl glucosaminyl transferase activity | Not Available |
| 1280 | Poly-beta-1, 6-N-acetyl-D-glucosamine synthase (Poly-beta-1, 6-GlcNAc synthase) (EC 2.4.1.-) | *Yersinia enterocolitica serotype O:8 / biotype 1B* | Strain NCTC 13174 / 8081 | A1JS96 | -- | 17173484 | Acetyl glucosaminyl transferase activity | Not Available |
| 1281 | Uncharacterized protein | *Acinetobacter baylyi* | Strain ATCC 33305 / BD413 / ADP1 | Q6FDD7 | -- | 15514110 | Not Available | Not Available |
| 1282 | Putative lipoprotein (Haemin storage system) (HmsF) | *Acinetobacter baylyi* | Strain ATCC 33305 / BD413 / ADP1 | Q6FDD9 | -- | 15514110 | Hydrolase activity, acting on carbon-nitrogen (but not peptide) bonds | Not Available |
| 1283 | Poly-beta-1, 6-N-acetyl-D-glucosamine biosynthesis protein PgaD (Fragment) | *Acidiferrobacteraceae bacterium* | | A0A7V2NLF0 | -- | 31911466 | Not Available | Not Available |
| 1284 | Poly-beta-1, 6-N-acetyl-D-glucosamine N-deacetylase PgaB | *Pseudomonas fluorescens* | | A0A7Z8NJ86 | -- | 31040301 | Hydrolase activity, acting on carbon-nitrogen (but not peptide) bonds | Not Available |
| 1285 | Poly-beta-1, 6-N-acetyl-D-glucosamine biosynthesis protein PgaD | *Pseudomonas fluorescens* | | A0A7Z8NJX9 | -- | 31040301 | Not Available | Not Available |
| 1286 | Poly-beta-1, 6-N-acetyl-D-glucosamine N-deacetylase PgaB | *Paraburkholderia azotifigens* | | A0A5C6VNL0 | -- | 29185955 | Hydrolase activity, acting on carbon-nitrogen (but not peptide) bonds | Not Available |
| 1287 | Poly-beta-1, 6-N-acetyl-D-glucosamine synthase (Poly-beta-1, 6-GlcNAc synthase) (EC 2.4.1.-) | *Pseudomonas fluorescens* | | A0A7Z8NJM0 | -- | 31040301 | Acetyl glucosaminyl transferase activity | Not Available |
| 1288 | Uncharacterized protein | *Gallionellales bacterium GWA2_59_43* | | A0A1G0CW50 | -- | 27774985 | Not Available | Not Available |
| 1289 | Uncharacterized protein | *Pseudomonas putida* | Strain DOT-T1E | I7BZQ0 | -- | 23815283 | Not Available | Not Available |
| 1290 | Two-component system response regulator RpfG | *Xanthomonas campestris pv. vesicatoria* | Strain 85-10 | Q3BUB5 | -- | 16237009 | phosphoric diester Hydrolase activity | Not Available |
| 1291 | Uncharacterized protein | *Sphingobium baderi LL03* | | T0HNL1 | -- | 24051322 | Not Available | Not Available |
| 1292 | Poly-beta-1, 6-N-acetyl-D-glucosamine synthase (Poly-beta-1, 6-GlcNAc synthase) (EC 2.4.1.-) | *Pseudogulbenkiania spp.* | Strain NH8B | G2J3U5 | -- | 22038961 | Acetyl glucosaminyl transferase activity | Not Available |
| 1293 | Diguanylate cyclase | *Chloroflexus aurantiacus* | Strain ATCC 29366 / DSM 635 / J-10-fl | A9WDH0 | -- | 21714912 | Diguanylate cyclase activity | Not Available |
| 1294 | Diguanylate cyclase | *Chloroflexus aurantiacus* | Strain ATCC 29366 / DSM 635 / J-10-fl | A9WJ93 | -- | 21714912 | Diguanylate cyclase activity | Not Available |
| 1295 | Diguanylate cyclase | *Chloroflexus aurantiacus* | Strain ATCC 29366 / DSM 635 / J-10-fl | A9WIR5 | -- | 21714912 | Diguanylate cyclase activity | Not Available |
| 1296 | Polysaccharide deacetylase | *Pseudogulbenkiania spp.* | Strain NH8B | G2J3U6 | -- | 22038961 | Hydrolase activity, acting on carbon-nitrogen (but not peptide) bonds | Not Available |
| 1297 | Poly-beta-1, 6-N-acetyl-D-glucosamine biosynthesis protein PgaD | *Citrobacter freundii complex spp. CFNIH3* | | A0A2I8SAW1 | -- | 29437920 | Not Available | Not Available |
| 1298 | Poly-beta-1, 6-N-acetyl-D-glucosamine synthase (Poly-beta-1, 6-GlcNAc synthase) (EC 2.4.1.-) | *Desulfotalea psychrophila* | Strain LSv54 / DSM 12343 | Q6AMQ9 | -- | 15305914 | Acetyl glucosaminyl transferase activity | Not Available |
| 1299 | Poly-beta-1, 6-N-acetyl-D-glucosamine synthase (Poly-beta-1, 6-GlcNAc synthase) (EC 2.4.1.-) | *Citrobacter freundii complex spp. CFNIH3* | | A0A2I8SBF6 | -- | 29437920 | Acetyl glucosaminyl transferase activity | Not Available |
| 1300 | Histidinekinase (EC 2.7.13.3) | *Bradyrhizobium diazoefficiens* | Strain JCM 10833 / BCRC 13528 / IAM 13628 / NBRC 14792 / USDA 110 | Q89FJ6 | -- | 12597275 | Diguanylate cyclase activity, protein histidine kinase activity | Not Available |
| 1301 | Polysaccharide deacetylase | *Pseudogulbenkiania spp.* | Strain NH8B | G2IVP1 | -- | 22038961 | Hydrolase activity, acting on carbon-nitrogen (but not peptide) bonds | Not Available |
| 1302 | Uncharacterized protein | *Actinobacteria bacterium HGW-Actinobacteria-8* | | A0A2N3F169 | -- | 28350393 | Not Available | Not Available |
| 1303 | Uncharacterized protein | *Pseudogulbenkiania spp.* | Strain NH8B | G2J3U4 | -- | 22038961 | Not Available | Not Available |
| 1304 | mRNA interferase MqsR | *Geobacter spp. OR-1* | | A0A0A8WX61 | -- | 23668621 | Not Available | Not Available |
| 1305 | Surface composition regulator | *Citrobacter freundii complex spp. CFNIH3* | | A0A2I8SBG3 | -- | 29437920 | Not Available | Not Available |
| 1306 | Poly-beta-1, 6-N-acetyl-D-glucosamine N-deacetylase PgaB | *Citrobacter freundii complex spp. CFNIH3* | | A0A2I8SBF0 | -- | 29437920 | Hydrolase activity, acting on carbon-nitrogen (but not peptide) bonds | Not Available |
| 1307 | Diguanylate cyclase | *Chloroflexus aurantiacus* | Strain ATCC 29366 / DSM 635 / J-10-fl | A9WBR3 | -- | 21714912 | Diguanylate cyclase activity | Not Available |
| 1308 | Diguanylate cyclase | *Chloroflexus aurantiacus* | Strain ATCC 29366 / DSM 635 / J-10-fl | A9WGU1 | -- | 21714912 | Diguanylate cyclase activity | Not Available |
| 1309 | Diguanylate cyclase | *Chloroflexus aurantiacus* | Strain ATCC 29366 / DSM 635 / J-10-fl | A9WCL4 | -- | 21714912 | Diguanylate cyclase activity | Not Available |
| 1310 | Diguanylate cyclase | *Chloroflexus aurantiacus* | Strain ATCC 29366 / DSM 635 / J-10-fl | A9WKG1 | -- | 21714912 | Diguanylate cyclase activity | Not Available |
| 1311 | Diguanylate cyclase | *Chloroflexus aurantiacus* | Strain ATCC 29366 / DSM 635 / J-10-fl | A9WC49 | -- | 21714912 | Diguanylate cyclase activity | Not Available |
| 1312 | Poly-beta-1, 6-N-acetyl-D-glucosamine synthase (Poly-beta-1, 6-GlcNAc synthase) (EC 2.4.1.-) | *Stenotrophomonas spp. SAU14A_naIMI4_5* | | A0A3Q8FC29 | -- | 29765358 | Acetyl glucosaminyl transferase activity | Not Available |
| 1313 | Poly-beta-1, 6-N-acetyl-D-glucosamine N-deacetylase PgaB | *Stenotrophomonas spp. SAU14A_naIMI4_5* | | A0A3Q8FQ27 | -- | 29765358 | Hydrolase activity, acting on carbon-nitrogen (but not peptide) bonds | Not Available |
| 1314 | Bll7113 protein | *Bradyrhizobium diazoefficiens* | Strain JCM 10833 / BCRC 13528 / IAM 13628 / NBRC 14792 / USDA 110 | Q89EH2 | -- | 12597275 | Diguanylate cyclase activity | Not Available |
| 1315 | Bll1500 protein | *Bradyrhizobium diazoefficiens* | Strain JCM 10833 / BCRC 13528 / IAM 13628 / NBRC 14792 / USDA 110 | Q89UB7 | -- | 12597275 | Diguanylate cyclase activity | Not Available |
| 1316 | Poly-beta-1, 6-N-acetyl-D-glucosamine synthase (Poly-beta-1, 6-GlcNAc synthase) (EC 2.4.1.-) | *Stenotrophomonas spp. SAU14A_naIMI4_5* | | A0A3S5GH39 | -- | 29765358 | Acetyl glucosaminyl transferase activity | Not Available |

| Target ID | Bacterial Protein Name | Organism Name | Strain | UniProt ID | PDB ID | PubMed ID | Classification as per UniProt GO Molecular Function | Classification of Function as per PDB |
|---|---|---|---|---|---|---|---|---|
| 1317 | Poly-beta-1, 6-N-acetyl-D-glucosamine synthase (Poly-beta-1, 6-GlcNAc synthase) (EC 2.4.1.-) | *Enterobacter cloacae subspp. cloacae* | Strain ATCC 13047 / DSM 30054 / NBRC 13535 / NCTC 10005 / WDCM 00083 / NCDC 279-56 | A0A0H3CRE9 | -- | 20207761 | Acetyl glucosaminyl transferase activity | Not Available |
| 1318 | GGDEF domain-containing protein | *Streptomyces coelicolor* | Strain ATCC BAA-471 / A3 2 / M145 | Q9ADC3 | -- | 12000953 | Diguanylate cyclase activity | Not Available |
| 1319 | Uncharacterized protein | *Polaromona naphthalenivorans* | Strain CJ2 | A1VWM5 | -- | 19453698 | Not Available | Not Available |
| 1320 | Poly-beta-1, 6-N-acetyl-D-glucosamine synthase (Poly-beta-1, 6-GlcNAc synthase) (EC 2.4.1.-) | *Staphylococcus aureus* | Strain USA300 | A0A0H2XJC1 | -- | 16517273 | Acetyl glucosaminyl transferase activity | Not Available |
| 1321 | Poly-beta-1, 6-N-acetyl-D-glucosamine synthase (Poly-beta-1, 6-GlcNAc synthase) (EC 2.4.1.-) | *Enterobacter ludwigii* | | W0BUX8 | -- | 25197457; 26251488 | Acetyl glucosaminyl transferase activity | Not Available |
| 1322 | Type IV major pilin (Type IV pilus assembly protein PilA) | *Acidovorax citrulli* | Acidovorax avenae subspp. citrulli | C0LEX7 | -- | 19589067 | Not Available | Not Available |
| 1323 | ABC transporter substrate-binding protein (Hemin Storage System Protein) (Poly-beta-1, 6-N-acetyl-D-glucosamine biosynthesis protein PgaD) | *Enterobacter ludwigii* | | G8LE15 | -- | 22675579; 25197457 | Not Available | Not Available |
| 1324 | Poly-beta-1, 6-N-acetyl-D-glucosamine biosynthesis protein PgaD | *Gammaproteobacteria bacterium* | | A0A2E6LPH3 | -- | 29337314 | Not Available | Not Available |
| 1325 | Protein LpfD | *Salmonella typhimurium* | Strain LT2 / SGSC1412 / ATCC 700720 | P43663 | -- | 7721701; 11677609 | Not Available | Not Available |
| 1326 | Poly-beta-1, 6-N-acetyl-D-glucosamine N-deacetylase (PGA N-deacetylase) (Poly-beta-1, 6-GlcNAc N-deacetylase) (EC 3.5.1.-) | *Escherichia coli O157:H7* | | Q8XAR3 | -- | 11206551; 11258796 | Hydrolase activity, acting on carbon-nitrogen (but not peptide) bonds | Not Available |
| 1327 | Poly-beta-1, 6-N-acetyl-D-glucosamine synthase (PGA synthase) (Poly-beta-1, 6-GlcNAc synthase) (EC 2.4.1.-) (Biofilm PGA synthesis protein PgaC) (N-acetylglucosaminylTransferase PgaC) | *Escherichia coli O157:H7* | | Q8XAR5 | -- | 11206551; 11258796 | Acetyl glucosaminyl transferase activity | Not Available |
| 1328 | Poly-beta-1, 6-N-acetyl-D-glucosamine synthase (PGA synthase) (Poly-beta-1, 6-GlcNAc synthase) (EC 2.4.1.-) (Biofilm PGA synthesis protein PgaC) (N-acetylglucosaminylTransferase PgaC) | *Escherichia coli* | Strain K12 | P75905 | -- | 8905232; 9278503; 16738553; 15090514; 15919996; 18359807 | Acetyl glucosaminyl transferase activity | Not Available |
| 1329 | Biofilm PGA synthesis protein PgaD | *Escherichia coli O157:H7* | | P69433 | -- | 11206551; 11258796 | Not Available | Not Available |
| 1330 | Biofilm PGA synthesis protein PgaD | *Escherichia coli* | Strain K12 | P69432 | -- | 8905232; 9278503; 16738553; 15090514; 15919996; 18359807; 19460094 | Not Available | Not Available |
| 1331 | DNA-binding HTH domain-containing protein | *Eggerthella spp.* | Strain YY7918 | F7V097 | -- | 21914883 | DNA binding | Not Available |
| 1332 | Uncharacterized protein | *Bordetella parapertussis* | Strain Bpp5 | K0MGY7 | -- | 23051057 | Not Available | Not Available |
| 1333 | Sensory box / ggdef domain protein | *Thermodesulfovibrio yellowstonii* | Strain ATCC 51303 / DSM 11347 / YP87 | B5YGD8 | -- | 25635016 | Diguanylate cyclase activity | Not Available |
| 1334 | Signaling protein | *Thermodesulfovibrio yellowstonii* | Strain ATCC 51303 / DSM 11347 / YP87 | B5YH87 | -- | 25635016 | Diguanylate cyclase activity | Not Available |
| 1335 | Haemin storage system, HmsS protein | *Yersinia pestis bv. Antiqua* | Strain Antiqua | A0A0E1NTX1 | -- | 16740952 | Not Available | Not Available |
| 1336 | Poly-beta-1, 6-N-acetyl-D-glucosamine biosynthesis protein PgaD | *Lysobacter gummosus* | | A0A0S2G4X3 | -- | 26597042 | Not Available | Not Available |
| 1337 | Uncharacterized protein | *Bordetella pertussis STO1-CHOC-0019* | | A0A6N3RZF6 | -- | 24356839 | Not Available | Not Available |
| 1338 | Putative hemin storage protein | *Bordetella pertussis* | Strain ATCC 9797 / DSM 5571 / NCTC 10739 / 18323 | A0A0T7CLL3 | -- | 23051057 | Hydrolase activity, acting on carbon-nitrogen (but not peptide) bonds | Not Available |
| 1339 | Poly-beta-1, 6-N-acetyl-D-glucosamine N-deacetylase PgaB (EC 3.5.1.-) | *Lysobacter gummosus* | | A0A0S2G4X4 | -- | 26597042 | Hydrolase activity, acting on carbon-nitrogen (but not peptide) bonds | Not Available |
| 1340 | Uncharacterized protein | *Bordetella pertussis* | Strain ATCC 9797 / DSM 5571 / NCTC 10739 / 18323 | A0A0T7CP04 | -- | 23051057 | Not Available | Not Available |
| 1341 | Ggdef domain protein | *Thermodesulfovibrio yellowstonii* | Strain ATCC 51303 / DSM 11347 / YP87 | B5YJP8 | -- | 25635016 | Diguanylate cyclase activity | Not Available |
| 1342 | Uncharacterized protein | *Yersinia pestis bv. antiqua* | Strain Antiqua | A0A0E1NW62 | -- | 16740952 | Not Available | Not Available |
| 1343 | Poly-beta-1, 6-N-acetyl-D-glucosamine synthase (Poly-beta-1, 6-GlcNAc synthase) (EC 2.4.1.-) | *Bordetella pertussis* | Strain ATCC 9797 / DSM 5571 / NCTC 10739 / 18323 | A0A0T7CLJ6 | -- | 23051057 | Acetyl glucosaminyl transferase activity | Not Available |
| 1344 | Surface composition regulator | *Salmonella enterica subspp. enterica serovar* | Adelaide Str. A4-669 | A0A6C8GIJ5 | -- | 21859443 | Not Available | Not Available |
| 1345 | Surface composition regulator | *Salmonella schwarzengrund* | Strain CVM19633 | A0A0N1QW65 | -- | 21602358 | Not Available | Not Available |
| 1346 | Uncharacterized protein | *Anaeromyxobacter spp. PSR-1* | | A0A0D6QFE1 | -- | 23709511 | Not Available | Not Available |
| 1347 | Poly-beta-1, 6-N-acetyl-D-glucosamine N-deacetylase PgaB | *Shigella spp.* | | A0A351P2X7 | -- | 30148503 | Hydrolase activity, acting on carbon-nitrogen (but not peptide) bonds | Not Available |
| 1348 | Poly-beta-1, 6-N-acetyl-D-glucosamine N-deacetylase PgaB | *Klebsiella pneumoniae* | | A0A2L0KTV2 | -- | 29437920 | Hydrolase activity, acting on carbon-nitrogen (but not peptide) bonds | Not Available |
| 1349 | Uncharacterized protein | *Gallionellales bacterium RBG_16_56_9* | | A0A1G0CZQ4 | -- | 27774985 | Not Available | Not Available |
| 1350 | Hemin storage system protein | *Enterobacter spp. DC4* | | W7P486 | -- | 24625871 | Not Available | Not Available |
| 1351 | Uncharacterized protein | *Ligilactobacillus apodemi* | DSM 16634 = JCM 16172 | A0A0R1TQZ9 | -- | 26415554 | Not Available | Not Available |
| 1352 | Poly-beta-1, 6-N-acetyl-D-glucosamine synthase (Poly-beta-1, 6-GlcNAc synthase) (EC 2.4.1.-) | *Enterobacter spp. DC4* | | W7P4H8 | -- | 24625871 | Acetyl glucosaminyl transferase activity | Not Available |
| 1353 | Poly-beta-1, 6-N-acetyl-D-glucosamine synthase (Poly-beta-1, 6-GlcNAc synthase) (EC 2.4.1.-) | *Xanthomonas axonopodis pv. citri* | Strain 306 | Q8PLI7 | -- | 12024217 | Acetyl glucosaminyl transferase activity | Not Available |
| 1354 | Poly-beta-1, 6-N-acetyl-D-glucosamine N-deacetylase (EC 3.5.1.-) | *Enterobacter spp. DC4* | | W7NUM0 | -- | 24625871 | Hydrolase activity, acting on carbon-nitrogen (but not peptide) bonds | Not Available |
| 1355 | mRNA interferase MqsR | *Photorhabdus spp. S8-52* | | A0A329WQK4 | -- | 29877789 | Not Available | Not Available |
| 1356 | Poly-beta-1, 6-N-acetyl-D-glucosamine synthase (Poly-beta-1, 6-GlcNAc synthase) (EC 2.4.1.-) | *Hahella chejuensis* | Strain KCTC 2396 | Q2SAS4 | -- | 16352867 | Acetyl glucosaminyl transferase activity | Not Available |
| 1357 | Predicted xylanase / chitin deacetylase | *Hahella chejuensis* | Strain KCTC 2396 | Q2SAS5 | -- | 16352867 | Hydrolase activity, acting on carbon-nitrogen (but not peptide) bonds, Hydrolase activity, acting on glycosyl bonds | Not Available |

| Target ID | Bacterial Protein Name | Organism Name | Strain | UniProt ID | PDB ID | PubMed ID | Classification as per UniProt GO Molecular Function | Classification of Function as per PDB |
|---|---|---|---|---|---|---|---|---|
| 1358 | Poly-beta-1, 6-N-acetyl-D-glucosamine N-deacetylase PgaB | *Nitrospiraceae bacterium* | | A0A354A3Q8 | -- | 30148503 | Hydrolase activity, acting on carbon-nitrogen (but not peptide) bonds | Not Available |
| 1359 | Poly-beta-1, 6-N-acetyl-D-glucosamine biosynthesis protein PgaD | *Nitrospiraceae bacterium* | | A0A419J381 | -- | 28644444 | Not Available | Not Available |
| 1360 | Surface composition regulator | *Salmonella gallinarum* | Strain 287 / 91 / NCTC 13346 | B5REF5 | -- | 18583645 | Not Available | Not Available |
| 1361 | Poly-beta-1, 6-N-acetyl-D-glucosamine synthase (Poly-beta-1, 6-GlcNAc synthase) (EC 2.4.1.-) | *Nitrospiraceae bacterium* | | A0A354A3Q7 | -- | 30148503 | Acetyl glucosaminyl transferase activity | Not Available |
| 1362 | Poly-beta-1, 6-N-acetyl-D-glucosamine N-deacetylase PgaB | *Nitrospiraceae bacterium* | | A0A419J3A1 | -- | 28644444 | Hydrolase activity, acting on carbon-nitrogen (but not peptide) bonds | Not Available |
| 1363 | Poly-beta-1, 6-N-acetyl-D-glucosamine biosynthesis protein PgaD | *Nitrospiraceae bacterium* | | A0A354A3Q6 | -- | 30148503 | Not Available | Not Available |
| 1364 | RNA polymerase sigma factor FliA (RNA polymerase sigma factor for flagellar operon) (Sigma F) (Sigma-28) | *Salmonella typhimurium* | Strain 14028S / SGSC 2262 | A0A0F6B2T8 | -- | 19897643 | DNA binding, DNA-directed 5'-3' RNA polymerase activity, sigma factor activity | Not Available |
| 1365 | Poly-beta-1, 6-N-acetyl-D-glucosamine synthase (Poly-beta-1, 6-GlcNAc synthase) (EC 2.4.1.-) | *Nitrospiraceae bacterium* | | A0A419J3A6 | -- | 28644444 | Acetyl glucosaminyl transferase activity | Not Available |
| 1366 | PgaD protein | *Escherichia coli* | Strain 55989 / EAEC | B7LFD1 | -- | 19165319 | Not Available | Not Available |
| 1367 | Hypothetical lipoprotein YcdR | *Escherichia coli O6:K15:H31* | Strain 536 / UPEC | A0A454A2Z0 | -- | 16879640 | Hydrolase activity, acting on carbon-nitrogen (but not peptide) bonds | Not Available |
| 1368 | Poly-beta-1, 6-N-acetyl-D-glucosamine synthase (Poly-beta-1, 6-GlcNAc synthase) (EC 2.4.1.-) | *Thiotrichales bacterium SG8_50* | | A0A0S8EMN7 | -- | 25922666 | Acetyl glucosaminyl transferase activity | Not Available |
| 1369 | Uncharacterized protein (Fragment) | *Rhodospirillales bacterium* | RIFCSPLOWO2_12_FULL_67_15 | A0A1G3IPC8 | -- | 27774985 | Not Available | Not Available |
| 1370 | Uncharacterized protein | *Escherichia coli O6:K15:H31* | Strain 536 / UPEC | A0A454A321 | -- | 16879640 | Not Available | Not Available |
| 1371 | Poly-beta-1, 6-N-acetyl-D-glucosamine synthase (Poly-beta-1, 6-GlcNAc synthase) (EC 2.4.1.-) | *Lysobacter antibioticus* | | A0A0S2FCT6 | -- | 26597042 | Acetyl glucosaminyl transferase activity | Not Available |
| 1372 | Poly-beta-1, 6-N-acetyl-D-glucosamine synthase (Poly-beta-1, 6-GlcNAc synthase) (EC 2.4.1.-) | *Macrococcus bohemicus* | | A0A328A4I7 | -- | 29951040 | Acetyl glucosaminyl transferase activity | Not Available |
| 1373 | Poly-beta-1, 6-N-acetyl-D-glucosamine synthase (Poly-beta-1, 6-GlcNAc synthase) (EC 2.4.1.-) | *Macrococcus caseolyticus* | Strain JCSC5402 | B9EAX5 | -- | 19074389 | Acetyl glucosaminyl transferase activity | Not Available |
| 1374 | Biofilm PGA synthesis lipoprotein PgaB | *Paraburkholderia spp. BL8N3* | | A0A4V2PL11 | -- | 26203337 | Hydrolase activity, acting on carbon-nitrogen (but not peptide) bonds | Not Available |
| 1375 | Poly-beta-1, 6-N-acetyl-D-glucosamine synthase (Poly-beta-1, 6-GlcNAc synthase) (EC 2.4.1.-) | *Paraburkholderia spp. BL8N3* | | A0A4R1IGQ8 | -- | 26203337 | Acetyl glucosaminyl transferase activity | Not Available |
| 1376 | Surface composition regulator | *Citrobacter amalonaticus Y19* | | A0A0F6RIS0 | -- | 23377788 | Not Available | Not Available |
| 1377 | Toxin-antitoxin system, toxin component | *Burkholderiales bacterium RIFOXYD12_FULL_59_19* | | A0A1F4MM23 | -- | 27774985 | Not Available | Not Available |
| 1378 | Type II toxin-antitoxin system MqsR family toxin | *Agrobacterium salinitolerans* | | A0A546WCU3 | -- | 31273398 | Not Available | Not Available |
| 1379 | Poly-beta-1, 6-N-acetyl-D-glucosamine biosynthesis protein PgaD | *Lysobacter antibioticus* | | A0A0S2FD23 | -- | 26597042 | Not Available | Not Available |
| 1380 | Poly-beta-1, 6-N-acetyl-D-glucosamine N-deacetylase PgaB (EC 3.5.1.-) | *Lysobacter antibioticus* | | A0A0S2FCY0 | -- | 26597042 | Hydrolase activity, acting on carbon-nitrogen (but not peptide) bonds | Not Available |
| 1381 | Polysaccharide deacetylase | *Xenorhabdus spp. KJ12.1* | | A0A2D0LA49 | -- | 28993611 | Hydrolase activity, acting on carbon-nitrogen (but not peptide) bonds | Not Available |
| 1382 | Hemin storage protein | *Stenotrophomonas maltophilia* | Pseudomonas maltophilia Xanthomonas maltophilia | A0A0K2IYF3 | -- | 26324280 | Hydrolase activity, acting on carbon-nitrogen (but not peptide) bonds | Not Available |
| 1383 | HmsS protein | *Pseudomonas fluorescens WH6* | | E2XJK0 | -- | 20920191 | Not Available | Not Available |
| 1384 | Poly-beta-1, 6-N-acetyl-D-glucosamine biosynthesis protein PgaD | *Salmonella typhimurium* | | A0A708ADF7 | -- | 30286803 | Not Available | Not Available |
| 1385 | Poly-beta-1, 6-N-acetyl-D-glucosamine synthase (Poly-beta-1, 6-GlcNAc synthase) (EC 2.4.1.-) | *Pseudomonas fluorescens WH6* | | E2XJJ9 | -- | 20920191 | Acetyl glucosaminyl transferase activity | Not Available |
| 1386 | Surface composition regulator | *Salmonella enterica subspp. enterica serovar* | Aberdeen | A0A5I4KB90 | -- | 30286803 | Not Available | Not Available |
| 1387 | Biofilm polysaccharide intercellular adhesin deacetylase (Intercellular adhesion protein B) (Poly-beta-1, 6-N-acetyl-D-glucosamine N-deacetylase) | *Staphylococcus haemolyticus* | | B8Y3Y6 | -- | 19144798 | Hydrolase activity, acting on carbon-nitrogen (but not peptide) bonds, in linear amides | Not Available |
| 1388 | Haemin storage system, HmsS protein | *Yersinia enterocolitica* | type O:9 Str. YE56 / 03 | A0A7U7FEY4 | -- | 24753568 | Not Available | Not Available |
| 1389 | Poly-beta-1, 6-N-acetyl-D-glucosamine synthase (Poly-beta-1, 6-GlcNAc synthase) (EC 2.4.1.-) | *Staphylococcus haemolyticus* | | B8Y3Y4 | -- | 19144798 | Acetyl glucosaminyl transferase activity | Not Available |
| 1390 | Biofilm operon icaADBC HTH-type negative Transcriptional regulator IcaR (Intercellular adhesion protein R) | *Staphylococcus haemolyticus* | | B8Y3Y3 | -- | 19144798 | DNA binding | Not Available |
| 1391 | Uncharacterized protein | *Yersinia enterocolitica* | type O:9 Str. YE56 / 03 | A0A7U7FBJ4 | -- | 24753568 | Not Available | Not Available |
| 1392 | Intercellular adhesion protein C | *Staphylococcus haemolyticus* | | B8Y3Y7 | -- | 19144798 | Acyltransferase activity, transferring groups other than amino-acyl groups | Not Available |
| 1393 | Polysaccharide deacetylase | *Pseudomonas fluorescens WH6* | | E2XJJ8 | -- | 20920191 | Hydrolase activity, acting on carbon-nitrogen (but not peptide) bonds | Not Available |
| 1394 | Surface composition regulator | *Salmonella enterica subspp. houtenae serovar* | 48:g, z51:- | A0A727TFY8 | -- | 30286803 | Not Available | Not Available |
| 1395 | Poly-beta-1, 6-N-acetyl-D-glucosamine synthase (Poly-beta-1, 6-GlcNAc synthase) (EC 2.4.1.-) (Fragment) | *Staphylococcus saprophyticus* | | Q5YFS7 | -- | 15472359 | Acetyl glucosaminyl transferase activity | Not Available |
| 1396 | Biofilm polysaccharide intercellular adhesin synthesis protein IcaD (Intercellular adhesion protein D) (Poly-beta-1, 6-N-acetyl-D-glucosamine synthesis protein IcaD) | *Staphylococcus haemolyticus* | | B8Y3Y5 | -- | 19144798 | Not Available | Not Available |
| 1397 | FucA | *Streptococcus gordonii* | | Q6QT50 | -- | 15205427 | Alpha-L-fucosidase activity | Not Available |
| 1398 | Uncharacterized protein | *Pseudomonas fluorescens WH6* | | E2XLZ9 | -- | 20920191 | Not Available | Not Available |
| 1399 | Motility quorum-sensing regulator MqsR | *Pseudomonas fluorescens Q8r1-96* | | I4JYJ9 | -- | 22792073 | Not Available | Not Available |
| 1400 | Uncharacterized protein | *Acinetobacter oleivorans* | Strain JCM 16667 / KCTC 23045 / DR1 | D3JTG6 | -- | 20629796 | Not Available | Not Available |
| 1401 | Putative biofilm PGA synthesis protein PgaD | *Pseudomonas fluorescens Q8r1-96* | | I4L127 | -- | 22792073 | Not Available | Not Available |
| 1402 | Biofilm PGA synthesis lipoprotein PgaB (EC 3.-.-.-) | *Pseudomonas fluorescens Q8r1-96* | | I4KZS7 | -- | 22792073 | Hydrolase activity, acting on carbon-nitrogen (but not peptide) bonds | Not Available |
| 1403 | Surface composition regulator | *Salmonella enterica subspp. salamae serovar 18:z10:z6* | | A0A732CTX5 | -- | 30286803 | Not Available | Not Available |

| Target ID | Bacterial Protein Name | Organism Name | Strain | UniProt ID | PDB ID | PubMed ID | Classification as per UniProt GO Molecular Function | Classification of Function as per PDB |
|---|---|---|---|---|---|---|---|---|
| 1404 | SpsR | *Serratia plymuthica* | | C4PDW9 | -- | 21284858 | DNA binding | Not Available |
| 1405 | Surface composition regulator | *Yersinia enterocolitica* | Type O:9 Str. YE56 / 03 | A0A7U7IP11 | -- | 24753568 | DNA binding, DNA-directed DNA polymerase activity | Not Available |
| 1406 | Haemin storage system, HmsF protein | *Yersinia enterocolitica* | Type O:9 Str. YE56 / 03 | A0A7U7FEF6 | -- | 24753568 | Hydrolase activity, acting on carbon-nitrogen (but not peptide) bonds | Not Available |
| 1407 | Surface composition regulator | *Salmonella typhimurium* | | A0A3Z1VB25 | -- | 30286803 | Not Available | Not Available |
| 1408 | Poly-beta-1, 6-N-acetyl-D-glucosamine synthase (Poly-beta-1, 6-GlcNAc synthase) (EC 2.4.1.-) | *Yersinia enterocolitica* | Type O:9 Str. YE56 / 03 | A0A7U7FED0 | -- | 24753568 | Acetyl glucosaminyl transferase activity | Not Available |
| 1409 | Surface composition regulator | *Salmonella typhimurium* | | A0A709Z2J5 | -- | 30286803 | Not Available | Not Available |
| 1410 | Biofilm polysaccharide intercellular adhesin synthesis protein IcaD (Intercellular adhesion protein D) (Poly-beta-1, 6-N-acetyl-D-glucosamine synthesis protein IcaD) | *Staphylococcus caprae* | | Q9AIQ7 | -- | 11159959 | Not Available | Not Available |
| 1411 | Surface composition regulator | *Salmonella bongori serovar* | 44:r:- | A0A702BVU7 | -- | 30286803 | Not Available | Not Available |
| 1412 | Poly-beta-1, 6-N-acetyl-D-glucosamine N-deacetylase PgaB (EC 3.5.1.-) | *Salmonella typhimurium* | | A0A708AH06 | -- | 30286803 | Hydrolase activity, acting on carbon-nitrogen (but not peptide) bonds | Not Available |
| 1413 | Biofilm operon icaADBC HTH-type negative Transcriptional regulator IcaR (Intercellular adhesion protein R) | *Staphylococcus caprae* | | Q9AIQ9 | -- | 11159959 | DNA binding | Not Available |
| 1414 | IcaA (Fragment) | *Staphylococcus cohnii* | | Q83ZI1 | -- | 12957956 | Not Available | Not Available |
| 1415 | Surface composition regulator | *Salmonella enterica subspp. salamae serovar* | 58:a:- | A0A701W280 | -- | 30286803 | Not Available | Not Available |
| 1416 | Poly-beta-1, 6-N-acetyl-D-glucosamine synthase (Poly-beta-1, 6-GlcNAc synthase) (EC 2.4.1.-) | *Staphylococcus caprae* | | Q9AIQ8 | -- | 11159959 | Acetyl glucosaminyl transferase activity | Not Available |
| 1417 | Outer membrane N-deacetylase | *Pseudomonas moraviensis R28-S* | | V8RE08 | -- | 24558233 | Hydrolase activity, acting on carbon-nitrogen (but not peptide) bonds | Not Available |
| 1418 | Poly-beta-1, 6-N-acetyl-D-glucosamine synthase (Poly-beta-1, 6-GlcNAc synthase) (EC 2.4.1.-) | *Salmonella typhimurium* | | A0A708A923 | -- | 30286803 | Acetyl glucosaminyl transferase activity | Not Available |
| 1419 | Poly-beta-1, 6-N-acetyl-D-glucosamine synthase (Poly-beta-1, 6-GlcNAc synthase) (EC 2.4.1.-) | *Aggregatibacter actinomycetemcomitans* | Actinobacillus actinomycetemcomitan s Haemophilus actinomycetemcomitan s | Q5VJB2 | -- | 15576769 | Acetyl glucosaminyl transferase activity | Not Available |
| 1420 | Poly-beta-1, 6-N-acetyl-D-glucosamine synthase (Poly-beta-1, 6-GlcNAc synthase) (EC 2.4.1.-) | *Actinobacillus pleuropneumoniae* | Haemophilus pleuropneumoniae | Q5QFG3 | -- | 15576769 | Acetyl glucosaminyl transferase activity | Not Available |
| 1421 | Poly-beta-1, 6-N-acetyl-D-glucosamine synthase (Poly-beta-1, 6-GlcNAc synthase) (EC 2.4.1.-) | *Aggregatibacter actinomycetemcomitans* | Actinobacillus actinomycetemcomitan s Haemophilus actinomycetemcomitan s | A5HJW9 | -- | 17851029 | Acetyl glucosaminyl transferase activity | Not Available |
| 1422 | Biofilm polysaccharide intercellular adhesin deacetylase (Intercellular adhesion protein B) (Poly-beta-1, 6-N-acetyl-D-glucosamine N-deacetylase) | *Staphylococcus caprae* | | Q9AIQ6 | -- | 11159959 | Hydrolase activity, acting on carbon-nitrogen (but not peptide) bonds, in linear amides | Not Available |
| 1423 | Toxin CdiA | *Neisseria meningitidis* | | C0KXJ0 | -- | 19289515 | Toxin activity | Not Available |
| 1424 | Acyl-homoserine-lactone synthase (EC 2.3.1.184) (Autoinducer synthesis protein) | *Ensifer adhaerens* | Sinorhizobium morelense | K4K1V0 | -- | 23090571 | N-acyl homoserine lactone synthase activity | Not Available |
| 1425 | Putative membrane protein | *Bordetella bronchiseptica* | Strain ATCC BAA-588 / NCTC 13252 / RB50 AlcaligeneS bronchiSepticuS | A0A0H3LTS4 | -- | 12910271 | Not Available | Not Available |
| 1426 | Toxin CdiA | *Neisseria meningitidis* | | C0KXI6 | -- | 19289515 | toxin activity | Not Available |
| 1427 | Toxin CdiA | *Neisseria meningitidis* | | C0KXI4 | -- | 19289515 | toxin activity | Not Available |
| 1428 | Uncharacterized protein | *Cupriavidus oxalaticus* | | Q84ES5 | -- | 12902278 | Not Available | Not Available |
| 1429 | Surface composition regulator | *Salmonella enterica subspp. enterica serovar* | Oslo | A0A5I0WQX2 | -- | 30286803 | Not Available | Not Available |
| 1430 | Acyl-homoserine-lactone synthase (EC 2.3.1.184) (Autoinducer synthesis protein) | *Ensifer adhaerens* | Sinorhizobium morelense | K4KF64 | -- | 23090571 | N-acyl homoserine lactone synthase activity | Not Available |
| 1431 | Uncharacterized protein | *Cupriavidus necator* | Alcaligenes eutrophus Ralstonia eutropha | Q9ZAL5 | -- | 9887305 | Not Available | Not Available |
| 1432 | Acyl-homoserine-lactone synthase (EC 2.3.1.184) (Autoinducer synthesis protein) | *Ensifer adhaerens* | Sinorhizobium morelense | K4K584 | -- | 23090571 | N-acyl homoserine lactone synthase activity | Not Available |
| 1433 | Surface composition regulator | *Salmonella enterica* | Salmonella choleraesuis | A0A750KGM0 | -- | 30286803 | Not Available | Not Available |
| 1434 | Surface composition regulator | *Salmonella enterica* | Salmonella choleraesuis | A0A753AFU1 | -- | 30286803 | Not Available | Not Available |
| 1435 | Metal-binding permease | *Streptococcus gordonii* | Str. Challis | Q842E9 | -- | 12700268 | zinc ion binding | Not Available |
| 1436 | Poly-beta-1, 6-N-acetyl-D-glucosamine synthase (Poly-beta-1, 6-GlcNAc synthase) (EC 2.4.1.-) | *Bordetella bronchiseptica* | Strain ATCC BAA-588 / NCTC 13252 / RB50 Alcaligenes bronchisepticus | A0A0H3LKJ5 | -- | 12910271 | Acetyl glucosaminyl transferase activity | Not Available |
| 1437 | Surface composition regulator | *Salmonella enterica* | Salmonella choleraesuis | A0A702CJE0 | -- | 30286803 | Not Available | Not Available |
| 1438 | Surface composition regulator | *Salmonella moscow* | | A0A5H7LB63 | -- | 30286803 | Not Available | Not Available |
| 1439 | Surface composition regulator | *Salmonella typhimurium* | | A0A735I5S7 | -- | 30286803 | Not Available | Not Available |
| 1440 | Surface composition regulator | *Salmonella newport* | | A0A3V7UC64 | -- | 30286803 | Not Available | Not Available |
| 1441 | Surface composition regulator | *Escherichia coli* | | C3SU77 | -- | 19439656 | Not Available | Not Available |
| 1442 | Poly-beta-1, 6-N-acetyl-D-glucosamine synthase (Poly-beta-1, 6-GlcNAc synthase) (EC 2.4.1.-) | *Escherichia coli* | | C3TEV2 | -- | 19439656 | Acetyl glucosaminyl transferase activity | Not Available |
| 1443 | MrkA | *Escherichia coli* | | B0LHG5 | -- | 18055599; 19717626 | Not Available | Not Available |
| 1444 | Surface composition regulator | *Salmonella enterica subspp. enterica serovar* | Corvallis | A0A4U7YCJ9 | -- | 30286803 | Not Available | Not Available |
| 1445 | Surface composition regulator | *Salmonella enterica subspp. enterica serovar* | Rissen | A0A5H7QG39 | -- | 30286803 | Not Available | Not Available |
| 1446 | IcaA (Fragment) | *Staphylococcus saprophyticus* | | Q83TU9 | -- | 12957956 | Not Available | Not Available |
| 1447 | Surface composition regulator | *Salmonella typhi* | | A0A4U8GBI9 | -- | 30286803 | Not Available | Not Available |
| 1448 | Acyl-homoserine-lactone synthase (EC 2.3.1.184) (Autoinducer synthesis protein) | *Pseudomonas putida* | Arthrobacter siderocapsulatus | Q8GEM0 | -- | 12450862; 15345437; 16585751 | N-acyl homoserine lactone synthase activity | Not Available |
| 1449 | Surface composition regulator | *Salmonella enterica* | Salmonella choleraesuis | A0A750MQQ2 | -- | 30286803 | Not Available | Not Available |
| 1450 | Surface composition regulator | *Salmonella abortus-equi* | | A0A738E8L1 | -- | 30286803 | Not Available | Not Available |
| 1451 | Surface composition regulator | *Salmonella enterica subspp. enterica serovar* | Brandenburg | A0A5I9JE21 | -- | 30286803 | Not Available | Not Available |
| 1452 | MrkD | *Escherichia coli* | | B0LHG8 | -- | 18055599; 19717626 | Not Available | Not Available |
| 1453 | Surface composition regulator | *Salmonella enterica subspp. enterica serovar* | Paratyphi C | A0A6Y5YMH6 | -- | 30286803 | Not Available | Not Available |
| 1454 | MrkC | *Escherichia coli* | | B0LHG7 | -- | 18055599; 19717626 | Fimbrial usher porin activity | Not Available |
| 1455 | SLATT_6 domain-containing protein | *Escherichia coli* | | B0LHG1 | -- | 18055599; 19717626 | Not Available | Not Available |

| Target ID | Bacterial Protein Name | Organism Name | Strain | UniProt ID | PDB ID | PubMed ID | Classification as per UniProt GO Molecular Function | Classification of Function as per PDB |
|---|---|---|---|---|---|---|---|---|
| 1456 | Surface composition regulator | *Salmonella typhimurium* | Strain SL1344 | A0A718QE43 | -- | 30286803 | Not Available | Not Available |
| 1457 | AGS_C domain-containing protein | *Escherichia coli* | | B0LHG2 | -- | 18055599; 19717626 | Nucleotidyltransferase activity | Not Available |
| 1458 | Poly-beta-1, 6-N-acetyl-D-glucosamine synthase (Poly-beta-1, 6-GlcNAc synthase) (EC 2.4.1.-) | *Staphylococcus capitis* | | G4Y1I6 | -- | 23052315 | Acetyl glucosaminyl transferase activity | Not Available |
| 1459 | Surface composition regulator | *Salmonella enterica subspp. enterica serovar* | Choleraesuis | A0A5I8Q4H1 | -- | 30286803 | Not Available | Not Available |
| 1460 | AagD (PgaD) | *Aggregatibacter actinomycetemcomitans* | Actinobacillus actinomycetemcomitans Haemophilus actinomycetemcomitans | Q5VJB1 | -- | 15576769; 17851029 | Not Available | Not Available |
| 1461 | Surface composition regulator | *Salmonella enterica subspp. salamae serovar 48:d:z6* | | A0A729JLQ4 | -- | 30286803 | Not Available | Not Available |
| 1462 | Acyl-homoserine-lactone synthase (EC 2.3.1.184) (Autoinducer synthesis protein) | *Edwardsiella tarda* | | Q60HM1 | -- | 16233705; 19283379 | N-acyl homoserine lactone synthase activity | Not Available |
| 1463 | Surface composition regulator | *Salmonella enterica subspp. enterica serovar* | Alachua | A0A5I4B3X4 | -- | 30286803 | Not Available | Not Available |
| 1464 | Surface composition regulator | *Salmonella enterica subspp. enterica serovar* | Altona | A0A3V4SJ84 | -- | 30286803 | Not Available | Not Available |
| 1465 | Surface composition regulator | *Salmonella enterica subspp. enterica serovar* | Agama | A0A3V8VN13 | -- | 30286803 | Not Available | Not Available |
| 1466 | Surface composition regulator | *Salmonella enterica subspp. salamae serovar* | 47:b:1, 5 | A0A735M8M7 | -- | 30286803 | Not Available | Not Available |
| 1467 | Surface composition regulator | *Salmonella enterica subspp. houtenae serovar* | 44:z36[z38]:- | A0A737M2X2 | -- | 30286803 | Not Available | Not Available |
| 1468 | Surface composition regulator | *Salmonella enterica subspp. enterica serovar* | Rough O:-:- | A0A730TM30 | -- | 30286803 | Not Available | Not Available |
| 1469 | Surface composition regulator | *Salmonella enteritidis PT4* | Strain P125109 | A0A725G1U5 | -- | 30286803 | Not Available | Not Available |
| 1470 | Surface composition regulator | *Salmonella enterica subspp. salamae serovar* | 58:d:z6 | A0A737SLZ3 | -- | 30286803 | Not Available | Not Available |
| 1471 | Poly-beta-1, 6-N-acetyl-D-glucosamine synthase (Poly-beta-1, 6-GlcNAc synthase) (EC 2.4.1.-) | *Yersinia pestis* | | Q56941 | -- | 9249062 | Acetyl glucosaminyl transferase activity | Not Available |
| 1472 | Surface composition regulator | *Salmonella enterica* | Salmonella choleraesuis | A0A742XT71 | -- | 30286803 | Not Available | Not Available |
| 1473 | RNA polymerase sigma factor (Fragment) | *Edwardsiella tarda* | | C3U1E8 | -- | 19283379 | DNA binding, sigma factor activity | Not Available |
| 1474 | HmsF | *Yersinia pestis* | | Q56940 | -- | 9249062 | Hydrolase activity, acting on carbon-nitrogen (but not peptide) bonds | Not Available |
| 1475 | IcaA (Fragment) | *Mammaliicoccus sciuri* | Staphylococcus sciuri | Q83ZI2 | -- | 12957956 | Not Available | Not Available |
| 1476 | Poly-beta-1, 6-N-acetyl-D-glucosamine synthase (Poly-beta-1, 6-GlcNAc synthase) (EC 2.4.1.-) | *Xenorhabdus nematophila* | Achromobacter nematophilus | B3GN23 | -- | 18515487 | Acetyl glucosaminyl transferase activity | Not Available |
| 1477 | MacB | *Pseudomonas putida* | Arthrobacter siderocapsulatus | A8MN40 | -- | 16585751; 18599835 | ATP binding | Not Available |
| 1478 | SuhB (Fragment) | *Pseudomonas putida* | Arthrobacter siderocapsulatus | Q3Y8H9 | -- | 16585751 | Not Available | Not Available |
| 1479 | PpuR | *Pseudomonas putida* | Arthrobacter siderocapsulatus | Q3Y8H6 | -- | 16585751 | DNA binding | Not Available |
| 1480 | Surface composition regulator | *Salmonella enterica subspp. enterica serovar* | Ank | A0A5I2X3G4 | -- | 30286803 | Not Available | Not Available |
| 1481 | OprM | *Pseudomonas putida* | Arthrobacter siderocapsulatus | A8MN35 | -- | 16585751; 18599835 | Efflux transmembrane transporter activity | Not Available |
| 1482 | Putative polysaccharide deacetylase associated with biofilm formation putative lipoprotein (PgaB) | *Ralstonia syzygii R24* | | G3A8J5 | -- | 21931687 | Hydrolase activity, acting on carbon-nitrogen (but not peptide) bonds | Not Available |
| 1483 | Poly-beta-1, 6-N-acetyl-D-glucosamine biosynthesis protein PgaD | *Shigella spp.* | | A0A352GZL1 | -- | 30148503 | Not Available | Not Available |
| 1484 | Surface composition regulator | *Shigella spp.* | | A0A358UM46 | -- | 30148503 | Not Available | Not Available |
| 1485 | Poly-beta-1, 6-N-acetyl-D-glucosamine biosynthesis protein PgaD | *Klebsiella pneumoniae* | | A0A2L0KL40 | -- | 29437920 | Not Available | Not Available |
| 1486 | HmsS protein | *Klebsiella pneumoniae* | | W9BJP9 | -- | 25341126 | Not Available | Not Available |
| 1487 | Poly-beta-1, 6-N-acetyl-D-glucosamine N-deacetylase | *Anaeromyxobacter spp. PSR-1* | | A0A0D6QE29 | -- | 23709511 | Hydrolase activity, acting on carbon-nitrogen (but not peptide) bonds | Not Available |
| 1488 | Poly-beta-1, 6-N-acetyl-D-glucosamine biosynthesis protein PgaD | *Acinetobacter seifertii* | | N8S8L3 | -- | 25563912 | Not Available | Not Available |
| 1489 | Poly-beta-1, 6-N-acetyl-D-glucosamine N-deacetylase PgaB | *Metakosakonia spp. MRY16-398* | | A0A348DF60 | -- | 30524415 | Hydrolase activity, acting on carbon-nitrogen (but not peptide) bonds | Not Available |
| 1490 | Poly-beta-1, 6-N-acetyl-D-glucosamine biosynthesis protein PgaD | *Metakosakonia spp. MRY16-398* | | A0A348DF58 | -- | 30524415 | Not Available | Not Available |
| 1491 | Poly-beta-1, 6-N-acetyl-D-glucosamine N-deacetylase PgaB | *Gammaproteobacteria bacterium* | | A0A7C5JBZ9 | -- | 31911466 | Hydrolase activity, acting on carbon-nitrogen (but not peptide) bonds | Not Available |
| 1492 | Surface composition regulator | *Metakosakonia spp. MRY16-398* | | A0A348DK62 | -- | 30524415 | Not Available | Not Available |
| 1493 | Poly-beta-1, 6-N-acetyl-D-glucosamine biosynthesis protein PgaD | *Gammaproteobacteria bacterium* | | A0A3D1RX59 | -- | 30148503 | Not Available | Not Available |
| 1494 | Uncharacterized protein | *Gammaproteobacteria bacterium* | | A0A3C0I6L8 | -- | 30148503 | Not Available | Not Available |
| 1495 | Poly-beta-1, 6-N-acetyl-D-glucosamine synthase (Poly-beta-1, 6-GlcNAc synthase) (EC 2.4.1.-) | *Gammaproteobacteria bacterium* | | A0A3D1RSI1 | -- | 30148503 | Acetyl glucosaminyl transferase activity | Not Available |
| 1496 | TonB_C domain-containing protein | *Lysobacter enzymogenes* | | A0A1J1EBZ8 | -- | 28065880 | Not Available | Not Available |
| 1497 | Motility quorum-sensing regulator MqsR | *Deltaproteobacteria bacterium RIFCSPLOWO2_02_FULL_53_8* | | A0A1F9IJA9 | -- | 27774985 | Not Available | Not Available |
| 1498 | Poly-beta-1, 6-N-acetyl-D-glucosamine synthase (Poly-beta-1, 6-GlcNAc synthase) (EC 2.4.1.-) | *Lentisphaerae bacterium GWF2_50_93* | | A0A1G0ZQ85 | -- | 27774985 | Acetyl glucosaminyl transferase activity | Not Available |
| 1499 | Uncharacterized protein | *Deltaproteobacteria bacterium RIFCSPLOWO2_02_FULL_53_8* | | A0A1F9IY36 | -- | 27774985 | Not Available | Not Available |
| 1500 | Biofilm formation regulator HmsP | *Enterobacter agglomerans* | Erwinia herbicola Pantoea agglomerans | A0A8I0LKD8 | -- | 32966572 | Not Available | Not Available |
| 1501 | Poly-beta-1, 6-N-acetyl-D-glucosamine biosynthesis protein PgaD | *Lentisphaerae bacterium GWF2_50_93* | | A0A1G0ZS47 | -- | 27774985 | Not Available | Not Available |
| 1502 | Poly-beta-1, 6-N-acetyl-D-glucosamine N-deacetylase PgaB | *Lentisphaerae bacterium GWF2_50_93* | | A0A1G0ZQG0 | -- | 27774985 | Hydrolase activity, acting on carbon-nitrogen (but not peptide) bonds | Not Available |
| 1503 | Poly-beta-1, 6-N-acetyl-D-glucosamine synthase (Poly-beta-1, 6-GlcNAc synthase) (EC 2.4.1.-) | *Achromobacter xylosoxidans* | Strain A8 | E3HKY0 | -- | 21097610 | Acetyl glucosaminyl transferase activity | Not Available |
| 1504 | Poly-beta-1, 6-N-acetyl-D-glucosamine synthase (Poly-beta-1, 6-GlcNAc synthase) (EC 2.4.1.-) | *Carnobacterium divergens* | Lactobacillus divergens | A0A5F0MFX7 | -- | 30081251 | Acetyl glucosaminyl transferase activity | Not Available |
| 1505 | Biofilm PGA synthesis lipoprotein PgaB (EC 3.-.-.-) | *Achromobacter xylosoxidans* | Strain A8 | E3HKY1 | -- | 21097610 | Hydrolase activity, acting on carbon-nitrogen (but not peptide) bonds | Not Available |

| Target ID | Bacterial Protein Name | Organism Name | Strain | UniProt ID | PDB ID | PubMed ID | Classification as per UniProt GO Molecular Function | Classification of Function as per PDB |
|---|---|---|---|---|---|---|---|---|
| 1506 | Poly-beta-1, 6-N-acetyl-D-glucosamine synthase (Poly-beta-1, 6-GlcNAc synthase) (EC 2.4.1.-) | *Pseudomonas moraviensis R28-S* | | V8RF67 | -- | 24558233 | Acetyl glucosaminyl transferase activity | Not Available |
| 1507 | Uncharacterized protein | *Caldanaerobacter subterraneus subspp. yonseiensis KB-1* | | U5CWT9 | -- | 24201201 | Not Available | Not Available |
| 1508 | Poly-beta-1, 6-N-acetyl-D-glucosamine synthase (Poly-beta-1, 6-GlcNAc synthase) (EC 2.4.1.-) | *Kangiella koreensis* | Strain DSM 16069 / KCTC 12182 / SW-125 | C7RBF1 | -- | 21304661 | Acetyl glucosaminyl transferase activity | Not Available |
| 1509 | Poly-beta-1, 6-N-acetyl-D-glucosamine synthase (Poly-beta-1, 6-GlcNAc synthase) (EC 2.4.1.-) | *Alicyclobacillus acidocaldarius subspp. acidocaldarius* | Strain ATCC 27009 / DSM 446 / BCRC 14685 / JCM 5260 / KCTC 1825 / NBRC 15652 / NCIMB 11725 / NRRL B-14509 / 104-IA Bacillus acidocaldarius | C8WT25 | -- | 21304673 | Acetyl glucosaminyl transferase activity | Not Available |
| 1510 | Poly-beta-1, 6-N-acetyl-D-glucosamine synthase (Poly-beta-1, 6-GlcNAc synthase) (EC 2.4.1.-) | *Escherichia coli O157:H7* | Strain EC869 | A0A0H3PPW9 | -- | 21421787 | Acetyl glucosaminyl transferase activity | Not Available |
| 1511 | Poly-beta-1, 6-N-acetyl-D-glucosamine N-deacetylase PgaB | *Stenotrophomonas spp. ESTM1D_MKCIP4_1* | | A0A3S7KT60 | -- | 29765358 | Hydrolase activity, acting on carbon-nitrogen (but not peptide) bonds | Not Available |
| 1512 | Poly-beta-1, 6-N-acetyl-D-glucosamine synthase (Poly-beta-1, 6-GlcNAc synthase) (EC 2.4.1.-) | *Comamonae bacterium* | | A0A4Q3R2D4 | -- | 30498029 | Acetyl glucosaminyl transferase activity | Not Available |
| 1513 | Poly-beta-1, 6-N-acetyl-D-glucosamine biosynthesis protein PgaD | *Comamonae bacterium* | | A0A4Q3QYG0 | -- | 30498029 | Not Available | Not Available |
| 1514 | Lipoprotein for biofilm formation | *Bacillus subtilis XF-1* | | M4KUX5 | -- | 23558530 | Not Available | Not Available |
| 1515 | Poly-beta-1, 6-N-acetyl-D-glucosamine synthase (Poly-beta-1, 6-GlcNAc synthase) (EC 2.4.1.-) | *Pectobacterium atrosepticum* | Strain SCRI 1043 / ATCC BAA-672 Erwinia carotovora subspp. atroseptica | Q6CYQ3 | -- | 15263089 | Acetyl glucosaminyl transferase activity | Not Available |
| 1516 | Surface composition regulator | *Pectobacterium atrosepticum* | Strain SCRI 1043 / ATCC BAA-672 Erwinia carotovora subspp. atroseptica | Q6D4M7 | -- | 15263089 | DNA binding, DNA-directed DNA polymerase activity | Not Available |
| 1517 | Putative hemin storage protein | *Pectobacterium atrosepticum* | Strain SCRI 1043 / ATCC BAA-672 Erwinia carotovora subspp. atroseptica | Q6CYQ4 | -- | 15263089 | Not Available | Not Available |
| 1518 | Poly-beta-1, 6-N-acetyl-D-glucosamine N-deacetylase PgaB | *Comamonae bacterium* | | A0A4Q3R554 | -- | 30498029 | Hydrolase activity, acting on carbon-nitrogen (but not peptide) bonds | Not Available |
| 1519 | Poly-beta-1, 6-N-acetyl-D-glucosamine biosynthesis protein PgaD | *Comamonae bacterium* | | A0A4Q3PH24 | -- | 30498029 | Not Available | Not Available |
| 1520 | Poly-beta-1, 6-N-acetyl-D-glucosamine biosynthesis protein PgaD | *Comamonae bacterium* | | A0A4Q3R2C1 | -- | 30498029 | Not Available | Not Available |
| 1521 | Uncharacterized protein | *Leptospira licerasiae serovar Varillal* | Str. VAR 10 | I0XRG0 | -- | 23145189 | Not Available | Not Available |
| 1522 | Uncharacterized protein | *Stenotrophomonas rhizophila* | | A0A4U3GLQ3 | -- | 31040301 | Not Available | Not Available |
| 1523 | Poly-beta-1, 6-N-acetyl-D-glucosamine synthase (Poly-beta-1, 6-GlcNAc synthase) (EC 2.4.1.-) | *Stenotrophomonas rhizophila* | | A0A4U3GSE5 | -- | 31040301 | Acetyl glucosaminyl transferase activity | Not Available |
| 1524 | Uncharacterized protein | *Pseudomonas brassicacearum* | Strain NFM421 | F2KL25 | -- | 21515771 | Not Available | Not Available |
| 1525 | Poly-beta-1, 6-N-acetyl-D-glucosamine N-deacetylase PgaB | *Stenotrophomonas rhizophila* | | A0A4U3GQQ3 | -- | 31040301 | Hydrolase activity, acting on carbon-nitrogen (but not peptide) bonds | Not Available |
| 1526 | Putative biofilm PNot AvailableG synthesis protein | *Pseudomonas brassicacearum* | Strain NFM421 | F2KD34 | -- | 21515771 | Not Available | Not Available |
| 1527 | Poly-beta-1, 6-N-acetyl-D-glucosamine biosynthesis protein PgaD | *Stenotrophomonas rhizophila* | | A0A4V5UEL4 | -- | 31040301 | Not Available | Not Available |
| 1528 | Uncharacterized protein | *Deltaproteobacteria bacterium RBG_16_58_17* | | A0A1F9CH81 | -- | 27774985 | Not Available | Not Available |
| 1529 | Poly-beta-1, 6-N-acetyl-D-glucosamine synthase (Poly-beta-1, 6-GlcNAc synthase) (EC 2.4.1.-) | *Comamonadaceae bacterium* | | A0A4Q3RBI5 | -- | 30498029 | Acetyl glucosaminyl transferase activity | Not Available |
| 1530 | Poly-beta-1, 6-N-acetyl-D-glucosamine N-deacetylase PgaB (Fragment) | *Comamonadaceae bacterium* | | A0A4V2ABY7 | -- | 30498029 | Hydrolase activity, acting on carbon-nitrogen (but not peptide) bonds | Not Available |
| 1531 | Poly-beta-1, 6-N-acetyl-D-glucosamine biosynthesis protein PgaD | *Proteobacteria bacterium ST_bin11* | | A0A1W9K6T1 | -- | 28390307 | Not Available | Not Available |
| 1532 | Putative hemin storage lipoprotein | *Pectobacterium atrosepticum* | Strain SCRI 1043 / ATCC BAA-672 Erwinia carotovora subspp. atroseptica | Q6CYQ2 | -- | 15263089 | Hydrolase activity, acting on carbon-nitrogen (but not peptide) bonds | Not Available |
| 1533 | Poly-beta-1, 6-N-acetyl-D-glucosamine N-deacetylase PgaB | *Proteobacteria bacterium ST_bin11* | | A0A1W9K6R2 | -- | 28390307 | Hydrolase activity, acting on carbon-nitrogen (but not peptide) bonds | Not Available |
| 1534 | Poly-beta-1, 6-N-acetyl-D-glucosamine N-deacetylase PgaB | *Acinetobacter spp.* | | A0A519JFC4 | -- | 30498029 | Hydrolase activity, acting on carbon-nitrogen (but not peptide) bonds | Not Available |
| 1535 | Poly-beta-1, 6-N-acetyl-D-glucosamine synthase (Poly-beta-1, 6-GlcNAc synthase) (EC 2.4.1.-) | *Acinetobacter spp.* | | A0A519JFC8 | -- | 30498029 | Acetyl glucosaminyl transferase activity | Not Available |
| 1536 | Poly-beta-1, 6-N-acetyl-D-glucosamine synthase (Poly-beta-1, 6-GlcNAc synthase) (EC 2.4.1.-) | *Klebsiella aerogenes* | Strain ATCC 13048 / DSM 30053 / JCM 1235 / KCTC 2190 / NBRC 13534 / NCIMB 10102 / NCTC 10006 Enterobacter aerogenes | A0A0H3FV53 | -- | 22493190 | Acetyl glucosaminyl transferase activity | Not Available |
| 1537 | Poly-beta-1, 6-N-acetyl-D-glucosamine biosynthesis protein PgaD | *Acinetobacter spp.* | | A0A519JGP9 | -- | 30498029 | Not Available | Not Available |
| 1538 | Poly-beta-1, 6-N-acetyl-D-glucosamine synthase (Poly-beta-1, 6-GlcNAc synthase) (EC 2.4.1.-) | *Acinetobacter spp.* | | A0A519JGN0 | -- | 30498029 | Acetyl glucosaminyl transferase activity | Not Available |
| 1539 | Hemin storage system protein | *Klebsiella aerogenes* | Strain ATCC 13048 / DSM 30053 / JCM 1235 / KCTC 2190 / NBRC 13534 / NCIMB 10102 / NCTC 10006 Enterobacter aerogenes | A0A0H3FMQ6 | -- | 22493190 | Not Available | Not Available |
| 1540 | Poly-beta-1, 6-N-acetyl-D-glucosamine synthase (Poly-beta-1, 6-GlcNAc synthase) (EC 2.4.1.-) | *Comamonae bacterium* | | A0A4Q3QVP0 | -- | 30498029 | Acetyl glucosaminyl transferase activity | Not Available |
| 1541 | Poly-beta-1, 6-N-acetyl-D-glucosamine N-deacetylase PgaB | *Acinetobacter spp.* | | A0A519JFC5 | -- | 30498029 | Hydrolase activity, acting on carbon-nitrogen (but not peptide) bonds | Not Available |
| 1542 | Poly-beta-1, 6-N-acetyl-D-glucosamine biosynthesis protein PgaD | *Comamonae bacterium* | | A0A4Q3Q3K8 | -- | 30498029 | Not Available | Not Available |

| Target ID | Bacterial Protein Name | Organism Name | Strain | UniProt ID | PDB ID | PubMed ID | Classification as per UniProt GO Molecular Function | Classification of Function as per PDB |
|---|---|---|---|---|---|---|---|---|
| 1543 | Poly-beta-1, 6-N-acetyl-D-glucosamine biosynthesis protein PgaD | *Acinetobacter spp.* | | A0A519JFE0 | -- | 30498029 | Not Available | Not Available |
| 1544 | Poly-beta-1, 6-N-acetyl-D-glucosamine synthase (Poly-beta-1, 6-GlcNAc synthase) (EC 2.4.1.-) | *Proteobacteria bacterium ST_bin11* | | A0A1W9K6Q3 | -- | 28390307 | Acetyl glucosaminyl transferase activity | Not Available |
| 1545 | Poly-beta-1, 6-N-acetyl-D-glucosamine N-deacetylase PgaB (Fragment) | *Comamonae bacterium* | | A0A4Q3NPB9 | -- | 30498029 | Hydrolase activity, acting on carbon-nitrogen (but not peptide) bonds | Not Available |
| 1546 | Poly-beta-1, 6-N-acetyl-D-glucosamine N-deacetylase PgaB (EC 3.5.1.-) | *Pseudomonas spp. BIOMIG1BAC* | | A0A6G6ZRV1 | -- | 26992451 | Hydrolase activity, acting on carbon-nitrogen (but not peptide) bonds | Not Available |
| 1547 | Surface composition regulator | *Klebsiella aerogenes* | Strain ATCC 13048 / DSM 30053 / JCM 1235 / KCTC 2190 / NBRC 13534 / NCIMB 10102 / NCTC 10006 Enterobacter aerogenes | A0A0H3FS42 | -- | 22493190 | Not Available | Not Available |
| 1548 | Type II toxin-antitoxin system MqsR family toxin | *Comamonae bacterium* | | A0A4Q3QZB8 | -- | 30498029 | Not Available | Not Available |
| 1549 | Poly-beta-1, 6-N-acetyl-D-glucosamine N-deacetylase PgaB | *Acinetobacter spp.* | | A0A519JGN1 | -- | 30498029 | Hydrolase activity, acting on carbon-nitrogen (but not peptide) bonds | Not Available |
| 1550 | Poly-beta-1, 6-N-acetyl-D-glucosamine biosynthesis protein PgaD | *Comamonadaceae bacterium* | | A0A4V1UAI0 | -- | 30498029 | Not Available | Not Available |
| 1551 | Lipoprotein for biofilm formation | *Bacillus spp. WP8* | | A0A0B4S988 | -- | 25614565 | Not Available | Not Available |
| 1552 | Poly-beta-1, 6-N-acetyl-D-glucosamine N-deacetylase PgaB | *Comamonadaceae bacterium* | | A0A4Q3RD47 | -- | 30498029 | Hydrolase activity, acting on carbon-nitrogen (but not peptide) bonds | Not Available |
| 1553 | Poly-beta-1, 6-N-acetyl-D-glucosamine biosynthesis protein PgaD | *Comamonadaceae bacterium* | | A0A4Q3PKV1 | -- | 30498029 | Not Available | Not Available |
| 1554 | Poly-beta-1, 6-N-acetyl-D-glucosamine biosynthesis protein PgaD | *Pseudomonas spp. BIOMIG1BAC* | | A0A6G6ZS19 | -- | 26992451 | Not Available | Not Available |
| 1555 | Poly-beta-1, 6-N-acetyl-D-glucosamine N-deacetylase PgaB (Fragment) | *Comamonadaceae bacterium* | | A0A4Q5X007 | -- | 30498029 | Hydrolase activity, acting on carbon-nitrogen (but not peptide) bonds | Not Available |
| 1556 | Putative biofilm PNot AvailableG synthesis protein | *Pseudomonas brassicacearum* | Strain NFM421 | F2KD32 | -- | 21515771 | Hydrolase activity, acting on carbon-nitrogen (but not peptide) bonds | Not Available |
| 1557 | Poly-beta-1, 6-N-acetyl-D-glucosamine N-deacetylase PgaB | *Pseudomonas viciae* | | A0A4P7PAA8 | -- | 30941113 | Hydrolase activity, acting on carbon-nitrogen (but not peptide) bonds | Not Available |
| 1558 | Poly-beta-1, 6-N-acetyl-D-glucosamine N-deacetylase PgaB | *Acinetobacter pittii* | Acinetobacter genomospp. 3 | A0A6H2VLN2 | -- | 30036696 | Hydrolase activity, acting on carbon-nitrogen (but not peptide) bonds | Not Available |
| 1559 | Polysaccharide deacetylase | *Burkholderia cenocepacia* | Strain ATCC BAA-245 / DSM 16553 / LMG 16656 / NCTC 13227 / J2315 / CF5610 Burkholderia cepacia Strain J2315 | B4EHA7 | -- | 18931103 | Hydrolase activity, acting on carbon-nitrogen (but not peptide) bonds | Not Available |
| 1560 | Poly-beta-1, 6-N-acetyl-D-glucosamine synthase (Poly-beta-1, 6-GlcNAc synthase) (EC 2.4.1.-) | *Burkholderia cenocepacia* | Strain ATCC BAA-245 / DSM 16553 / LMG 16656 / NCTC 13227 / J2315 / CF5610 Burkholderia cepacia Strain J2315 | B4EHA6 | -- | 18931103 | Acetyl glucosaminyl transferase activity | Not Available |
| 1561 | Poly-beta-1, 6-N-acetyl-D-glucosamine biosynthesis protein PgaD | *Acinetobacter pittii* | Acinetobacter genomospp. 3 | A0A6H2VP90 | -- | 30036696 | Not Available | Not Available |
| 1562 | Polysaccharide deacetylase | *Xenorhabdus stockiae* | | A0A2D0KWI4 | -- | 28993611 | Hydrolase activity, acting on carbon-nitrogen (but not peptide) bonds | Not Available |
| 1563 | GGDEF family protein | *Vibrio cholerae serotype O1* | Strain ATCC 39315 / El Tor Inaba N16961 | Q9KPY8 | -- | 10952301 | Diguanylate cyclase activity | Not Available |
| 1564 | GGDEF family protein | *Vibrio cholerae serotype O1* | Strain ATCC 39315 / El Tor Inaba N16961 | Q9KN01 | -- | 10952301 | Diguanylate cyclase activity | Not Available |
| 1565 | GGDEF family protein | *Vibrio cholerae serotype O1* | Strain ATCC 39315 / El Tor Inaba N16961 | Q9KS83 | -- | 10952301 | Diguanylate cyclase activity | Not Available |
| 1566 | Poly-beta-1, 6-N-acetyl-D-glucosamine synthase (Poly-beta-1, 6-GlcNAc synthase) (EC 2.4.1.-) | *Lachnospiraceae bacterium* | | A0A3D0DPD1 | -- | 30148503 | Acetyl glucosaminyl transferase activity | Not Available |
| 1567 | GGDEF family protein | *Vibrio cholerae serotype O1* | Strain ATCC 39315 / El Tor Inaba N16961 | Q9KN91 | -- | 10952301 | Diguanylate cyclase activity | Not Available |
| 1568 | GGDEF family protein | *Vibrio cholerae serotype O1* | Strain ATCC 39315 / El Tor Inaba N16961 | Q9KNB6 | -- | 10952301 | Diguanylate cyclase activity | Not Available |
| 1569 | GGDEF family protein | *Vibrio cholerae serotype O1* | Strain ATCC 39315 / El Tor Inaba N16961 | Q9KPS7 | -- | 10952301 | Diguanylate cyclase activity | Not Available |
| 1570 | Uncharacterized protein | *Cupriavidus necator* | Strain ATCC 17699 / DSM 428 / KCTC 22496 / NCIMB 10442 / H16 / stanier 337 Ralstonia eutropha | Q0KF48 | -- | 16964242 | Not Available | Not Available |
| 1571 | GGDEF family protein | *Vibrio cholerae serotype O1* | Strain ATCC 39315 / El Tor Inaba N16961 | Q9KS92 | -- | 10952301 | Diguanylate cyclase activity | Not Available |
| 1572 | Surface composition regulator | *Serratia spp. AS13* | | A0A7U3Z3G0 | -- | 23450001 | Not Available | Not Available |
| 1573 | mRNA interferase MqsR | *Betaproteobacteria bacterium RBG_16_58_11* | | A0A1F3X5Y1 | -- | 27774985 | Not Available | Not Available |
| 1574 | Uncharacterized protein | *Azospirillum lipoferum* | Strain 4B | G7Z3P9 | -- | 22216014 | Not Available | Not Available |
| 1575 | Poly-beta-1, 6-N-acetyl-D-glucosamine N-deacetylase PgaB | *Acinetobacter pittii* | Acinetobacter genomospp. 3 | A0A6H2VGR7 | -- | 30036696 | Hydrolase activity, acting on carbon-nitrogen (but not peptide) bonds | Not Available |
| 1576 | Regulator of the extracellular matrix, regulation of biofilm formation | *Bacillus methanolicus PB1* | | I3E0X0 | -- | 22610424 | Not Available | Not Available |
| 1577 | Poly-beta-1, 6-N-acetyl-D-glucosamine N-deacetylase PgaB | *Chromobacterium haemolyticum* | | A0A2T5P0U5 | -- | 29670144 | Hydrolase activity, acting on carbon-nitrogen (but not peptide) bonds | Not Available |
| 1578 | Lipoprotein for biofilm formation | *Bacillus spizizenii* | Strain ATCC 23059 / NRRL B-14472 / W23 Bacillus Subtilis subspp. spizizenii | E0U410 | -- | 21527469 | Not Available | Not Available |
| 1579 | Poly-beta-1, 6-N-acetyl-D-glucosamine N-deacetylase PgaB | *Acinetobacter seifertii* | | N8SA96 | -- | 25563912 | Hydrolase activity, acting on carbon-nitrogen (but not peptide) bonds | Not Available |
| 1580 | Uncharacterized protein | *Gammaproteobacteria bacterium HGW-Gammaproteobacteria-14* | | A0A2N1YKL6 | -- | 28350393 | Not Available | Not Available |
| 1581 | Poly-beta-1, 6-N-acetyl-D-glucosamine biosynthesis protein PgaD | *Enterobacter chengduensis* | | A0A5C1C3Z6 | -- | 30302649 | Not Available | Not Available |
| 1582 | Poly-beta-1, 6-N-acetyl-D-glucosamine synthase (Poly-beta-1, 6-GlcNAc synthase) (EC 2.4.1.-) | *Enterobacter chengduensis* | | A0A5C1C402 | -- | 30302649 | Acetyl glucosaminyl transferase activity | Not Available |

| Target ID | Bacterial Protein Name | Organism Name | Strain | UniProt ID | PDB ID | PubMed ID | Classification as per UniProt GO Molecular Function | Classification of Function as per PDB |
|---|---|---|---|---|---|---|---|---|
| 1583 | Poly-beta-1, 6-N-acetyl-D-glucosamine synthase (Poly-beta-1, 6-GlcNAc synthase) (EC 2.4.1.-) | *Pseudomonas fluorescens* | Strain Pf0-1 | Q3KJY3 | -- | 19432983 | Acetyl glucosaminyl transferase activity | Not Available |
| 1584 | Hemin storage system, HmsF protein | *Pseudomonas fluorescens* | Strain Pf0-1 | Q3KJY4 | -- | 19432983 | Hydrolase activity, acting on carbon-nitrogen (but not peptide) bonds | Not Available |
| 1585 | Putative hemin storage system | *Pseudomonas fluorescens* | Strain Pf0-1 | Q3KJY2 | -- | 19432983 | Not Available | Not Available |
| 1586 | Type II toxin-antitoxin system MqsR family toxin | *Pseudomonas spp. WS 5019* | | A0A7Y1FEC9 | -- | 32587583 | Not Available | Not Available |
| 1587 | Uncharacterized protein | *Pseudomonas putida ND6* | | I3V0P5 | -- | 22933774 | Not Available | Not Available |
| 1588 | Type II toxin-antitoxin system MqsR family toxin | *Oxalobacteraceae bacterium* | | A0A4Q3HUY0 | -- | 30498029 | Not Available | Not Available |
| 1589 | Poly-beta-1, 6-N-acetyl-D-glucosamine N-deacetylase PgaB (Fragment) | *Oxalobacteraceae bacterium* | | A0A4Q3K4V3 | -- | 30498029 | Hydrolase activity, acting on carbon-nitrogen (but not peptide) bonds | Not Available |
| 1590 | Poly-beta-1, 6-N-acetyl-D-glucosamine N-deacetylase PgaB | *Pelosinus spp. UFO1* | | A0A075KI64 | -- | 25189589 | Hydrolase activity, acting on carbon-nitrogen (but not peptide) bonds | Not Available |
| 1591 | Poly-beta-1, 6-N-acetyl-D-glucosamine N-deacetylase PgaB | *Pelosinus spp. UFO1* | | A0A075KDY1 | -- | 25189589 | Hydrolase activity, acting on carbon-nitrogen (but not peptide) bonds | Not Available |
| 1592 | HmsF protein | *Klebsiella pneumoniae subspp. rhinoscleromatis SB3432* | | R4Y4I7 | -- | 21629720 | Hydrolase activity, acting on carbon-nitrogen (but not peptide) bonds | Not Available |
| 1593 | HmsS protein | *Klebsiella pneumoniae subspp. rhinoscleromatis SB3432* | | R4YAK7 | -- | 21629720 | Not Available | Not Available |
| 1594 | Surface composition regulator | *Salmonella enterica subspp. enterica serovar* | Uganda Str. R8-3404 | A0A6C8GYH9 | -- | 21859443 | Not Available | Not Available |
| 1595 | Poly-beta-1, 6-N-acetyl-D-glucosamine synthase (Poly-beta-1, 6-GlcNAc synthase) (EC 2.4.1.-) | *Pelosinus spp. UFO1* | | A0A075KH19 | -- | 25189589 | Acetyl glucosaminyl transferase activity | Not Available |
| 1596 | Type II toxin-antitoxin system MqsR family toxin | *Lentisphaerae bacterium* | | A0A7X8XB07 | -- | 32123542 | Not Available | Not Available |
| 1597 | Poly-beta-1, 6-N-acetyl-D-glucosamine N-deacetylase PgaB | *Lentisphaerae bacterium GWF2_52_8* | | A0A1G0ZGS0 | -- | 27774985 | Hydrolase activity, acting on carbon-nitrogen (but not peptide) bonds | Not Available |
| 1598 | Outer membrane N-deacetylase | *Pseudomonas aestus* | | U6ZQ67 | -- | 24285649 | Hydrolase activity, acting on carbon-nitrogen (but not peptide) bonds | Not Available |
| 1599 | Poly-beta-1, 6-N-acetyl-D-glucosamine biosynthesis protein PgaD | *Acinetobacter spp. ABNIH27* | | A0A2S4T987 | -- | 29437920 | Not Available | Not Available |
| 1600 | Poly-beta-1, 6-N-acetyl-D-glucosamine synthase (Poly-beta-1, 6-GlcNAc synthase) (EC 2.4.1.-) | *Pseudomonas aestus* | | U6ZPI1 | -- | 24285649 | Acetyl glucosaminyl transferase activity | Not Available |
| 1601 | Biofilm PGA synthesis protein PgaD | *Pseudomonas aestus* | | U6ZVB6 | -- | 24285649 | Not Available | Not Available |
| 1602 | Poly-beta-1, 6-N-acetyl-D-glucosamine biosynthesis protein PgaD | *Lentisphaerae bacterium GWF2_52_8* | | A0A1G0ZGL6 | -- | 27774985 | Not Available | Not Available |
| 1603 | Poly-beta-1, 6-N-acetyl-D-glucosamine N-deacetylase PgaB | *Acinetobacter spp. ABNIH27* | | A0A2S4TAA7 | -- | 29437920 | Hydrolase activity, acting on carbon-nitrogen (but not peptide) bonds | Not Available |
| 1604 | Poly-beta-1, 6-N-acetyl-D-glucosamine N-deacetylase PgaB | *Lentisphaerae bacterium* | | A0A7V4HHC5 | -- | 31911466 | Hydrolase activity, acting on carbon-nitrogen (but not peptide) bonds | Not Available |
| 1605 | Poly-beta-1, 6-N-acetyl-D-glucosamine biosynthesis protein PgaD | *Lentisphaerae bacterium GWF2_49_21* | | A0A1G0Y752 | -- | 27774985 | Not Available | Not Available |
| 1606 | Poly-beta-1, 6-N-acetyl-D-glucosamine biosynthesis protein PgaD | *Lentisphaerae bacterium* | | A0A7C5AG76 | -- | 31911466 | Not Available | Not Available |
| 1607 | Poly-beta-1, 6-N-acetyl-D-glucosamine biosynthesis protein PgaD | *Escherichia spp. ESNIH1* | | A0A2S4QVY0 | -- | 29437920 | Not Available | Not Available |
| 1608 | Poly-beta-1, 6-N-acetyl-D-glucosamine synthase (Poly-beta-1, 6-GlcNAc synthase) (EC 2.4.1.-) | *Lentisphaerae bacterium GWF2_49_21* | | A0A1G0Y743 | -- | 27774985 | Acetyl glucosaminyl transferase activity | Not Available |
| 1609 | Poly-beta-1, 6-N-acetyl-D-glucosamine synthase (Poly-beta-1, 6-GlcNAc synthase) (EC 2.4.1.-) | *Escherichia spp. ESNIH1* | | A0A2S4QVY8 | -- | 29437920 | Acetyl glucosaminyl transferase activity | Not Available |
| 1610 | Uncharacterized protein | *Lentisphaerae bacterium* | | A0A7X9AUX7 | -- | 32123542 | Not Available | Not Available |
| 1611 | Poly-beta-1, 6-N-acetyl-D-glucosamine synthase (Poly-beta-1, 6-GlcNAc synthase) (EC 2.4.1.-) | *Acinetobacter spp. ABNIH27* | | A0A2S4T939 | -- | 29437920 | Acetyl glucosaminyl transferase activity | Not Available |
| 1612 | Poly-beta-1, 6-N-acetyl-D-glucosamine N-deacetylase PgaB | *Acinetobacter spp. ABNIH27* | | A0A2S4T902 | -- | 29437920 | Hydrolase activity, acting on carbon-nitrogen (but not peptide) bonds | Not Available |
| 1613 | Poly-beta-1, 6-N-acetyl-D-glucosamine synthase (Poly-beta-1, 6-GlcNAc synthase) (EC 2.4.1.-) | *Acinetobacter spp. ABNIH27* | | A0A2S4TAB3 | -- | 29437920 | Acetyl glucosaminyl transferase activity | Not Available |
| 1614 | Poly-beta-1, 6-N-acetyl-D-glucosamine N-deacetylase PgaB | *Lentisphaerae bacterium GWF2_49_21* | | A0A1G0Y7B1 | -- | 27774985 | Hydrolase activity, acting on carbon-nitrogen (but not peptide) bonds | Not Available |
| 1615 | Poly-beta-1, 6-N-acetyl-D-glucosamine biosynthesis protein PgaD | *Acinetobacter spp. ABNIH27* | | A0A2S4TAC3 | -- | 29437920 | Not Available | Not Available |
| 1616 | Poly-beta-1, 6-N-acetyl-D-glucosamine biosynthesis protein PgaD | *Lentisphaerae bacterium* | | A0A7V4HI93 | -- | 31911466 | Not Available | Not Available |
| 1617 | Type II toxin-antitoxin system MqsR family toxin | *Lentisphaerae bacterium* | | A0A7X7M3S1 | -- | 32123542 | Not Available | Not Available |
| 1618 | Response regulator | *Xanthomonas campestris pv. campestris* | Strain ATCC 33913 / DSM 3586 / NCPPB 528 / LMG 568 / P 25 | Q8P4H8 | -- | 12024217 | Diguanylate cyclase activity | Not Available |
| 1619 | Biofilm PGA synthesis lipoprotein PgaB | *Aggregatibacter actinomycetemcomitans RhAA1* | | H0KC42 | -- | 22328766 | Hydrolase activity, acting on carbon-nitrogen (but not peptide) bonds | Not Available |
| 1620 | Surface composition regulator | *Cronobacter condimenti 1330* | | K8ACP6 | -- | 23166675 | Not Available | Not Available |
| 1621 | Sensor histidinekinase | *Xanthomonas campestris pv. campestris* | Strain ATCC 33913 / DSM 3586 / NCPPB 528 / LMG 568 / P 25 | Q8PAN4 | -- | 12024217 | Diguanylate cyclase activity, kinase activity | Not Available |
| 1622 | GGDEF domain-containing protein | *Xanthomonas campestris pv. campestris* | Strain ATCC 33913 / DSM 3586 / NCPPB 528 / LMG 568 / P 25 | Q8PDD3 | -- | 12024217 | Diguanylate cyclase activity | Not Available |
| 1623 | Uncharacterized protein | *Sulfuricurvum kujiense* | Strain ATCC BAA-921 / DSM 16994 / JCM 11577 / YK-1 | E4U029 | -- | 22675602 | Not Available | Not Available |
| 1624 | Biofilm formation protein | *Bacillus licheniformis CG-B52* | | T5HTA9 | -- | 27174263 | Not Available | Not Available |
| 1625 | Type II toxin-antitoxin system MqsR family toxin | *Sinorhizobium terangae* | | A0A6N7LAM3 | -- | 23425606 | Not Available | Not Available |
| 1626 | Sensor histidinekinase | *Xanthomonas campestris pv. campestris* | Strain ATCC 33913 / DSM 3586 / NCPPB 528 / LMG 568 / P 25 | Q8PB32 | -- | 12024217 | Diguanylate cyclase activity, kinase activity | Not Available |
| 1627 | GGDEF domain-containing protein | *Xanthomonas campestris pv. campestris* | Strain ATCC 33913 / DSM 3586 / NCPPB 528 / LMG 568 / P 25 | Q8PAN6 | -- | 12024217 | Diguanylate cyclase activity | Not Available |

| Target ID | Bacterial Protein Name | Organism Name | Strain | UniProt ID | PDB ID | PubMed ID | Classification as per UniProt GO Molecular Function | Classification of Function as per PDB |
|---|---|---|---|---|---|---|---|---|
| 1628 | GGDEF family protein | *Xanthomonas campestris pv. campestris* | Strain ATCC 33913 / DSM 3586 / NCPPB 528 / LMG 568 / P 25 | Q8P9F2 | -- | 12024217 | Diguanylate cyclase activity | Not Available |
| 1629 | Polysaccharide deacetylase | *Sulfuricurvum kujiense* | Strain ATCC BAA-921 / DSM 16994 / JCM 11577 / YK-1 | E4U031 | -- | 22675602 | Hydrolase activity, acting on carbon-nitrogen (but not peptide) bonds | Not Available |
| 1630 | GGDEF domain-containing protein | *Xanthomonas campestris pv. campestris* | Strain ATCC 33913 / DSM 3586 / NCPPB 528 / LMG 568 / P 25 | Q8P502 | -- | 12024217 | Diguanylate cyclase activity | Not Available |
| 1631 | Uncharacterized protein | *Methylotenera versatilis* | Strain 301 | D7DJ79 | -- | 21622745 | Not Available | Not Available |
| 1632 | Poly-beta-1, 6-N-acetyl-D-glucosamine synthase (Poly-beta-1, 6-GlcNAc synthase) (EC 2.4.1.-) | *Methylotenera versatilis* | Strain 301 | D7DK50 | -- | 21622745 | Acetyl glucosaminyl transferase activity | Not Available |
| 1633 | Polysaccharide deacetylase | *Methylotenera versatilis* | Strain 301 | D7DJ81 | -- | 21622745 | Hydrolase activity, acting on carbon-nitrogen (but not peptide) bonds | Not Available |
| 1634 | GGDEF family protein | *Xanthomonas campestris pv. campestris* | Strain ATCC 33913 / DSM 3586 / NCPPB 528 / LMG 568 / P 25 | Q8P528 | -- | 12024217 | Diguanylate cyclase activity | Not Available |
| 1635 | Uncharacterized protein | *Anaeromyxobacter spp. RBG_16_69_14* | | A0A1F3C6Q7 | -- | 27774985 | Not Available | Not Available |
| 1636 | Poly-beta-1, 6-N-acetyl-D-glucosamine synthase (Poly-beta-1, 6-GlcNAc synthase) (EC 2.4.1.-) | *Pseudomonas brassicacearum* | Strain NFM421 | F2KD33 | -- | 21515771 | Acetyl glucosaminyl transferase activity | Not Available |
| 1637 | Poly-beta-1, 6-N-acetyl-D-glucosamine N-deacetylase | *Escherichia spp. ESNIH1* | | A0A2S4QVZ5 | -- | 29437920 | Hydrolase activity, acting on carbon-nitrogen (but not peptide) bonds | Not Available |
| 1638 | Type II toxin-antitoxin system MqsR family toxin | *Verrucomicrobiales bacterium* | | A0A354TEC8 | -- | 30148503 | Not Available | Not Available |
| 1639 | Uncharacterized protein | *Candidatus Rokubacteria bacterium GWC2_70_16* | | A0A1F7LKG1 | -- | 27774985 | Not Available | Not Available |
| 1640 | Poly-beta-1, 6-N-acetyl-D-glucosamine biosynthesis protein PgaD | *Clostridium spp.* | | A0A7X7XZA3 | -- | 32123542 | Not Available | Not Available |
| 1641 | NodB homology domain-containing protein | *Geoalkalibacter subterraneus* | | A0A0B5FGS1 | -- | 25767222 | Hydrolase activity, acting on carbon-nitrogen (but not peptide) bonds | Not Available |
| 1642 | Poly-beta-1, 6-N-acetyl-D-glucosamine synthase (Poly-beta-1, 6-GlcNAc synthase) (EC 2.4.1.-) | *Klebsiella pneumoniae subspp. pneumoniae DSM 30104 = JCM 1662 = NBRC 14940* | | J2XCJ6 | -- | 23012294 | Acetyl glucosaminyl transferase activity | Not Available |
| 1643 | Poly-beta-1, 6-N-acetyl-D-glucosamine N-deacetylase PgaB (EC 3.5.1.-) | *Lysobacter capsici* | | A0A0S2FSL6 | -- | 26597042 | Hydrolase activity, acting on carbon-nitrogen (but not peptide) bonds | Not Available |
| 1644 | Uncharacterized protein | *Geoalkalibacter subterraneus* | | A0A0B5FT76 | -- | 25767222 | Not Available | Not Available |
| 1645 | Poly-beta-1, 6-N-acetyl-D-glucosamine biosynthesis protein PgaD | *Lysobacter capsici* | | A0A0S2FSG4 | -- | 26597042 | Not Available | Not Available |
| 1646 | Uncharacterized protein | *Romboutsia maritimum* | | A0A371IQ26 | -- | 29025937 | Not Available | Not Available |
| 1647 | Type II toxin-antitoxin system MqsR family toxin | *Dehalococcoidia bacterium* | | A0A7X7GXM7 | -- | 32123542 | Not Available | Not Available |
| 1648 | Poly-beta-1, 6-N-acetyl-D-glucosamine synthase (Poly-beta-1, 6-GlcNAc synthase) (EC 2.4.1.-) | *Lysobacter capsici* | | A0A0S2FSU4 | -- | 26597042 | Acetyl glucosaminyl transferase activity | Not Available |
| 1649 | Poly-beta-1, 6-N-acetyl-D-glucosamine synthase (Poly-beta-1, 6-GlcNAc synthase) (EC 2.4.1.-) | *Geoalkalibacter subterraneus* | | A0A0B5FUS7 | -- | 25767222 | Acetyl glucosaminyl transferase activity | Not Available |
| 1650 | Surface composition regulator | *Serratia inhibens PRI-2C* | | A0A1B1KT89 | -- | 22815440 | Not Available | Not Available |
| 1651 | Uncharacterized protein | *Strain Marseille Minibacterium maSilienSiS* | | J3G480 | -- | 23045501 | Not Available | Not Available |
| 1652 | Hemin storage protein | *Janthinobacterium spp.* | Strain Marseille Minibacterium masiliensis | A6T1E1 | -- | 17722982 | Hydrolase activity, acting on carbon-nitrogen (but not peptide) bonds | Not Available |
| 1653 | Poly-beta-1, 6-N-acetyl-D-glucosamine synthase (Poly-beta-1, 6-GlcNAc synthase) (EC 2.4.1.-) | *Janthinobacterium spp.* | v | A6T1E0 | -- | 17722982 | Acetyl glucosaminyl transferase activity | Not Available |
| 1654 | Uncharacterized protein | *Dehalococcoidia bacterium* | | A0A7C2WC69 | -- | 32372496 | Not Available | Not Available |
| 1655 | Poly-beta-1, 6-N-acetyl-D-glucosamine N-deacetylase PgaB | *Proteobacteria bacterium* | | A0A6H9I2Q0 | -- | 31478060 | Hydrolase activity, acting on carbon-nitrogen (but not peptide) bonds | Not Available |
| 1656 | Outer membrane N-deacetylase | *Klebsiella pneumoniae subspp. pneumoniae DSM 30104 = JCM 1662 = NBRC 14940* | | J2DQS3 | -- | 23012294 | Hydrolase activity, acting on carbon-nitrogen (but not peptide) bonds | Not Available |
| 1657 | Poly-beta-1, 6-N-acetyl-D-glucosamine biosynthesis protein PgaD | *Proteobacteria bacterium* | | A0A6H9ICA2 | -- | 31478060 | Not Available | Not Available |
| 1658 | Poly-beta-1, 6-N-acetyl-D-glucosamine synthase (Poly-beta-1, 6-GlcNAc synthase) (EC 2.4.1.-) | *Proteobacteria bacterium* | | A0A6H9IFG5 | -- | 31478060 | Acetyl glucosaminyl transferase activity | Not Available |
| 1659 | Uncharacterized conserved protein | *Janthinobacterium spp.* | Strain Marseille Minibacterium masiliensis | A6T1D9 | -- | 17722982 | Not Available | Not Available |
| 1660 | Biofilm PGA synthesis lipoprotein PgaB | *Actinobacillus pleuropneumoniae serovar 10* | Str. D13039 | E0F777 | -- | 20802045 | Hydrolase activity, acting on carbon-nitrogen (but not peptide) bonds | Not Available |
| 1661 | Biofilm PGA synthesis auxiliary protein PgaD | *Burkholderia pyrrocinia* | Pseudomonas pyrrocinia | A0A0G3WZX8 | -- | 26150017 | Not Available | Not Available |
| 1662 | Hemin storage protein | *Burkholderia pyrrocinia* | Pseudomonas pyrrocinia | A0A0G3X357 | -- | 26150017 | Hydrolase activity, acting on carbon-nitrogen (but not peptide) bonds | Not Available |
| 1663 | Uncharacterized protein | *Tistrella mobilis* | Strain KA081020-065 | I3TVX0 | -- | 22458477 | Not Available | Not Available |
| 1664 | Uncharacterized protein | *Acinetobacter oleivorans* | Strain JCM 16667 / KCTC 23045 / DR1 | D8JE02 | -- | 20639327 | Not Available | Not Available |
| 1665 | Poly-beta-1, 6-N-acetyl-D-glucosamine synthase (Poly-beta-1, 6-GlcNAc synthase) (EC 2.4.1.-) | *Deltaproteobacteria bacterium* | | A0A7V3KXN3 | -- | 31911466 | Acetyl glucosaminyl transferase activity | Not Available |
| 1666 | Poly-beta-1, 6-N-acetyl-D-glucosamine synthase (Poly-beta-1, 6-GlcNAc synthase) (EC 2.4.1.-) | *Shigella flexneri serotype 5b* | Strain 8401 | Q0T614 | -- | 16822325 | Acetyl glucosaminyl transferase activity | Not Available |
| 1667 | Poly-beta-1, 6-N-acetyl-D-glucosamine biosynthesis protein PgaD | *Deltaproteobacteria bacterium* | | A0A7X6PVS7 | -- | 32123542 | Not Available | Not Available |
| 1668 | Poly-beta-1, 6-N-acetyl-D-glucosamine N-deacetylase PgaB | *Deltaproteobacteria bacterium* | | A0A7V3NYI2 | -- | 31911466 | Hydrolase activity, acting on carbon-nitrogen (but not peptide) bonds | Not Available |
| 1669 | Uncharacterized protein | *Shigella flexneri serotype 5b* | Strain 8401 | Q0T615 | -- | 16822325 | Not Available | Not Available |
| 1670 | Uncharacterized protein | *Deltaproteobacteria bacterium* | | A0A7V0QJM4 | -- | 31911466 | Not Available | Not Available |
| 1671 | Type II toxin-antitoxin system MqsR family toxin | *Thermodesulfatator atlanticus* | | A0A7V5NYK0 | -- | 31911466 | Not Available | Not Available |
| 1672 | Outer membrane N-deacetylase | *Pseudomonas fluorescens LMG 5329* | | A0A0A1Z5R3 | -- | 23887909 | Hydrolase activity, acting on carbon-nitrogen (but not peptide) bonds | Not Available |

| Target ID | Bacterial Protein Name | Organism Name | Strain | UniProt ID | PDB ID | PubMed ID | Classification as per UniProt GO Molecular Function | Classification of Function as per PDB |
|---|---|---|---|---|---|---|---|---|
| 1673 | Poly-beta-1, 6-N-acetyl-D-glucosamine synthase (Poly-beta-1, 6-GlcNAc synthase) (EC 2.4.1.-) | *Pseudomonas fluorescens LMG 5329* | | A0A0A1Z9D1 | -- | 23887909 | Acetyl glucosaminyl transferase activity | Not Available |
| 1674 | Biofilm PGA synthesis protein PgaD | *Pseudomonas fluorescens LMG 5329* | | A0A0A1Z9Y2 | -- | 23887909 | Not Available | Not Available |
| 1675 | Poly-beta-1, 6-N-acetyl-D-glucosamine N-deacetylase PgaB | *Deltaproteobacteria bacterium* | | A0A7V0JQ66 | -- | 31911466 | Hydrolase activity, acting on carbon-nitrogen (but not peptide) bonds | Not Available |
| 1676 | Poly-beta-1, 6-N-acetyl-D-glucosamine N-deacetylase PgaB (Fragment) | *Deltaproteobacteria bacterium* | | A0A7X6PV65 | -- | 32123542 | Hydrolase activity, acting on carbon-nitrogen (but not peptide) bonds | Not Available |
| 1677 | Surface composition regulator | *Salmonella enterica subspp. enterica serovar* | WandSworth Str. A4-580 | G5SH05 | -- | 21859443 | Not Available | Not Available |
| 1678 | Motility quorum-sensing regulator MqsR | *Pseudomonas fluorescens LMG 5329* | | A0A0A1YWG5 | -- | 23887909 | Not Available | Not Available |
| 1679 | Uncharacterized protein | *Companilactobacillus farciminis* | | A0A4V3A398 | -- | 30830251 | Not Available | Not Available |
| 1680 | Poly-beta-1, 6-N-acetyl-D-glucosamine synthase (Poly-beta-1, 6-GlcNAc synthase) (EC 2.4.1.-) | *Deltaproteobacteria bacterium* | | A0A7V0JR38 | -- | 31911466 | Acetyl glucosaminyl transferase activity | Not Available |
| 1681 | Poly-beta-1, 6-N-acetyl-D-glucosamine synthase (Poly-beta-1, 6-GlcNAc synthase) (EC 2.4.1.-) | *Deltaproteobacteria bacterium* | | A0A7C1B4X9 | -- | 31911466 | Acetyl glucosaminyl transferase activity | Not Available |
| 1682 | Poly-beta-1, 6-N-acetyl-D-glucosamine synthase (Poly-beta-1, 6-GlcNAc synthase) (EC 2.4.1.-) | *Stenotrophomonas maltophilia* | Pseudomonas maltophilia Xanthomonas maltophilia | A0A7Z1RIE6 | -- | 29170656 | Acetyl glucosaminyl transferase activity | Not Available |
| 1683 | Surface composition regulator | *Salmonella enterica subspp. enterica serovar Rubislaw str. A4-653* | | G5QPX3 | -- | 21859443 | Not Available | Not Available |
| 1684 | Poly-beta-1, 6-N-acetyl-D-glucosamine biosynthesis protein PgaD | *Stenotrophomonas maltophilia* | Pseudomonas maltophilia Xanthomonas maltophilia | A0A7Z1RIY8 | -- | 29170656 | Not Available | Not Available |
| 1685 | Poly-beta-1, 6-N-acetyl-D-glucosamine biosynthesis protein PgaD | *Deltaproteobacteria bacterium* | | A0A7C1BIY6 | -- | 31911466 | Not Available | Not Available |
| 1686 | Uncharacterized protein | *Polaromona spp.* | Strain JS666 / ATCC BAA-500 | Q11Z97 | -- | 18723656 | Not Available | Not Available |
| 1687 | Poly-beta-1, 6-N-acetyl-D-glucosamine biosynthesis protein PgaD | *Stenotrophomonas maltophilia* | Pseudomonas maltophilia Xanthomonas maltophilia | A0A2Y9U8G9 | -- | 29170656 | Not Available | Not Available |
| 1688 | Poly-beta-1, 6-N-acetyl-D-glucosamine N-deacetylase PgaB | *Stenotrophomonas maltophilia* | Pseudomonas maltophilia Xanthomonas maltophilia | A0A7Z1RJ06 | -- | 29170656 | Hydrolase activity, acting on carbon-nitrogen (but not peptide) bonds | Not Available |
| 1689 | Poly-beta-1, 6-N-acetyl-D-glucosamine biosynthesis protein PgaD | *Chromobacterium spp. Panama* | | A0A2T5NK43 | -- | 29670144 | Not Available | Not Available |
| 1690 | Poly-beta-1, 6-N-acetyl-D-glucosamine N-deacetylase PgaB | *Deltaproteobacteria bacterium* | | A0A7C1B8G0 | -- | 31911466 | Hydrolase activity, acting on carbon-nitrogen (but not peptide) bonds | Not Available |
| 1691 | Type II toxin-antitoxin system MqsR family toxin | *Deltaproteobacteria bacterium* | | A0A3A5ARM9 | -- | 28644444 | Not Available | Not Available |
| 1692 | Poly-beta-1, 6-N-acetyl-D-glucosamine synthase (Poly-beta-1, 6-GlcNAc synthase) (EC 2.4.1.-) | *Escherichia coli O55:H7* | Strain CB9615 / EPEC | D3QPZ6 | -- | 20090843 | Acetyl glucosaminyl transferase activity | Not Available |
| 1693 | Poly-beta-1, 6-N-acetyl-D-glucosamine synthase (Poly-beta-1, 6-GlcNAc synthase) (EC 2.4.1.-) | *Chromobacterium spp. Panama* | | A0A2T5NQP9 | -- | 29670144 | Acetyl glucosaminyl transferase activity | Not Available |
| 1694 | Poly-beta-1, 6-N-acetyl-D-glucosamine N-deacetylase PgaB | *Chromobacterium spp. Panama* | | A0A2T5NK52 | -- | 29670144 | Hydrolase activity, acting on carbon-nitrogen (but not peptide) bonds | Not Available |
| 1695 | Biofilm PGA synthesis protein pgaD | *Escherichia coli O55:H7* | Strain CB9615 / EPEC | D3QPZ5 | -- | 20090843 | Not Available | Not Available |
| 1696 | Surface composition regulator | *Escherichia coli O55:H7* | Strain CB9615 / EPEC | D3QRZ4 | -- | 20090843 | Not Available | Not Available |
| 1697 | Bll4347 protein | *Bradyrhizobium diazoefficiens* | Strain JCM 10833 / BCRC 13528 / IAM 13628 / NBRC 14792 / USDA 110 | Q89M46 | -- | 12597275 | Diguanylate cyclase activity | Not Available |
| 1698 | Bll7340 protein | *Bradyrhizobium diazoefficiens* | Strain JCM 10833 / BCRC 13528 / IAM 13628 / NBRC 14792 / USDA 110 | Q89DU8 | -- | 12597275 | Diguanylate cyclase activity | Not Available |
| 1699 | Blr2401 protein | *Bradyrhizobium diazoefficiens* | Strain JCM 10833 / BCRC 13528 / IAM 13628 / NBRC 14792 / USDA 110 | Q89SJ9 | -- | 12597275 | Diguanylate cyclase activity | Not Available |
| 1700 | Bll3041 protein | *Bradyrhizobium diazoefficiens* | Strain JCM 10833 / BCRC 13528 / IAM 13628 / NBRC 14792 / USDA 110 | Q89QT3 | -- | 12597275 | Diguanylate cyclase activity | Not Available |
| 1701 | Surface composition regulator | *Citrobacter rodentium* | Strain ICC168 Citrobacter freundii biotype 4280 | D2TRP2 | -- | 19897651 | Not Available | Not Available |
| 1702 | Poly-beta-1, 6-N-acetyl-D-glucosamine biosynthesis protein PgaD | *Stenotrophomonas spp. SAU14A_naIMI4_5* | | A0A3S7KI35 | -- | 29765358 | Not Available | Not Available |
| 1703 | Blr0624 protein | *Bradyrhizobium diazoefficiens* | Strain JCM 10833 / BCRC 13528 / IAM 13628 / NBRC 14792 / USDA 110 | Q89WQ4 | -- | 12597275 | Diguanylate cyclase activity | Not Available |
| 1704 | Bll1362 protein | *Bradyrhizobium diazoefficiens* | Strain JCM 10833 / BCRC 13528 / IAM 13628 / NBRC 14792 / USDA 110 | Q89UP9 | -- | 12597275 | Diguanylate cyclase activity | Not Available |
| 1705 | Bll2326 protein | *Bradyrhizobium diazoefficiens* | Strain JCM 10833 / BCRC 13528 / IAM 13628 / NBRC 14792 / USDA 110 | Q89SS4 | -- | 12597275 | Diguanylate cyclase activity | Not Available |
| 1706 | Poly-beta-1, 6-N-acetyl-D-glucosamine biosynthesis protein PgaD | *Stenotrophomonas spp. SAU14A_naIMI4_5* | | A0A3S7KGX6 | -- | 29765358 | Not Available | Not Available |
| 1707 | Bll4961 protein | *Bradyrhizobium diazoefficiens* | Strain JCM 10833 / BCRC 13528 / IAM 13628 / NBRC 14792 / USDA 110 | Q89KE6 | -- | 12597275 | Diguanylate cyclase activity | Not Available |
| 1708 | Motility quorum-sensing regulator MqsR | *Gallionellales bacterium RIFCSPLOWO2_02_58_13* | | A0A1G0DUP1 | -- | 27774985 | Not Available | Not Available |
| 1709 | Poly-beta-1, 6-N-acetyl-D-glucosamine biosynthesis protein PgaD | *Xenorhabdus spp. KJ12.1* | | A0A2D0LA77 | -- | 28993611 | Not Available | Not Available |
| 1710 | Uncharacterized protein | *Gallionellales bacterium RIFCSPLOWO2_02_58_13* | | A0A1G0DV73 | -- | 27774985 | Not Available | Not Available |

| Target ID | Bacterial Protein Name | Organism Name | Strain | UniProt ID | PDB ID | PubMed ID | Classification as per UniProt GO Molecular Function | Classification of Function as per PDB |
|---|---|---|---|---|---|---|---|---|
| 1711 | Poly-beta-1, 6-N-acetyl-D-glucosamine N-deacetylase PgaB | *Stenotrophomonas maltophilia* | Pseudomonas maltophilia Xanthomonas maltophilia | A0A2J0SHH4 | -- | 29170656 | Hydrolase activity, acting on carbon-nitrogen (but not peptide) bonds | Not Available |
| 1712 | Poly-beta-1, 6-N-acetyl-D-glucosamine synthase (Poly-beta-1, 6-GlcNAc synthase) (EC 2.4.1.-) | *Xenorhabdus spp. KJ12.1* | | A0A2D0LAE0 | -- | 28993611 | Acetyl glucosaminyl transferase activity | Not Available |
| 1713 | Poly-beta-1, 6-N-acetyl-D-glucosamine biosynthesis protein PgaD | *Stenotrophomonas maltophilia* | Pseudomonas maltophilia Xanthomonas maltophilia | A0A2J0TWJ3 | -- | 29170656 | Not Available | Not Available |
| 1714 | Poly-beta-1, 6-N-acetyl-D-glucosamine N-deacetylase PgaB | *Stenotrophomonas maltophilia* | Pseudomonas maltophilia Xanthomonas maltophilia | A0A2J0TWJ1 | -- | 29170656 | Hydrolase activity, acting on carbon-nitrogen (but not peptide) bonds | Not Available |
| 1715 | Poly-beta-1, 6-N-acetyl-D-glucosamine N-deacetylase PgaB | *Stenotrophomonas maltophilia* | Pseudomonas maltophilia Xanthomonas maltophilia | A0A2J0UEW1 | -- | 29170656 | Hydrolase activity, acting on carbon-nitrogen (but not peptide) bonds | Not Available |
| 1716 | Poly-beta-1, 6-N-acetyl-D-glucosamine synthase (Poly-beta-1, 6-GlcNAc synthase) (EC 2.4.1.-) | *Pseudomonas fluorescens Q8r1-96* | | I4L1A4 | -- | 22792073 | Acetyl glucosaminyl transferase activity | Not Available |
| 1717 | Surface composition regulator | *Salmonella enterica subspp. salamae serovar* | 42:r:- | A0A731NDC1 | -- | 30286803 | Not Available | Not Available |
| 1718 | Acyl-homoserine-lactone synthase (EC 2.3.1.184) (Autoinducer synthesis protein) | *Serratia plymuthica* | | C4PDX0 | -- | 21284858 | N-acyl homoserine lactone synthase activity | Not Available |
| 1719 | SplR | *Serratia plymuthica* | | C4PDW8 | -- | 21284858 | DNA binding | Not Available |
| 1720 | Surface composition regulator | *Salmonella enterica subspp. houtenae serovar* | 21:z4, z23:- | A0A752IW31 | -- | 30286803 | Not Available | Not Available |
| 1721 | IcaA (Fragment) | *Staphylococcus saprophyticus* | | Q83ZH7 | -- | 12957956 | Not Available | Not Available |
| 1722 | Putative transporter protein | *Salmonella typhimurium* | Strain LT2 / SGSC1412 / ATCC 700720 | Q8RSN8 | -- | 11260463; 11929533 | Not Available | Not Available |
| 1723 | Acyl-homoserine-lactone synthase (EC 2.3.1.184) (Autoinducer synthesis protein) | *Serratia plymuthica* | | C4PDW7 | -- | 21284858 | N-acyl homoserine lactone synthase activity | Not Available |
| 1724 | Uncharacterized protein | *Bordetella bronchiseptica* | Strain ATCC BAA-588 / NCTC 13252 / RB50 Alcaligenes bronchisepticus | A0A0H3LPJ8 | -- | 12910271 | Not Available | Not Available |
| 1725 | Surface composition regulator | *Salmonella enterica subspp. enterica serovar* | Duisburg | A0A5X0EQI5 | -- | 30286803 | Not Available | Not Available |
| 1726 | Surface composition regulator | *Salmonella typhisuis* | | A0A735MWU7 | -- | 30286803 | Not Available | Not Available |
| 1727 | Poly-beta-1, 6-N-acetyl-D-glucosamine synthase (Poly-beta-1, 6-GlcNAc synthase) (EC 2.4.1.-) | *Staphylococcus lugdunensis* | | A8DN18 | -- | 17635864 | Acetyl glucosaminyl transferase activity | Not Available |
| 1728 | Poly-beta-1, 6-N-acetyl-D-glucosamine synthase (Poly-beta-1, 6-GlcNAc synthase) (EC 2.4.1.-) | *Pseudomonas synxantha BG33R* | | I4KXU6 | -- | 22792073 | Acetyl glucosaminyl transferase activity | Not Available |
| 1729 | Surface composition regulator | *Salmonella enterica subspp. arizonae serovar* | 18:z4, z32:- | A0A737EZC4 | -- | 30286803 | Not Available | Not Available |
| 1730 | Uncharacterized protein | *Yersinia enterocolitica* | Type O:5, 27 Str. YE149 / 02 | A0A7U7FJ92 | -- | 24753568 | Not Available | Not Available |
| 1731 | Surface composition regulator | *Salmonella enterica subspp. enterica serovar* | Decatur | A0A6V9XHS2 | -- | 30286803 | Not Available | Not Available |
| 1732 | Poly-beta-1, 6-N-acetyl-D-glucosamine synthase (Poly-beta-1, 6-GlcNAc synthase) (EC 2.4.1.-) | *Xanthomonas hortorum pv. carotae* | Str. M081 | V7ZDU8 | -- | 21722296 | Acetyl glucosaminyl transferase activity | Not Available |
| 1733 | Surface composition regulator | *Salmonella enterica subspp. arizonae serovar* | 48:z4, z24:- | A0A738XIB5 | -- | 30286803 | Not Available | Not Available |
| 1734 | Poly-beta-1, 6-N-acetyl-D-glucosamine synthase (Poly-beta-1, 6-GlcNAc synthase) (EC 2.4.1.-) | *Yersinia enterocolitica* | Type O:5, 27 Str. YE149 / 02 | A0A7U7IS30 | -- | 24753568 | Acetyl glucosaminyl transferase activity | Not Available |
| 1735 | Haemin storage system, HmsF protein | *Yersinia enterocolitica* | Type O:5, 27 Str. YE149 / 02 | A0A7U7FN73 | -- | 24753568 | Hydrolase activity, acting on carbon-nitrogen (but not peptide) bonds | Not Available |
| 1736 | AI-2E family transporter | *Derxia gummosa DSM 723* | | A0A8B6XBG3 | -- | 16385049; 20559013 | Not Available | Not Available |
| 1737 | HmsF protein | *Xanthomonas hortorum pv. carotae* | Str. M081 | V7ZCS1 | -- | 21722296 | Hydrolase activity, acting on carbon-nitrogen (but not peptide) bonds | Not Available |
| 1738 | Surface composition regulator | *Salmonella enterica subspp. houtenae serovar* | 18:z36, z38:- | A0A729K7U1 | -- | 30286803 | Not Available | Not Available |
| 1739 | Poly-beta-1, 6-N-acetyl-D-glucosamine synthase (Poly-beta-1, 6-GlcNAc synthase) (EC 2.4.1.-) | *Yersinia enterocolitica* | Type O:3 Str. YE12 / 03 | A0A7U7IUF3 | -- | 24753568 | Acetyl glucosaminyl transferase activity | Not Available |
| 1740 | Uncharacterized protein | *uncultured bacterium* | | K1YAI8 | -- | 23019650 | Not Available | Not Available |
| 1741 | Haemin storage system, HmsF protein | *Yersinia enterocolitica* | Type O:3 Str. YE12 / 03 | A0A7U7ITX8 | -- | 24753568 | Hydrolase activity, acting on carbon-nitrogen (but not peptide) bonds | Not Available |
| 1742 | Biofilm PGA synthesis lipoprotein PgaB (EC 3.-.-.-) | *Pseudomonas synxantha BG33R* | | I4KSD4 | -- | 22792073 | Hydrolase activity, acting on carbon-nitrogen (but not peptide) bonds | Not Available |
| 1743 | Haemin storage system, HmsS protein | *Yersinia enterocolitica* | Type O:3 Str. YE12 / 03 | A0A7U7IUU3 | -- | 24753568 | Not Available | Not Available |
| 1744 | Surface composition regulator | *Yersinia enterocolitica* | Type O:3 Str. YE12 / 03 | A0A7U7ITX3 | -- | 24753568 | DNA binding, DNA-directed DNA polymerase activity | Not Available |
| 1745 | Surface composition regulator | *Salmonella enterica subspp. salamae serovar* | 42:b:1, 5 | A0A737IX99 | -- | 30286803 | Not Available | Not Available |
| 1746 | S-ribosylhomocysteine lyase (EC 4.4.1.21) (AI-2 synthesis protein) (Autoinducer-2 production protein LuxS) | *Eikenella corrodens* | | Q1HAV9 | -- | 17027872 | Iron ion binding, S-ribosylhomocysteine lyase activity | Not Available |
| 1747 | Putative biofilm PGA synthesis protein PgaD | *Pseudomonas synxantha BG33R* | | I4KVN4 | -- | 22792073 | Not Available | Not Available |
| 1748 | Motility quorum-sensing regulator | *uncultured bacterium* | | K1YWV6 | -- | 23019650 | Not Available | Not Available |
| 1749 | Surface composition regulator | *Salmonella enterica subspp. enterica serovar* | Saintpaul | A0A1S0ZD14 | -- | 30286803 | Not Available | Not Available |
| 1750 | Uncharacterized protein | *Yersinia enterocolitica* | Type O:3 Str. YE12 / 03 | A0A7U7FV97 | -- | 24753568 | Not Available | Not Available |
| 1751 | Surface composition regulator | *Salmonella muenchen* | | A0A3T3ENA5 | -- | 30286803; 30902850 | Not Available | Not Available |
| 1752 | Surface composition regulator | *Salmonella enterica subspp. enterica serovar* | Cerro | A0A2T9QB58 | -- | 30286803; 31527278 | Not Available | Not Available |
| 1753 | Poly-beta-1, 6-N-acetyl-D-glucosamine synthase (Poly-beta-1, 6-GlcNAc synthase) (EC 2.4.1.-) | *Pseudomonas fluorescens* | | A0A0N9WUK7 | -- | 29769716 | Acetyl glucosaminyl transferase activity | Not Available |
| 1754 | Surface composition regulator | *Salmonella enterica subspp. enterica serovar* | Carrau | A0A752RN04 | -- | 30286803 | Not Available | Not Available |
| 1755 | PpuA (Fragment) | *Pseudomonas putida* | Arthrobacter siderocapsulatus | Q3Y8H5 | -- | 16585751 | Not Available | Not Available |
| 1756 | PsoB | *Pseudomonas putida* | Arthrobacter siderocapsulatus | A8MN37 | -- | 16585751; 18599835 | Catalytic activity, phosphopantetheine binding | Not Available |
| 1757 | IcaA (Fragment) | *Staphylococcus epidermidis* | | Q83ZH6 | -- | 12957956 | Not Available | Not Available |

| Target ID | Bacterial Protein Name | Organism Name | Strain | UniProt ID | PDB ID | PubMed ID | Classification as per UniProt GO Molecular Function | Classification of Function as per PDB |
|---|---|---|---|---|---|---|---|---|
| 1758 | Surface composition regulator | *Salmonella enterica subspp. enterica serovar* | Abortusovis | A0A738AI74 | -- | 30286803 | Not Available | Not Available |
| 1759 | NodB homology domain-containing protein | *Escherichia coli* | | C3TEU7 | -- | 19439656 | Hydrolase activity, acting on carbon-nitrogen (but not peptide) bonds | Not Available |
| 1760 | IcaA (Fragment) | *Macrococcus caseolyticus* | | Q83ZI3 | -- | 12957956 | Not Available | Not Available |
| 1761 | Surface composition regulator | *Salmonella enterica subspp. VII serovar* | 40:z4, z24:[z39] | A0A731XYY4 | -- | 30286803 | Not Available | Not Available |
| 1762 | Fimbrial protein FimH (Fragment) | *Escherichia coli* | | A0A075Q3Y2 | -- | 25012525 | Not Available | Not Available |
| 1763 | PsoC | *Pseudomonas putida* | Arthrobacter siderocapsulatus | A8MN38 | -- | 16585751; 18599835 | Catalytic activity, phosphopantetheine binding | Not Available |
| 1764 | MacA | *Pseudomonas putida* | Arthrobacter siderocapsulatus | A8MN39 | -- | 16585751; 18599835 | Transmembrane transporter activity | Not Available |
| 1765 | Poly-beta-1, 6-N-acetyl-D-glucosamine synthase (Poly-beta-1, 6-GlcNAc synthase) (EC 2.4.1.-) | *Ralstonia syzygii R24* | | G3A8J6 | -- | 21931687 | Acetyl glucosaminyl transferase activity | Not Available |
| 1766 | Surface composition regulator | *Salmonella enterica subspp. enterica serovar* | Sendai | A0A737RJS4 | -- | 30286803 | Not Available | Not Available |
| 1767 | HmsS | *Xenorhabdus nematophila* | Achromobacter nematophilus | B3GN24 | -- | 18515487 | Not Available | Not Available |
| 1768 | Fimbrial protein FimF (Fragment) | *Escherichia coli* | | A0A075QDV6 | -- | 25012525 | Not Available | Not Available |
| 1769 | Uncharacterized protein | *Escherichia coli* | | C3TEV7 | -- | 19439656 | Not Available | Not Available |
| 1770 | Fimbrial protein FimG | *Escherichia coli* | | A0A075Q6V7 | -- | 25012525 | Not Available | Not Available |
| 1771 | HmsF | *Xenorhabdus nematophila* | Achromobacter nematophilus | B3GN22 | -- | 18515487 | Hydrolase activity, acting on carbon-nitrogen (but not peptide) bonds | Not Available |
| 1772 | Surface composition regulator | *Salmonella enterica subspp. salamae serovar* | 42:f, g, t:-- | A0A737HCW8 | -- | 30286803 | Not Available | Not Available |
| 1773 | NodB homology domain-containing protein | *Escherichia coli* | | C3TEU8 | -- | 19439656 | Hydrolase activity, acting on carbon-nitrogen (but not peptide) bonds | Not Available |
| 1774 | Poly-beta-1, 6-N-acetyl-D-glucosamine synthase (Poly-beta-1, 6-GlcNAc synthase) (EC 2.4.1.-) | *Escherichia coli* | | A0A6D1ART7 | -- | 16943019 | Acetyl glucosaminyl transferase activity | Not Available |
| 1775 | Uncharacterized protein | *Escherichia coli* | | C3TEV0 | -- | 19439656 | Hydrolase activity, acting on carbon-nitrogen (but not peptide) bonds | Not Available |
| 1776 | Surface composition regulator | *Escherichia coli* | | A0A7A2XRI7 | -- | 30286803 | Not Available | Not Available |
| 1777 | LuxR-like protein | *Pseudomonas putida* | Arthrobacter siderocapsulatus | A8MN41 | -- | 16585751; 18599835 | DNA binding | Not Available |
| 1778 | Ribosome maturation factor RimP | *Bacillus cereus* | | A0A063CC81 | -- | 25301660 | Not Available | Not Available |
| 1779 | S-ribosylhomocysteine lyase (EC 4.4.1.21) (AI-2 synthesis protein) (Autoinducer-2 production protein LuxS) | *Bacillus cereus* | | A0A063CL35 | -- | 25301660 | Iron ion binding, S-ribosylhomocysteine lyase activity | Not Available |
| 1780 | Surface composition regulator | *Pectobacterium parmentieri* | | A0A0H3I8L4 | -- | 23045508; 30326842 | DNA binding, DNA-directed DNA polymerase activity | Not Available |
| 1781 | S-ribosylhomocysteine lyase (EC 4.4.1.21) (AI-2 synthesis protein) (Autoinducer-2 production protein LuxS) | *Edwardsiella tarda* | Strain EIB202 | C5IWN6 | -- | 19283379; 19865481 | Iron ion binding, S-ribosylhomocysteine lyase activity | Not Available |
| 1782 | Poly-beta-1, 6-N-acetyl-D-glucosamine synthase (Poly-beta-1, 6-GlcNAc synthase) (EC 2.4.1.-) | *Pectobacterium parmentieri* | | A0A0H3ICJ6 | -- | 23045508; 30326842 | Acetyl glucosaminyl transferase activity | Not Available |
| 1783 | Poly-beta-1, 6-N-acetyl-D-glucosamine N-deacetylase (Poly-beta-1, 6-N-acetyl-D-glucosamine N-deacetylase PgaB) (EC 3.5.1.-) | *Pseudomonas panacis* | | A0A5M9I3H1 | -- | 32587583 | Hydrolase activity, acting on carbon-nitrogen (but not peptide) bonds | Not Available |
| 1784 | Motility quorum-sensing regulator MqsR | *Pseudomonas fluorescens* | | A0A0N9WTK9 | -- | 29769716 | Not Available | Not Available |
| 1785 | Ribosome maturation factor RimM | *Bacillus cereus* | | A0A063CCA6 | -- | 25301660 | ribosome binding | Not Available |
| 1786 | Poly-beta-1, 6-N-acetyl-D-glucosamine synthase (Poly-beta-1, 6-GlcNAc synthase) (EC 2.4.1.-) | *Staphylococcus aureus* | | W8TQA2 | -- | 26048971; 32562543; 33052925 | Acetyl glucosaminyl transferase activity | Not Available |
| 1787 | Poly-beta-1, 6-N-acetyl-D-glucosamine synthase (Poly-beta-1, 6-GlcNAc synthase) (EC 2.4.1.-) | *Trabulsiella odontotermitis* | | A0A0L0H1H9 | -- | 26162887 | Acetyl glucosaminyl transferase activity | Not Available |
| 1788 | Haemin storage system, HmsS protein | *Yersinia enterocolitica serotype O:8 / biotype 1B* | Strain NCTC 13174 / 8081 | A1JS92 | -- | 17173484 | Not Available | Not Available |
| 1789 | Poly-beta-1, 6-N-acetyl-D-glucosamine biosynthesis protein PgaD | *Lentisphaeria bacterium* | | A0A3D1KDG4 | -- | 30148503 | Not Available | Not Available |
| 1790 | Poly-beta-1, 6-N-acetyl-D-glucosamine synthase (Poly-beta-1, 6-GlcNAc synthase) (EC 2.4.1.-) | *Acinetobacter baylyi* | Strain ATCC 33305 / BD413 / ADP1 | Q6FDD8 | -- | 15514110 | Acetyl glucosaminyl transferase activity | Not Available |
| 1791 | Poly-beta-1, 6-N-acetyl-D-glucosamine synthase (Poly-beta-1, 6-GlcNAc synthase) (EC 2.4.1.-) | *Candidatus Muproteobacteria bacterium RBG_16_62_13* | | A0A1F6T856 | -- | 27774985 | Acetyl glucosaminyl transferase activity | Not Available |
| 1792 | Motility quorum-sensing regulator MqsR | *Gallionellales bacterium RIFCSPLOWO2_12_FULL_59_22* | | A0A1G0EM28 | -- | 27774985 | Not Available | Not Available |
| 1793 | Outer membrane N-deacetylase | *Trabulsiella odontotermitis* | | A0A0L0H2S7 | -- | 26162887 | Hydrolase activity, acting on carbon-nitrogen (but not peptide) bonds | Not Available |
| 1794 | Haemin storage system, HmsF protein | *Yersinia enterocolitica serotype O:8 / biotype 1B* | Strain NCTC 13174 / 8081 | A1JSA0 | -- | 17173484 | Hydrolase activity, acting on carbon-nitrogen (but not peptide) bonds | Not Available |
| 1795 | Hemin storage protein | *Trabulsiella odontotermitis* | | A0A0L0H272 | -- | 26162887 | Not Available | Not Available |
| 1796 | Surface composition regulator | *Trabulsiella odontotermitis* | | A0A0L0GTT1 | -- | 26162887 | Not Available | Not Available |
| 1797 | Uncharacterized protein | *Gallionellales bacterium RIFCSPLOWO2_12_FULL_59_22* | | A0A1G0EIH8 | -- | 27774985 | Not Available | Not Available |
| 1798 | Poly-beta-1, 6-N-acetyl-D-glucosamine N-deacetylase PgaB | *Lentisphaeria bacterium* | | A0A3526 | -- | 30148503 | Not Available | Not Available |
| 1799 | Poly-beta-1, 6-N-acetyl-D-glucosamine synthase (Poly-beta-1, 6-GlcNAc synthase) (EC 2.4.1.-) | *Lentisphaeria bacterium* | | A0A3525 | -- | 30148503 | Not Available | Not Available |
| 1800 | Poly-beta-1, 6-N-acetyl-D-glucosamine N-deacetylase PgaB | *Lentisphaeria bacterium* | | A0A3D1KEA7 | -- | 30148503 | Hydrolase activity, acting on carbon-nitrogen (but not peptide) bonds | Not Available |
| 1801 | Poly-beta-1, 6-N-acetyl-D-glucosamine biosynthesis protein PgaD | *Lentisphaeria bacterium* | | A0A3524 | -- | 30148503 | Not Available | Not Available |
| 1802 | Poly-beta-1, 6-N-acetyl-D-glucosamine synthase (Poly-beta-1, 6-GlcNAc synthase) (EC 2.4.1.-) | *Lentisphaeria bacterium* | | A0A3D1KDU7 | -- | 30148503 | Acetyl glucosaminyl transferase activity | Not Available |
| 1803 | Poly-beta-1, 6-N-acetyl-D-glucosamine synthase (Poly-beta-1, 6-GlcNAc synthase) (EC 2.4.1.-) | *Chromobacterium haemolyticum* | | A0A2T5P4D0 | -- | 29670144 | Acetyl glucosaminyl transferase activity | Not Available |
| 1804 | Poly-beta-1, 6-N-acetyl-D-glucosamine synthase (Poly-beta-1, 6-GlcNAc synthase) (EC 2.4.1.-) | *Actinobacillus suis H91-0380* | | K0GD63 | -- | 23144422 | Acetyl glucosaminyl transferase activity | Not Available |
| 1805 | Biofilm PGA synthesis lipoprotein PgaB | *Actinobacillus suis H91-0380* | | K0G5L0 | -- | 23144422 | Hydrolase activity, acting on carbon-nitrogen (but not peptide) bonds | Not Available |
| 1806 | Putative inner membrane protein associated with biofilm formation | *Escherichia coli O7:K1* | Strain IAI39 / ExPEC | A0A0H3MI95 | -- | 19165319 | Not Available | Not Available |

| Target ID | Bacterial Protein Name | Organism Name | Strain | UniProt ID | PDB ID | PubMed ID | Classification as per UniProt GO Molecular Function | Classification of Function as per PDB |
|---|---|---|---|---|---|---|---|---|
| 1807 | Uncharacterized protein | *Cupriavidus taiwanensis* | Strain DSM 17343 / BCRC 17206 / CCUG 44338 / CIP 107171 / LMG 19424 / R1 Ralstonia taiwanensis strain LMG 19424 | B2AGD9 | -- | 18490699 | Not Available | Not Available |
| 1808 | Poly-beta-1, 6-N-acetyl-D-glucosamine synthase (Poly-beta-1, 6-GlcNAc synthase) (EC 2.4.1.-) | *Escherichia coli O7:K1* | Strain IAI39 / ExPEC | A0A0H3MLZ7 | -- | 19165319 | Acetyl glucosaminyl transferase activity | Not Available |
| 1809 | Surface composition regulator | *Yersinia enterocolitica serotype O:8 / biotype 1B* | Strain NCTC 13174 / 8081 | A1JPD4 | -- | 17173484 | DNA binding, DNA-directed DNA polymerase activity | Not Available |
| 1810 | Poly-beta-1, 6-N-acetyl-D-glucosamine synthase (Poly-beta-1, 6-GlcNAc synthase) (EC 2.4.1.-) | *Ralstonia solanaarum CFBP2957* | | D8P2G4 | -- | 20550686 | Acetyl glucosaminyl transferase activity | Not Available |
| 1811 | Putative polysaccharide deacetylase associated with biofilm formation putative lipoprotein (PgaB) | *Ralstonia solanaarum CFBP2957* | | D8P2G3 | -- | 20550686 | Hydrolase activity, acting on carbon-nitrogen (but not peptide) bonds | Not Available |
| 1812 | Uncharacterized protein | *Azospirillum lipoferum* | Strain 4B | G7Z9K0 | -- | 22216014 | Not Available | Not Available |
| 1813 | Putative intercellular adhesion protein A (Biofilm formation) | *Vibrio alginolyticus* | Strain ATCC 17749 / DSM 2171 / NBRC 15630 / NCIMB 1903 / NCTC 12160 / XII-53 | A0A2I3C1P9 | -- | 25635021 | Not Available | Not Available |
| 1814 | Poly-beta-1, 6-N-acetyl-D-glucosamine synthase (Poly-beta-1, 6-GlcNAc synthase) (EC 2.4.1.-) | *Firmicutes bacterium* | | A0A350X1A8 | -- | 30148503 | Acetyl glucosaminyl transferase activity | Not Available |
| 1815 | Uncharacterized protein | *Firmicutes bacterium* | | A0A3D5Z757 | -- | 30148503 | Not Available | Not Available |
| 1816 | Poly-beta-1, 6-N-acetyl-D-glucosamine synthase (Poly-beta-1, 6-GlcNAc synthase) (EC 2.4.1.-) | *Xanthomonas oryzae pv. oryzae* | Strain PXO99A | A0A0K0GHM1 | -- | 18452608 | Acetyl glucosaminyl transferase activity | Not Available |
| 1817 | Uncharacterized protein | *Xanthomonas oryzae pv. oryzae* | Strain PXO99A | A0A0K0GHA3 | -- | 18452608 | Not Available | Not Available |
| 1818 | Uncharacterized protein | *Firmicutes bacterium* | | A0A3B8K032 | -- | 30148503 | Not Available | Not Available |
| 1819 | ABC transporter substrate-binding protein | *Enterobacter asburiae* | | A0A0F3XQA0 | -- | 27284137 | Not Available | Not Available |
| 1820 | Biofilm PGA synthesis protein PgaD | *Escherichia coli O9:H4* | Strain Hs | A0A7I6GYH4 | -- | 18676672 | Not Available | Not Available |
| 1821 | Toxin-antitoxin system, toxin component | *Lentilactobacillus kisonensis DSM 19906 = JCM 15041* | | A0A0R1NQW4 | -- | 26415554 | Not Available | Not Available |
| 1822 | Poly-beta-1, 6-N-acetyl-D-glucosamine synthase (Poly-beta-1, 6-GlcNAc synthase) (EC 2.4.1.-) | *Escherichia coli O9:H4* | Strain Hs | A0A7I6H0P8 | -- | 18676672 | Acetyl glucosaminyl transferase activity | Not Available |
| 1823 | Uncharacterized protein | *Escherichia coli O9:H4* | Strain Hs | A0A7I6H8P9 | -- | 18676672 | Not Available | Not Available |
| 1824 | Outer membrane N-deacetylase | *Enterobacter asburiae* | | A0A143I3T0 | -- | 27284137 | Hydrolase activity, acting on carbon-nitrogen (but not peptide) bonds | Not Available |
| 1825 | Poly-beta-1, 6-N-acetyl-D-glucosamine biosynthesis protein PgaD | *Firmicutes bacterium* | | A0A350X1A9 | -- | 30148503 | Not Available | Not Available |
| 1826 | Uncharacterized protein | *Xanthomonadaceae bacterium* | | A0A3C0HHE3 | -- | 30148503 | Not Available | Not Available |
| 1827 | Poly-beta-1, 6-N-acetyl-D-glucosamine biosynthesis protein PgaD (Fragment) | *Xanthomonadaceae bacterium* | | A0A354YBS0 | -- | 30148503 | Not Available | Not Available |
| 1828 | Poly-beta-1, 6-N-acetyl-D-glucosamine N-deacetylase PgaB | *Xanthomonadaceae bacterium* | | A0A354YE77 | -- | 30148503 | Hydrolase activity, acting on carbon-nitrogen (but not peptide) bonds | Not Available |
| 1829 | Poly-beta-1, 6-N-acetyl-D-glucosamine synthase (Poly-beta-1, 6-GlcNAc synthase) (EC 2.4.1.-) | *Xanthomonadaceae bacterium* | | A0A354YDH4 | -- | 30148503 | Acetyl glucosaminyl transferase activity | Not Available |
| 1830 | Biofilm PGA synthesis lipoprotein pgaB (EC 3.-.-.-) | *Escherichia coli O9:H4* | Strain Hs | A0A7I6H1T4 | -- | 18676672 | Hydrolase activity, acting on carbon-nitrogen (but not peptide) bonds | Not Available |
| 1831 | Surface composition regulator | *Cronobacter sakazakii 701* | | K8CP83 | -- | 23166675 | Not Available | Not Available |
| 1832 | Poly-beta-1, 6-N-acetyl-D-glucosamine synthase (Poly-beta-1, 6-GlcNAc synthase) (EC 2.4.1.-) | *Enterobacter asburiae* | | A0A143I3I2 | -- | 27284137 | Acetyl glucosaminyl transferase activity | Not Available |
| 1833 | Uncharacterized protein | *Pusillimona spp.* | Strain T7-7 | F4GTJ2 | -- | 21622753 | Not Available | Not Available |
| 1834 | Motility quorum-sensing regulator MqsR | *Betaproteobacteria bacterium RBG_16_56_24* | | A0A1F3WWN2 | -- | 27774985 | Not Available | Not Available |
| 1835 | Uncharacterized protein | *Rahnella spp.* | Strain Y9602 | A0A0H3FP36 | -- | 22461551 | Not Available | Not Available |
| 1836 | Poly-beta-1, 6-N-acetyl-D-glucosamine synthase (Poly-beta-1, 6-GlcNAc synthase) (EC 2.4.1.-) | *Pusillimona spp.* | Strain T7-7 | F4GTJ1 | -- | 21622753 | Acetyl glucosaminyl transferase activity | Not Available |
| 1837 | Polysaccharide deacetylase | *Pusillimona spp.* | Strain T7-7 | F4GTJ0 | -- | 21622753 | Hydrolase activity, acting on carbon-nitrogen (but not peptide) bonds | Not Available |
| 1838 | Type II toxin-antitoxin system MqsR family toxin | *Firmicutes bacterium* | | A0A7C7D9P2 | -- | 32123542 | Not Available | Not Available |
| 1839 | Poly-beta-1, 6-N-acetyl-D-glucosamine synthase (Poly-beta-1, 6-GlcNAc synthase) (EC 2.4.1.-) | *Xanthomonadaceae bacterium SCN 69-123* | | A0A1E4L167 | -- | 26031303 | Acetyl glucosaminyl transferase activity | Not Available |
| 1840 | Poly-beta-1, 6-N-acetyl-D-glucosamine synthase (Poly-beta-1, 6-GlcNAc synthase) (EC 2.4.1.-) (Fragment) | *Alphaproteobacteria bacterium* | | A0A537MT94 | -- | 31110364 | Acetyl glucosaminyl transferase activity | Not Available |
| 1841 | Surface composition regulator | *Salmonella enterica subspp. indica serovar* | 6, 14, 25:z10:1, 2, 7 Str. 1121 | V1H6C0 | -- | 24158624 | Not Available | Not Available |
| 1842 | Poly-beta-1, 6-N-acetyl-D-glucosamine N-deacetylase PgaB | *Chromatiales bacterium* | | A0A7C1WNC0 | -- | 31911466 | Hydrolase activity, acting on carbon-nitrogen (but not peptide) bonds | Not Available |
| 1843 | Poly-beta-1, 6-N-acetyl-D-glucosamine synthase (Poly-beta-1, 6-GlcNAc synthase) (EC 2.4.1.-) (Fragment) | *Chromatiales bacterium* | | A0A7C2AD10 | -- | 31911466 | Acetyl glucosaminyl transferase activity | Not Available |
| 1844 | Response receiver-modulated diguanylate cyclase | *Geobacter sulfurreducens* | Strain ATCC 51573 / DSM 12127 / PCA | Q746Z1 | -- | 14671304 | Diguanylate cyclase activity | Not Available |
| 1845 | Poly-beta-1, 6-N-acetyl-D-glucosamine synthase (Poly-beta-1, 6-GlcNAc synthase) (EC 2.4.1.-) | *Xanthomonas gardneri ATCC 19865* | | F0CCM8 | -- | 21396108 | Acetyl glucosaminyl transferase activity | Not Available |
| 1846 | Response receiver-modulated diguanylate cyclase | *Geobacter sulfurreducens* | Strain ATCC 51573 / DSM 12127 / PCA | Q74AN9 | -- | 14671304 | Diguanylate cyclase activity | Not Available |
| 1847 | Sensor diguanylate cyclase, GAF domain-containing | *Geobacter sulfurreducens* | Strain ATCC 51573 / DSM 12127 / PCA | Q74ER5 | -- | 14671304 | Diguanylate cyclase activity | Not Available |
| 1848 | Response receiver-modulated diguanylate cyclase | *Geobacter sulfurreducens* | Strain ATCC 51573 / DSM 12127 / PCA | Q74CL4 | -- | 14671304 | Diguanylate cyclase activity | Not Available |
| 1849 | Sensor diguanylate cyclase, GAF domain-containing | *Geobacter sulfurreducens* | Strain ATCC 51573 / DSM 12127 / PCA | Q74C06 | -- | 14671304 | Diguanylate cyclase activity | Not Available |
| 1850 | Poly-beta-1, 6-N-acetyl-D-glucosamine synthase (Poly-beta-1, 6-GlcNAc synthase) (EC 2.4.1.-) | *Sporolactobacillus laevolacticus DSM 442* | | V6J0L8 | -- | 24371202 | Acetyl glucosaminyl transferase activity | Not Available |
| 1851 | Surface composition regulator | *Salmonella enterica subspp. indica serovar* | 6, 14, 25:z10:1, 2, 7 Str. 1121 | V1GMT8 | -- | 24158624 | Not Available | Not Available |
| 1852 | Diguanylate cyclase, HAMP domain-containing | *Geobacter sulfurreducens* | Strain ATCC 51573 / DSM 12127 / PCA | Q747B1 | -- | 14671304 | Diguanylate cyclase activity | Not Available |
| 1853 | Diguanylate cyclase, CAP_ED domain-containing | *Geobacter sulfurreducens* | Strain ATCC 51573 / DSM 12127 / PCA | Q74F01 | -- | 14671304 | Diguanylate cyclase activity | Not Available |
| 1854 | Diguanylate cyclase | *Geobacter sulfurreducens* | Strain ATCC 51573 / DSM 12127 / PCA | Q74CW7 | -- | 14671304 | Diguanylate cyclase activity | Not Available |
| 1855 | Uncharacterized protein | *Xanthomonas gardneri ATCC 19865* | | F0CCM7 | -- | 21396108 | Not Available | Not Available |
| 1856 | Hemin storage system protein | *Enterobacter spp. Ag1* | | J1GD16 | -- | 22965099 | Not Available | Not Available |
| 1857 | Putative transmembrane protein | *Ralstonia solanacearum FQY_4* | | M4UJ22 | -- | 23661471 | Not Available | Not Available |

| Target ID | Bacterial Protein Name | Organism Name | Strain | UniProt ID | PDB ID | PubMed ID | Classification as per UniProt GO Molecular Function | Classification of Function as per PDB |
|---|---|---|---|---|---|---|---|---|
| 1858 | Biofilm PGA synthesis deacetylase PgaB (EC 3.-.-.-) | *Ralstonia solanacearum FQY_4* | | M4ULZ0 | -- | 23661471 | Hydrolase activity, acting on carbon-nitrogen (but not peptide) bonds | Not Available |
| 1859 | Toxin-antitoxin system, toxin component | *Candidatus melainabacteria bacterium GWF2_37_15* | | A0A1F6PUW9 | -- | 27774985 | Not Available | Not Available |
| 1860 | Poly-beta-1, 6-N-acetyl-D-glucosamine synthase (Poly-beta-1, 6-GlcNAc synthase) (EC 2.4.1.-) | *Enterobacter spp. Ag1* | | J0M9E9 | -- | 22965099 | Acetyl glucosaminyl transferase activity | Not Available |
| 1861 | Outer membrane N-deacetylase | *Enterobacter spp. Ag1* | | J1GD54 | -- | 22965099 | Hydrolase activity, acting on carbon-nitrogen (but not peptide) bonds | Not Available |
| 1862 | Uncharacterized protein | *Caldanaerobacter subterraneus* | | A0A357VJY1 | -- | 30148503 | Not Available | Not Available |
| 1863 | PleD | *Synechocystis spp.* | Strain PCC 6803 / Kazusa | P73011 | -- | 8905231 | Diguanylate cyclase activity | Not Available |
| 1864 | Surface composition regulator | *Salmonella typhimurium* | Strain ATCC 68169 / UK-1 | A0A3Z6P281 | -- | 21622747 | Not Available | Not Available |
| 1865 | Sll0779 protein | *Synechocystis spp.* | Strain PCC 6803 / Kazusa | Q55955 | -- | 8905231 | Diguanylate cyclase activity | Not Available |
| 1866 | Regulatory components of sensory transduction system | *Synechocystis spp.* | Strain PCC 6803 / Kazusa | P72790 | -- | 8905231 | Diguanylate cyclase activity | Not Available |
| 1867 | Uncharacterized protein | *Cupriavidus spp. GA3-3* | | R7XF68 | -- | 23833131 | Not Available | Not Available |
| 1868 | Slr0302 protein | *Synechocystis spp.* | Strain PCC 6803 / Kazusa | Q55906 | -- | 8905231 | Diguanylate cyclase activity | Not Available |
| 1869 | Slr0829 protein | *Synechocystis spp.* | Strain PCC 6803 / Kazusa | Q55414 | -- | 8905231 | Diguanylate cyclase activity | Not Available |
| 1870 | Uncharacterized protein | *Candidatus Rokubacteria bacterium RIFCSPLOWO2_12_FULL_71_19* | | A0A1F7PI81 | -- | 27774985 | Not Available | Not Available |
| 1871 | Poly-beta-1, 6-N-acetyl-D-glucosamine biosynthesis protein PgaD | *Chromatiales bacterium* | | A0A7C2B5U1 | -- | 31911466 | Not Available | Not Available |
| 1872 | Regulatory components of sensory transduction system | *Synechocystis spp.* | Strain PCC 6803 / Kazusa | P73036 | -- | 8905231 | Diguanylate cyclase activity | Not Available |
| 1873 | Putative bacteriophage related protein | *Lactobacillus acidophilus* | Strain ATCC 700396 / NCK56 / N2 / NCFM | Q5FJC7 | -- | 15671160 | Not Available | Not Available |
| 1874 | PleD gene product | *Synechocystis spp.* | Strain PCC 6803 / Kazusa | P74658 | -- | 8905231 | Diguanylate cyclase activity | Not Available |
| 1875 | Poly-beta-1, 6-N-acetyl-D-glucosamine synthase (Poly-beta-1, 6-GlcNAc synthase) (EC 2.4.1.-) | *Ralstonia solanacearum FQY_4* | | M4UIL5 | -- | 23661471 | Acetyl glucosaminyl transferase activity | Not Available |
| 1876 | Uncharacterized protein | *Mammaliicoccus lentus* | Staphylococcus lentus | A0A178PEY6 | -- | 27375578 | Not Available | Not Available |
| 1877 | Poly-beta-1, 6-N-acetyl-D-glucosamine synthase (Poly-beta-1, 6-GlcNAc synthase) (EC 2.4.1.-) | *Achromobacter spp. AONIH1* | | A0A2I8DJJ4 | -- | 29437920 | Acetyl glucosaminyl transferase activity | Not Available |
| 1878 | Poly-beta-1, 6-N-acetyl-D-glucosamine biosynthesis protein PgaD | *Acinetobacter spp. COS3* | | U7GZZ6 | -- | 24356826 | Not Available | Not Available |
| 1879 | Surface composition regulator | *Yersinia pseudotuberculosis serotype O:1b* | Strain IP 31758 | A0A0U1R214 | -- | 17784789 | DNA binding, DNA-directed DNA polymerase activity | Not Available |
| 1880 | Poly-beta-1, 6-N-acetyl-D-glucosamine N-deacetylase PgaB | *Acinetobacter baumannii* | Strain MDR-ZJ06 | A0A654L1R9 | -- | 21788470 | Hydrolase activity, acting on carbon-nitrogen (but not peptide) bonds | Not Available |
| 1881 | Putative biofilm PGA synthesis protein PgaD | *Yersinia pseudotuberculosis serotype O:1b* | Strain IP 31758 | A0A0U1QUY6 | -- | 17784789 | Not Available | Not Available |
| 1882 | Type II toxin-antitoxin system MqsR family toxin | *Pseudomonas spp. MPC6* | | A0A5B7VRQ1 | -- | 31178851; 31089984 | Not Available | Not Available |
| 1883 | Poly-beta-1, 6-N-acetyl-D-glucosamine synthase (Poly-beta-1, 6-GlcNAc synthase) (EC 2.4.1.-) | *Yersinia pseudotuberculosis serotype O:1b* | Strain IP 31758 | A0A0U1QVU7 | -- | 17784789 | Acetyl glucosaminyl transferase activity | Not Available |
| 1884 | Poly-beta-1, 6-N-acetyl-D-glucosamine N-deacetylase PgaB | *Paraburkholderia phytofirmans OLGA172* | | A0A160FHA6 | -- | 27063562 | Hydrolase activity, acting on carbon-nitrogen (but not peptide) bonds | Not Available |
| 1885 | Poly-beta-1, 6-N-acetyl-D-glucosamine N-deacetylase | *Serratia marcescens* | | A0A221DT18 | -- | 28536292 | Hydrolase activity, acting on carbon-nitrogen (but not peptide) bonds | Not Available |
| 1886 | Uncharacterized protein | *Thioalkalivibrio sulfidiphilus* | Strain HL-EbGR7 | B8GT53 | -- | 21475584 | Not Available | Not Available |
| 1887 | Toxin | *Desulfobacteraceae bacterium* | | A0A3N5RWQ1 | -- | 30086797 | Not Available | Not Available |
| 1888 | Poly-beta-1, 6-N-acetyl-D-glucosamine biosynthesis protein PgaD | *Variovorax spp. SCN 67-85* | | A0A1E4IRH5 | -- | 26031303 | Not Available | Not Available |
| 1889 | Uncharacterized protein | *Acinetobacter venetianus* | | A0A150HS68 | -- | 26902269 | Not Available | Not Available |
| 1890 | GGDEF domain-containing protein | *Aquifex aeolicus* | Strain VF5 | O66743 | -- | 9537320 | Diguanylate cyclase activity | Not Available |
| 1891 | Poly-beta-1, 6-N-acetyl-D-glucosamine N-deacetylase PgaB | *Variovorax spp. SCN 67-85* | | A0A1E4IR53 | -- | 26031303 | Hydrolase activity, acting on carbon-nitrogen (but not peptide) bonds | Not Available |
| 1892 | GGDEF domain-containing protein | *Aquifex aeolicus* | Strain VF5 | O66912 | -- | 9537320 | Diguanylate cyclase activity | Not Available |
| 1893 | Poly-beta-1, 6-N-acetyl-D-glucosamine synthase (Poly-beta-1, 6-GlcNAc synthase) (EC 2.4.1.-) | *Variovorax spp. SCN 67-85* | | A0A1E4IR81 | -- | 26031303 | Acetyl glucosaminyl transferase activity | Not Available |
| 1894 | Poly-beta-1, 6-N-acetyl-D-glucosamine synthase (Poly-beta-1, 6-GlcNAc synthase) (EC 2.4.1.-) | *Variovorax spp. CF313* | | J3CAZ9 | -- | 23045501 | Acetyl glucosaminyl transferase activity | Not Available |
| 1895 | GGDEF domain-containing protein | *Aquifex aeolicus* | Strain VF5 | O66623 | -- | 9537320 | Diguanylate cyclase activity | Not Available |
| 1896 | Hemin storage protein | *Thioalkalivibrio sulfidiphilus* | Strain HL-EbGR7 | B8GTH8 | -- | 21475584 | Hydrolase activity, acting on carbon-nitrogen (but not peptide) bonds | Not Available |
| 1897 | Poly-beta-1, 6-N-acetyl-D-glucosamine N-deacetylase PgaB | *Serratia marcescens* | | A0A7D6C0J8 | -- | 21967170 | Hydrolase activity, acting on carbon-nitrogen (but not peptide) bonds | Not Available |
| 1898 | Putative hemin storage signal peptide protein | *Ralstonia solanacearum* | Strain GMI1000 Pseudomonas solanacearum | Q8XT28 | -- | 11823852 | Hydrolase activity, acting on carbon-nitrogen (but not peptide) bonds | Not Available |
| 1899 | Poly-beta-1, 6-N-acetyl-D-glucosamine synthase (Poly-beta-1, 6-GlcNAc synthase) (EC 2.4.1.-) | *Desulfobacteraceae bacterium* | | A0A3A4RX93 | -- | 28644444 | Acetyl glucosaminyl transferase activity | Not Available |
| 1900 | Poly-beta-1, 6-N-acetyl-D-glucosamine synthase (Poly-beta-1, 6-GlcNAc synthase) (EC 2.4.1.-) | *Pseudomonas spp. FH1* | | W2DS81 | -- | 24286439 | Acetyl glucosaminyl transferase activity | Not Available |
| 1901 | Uncharacterized protein | *Marine gamma proteobacterium HTCC2143* | | A0YHD9 | -- | 20601481 | Not Available | Not Available |
| 1902 | Uncharacterized protein | *Desulfobacteraceae bacterium* | | A0A7C1DMQ6 | -- | 31911466 | Not Available | Not Available |
| 1903 | Poly-beta-1, 6-N-acetyl-D-glucosamine synthase (Poly-beta-1, 6-GlcNAc synthase) (EC 2.4.1.-) | *Syntrophotalea carbinolica* | Strain DSM 2380 / NBRC 103641 / GraBd1 Pseudomonas solanacearum | Q3A0T2 | -- | 23227809 | Acetyl glucosaminyl transferase activity | Not Available |
| 1904 | PgaD | *Syntrophotalea carbinolica* | Strain DSM 2380 / NBRC 103641 / GraBd1 Pseudomonas solanacearum | Q3A0T3 | -- | 23227809 | Not Available | Not Available |

| Target ID | Bacterial Protein Name | Organism Name | Strain | UniProt ID | PDB ID | PubMed ID | Classification as per UniProt GO Molecular Function | Classification of Function as per PDB |
|---|---|---|---|---|---|---|---|---|
| 1905 | Uncharacterized protein | *Pseudomonas spp. FH1* | | W2D7M0 | -- | 24286439 | Not Available | Not Available |
| 1906 | Type II toxin-antitoxin system MqsR family toxin | *Xylella fastidiosa subspp. multiplex* | | A0A6I2KF74 | -- | 31704683 | Not Available | Not Available |
| 1907 | Outer membrane N-deacetylase | *Pseudomonas spp. FH1* | | W2DUA0 | -- | 24286439 | Hydrolase activity, acting on carbon-nitrogen (but not peptide) bonds | Not Available |
| 1908 | Uncharacterized protein | *Pseudomonas spp. FH1* | | W2DS47 | -- | 24286439 | Not Available | Not Available |
| 1909 | Uncharacterized protein | *Nitrospirae bacterium RBG_16_64_22* | | A0A1G1HY41 | -- | 27774985 | Not Available | Not Available |
| 1910 | Outer membrane polysaccharide N-deacetylase | *Syntrophotalea carbinolica* | Strain DSM 2380 / NBRC 103641 / GraBd1 Pseudomonas solanacearum | Q3A0T1 | -- | 23227809 | Hydrolase activity, acting on carbon-nitrogen (but not peptide) bonds | Not Available |
| 1911 | Surface composition regulator | *Edwardsiella anguillarum ET080813* | | A0A076LXT7 | -- | 22590641 | Not Available | Not Available |
| 1912 | Uncharacterized protein | *Spirochaetae bacterium HGW-Spirochaetae-9* | | A0A2N1R0F0 | -- | 28350393 | Not Available | Not Available |
| 1913 | Uncharacterized protein | *Pseudomonas spp. GM24* | | J3FKT9 | -- | 23045501 | Not Available | Not Available |
| 1914 | Uncharacterized protein | *Deltaproteobacteria bacterium RIFOXYD12_FULL_55_16* | | A0A1F9MB73 | -- | 27774985 | Not Available | Not Available |
| 1915 | Probable transmembrane protein | *Ralstonia solanacearum* | Strain GMI1000 Pseudomonas solanacearum | Q8XT26 | -- | 11823852 | Not Available | Not Available |
| 1916 | Poly-beta-1, 6-N-acetyl-D-glucosamine synthase (Poly-beta-1, 6-GlcNAc synthase) (EC 2.4.1.-) | *Pseudomonadales bacterium RIFCSPLOWO2_12_FULL_59_450* | | A0A1G3F5W8 | -- | 27774985 | Acetyl glucosaminyl transferase activity | Not Available |
| 1917 | Surface composition regulator | *Salmonella enterica subspp. enterica serovar Inverness* | Str. R8-3668 | G5NIA2 | -- | 21859443 | Not Available | Not Available |
| 1918 | Poly-beta-1, 6-N-acetyl-D-glucosamine synthase (Poly-beta-1, 6-GlcNAc synthase) (EC 2.4.1.-) | *Citrobacter freundii complex spp. CFNIH9* | | A0A2I8THL0 | -- | 29437920 | Acetyl glucosaminyl transferase activity | Not Available |
| 1919 | Poly-beta-1, 6-N-acetyl-D-glucosamine N-deacetylase PgaB | *Citrobacter freundii complex spp. CFNIH9* | | A0A2I8TH49 | -- | 29437920 | Hydrolase activity, acting on carbon-nitrogen (but not peptide) bonds | Not Available |
| 1920 | Uncharacterized protein | *Sporolactobacillus laevolacticus DSM 442* | | V6IZ98 | -- | 24371202 | Not Available | Not Available |
| 1921 | Putative xylanase / chitin deacetylase | *Xanthomonas gardneri ATCC 19865* | | F0CCM9 | -- | 21396108 | Hydrolase activity, acting on carbon-nitrogen (but not peptide) bonds, Hydrolase activity, acting on glycosyl bonds | Not Available |
| 1922 | Nucleotide cyclase, HAMP and GGDEF-related domain-containing | *Geobacter sulfurreducens* | Strain ATCC 51573 / DSM 12127 / PCA | Q74BT9 | -- | 14671304 | Diguanylate cyclase activity | Not Available |
| 1923 | Poly-beta-1, 6-N-acetyl-D-glucosamine biosynthesis protein PgaD | *Citrobacter freundii complex spp. CFNIH9* | | A0A2I8THC0 | -- | 29437920 | Not Available | Not Available |
| 1924 | Surface composition regulator | *Serratia spp. 14-2641* | | A0A1C7WFU5 | -- | 27547538 | Not Available | Not Available |
| 1925 | Poly-beta-1, 6-N-acetyl-D-glucosamine N-deacetylase | *Pseudomonadales bacterium RIFCSPLOWO2_12_FULL_59_450* | | A0A1G3F604 | -- | 27774985 | Hydrolase activity, acting on carbon-nitrogen (but not peptide) bonds | Not Available |
| 1926 | Surface composition regulator | *Citrobacter freundii complex spp. CFNIH9* | | A0A2I8TGS9 | -- | 29437920 | Not Available | Not Available |
| 1927 | Motility quorum-sensing regulator MqsR | *Pseudomonadales bacterium RIFCSPLOWO2_12_FULL_59_450* | | A0A1G3FBL9 | -- | 27774985 | Not Available | Not Available |
| 1928 | Type II toxin-antitoxin system MqsR family toxin | *Mesorhizobium huakuii* | | A0A7G6T3G4 | -- | 32686981 | Not Available | Not Available |
| 1929 | Poly-beta-1, 6-N-acetyl-D-glucosamine biosynthesis protein PgaD | *Pseudomonadales bacterium RIFCSPLOWO2_12_FULL_59_450* | | A0A1G3F5W0 | -- | 27774985 | Not Available | Not Available |
| 1930 | Poly-beta-1, 6-N-acetyl-D-glucosamine biosynthesis protein PgaD | *Acinetobacter baumannii* | Strain MDR-ZJ06 | A0A654L1G9 | -- | 21788470 | Not Available | Not Available |
| 1931 | Uncharacterized protein | *Yersinia pseudotuberculosis serotype O:1b* | Strain IP 31758 | A0A0U1R210 | -- | 17784789 | Not Available | Not Available |
| 1932 | Poly-beta-1, 6-N-acetyl-D-glucosamine synthase (Poly-beta-1, 6-GlcNAc synthase) (EC 2.4.1.-) | *Acinetobacter spp. COS3* | | U7H247 | -- | 24356826 | Acetyl glucosaminyl transferase activity | Not Available |
| 1933 | Poly-beta-1, 6-N-acetyl-D-glucosamine synthase (Poly-beta-1, 6-GlcNAc synthase) (EC 2.4.1.-) | *Lysobacter capsici AZ78* | | A0A125U042 | -- | 24762937 | Acetyl glucosaminyl transferase activity | Not Available |
| 1934 | Poly-beta-1, 6-N-acetyl-D-glucosamine N-deacetylase PgaB | *Acinetobacter baumannii* | Strain MDR-ZJ06 | A0A654KXD6 | -- | 21788470 | Hydrolase activity, acting on carbon-nitrogen (but not peptide) bonds | Not Available |
| 1935 | Poly-beta-1, 6-N-acetyl-D-glucosamine N-deacetylase | *Achromobacter spp. AONIH1* | | A0A2I8DLA9 | -- | 29437920 | Hydrolase activity, acting on carbon-nitrogen (but not peptide) bonds | Not Available |
| 1936 | Biofilm PGA synthesis deacetylase PgaB | *Lysobacter capsici AZ78* | | A0A125U043 | -- | 24762937 | Hydrolase activity, acting on carbon-nitrogen (but not peptide) bonds | Not Available |
| 1937 | Biofilm PGA synthesis lipoprotein PgaB (EC 3.-.-.-) | *Yersinia pseudotuberculosis serotype O:1b* | Strain IP 31758 | A0A0U1R215 | -- | 17784789 | Hydrolase activity, acting on carbon-nitrogen (but not peptide) bonds | Not Available |
| 1938 | Polysaccharide deacetylase | *Acinetobacter spp. COS3* | | U7GXQ6 | -- | 24356826 | Hydrolase activity, acting on carbon-nitrogen (but not peptide) bonds | Not Available |
| 1939 | Poly-beta-1, 6-N-acetyl-D-glucosamine biosynthesis protein PgaD | *Achromobacter spp. AONIH1* | | A0A2I8DK81 | -- | 29437920 | Not Available | Not Available |
| 1940 | Surface composition regulator | *Salmonella enterica subspp. enterica serovar* | Cubana Str. 76814 | V7IJA1 | -- | 24459266 | Not Available | Not Available |
| 1941 | Ferric siderophore transport system, periplasmic binding protein TonB | *Lysobacter capsici AZ78* | | A0A125U041 | -- | 24762937 | Not Available | Not Available |
| 1942 | Surface composition regulator | *Salmonella enterica subspp. enterica serovar* | Virchow Str. SL491 | A0A6C8EY32 | -- | 21602358 | Not Available | Not Available |
| 1943 | Poly-beta-1, 6-N-acetyl-D-glucosamine N-deacetylase PgaB | *Pseudomonas spp. MPC6* | | A0A5B7VT08 | -- | 31178851; 31089984 | Hydrolase activity, acting on carbon-nitrogen (but not peptide) bonds | Not Available |
| 1944 | Uncharacterized protein | *Xylella fastidiosa* | Strain 9a5c | Q9PAL4 | -- | 10910347 | Not Available | Not Available |
| 1945 | Poly-beta-1, 6-N-acetyl-D-glucosamine biosynthesis protein PgaD | *Pseudomonas spp. MPC6* | | A0A5B7VSW4 | -- | 31178851; 31089984 | Not Available | Not Available |
| 1946 | Type II toxin-antitoxin system MqsR family toxin | *Pseudomonas spp. MPC6* | | A0A5B7W3T4 | -- | 31178851; 31089984 | Not Available | Not Available |
| 1947 | Surface composition regulator | *Yersinia pestis bv. Antiqua* | Strain Nepal516 | A0A0H2YFC4 | -- | 16740952 | DNA binding, DNA-directed DNA polymerase activity | Not Available |

| Target ID | Bacterial Protein Name | Organism Name | Strain | UniProt ID | PDB ID | PubMed ID | Classification as per UniProt GO Molecular Function | Classification of Function as per PDB |
|---|---|---|---|---|---|---|---|---|
| 1948 | Lmo2174 protein | *Listeria monocytogenes serovar 1 / 2a* | Strain ATCC BAA-679 / EGD-e | Q8Y595 | -- | 11679669 | Diguanylate cyclase activity, phosphorelay sensor kinase activity | Not Available |
| 1949 | Biofilm PGA synthesis deacetylase PgaB | *Pseudomonas yamanorum* | | A0A145XGS6 | -- | 27231373 | Hydrolase activity, acting on carbon-nitrogen (but not peptide) bonds | Not Available |
| 1950 | Lmo1911 protein | *Listeria monocytogenes serovar 1 / 2a* | Strain ATCC BAA-679 / EGD-e | Q8Y5Z2 | -- | 11679669 | Diguanylate cyclase activity | Not Available |
| 1951 | Uncharacterized protein | *Yersinia pestis bv. Antiqua* | Strain Nepal516 | A0A0H2YLS5 | -- | 16740952 | Not Available | Not Available |
| 1952 | Lmo1912 protein | *Listeria monocytogenes serovar 1 / 2a* | Strain ATCC BAA-679 / EGD-e | Q8Y5Z1 | -- | 11679669 | Diguanylate cyclase activity, phosphorelay sensor kinase activity | Not Available |
| 1953 | Poly-beta-1, 6-N-acetyl-D-glucosamine synthase (Poly-beta-1, 6-GlcNAc synthase) (EC 2.4.1.-) | *Tissierellia bacterium* | | A0A7X8I1F8 | -- | 32123542 | Acetyl glucosaminyl transferase activity | Not Available |
| 1954 | Poly-beta-1, 6-N-acetyl-D-glucosamine synthase (Poly-beta-1, 6-GlcNAc synthase) (EC 2.4.1.-) | *Pseudomonas spp. MPC6* | | A0A5B7VSI7 | -- | 31178851; 31089984 | Acetyl glucosaminyl transferase activity | Not Available |
| 1955 | Poly-beta-1, 6-N-acetyl-D-glucosamine biosynthesis protein PgaD | *Xenorhabdus ishibashii* | | A0A2D0KJM4 | -- | 28993611 | Not Available | Not Available |
| 1956 | Poly-beta-1, 6-N-acetyl-D-glucosamine N-deacetylase PgaB | *Pseudomonas nabeulensis* | | A0A4Z0APQ9 | -- | 30910423 | Hydrolase activity, acting on carbon-nitrogen (but not peptide) bonds | Not Available |
| 1957 | Poly-beta-1, 6-N-acetyl-D-glucosamine synthase (Poly-beta-1, 6-GlcNAc synthase) (EC 2.4.1.-) | *Xenorhabdus ishibashii* | | A0A2D0KJN1 | -- | 28993611 | Acetyl glucosaminyl transferase activity | Not Available |
| 1958 | Poly-beta-1, 6-N-acetyl-D-glucosamine synthase (Poly-beta-1, 6-GlcNAc synthase) (EC 2.4.1.-) | *Acinetobacter baumannii* | Strain MDR-ZJ06 | A0A654KXG3 | -- | 21788470 | Acetyl glucosaminyl transferase activity | Not Available |
| 1959 | Poly-beta-1, 6-N-acetyl-D-glucosamine biosynthesis protein PgaD | *Acinetobacter baumannii* | Strain MDR-ZJ06 | A0A654KXA5 | -- | 21788470 | Not Available | Not Available |
| 1960 | Poly-beta-1, 6-N-acetyl-D-glucosamine synthase (Poly-beta-1, 6-GlcNAc synthase) (EC 2.4.1.-) | *Acinetobacter baumannii* | Strain MDR-ZJ06 | A0A654L1A0 | -- | 21788470 | Acetyl glucosaminyl transferase activity | Not Available |
| 1961 | Polysaccharide deacetylase | *Xenorhabdus ishibashii* | | A0A2D0KK07 | -- | 28993611 | Hydrolase activity, acting on carbon-nitrogen (but not peptide) bonds | Not Available |
| 1962 | Motility quorum-sensing regulator MqsR | *Pseudomonas spp. Os17* | | A0A0D6BDD9 | -- | 25496595 | Not Available | Not Available |
| 1963 | Putative biofilm PGA synthesis protein PgaD | *Pseudomonas fluorescens* | Strain ATCC BAA-477 / NRRL B-23932 / Pf-5 | Q4KKC2 | -- | 15980861 | Not Available | Not Available |
| 1964 | Poly-beta-1, 6-N-acetyl-D-glucosamine synthase (Poly-beta-1, 6-GlcNAc synthase) (EC 2.4.1.-) | *Actinobacillus pleuropneumoniae serotype 5b* | Strain L20 | A3N3L7 | -- | 18065534 | Acetyl glucosaminyl transferase activity | Not Available |
| 1965 | Poly-beta-1, 6-N-acetyl-D-glucosamine biosynthesis protein PgaD | *Pseudomonas fuscovaginae* | | A0A0M9GIL7 | -- | 26422147 | Not Available | Not Available |
| 1966 | Poly-beta-1, 6-N-acetyl-D-glucosamine synthase (Poly-beta-1, 6-GlcNAc synthase) (EC 2.4.1.-) | *Enterobacter hormaechei* | | A0A6G3F3P9 | -- | 31477907 | Acetyl glucosaminyl transferase activity | Not Available |
| 1967 | Uncharacterized protein | *Alphaproteobacteria bacterium HGW-Alphaproteobacteria-3* | | A0A2N3BZZ5 | -- | 28350393 | Not Available | Not Available |
| 1968 | Poly-beta-1, 6-N-acetyl-D-glucosamine biosynthesis protein PgaD (Fragment) | *Acinetobacter ursingii* | | A0A3D2SJF3 | -- | 30148503 | Not Available | Not Available |
| 1969 | Poly-beta-1, 6-N-acetyl-D-glucosamine biosynthesis protein PgaD | *Xanthomonadaceae bacterium SCN 69-123* | | A0A1E4L188 | -- | 26031303 | Not Available | Not Available |
| 1970 | Outer membrane N-deacetylase | *Achromobacter marplatensis* | | J4P4T1 | -- | 23105084 | Hydrolase activity, acting on carbon-nitrogen (but not peptide) bonds | Not Available |
| 1971 | Surface composition regulator | *Serratia fonticola AU-P3* | 3 | U2LWW9 | -- | 24233592 | Not Available | Not Available |
| 1972 | Surface composition regulator | *Enterobacteriaceae bacterium* | Strain FGI 57 | L0LYH9 | -- | 23469353 | Not Available | Not Available |
| 1973 | Poly-beta-1, 6-N-acetyl-D-glucosamine biosynthesis protein PgaD | *Escherichia coli* | | A0A5P0JEL1 | -- | 31642700 | Not Available | Not Available |
| 1974 | Uncharacterized protein | *Pluralibacter gergoviae* | Enterobacter gergoviae | A0A1S2B9G4 | -- | 25502672 | Not Available | Not Available |
| 1975 | Surface composition regulator | *Cronobacter sakazakii* | Strain ATCC BAA-894 Enterobacter sakazakii | A7MF81 | -- | 20221447 | Not Available | Not Available |
| 1976 | Surface composition regulator | *Escherichia coli* | | A0A5D1FAL4 | -- | 31477907 | Not Available | Not Available |
| 1977 | Poly-beta-1, 6-N-acetyl-D-glucosamine synthase (Poly-beta-1, 6-GlcNAc synthase) (EC 2.4.1.-) | *Escherichia coli* | | A0A246NPP4 | -- | 30286803 | Acetyl glucosaminyl transferase activity | Not Available |
| 1978 | Surface composition regulator | *Pectobacterium polonicum* | | A0A4U7KK94 | -- | 30973319 | DNA binding, DNA-directed DNA polymerase activity | Not Available |
| 1979 | Poly-beta-1, 6-N-acetyl-D-glucosamine synthase (Poly-beta-1, 6-GlcNAc synthase) (EC 2.4.1.-) | *Macrococcus epidermidis* | | A0A328A136 | -- | 29951040 | Acetyl glucosaminyl transferase activity | Not Available |
| 1980 | Surface composition regulator | *Escherichia coli* | | A0A6M0PTC6 | -- | 16943019 | Not Available | Not Available |
| 1981 | Poly-beta-1, 6-N-acetyl-D-glucosamine synthase (Poly-beta-1, 6-GlcNAc synthase) (EC 2.4.1.-) | *Escherichia coli* | | A0A6M0PSJ4 | -- | 16943019 | Acetyl glucosaminyl transferase activity | Not Available |
| 1982 | Lipoprotein (HmsF protein of hmsHFRS Operon) | *Xenorhabdus nematophila* | Strain ATCC 19061 / DSM 3370 / CCUG 14189 / LMG 1036 / NCIMB 9965 / AN6 | D3VJ37 | -- | 22125637 | Hydrolase activity, acting on carbon-nitrogen (but not peptide) bonds | Not Available |
| 1983 | Poly-beta-1, 6-N-acetyl-D-glucosamine synthase (Poly-beta-1, 6-GlcNAc synthase) (EC 2.4.1.-) | *Xenorhabdus nematophila* | Strain ATCC 19061 / DSM 3370 / CCUG 14189 / LMG 1036 / NCIMB 9965 / AN6 | D3VJ36 | -- | 22125637 | Acetyl glucosaminyl transferase activity | Not Available |
| 1984 | Poly-beta-1, 6-N-acetyl-D-glucosamine N-deacetylase PgaB | *Xanthomonadaceae bacterium SCN 69-123* | | A0A1E4L199 | -- | 26031303 | Hydrolase activity, acting on carbon-nitrogen (but not peptide) bonds | Not Available |
| 1985 | Haemin storage system (HmsS protein of hmsHFRS Operon) | *Xenorhabdus nematophila* | Strain ATCC 19061 / DSM 3370 / CCUG 14189 / LMG 1036 / NCIMB 9965 / AN6 | D3VJ35 | -- | 22125637 | Not Available | Not Available |
| 1986 | Poly-beta-1, 6-N-acetyl-D-glucosamine synthase (Poly-beta-1, 6-GlcNAc synthase) (EC 2.4.1.-) | *Escherichia coli* | | A0A792HM93 | -- | 30286803 | Acetyl glucosaminyl transferase activity | Not Available |
| 1987 | Poly-beta-1, 6-N-acetyl-D-glucosamine biosynthesis protein PgaD | *Pectobacterium polonicum* | | A0A4U7KCH9 | -- | 30973319 | Not Available | Not Available |
| 1988 | Poly-beta-1, 6-N-acetyl-D-glucosamine biosynthesis protein PgaD | *Escherichia coli* | | A0A5P0J1W1 | -- | 31642700 | Not Available | Not Available |
| 1989 | mRNA interferase MqsR | *Escherichia coli* | | A0A0P7PX62 | -- | 26441869 | Not Available | Not Available |
| 1990 | Poly-beta-1, 6-N-acetyl-D-glucosamine biosynthesis protein PgaD | *Clostridium spp. Maddingley MBC34-26* | | K6T4C8 | -- | 23405323 | Not Available | Not Available |
| 1991 | Poly-beta-1, 6-N-acetyl-D-glucosamine synthase (Poly-beta-1, 6-GlcNAc synthase) (EC 2.4.1.-) | *Pectobacterium polonicum* | | A0A4U7KDL9 | -- | 30973319 | Acetyl glucosaminyl transferase activity | Not Available |
| 1992 | Poly-beta-1, 6-N-acetyl-D-glucosamine N-deacetylase PgaB (EC 3.5.1.-) | *Escherichia coli* | | A0A7B3MEP4 | -- | 30286803 | Hydrolase activity, acting on carbon-nitrogen (but not peptide) bonds | Not Available |
| 1993 | Poly-beta-1, 6-N-acetyl-D-glucosamine synthase (Poly-beta-1, 6-GlcNAc synthase) (EC 2.4.1.-) | *Escherichia coli* | | A0A5P0JF91 | -- | 31642700 | Acetyl glucosaminyl transferase activity | Not Available |
| 1994 | Poly-beta-1, 6-N-acetyl-D-glucosamine synthase (Poly-beta-1, 6-GlcNAc synthase) (EC 2.4.1.-) | *Clostridium spp. Maddingley MBC34-26* | | K6U9Y4 | -- | 23405323 | Acetyl glucosaminyl transferase activity | Not Available |

| Target ID | Bacterial Protein Name | Organism Name | Strain | UniProt ID | PDB ID | PubMed ID | Classification as per UniProt GO Molecular Function | Classification of Function as per PDB |
|---|---|---|---|---|---|---|---|---|
| 1995 | Poly-beta-1, 6-N-acetyl-D-glucosamine N-deacetylase PgaB | *Escherichia coli* | | A0A5P0JEP5 | -- | 31642700 | Hydrolase activity, acting on carbon-nitrogen (but not peptide) bonds | Not Available |
| 1996 | Poly-beta-1, 6-N-acetyl-D-glucosamine N-deacetylase PgaB | *Escherichia coli* | | A0A6M0PSS0 | -- | 16943019 | Hydrolase activity, acting on carbon-nitrogen (but not peptide) bonds | Not Available |
| 1997 | Sll1687 protein | *Synechocystis spp.* | Strain PCC 6803 / Kazusa | P73713 | -- | 8905231 | Diguanylate cyclase activity | Not Available |
| 1998 | Poly-beta-1, 6-N-acetyl-D-glucosamine synthase (Poly-beta-1, 6-GlcNAc synthase) (EC 2.4.1.-) | *Halomonas spp. HAL1* | | G4F1K5 | -- | 22156396 | Acetyl glucosaminyl transferase activity | Not Available |
| 1999 | Xylanae / chitin deacetylase | *Halomonas spp. HAL1* | | G4F1K6 | -- | 22156396 | Hydrolase activity, acting on carbon-nitrogen (but not peptide) bonds, Hydrolase activity, acting on glycosyl bonds | Not Available |
| 2000 | Diguanylate cyclase (EC 2.7.7.65) | *Shewanella oneidensis* | Strain MR-1 | Q8EF08 | -- | 12368813 | Diguanylate cyclase activity | Not Available |
| 2001 | Diguanylate cyclase (EC 2.7.7.65) | *Shewanella oneidensis* | Strain MR-1 | Q8EFD2 | -- | 12368813 | Diguanylate cyclase activity | Not Available |
| 2002 | Diguanylate cyclase with PAS sensory domain (EC 2.7.7.65) | *Shewanella oneidensis* | Strain MR-1 | Q8EJ99 | -- | 12368813 | Diguanylate cyclase activity | Not Available |
| 2003 | Diguanylate cyclase (EC 2.7.7.65) | *Shewanella oneidensis* | Strain MR-1 | Q8EIP1 | -- | 12368813 | Diguanylate cyclase activity | Not Available |
| 2004 | Surface composition regulator | *Citrobacter spp. CFNIH10* | | A0A3Q8DFL3 | -- | 29437920 | Not Available | Not Available |
| 2005 | Diguanylate cyclase with PAS sensory domain (EC 2.7.7.65) | *Shewanella oneidensis* | Strain MR-1 | Q8EF28 | -- | 12368813 | Diguanylate cyclase activity, phosphoprotein phosphatase activity | Not Available |
| 2006 | Poly-beta-1, 6-N-acetyl-D-glucosamine synthase (Poly-beta-1, 6-GlcNAc synthase) (EC 2.4.1.-) | *Nitrospira bacterium SG8_35_1* | | A0A0S8CZY6 | -- | 25922666 | Acetyl glucosaminyl transferase activity | Not Available |
| 2007 | Type II toxin-antitoxin system MqsR family toxin | *Pseudomonas spp. WS 5078* | | A0A7Y1FKF5 | -- | 32587583 | Not Available | Not Available |
| 2008 | Diguanylate cyclase with HAMP domain (EC 2.7.7.65) | *Shewanella oneidensis* | Strain MR-1 | Q8EC11 | -- | 12368813 | Diguanylate cyclase activity | Not Available |
| 2009 | Poly-beta-1, 6-N-acetyl-D-glucosamine synthase (Poly-beta-1, 6-GlcNAc synthase) (EC 2.4.1.-) | *Enterobacter ludwigii* | | G8LE14 | -- | 22675579 | Acetyl glucosaminyl transferase activity | Not Available |
| 2010 | Diguanylate cyclase (EC 2.7.7.65) | *Shewanella oneidensis* | Strain MR-1 | Q8EGF8 | -- | 12368813 | Diguanylate cyclase activity | Not Available |
| 2011 | Diguanylate cyclase (EC 2.7.7.65) | *Shewanella oneidensis* | Strain MR-1 | Q8EIB5 | -- | 12368813 | Diguanylate cyclase activity | Not Available |
| 2012 | Outer membrane N-deacetylase | *Enterobacter ludwigii* | | W0BV21 | -- | 25197457 | Hydrolase activity, acting on carbon-nitrogen (but not peptide) bonds | Not Available |
| 2013 | Outer membrane N-deacetylase | *Enterobacter ludwigii* | | A0A7U4N614 | -- | 26251488 | Hydrolase activity, acting on carbon-nitrogen (but not peptide) bonds | Not Available |
| 2014 | P pilus subunit FimA | *Shewanella oneidensis* | Strain MR-1 | Q8E8S9 | -- | 12368813 | Not Available | Not Available |
| 2015 | Diguanylate cyclase (EC 2.7.7.65) | *Shewanella oneidensis* | Strain MR-1 | Q8EAN9 | -- | 12368813 | Diguanylate cyclase activity | Not Available |
| 2016 | Diguanylate cyclase (EC 2.7.7.65) | *Shewanella oneidensis* | Strain MR-1 | Q8EGP7 | -- | 12368813 | Diguanylate cyclase activity | Not Available |
| 2017 | 7TM intracellular sigNot Availablelling protein with GGDEF domain (EC 2.7.7.65) | *Shewanella oneidensis* | Strain MR-1 | Q8EGN0 | -- | 12368813 | Diguanylate cyclase activity | Not Available |
| 2018 | NodB homology domain-containing protein | *Nitrospira bacterium SG8_35_1* | | A0A0S8CYX5 | -- | 25922666 | Hydrolase activity, acting on carbon-nitrogen (but not peptide) bonds | Not Available |
| 2019 | Diguanylate cyclase with response regulator receiver modulation (EC 2.7.7.65) | *Shewanella oneidensis* | Strain MR-1 | Q8EF60 | -- | 12368813 | Diguanylate cyclase activity, protein kinase activity | Not Available |
| 2020 | Diguanylate cyclase (EC 2.7.7.65) | *Shewanella oneidensis* | Strain MR-1 | Q8EGW1 | -- | 12368813 | Diguanylate cyclase activity | Not Available |
| 2021 | Diguanylate cyclase | *Shewanella oneidensis* | Strain MR-1 | Q8EBM2 | -- | 12368813 | Diguanylate cyclase activity | Not Available |
| 2022 | ABC transporter substrate-binding protein | *Enterobacter ludwigii* | | A0A7U4N650 | -- | 26251488 | Not Available | Not Available |
| 2023 | BifunctioNot Availablel periplasmic substrate binding protein / cytoplasmic diguanylate cyclase | *Shewanella oneidensis* | Strain MR-1 | Q8EB35 | -- | 12368813 | Diguanylate cyclase activity | Not Available |
| 2024 | Diguanylate cyclase (EC 2.7.7.65) | *Shewanella oneidensis* | Strain MR-1 | Q8E979 | -- | 12368813 | Diguanylate cyclase activity | Not Available |
| 2025 | Diguanylate cyclase (EC 2.7.7.65) | *Shewanella oneidensis* | Strain MR-1 | Q8E963 | -- | 12368813 | Diguanylate cyclase activity | Not Available |
| 2026 | Diguanylate cyclase (EC 2.7.7.65) | *Shewanella oneidensis* | Strain MR-1 | Q8E936 | -- | 12368813 | Diguanylate cyclase activity | Not Available |
| 2027 | SigNot Availablelling protein with GGDEF domain | *Shewanella oneidensis* | Strain MR-1 | Q8EDJ0 | -- | 12368813 | Diguanylate cyclase activity | Not Available |
| 2028 | BifunctioNot Availablel periplasmic substrate binding protein / cytoplasmic diguanylate cyclase (EC 2.7.7.65) | *Shewanella oneidensis* | Strain MR-1 | Q8EIJ8 | -- | 12368813 | Diguanylate cyclase activity | Not Available |
| 2029 | Diguanylate cyclase with GAF sensory domain | *Shewanella oneidensis* | Strain MR-1 | Q8EBG2 | -- | 12368813 | Diguanylate cyclase activity, phosphoprotein phosphatase activity | Not Available |
| 2030 | SigNot Availablelling protein with GGDEF domain | *Shewanella oneidensis* | Strain MR-1 | Q8EE69 | -- | 12368813 | Diguanylate cyclase activity | Not Available |
| 2031 | Poly-beta-1, 6-N-acetyl-D-glucosamine N-deacetylase | *Enterobacter ludwigii* | | G8LE13 | -- | 22675579 | Hydrolase activity, acting on carbon-nitrogen (but not peptide) bonds | Not Available |
| 2032 | Putative membrane protein | *Acinetobacter baumannii* | Strain ACICU | A0A7U3XX92 | -- | 18411315 | Not Available | Not Available |
| 2033 | Poly-beta-1, 6-N-acetyl-D-glucosamine synthase (Poly-beta-1, 6-GlcNAc synthase) (EC 2.4.1.-) | *Acinetobacter baumannii* | Strain ACICU | A0A7U4DDW4 | -- | 18411315 | Acetyl glucosaminyl transferase activity | Not Available |
| 2034 | Poly-beta-1, 6-N-acetyl-D-glucosamine synthase (Poly-beta-1, 6-GlcNAc synthase) (EC 2.4.1.-) | *Cupriavidus basilensis* | | A0A0C4Y8S7 | -- | 25977418 | Acetyl glucosaminyl transferase activity | Not Available |
| 2035 | Uncharacterized protein | *Acinetobacter baumannii* | Strain ACICU | A0A7U3Y161 | -- | 18411315 | Not Available | Not Available |
| 2036 | Predicted xylanase / chitin deacetylase | *Acinetobacter baumannii* | Strain ACICU | A0A7U3Y173 | -- | 18411315 | Hydrolase activity, acting on carbon-nitrogen (but not peptide) bonds, Hydrolase activity, acting on glycosyl bonds | Not Available |
| 2037 | Surface composition regulator | *Cronobacter dublinensis subspp. dublinensis LMG 23823* | | A0A0K2P1G4 | -- | 27013041 | Not Available | Not Available |
| 2038 | Poly-beta-1, 6-N-acetyl-D-glucosamine synthase (Poly-beta-1, 6-GlcNAc synthase) (EC 2.4.1.-) | *Acinetobacter baumannii* | Strain ACICU | A0A7U3XXA6 | -- | 18411315 | Acetyl glucosaminyl transferase activity | Not Available |
| 2039 | Uncharacterized protein | *Methylocucumis oryzae* | | A0A0F3IJ44 | -- | 26547566 | Not Available | Not Available |
| 2040 | Biofilm PGA synthesis deacetylase PgaB (EC 3.-.-.-) | *Cupriavidus basilensis* | | A0A0C4YAV1 | -- | 25977418 | Hydrolase activity, acting on carbon-nitrogen (but not peptide) bonds | Not Available |
| 2041 | Predicted xylanase / chitin deacetylase | *Acinetobacter baumannii* | Strain ACICU | A0A7U3XXD6 | -- | 18411315 | Hydrolase activity, acting on carbon-nitrogen (but not peptide) bonds, Hydrolase activity, acting on glycosyl bonds | Not Available |

| Target ID | Bacterial Protein Name | Organism Name | Strain | UniProt ID | PDB ID | PubMed ID | Classification as per UniProt GO Molecular Function | Classification of Function as per PDB |
|---|---|---|---|---|---|---|---|---|
| 2042 | Polysaccharide deacetylase | *Methylocucumis oryzae* | | A0A0F3IGY7 | -- | 26547566 | Hydrolase activity, acting on carbon-nitrogen (but not peptide) bonds | Not Available |
| 2043 | Surface composition regulator | *Salmonella dublin* | Strain CT_02021853 | A0A6C7A0T4 | -- | 21602358 | Not Available | Not Available |
| 2044 | Putative xylanase / chitin deacetylase | *Xanthomonas vesicatoria ATCC 35937* | | F0BJX6 | -- | 21396108 | Hydrolase activity, acting on carbon-nitrogen (but not peptide) bonds, Hydrolase activity, acting on glycosyl bonds | Not Available |
| 2045 | Poly-beta-1, 6-N-acetyl-D-glucosamine synthase (Poly-beta-1, 6-GlcNAc synthase) (EC 2.4.1.-) | *Methylocucumis oryzae* | | A0A0F3IH37 | -- | 26547566 | Acetyl glucosaminyl transferase activity | Not Available |
| 2046 | Uncharacterized protein | *Xanthomonas vesicatoria ATCC 35937* | | F0BJX8 | -- | 21396108 | Not Available | Not Available |
| 2047 | Uncharacterized protein | *Methylocucumis oryzae* | | A0A0F3IN01 | -- | 26547566 | Not Available | Not Available |
| 2048 | Uncharacterized protein | *Koribacter versatilis* | Strain Ellin345 | Q1IJ03 | -- | 19201974 | Not Available | Not Available |
| 2049 | Poly-beta-1, 6-N-acetyl-D-glucosamine biosynthesis protein PgaD | *Oxalobacteraceae bacterium* | | A0A4Q3IWL6 | -- | 30498029 | Not Available | Not Available |
| 2050 | Poly-beta-1, 6-N-acetyl-D-glucosamine biosynthesis protein PgaD | *Pelosinus spp. UFO1* | | A0A075KH26 | -- | 25189589 | Not Available | Not Available |
| 2051 | Poly-beta-1, 6-N-acetyl-D-glucosamine synthase (Poly-beta-1, 6-GlcNAc synthase) (EC 2.4.1.-) | *Oxalobacteraceae bacterium* | | A0A4Q3IX21 | -- | 30498029 | Acetyl glucosaminyl transferase activity | Not Available |
| 2052 | Poly-beta-1, 6-N-acetyl-D-glucosamine synthase (Poly-beta-1, 6-GlcNAc synthase) (EC 2.4.1.-) | *Pelosinus spp. UFO1* | | A0A075KDU5 | -- | 25189589 | Acetyl glucosaminyl transferase activity | Not Available |
| 2053 | Uncharacterized protein | *Helicobacteraceae bacterium CG1_02_36_14* | | A0A1J4U2J2 | -- | 27112493 | Not Available | Not Available |
| 2054 | Poly-beta-1, 6-N-acetyl-D-glucosamine synthase (Poly-beta-1, 6-GlcNAc synthase) (EC 2.4.1.-) | *Xanthomonas vesicatoria ATCC 35937* | | F0BJX7 | -- | 21396108 | Acetyl glucosaminyl transferase activity | Not Available |
| 2055 | Poly-beta-1, 6-N-acetyl-D-glucosamine synthase (Poly-beta-1, 6-GlcNAc synthase) (EC 2.4.1.-) | *Escherichia coli O1:K1 / APEC* | | A0A0H2YXS0 | -- | 17293413 | Acetyl glucosaminyl transferase activity | Not Available |
| 2056 | Poly-beta-1, 6-N-acetyl-D-glucosamine synthase (Poly-beta-1, 6-GlcNAc synthase) (EC 2.4.1.-) | *Staphylococcus aureus subspp. aureus M013* | | A0A7U4AW02 | -- | 22328755 | Acetyl glucosaminyl transferase activity | Not Available |
| 2057 | Predicted inner Membrane protein | *Escherichia coli O78:H11* | Strain H10407 / ETEC | E3PJI8 | -- | 20802035 | Not Available | Not Available |
| 2058 | Putative polysaccharide deacetylase associated with biofilm formation putative lipoprotein | *Escherichia fergusonii* | Strain ATCC 35469 / DSM 13698 / CCUG 18766 / IAM 14443 / JCM 21226 / LMG 7866 / NBRC 102419 / NCTC 12128 / CDC 0568-73 | B7LTB0 | -- | 19165319 | Hydrolase activity, acting on carbon-nitrogen (but not peptide) bonds | Not Available |
| 2059 | Poly-beta-1, 6-N-acetyl-D-glucosamine synthase (Poly-beta-1, 6-GlcNAc synthase) (EC 2.4.1.-) | *Escherichia fergusonii* | Strain ATCC 35469 / DSM 13698 / CCUG 18766 / IAM 14443 / JCM 21226 / LMG 7866 / NBRC 102419 / NCTC 12128 / CDC 0568-73 | B7LTB1 | -- | 19165319 | Acetyl glucosaminyl transferase activity | Not Available |
| 2060 | Uncharacterized protein | *Escherichia coli O78:H11* | Strain H10407 / ETEC | E3PJE2 | -- | 20802035 | Not Available | Not Available |
| 2061 | Biofilm PGA synthesis protein pgaD | *Escherichia fergusonii* | Strain ATCC 35469 / DSM 13698 / CCUG 18766 / IAM 14443 / JCM 21226 / LMG 7866 / NBRC 102419 / NCTC 12128 / CDC 0568-73 | B7LTB2 | -- | 19165319 | Not Available | Not Available |
| 2062 | Surface composition regulator | *Klebsiella pneumoniae subspp. pneumoniae KpQ3* | | A0A7U9CYT9 | -- | 23469341 | Not Available | Not Available |
| 2063 | Hemin storage system protein | *Klebsiella pneumoniae subspp. pneumoniae KpQ3* | | A0A7U9GZW8 | -- | 23469341 | Not Available | Not Available |
| 2064 | Poly-beta-1, 6-N-acetyl-D-glucosamine synthase (Poly-beta-1, 6-GlcNAc synthase) (EC 2.4.1.-) | *Klebsiella pneumoniae subspp. pneumoniae KpQ3* | | A0A7U9D807 | -- | 23469341 | Acetyl glucosaminyl transferase activity | Not Available |
| 2065 | Outer membrane N-deacetylase | *Klebsiella pneumoniae subspp. pneumoniae KpQ3* | | A0A7U9D765 | -- | 23469341 | Hydrolase activity, acting on carbon-nitrogen (but not peptide) bonds | Not Available |
| 2066 | Putative hemin storage protein | *Bordetella parapertussis* | Strain 12822 / ATCC BAA-587 / NCTC 13253 | Q7W830 | -- | 12910271 | Hydrolase activity, acting on carbon-nitrogen (but not peptide) bonds | Not Available |
| 2067 | Uncharacterized protein | *Desulfovibrio spp. A2* | | G2H343 | -- | 22072648 | Not Available | Not Available |
| 2068 | Poly-beta-1, 6-N-acetyl-D-glucosamine synthase (Poly-beta-1, 6-GlcNAc synthase) (EC 2.4.1.-) | *Pseudomonas spp. GM80* | | J2Y7T2 | -- | 23045501 | Acetyl glucosaminyl transferase activity | Not Available |
| 2069 | Poly-beta-1, 6-N-acetyl-D-glucosamine N-deacetylase PgaB | *Pseudomonas spp. GM102* | | J2MA13 | -- | 23045501 | Hydrolase activity, acting on carbon-nitrogen (but not peptide) bonds | Not Available |
| 2070 | Poly-beta-1, 6-N-acetyl-D-glucosamine synthase (Poly-beta-1, 6-GlcNAc synthase) (EC 2.4.1.-) | *Pseudomonas spp. GM102* | | J3DYT0 | -- | 23045501 | Acetyl glucosaminyl transferase activity | Not Available |
| 2071 | Biofilm PGA synthesis lipoprotein PgaB | *Bibersteinia trehalosi USDA-ARS-USMARC-188* | | A0A4V7IAU5 | -- | 24526647 | Hydrolase activity, acting on carbon-nitrogen (but not peptide) bonds | Not Available |
| 2072 | Poly-beta-1, 6-N-acetyl-D-glucosamine synthase (Poly-beta-1, 6-GlcNAc synthase) (EC 2.4.1.-) | *Bibersteinia trehalosi USDA-ARS-USMARC-188* | | A0A4V7IB85 | -- | 24526647 | Acetyl glucosaminyl transferase activity | Not Available |
| 2073 | Uncharacterized protein | *Bordetella parapertussis* | Strain 12822 / ATCC BAA-587 / NCTC 13253 | Q7WB14 | -- | 12910271 | Not Available | Not Available |
| 2074 | Poly-beta-1, 6-N-acetyl-D-glucosamine N-deacetylase PgaB | *Pseudomonas spp. GM80* | | J3IPL5 | -- | 23045501 | Hydrolase activity, acting on carbon-nitrogen (but not peptide) bonds | Not Available |
| 2075 | Poly-beta-1, 6-N-acetyl-D-glucosamine synthase (Poly-beta-1, 6-GlcNAc synthase) (EC 2.4.1.-) | *Pseudomonas brassicacearum* | | W8PKM7 | -- | 24812222 | Acetyl glucosaminyl transferase activity | Not Available |
| 2076 | Poly-beta-1, 6-N-acetyl-D-glucosamine synthase (Poly-beta-1, 6-GlcNAc synthase) (EC 2.4.1.-) | *Pseudomonas spp. GM18* | | J2NNQ5 | -- | 23045501 | Acetyl glucosaminyl transferase activity | Not Available |
| 2077 | Outer membrane N-deacetylase | *Pseudomonas brassicacearum* | | W8PPL1 | -- | 24812222 | Hydrolase activity, acting on carbon-nitrogen (but not peptide) bonds | Not Available |
| 2078 | Biofilm PGA synthesis protein PgaD | *Pseudomonas brassicacearum* | | W8PCX6 | -- | 24812222 | Not Available | Not Available |
| 2079 | Poly-beta-1, 6-N-acetyl-D-glucosamine biosynthesis protein PgaD | *Pseudomonas spp. GM18* | | J3EH54 | -- | 23045501 | Not Available | Not Available |
| 2080 | Type II toxin-antitoxin system MqsR family toxin | *Geobacter spp.* | | A0A351WRQ6 | -- | 30148503 | Not Available | Not Available |
| 2081 | Poly-beta-1, 6-N-acetyl-D-glucosamine synthase (Poly-beta-1, 6-GlcNAc synthase) (EC 2.4.1.-) | *Bordetella parapertussis* | Strain 12822 / ATCC BAA-587 / NCTC 13253 | Q7W831 | -- | 12910271 | Acetyl glucosaminyl transferase activity | Not Available |
| 2082 | Uncharacterized protein | *Pediococcus damnosus* | | A0A2G9VE09 | -- | 29126241 | Not Available | Not Available |
| 2083 | Uncharacterized protein | *Pseudomonas spp. GM102* | | J3DNY8 | -- | 23045501 | Not Available | Not Available |

| Target ID | Bacterial Protein Name | Organism Name | Strain | UniProt ID | PDB ID | PubMed ID | Classification as per UniProt GO Molecular Function | Classification of Function as per PDB |
|---|---|---|---|---|---|---|---|---|
| 2084 | Putative membrane protein | *Bordetella parapertussis* | Strain 12822 / ATCC BAA-587 / NCTC 13253 | Q7W832 | -- | 12910271 | Not Available | Not Available |
| 2085 | Poly-beta-1, 6-N-acetyl-D-glucosamine biosynthesis protein PgaD | *Pseudomonas spp. GM102* | | J2VDN2 | -- | 23045501 | Not Available | Not Available |
| 2086 | Poly-beta-1, 6-N-acetyl-D-glucosamine synthase (Poly-beta-1, 6-GlcNAc synthase) (EC 2.4.1.-) | *Escherichia coli* | Strain UTI89 / UPEC | Q1RDJ5 | -- | 16585510 | Acetyl glucosaminyl transferase activity | Not Available |
| 2087 | Poly-beta-1, 6-N-acetyl-D-glucosamine synthase (Poly-beta-1, 6-GlcNAc synthase) (EC 2.4.1.-) (Fragment) | *Enterobacteriaceae bacterium* | | A0A3C2BGL3 | -- | 30148503 | Acetyl glucosaminyl transferase activity | Not Available |
| 2088 | Poly-beta-1, 6-N-acetyl-D-glucosamine synthase (Poly-beta-1, 6-GlcNAc synthase) (EC 2.4.1.-) | *Halomonas spp. TD01* | | F7SRR7 | -- | 22040376 | Acetyl glucosaminyl transferase activity | Not Available |
| 2089 | Poly-beta-1, 6-N-acetyl-D-glucosamine N-deacetylase (EC 3.5.1.-) | *Acinetobacter baumannii* | Strain AB307-0294 | A0A5K6CQ92 | -- | 18931120 | Hydrolase activity, acting on carbon-nitrogen (but not peptide) bonds | Not Available |
| 2090 | Putative major fimbrial subunit | *Salmonella typhimurium* | Strain LT2 / SGSC1412 / ATCC 700720 | Q8ZJU3 | -- | 11677609 | Not Available | Not Available |
| 2091 | Two-component response regulator | *Leptospira interrogans serogroup Icterohaemorrhagiae serovar Lai* | Strain 56601 | Q8F2R4 | -- | 12712204 | Diguanylate cyclase activity | Not Available |
| 2092 | Response regulator | *Leptospira interrogans serogroup Icterohaemorrhagiae serovar Lai* | Strain 56601 | Q8F629 | -- | 12712204 | Diguanylate cyclase activity | Not Available |
| 2093 | Poly-beta-1, 6-N-acetyl-D-glucosamine synthase (Poly-beta-1, 6-GlcNAc synthase) (EC 2.4.1.-) | *Acinetobacter baumannii* | Strain AB307-0294 | A0A5K6CPH3 | -- | 18931120 | Acetyl glucosaminyl transferase activity | Not Available |
| 2094 | Putative fimbriae major subunit | *Salmonella typhimurium* | Strain LT2 / SGSC1412 / ATCC 700720 | Q8ZRH9 | -- | 11677609 | Not Available | Not Available |
| 2095 | Putative fimbrial-like protein | *Salmonella typhimurium* | Strain LT2 / SGSC1412 / ATCC 700720 | Q7CQ93 | -- | 11677609 | Not Available | Not Available |
| 2096 | Putative fimbrial-like protein | *Salmonella typhimurium* | Strain LT2 / SGSC1412 / ATCC 700720 | Q8ZM89 | -- | 11677609 | Not Available | Not Available |
| 2097 | Fimbrial subunit | *Salmonella typhimurium* | Strain LT2 / SGSC1412 / ATCC 700720 | H9L412 | -- | 11677609 | Not Available | Not Available |
| 2098 | Surface composition regulator | *Escherichia fergusonii* | Strain ATCC 35469 / DSM 13698 / CCUG 18766 / IAM 14443 / JCM 21226 / LMG 7866 / NBRC 102419 / NCTC 12128 / CDC 0568-73 | B7LQC5 | -- | 19165319 | Not Available | Not Available |
| 2099 | Uncharacterized protein | *Priestia megaterium* | Strain ATCC 12872 / QMB1551 Bacillus megaterium | D5E339 | -- | 21705586 | Not Available | Not Available |
| 2100 | Poly-beta-1, 6-N-acetyl-D-glucosamine synthase (Poly-beta-1, 6-GlcNAc synthase) (EC 2.4.1.-) | *Escherichia coli O104:H4* | Strain 2011C-3493 | A0A0E0Y0U3 | -- | 23133618 | Acetyl glucosaminyl transferase activity | Not Available |
| 2101 | Putative PGA biosynthesis protein | *Escherichia coli O104:H4* | Strain 2011C-3493 | A0A0E0Y2B1 | -- | 23133618 | Not Available | Not Available |
| 2102 | Poly-beta-1, 6-N-acetyl-D-glucosamine synthase (Poly-beta-1, 6-GlcNAc synthase) (EC 2.4.1.-) | *Priestia megaterium* | Strain ATCC 12872 / QMB1551 Bacillus megaterium | D5E340 | -- | 21705586 | Acetyl glucosaminyl transferase activity | Not Available |
| 2103 | Poly-beta-1, 6-N-acetyl-D-glucosamine synthase (Poly-beta-1, 6-GlcNAc synthase) (EC 2.4.1.-) | *Klebsiella pneumoniae 30684 / NJST258_2* | | W8URY0 | -- | 24639510 | Acetyl glucosaminyl transferase activity | Not Available |
| 2104 | Secreted biofilm formation protein YqxM | *Bacillus sonorensis L12* | | M5P5Z6 | -- | 23538904 | Not Available | Not Available |
| 2105 | Outer membrane N-deacetylase | *Escherichia coli O104:H4* | Strain 2011C-3493 | A0A0E0Y2J4 | -- | 23133618 | Hydrolase activity, acting on carbon-nitrogen (but not peptide) bonds | Not Available |
| 2106 | Hemin storage protein HmsS | *Klebsiella pneumoniae 30684 / NJST258_2* | | W8VIB3 | -- | 24639510 | Not Available | Not Available |
| 2107 | Surface composition regulator | *Klebsiella pneumoniae 30684 / NJST258_2* | | W8UUA8 | -- | 24639510 | Not Available | Not Available |
| 2108 | Surface composition regulator | *Escherichia coli O104:H4* | Strain 2011C-3493 | A0A0E0XXK7 | -- | 23133618 | Not Available | Not Available |
| 2109 | mRNA interferase MqsR | *Gallionella spp.* | | A0A3D2NL57 | -- | 30148503 | Not Available | Not Available |
| 2110 | HmsF protein | *Klebsiella pneumoniae 30684 / NJST258_2* | | W8UPN5 | -- | 24639510 | Hydrolase activity, acting on carbon-nitrogen (but not peptide) bonds | Not Available |
| 2111 | Poly-beta-1, 6-N-acetyl-D-glucosamine synthase (Poly-beta-1, 6-GlcNAc synthase) (EC 2.4.1.-) | *Syntrophobacter spp. DG_60* | | A0A0S7XPJ3 | -- | 25922666 | Acetyl glucosaminyl transferase activity | Not Available |
| 2112 | Poly-beta-1, 6-N-acetyl-D-glucosamine biosynthesis protein PgaD | *Lentisphaerae bacterium GWF2_44_16* | | A0A1G0YRT2 | -- | 27774985 | Not Available | Not Available |
| 2113 | Poly-beta-1, 6-N-acetyl-D-glucosamine N-deacetylase PgaB | *Acinetobacter baumannii* | Strain AB0057 | A0A7U3Y5V9 | -- | 18931120 | Hydrolase activity, acting on carbon-nitrogen (but not peptide) bonds | Not Available |
| 2114 | Poly-beta-1, 6-N-acetyl-D-glucosamine synthase (Poly-beta-1, 6-GlcNAc synthase) (EC 2.4.1.-) | *Acinetobacter baumannii* | Strain AB0057 | A0A7U4DHS5 | -- | 18931120 | Acetyl glucosaminyl transferase activity | Not Available |
| 2115 | Uncharacterized protein | *Pseudomonas fluorescens* | Strain SBW25 | C3K4N7 | -- | 19432983 | Not Available | Not Available |
| 2116 | Poly-beta-1, 6-N-acetyl-D-glucosamine synthase (Poly-beta-1, 6-GlcNAc synthase) (EC 2.4.1.-) | *Staphylococcus aureus subspp. aureus 71193* | | A0A0E0VSR0 | -- | 22375071 | Acetyl glucosaminyl transferase activity | Not Available |
| 2117 | Poly-beta-1, 6-N-acetyl-D-glucosamine biosynthesis protein PgaD | *Acinetobacter baumannii* | Strain AB0057 | A0A7U3Y617 | -- | 18931120 | Not Available | Not Available |
| 2118 | Poly-beta-1, 6-N-acetyl-D-glucosamine N-deacetylase PgaB | *Acinetobacter baumannii* | Strain AB0057 | A0A7U3Y934 | -- | 18931120 | Hydrolase activity, acting on carbon-nitrogen (but not peptide) bonds | Not Available |
| 2119 | Poly-beta-1, 6-N-acetyl-D-glucosamine synthase (Poly-beta-1, 6-GlcNAc synthase) (EC 2.4.1.-) | *Pseudomonas fluorescens* | Strain SBW25 | C3KAF9 | -- | 19432983 | Acetyl glucosaminyl transferase activity | Not Available |
| 2120 | Poly-beta-1, 6-N-acetyl-D-glucosamine synthase (Poly-beta-1, 6-GlcNAc synthase) (EC 2.4.1.-) | *Paraburkholderia caribensis MBA4* | | A0A0P0RAV3 | -- | 24558235 | Acetyl glucosaminyl transferase activity | Not Available |
| 2121 | Poly-beta-1, 6-N-acetyl-D-glucosamine biosynthesis protein PgaD | *Acinetobacter baumannii* | Strain AB0057 | A0A7U3Y8Z6 | -- | 18931120 | Not Available | Not Available |
| 2122 | Type II toxin-antitoxin system MqsR family toxin | *Geobacter spp.* | | A0A3N9VGQ2 | -- | 30086797 | Not Available | Not Available |
| 2123 | Type II toxin-antitoxin system MqsR family toxin | *Geobacter spp.* | | A0A353MF81 | -- | 30148503 | Not Available | Not Available |
| 2124 | Poly-beta-1, 6-N-acetyl-D-glucosamine synthase (Poly-beta-1, 6-GlcNAc synthase) (EC 2.4.1.-) | *Clostridiaceae bacterium* | | A0A7C6H2P8 | -- | 32123542 | Acetyl glucosaminyl transferase activity | Not Available |
| 2125 | Poly-beta-1, 6-N-acetyl-D-glucosamine N-deacetylase PgaB | *Pseudomonas spp. GM18* | | J2W8M4 | -- | 23045501 | Hydrolase activity, acting on carbon-nitrogen (but not peptide) bonds | Not Available |
| 2126 | HmsF protein | *Klebsiella pneumoniae* | | W9BQH1 | -- | 25341126 | Hydrolase activity, acting on carbon-nitrogen (but not peptide) bonds | Not Available |
| 2127 | Putative membrane protein | *Bordetella parapertussis* | Strain Bpp5 | K0MBZ6 | -- | 23051057 | Not Available | Not Available |

| Target ID | Bacterial Protein Name | Organism Name | Strain | UniProt ID | PDB ID | PubMed ID | Classification as per UniProt GO Molecular Function | Classification of Function as per PDB |
|---|---|---|---|---|---|---|---|---|
| 2128 | AI-2 synthesis protein (EC 4.4.1.21) (Autoinducer-2 production protein LuxS) (S-ribosylhomocysteine lyase) | *Capnocytophaga ochracea* | Strain ATCC 27872 / DSM 7271 / JCM 12966 / NCTC 12371 / VPI 2845 Bacteroides ochraceus | C7M4X6 | -- | 21304645 | Iron ion binding, S-ribosylhomocysteine lyase activity | Not Available |
| 2129 | Putative hemin storage protein | *Bordetella parapertussis* | Strain Bpp5 | K0MFJ0 | -- | 23051057 | Hydrolase activity, acting on carbon-nitrogen (but not peptide) bonds | Not Available |
| 2130 | Poly-beta-1, 6-N-acetyl-D-glucosamine synthase (Poly-beta-1, 6-GlcNAc synthase) (EC 2.4.1.-) | *Bordetella parapertussis* | Strain Bpp5 | K0MDE7 | -- | 23051057 | Acetyl glucosaminyl transferase activity | Not Available |
| 2131 | Poly-beta-1, 6-N-acetyl-D-glucosamine synthase (Poly-beta-1, 6-GlcNAc synthase) (EC 2.4.1.-) | *Acinetobacter calcoaceticus* | Strain PHEA-2 | F0KG89 | -- | 21441526 | Acetyl glucosaminyl transferase activity | Not Available |
| 2132 | Uncharacterized protein | *Ralstonia solanacearum* | Strain UW551 | A3RPC6 | -- | 16404955 | Not Available | Not Available |
| 2133 | Uncharacterized protein | *Gordonibacter pamelaeae* | | A0A369M219 | -- | 29761785 | Not Available | Not Available |
| 2134 | Polysaccharide deacetylase | *Methylomona methanica* | Strain MC09 | G0A379 | -- | 22123758 | Hydrolase activity, acting on carbon-nitrogen (but not peptide) bonds | Not Available |
| 2135 | HmsF protein | *Ralstonia solanacearum* | Strain UW551 | A3RPC4 | -- | 16404955 | Hydrolase activity, acting on carbon-nitrogen (but not peptide) bonds | Not Available |
| 2136 | Poly-beta-1, 6-N-acetyl-D-glucosamine synthase (Poly-beta-1, 6-GlcNAc synthase) (EC 2.4.1.-) | *Acinetobacter calcoaceticus* | Strain PHEA-2 | F0KHT9 | -- | 21441526 | Acetyl glucosaminyl transferase activity | Not Available |
| 2137 | Biofilm PGA synthesis lipoprotein PgaB | *Acinetobacter calcoaceticus* | Strain PHEA-2 | F0KG88 | -- | 21441526 | Hydrolase activity, acting on carbon-nitrogen (but not peptide) bonds | Not Available |
| 2138 | Poly-beta-1, 6-N-acetyl-D-glucosamine synthase (Poly-beta-1, 6-GlcNAc synthase) (EC 2.4.1.-) | *Ralstonia solanacearum* | Strain UW551 | A3RPC5 | -- | 16404955 | Acetyl glucosaminyl transferase activity | Not Available |
| 2139 | Poly-beta-1, 6-N-acetyl-D-glucosamine synthase (Poly-beta-1, 6-GlcNAc synthase) (EC 2.4.1.-) | *Oceanisphaera profunda* | | A0A1Y0D3Z0 | -- | 24425742 | Acetyl glucosaminyl transferase activity | Not Available |
| 2140 | Uncharacterized protein | *Acinetobacter calcoaceticus* | Strain PHEA-2 | F0KHT8 | -- | 21441526 | Not Available | Not Available |
| 2141 | Poly-beta-1, 6-N-acetyl-D-glucosamine synthase (Poly-beta-1, 6-GlcNAc synthase) (EC 2.4.1.-) | *Bordetella pertussis STO1-CHOC-0019* | | A0A6N3RYK7 | -- | 24356839 | Acetyl glucosaminyl transferase activity | Not Available |
| 2142 | Poly-beta-1, 6-N-acetyl-D-glucosamine synthase (Poly-beta-1, 6-GlcNAc synthase) (EC 2.4.1.-) | *Yersinia pestis bv. Antiqua* | Strain Antiqua | A0A0H2Y673 | -- | 16740952 | Acetyl glucosaminyl transferase activity | Not Available |
| 2143 | Biofilm PGA synthesis lipoprotein PgaB | *Acinetobacter calcoaceticus* | Strain PHEA-2 | F0KHU0 | -- | 21441526 | Hydrolase activity, acting on carbon-nitrogen (but not peptide) bonds | Not Available |
| 2144 | Uncharacterized protein | *Acinetobacter calcoaceticus* | Strain PHEA-2 | F0KG90 | -- | 21441526 | Not Available | Not Available |
| 2145 | Poly-beta-1, 6-N-acetyl-D-glucosamine synthase (Poly-beta-1, 6-GlcNAc synthase) (EC 2.4.1.-) | *Rubrivivax gelatinosus* | Strain NBRC 100245 / IL144 | I0HLL7 | -- | 22689232 | Acetyl glucosaminyl transferase activity | Not Available |
| 2146 | Surface composition regulator | *Yersinia enterocolitica subspp. palearctica serotype O:3* | Strain DSM 13030 / CIP 106945 / Y11 | A0A0H3NPC2 | -- | 21296963 | DNA binding, DNA-directed DNA polymerase activity | Not Available |
| 2147 | Uncharacterized protein | *Pseudomonas spp. GM50* | | J3AUJ3 | -- | 23045501 | Not Available | Not Available |
| 2148 | Poly-beta-1, 6-N-acetyl-D-glucosamine synthase (Poly-beta-1, 6-GlcNAc synthase) (EC 2.4.1.-) | *Yersinia enterocolitica subspp. palearctica serotype O:3* | Strain DSM 13030 / CIP 106945 / Y11 | A0A0H3NQX9 | -- | 21296963 | Acetyl glucosaminyl transferase activity | Not Available |
| 2149 | Type II toxin-antitoxin system MqsR family toxin | *Pseudomonas mandelii* | | A0A502IHF4 | -- | 31206918 | Not Available | Not Available |
| 2150 | Poly-beta-1, 6-N-acetyl-D-glucosamine synthase (Poly-beta-1, 6-GlcNAc synthase) (EC 2.4.1.-) | *Pseudomonas spp. GM50* | | J2SHC7 | -- | 23045501 | Acetyl glucosaminyl transferase activity | Not Available |
| 2151 | Poly-beta-1, 6-N-acetyl-D-glucosamine biosynthesis protein PgaD | *Pseudomonas spp. GM50* | | J3GJ92 | -- | 23045501 | Not Available | Not Available |
| 2152 | Poly-beta-1, 6-N-acetyl-D-glucosamine N-deacetylase PgaB | *Pseudomonas spp. GM50* | | J3GJ29 | -- | 23045501 | Hydrolase activity, acting on carbon-nitrogen (but not peptide) bonds | Not Available |
| 2153 | Uncharacterized protein | *Pseudomonas furukawaii* | | L8MFN8 | -- | 22843571 | Not Available | Not Available |
| 2154 | Uncharacterized protein | *Acidithiobacillales bacterium SM23_46* | | A0A0S8GIC3 | -- | 25922666 | Not Available | Not Available |
| 2155 | Uncharacterized protein | *Methylococcaceae bacterium* | | A0A351V4P3 | -- | 30148503 | Not Available | Not Available |
| 2156 | Poly-beta-1, 6-N-acetyl-D-glucosamine N-deacetylase PgaB (Fragment) | *Methylococcaceae bacterium* | | A0A351KVS9 | -- | 30148503 | Hydrolase activity, acting on carbon-nitrogen (but not peptide) bonds | Not Available |
| 2157 | Poly-beta-1, 6-N-acetyl-D-glucosamine synthase (Poly-beta-1, 6-GlcNAc synthase) (EC 2.4.1.-) | *Acidithiobacillales bacterium SM23_46* | | A0A0S8GIQ5 | -- | 25922666 | Acetyl glucosaminyl transferase activity | Not Available |
| 2158 | Poly-beta-1, 6-N-acetyl-D-glucosamine biosynthesis protein PgaD (Fragment) | *Methylococcaceae bacterium* | | A0A351KVS6 | -- | 30148503 | Not Available | Not Available |
| 2159 | Poly-beta-1, 6-N-acetyl-D-glucosamine N-deacetylase PgaB | *Lautropia spp. SCN 69-89* | | A0A1D2SJQ0 | -- | 26031303 | Hydrolase activity, acting on carbon-nitrogen (but not peptide) bonds | Not Available |
| 2160 | Uncharacterized protein | *Burkholderia multivorans CGD2* | | B9BI89 | -- | 23105085 | Not Available | Not Available |
| 2161 | Poly-beta-1, 6-N-acetyl-D-glucosamine synthase (Poly-beta-1, 6-GlcNAc synthase) (EC 2.4.1.-) | *Mannheimia succiniciproducens* | Strain MBEL55E | Q65TJ1 | -- | 15378067 | Acetyl glucosaminyl transferase activity | Not Available |
| 2162 | Poly-beta-1, 6-N-acetyl-D-glucosamine synthase (Poly-beta-1, 6-GlcNAc synthase) (EC 2.4.1.-) | *Lautropia spp. SCN 69-89* | | A0A1D2SJP8 | -- | 26031303 | Acetyl glucosaminyl transferase activity | Not Available |
| 2163 | Outer membrane N-deacetylase | *Yersinia enterocolitica subspp. palearctica serotype O:3* | Strain YE-P4 | A0A7H4ZKR3 | -- | 23846271 | Hydrolase activity, acting on carbon-nitrogen (but not peptide) bonds | Not Available |
| 2164 | Outer membrane N-deacetylase | *Pectobacterium peruviense* | | A0A0J5XC35 | -- | 26251497 | Hydrolase activity, acting on carbon-nitrogen (but not peptide) bonds | Not Available |
| 2165 | Poly-beta-1, 6-N-acetyl-D-glucosamine synthase (Poly-beta-1, 6-GlcNAc synthase) (EC 2.4.1.-) | *Priestia megaterium* | Bacillus megaterium | A0A5B8PY98 | -- | 31228823 | Acetyl glucosaminyl transferase activity | Not Available |
| 2166 | Surface composition regulator | *Shimwellia blattae* | Strain ATCC 29907 / DSM 4481 / JCM 1650 / NBRC 105725 / CDC 9005-74 Escherichia blattae | I2B566 | -- | 22843577 | Not Available | Not Available |
| 2167 | Poly-beta-1, 6-N-acetyl-D-glucosamine biosynthesis protein PgaD | *Oceanisphaera profunda* | | A0A1Y0D3X8 | -- | 24425742 | Not Available | Not Available |
| 2168 | Surface composition regulator | *Cronobacter dublinensis 582* | | K8AVI0 | -- | 23166675 | Not Available | Not Available |
| 2169 | Polysaccharide deacetylase domain protein | *Burkholderia multivorans CGD2* | | B9BJ72 | -- | 23105085 | Hydrolase activity, acting on carbon-nitrogen (but not peptide) bonds | Not Available |
| 2170 | Poly-beta-1, 6-N-acetyl-D-glucosamine N-deacetylase PgaB | *Pseudomonas chlororaphis* | | A0A0D5Y8P3 | -- | 25679537 | Hydrolase activity, acting on carbon-nitrogen (but not peptide) bonds | Not Available |
| 2171 | Motility quorum-sensing regulator MqsR | *Pseudomonas chlororaphis* | | A0A0D5Y591 | -- | 25679537 | Not Available | Not Available |
| 2172 | GGDEF domain-containing protein | *Thermotoga maritima* | Strain ATCC 43589 / DSM 3109 / JCM 10099 / NBRC 100826 / MSB8 | Q9X0P6 | -- | 10360571 | Diguanylate cyclase activity | Not Available |
| 2173 | Biofilm PGA synthesis protein PgaD | *Pseudomonas chlororaphis* | | A0A0G3GEA7 | -- | 26634018 | Not Available | Not Available |
| 2174 | Poly-beta-1, 6-N-acetyl-D-glucosamine synthase (Poly-beta-1, 6-GlcNAc synthase) (EC 2.4.1.-) | *Stenotrophomonas spp. YAU14A_MKIMI4_1* | | A0A7U6AEA2 | -- | 29765358 | Acetyl glucosaminyl transferase activity | Not Available |

| Target ID | Bacterial Protein Name | Organism Name | Strain | UniProt ID | PDB ID | PubMed ID | Classification as per UniProt GO Molecular Function | Classification of Function as per PDB |
|---|---|---|---|---|---|---|---|---|
| 2175 | Biofilm PGA synthesis protein PgaD | *Pseudomonas chlororaphis* | | A0A0D5Y8I0 | -- | 25679537 | Not Available | Not Available |
| 2176 | Poly-beta-1, 6-N-acetyl-D-glucosamine synthase (Poly-beta-1, 6-GlcNAc synthase) (EC 2.4.1.-) | *Pseudomonas chlororaphis* | | A0A0G3G9K3 | -- | 26634018 | Acetyl glucosaminyl transferase activity | Not Available |
| 2177 | Outer membrane N-deacetylase | *Pseudomonas chlororaphis* | | A0A0G3GBF9 | -- | 26634018 | Hydrolase activity, acting on carbon-nitrogen (but not peptide) bonds | Not Available |
| 2178 | Uncharacterized protein | *Companilactobacillus mindensis DSM 14500* | | A0A0R1QRU5 | -- | 26415554 | Not Available | Not Available |
| 2179 | Motility quorum-sensing regulator MqsR | *candidate division NC10 bacterium* | | A0A2U1RPS3 | -- | 29662147 | Not Available | Not Available |
| 2180 | Outer membrane N-deacetylase | *Pseudomonas fluorescens HK44* | | A0A010T0D0 | -- | 21742869 | Hydrolase activity, acting on carbon-nitrogen (but not peptide) bonds | Not Available |
| 2181 | Poly-beta-1, 6-N-acetyl-D-glucosamine synthase (Poly-beta-1, 6-GlcNAc synthase) (EC 2.4.1.-) | *Tetragenococcus koreensis* | | A0A3G5F5R6 | -- | 16014460 | Acetyl glucosaminyl transferase activity | Not Available |
| 2182 | Poly-beta-1, 6-N-acetyl-D-glucosamine N-deacetylase | *Oceanisphaera profunda* | | A0A1Y0D9J9 | -- | 24425742 | Hydrolase activity, acting on carbon-nitrogen (but not peptide) bonds | Not Available |
| 2183 | Poly-beta-1, 6-N-acetyl-D-glucosamine synthase (Poly-beta-1, 6-GlcNAc synthase) (EC 2.4.1.-) | *Pseudomonas fluorescens HK44* | | A0A010RED1 | -- | 21742869 | Acetyl glucosaminyl transferase activity | Not Available |
| 2184 | Motility quorum-sensing regulator MqsR | *Pseudomonas fluorescens HK44* | | A0A010SPS9 | -- | 21742869 | Not Available | Not Available |
| 2185 | Lipoprotein for biofilm formation | *Bacillus subtilis subspp. subtilis* | Str. SC-8 | G4ETW0 | -- | 22207744 | Not Available | Not Available |
| 2186 | Biofilm PGA synthesis protein PgaD | *Pseudomonas fluorescens HK44* | | A0A010SD71 | -- | 21742869 | Not Available | Not Available |
| 2187 | Poly-beta-1, 6-N-acetyl-D-glucosamine synthase (Poly-beta-1, 6-GlcNAc synthase) (EC 2.4.1.-) | *Actinobacillus pleuropneumoniae serotype 3* | Strain JL03 | B0BTK4 | -- | 18197260 | Acetyl glucosaminyl transferase activity | Not Available |
| 2188 | Poly-beta-1, 6-N-acetyl-D-glucosamine synthase (Poly-beta-1, 6-GlcNAc synthase) (EC 2.4.1.-) | *Pseudomonas spp. WS 5027* | | A0A7Y1B5C2 | -- | 32587583 | Acetyl glucosaminyl transferase activity | Not Available |
| 2189 | Uncharacterized protein | *Aquitalea magnusonii* | | A0A3G9GES3 | -- | 28293292 | Not Available | Not Available |
| 2190 | Surface composition regulator | *Salmonella typhimurium* | Strain 14028S / SGSC 2262 | A0A0F6B6X7 | -- | 19897643 | Not Available | Not Available |
| 2191 | Flagellar Transcriptional regulator FlhC | *Salmonella typhimurium* | Strain 14028S / SGSC 2262 | A0A0F6B2Q8 | -- | 19897643 | DNA binding, zinc ion binding | Not Available |
| 2192 | Flagellar Transcriptional regulator FlhD | *Salmonella typhimurium* | Strain 14028S / SGSC 2262 | A0A0F6B2Q9 | -- | 19897643 | DNA binding | Not Available |
| 2193 | Poly-beta-1, 6-N-acetyl-D-glucosamine biosynthesis protein PgaD | *Pseudomonas spp. WS 5027* | | A0A7Y1B5A3 | -- | 32587583 | Not Available | Not Available |
| 2194 | Poly-beta-1, 6-N-acetyl-D-glucosamine synthase (Poly-beta-1, 6-GlcNAc synthase) (EC 2.4.1.-) | *Thiotrichales bacterium SG8_50* | | A0A0S8ELF2 | -- | 25922666 | Acetyl glucosaminyl transferase activity | Not Available |
| 2195 | Uncharacterized protein | *Thiotrichales bacterium SG8_50* | | A0A0S8EME4 | -- | 25922666 | Not Available | Not Available |
| 2196 | Transcriptional regulatory protein SrrA (Staphylococcal respiratory response protein A) | *Staphylococcus aureus* | Strain Mu50 / ATCC 700699 | Q7A2R6 | -- | 11418146 | DNA binding | Not Available |
| 2197 | Sensor protein SrrB (EC 2.7.13.3) (Staphylococcal respiratory response protein B) | *Staphylococcus aureus* | Strain Mu50 / ATCC 700699 | Q99TZ9 | -- | 11418146 | ATP binding, phosphorelay sensor kinase activity | Not Available |
| 2198 | Transcriptional regulatory protein SrrA (Staphylococcal respiratory response protein A) | *Staphylococcus aureus* | Strain MSSA476 | Q6G972 | -- | 15213324 | DNA binding | Not Available |
| 2199 | Transcriptional regulatory protein SrrA (Staphylococcal respiratory response protein A) | *Staphylococcus aureus* | | Q9L524 | -- | 11157922; 15060046 | DNA binding | Not Available |
| 2200 | Transcriptional regulatory protein SrrA (Staphylococcal respiratory response protein A) | *Staphylococcus aureus* | Strain MW2 | Q7A0U4 | -- | 12044378 | DNA binding | Not Available |
| 2201 | Diguanylate cyclase VdcA (DGC) (EC 2.7.7.65) | *Vibrio cholerae serotype O1* | Strain ATCC 39315 / El Tor Inaba N16961 | Q9KKZ4 | -- | 10952301; 18227161 | Diguanylate cyclase activity, GTP binding, Metal ion binding | Not Available |
| 2202 | signaltransduction protein TRAP (Target of RNAIII-activating protein) | *Staphylococcus aureus* | Strain USA300 | Q2FFR1 | -- | 16517273 | Not Available | Not Available |
| 2203 | signaltransduction protein TRAP (Target of RNAIII-activating protein) | *Staphylococcus aureus* | Strain bovine RF122 / ET3-1 | Q2YU04 | -- | 17971880 | Not Available | Not Available |
| 2204 | signaltransduction protein TRAP (Target of RNAIII-activating protein) | *Staphylococcus aureus* | Strain COL | Q5HEU0 | -- | 15774886 | Not Available | Not Available |
| 2205 | signaltransduction protein TRAP (Target of RNAIII-activating protein) | *Staphylococcus aureus* | Strain Mu50 / ATCC 700699 | Q7A2Q4 | -- | 11418146 | Not Available | Not Available |
| 2206 | signaltransduction protein TRAP (Target of RNAIII-activating protein) | *Staphylococcus aureus* | Strain N315 | Q7A4W3 | -- | 11418146 | Not Available | Not Available |
| 2207 | signaltransduction protein TRAP (Target of RNAIII-activating protein) | *Staphylococcus aureus* | Strain MRSA252 | Q6GFM2 | -- | 15213324 | Not Available | Not Available |
| 2208 | signaltransduction protein TRAP (Target of RNAIII-activating protein) | *Staphylococcus aureus* | Strain MSSA476 | Q6G8A1 | -- | 15213324 | Not Available | Not Available |
| 2209 | signaltransduction protein TRAP (Target of RNAIII-activating protein) | *Staphylococcus aureus* | Strain MW2 | Q8NVW1 | -- | 12044378 | Not Available | Not Available |
| 2210 | signaltransduction protein TRAP (Target of RNAIII-activating protein) | *Staphylococcus epidermidis* | | Q8GQQ1 | -- | 12599079 | Not Available | Not Available |
| 2211 | signaltransduction protein TRAP (Target of RNAIII-activating protein) | *Staphylococcus epidermidis* | Strain ATCC 35984 / RP62A | Q5HNA3 | -- | 15774886; 16842355 | Not Available | Not Available |
| 2212 | RNA polymerase sigma-54 factor | *Pseudomonas aeruginosa* | Strain ATCC 15692 / DSM 22644 / CIP 104116 / JCM 14847 / LMG 12228 / 1C / PRS 101 / PAO1 | P49988 | -- | 8113171; 10984043; 2152909; 9595661; 12644493; 10792721; 26633362; 29760208 | DNA-binding Transcription activator activity, DNA-directed 5'-3' RNA polymerase activity, sigma factor activity, Transcription cis-regulatory region binding | Not Available |
| 2213 | Uncharacterized fimbrial-like protein SfmA (Type-1A pilin) | *Escherichia coli* | Strain K12 | P0ABW5 | -- | 9278503; 16738553; 20345943 | Not Available | Not Available |
| 2214 | Protein translocase subunit SecA 2 (EC 7.4.2.8) | *Streptococcus pneumoniae serotype 4* | Strain ATCC BAA-334 / TIGR4 | Q97P83 | -- | 11463916; 16861665; 18507531; 20714350; 28456649 | ATP binding, protein-exporting ATPase activity | Not Available |
| 2215 | Sensor protein SrrB (EC 2.7.13.3) (Staphylococcal respiratory response protein B) | *Staphylococcus aureus* | Strain COL | Q5HFT1 | -- | 15774886; 16859493; 17384184; 24222487 | ATP binding, phosphorelay sensor kinase activity | Not Available |
| 2216 | Transcriptional regulatory protein LytR (Sensory transduction protein LytR) | *Staphylococcus aureus* | Strain USA300 | Q2FK09 | -- | 16517273 | DNA binding | Not Available |
| 2217 | Transcriptional regulatory protein LytR (Sensory transduction protein LytR) | *Staphylococcus aureus* | Strain bovine RF122 / ET3-1 | Q2YV67 | -- | 17971880 | DNA binding | Not Available |
| 2218 | Transcriptional regulatory protein LytR (Sensory transduction protein LytR) | *Staphylococcus aureus* | Strain COL | Q5HJB5 | -- | 15774886 | DNA binding | Not Available |
| 2219 | Transcriptional regulatory protein LytR (Sensory transduction protein LytR) | *Staphylococcus aureus* | Strain Mu50 / ATCC 700699 | P60609 | -- | 11418146; 27127614 | DNA binding | Not Available |
| 2220 | Transcriptional regulatory protein LytR (Sensory transduction protein LytR) | *Staphylococcus aureus* | Strain N315 | P60610 | -- | 11418146 | DNA binding | Not Available |

| Target ID | Bacterial Protein Name | Organism Name | Strain | UniProt ID | PDB ID | PubMed ID | Classification as per UniProt GO Molecular Function | Classification of Function as per PDB |
|---|---|---|---|---|---|---|---|---|
| 2221 | Transcriptional regulatory protein LytR (Sensory transduction protein LytR) | *Staphylococcus aureus* | Strain MRSA252 | Q6GK51 | -- | 15213324 | DNA binding | Not Available |
| 2222 | Transcriptional regulatory protein LytR (Sensory transduction protein LytR) | *Staphylococcus aureus* | Strain MSSA476 | Q6GCL1 | -- | 15213324 | DNA binding | Not Available |
| 2223 | Transcriptional regulatory protein LytR (Sensory transduction protein LytR) | *Staphylococcus aureus* | Strain MW2 | Q8NYH3 | -- | 12044378 | DNA binding | Not Available |
| 2224 | Sensor histidinekinase / phosphatase LytS (EC 2.7.13.3) (EC 3.1.3.-) (Autolysin sensorkinase ) | *Staphylococcus aureus* | Strain USA300 | Q2FK10 | -- | 16517273 | ATP binding, Hydrolase activity, phosphorelay sensor kinase activity | Not Available |
| 2225 | Sensor histidinekinase / phosphatase LytS (EC 2.7.13.3) (EC 3.1.3.-) (Autolysin sensorkinase ) | *Staphylococcus aureus* | Strain NCTC 8325 / PS 47 | Q53705 | -- | 8550490; 9308171; 19502411; 23733465; 25491472 | ATP binding, Hydrolase activity, phosphorelay sensor kinase activity | Not Available |
| 2226 | Sensor histidinekinase / phosphatase LytS (EC 2.7.13.3) (EC 3.1.3.-) (Autolysin sensorkinase ) | *Staphylococcus aureus* | Strain bovine RF122 / ET3-1 | Q2YV68 | -- | 17971880 | ATP binding, Hydrolase activity, phosphorelay sensor kinase activity | Not Available |
| 2227 | Sensor histidinekinase / phosphatase LytS (EC 2.7.13.3) (EC 3.1.3.-) (Autolysin sensorkinase ) | *Staphylococcus aureus* | Strain COL | Q5HJB6 | -- | 15774886 | ATP binding, Hydrolase activity, phosphorelay sensor kinase activity | Not Available |
| 2228 | Sensor histidinekinase / phosphatase LytS (EC 2.7.13.3) (EC 3.1.3.-) (Autolysin sensorkinase ) | *Staphylococcus aureus* | Strain Mu50 / ATCC 700699 | P60612 | -- | 11418146; 27127614 | ATP binding, Hydrolase activity, phosphorelay sensor kinase activity | Not Available |
| 2229 | Sensor histidinekinase / phosphatase LytS (EC 2.7.13.3) (EC 3.1.3.-) (Autolysin sensorkinase ) | *Staphylococcus aureus* | Strain N315 | P60613 | -- | 11418146 | ATP binding, Hydrolase activity, phosphorelay sensor kinase activity | Not Available |
| 2230 | Sensor histidinekinase / phosphatase LytS (EC 2.7.13.3) (EC 3.1.3.-) (Autolysin sensorkinase ) | *Staphylococcus aureus* | Strain MRSA252 | Q6GK52 | -- | 15213324 | ATP binding, Hydrolase activity, phosphorelay sensor kinase activity | Not Available |
| 2231 | Sensor histidinekinase / phosphatase LytS (EC 2.7.13.3) (EC 3.1.3.-) (Autolysin sensorkinase ) | *Staphylococcus aureus* | Strain MSSA476 | Q6GCL2 | -- | 15213324 | ATP binding, Hydrolase activity, phosphorelay sensor kinase activity | Not Available |
| 2232 | Sensor histidinekinase / phosphatase LytS (EC 2.7.13.3) (EC 3.1.3.-) (Autolysin sensorkinase ) | *Staphylococcus aureus* | Strain MW2 | P60614 | -- | 12044378 | ATP binding, Hydrolase activity, phosphorelay sensor kinase activity | Not Available |
| 2233 | Major fimbrium subunit FimA type-2 (Fimbrillin) (Fimbrilin) (Major fimbrial subunit protein type II) | *Porphyromonas gingivalis* |  | Q51822 | -- | 7902712; 1987052; 11748193; 23809984 | Structural protein | Not Available |
| 2234 | Major fimbrium anchoring subunit FimB | *Porphyromonas gingivalis* | Strain ATCC BAA-308 / W83 | Q7MT55 | -- | 12949112; 20530728 | Not Available | Not Available |
| 2235 | Transcriptionalactivator protein AnoR | *Acinetobacter nosocomialis* |  | A0A0N8YGA2 | -- | 25975610 | DNA binding | Not Available |
| 2236 | Major fimbrium tip subunit FimD | *Porphyromonas gingivalis* |  | O32389 | -- | 8981345; 17526848 | Host cell extracellular matrix binding | Not Available |
| 2237 | Minor curlin subunit | *Escherichia coli* | Strain K12 | P0ABK7 | -- | 8817489; 8905232; 9278503; 16738553; 7854117; 17010156; 24097954 | Identical protein binding | Not Available |
| 2238 | Fimbrial subunit ElfA | *Escherichia coli* | Strain K12 | P75855 | -- | 8905232; 9278503; 16738553; 20345943 | Not Available | Not Available |
| 2239 | Uncharacterized fimbrial-like protein ElfG | *Escherichia coli* | Strain K12 | P75858 | -- | 8905232; 9278503; 16738553; 20345943 | Not Available | Not Available |
| 2240 | Major fimbrium subunit FimA type-1 (FimA1) (Fimbrillin) (Fimbrilin) (Major fimbrial subunit protein type I) | *Porphyromonas gingivalis* |  | P0C940 | -- | 2895100; 7902712; 8981345; 1987052; 11748193; 20530728; 23809984 | Structural protein | Not Available |
| 2241 | Probable diguanylate cyclase DgcC (DGC) (EC 2.7.7.65) | *Escherichia coli* | Strain K12 | P0AAP1 | -- | 9278503; 16738553; 3533724; 11260463; 16513732; 17010156; 19332833; 19707751; 26148715 | Diguanylate cyclase activity, GTP binding, Metal ion binding | Not Available |
| 2242 | Probable diguanylate cyclase DgcE (DGC) (EC 2.7.7.65) | *Escherichia coli* | Strain K12 | P38097 | -- | 9097040; 9278503; 16738553; 6094528; 7984428; 18765794; 19332833; 20303158; 23708798; 26148715 | Diguanylate cyclase activity, GTP binding, Metal ion binding | Not Available |
| 2243 | Probable diguanylate cyclase DgcF (DGC) (EC 2.7.7.65) | *Escherichia coli* | Strain K12 | P76147 | -- | 9097039; 9278503; 16738553; 21181144; 26148715 | Diguanylate cyclase activity, GTP binding, Metal ion binding | Not Available |
| 2244 | Probable diguanylate cyclase DgcI (DGC) (EC 2.7.7.65) (Putative lipoprotein DgcI) | *Escherichia coli* | Strain K12 | P75801 | -- | 8905232; 9278503; 16738553; 26148715 | Diguanylate cyclase activity, GTP binding, Metal ion binding | Not Available |
| 2245 | Probable diguanylate cyclase DgcJ (DGC) (EC 2.7.7.65) | *Escherichia coli* | Strain K12 | P76237 | -- | 9097040; 9278503; 16738553; 18765794; 19332833; 21181144; 26148715 | Diguanylate cyclase activity, GTP binding, Metal ion binding | Not Available |
| 2246 | Diguanylate cyclase DgcM (DGC) (EC 2.7.7.65) | *Escherichia coli* | Strain K12 | P77302 | -- | 9097039; 9278503; 16738553; 17010156; 19332833; 23708798; 26148715 | Diguanylate cyclase activity, GTP binding, Identical protein binding, Metal ion binding | Not Available |

| Target ID | Bacterial Protein Name | Organism Name | Strain | UniProt ID | PDB ID | PubMed ID | Classification as per UniProt GO Molecular Function | Classification of Function as per PDB |
|---|---|---|---|---|---|---|---|---|
| 2247 | Diguanylate cyclase DgcN (DGC) (EC 2.7.7.65) | *Escherichia coli* | Strain K12 | P46139 | -- | 9205837; 9278503; 16738553; 6357787; 7567469; 15919996; 19332833; 20303158; 21181144; 26148715; 27507823 | Diguanylate cyclase activity, GTP binding, Identical protein binding, Metal ion binding | Not Available |
| 2248 | Diguanylate cyclase DgcP (EC 2.7.7.65) | *Pseudomonas aeruginosa* | Strain ATCC 15692 / DSM 22644 / CIP 104116 / JCM 14847 / LMG 12228 / 1C / PRS 101 / PAO1 | Q9HT84 | -- | 10984043; 25118352; 25809128, 2152909 | Diguanylate cyclase activity, GTP binding, Metal ion binding | Not Available |
| 2249 | Probable diguanylate cyclase DgcQ (DGC) (EC 2.7.7.65) (Cellulose synthesis regulatory protein) | *Salmonella typhimurium* | Strain LT2 / SGSC1412 / ATCC 700720 | Q8ZNT5 | -- | 11677609 | Diguanylate cyclase activity, GTP binding, Metal ion binding | Not Available |
| 2250 | Putative glycosylTransferase EpsH (EC 2.4.-.-) | *Bacillus subtilis* | Strain 168 | P71057 | -- | 8969506; 9384377; 15083514; 15175311; 15661000 | | Not Available |
| 2251 | Putative glycosylTransferase EpsD (EC 2.4.-.-) | *Bacillus subtilis* | Strain 168 | P71053 | -- | 8969506; 9384377; 15175311; 15661000 | | Not Available |
| 2252 | Minor fimbrium subunit Mfa1 (53 kDa major outer Membrane protein) | *Porphyromonas gingivalis* | | P81363 | -- | 9569663; 8208139; 26001707 | Not Available | Not Available |
| 2253 | Probable diguanylate cyclase CdgI (DGC) (EC 2.7.7.65) | *Escherichia coli* | Strain K12 | P76236 | -- | 9097040; 9278503; 16738553; 17010156; 15919996; 21181144; 26148715 | Diguanylate cyclase activity, GTP binding, Metal ion binding | Not Available |
| 2254 | TransMembrane protein EpsG | *Bacillus subtilis* | Strain 168 | P71056 | -- | 8969506; 9384377; 15175311; 15661000 | Not Available | Not Available |
| 2255 | Fimbrin-like protein FimI | *Salmonella typhimurium* | Strain LT2 / SGSC1412 / ATCC 700720 | P37922 | -- | 11677609 | Not Available | Not Available |
| 2256 | Fimbrial-like protein FimF | *Salmonella typhimurium* | Strain LT2 / SGSC1412 / ATCC 700720 | P37926 | -- | 11677609 | Not Available | Not Available |
| 2257 | Major fimbrium subunit FimC | *Porphyromonas gingivalis* | | O32388 | -- | 8981345; 17526848 | Host cell extracellular matrix binding | Not Available |
| 2258 | CRISPR-associated endonuclease Cas6 / Csy4 (EC 3.1.-.-) (crRNA endonuclease) | *Escherichia coli* | Strain UTI89 / UPEC | Q1RE32 | -- | 16585510; 21398535 | Endonuclease activity, RNA binding | Not Available |
| 2259 | Fibronectin-binding protein B | *Staphylococcus aureus* | Strain USA300 | A0A0H2XKG3 | -- | 16517273; 15234962; 21569203; 24628034; 27387503; 30622139 | Not Available | Not Available |
| 2260 | Type-1 fimbrial protein, A chain (Type-1A pilin) | *Salmonella typhimurium* | Strain LT2 / SGSC1412 / ATCC 700720 | P37921 | -- | 11677609 | Not Available | Not Available |
| 2261 | DEAD-box ATP-dependent RNA helicase CshA (EC 3.6.4.13) | *Staphylococcus aureus* | Strain NCTC 8325 / PS 47 | Q2FWH5 | -- | 21764917; 22447609; 23229022 | ATP binding, ATP hydrolysis activity, RNA binding, RNA helicase activity, RNA strand annealing activity | Not Available |
| 2262 | ComX pheromone (Competence pheromone) | *Bacillus subtilis subspp. natto* | Strain BEST195 | D4G0R3 | -- | 20398357; 25329997; 25855042; 26276536; 29665236 | Not Available | Not Available |
| 2263 | Putative glycosylTransferase EpsE (EC 2.4.-.-) | *Bacillus subtilis* | Strain 168 | P71054 | -- | 8969506; 9384377; 19383706; 15175311; 15661000 | | Not Available |
| 2264 | Putative pyridoxal phosphate-dependent aminoTransferase EpsN (EC 2.6.1.-) | *Bacillus subtilis* | Strain 168 | Q795J3 | -- | 8969506; 9384377; 15661000 | Pyridoxal phosphate binding, transaminae activity | Not Available |
| 2265 | Type 1 fimbrin D-mannose specific adhesin (Protein FimH) | *Salmonella typhimurium* | Strain LT2 / SGSC1412 / ATCC 700720 | P37925 | -- | 11677609 | Not Available | Not Available |
| 2266 | Sensor histidinekinase ArsS (EC 2.7.13.3) | *Helicobacter pylori* | Strain ATCC 700392 / 26695 Campylobacter pylori | O24972 | -- | 9252185; 10735847; 15109719; 16672598; 22865848; 25997502; 27432830 | Phosphorelay sensor kinase activity | Not Available |
| 2267 | Major fimbrium subunit FimC | *Porphyromonas gingivalis* | Strain ATCC 33277 / DSM 20709 / CIP 103683 / JCM 12257 / NCTC 11834 / 2561 | B2RH57 | -- | 18524787; 17675496; 17526848; 19506009 | Host cell extracellular matrix binding | Not Available |
| 2268 | Response regulator GacA (Global activator) | *Pseudomonas aeruginosa* | Strain ATCC 15692 / DSM 22644 / CIP 104116 / JCM 14847 / LMG 12228 / 1C / PRS 101 / PAO1 | Q51373 | -- | 9159518; 10984043, 2152909 | DNA binding | Not Available |
| 2269 | Major fimbrium tip subunit FimE | *Porphyromonas gingivalis* | Strain ATCC 33277 / DSM 20709 / CIP 103683 / JCM 12257 / NCTC 11834 / 2561 | B2RH59 | -- | 18524787; 17675496; 17526848; 19506009 | Host cell extracellular matrix binding | Not Available |
| 2270 | Long polar fimbria protein A | *Salmonella typhimurium* | Strain LT2 / SGSC1412 / ATCC 700720 | P43660 | -- | 7721701; 11677609 | Not Available | Not Available |

| Target ID | Bacterial Protein Name | Organism Name | Strain | UniProt ID | PDB ID | PubMed ID | Classification as per UniProt GO Molecular Function | Classification of Function as per PDB |
|---|---|---|---|---|---|---|---|---|
| 2271 | Putative pyruvyl Transferase EpsO (EC 2.-.-.-) | *Bacillus subtilis* | Strain 168 | P71065 | -- | 8969506; 9384377; 15661000 | Transferase activity | Not Available |
| 2272 | Response regulator protein FleR | *Pseudomonas aeruginosa* | Strain ATCC 15692 / DSM 22644 / CIP 104116 / JCM 14847 / LMG 12228 / 1C / PRS 101 / PAO1 | Q9I4N3 | -- | 10984043; 7591148; 8876537; 30877999, 2152909 | ATP binding, Sequence-specific DNA binding | Not Available |
| 2273 | Major fimbrium anchoring subunit FimB | *Porphyromonas gingivalis* | Strain ATCC 33277 / DSM 20709 / CIP 103683 / JCM 12257 / NCTC 11834 / 2561 | A0PA81 | -- | 17081195; 20530728 | Not Available | Not Available |
| 2274 | Putative glycosylTransferase EpsF (EC 2.4.-.-) | *Bacillus subtilis* | Strain 168 | P71055 | -- | 8969506; 9384377; 15175311; 15661000 | | Not Available |
| 2275 | Major fimbrium subunit FimA type-3 (Fimbrillin) (Fimbrilin) (Major fimbrial subunit protein type III) | *Porphyromonas gingivalis* | | Q51826 | -- | 7902712; 11748193 | Structural protein | Not Available |
| 2276 | Major fimbrium subunit FimA type-2 (Fimbrillin) (Fimbrilin) (Major fimbrial subunit protein type II) | *Porphyromonas gingivalis* | | Q51825 | -- | 7902712; 23809984 | Structural protein | Not Available |
| 2277 | Major fimbrium tip subunit FimD | *Porphyromonas gingivalis* | Strain ATCC 33277 / DSM 20709 / CIP 103683 / JCM 12257 / NCTC 11834 / 2561 | B2RH58 | -- | 18524787; 17675496; 17526848; 19506009 | Host cell extracellular matrix binding | Not Available |
| 2278 | AcylTransferase PE (EC 2.3.1.-) (Proline-glutamic acid-rich protein) (PE protein) | *Mycolicibacterium smegmatis* | Strain ATCC 700084 / mc2155 Mycobacterium smegmatis | Q2M5K2 | -- | 17295914; 18955433; 32554804; 27825305; 27028886 | AcylTransferase activity | Not Available |
| 2279 | RNA polymerase sigma-54 factor | *Pseudomonas aeruginosa* | Strain UCBPP-PA14 | A0A0H2ZGR9 | -- | 17038190; 11717271 | DNA binding, DNA-binding Transcription activator activity, DNA-directed 5'-3' RNA polymerase activity, sigma factor activity | Not Available |
| 2280 | Major fimbrium subunit FimA type-5 (Fimbrillin) (Fimbrilin) (Major fimbrial subunit protein type V) | *Porphyromonas gingivalis* | | Q9S0W8 | -- | 10790120; 11748193 | Structural protein | Not Available |
| 2281 | Pneumolysin (PLY) (Thiol-activated cytolysin) | *Streptococcus pneumoniae serotype 4* | Strain ATCC BAA-334 / TIGR4 | P0C2J9 | -- | 11463916; 28456649 | Cholesterol binding, toxin activity | Not Available |
| 2282 | Rqc2 homolog RqcH (RqcH) (Fibronectin-binding protein YloA) | *Bacillus subtilis subspp. natto* | Strain BEST195 | D4FWG0 | -- | 20398357; 25329997; 28294433 | Ribosomal large subunit binding, tRNA binding | Not Available |
| 2283 | Transcriptional regulatory protein SrrA (Staphylococcal respiratory response protein A) | *Staphylococcus aureus* | Strain MRSA252 | Q6GGK6 | -- | 15213324 | DNA binding | Not Available |
| 2284 | Serine-aspartate repeat-containing protein C | *Staphylococcus aureus* | Strain Newman | O86487 | -- | 9884231; 17951380; 20090838; 25115812; 28320940 | Not Available | Not Available |
| 2285 | Global Transcriptional regulator Spx | *Staphylococcus aureus* | Strain NCTC 8325 / PS 47 | Q2G1U6 | -- | 16788195; 23629700; 32117138 | Not Available | Not Available |
| 2286 | Transcription antitermiNot Availabletion protein RfaH | *Escherichia coli O6:K15:H31* | Strain 536 / UPEC | Q0TAL4 | -- | 16879640; 16452414 | DNA binding, Transcription antitermination factor activity, DNA binding | Not Available |
| 2287 | HTH-type Transcriptional regulator MlrA (MerR-like regulator A) | *Escherichia coli* | Strain K12 | P33358 | -- | 11489123; 9278503; 16738553; 20874755; 23708798 | DNA binding, DNA-binding Transcription factor activity | Not Available |
| 2288 | Uncharacterized fimbrial-like protein SfmF | *Escherichia coli* | Strain K12 | P38052 | -- | 9278503; 16738553; 2183007; 7984428; 20345943 | Not Available | Not Available |
| 2289 | Transcriptional regulatory protein SrrA (Staphylococcal respiratory response protein A) | *Staphylococcus aureus* | Strain COL | Q5HFT0 | -- | 15774886; 16859493; 17384184; 24222487 | DNA binding | Not Available |
| 2290 | Transcriptional regulatory protein SrrA (Staphylococcal respiratory response protein A) | *Staphylococcus aureus* | Strain N315 | Q7A5H6 | -- | 11418146; 15590099 | DNA binding | Not Available |
| 2291 | Uncharacterized fimbrial-like protein YraH | *Escherichia coli* | Strain K12 | P42913 | -- | 9278503; 16738553; 20345943 | Not Available | Not Available |
| 2292 | Uncharacterized fimbrial-like protein SfmH | *Escherichia coli* | Strain K12 | P75715 | -- | 9278503; 16738553; 20345943 | mannose binding | Not Available |
| 2293 | Accessory Sec system protein translocase subunit SecY2 | *Streptococcus pneumoniae serotype 4* | Strain ATCC BAA-334 / TIGR4 | Q97P79 | -- | 11463916; 16861665; 18507531; 20714350 | Not Available | Not Available |
| 2294 | Sensor protein SrrB (EC 2.7.13.3) (Staphylococcal respiratory response protein B) | *Staphylococcus aureus* | Strain MW2 | Q8NWF3 | -- | 12044378 | ATP binding, phosphorelay sensor kinase activity | Not Available |
| 2295 | Sensor protein SrrB (EC 2.7.13.3) (Staphylococcal respiratory response protein B) | *Staphylococcus aureus* | Strain N315 | Q7A5H7 | -- | 11418146 | ATP binding, phosphorelay sensor kinase activity | Not Available |
| 2296 | Sensor protein SrrB (EC 2.7.13.3) (Staphylococcal respiratory response protein B) | *Staphylococcus aureus* | Strain MRSA252 | Q6GGK7 | -- | 15213324 | ATP binding, phosphorelay sensor kinase activity | Not Available |
| 2297 | Major fimbrium anchoring subunit FimB | *Porphyromonas gingivalis* | | A0PA72 | -- | 17081195; 20530728 | Not Available | Not Available |
| 2298 | Minor fimbrium subunit Mfa1 (53 kDa fimbrilin) | *Porphyromonas gingivalis* | | P0DOA1 | -- | 26001707 | Not Available | Not Available |
| 2299 | Sensor protein VraS (EC 2.7.13.3) | *Staphylococcus aureus* | Strain MRSA252 | Q6GFH2 | -- | 15213324 | ATP binding, phosphorelay sensor kinase activity, protein dimerization activity | Not Available |
| 2300 | 23S rRNA (cytidine-2'-O)-methylTransferase TlyA (EC 2.1.1.226) (23S rRNA (cytidine1920-2'-O)-methylTransferase) (Hemolysin A) | *Campylobacter jejuni subspp. jejuni serotype O:23 / 36* | Strain 81-176 | A0A0H3PEK7 | -- | 18389311; 23971210; 24796671; 29404277; 32134554 | MethylTransferase activity, RNA binding | Not Available |
| 2301 | Sensor protein SrrB (EC 2.7.13.3) (Staphylococcal respiratory response protein B) | *Staphylococcus aureus* | Strain MSSA476 | Q6G973 | -- | 15213324 | ATP binding, phosphorelay sensor kinase activity | Not Available |

| Target ID | Bacterial Protein Name | Organism Name | Strain | UniProt ID | PDB ID | PubMed ID | Classification as per UniProt GO Molecular Function | Classification of Function as per PDB |
|---|---|---|---|---|---|---|---|---|
| 2302 | Probable autotransporter YfaL [Cleaved into: Probable secreted autotransporter protein YfaL; Probable autotransporter YfaL translocator] | *Escherichia coli* | Strain K12 | P45508 | -- | 9205837; 9278503; 16738553; 6087316; 2834621; 7567469; 16522795; 22344647 | serine-type peptidase activity | Not Available |
| 2303 | Uncharacterized fimbrial-like protein YadM | *Escherichia coli* | Strain K12 | P37018 | -- | 8202364; 9278503; 16738553; 20345943 | Not Available | Not Available |
| 2304 | Uncharacterized protein YhcK | *Bacillus subtilis* | Strain 168 | P54595 | -- | 8969498; 9384377 | Diguanylate cyclase activity, phosphorelay sensor kinase activity | Not Available |
| 2305 | Uncharacterized fimbrial-like protein YbgD | *Escherichia coli* | Strain K12 | P37909 | -- | 8905232; 9278503; 16738553; 7984428 | Not Available | Not Available |
| 2306 | Uncharacterized fimbrial-like protein YfcQ | *Escherichia coli* | Strain K12 | P76500 | -- | 9278503; 16738553; 20345943 | Not Available | Not Available |
| 2307 | Autotransporter adhesin UpaG (Type 5 secretion system autotransporter YadA) | *Escherichia coli O6:H1* | Strain CFT073 / ATCC 700928 / UPEC | A0A0H2VCA1 | -- | 12471157; 18424525; 22286983 | Not Available | Not Available |
| 2308 | Uncharacterized fimbrial-like protein YadL | *Escherichia coli* | Strain K12 | P37017 | -- | 8202364; 9278503; 16738553; 20345943 | Not Available | Not Available |
| 2309 | Uncharacterized fimbrial-like protein YcbV | *Escherichia coli* | Strain K12 | P75860 | -- | 8905232; 9278503; 16738553; 20345943 | Not Available | Not Available |
| 2310 | Sensor protein VraS (EC 2.7.13.3) | *Staphylococcus aureus* | Strain MW2 | Q7A0H9 | -- | 12044378 | ATP binding, phosphorelay sensor kinase activity, protein dimerization activity | Not Available |
| 2311 | Uncharacterized fimbrial-like protein YehD | *Escherichia coli* | Strain K12 | P33343 | -- | 9278503; 16738553; 20345943 | Not Available | Not Available |
| 2312 | Uncharacterized fimbrial-like protein YadN | *Escherichia coli* | Strain K12 | P37050 | -- | 8202364; 9278503; 16738553; 20345943 | Not Available | Not Available |
| 2313 | Uncharacterized fimbrial-like protein YgiL | *Escherichia coli* | Strain K12 | P39834 | -- | 9278503; 16738553; 1917833; 7984428 | Not Available | Not Available |
| 2314 | Probable two-component-system connector protein YcgZ | *Escherichia coli* | Strain K12 | P75991 | -- | 9278503; 16738553; 14527658; 18174134; 19172264; 19240136 | Not Available | Not Available |
| 2315 | Uncharacterized fimbrial-like protein YehA | *Escherichia coli* | Strain K12 | P33340 | -- | 9097040; 9278503; 16738553; 20345943 | Not Available | Not Available |
| 2316 | Protein Veg | *Bacillus subtilis* | Strain 168 | P37466 | -- | 7584024; 9384377; 6181373; 6790516; 12761295; 23378512 | Not Available | Not Available |
| 2317 | Uncharacterized fimbrial-like protein YadK | *Escherichia coli* | Strain K12 | P37016 | -- | 8202364; 9278503; 16738553; 20345943 | Not Available | Not Available |
| 2318 | Probable two-component-system connector protein YmgA | *Escherichia coli* | Strain K12 | P75992 | -- | 9278503; 16738553; 18174134; 19172264; 19240136 | Not Available | Not Available |
| 2319 | Sensor protein VraS (EC 2.7.13.3) | *Staphylococcus aureus* | Strain COL | Q5HEN9 | -- | 15774886 | ATP binding, phosphorelay sensor kinase activity, protein dimerization activity | Not Available |
| 2320 | Sensor protein VraS (EC 2.7.13.3) | *Staphylococcus aureus* | Strain Mu3 / ATCC 700698 | Q9KWK8 | -- | 10708580; 17954695 | ATP binding, phosphorelay sensor kinase activity, protein dimerization activity | Not Available |
| 2321 | Putative iNot Availablective recombiNot Availabletion-promoting nuclease-like protein YjiP | *Escherichia coli* | Strain K12 | P0DP21 | -- | 7610040; 9278503; 16738553; 16385049; 26162088; 26618540; 28096446 | Not Available | Not Available |
| 2322 | Uncharacterized protein YcgV | *Escherichia coli* | Strain K12 | P76017 | -- | 9278503; 16738553; 8905232 | Not Available | Not Available |
| 2323 | Sensor protein VraS (EC 2.7.13.3) | *Staphylococcus aureus* | Strain MSSA476 | Q6G849 | -- | 15213324 | ATP binding, phosphorelay sensor kinase activity, protein dimerization activity | Not Available |
| 2324 | Uncharacterized fimbrial-like protein YfcP | *Escherichia coli* | Strain K12 | P76499 | -- | 9278503; 16738553; 20345943 | Not Available | Not Available |
| 2325 | Uncharacterized fimbrial-like protein YfcR | *Escherichia coli* | Strain K12 | P76501 | -- | 9278503; 16738553; 20345943 | Not Available | Not Available |
| 2326 | Uncharacterized fimbrial-like protein YcbU | *Escherichia coli* | Strain K12 | P75859 | -- | 8905232; 9278503; 16738553; 20345943 | Not Available | Not Available |
| 2327 | Uncharacterized fimbrial-like protein YfcV | *Escherichia coli* | Strain K12 | P77288 | -- | 9205837; 9278503; 16738553; 20345943 | Not Available | Not Available |

| Target ID | Bacterial Protein Name | Organism Name | Strain | UniProt ID | PDB ID | PubMed ID | Classification as per UniProt GO Molecular Function | Classification of Function as per PDB |
|---|---|---|---|---|---|---|---|---|
| 2328 | Uncharacterized fimbrial-like protein YdeR | *Escherichia coli* | Strain K12 | P77294 | -- | 9097039; 9278503; 16738553 | Not Available | Not Available |
| 2329 | Uncharacterized fimbrial-like protein YraK | *Escherichia coli* | Strain K12 | P43319 | -- | 9278503; 16738553; 20345943 | Not Available | Not Available |
| 2330 | Uncharacterized protein YtrP | *Bacillus subtilis* | Strain 168 | O34325 | -- | 1699930; 9387221; 9384377 | Diguanylate cyclase activity | Not Available |
| 2331 | AI-2 transport protein TqsA (Transport of quorum-sensing signalprotein) | *Shigella flexneri* | | P0AFS6 | -- | 12384590; 12704152 | Not Available | Not Available |
| 2332 | Uncharacterized protein YeeJ | *Escherichia coli* | Strain K12 | P76347 | -- | 9097040; 9278503; 16397293; 16738553 | Not Available | Not Available |
| 2333 | Sensor protein VraS (EC 2.7.13.3) | *Staphylococcus aureus* | Strain N315 | Q99SZ7 | -- | 11418146; 12864861; 21173175; 25899124 | ATP binding, phosphorelay sensor kinase activity, protein dimerization activity | Not Available |
| 2334 | Uncharacterized fimbrial-like protein YadC | *Escherichia coli* | Strain K12 | P31058 | -- | 8202364; 9278503; 16738553; 8096212; 20345943 | Not Available | Not Available |
| 2335 | Uncharacterized fimbrial-like protein YdeS | *Escherichia coli* | Strain K12 | P77789 | -- | 9097039; 9278503; 16738553 | Not Available | Not Available |
| 2336 | Protein LpfE | *Salmonella typhimurium* | Strain LT2 / SGSC1412 / ATCC 700720 | P43664 | -- | 7721701; 11677609 | Not Available | Not Available |
| 2337 | Transcriptional regulatory protein QseB | *Escherichia coli* | Strain K12 | P52076 | -- | 9278503; 16738553; 11929534; 16352847 | DNA-binding Transcription activator activity, phosphorelay response regulator activity, Transcription cis-regulatory region binding | Not Available |
| 2338 | Extracellular matrix regulatory protein A (Regulator of extracellular matrix A) | *Bacillus subtilis* | Strain 168 | Q7WY72 | -- | 9384377; 19363116; 23396918; 23646920 | DNA binding | Not Available |
| 2339 | Extracellular matrix regulatory protein B (Regulator of the extracellular matrix B) | *Bacillus subtilis* | Strain 168 | P37525 | -- | 2987847; 7584024; 9384377; 19383706; 19363116 | Not Available | Not Available |
| 2340 | Cyclic di-GMP-binding protein (Cellulose synthase regulatory subunit) | *Salmonella typhimurium* | Strain LT2 / SGSC1412 / ATCC 700720 | Q93IN1 | -- | 11260463; 11929533; 11677609 | Not Available | Not Available |
| 2341 | Quorum-quenching protein AidA (Quorum-quenching enzyme) (QQ enzyme) | *Acinetobacter baumannii* | Strain MDR-ZJ06 | G2JHL6 | -- | 21788470; 28328989 | Hydrolase activity | Not Available |
| 2342 | Toxin CdiA (CdiA-EC93) | *Escherichia coli* | | Q3YL96 | -- | 16109881; 19124575; 21085179; 23882017; 25174572; 28351921; 30388452 | toxin activity | Not Available |
| 2343 | Fosfomycin resistance protein AbaF (Acinetobacter baumannii fosfomycin efflux) | *Acinetobacter baumannii* | Strain ATCC 17978 / CIP 53.77 / LMG 1025 / NCDC KC755 / 5377 | P0DPR5 | -- | 17344419; 27650185 | Transmembrane transporter activity | Not Available |
| 2344 | Transcriptional regulatory protein ArsR | *Helicobacter pylori* | Strain ATCC 700392 / 26695 Campylobacter pylori | O24973 | -- | 9252185; 11751811; 16672598; 22865848; 25997502; 27432830 | DNA-binding Transcription activator activity, DNA-binding Transcription repressor activity, phosphorelay response regulator activity, Transcription cis-regulatory region binding | Not Available |
| 2345 | Uncharacterized Membrane protein EpsK | *Bacillus subtilis* | Strain 168 | P71060 | -- | 8969506; 9384377; 19383706; 15661000 | Not Available | Not Available |
| 2346 | Poly-beta-1, 6-N-acetyl-D-glucosamine synthase (PNot AvailableG synthase) (Poly-beta-1, 6-GlcNAc synthase) (EC 2.4.1.-) (Biofilm polysaccharide intercellular adhesin synthesis protein IcaA) (Biofilm PIA synthesis protein IcaA) (Intercellular adhesion protein A) (N-acetylglucosaminylTransferase IcaA) | *Staphylococcus aureus* | Strain NCTC 8325 / PS 47 | Q9RQP9 | -- | 10496925 | Acetyl glucosaminyl transferase activity | Not Available |
| 2347 | Poly-beta-1, 6-N-acetyl-D-glucosamine synthase (PNot AvailableG synthase) (Poly-beta-1, 6-GlcNAc synthase) (EC 2.4.1.-) (Biofilm polysaccharide intercellular adhesin synthesis protein IcaA) (Biofilm PIA synthesis protein IcaA) (Intercellular adhesion protein A) (N-acetylglucosaminylTransferase IcaA) | *Staphylococcus aureus* | Strain COL | Q5HCN1 | -- | 15774886 | Acetyl glucosaminyl transferase activity | Not Available |
| 2348 | Poly-beta-1, 6-N-acetyl-D-glucosamine synthase (PNot AvailableG synthase) (Poly-beta-1, 6-GlcNAc synthase) (EC 2.4.1.-) (Biofilm polysaccharide intercellular adhesin synthesis protein IcaA) (Biofilm PIA synthesis protein IcaA) (Intercellular adhesion protein A) (N-acetylglucosaminylTransferase IcaA) | *Staphylococcus aureus* | Strain Mu50 / ATCC 700699 | Q99QX3 | -- | 11418146 | Acetyl glucosaminyl transferase activity | Not Available |
| 2349 | Poly-beta-1, 6-N-acetyl-D-glucosamine synthase (PNot AvailableG synthase) (Poly-beta-1, 6-GlcNAc synthase) (EC 2.4.1.-) (Biofilm polysaccharide intercellular adhesin synthesis protein IcaA) (Biofilm PIA synthesis protein IcaA) (Intercellular adhesion protein A) (N-acetylglucosaminylTransferase IcaA) | *Staphylococcus aureus* | Strain N315 | Q7A351 | -- | 11418146 | Acetyl glucosaminyl transferase activity | Not Available |
| 2350 | Poly-beta-1, 6-N-acetyl-D-glucosamine synthase (PNot AvailableG synthase) (Poly-beta-1, 6-GlcNAc synthase) (EC 2.4.1.-) (Biofilm polysaccharide intercellular adhesin synthesis protein IcaA) (Biofilm PIA synthesis protein IcaA) (Intercellular adhesion protein A) (N-acetylglucosaminylTransferase IcaA) | *Staphylococcus aureus* | Strain MRSA252 | Q6GDD8 | -- | 15213324 | Acetyl glucosaminyl transferase activity | Not Available |

| Target ID | Bacterial Protein Name | Organism Name | Strain | UniProt ID | PDB ID | PubMed ID | Classification as per UniProt GO Molecular Function | Classification of Function as per PDB |
|---|---|---|---|---|---|---|---|---|
| 2351 | Poly-beta-1, 6-N-acetyl-D-glucosamine synthase (PNot AvailableG synthase) (Poly-beta-1, 6-GlcNAc synthase) (EC 2.4.1.-) (Biofilm polysaccharide intercellular adhesin synthesis protein IcaA) (Biofilm PIA synthesis protein IcaA) (Intercellular adhesion protein A) (N-acetylglucosaminylTransferase IcaA) | *Staphylococcus aureus* | Strain MSSA476 | Q6G608 | -- | 15213324 | Acetyl glucosaminyl transferase activity | Not Available |
| 2352 | Poly-beta-1, 6-N-acetyl-D-glucosamine synthase (PNot AvailableG synthase) (Poly-beta-1, 6-GlcNAc synthase) (EC 2.4.1.-) (Biofilm polysaccharide intercellular adhesin synthesis protein IcaA) (Biofilm PIA synthesis protein IcaA) (Intercellular adhesion protein A) (N-acetylglucosaminylTransferase IcaA) | *Staphylococcus aureus* | Strain MW2 | Q8NUI7 | -- | 12044378 | Acetyl glucosaminyl transferase activity | Not Available |
| 2353 | Poly-beta-1, 6-N-acetyl-D-glucosamine synthase (PNot AvailableG synthase) (Poly-beta-1, 6-GlcNAc synthase) (EC 2.4.1.-) (Biofilm polysaccharide intercellular adhesin synthesis protein IcaA) (Biofilm PIA synthesis protein IcaA) (Intercellular adhesion protein A) (N-acetylglucosaminylTransferase IcaA) | *Staphylococcus epidermidis* | | Q8GLC5 | -- | 15096544 | Acetyl glucosaminyl transferase activity | Not Available |
| 2354 | Poly-beta-1, 6-N-acetyl-D-glucosamine synthase (PNot AvailableG synthase) (Poly-beta-1, 6-GlcNAc synthase) (EC 2.4.1.-) (Biofilm polysaccharide intercellular adhesin synthesis protein IcaA) (Biofilm PIA synthesis protein IcaA) (Intercellular adhesion protein A) (N-acetylglucosaminylTransferase IcaA) | *Staphylococcus epidermidis* | Strain ATCC 35984 / RP62A | Q5HKQ0 | -- | 8809760; 15774886; 9660830 | Acetyl glucosaminyl transferase activity | Not Available |
| 2355 | Poly-beta-1, 6-N-acetyl-D-glucosamine N-deacetylase (PNot AvailableG N-deacetylase) (Poly-beta-1, 6-GlcNAc N-deacetylase) (EC 3.5.1.-) (Biofilm polysaccharide intercellular adhesin deacetylase) (Biofilm PIA deacetylase) (Intercellular adhesion protein B) | *Staphylococcus aureus* | Strain NCTC 8325 / PS 47 | Q9RQP7 | -- | 10496925 | Hydrolase activity, acting on carbon-nitrogen (but not peptide) bonds, in linear amides | Not Available |
| 2356 | Poly-beta-1, 6-N-acetyl-D-glucosamine N-deacetylase (PNot AvailableG N-deacetylase) (Poly-beta-1, 6-GlcNAc N-deacetylase) (EC 3.5.1.-) (Biofilm polysaccharide intercellular adhesin deacetylase) (Biofilm PIA deacetylase) (Intercellular adhesion protein B) | *Staphylococcus epidermidis* | | Q6TYB1 | -- | 15096544; 15501828 | Hydrolase activity, acting on carbon-nitrogen (but not peptide) bonds, in linear amides | Not Available |
| 2357 | Poly-beta-1, 6-N-acetyl-D-glucosamine N-deacetylase (PNot AvailableG N-deacetylase) (Poly-beta-1, 6-GlcNAc N-deacetylase) (EC 3.5.1.-) (Biofilm polysaccharide intercellular adhesin deacetylase) (Biofilm PIA deacetylase) (Intercellular adhesion protein B) | *Staphylococcus epidermidis* | Strain ATCC 35984 / RP62A | Q5HKP8 | -- | 8809760; 15774886 | Hydrolase activity, acting on carbon-nitrogen (but not peptide) bonds, in linear amides | Not Available |
| 2358 | Probable poly-beta-1, 6-N-acetyl-D-glucosamine export protein (PGA export protein) (Poly-beta-1, 6-GlcNAc export protein) (Biofilm polysaccharide intercellular adhesin export protein) (Biofilm PIA export protein) (Intercellular adhesion protein C) | *Staphylococcus aureus* | Strain NCTC 8325 / PS 47 | Q9RQP6 | -- | 10496925 | Acyltransferase activity, transferring groups other than amino-acyl groups | Not Available |
| 2359 | Probable poly-beta-1, 6-N-acetyl-D-glucosamine export protein (PGA export protein) (Poly-beta-1, 6-GlcNAc export protein) (Biofilm polysaccharide intercellular adhesin export protein) (Biofilm PIA export protein) (Intercellular adhesion protein C) | *Staphylococcus epidermidis* | | P69518 | -- | 15096544; 15501828 | Acyltransferase activity, transferring groups other than amino-acyl groups | Not Available |
| 2360 | Probable poly-beta-1, 6-N-acetyl-D-glucosamine export protein (PGA export protein) (Poly-beta-1, 6-GlcNAc export protein) (Biofilm polysaccharide intercellular adhesin export protein) (Biofilm PIA export protein) (Intercellular adhesion protein C) | *Staphylococcus epidermidis* | Strain ATCC 35984 / RP62A | Q5HKP7 | -- | 8809760; 15774886; 9660830 | Acyltransferase activity, transferring groups other than amino-acyl groups | Not Available |
| 2361 | Poly-beta-1, 6-N-acetyl-D-glucosamine synthesis protein IcaD (PGA synthesis protein IcaD) (Poly-beta-1, 6-GlcNAc synthesis protein IcaD) (Biofilm polysaccharide intercellular adhesin synthesis protein IcaD) (Biofilm PIA synthesis protein IcaD) (Intercellular adhesion protein D) | *Staphylococcus aureus* | Strain NCTC 8325 / PS 47 | Q9RQP8 | -- | 10496925 | Not Available | Not Available |
| 2362 | Biofilm operon icaADBC HTH-type negative Transcriptional regulator IcaR (Intercellular adhesion protein R) | *Staphylococcus aureus* | Strain NCTC 8325 / PS 47 | Q9RQQ0 | -- | 10496925 | DNA-binding Transcription factor activity, Transcription cis-regulatory region binding | Not Available |
| 2363 | signalpeptidase I W (SPase I) (EC 3.4.21.89) (Leader peptidase I) | *Bacillus subtilis* | Strain 168 | P54506 | -- | 8969508; 9384377; 9694797; 9823656; 10049401; 10464223; 10559173; 16430695; 23646920 | Serine-type endopeptidase activity | Not Available |
| 2364 | Endoglucanae (EC 3.2.1.4) (Carboxymethylcellulase) (CMCase) (Cellulase) (Endo-1, 4-beta-glucanae) | *Salmonella typhimurium* | Strain LT2 / SGSC1412 / ATCC 700720 | Q8ZLB7 | -- | 11260463; 11929533; 11677609 | Sellulase activity | Not Available |
| 2365 | Major fimbrium subunit FimA type-4 (Fimbrillin) (Fimbrilin) (Major fimbrial subunit protein type IV) | *Porphyromonas gingivalis* | | Q51827 | -- | 7902712; 7851739; 1987052; 11748193 | Structural protein | Not Available |
| 2366 | Uncharacterized sugar Transferase EpsL (EC 2.-.-.-) | *Bacillus subtilis* | Strain 168 | P71062 | -- | 8969506; 9384377; 15661000 | PhosphoTransferase activity, for other substituted phosphate groups | Not Available |
| 2367 | Biofilm architecture maintece protein MbaA | *Vibrio cholerae serotype O1* | Strain ATCC 39315 / El Tor Iba N16961 | Q9KU26 | -- | 10952301; 12562809; 16237027 | Not Available | Not Available |
| 2368 | Cellulose synthase catalytic subunit [UDP-forming] (EC 2.4.1.12) | *Salmonella typhimurium* | Strain LT2 / SGSC1412 / ATCC 700720 | Q93IN2 | -- | 11260463; 11929533; 11677609 | Cellulose synthase (UDP-forming) activity, cyclic-di-GMP binding, hexosylTransferase activity | Not Available |
| 2369 | CsgBAC operon Transcriptional regulatory protein | *Escherichia coli* | Strain K12 | P52106 | -- | 8817489; 8905232; 9278503; 16738553; 16513732; 17010156; 24212724 | DNA-binding Transcription activator activity, DNA-binding Transcription repressor activity, Identical protein binding, Transcription cis-regulatory region binding | Not Available |
| 2370 | Surface composition regulator | *Escherichia coli O139:H28* | Strain E24377A / ETEC | A7ZRT0 | -- | 18676672 | Not Available | Not Available |

| Target ID | Bacterial Protein Name | Organism Name | Strain | UniProt ID | PDB ID | PubMed ID | Classification as per UniProt GO Molecular Function | Classification of Function as per PDB |
|---|---|---|---|---|---|---|---|---|
| 2371 | Surface composition regulator | *Escherichia coli O127:H6* | Strain E2348 / 69 / EPEC | B7UIV7 | -- | 18952797 | Not Available | Not Available |
| 2372 | Surface composition regulator | *Escherichia coli O45:K1* | Strain S88 / ExPEC | B7MAC5 | -- | 19165319 | Not Available | Not Available |
| 2373 | Surface composition regulator | *Escherichia coli* | Strain 55989 / EAEC | B7LGY3 | -- | 19165319 | Not Available | Not Available |
| 2374 | Surface composition regulator | *Escherichia coli O157:H7* | | P64244 | -- | 11206551; 11258796 | Not Available | Not Available |
| 2375 | Surface composition regulator | *Escherichia coli O157:H7* | Strain EC4115 / EHEC | B5YR89 | -- | 22135463 | Not Available | Not Available |
| 2376 | Surface composition regulator | *Escherichia coli O7:K1* | Strain IAI39 / ExPEC | B7NJR2 | -- | 19165319 | Not Available | Not Available |
| 2377 | Surface composition regulator | *Escherichia coli O81* | Strain ED1a | B7N052 | -- | 19165319 | Not Available | Not Available |
| 2378 | Surface composition regulator | *Escherichia coli O8* | Strain IAI1 | B7LZJ8 | -- | 19165319 | Not Available | Not Available |
| 2379 | Surface composition regulator | *Escherichia coli* | Strain K12 / MC4100 / BW2952 | C4ZQW6 | -- | 19376874 | Not Available | Not Available |
| 2380 | Surface composition regulator | *Escherichia coli* | Strain K12 / DH10B | B1XG54 | -- | 18245285 | Not Available | Not Available |
| 2381 | Surface composition regulator | *Escherichia coli O9:H4* | Strain HS | A8A4K4 | -- | 18676672 | Not Available | Not Available |
| 2382 | Surface composition regulator | *Escherichia coli O6:K15:H31* | Strain 536 / UPEC | Q0TD58 | -- | 16879640 | Not Available | Not Available |
| 2383 | Surface composition regulator | *Escherichia coli O6:H1* | Strain CFT073 / ATCC 700928 / UPEC | P64243 | -- | 12471157 | Not Available | Not Available |
| 2384 | Surface composition regulator | *Escherichia coli O17:K52:H18* | Strain UMN026 / ExPEC | B7ND37 | -- | 19165319 | Not Available | Not Available |
| 2385 | Surface composition regulator | *Escherichia coli* | Strain SE11 | B6I419 | -- | 18931093 | Not Available | Not Available |
| 2386 | Surface composition regulator | *Escherichia coli* | Strain SMS-3-5 / SECEC | B1LF40 | -- | 18708504 | Not Available | Not Available |
| 2387 | Surface composition regulator | *Escherichia coli* | Strain UTI89 / UPEC | Q1R6T2 | -- | 16585510 | Not Available | Not Available |
| 2388 | Surface composition regulator | *Salmonella choleraesuis* | Strain SC-B67 | Q57JR3 | -- | 15781495 | Not Available | Not Available |
| 2389 | Surface composition regulator | *Salmonella enteritidis PT4* | Strain P125109 | B5QZ33 | -- | 18583645 | Not Available | Not Available |
| 2390 | Surface composition regulator | *Salmonella paratyphi A* | Strain ATCC 9150 / SARB42 | Q5PC89 | -- | 15531882 | Not Available | Not Available |
| 2391 | Surface composition regulator | *Salmonella paratyphi A* | Strain AKU_12601 | B5BG08 | -- | 19159446 | Not Available | Not Available |
| 2392 | Surface composition regulator | *Salmonella typhi* | | P58615 | -- | 11677608; 12644504 | Not Available | Not Available |
| 2393 | Surface composition regulator | *Salmonella typhimurium* | Strain LT2 / SGSC1412 / ATCC 700720 | P58614 | -- | 11677609 | Not Available | Not Available |
| 2394 | Surface composition regulator | *Shigella dysenteriae serotype 1* | Strain Sd197 | Q32BR9 | -- | 16275786 | Not Available | Not Available |
| 2395 | Surface composition regulator | *Shigella flexneri* | | Q7UBJ0 | -- | 12384590; 12704152 | Not Available | Not Available |
| 2396 | Surface composition regulator | *Shigella sonnei* | Strain SS046 | Q3YXJ3 | -- | 16275786 | Not Available | Not Available |
| 2397 | Cellulose synthase operon protein C | *Salmonella typhimurium* | Strain LT2 / SGSC1412 / ATCC 700720 | Q8ZLB8 | -- | 11260463; 11929533; 11677609 | Not Available | Not Available |
| 2398 | Major fimbrium tip subunit FimE | *Porphyromonas gingivalis* | | O32390 | -- | 8981345; 17526848 | Host cell extracellular matrix binding | Not Available |
| 2399 | Probable diguanylate cyclase DgcQ (DGC) (EC 2.7.7.65) (Cellulose synthesis regulatory protein) | *Escherichia coli* | Strain K12 | P76330 | -- | 9097040; 9278503; 16738553; 16585767; 17010156; 19332833; 20303158; 21181144; 26148715 | Diguanylate cyclase activity, GTP binding, Metal ion binding | Not Available |
| 2400 | Autotransporter adhesin EhaG (Type 5 secretion system autotransporter EhaG) | *Escherichia coli O157:H7* | | Q7DJ60 | -- | 11206551; 11258796; 27056233; 22286983 | Not Available | Not Available |
| 2401 | Uncharacterized glycosylTransferase EpsJ (EC 2.4.-.-) | *Bacillus subtilis* | Strain 168 | P71059 | -- | 8969506; 9384377; 15661000 | | Not Available |
| 2402 | Putative pyruvyl Transferase EpsI (EC 2.-.-.-) | *Bacillus subtilis* | Strain 168 | P71058 | -- | 8969506; 9384377; 15175311; 15661000 | Transferase activity | Not Available |
| 2403 | Hemin storage protein | *Lysobacter arseniciresistens ZS79* | | A0A0A0F1X8 | -- | 26516404 | Hydrolase activity, acting on carbon-nitrogen (but not peptide) bonds | Not Available |
| 2404 | Poly-beta-1, 6-N-acetyl-D-glucosamine synthase (Poly-beta-1, 6-GlcNAc synthase) (EC 2.4.1.-) | *Lysobacter arseniciresistens ZS79* | | A0A0A0EWS4 | -- | 26516404 | Acetyl glucosaminyl transferase activity | Not Available |
| 2405 | Type II toxin-antitoxin system MqsR family toxin | *Pseudomonas panacis* | | A0A7Y1IL75 | -- | 32587583 | Not Available | Not Available |
| 2406 | Poly-beta-1, 6-N-acetyl-D-glucosamine N-deacetylase PgaB | *Desulfobulbaceae bacterium* | | A0A7V5KUV9 | -- | 31911466 | Hydrolase activity, acting on carbon-nitrogen (but not peptide) bonds | Not Available |
| 2407 | Poly-beta-1, 6-N-acetyl-D-glucosamine synthase (Poly-beta-1, 6-GlcNAc synthase) (EC 2.4.1.-) | *Desulfobulbaceae bacterium* | | A0A7V5V5A8 | -- | 31911466 | Acetyl glucosaminyl transferase activity | Not Available |
| 2408 | Toxin mRNA interferase YgiU | *Pseudomonas furukawaii* | | L8MLK2 | -- | 22843571 | Not Available | Not Available |
| 2409 | Poly-beta-1, 6-N-acetyl-D-glucosamine synthase (Poly-beta-1, 6-GlcNAc synthase) (EC 2.4.1.-) | *Desulfobulbaceae bacterium* | | A0A7V5FPJ1 | -- | 31911466 | Acetyl glucosaminyl transferase activity | Not Available |
| 2410 | Poly-beta-1, 6-N-acetyl-D-glucosamine biosynthesis protein PgaD | *Desulfobulbaceae bacterium* | | A0A7V5FPJ3 | -- | 31911466 | Not Available | Not Available |
| 2411 | Poly-beta-1, 6-N-acetyl-D-glucosamine N-deacetylase PgaB | *Desulfobulbaceae bacterium* | | A0A7V5KV26 | -- | 31911466 | Hydrolase activity, acting on carbon-nitrogen (but not peptide) bonds | Not Available |
| 2412 | Poly-beta-1, 6-N-acetyl-D-glucosamine synthase (Poly-beta-1, 6-GlcNAc synthase) (EC 2.4.1.-) | *Pseudomonas panacis* | | A0A7Y1IRF0 | -- | 32587583 | Acetyl glucosaminyl transferase activity | Not Available |
| 2413 | Poly-beta-1, 6-N-acetyl-D-glucosamine synthase (Poly-beta-1, 6-GlcNAc synthase) (EC 2.4.1.-) | *Aeromona salmonicida* | Strain A449 | A4SIQ6 | -- | 18801193 | Acetyl glucosaminyl transferase activity | Not Available |
| 2414 | Surface composition regulator | *Cronobacter sakazakii 696* | | K8CNU6 | -- | 23166675 | Not Available | Not Available |
| 2415 | Poly-beta-1, 6-N-acetyl-D-glucosamine biosynthesis protein PgaD | *Pseudomonas panacis* | | A0A7Y1IRD1 | -- | 32587583 | Not Available | Not Available |
| 2416 | Poly-beta-1, 6-N-acetyl-D-glucosamine biosynthesis protein PgaD | *Desulfobulbaceae bacterium* | | A0A7V5V4Z0 | -- | 31911466 | Not Available | Not Available |
| 2417 | Poly-beta-1, 6-N-acetyl-D-glucosamine biosynthesis protein PgaD (Fragment) | *Enterobacter spp.* | | A0A3D2B4K2 | -- | 30148503 | Not Available | Not Available |
| 2418 | Poly-beta-1, 6-N-acetyl-D-glucosamine synthase (Poly-beta-1, 6-GlcNAc synthase) (EC 2.4.1.-) | *Acinetobacter wuhouensis* | | A0A385C658 | -- | 30117799 | Acetyl glucosaminyl transferase activity | Not Available |
| 2419 | Uncharacterized protein | *Photorhabdus temperata subspp. temperata M1021* | | T0PA00 | -- | 24029767 | Not Available | Not Available |
| 2420 | Biofilm PGA synthesis deacetylase PgaB | *Yersinia enterocolitica subspp. palearctica serotype O:3* | Strain DSM 13030 / CIP 106945 / Y11 | A0A0H3NVK7 | -- | 21296963 | Hydrolase activity, acting on carbon-nitrogen (but not peptide) bonds | Not Available |
| 2421 | Biofilm PGA synthesis auxiliary protein PgaD | *Yersinia enterocolitica subspp. palearctica serotype O:3* | Strain DSM 13030 / CIP 106945 / Y11 | A0A0H3NSA5 | -- | 21296963 | Not Available | Not Available |
| 2422 | Uncharacterized protein | *Rhodocyclales bacterium GWA2_65_19* | | A0A1G3H3U9 | -- | 27774985 | Not Available | Not Available |

| Target ID | Bacterial Protein Name | Organism Name | Strain | UniProt ID | PDB ID | PubMed ID | Classification as per UniProt GO Molecular Function | Classification of Function as per PDB |
|---|---|---|---|---|---|---|---|---|
| 2423 | Uncharacterized protein | *Syntrophomona wolfei subspp. wolfei* | Strain DSM 2245B / Goettingen | Q0AU88 | -- | 21966920 | Not Available | Not Available |
| 2424 | Toxin mRNA interferase YgiU | *Yersinia enterocolitica subspp. palearctica serotype O:3* | Strain DSM 13030 / CIP 106945 / Y11 | A0A0H3NTG7 | -- | 21296963 | Not Available | Not Available |
| 2425 | DUF1471 domain-containing protein (Multiple stress resistance protein BhsA) (Outer Membrane protein) (Putative outer Membrane protein) | *Salmonella typhimurium* | | A0A0F7J832 | -- | 26089426; 30286803 | Not Available | Not Available |
| 2426 | Surface composition regulator | *Serratia marcescens* | | A0A0F6KTF3 | -- | 28536292 | Not Available | Not Available |
| 2427 | Surface composition regulator | *Serratia marcescens* | | A0A0A5LSU7 | -- | 28536292 | Not Available | Not Available |
| 2428 | Poly-beta-1, 6-N-acetyl-D-glucosamine biosynthesis protein PgaD | *Serratia marcescens* | | A0A2V4G8U1 | -- | 21967170; 30286803 | Not Available | Not Available |
| 2429 | Poly-beta-1, 6-N-acetyl-D-glucosamine synthase (Poly-beta-1, 6-GlcNAc synthase) (EC 2.4.1.-) | *Pseudomonas yamanorum* | | A0A145XGR5 | -- | 27231373 | Acetyl glucosaminyl transferase activity | Not Available |
| 2430 | Surface composition regulator | *Salmonella enterica subspp. enterica serovar* | Poona | A0A4Z0KU92 | -- | 30286803 | Not Available | Not Available |
| 2431 | Poly-beta-1, 6-N-acetyl-D-glucosamine synthase (Poly-beta-1, 6-GlcNAc synthase) (EC 2.4.1.-) | *Acinetobacter venetianus* | | A0A150HR71 | -- | 26902269 | Acetyl glucosaminyl transferase activity | Not Available |
| 2432 | Biofilm PGA synthesis auxiliary protein PgaD (Biofilm PGA synthesis protein PgaD) (Poly-beta-1, 6-N-acetyl-D-glucosamine biosynthesis protein PgaD) | *Pseudomonas yamanorum* | | A0A145XGQ4 | -- | 27231373 | Not Available | Not Available |
| 2433 | Toxin CdiA | *Neisseria meningitidis* | | Q9JPE3 | -- | 10722605; 19289515; 28505234; 30485280; 10722605 | Toxin activity | Not Available |
| 2434 | Surface composition regulator | *Salmonella typhimurium* | | A0A0J5E264 | -- | 30286803 | Not Available | Not Available |
| 2435 | Surface composition regulator | *Serratia marcescens* | | A0A7D6C0Q8 | -- | 21967170; 30286803 | Not Available | Not Available |
| 2436 | Poly-beta-1, 6-N-acetyl-D-glucosamine synthase (Poly-beta-1, 6-GlcNAc synthase) (EC 2.4.1.-) | *Serratia marcescens* | | A0A7D6BXB5 | -- | 21967170; 30286803 | Acetyl glucosaminyl transferase activity | Not Available |
| 2437 | Poly-beta-1, 6-N-acetyl-D-glucosamine synthase (Poly-beta-1, 6-GlcNAc synthase) (EC 2.4.1.-) | *Serratia marcescens* | | A0A0A5LT03 | -- | 28536292; 28636609 | Acetyl glucosaminyl transferase activity | Not Available |
| 2438 | Poly-beta-1, 6-N-acetyl-D-glucosamine biosynthesis protein PgaD | *Acinetobacter seifertii* | | N8QXR6 | -- | 25563912 | Not Available | Not Available |
| 2439 | Poly-beta-1, 6-N-acetyl-D-glucosamine synthase (Poly-beta-1, 6-GlcNAc synthase) (EC 2.4.1.-) | *Acinetobacter seifertii* | | N8QY22 | -- | 25563912 | Acetyl glucosaminyl transferase activity | Not Available |
| 2440 | Poly-beta-1, 6-N-acetyl-D-glucosamine N-deacetylase (Poly-beta-1, 6-N-acetyl-D-glucosamine N-deacetylase PgaB) | *Serratia marcescens* | | A0A0G3SWE7 | -- | 28636609 | Hydrolase activity, acting on carbon-nitrogen (but not peptide) bonds | Not Available |
| 2441 | Guanosine-3', 5'-bis(diphosphate) 3'-diphosphatase (EC 3.1.7.2) | *Salmonella enteritidis* | | K0A192 | -- | 30286803 | Guanosine-3', 5'-bis(diphosphate) 3'-diphosphatase activity, kinase activity | Not Available |
| 2442 | Poly-beta-1, 6-N-acetyl-D-glucosamine synthase (Poly-beta-1, 6-GlcNAc synthase) (EC 2.4.1.-) | *Acinetobacter seifertii* | | N8S570 | -- | 25563912 | Acetyl glucosaminyl transferase activity | Not Available |
| 2443 | Surface composition regulator | *Salmonella enterica subspp. enterica serovar* | Heidelberg | A0A265B3W8 | -- | 30286803 | Not Available | Not Available |
| 2444 | Type II toxin-antitoxin system MqsR family toxin | *Pseudomonas putida* | Arthrobacter siderocapsulatus | A0A550F6I7 | -- | 19827713 | Not Available | Not Available |
| 2445 | Surface composition regulator | *Salmonella enterica subspp. enterica serovar* | Typhimurium var. 5- | A0A5W1BAQ5 | -- | 30286803 | Not Available | Not Available |
| 2446 | Surface composition regulator | *Salmonella enterica subspp. arizonae* | | A0A3Z4JNS2 | -- | 30286803 | Not Available | Not Available |
| 2447 | Surface composition regulator | *Salmonella paratyphi A* | | A0A3U5EDJ2 | -- | 22190395; 30286803 | Not Available | Not Available |
| 2448 | Flagellin | *Acidovorax citrulli* | Acidovorax avenae subspp. citrulli | A5HIK5 | -- | 20807236; 21554180 | Structural protein | Not Available |
| 2449 | Poly-beta-1, 6-N-acetyl-D-glucosamine biosynthesis protein PgaD | *Deltaproteobacteria bacterium HGW-Deltaproteobacteria-4* | | A0A2N2FUX1 | -- | 28350393 | Not Available | Not Available |
| 2450 | Toxin mRNA interferase YgiU | *Yersinia enterocolitica subspp. palearctica serotype O:3* | Strain YE-P4 | A0A7H4ZBT2 | -- | 23846271 | Not Available | Not Available |
| 2451 | Poly-beta-1, 6-N-acetyl-D-glucosamine synthase (Poly-beta-1, 6-GlcNAc synthase) (EC 2.4.1.-) | *Yersinia enterocolitica subspp. palearctica serotype O:3* | Strain YE-P4 | A0A7H4ZKR9 | -- | 23846271 | Acetyl glucosaminyl transferase activity | Not Available |
| 2452 | Type II toxin-antitoxin system MqsR family toxin | *Pseudomonas spp. WS 5532* | | A0A7Y0Z345 | -- | 32587583 | Not Available | Not Available |
| 2453 | Poly-beta-1, 6-N-acetyl-D-glucosamine biosynthesis protein PgaD | *Pseudomonas spp. WS 5532* | | A0A7Y0Z6M5 | -- | 32587583 | Not Available | Not Available |
| 2454 | Poly-beta-1, 6-N-acetyl-D-glucosamine synthase (Poly-beta-1, 6-GlcNAc synthase) (EC 2.4.1.-) | *Acinetobacter spp. ACNIH4* | | A0A2S4TAE5 | -- | 29437920 | Acetyl glucosaminyl transferase activity | Not Available |
| 2455 | Surface composition regulator | *Yersinia enterocolitica subspp. palearctica serotype O:3* | Strain YE-P4 | A0A7H4ZIG3 | -- | 23846271 | DNA binding, DNA-directed DNA polymerase activity | Not Available |
| 2456 | Poly-beta-1, 6-N-acetyl-D-glucosamine synthase (Poly-beta-1, 6-GlcNAc synthase) (EC 2.4.1.-) | *Pseudomonas spp. WS 5532* | | A0A7Y0Z6L1 | -- | 32587583 | Acetyl glucosaminyl transferase activity | Not Available |
| 2457 | Poly-beta-1, 6-N-acetyl-D-glucosamine N-deacetylase PgaB | *Deltaproteobacteria bacterium HGW-Deltaproteobacteria-4* | | A0A2N2FUZ1 | -- | 28350393 | Hydrolase activity, acting on carbon-nitrogen (but not peptide) bonds | Not Available |
| 2458 | Surface composition regulator | *Cronobacter dublinensis 1210* | | K8ANJ5 | -- | 23166675 | Not Available | Not Available |
| 2459 | Poly-beta-1, 6-N-acetyl-D-glucosamine synthase (Poly-beta-1, 6-GlcNAc synthase) (EC 2.4.1.-) | *Deltaproteobacteria bacterium HGW-Deltaproteobacteria-4* | | A0A2N2FUX4 | -- | 28350393 | Acetyl glucosaminyl transferase activity | Not Available |
| 2460 | Poly-beta-1, 6-N-acetyl-D-glucosamine N-deacetylase PgaB (EC 3.5.1.-) | *Pseudomonas spp. WS 5532* | | A0A7Y0Z6L3 | -- | 32587583 | Hydrolase activity, acting on carbon-nitrogen (but not peptide) bonds | Not Available |
| 2461 | Hemin storage system protein | *Yersinia enterocolitica subspp. palearctica serotype O:3* | Strain YE-P4 | A0A7H4ZKV9 | -- | 23846271 | Not Available | Not Available |
| 2462 | mRNA interferase MqsR | *Photorhabdus thracensis* | | A0A0F7LS53 | -- | 26415660 | Not Available | Not Available |
| 2463 | Type II toxin-antitoxin system MqsR family toxin | *Rhizobium spp.* | | A0A3D3BHM1 | -- | 30148503 | Not Available | Not Available |
| 2464 | Secreted biofilm formation protein YqxM | *Bacillus licheniformis* | Strain ATCC 14580 / DSM 13 / JCM 2505 / CCUG 7422 / NBRC 12200 / NCIMB 9375 / NCTC 10341 / NRRL NRS-1264 / Gibson 46 | Q65HF1 | -- | 15461803 | Not Available | Not Available |
| 2465 | Surface composition regulator | *Enterobacteriaceae bacterium ENNIH1* | | A0A2I8Q9C8 | -- | 29437920 | Not Available | Not Available |
| 2466 | Poly-beta-1, 6-N-acetyl-D-glucosamine biosynthesis protein PgaD | *Acinetobacter spp. ACNIH4* | | A0A2S4TAG1 | -- | 29437920 | Not Available | Not Available |

| Target ID | Bacterial Protein Name | Organism Name | Strain | UniProt ID | PDB ID | PubMed ID | Classification as per UniProt GO Molecular Function | Classification of Function as per PDB |
|---|---|---|---|---|---|---|---|---|
| 2467 | mRNA interferase MqsR | *Photorhabdus khanii subspp. guanajuatensis* | | A0A4R4JWB8 | -- | 30688647 | Not Available | Not Available |
| 2468 | Poly-beta-1, 6-N-acetyl-D-glucosamine biosynthesis protein PgaD | *Stenotrophomonas spp. ESTM1D_MKCIP4_1* | | A0A3Q8FUZ0 | -- | 29765358 | Not Available | Not Available |
| 2469 | Poly-beta-1, 6-N-acetyl-D-glucosamine synthase (Poly-beta-1, 6-GlcNAc synthase) (EC 2.4.1.-) | *Stenotrophomonas spp. ESTM1D_MKCIP4_1* | | A0A3S7KXH2 | -- | 29765358 | Acetyl glucosaminyl transferase activity | Not Available |
| 2470 | Type II toxin-antitoxin system MqsR family toxin | *Rhizobium radiobacter* | Agrobacterium tumefaciens Agrobacterium radiobacter | A0A546XJ46 | -- | 31273398 | Not Available | Not Available |
| 2471 | Uncharacterized protein | *Alicyclobacillus acidocaldarius subspp. acidocaldarius* | Strain ATCC 27009 / DSM 446 / BCRC 14685 / JCM 5260 / KCTC 1825 / NBRC 15652 / NCIMB 11725 / NRRL B-14509 / 104-IA Bacillus acidocaldarius | C8WT26 | -- | 21304673 | Not Available | Not Available |
| 2472 | Biofilm PGA synthesis protein PgaD | *Escherichia coli O157:H7* | Strain EC869 | A0A0H3PN65 | -- | 21421787 | Not Available | Not Available |
| 2473 | Poly-beta-1, 6-N-acetyl-D-glucosamine biosynthesis protein PgaD | *Candidatus Manganitrophus noduliformans* | | A0A7X6DPU5 | -- | 32669693 | Not Available | Not Available |
| 2474 | Poly-beta-1, 6-N-acetyl-D-glucosamine biosynthesis protein PgaD | *Stenotrophomonas spp. ESTM1D_MKCIP4_1* | | A0A3S5GHZ3 | -- | 29765358 | Not Available | Not Available |
| 2475 | Poly-beta-1, 6-N-acetyl-D-glucosamine N-deacetylase PgaB | *Acinetobacter spp. ACNIH2* | | A0A2L0HWN8 | -- | 29437920 | Hydrolase activity, acting on carbon-nitrogen (but not peptide) bonds | Not Available |
| 2476 | Uncharacterized protein | *Desulfovibrio magneticus* | Strain ATCC 700980 / DSM 13731 / RS-1 | C4XJ49 | -- | 19675025 | Not Available | Not Available |
| 2477 | Poly-beta-1, 6-N-acetyl-D-glucosamine synthase (Poly-beta-1, 6-GlcNAc synthase) (EC 2.4.1.-) | *Aeromona sobria* | | A0A2N3IZW3 | -- | 29276504 | Acetyl glucosaminyl transferase activity | Not Available |
| 2478 | Poly-beta-1, 6-N-acetyl-D-glucosamine biosynthesis protein PgaD (Fragment) | *Oceanospirillales bacterium* | | A0A7V2SRU5 | -- | 31911466 | Not Available | Not Available |
| 2479 | Chitin deacetylase | *Aeromona sobria* | | A0A2N3IZH4 | -- | 29276504 | Hydrolase activity, acting on carbon-nitrogen (but not peptide) bonds | Not Available |
| 2480 | Uncharacterized protein | *Candidatus Eisenbacteria bacterium* | | A0A538U948 | -- | 31110364 | Not Available | Not Available |
| 2481 | Poly-beta-1, 6-N-acetyl-D-glucosamine synthase (Poly-beta-1, 6-GlcNAc synthase) (EC 2.4.1.-) | *Stenotrophomonas spp. ESTM1D_MKCIP4_1* | | A0A3S5GHV1 | -- | 29765358 | Acetyl glucosaminyl transferase activity | Not Available |
| 2482 | Surface composition regulator | *Salmonella typhimurium* | Strain 4 / 74 | E8XBC6 | -- | 21478351 | Not Available | Not Available |
| 2483 | Surface composition regulator | *Escherichia coli O157:H7* | Strain EC869 | A0A0H3PT67 | -- | 21421787 | Not Available | Not Available |
| 2484 | Poly-beta-1, 6-N-acetyl-D-glucosamine biosynthesis protein PgaD | *Methylococcaceae bacterium* | | A0A351V596 | -- | 30148503 | Not Available | Not Available |
| 2485 | Biofilm PGA synthesis lipoprotein PgaB | *Pectobacterium parmentieri* | | A0A0H3ICP1 | -- | 23045508 | Hydrolase activity, acting on carbon-nitrogen (but not peptide) bonds | Not Available |
| 2486 | Hemin storage system, HmsS protein | *Pectobacterium parmentieri* | | A0A0H3IBQ4 | -- | 23045508 | Not Available | Not Available |
| 2487 | Poly-beta-1, 6-N-acetyl-D-glucosamine N-deacetylase PgaB | *Pseudomonas spp.* | | A0A3D4NT91 | -- | 30148503 | Hydrolase activity, acting on carbon-nitrogen (but not peptide) bonds | Not Available |
| 2488 | Poly-beta-1, 6-N-acetyl-D-glucosamine biosynthesis protein PgaD | *Xenorhabdus budapestensis* | | A0A2D0J1F6 | -- | 28993611 | Not Available | Not Available |
| 2489 | Polysaccharide deacetylase | *Xenorhabdus budapestensis* | | A0A2D0J1D9 | -- | 28993611 | Hydrolase activity, acting on carbon-nitrogen (but not peptide) bonds | Not Available |
| 2490 | Poly-beta-1, 6-N-acetyl-D-glucosamine synthase (Poly-beta-1, 6-GlcNAc synthase) (EC 2.4.1.-) | *Methylococcaceae bacterium* | | A0A351KVS7 | -- | 30148503 | Acetyl glucosaminyl transferase activity | Not Available |
| 2491 | Surface composition regulator | *Salmonella enteritidis* | | A0A0M0QID5 | -- | 30286803 | Not Available | Not Available |
| 2492 | Surface composition regulator | *Salmonella enterica subspp. enterica serovar* | Agona | A0A2T9HZU6 | -- | 30286803 | Not Available | Not Available |
| 2493 | Surface composition regulator | *Salmonella infantis* | | A0A402RIU0 | -- | 30286803; 31476731 | Not Available | Not Available |
| 2494 | Poly-beta-1, 6-N-acetyl-D-glucosamine biosynthesis protein PgaD | *Serratia marcescens* | | A0A658C2L5 | -- | 28636609 | Not Available | Not Available |
| 2495 | Surface composition regulator | *Salmonella enterica subspp. enterica serovar* | Indiana | A0A4Y6N1C4 | -- | 30286803 | Not Available | Not Available |
| 2496 | Surface composition regulator | *Pectobacterium brasiliense* | | A0A086EFR3 | -- | 32193231 | DNA binding, DNA-directed DNA polymerase activity | Not Available |
| 2497 | Poly-beta-1, 6-N-acetyl-D-glucosamine biosynthesis protein PgaD | *Pectobacterium brasiliense* | Pseudomonas maltophilia Xanthomonas maltophilia | A0A2J0SHJ3 | -- | 29170656 | Not Available | Not Available |
| 2498 | Poly-beta-1, 6-N-acetyl-D-glucosamine biosynthesis protein PgaD | *Stenotrophomonas maltophilia* | Pseudomonas maltophilia Xanthomonas maltophilia | A0A1S2B2J2 | -- | 29170656 | Not Available | Not Available |
| 2499 | Surface composition regulator | *Salmonella houtenae* | | A0A5Y6MAD3 | -- | 30286803 | Not Available | Not Available |
| 2500 | Surface composition regulator | *Salmonella typhimurium* | | A0A0F7JA38 | -- | 26089426 | Not Available | Not Available |
| 2501 | Poly-beta-1, 6-N-acetyl-D-glucosamine synthase (Poly-beta-1, 6-GlcNAc synthase) (EC 2.4.1.-) | *Burkholderia ambifaria* | Strain ATCC BAA-244 / AMMD Burkholderia cepacia Strain AMMD | Q0B9V6 | -- | 25931592 | Acetyl glucosaminyl transferase activity | Not Available |
| 2502 | S-ribosylhomocysteine lyase (EC 4.4.1.21) (AI-2 synthesis protein) (Autoinducer-2 production protein LuxS) | *Lacticaseibacillus rhamnosus* | Lactobacillus rhamnosus | A1XC58 | -- | 17098890; 31477907 | Iron ion binding, S-ribosylhomocysteine lyase activity | Not Available |
