## Supplementary material for "‘Targeting’ the Search: An Upgraded Structural and Functional Repository of Antimicrobial Peptides for Biofilm Studies (B-AMP v2.0) with a Focus on Biofilm Protein Targets": Suppl File 6

**Full list of PDB target functions with source organisms and names of biofilm protein targets**

| Source organism → PDB Function ↓ | *Pseudomonas aeruginosa* | *Escherichia coli* | *Staphylococcus aureus* | *Porphyromonas gingivalis* | *Salmonella typhimurium* | *Bacillus subtilis* | *Streptococcus gordonii* | *Streptococcus pyogenes serotype M1* | *Parabacteroides distasonis* | *Streptococcus mutans* | *Acinetobacter spp.* | *Aggregatibacter actinomycetemcomitans* |
|---|---|---|---|---|---|---|---|---|---|---|---|---|
| **Cell adhesion** | Fimbrial subunit CupB6 (Target 37) | Type 1 fimbrin D-mannose specific adhesin (Protein FimH) (Target 2); Protein FimG (Target 33); Adhesin / invasin TibA autotransporter [Cleaved into: Adhesin / invasin TibA (Glycoprotein TibA); Adhesin / invasin TibA translocator] (Target 51); Protein FimF (Target 92) | Biofilm-associated surface protein (Target1) | Minor fimbrium subunit Mfa5 (Target 5); Major fimbrium subunit FimA type-1 (Fimbrillin) (Fimbrilin) (Major fimbrial subunit protein type I) (Target 12); Minor fimbrium anchoring subunit Mfa2 (Minor fimbrial antigen 2)(Target 27); Minor fimbrium tip subunit Mfa3 (Target 28); Minor fimbrium subunit Mfa1 (Pg-II fim a) (Target 29); Minor fimbrium tip subunit MfA4 (Immunoreactive 32 kDa antigen) (Target 36); Major fimbrium subunit FimA type-4 (Fimbrillin) (Fimbrilin) (Major fimbrial subunit protein type-4) (FimA4) (Target 50) | Putative outer Membrane protein (Target 19) Putative fimbriae subunit (Target 20) | | Streptococcal hemagglutinin (Hs antigen) (Sialic acid-binding adhesin) (Target 16) | | Putative fimbrium tip subunit Fim1C (Target 57); Putative fimbrium tip subunit Fim1F (Target 70) | | Trimeric autotransporter adhesin AtaA (TAA AtaA) (Type 5 secretion system autotransporter AtaA) (Target 65) | |
| **Transferase** | 2-heptyl-4(1H)-quinolone synthase subunit PqsB (Target 34); 2-heptyl-4(1H)-quinolone synthase subunit PqsC (EC 2.3.1.230) (Target 35); GGDEF domain-containing protein (Target 41); Anthraniloyl-CoA anthraniloylTransferase (EC 2.3.1.262) (2-heptyl-4(1H)-quinolone synthase PqsD) (PqsD) (Target 77) | Sensor histidine kinase CpxA (EC 2.7.13.3) (Target 61); Diguanylate cyclase DgcZ (DGC) (EC 2.7.7.65) (Zinc-sensory diguanylate cyclase) (Target 67); Sensor protein QseC (EC 2.7.13.3) (Target 91) | Sensor protein kinase WalK (EC 2.7.13.3) (Target 32); CapA (Capsular polysaccharide biosynthesis protein CapA) (Capsular polysaccharide type 8 biosynthesis protein cap8A) (Capsule biosynthesis protein CapA) (Chain length regulator (Capsular polysaccharide biosynthesis)) (Membrane protein CapB2) (Target 56); Sensor protein VraS (EC 2.7.13.3) (Target 58) | | | | | | | | | |
| **Hydrolase/ Hydrolase inhibitor** | PslG (Target 40); Acyl-homoserine lactone acylase PvdQ (AHL acylase PvdQ) (Acyl-HSL acylase PvdQ) (EC 3.5.1.97) [Cleaved into: Acyl-homoserine lactone acylase PvdQ subunit alpha (Acyl-HSL acylase PvdQ subunit alpha); Acyl-homoserine lactone acylase PvdQ subunit beta (Acyl-HSL acylase PvdQ subunit beta)] (Target 68); CRISPR-associated nuclease / helicase Cas3 subtype I-F / YPEST (EC 3.1.-.-) (EC 3.6.4.-) (Target 38); 2-aminobenzoylacetyl-CoA thioesterase (EC | Poly-beta-1, 6-N-acetyl-D-glucosamine N-deacetylase (PGA N-deacetylase) (Poly-beta-1, 6-GlcNAc N-deacetylase) (EC 3.5.1.-) (Target 66); Endoribonuclease toxin MazF (EC 3.1.27.-) (Toxin MazF) (mRNA interferase MazF) (Target 99) | | | | | | | | Putative ABC transporter, ATP-binding protein ComA (Target 21) | | PgaB (Target 46) |
| **Signaling protein/ inhibitor** | Methyl-accepting chemotaxis protein PctA (Target 30); Probable two-component response regulator (Target 74) | Diguanylate cyclase DosC (DGC) (EC 2.7.7.65) (Direct oxygen-sensing cyclase) (Target 39) | Transcriptional regulatory protein LytR (Target 10); signaltransduction protein TRAP (Target of RNAIII-activating protein) (Target 63) | Methyl-accepting chemotaxis protein TlpQ (TlpQ chemoreceptor) (Target 15) | | | | | | Competence stimulating peptide (Competence stimulating protein) (Target 93) | | |
| **Structural protein** | | Fimbrin-like protein FimI (Target 6); Autoinducer-2 kinase (AI-2 kinase) (EC 2.7.1.189) (Target 18); Type-1 fimbrial protein, A chain (Type-1A pilin) (Target 90); O-acetyl-ADP-ribose deacetylase (EC 3.1.1.106) (Regulator of RNAse III activity) (Target 100); Surface composition regulator (Target 101) | Surface protein G (Target 43) | Minor fimbrium tip subunit MfA4 (Immunoreactive 32 kDa antigen PG49) (Target 48) | | Biofilm-surface layer protein A (ORF-1) (Target 62); High-affinity zinc uptake system binding-protein ZnuA (Target 88) | | Pilin (Lancefield T antigen) (Pilus backbone Structural protein) (Target 80) | | | | |
| **Transcription** | | HTH-type Transcriptional regulator RcdA (Regulator of csgD) (Target 22); Transcriptional regulatory protein CpxR (Target 45); Transcription antitermination protein RfaH (Target 87) | | RNA polymerase sigma-H factor (Sigma-30) (Target 13) | | | | | | | | |
| **Immune system / RNA** | CRISPR-associated protein Csy1 (Target 23); CRISPR-associated protein Csy2 (Target 24); CRISPR-associated protein Csy3 (Target 25); CRISPR-associated endonuclease Cas6 / Csy4 (EC 3.1.-.-) (Target 83); | | | | | | | | | | | |
| **Protein fibril** | | Major curlin subunit (Target 14) | | | | TasA anchoring / assembly protein (Biofilm assembly accessory protein TapA) (Target 7); Major biofilm matrix component (Spore coat-associated protein N) (Translocation-dependent antimicrobial spore component) (Target 26) | | | | | | |
| **Sugar binding protein/ inhibitor** | Fucose-binding lectin PA-IIL (Target 79) | Uncharacterized fimbrial-like protein YdeQ (Target 9) | | | | | Platelet binding protein GspB (Adhesin GspB) (Serine-rich adhesin for platelets) (Serine-rich repeat protein GspB) (Target 72) | | | | | |
| **Gene regulation** | Regulatory protein RsaL (RsaL protein) (Virulence gene repressor RsaL) (Target 31) | Probable two-component-system connector protein AriR (Target 86) | | | | | | | | | | |
| **Protein binding** | | AIDA-I autotransporter (AIDA) [Cleaved into: Adhesin AIDA-I; AIDA-I translocator (AIDAc)] (Target 53) | | | | Global transcriptional regulator Spx (Redox-responsive Transcription factor Spx) (Suppressor of clpP and clpX) (Spx) (Target 96) | | | | | | |
| **Membrane protein** | | | Sensor protein SrrB (EC 2.7.13.3) (Staphylococcal respiratory response protein B) (Target 8) | | Autotransporter adhesin SadA (Salmonella adhesin A) (Type 5 secretion system autotransporter SadA) (Target 84) | | | | | | | |
| **RNA binding protein/ chaperone** | Translational regulator CsrA (Carbon storage regulator) (Global Translational regulatory protein RsmA) (Target 98) | Carbon storage regulator (Translational dual regulator CsrA) (Target 97) | | | | | | | | | | |

| Source organism → PDB Function ↓ | *Bacteroides ovatus* | *Bacteroides uniformis* | *Bordetella bronchiseptica* | *Geobacter sulfurreducens* | *Marinobacter nauticus* | *Neisseria meningitidis serogroup B* | *Phocaeicola vulgatus* | *Proteus mirabilis* | *Shewanella oneidensis* | *Staphylococcus epidermidis* | *Streptococcus parasanguinis* | *Streptococcus pneumoniae serotype 4* | *Xanthomonas campestris pv. campestris* |
|---|---|---|---|---|---|---|---|---|---|---|---|---|---|
| **Cell adhesion** | | Putative fimbrium subunit Fim1C (Target 55) | | | | Fimbrial protein (Pilin) (Target 44) | Fimbrium subunit Fim1C (Target 49) | Fimbrial adhesin (Target 3) | | | | | |
| **Transferase** | | | | | | | | | | | | | |
| **Hydrolase/ Hydrolase inhibitor** | | | Putative hemin storage protein (Target 17) | | | | | | | | | | |
| **Signaling protein/ inhibitor** | | | | Diguanylate cyclase (Target 78) | | | | | | | | | Response regulator (Target 71) |
| **Structural protein** | Putative fimbrium tip subunit Fim1C (Target 60) | | | | | | | | | | Fap1 adhesin (Adhesin Fap1) (Fimbriae-associated protein Fap1) (Serine-rich repeat protein Fap1) (Target 69) | Pneumococcal serine-rich repeat protein (PsrP) (Adhesin PsrP) (Serine-rich repeat protein PsrP) (Target 64) | |
| **Transcription** | | | | | | | | | Diguanylate cyclase with PAS sensory domain (EC 2.7.7.65) (Target 73) | Biofilm operon icaADBC HTH-type negative Transcriptional regulator IcaR (Intercellular adhesion protein R) (Target 81) | | | |
| **Immune system / RNA** | | | | | | | | | | | | | |
| **Protein fibril** | | | | | | | | | | | | | |
| **Sugar binding protein/ inhibitor** | | | | | | | | | | | | | |
| **Gene regulation** | | | | | | | | | | | | | |
| **Protein binding** | | | | | | | | | | | | | |
| **Membrane protein** | | | | | | | | | | | | | |
| **RNA binding protein/ chaperone** | | | | | | | | | | | | | |

| Source organism → / PDB Function ↓ | *Pseudomonas aeruginosa* | *Escherichia coli* | *Staphylococcus aureus* | *Porphyromonas gingivalis* | *Salmonella typhimurium* | *Bacillus subtilis* | *Streptococcus gordonii* | *Streptococcus pyogenes serotype M1* | *Parabacteroides distasonis* | *Streptococcus mutans* | *Acinetobacter spp.* | *Aggregatibacter actinomycetemcomitans* | *Bacteroides ovatus* | *Bacteroides uniformis* | *Bordetella bronchiseptica* | *Geobacter sulfurreducens* | *Marinobacter nauticus* | *Neisseria meningitidis serogroup B* | *Phocaeicola vulgatus* | *Proteus mirabilis* | *Shewanella oneidensis* | *Staphylococcus epidermidis* | *Streptococcus parasanguinis* | *Streptococcus pneumoniae serotype 4* | *Xanthomonas campestris pv. campestris* |
|---|---|---|---|---|---|---|---|---|---|---|---|---|---|---|---|---|---|---|---|---|---|---|---|---|---|
| **DNA binding protein / toxin** | | mRNA interferase toxin MqsR (EC 3.1.-.-) (Endoribonuclease MqsR) (Motility quorum-sensing regulator MqsR) (Toxin MqsR) (Target 75); Antitoxin MqsA (Target 76) | | | | | | | | | | | | | | | | | | | | | | | |
| **Transcriptional regulator** | Transcriptional regulator FleQ (Target 42); Exodeoxyribonuclease III (EC 3.1.11.2) (Target 59) | | | | | | | | | | | | | | | | | | | | | | | | |
| **Transport protein** | | | | Phosphate-binding protein PstS (Target 52) | | | | | | | | | | | | | | | | | | | | | |
| **Fimbrial protein** | Type IV major pilin protein PilA (Pilin) (Target 102) | | | | | | | | | | | | | | | | | | | | | | | | |
| **Protein transport** | | Curli production assembly / transport component CsgG (Target 11) | | | | | | | | | | | | | | | | | | | | | | | |
| **Metal binding protein** | | | | | | | | | | | | | | | | | Putative large adhesion protein (Lap) involved in biofilm formation (Target 4) | | | | | | | | |
| **Signaling protein activator** | | Lipoprotein NlpE (Copper homeostasis protein CutF) (Target 82) | | | | | | | | | | | | | | | | | | | | | | | |
| **Peptide binding protein** | Outer membrane porin F (Target 47) | | | | | | | | | | | | | | | | | | | | | | | | |
| **Toxin inhibhitor** | | Antitoxin YefM (Target 95) | | | | | | | | | | | | | | | | | | | | | | | |
| **Calcium binding protein** | | | Serine-rich adhesin for platelets (Adhesin SraP) (Staphylococcus aureus surface protein A) (SasA) (Target 54) | | | | | | | | | | | | | | | | | | | | | | |
| **Ribosome** | | Toxin YoeB (EC 3.1.-.-) (Putative endoribonuclease YoeB) (Putative mRNA interferase Yoeb) (Target 94) | | | | | | | | | | | | | | | | | | | | | | | |
| **Chaperone / protein transport** | | Curli production assembly / transport component CsgE (Target 89) | | | | | | | | | | | | | | | | | | | | | | | |
| **Structural genomics / biosynthetic protein** | | | | | | | | | | | | | | | | | | | | | | | | | |
