## Supplementary material for "‘Targeting’ the Search: An Upgraded Structural and Functional Repository of Antimicrobial Peptides for Biofilm Studies (B-AMP v2.0) with a Focus on Biofilm Protein Targets": Suppl File 10

**Supplementary File 10: Full list of AMP-sortase C docking models for *C. striatum* analyzed using MD simulations**

**Standard**

**Molecular Dynamics trajectory movie:** https://drive.google.com/drive/folders/1mNLQzoKsZHoPzKCD_f8yiQPefJg0EZv4?usp=sharing

| Pep ID | 0ns snapshot of AMP bound with mutated Class C Sortase protein | 20ns snapshot of AMP bound with mutated Class C Sortase protein | AMP-mutated Class C Sortase interactions at 0ns of the MD simulation | AMP-mutated Class C Sortase interactions at 20ns of the MD simulation |
|---|---|---|---|---|
| Pep1 | 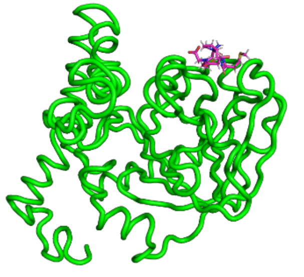 | 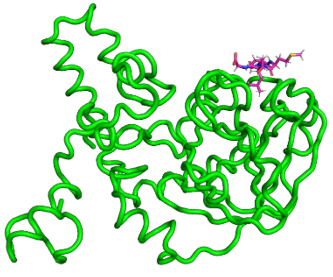 | 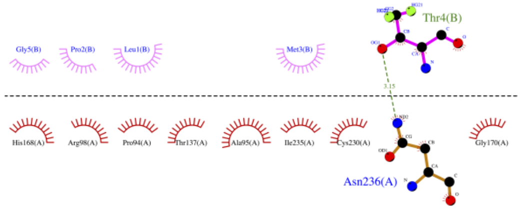 | 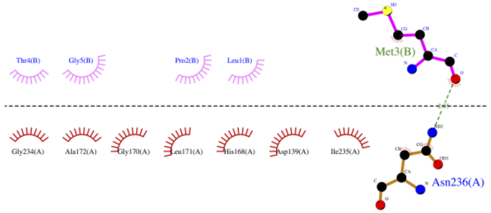 |

**Preference score 10**

**Molecular Dynamics trajectory movies:**
https://drive.google.com/drive/folders/11Wk2zU27oV5Y5mAXLUJ3UgBmtSMkkcdP?usp=sharing

| Pep ID | 0ns snapshot of AMP bound with mutated Class C Sortase protein | 20ns snapshot of AMP bound with mutated Class C Sortase protein | AMP-mutated Class C Sortase interactions at 0ns of the MD simulation | AMP-mutated Class C Sortase interactions at 20ns of the MD simulation |
|---|---|---|---|---|
| Pep1437 | 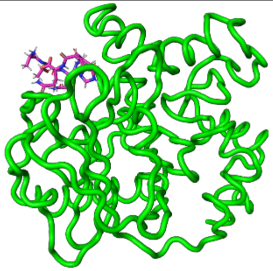 | 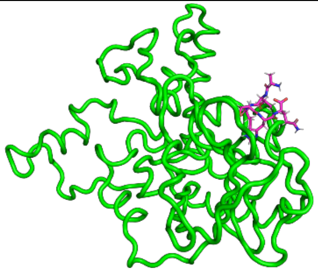 | 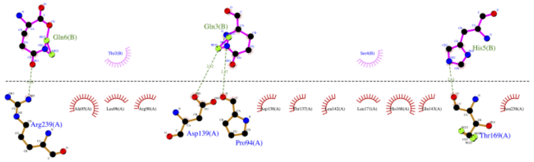 | 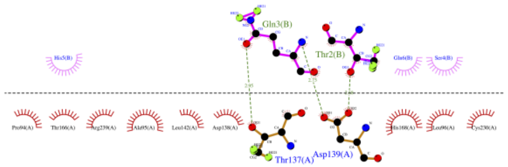 |
| Pep4021 | 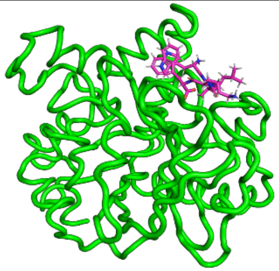 | 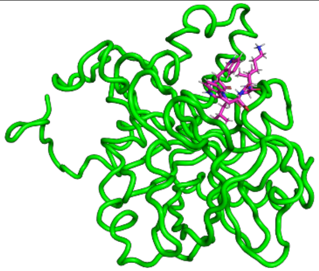 | 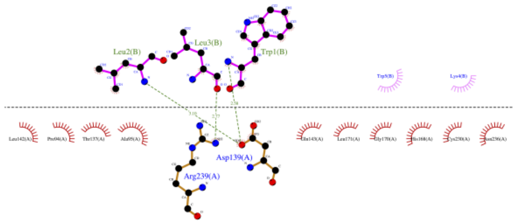 | 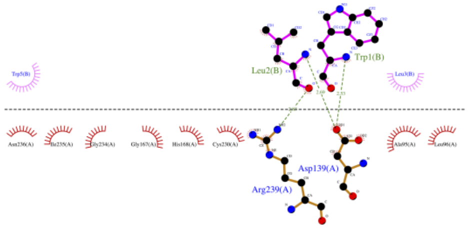 |
| Pep4583 | 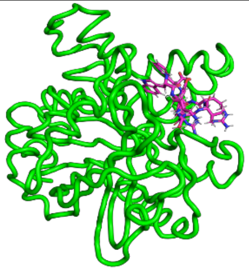 | 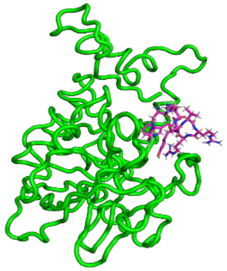 | 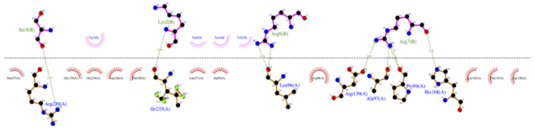 | 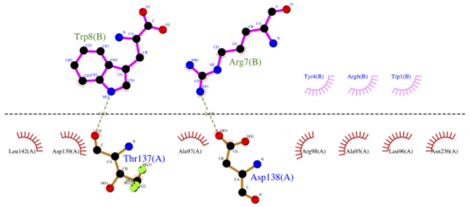 |

| Pep ID | 0ns snapshot of AMP bound with mutated Class C Sortase protein | 20ns snapshot of AMP bound with mutated Class C Sortase protein | AMP-mutated Class C Sortase interactions at 0ns of the MD simulation | AMP-mutated Class C Sortase interactions at 20ns of the MD simulation |
|---|---|---|---|---|
| Pep4707 | 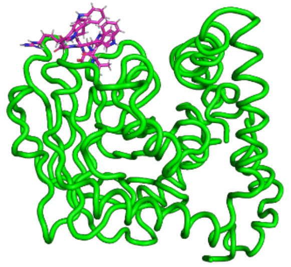 | 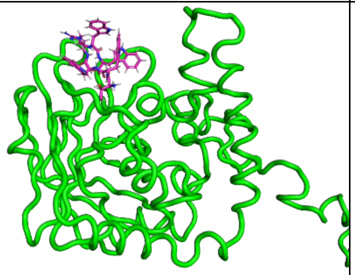 | 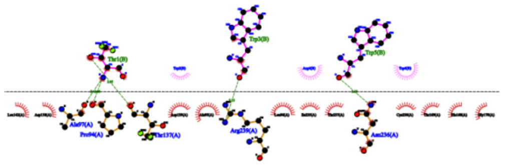 | 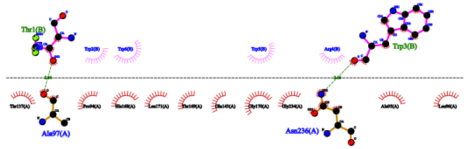 |
| Pep4785 | 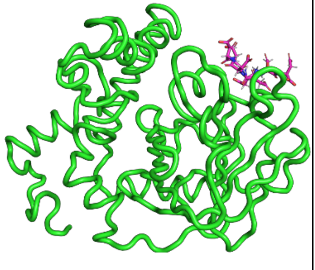 | 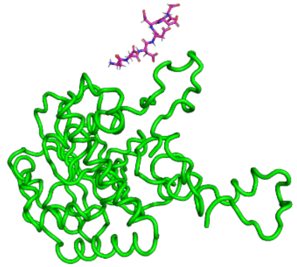 | 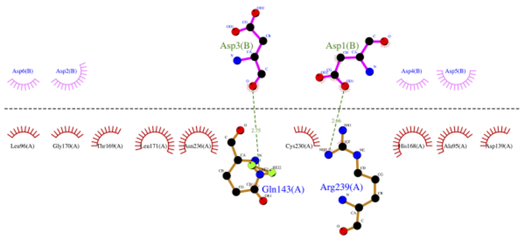  | **AMP drifts away from the complex** |
| Pep5494 | 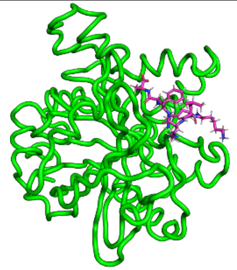 | 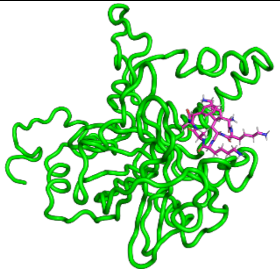 | 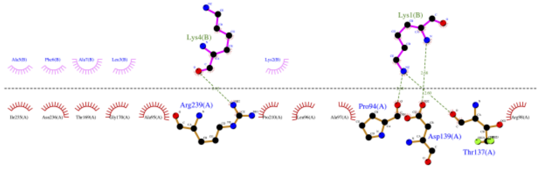  | 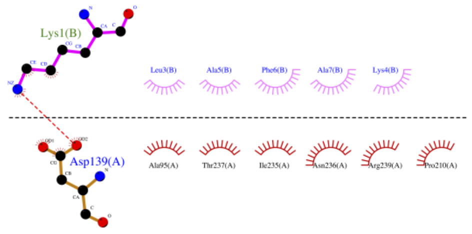  |
| Pep5496 | 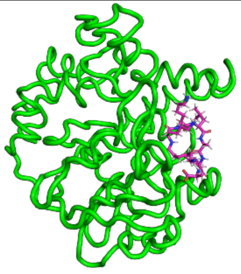 | 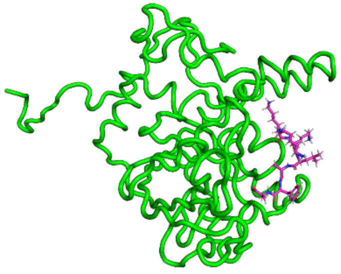 | 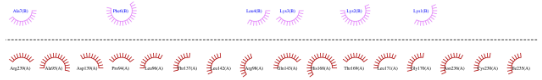 | 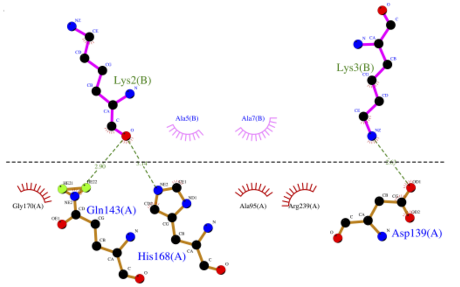  |

**Preference Score 9**

**Molecular Dynamics trajectory movies: https://drive.google.com/drive/folders/1eZxdvx22BRFIOHNBrk5rkGDaTk49ZWH1?usp=sharing**

| Pep ID | 0ns snapshot of AMP bound with mutated Class C Sortase protein | 20ns snapshot of AMP bound with mutated Class C Sortase protein | AMP-mutated Class C Sortase interactions at 0ns of the MD simulation | AMP-mutated Class C Sortase interactions at 20ns of the MD simulation |
|---|---|---|---|---|
| Pep1218 | 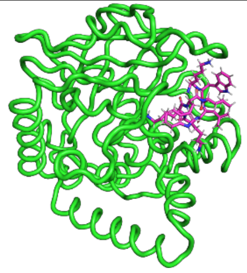 | 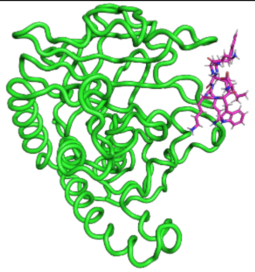 | 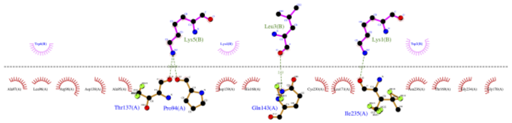 | 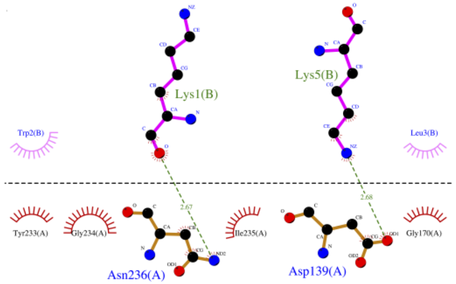 |
| Pep4708 | 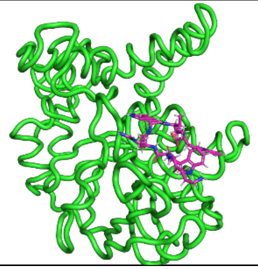 | 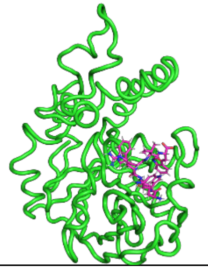 | 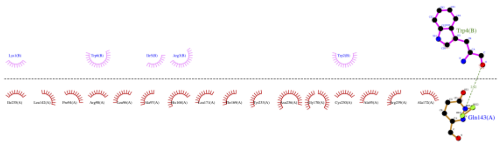 | 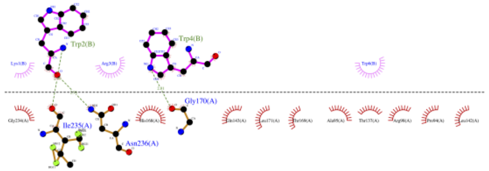 |
| Pep4931 | 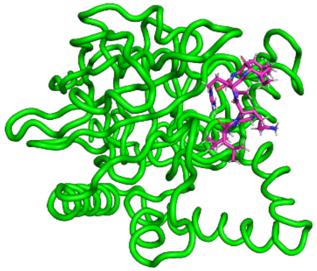 | 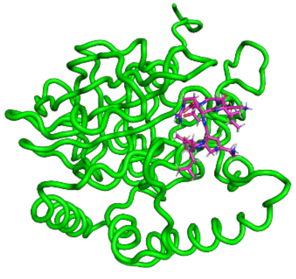 | 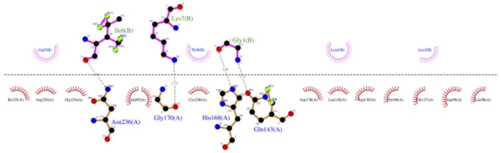 | 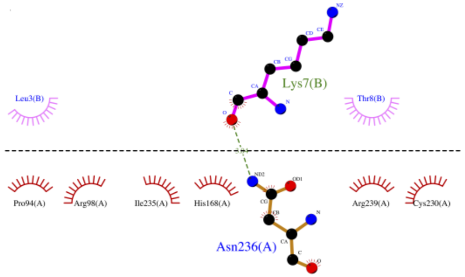 |

| Pep ID | 0ns snapshot of AMP bound with mutated Class C Sortase protein | 20ns snapshot of AMP bound with mutated Class C Sortase protein | AMP-mutated Class C Sortase interactions at 0ns of the MD simulation | AMP-mutated Class C Sortase interactions at 20ns of the MD simulation |
|---|---|---|---|---|
| Pep4934 | 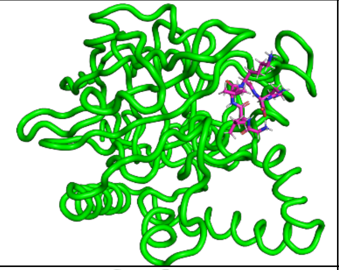 | 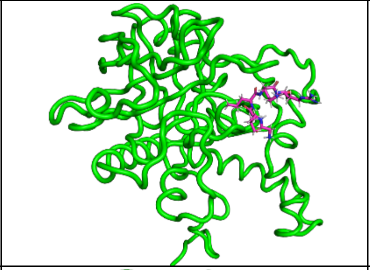 | 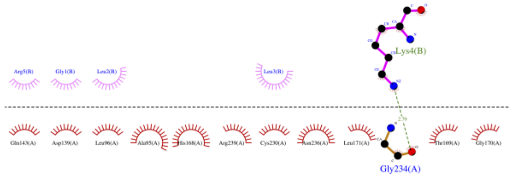  | 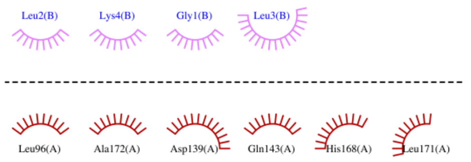  |
| Pep5482 | 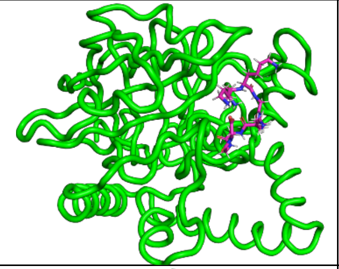 | 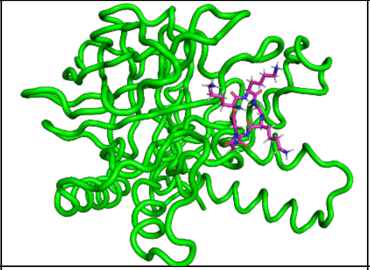 | 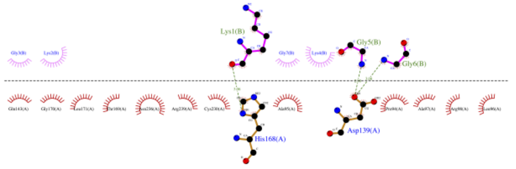  | 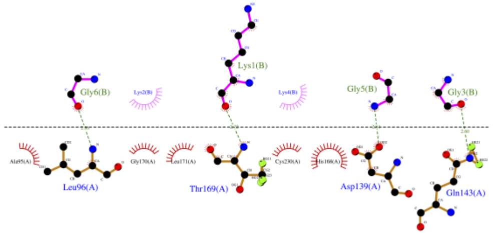  |
| Pep5508 | 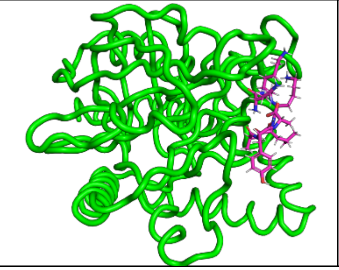 | 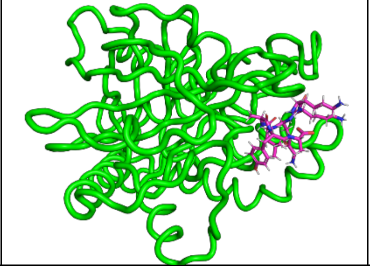 | 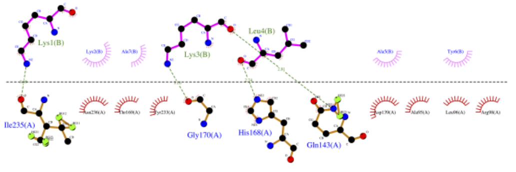  | 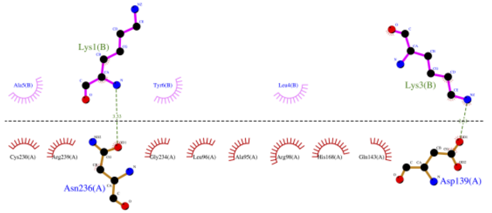  |

### Preference score 8

**Molecular Dynamics trajectory movies:** https://drive.google.com/drive/folders/1uw3buKTkkfMW2E5G_8rweyftcupmQheP?usp=sharing

| Pep ID | 0ns snapshot of AMP bound with mutated Class C Sortase protein | 20ns snapshot of AMP bound with mutated Class C Sortase protein | AMP-mutated Class C Sortase interactions at 0ns of the MD simulation | AMP-mutated Class C Sortase interactions at 20ns of the MD simulation |
|---|---|---|---|---|
| Pep51 | 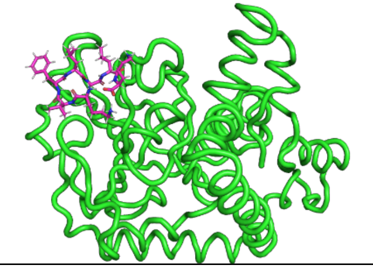 | 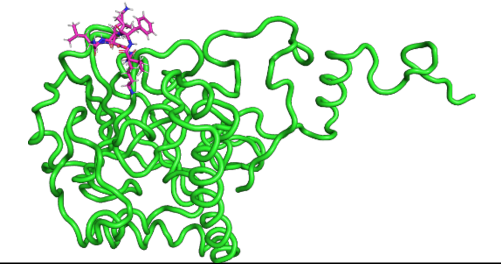 | 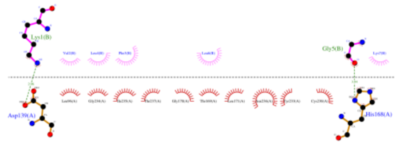 | 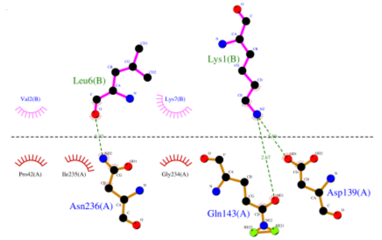  |
| Pep168 | 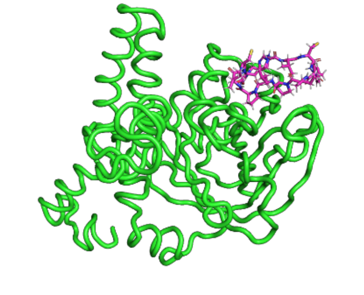 | 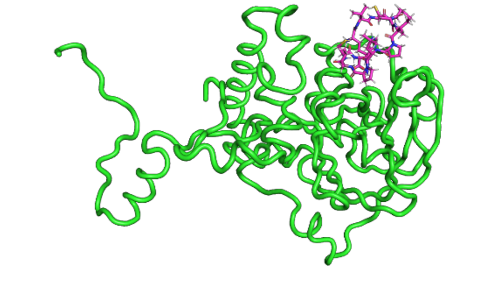 | 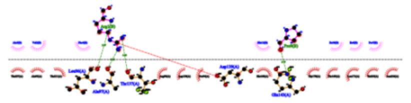 | 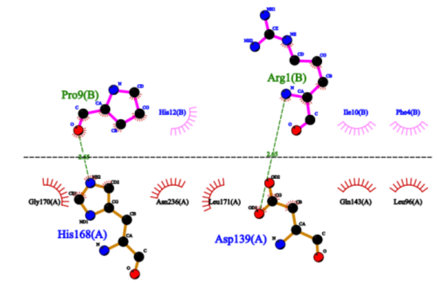  |
| Pep696 | 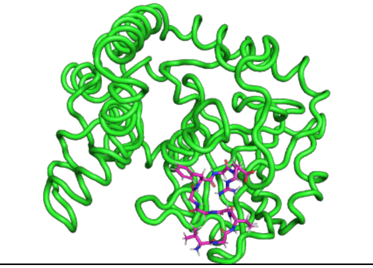 | 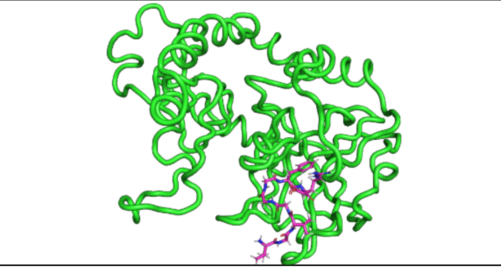 | 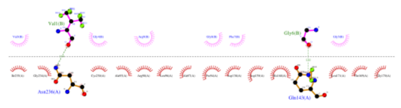 | 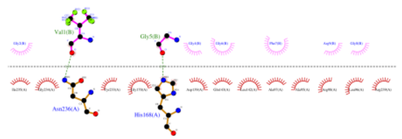 |

| Pep ID | 0ns snapshot of AMP bound with mutated Class C Sortase protein | 20ns snapshot of AMP bound with mutated Class C Sortase protein | AMP-mutated Class C Sortase interactions at 0ns of the MD simulation | AMP-mutated Class C Sortase interactions at 20ns of the MD simulation |
|---|---|---|---|---|
| Pep782 | 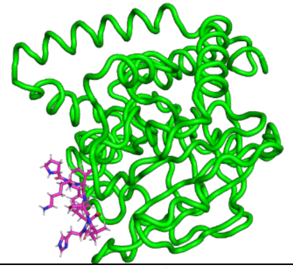 | 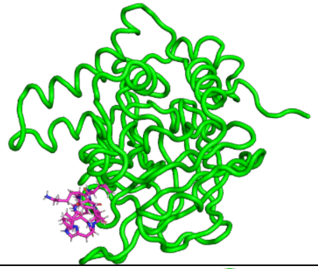 | 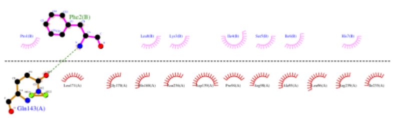 | 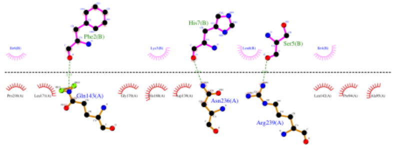 |
| Pep996 | 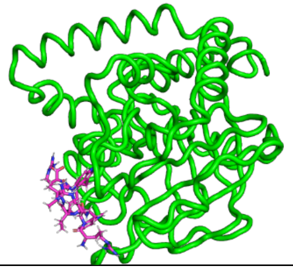 | 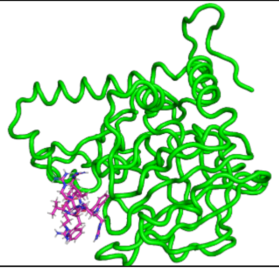 | 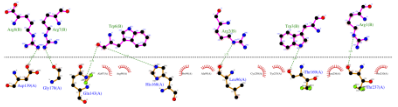 | 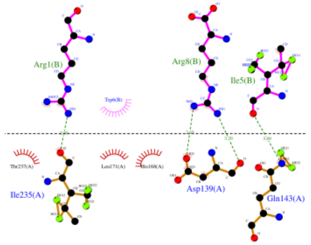 |
| Pep1005 | 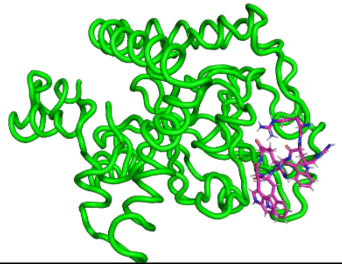 | 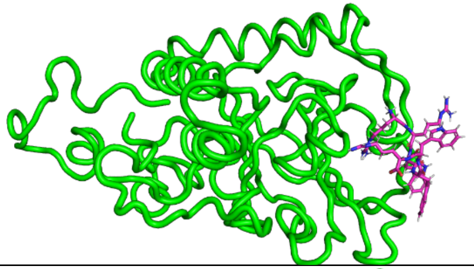 | 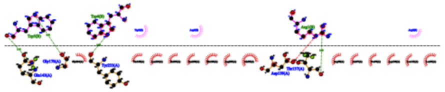 | 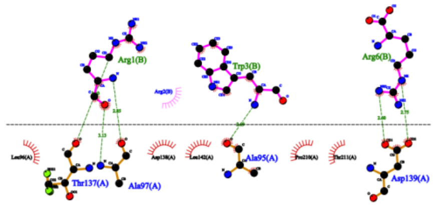 |
| Pep1087 | 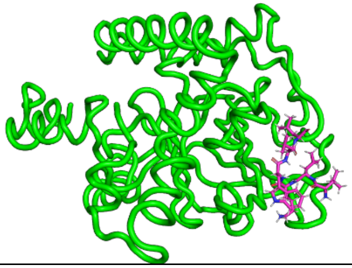 | 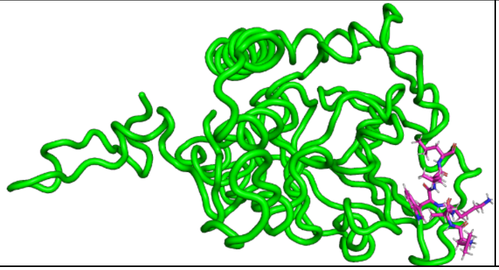 | 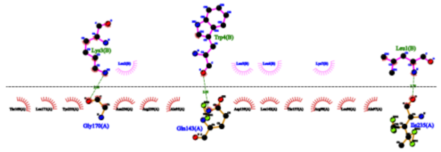 | 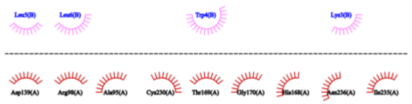 |

| Pep ID | 0ns snapshot of AMP bound with mutated Class C Sortase protein | 20ns snapshot of AMP bound with mutated Class C Sortase protein | AMP-mutated Class C Sortase interactions at 0ns of the MD simulation | AMP-mutated Class C Sortase interactions at 20ns of the MD simulation |
|---|---|---|---|---|
| Pep1219 | 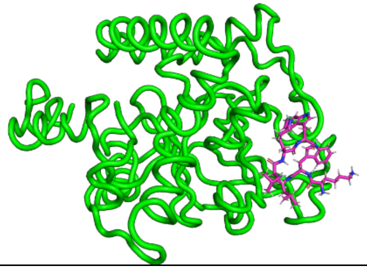 | 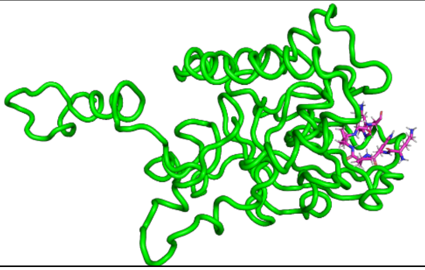 | 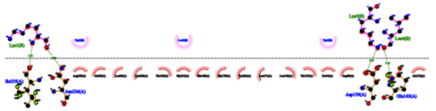 | 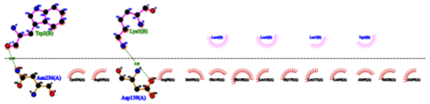 |
| Pep1245 | 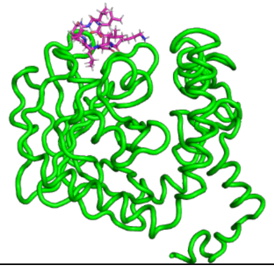 | 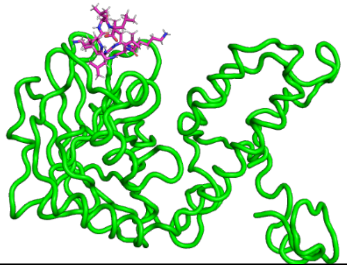 | 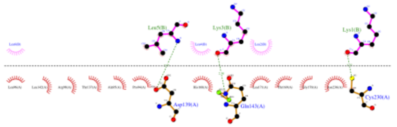 | 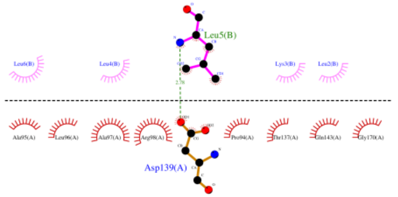 |
| Pep2442 | 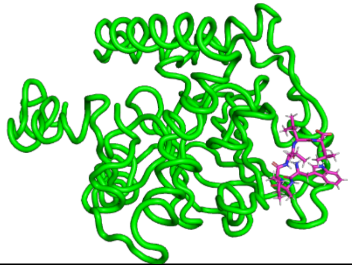 | 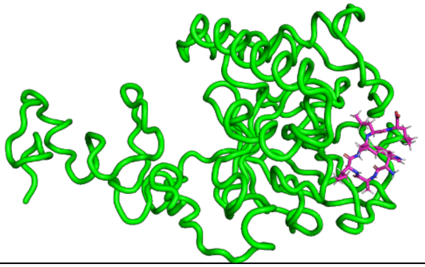 | 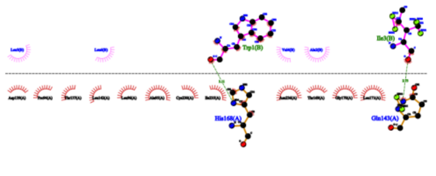 | 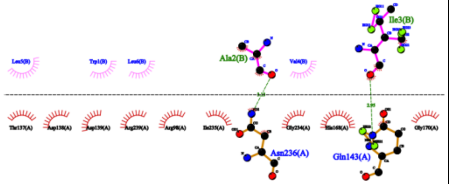 |
| Pep3059 | 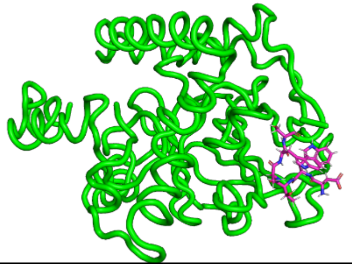 | 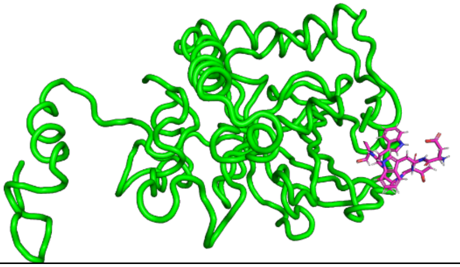 | 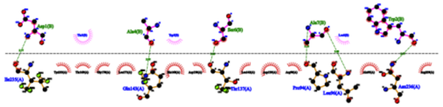 | 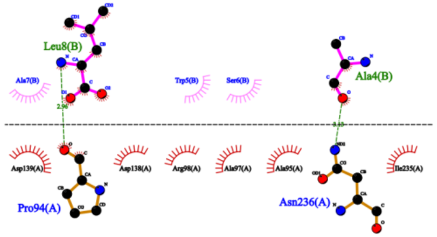 |

| Pep ID | 0ns snapshot of AMP bound with mutated Class C Sortase protein | 20ns snapshot of AMP bound with mutated Class C Sortase protein | AMP-mutated Class C Sortase interactions at 0ns of the MD simulation | AMP-mutated Class C Sortase interactions at 20ns of the MD simulation |
|---|---|---|---|---|
| Pep3219 | 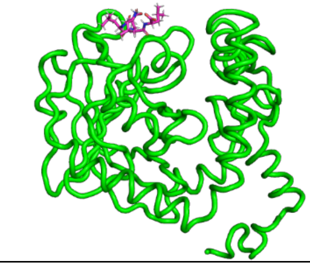 | 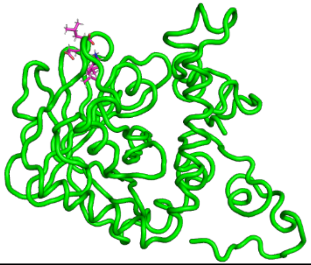 | 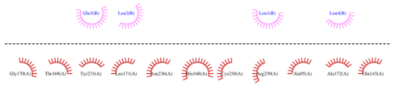 | 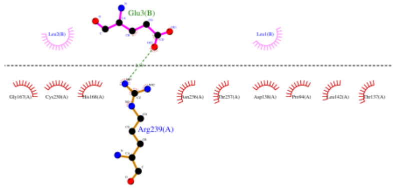 |
| Pep3239 | 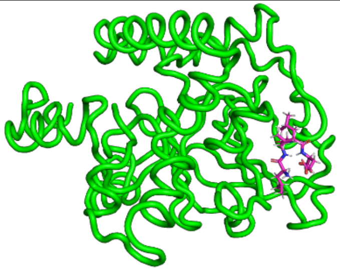 | 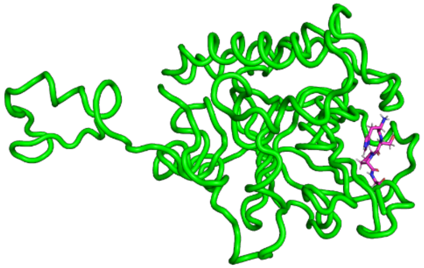 | 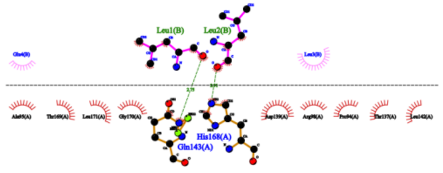 | 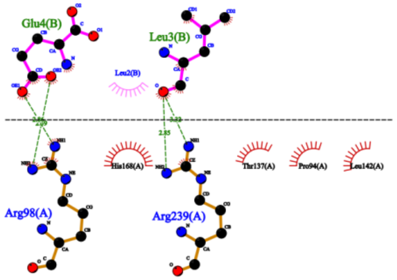 |
| Pep3240 | 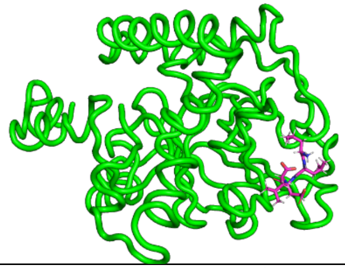 | 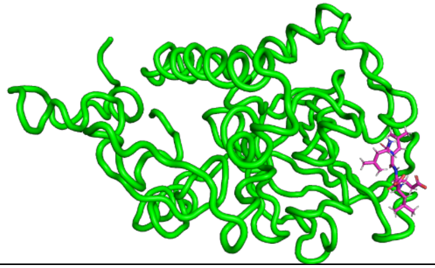 | 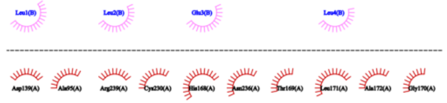 | 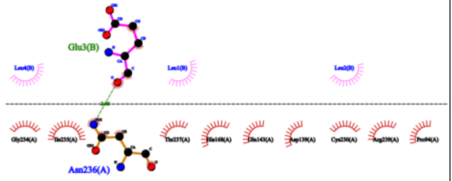 |
| Pep3292 | 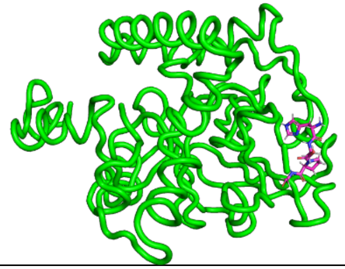 | 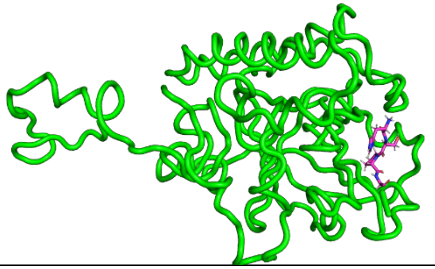 | 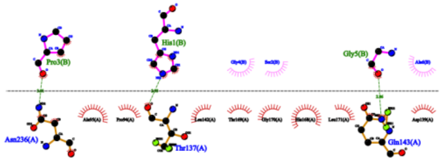 | 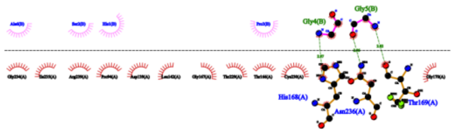 |

| Pep ID | 0ns snapshot of AMP bound with mutated Class C Sortase protein | 20ns snapshot of AMP bound with mutated Class C Sortase protein | AMP-mutated Class C Sortase interactions at 0ns of the MD simulation | AMP-mutated Class C Sortase interactions at 20ns of the MD simulation |
|---|---|---|---|---|
| Pep3296 | 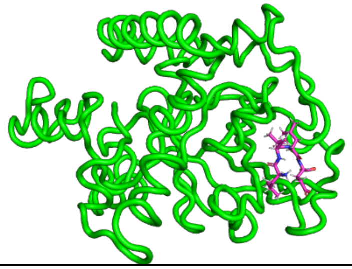 | 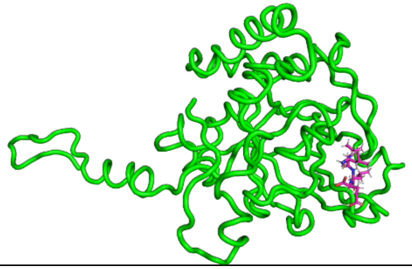 | 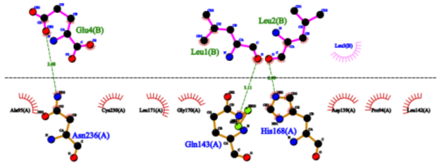 | 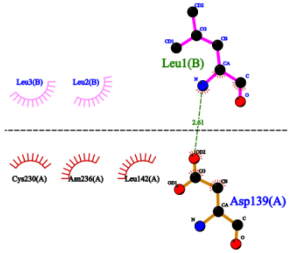 |
| Pep3297 | 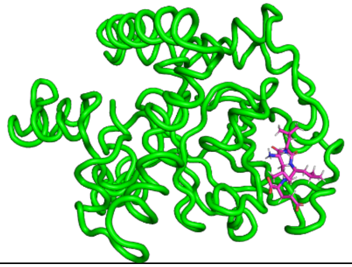 | 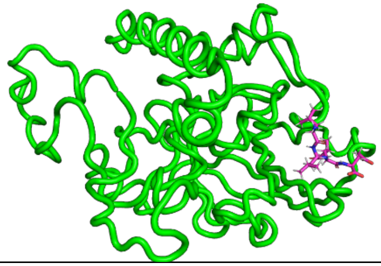 | 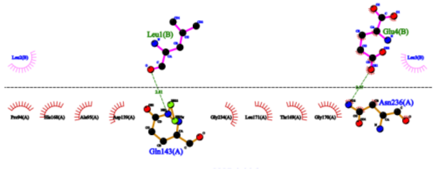 | 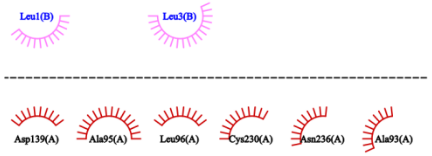 |
| Pep3298 | 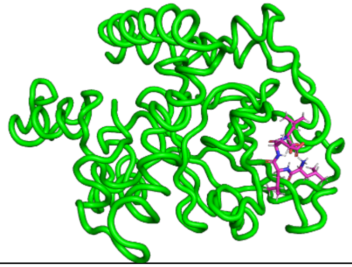 | 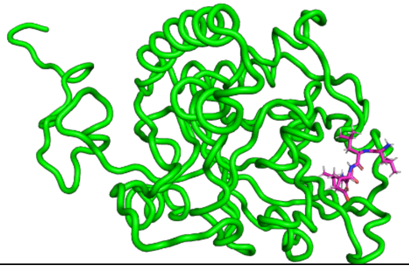 | 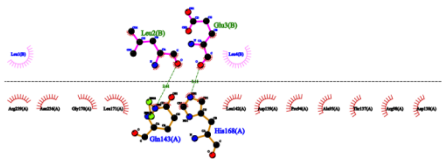 | 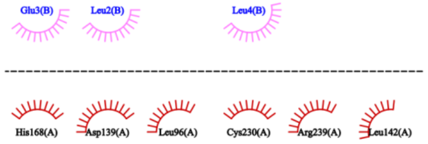 |
| Pep3308 | 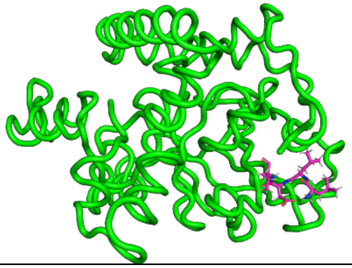 | 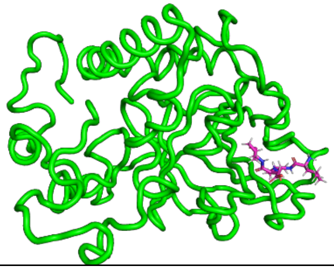 | 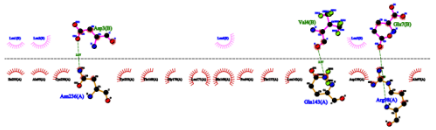 | 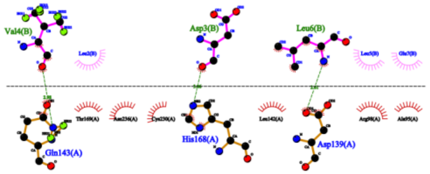 |

| Pep ID | 0ns snapshot of AMP bound with mutated Class C Sortase protein | 20ns snapshot of AMP bound with mutated Class C Sortase protein | AMP-mutated Class C Sortase interactions at 0ns of the MD simulation | AMP-mutated Class C Sortase interactions at 20ns of the MD simulation |
|---|---|---|---|---|
| Pep3366 | 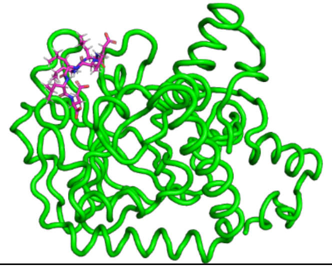 | 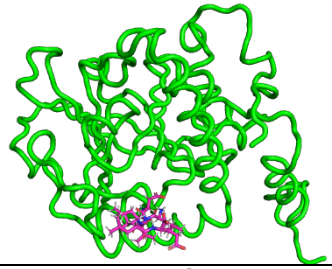 | 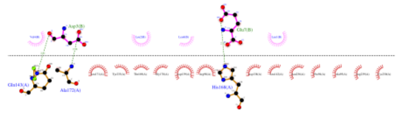 | **No interactions** |
| Pep3461 | 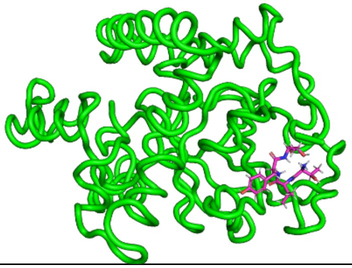 | 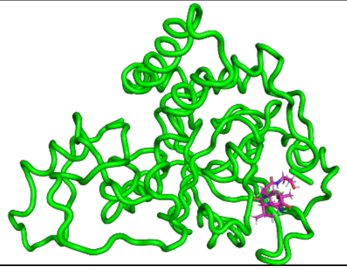 | 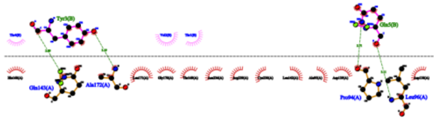 | 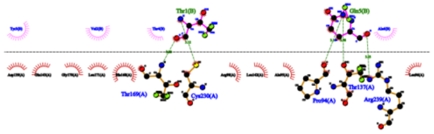 |
| Pep3475 | 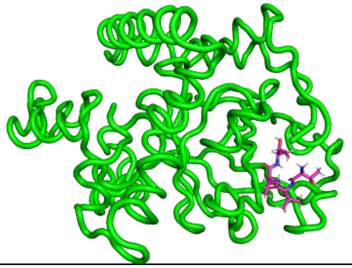 | 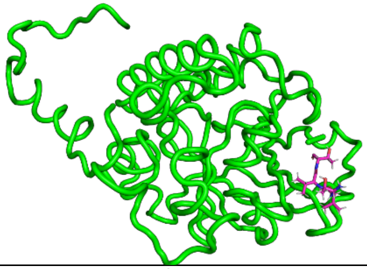 | 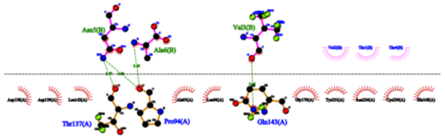 | 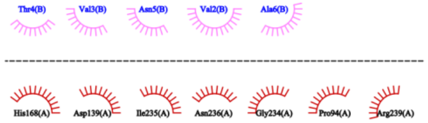 |
| Pep4020 | 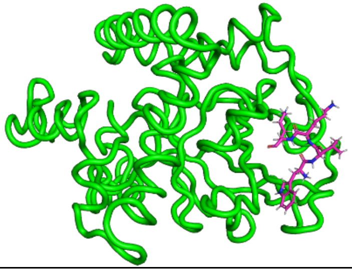 | 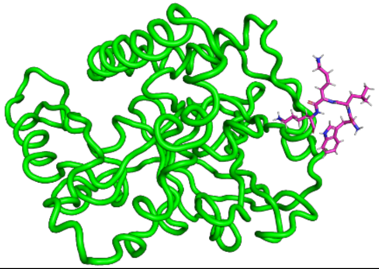 | 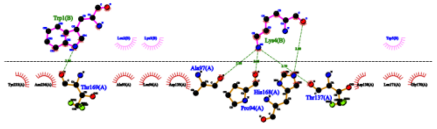 | 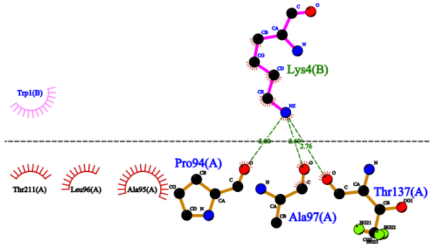 |

| Pep ID | 0ns snapshot of AMP bound with mutated Class C Sortase protein | 20ns snapshot of AMP bound with mutated Class C Sortase protein | AMP-mutated Class C Sortase interactions at 0ns of the MD simulation | AMP-mutated Class C Sortase interactions at 20ns of the MD simulation |
|---|---|---|---|---|
| Pep4674 | 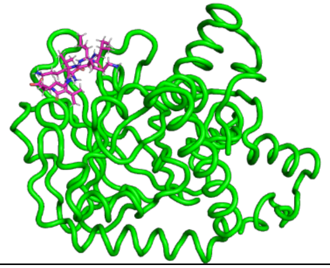 | 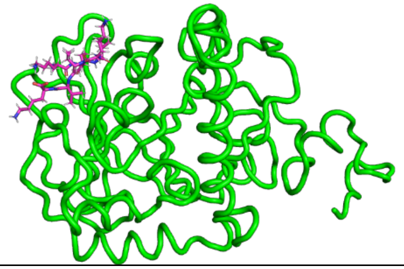 | 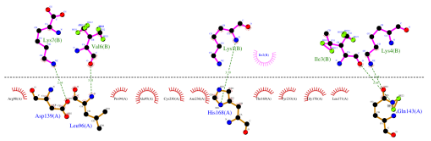 | 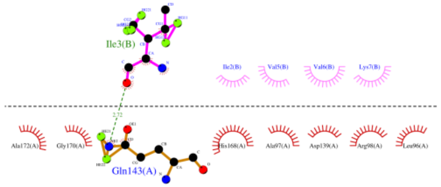 |
| Pep4709 | 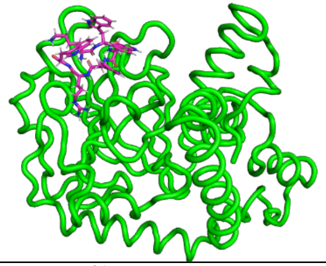 | 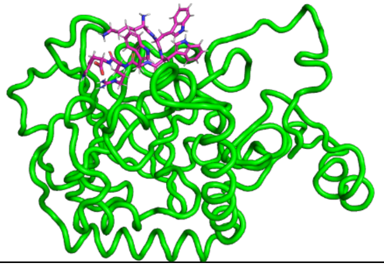 | 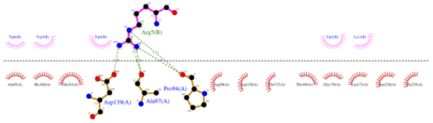 | 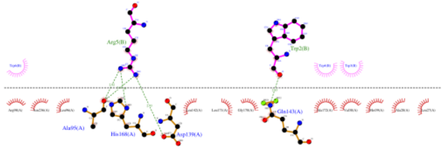 |
| Pep4710 | 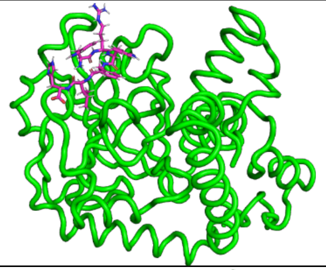 | 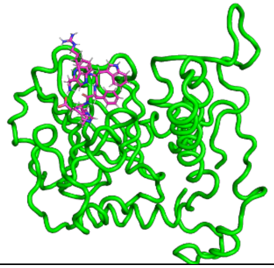 | 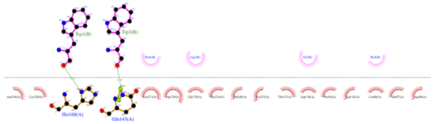 | 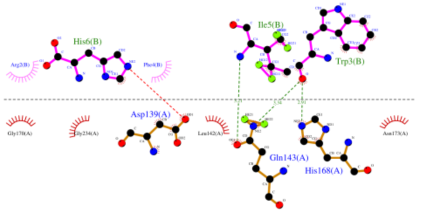 |
| Pep4932 | 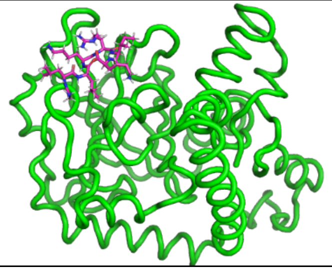 | 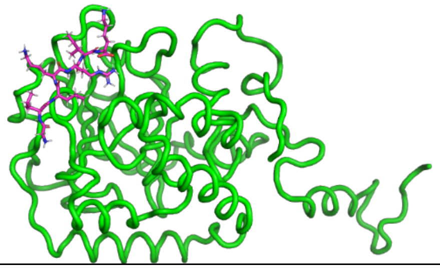 | 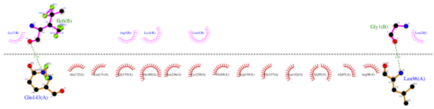 | 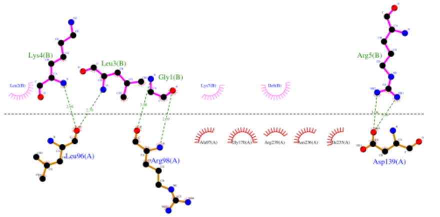 |

| Pep ID | 0ns snapshot of AMP bound with mutated Class C Sortase protein | 20ns snapshot of AMP bound with mutated Class C Sortase protein | AMP-mutated Class C Sortase interactions at 0ns of the MD simulation | AMP-mutated Class C Sortase interactions at 20ns of the MD simulation |
|---|---|---|---|---|
| Pep5241 | 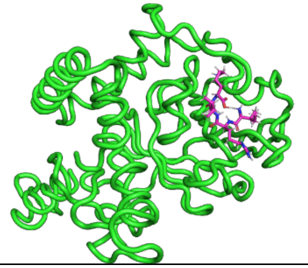 | 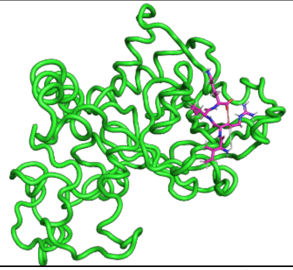 | 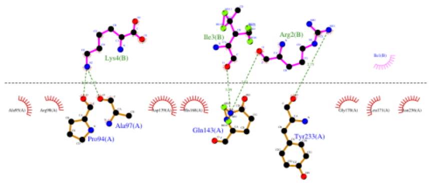 | 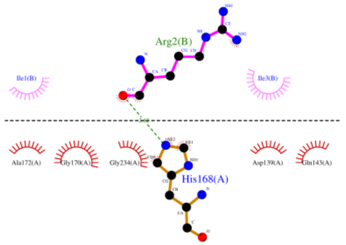 |
| Pep5244 | 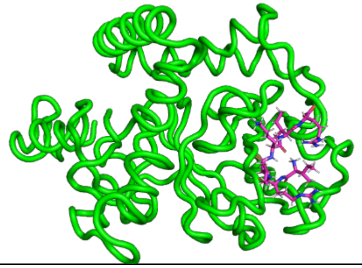 | 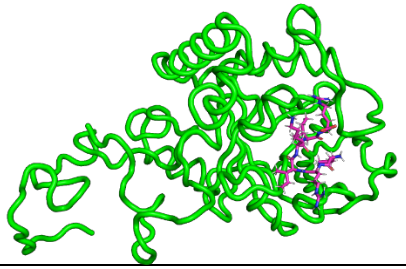 | 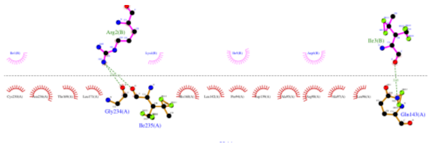 | 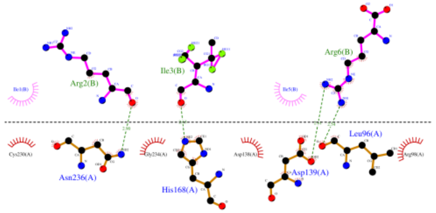 |
| Pep5363 | 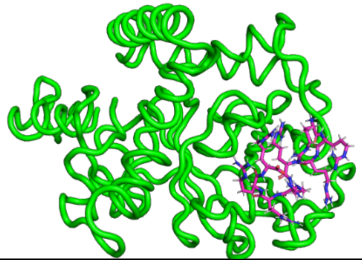 | 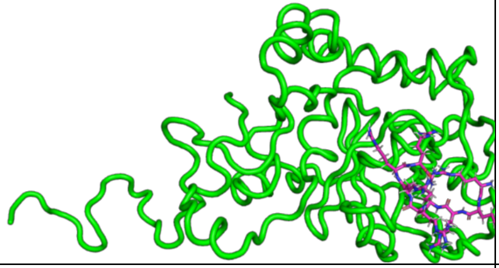 | 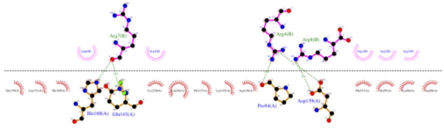 | 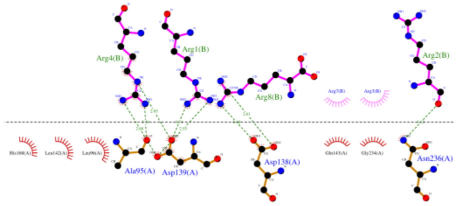 |
| Pep5483 | 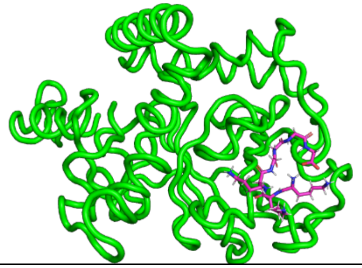 | 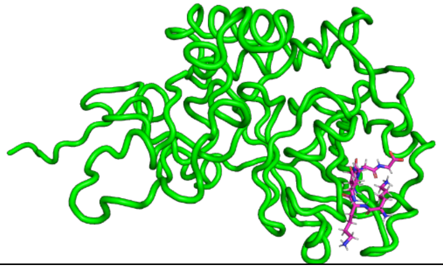 | 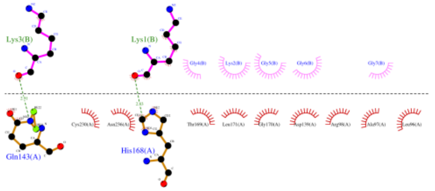 | 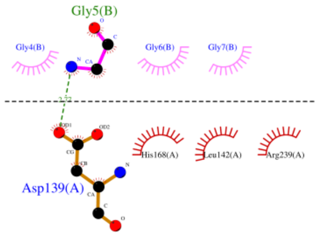 |

| Pep ID | 0ns snapshot of AMP bound with mutated Class C Sortase protein | 20ns snapshot of AMP bound with mutated Class C Sortase protein | AMP-mutated Class C Sortase interactions at 0ns of the MD simulation | AMP-mutated Class C Sortase interactions at 20ns of the MD simulation |
|---|---|---|---|---|
| Pep5485 | 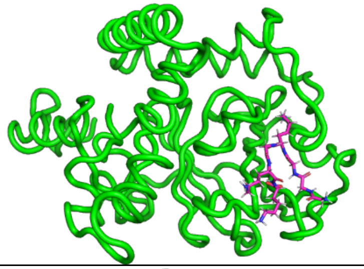 | 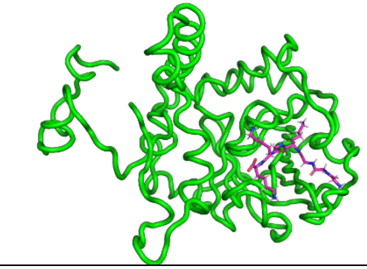 | 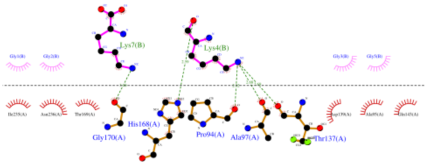 | 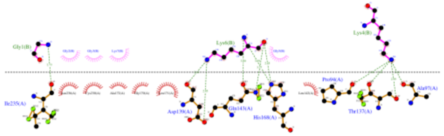 |
| Pep5487 | 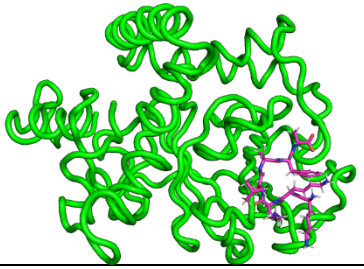 | 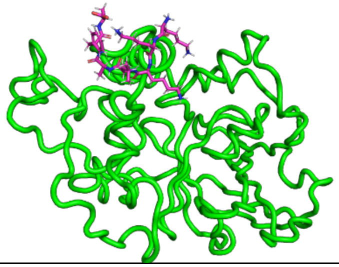 | 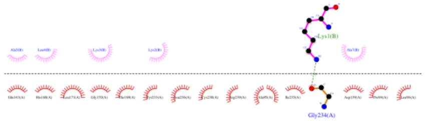 | **AMP drifts away from the complex** |
| Pep5489 | 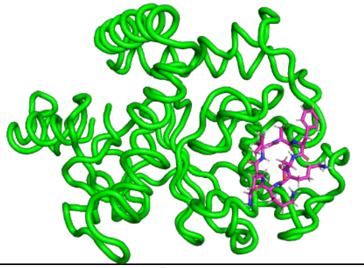 | 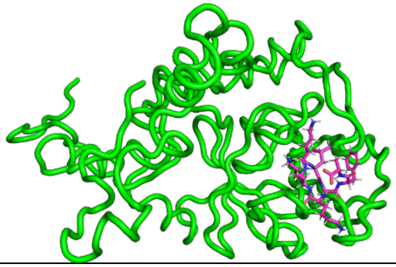 | 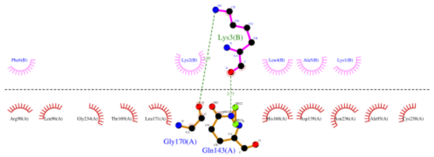 | 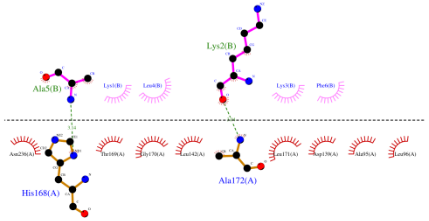 |
| Pep5491 | 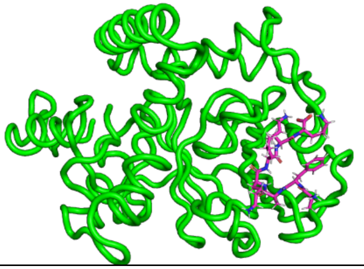 | 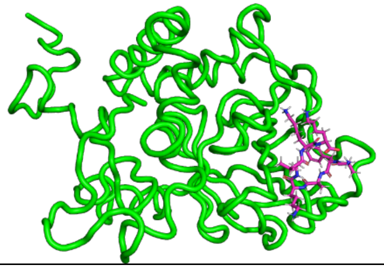 | 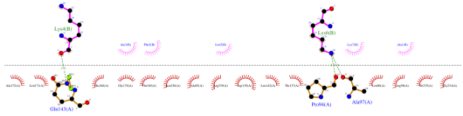 | 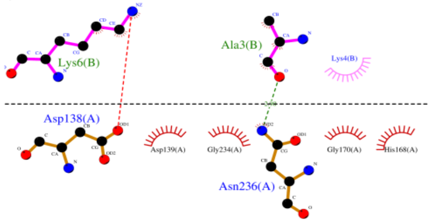  |

| Pep ID | 0ns snapshot of AMP bound with mutated Class C Sortase protein | 20ns snapshot of AMP bound with mutated Class C Sortase protein | AMP-mutated Class C Sortase interactions at 0ns of the MD simulation | AMP-mutated Class C Sortase interactions at 20ns of the MD simulation |
|---|---|---|---|---|
| Pep5492 | 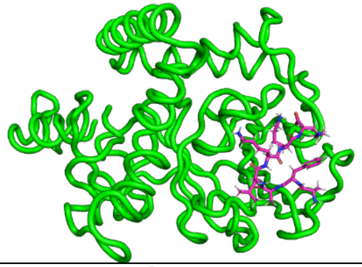 | 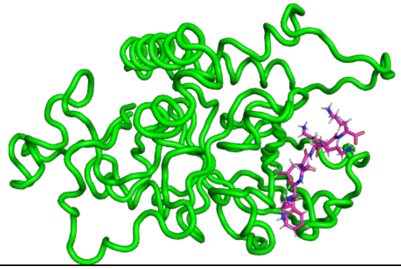 | 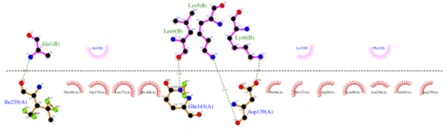 | 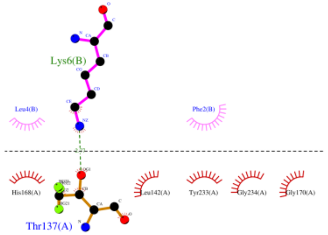 |
| Pep5497 | 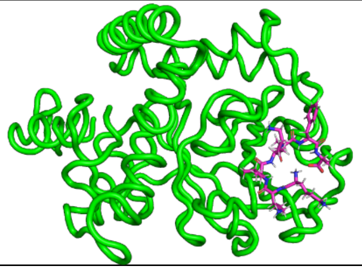 | 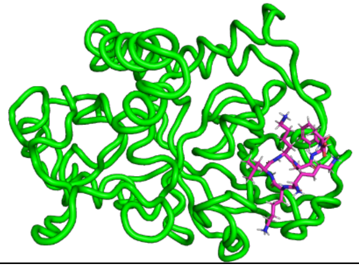 | 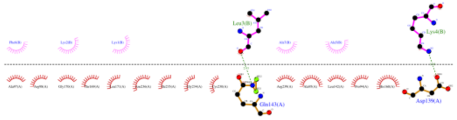 | 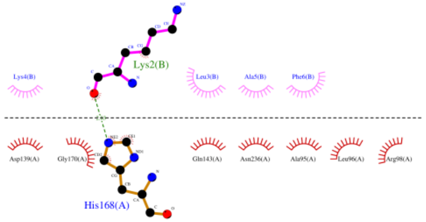 |
| Pep5499 | 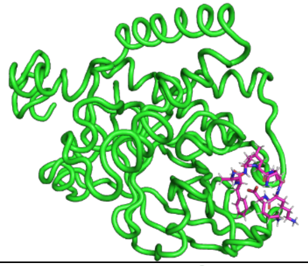 | 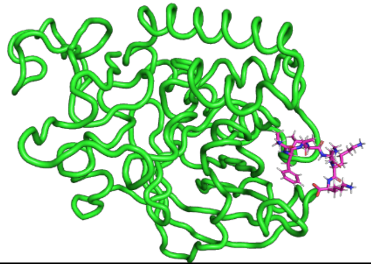 | 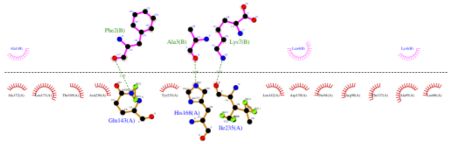 | 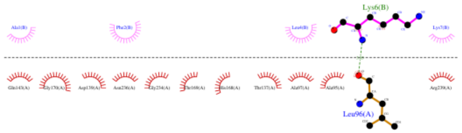 |
| Pep5501 | 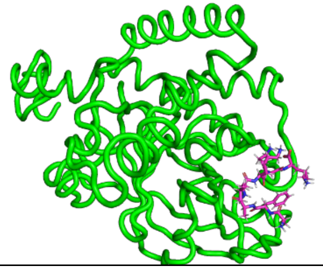 | 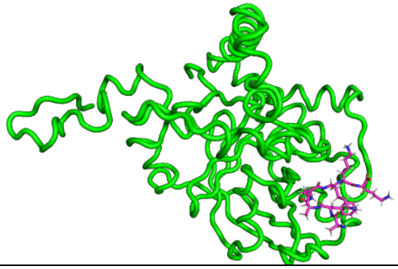 | 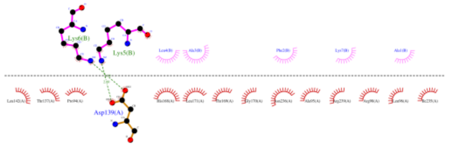 | 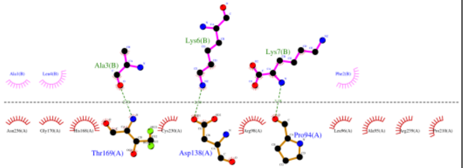 |

| Pep ID | 0ns snapshot of AMP bound with mutated Class C Sortase protein | 20ns snapshot of AMP bound with mutated Class C Sortase protein | AMP-mutated Class C Sortase interactions at 0ns of the MD simulation | AMP-mutated Class C Sortase interactions at 20ns of the MD simulation |
|---|---|---|---|---|
| Pep5502 | 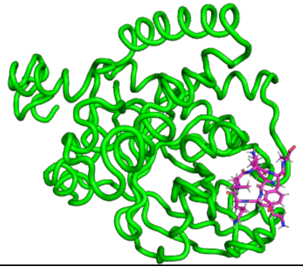 | 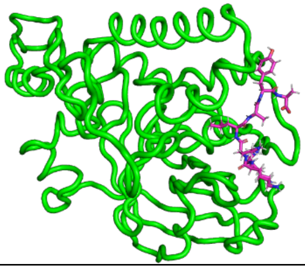 | 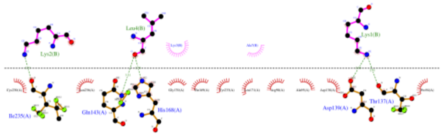 | 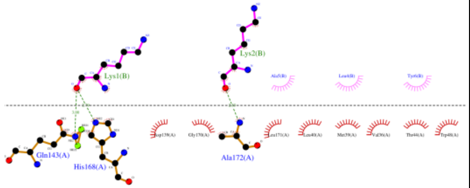 |
| Pep5505 | 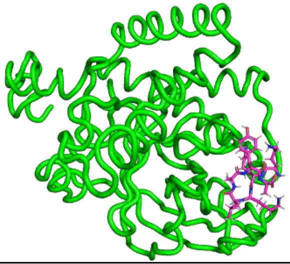 | 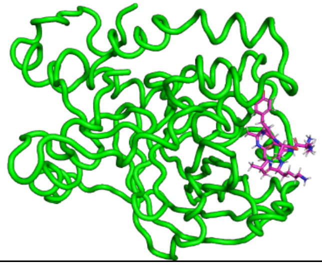 | 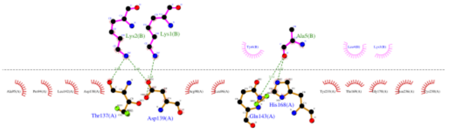 | 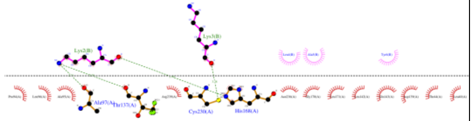 |
| Pep5509 | 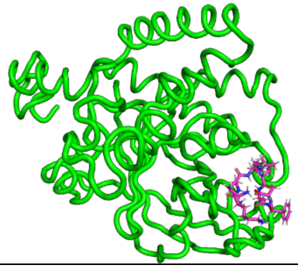 | 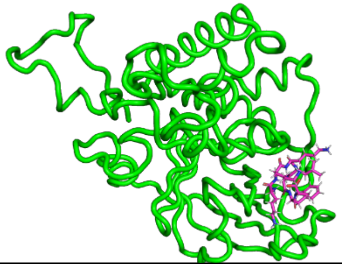 | 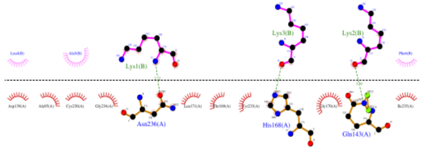 | 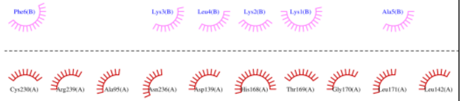 |

**Preference Score 1**

**Molecular Dynamics trajectory movies: https://drive.google.com/drive/folders/1T24rbeKqzZcbryf5HiSG3wcE_fzgFeGu?usp=sharing**

| Pep ID | 0ns snapshot of AMP bound with mutated Class C Sortase protein | 20ns snapshot of AMP bound with mutated Class C Sortase protein | AMP-mutated Class C Sortase interactions at 0ns of the MD simulation | AMP-mutated Class C Sortase interactions at 20ns of the MD simulation |
|---|---|---|---|---|
| Pep3309 | 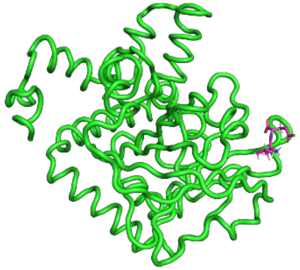 | 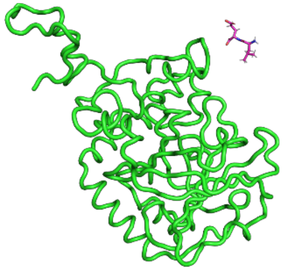 | 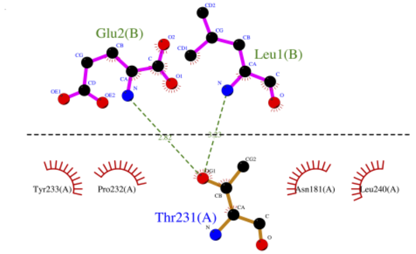 | AMP drifts away from the complex |
| Pep5236 | 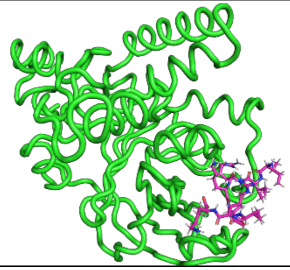 | 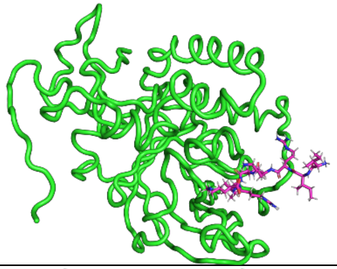 | 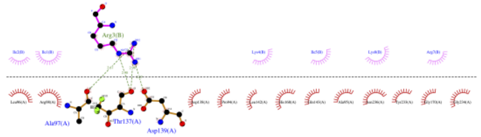 | 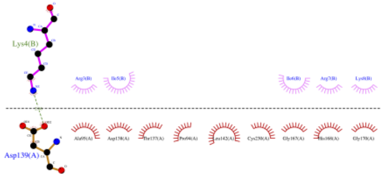 |
| Pep5493 | 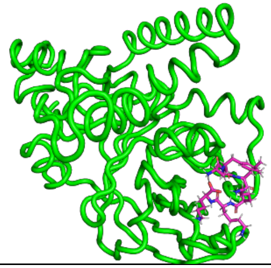 | 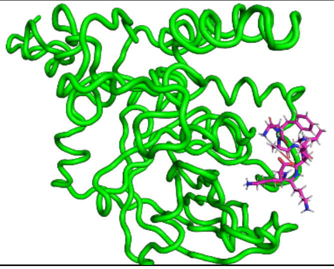 | 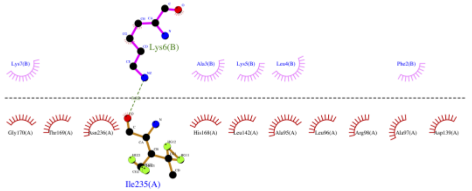 | 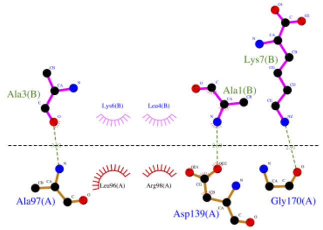 |

**Preference Score 0**

**Molecular Dynamics trajectory movie: [https://drive.google.com/drive/folders/1MRLEgSEre-ixhwyXEjFiBRyxlfFPmj1u?usp=sharing](https://drive.google.com/drive/folders/1MRLEgSEre-ixhwyXEjFiBRyxlfFPmj1u?usp=sharing)**

| **Pep ID** | **0ns snapshot of AMP bound with mutated Class C Sortase protein** | **20ns snapshot of AMP bound with mutated Class C Sortase protein** | **AMP-mutated Class C Sortase interactions at 0ns of the MD simulation** | **AMP-mutated Class C Sortase interactions at 20ns of the MD simulation** |
|---|---|---|---|---|
| Pep993 | 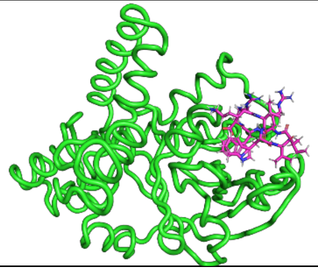 | 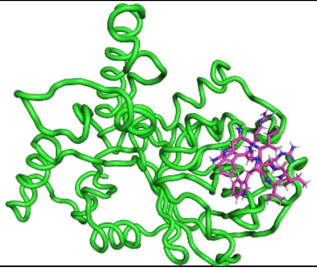 | 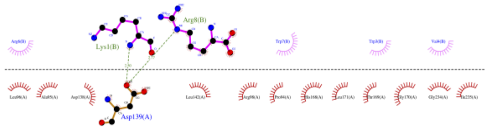 | 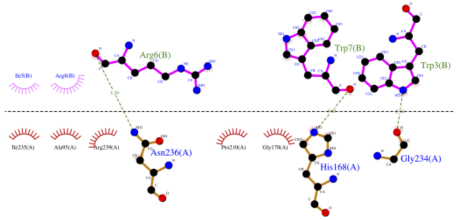 |
