## Supplementary material for "‘Targeting’ the Search: An Upgraded Structural and Functional Repository of Antimicrobial Peptides for Biofilm Studies (B-AMP v2.0) with a Focus on Biofilm Protein Targets": Suppl File 11

| S.no. | PepID | DRAMP_ID | Name of the AMP | Activity |
|---|---|---|---|---|
| 1 | 9 | DRAMP00089 | Bacteriocin E50-52 (Preclinical) | Antibacterial, Anti-Gram+, Anti-Gram-, Antimicrobial |
| 2 | 13 | DRAMP00107 | Bacteriocin L-1077 | Antibacterial, Anti-Gram+, Anti-Gram-, Antimicrobial |
| 3 | 18 | DRAMP00136 | Enterocin E-760 (Bacteriocin) | Antibacterial, Anti-Gram+, Anti-Gram-, Antimicrobial |
| 4 | 19 | DRAMP00171 | Lactocyclicin Q (Bacteriocin) | Antibacterial, Anti-Gram+, Anti-Gram-, Antimicrobial |
| 5 | 34 | DRAMP00336 | ChaC7 (Chassatide C7; uncyclotides; Plant defensin) | Antibacterial, Anti-Gram+, Anti-Gram-, Antimicrobial |
| 6 | 35 | DRAMP00337 | ChaC8 (Chassatide C8; uncyclotides; Plant defensin) | Antibacterial, Anti-Gram+, Anti-Gram-, Antimicrobial |
| 7 | 36 | DRAMP00338 | ChaC11 (Chassatide C11; uncyclotides; Plant defensin) | Antibacterial, Anti-Gram+, Anti-Gram-, Antimicrobial |
| 8 | 44 | DRAMP00431 | Defensin-like protein 2 (Cp-thionin II; Cp-thionin-2; Gamma- thionin | Antibacterial, Anti-Gram+, Anti-Gram-, Antimicrobial |
| 9 | 49 | DRAMP00764 | Piceain 1 (Plants) | Antibacterial, Antifungal, Anti-Gram+, Anti-Gram-, Antimicrobial |
| 10 | 50 | DRAMP00765 | Piceain 2 (Plants) | Antibacterial, Antifungal, Anti-Gram+, Anti-Gram-, Antimicrobial |
| 11 | 51 | DRAMP00766 | JCpep7 (Plants) | Antibacterial, Anti-Gram+, Anti-Gram-, Antimicrobial |
| 12 | 52 | DRAMP00774 | Hedyotide B1 (hB1; Plants) | Antibacterial, Anti-Gram+, Anti-Gram-, Antimicrobial |
| 13 | 53 | DRAMP00795 | Cliotide T1 (cT1; Plant defensin) | Antibacterial, Anticancer, Anti-Gram+, Anti-Gram-, Antimicrobial |
| 14 | 54 | DRAMP00798 | Cliotide T4 (cT4; Plant defensin) | Antibacterial, Anticancer, Anti-Gram+, Anti-Gram-, Antimicrobial |
| 15 | 55 | DRAMP00856 | Kalata-B1 (Plant defensin) | Antibacterial, Antifungal, Insecticidal, Anti-Gram+, Anti-Gram-, Antimicrobial |
| 16 | 56 | DRAMP00877 | Circulin-A (CIRA; Plant defensin) | Antibacterial, Antifungal, Antiviral, Anti-Gram+, Anti-Gram-, Antimicrobial |
| 17 | 57 | DRAMP00878 | Circulin-B (CIRB; Plant defensin) | Antibacterial, Antifungal, Antiviral, Anti-Gram+, Anti-Gram-, Antimicrobial |
| 18 | 58 | DRAMP01374 | Odorranain-D1 (OdD1; Frogs, amphibians, animals) | Antimicrobial, Antibacterial, Antifungal, Anti-Gram+, Anti-Gram-, |
| 19 | 59 | DRAMP01373 | Odorranain-C1 (OdC1; Frogs, amphibians, animals) | Antimicrobial, Antibacterial, Antifungal, Anti-Gram+, Anti-Gram-, |
| 20 | 60 | DRAMP01372 | Odorranain-B1 (Frogs, amphibians, animals) | Antimicrobial, Antibacterial, Antifungal, Anti-Gram+, Anti- Gram-, |
| 21 | 66 | DRAMP01018 | Cyclopsychotride-A (CPT; Plant defensin) | Antibacterial, Antifungal, Anti-Gram+, Anti-Gram-, Antimicrobial |
| 22 | 68 | DRAMP18193 | Cathelicidin-related peptide crotalicidin | Antibacterial, Anti-Gram+, Anti-Gram-, Antimicrobial |
| 23 | 69 | DRAMP01064 | Anticancerous peptide 1 (Cr-ACP1; Plants) | Anticancer, Antibacterial, Anti-Gram+, Anti-Gram-, Antimicrobial |
| 24 | 75 | DRAMP01088 | Alyteserin-1Ma (toads, amphibians, animals) | Antibacterial, Antifungal, Anti-Gram+, Anti-Gram-, Antimicrobial |
| 25 | 76 | DRAMP01089 | Alyteserin-1Mb (toads, amphibians, animals) | Antibacterial, Antifungal, Anti-Gram+, Anti-Gram-, Antimicrobial |
| 26 | 77 | DRAMP01090 | Alyteserin-2Ma (toads, amphibians, animals) | Antibacterial, Antifungal, Anti-Gram+, Anti-Gram-, Antimicrobial |
| 27 | 83 | DRAMP02090 | Brevinin-1Lb (Frogs, amphibians, animals) | Antibacterial, Anti-Gram+, Anti-Gram-, Antimicrobial |
| 28 | 87 | DRAMP01518 | Esculentin-2L (Frogs, amphibians, animals) | Antimicrobial, Antibacterial, Antifungal, Anti-Gram+, Anti- Gram-, |
| 29 | 90 | DRAMP01516 | Esculentin-2B (Frogs, amphibians, animals) | Antimicrobial, Antibacterial, Antifungal, Anti-Gram+, Anti-Gram-, |
| 30 | 91 | DRAMP02077 | Brevinin-1Pb (Frogs, amphibians, animals) | Antimicrobial, Antibacterial, Antifungal, Anti-Gram+, Anti-Gram-, |
| 31 | 98 | DRAMP01162 | Buforin-1 (Buforin I; Fragment of Histone H2A; toads, amphibians, | Antibacterial, Antifungal, Anti-Gram+, Anti-Gram-, Antimicrobial |
| 32 | 99 | DRAMP01163 | Buforin-2 (Buforin II; Fragment of Histone H2A; toads, amphibians, | Antibacterial, Antifungal, Anti-Gram+, Anti-Gram-, Antimicrobial |
| 33 | 100 | DRAMP01164 | Bombinin (toads, amphibians, animals) | Antibacterial, Anti-Gram+, Anti-Gram-, Antimicrobial |
| 34 | 102 | DRAMP01167 | Hylaseptin-P1 (HSP1) | Antibacterial, Anti-Gram+, Anti-Gram-, Antimicrobial |
| 35 | 103 | DRAMP01170 | Distinctin 2 (Frogs, amphibians, animals) | Antibacterial, Anti-Gram+, Anti-Gram-, Antimicrobial |
| 36 | 104 | DRAMP01174 | Ocellatin-4 (Frogs, amphibians, animals) | Antibacterial, Anti-Gram+, Anti-Gram-, Antimicrobial |
| 37 | 106 | DRAMP01182 | Ocellatin-P1 (Pentadactylin; Frogs, amphibians, animals) | Antibacterial, Anti-Gram+, Anti-Gram-, Antimicrobial |
| 38 | 107 | DRAMP01184 | SPX(1-22)(truncated peptide of Syphaxin; Frogs, amphibians, | Antibacterial, Anti-Gram+, Anti-Gram-, Antimicrobial |
| 39 | 108 | DRAMP01185 | SPX(1-16)(truncated peptide of Syphaxin; Frogs, amphibians, | Antibacterial, Anti-Gram+, Anti-Gram-, Antimicrobial |
| 40 | 109 | DRAMP01188 | Chensinin-1ZHa (Frogs, amphibians, animals) | Antibacterial, Antifungal, Anti-Gram+, Anti-Gram-, Antimicrobial |
| 41 | 110 | DRAMP01189 | Andersonin-W1 (Frogs, amphibians, animals) | Antibacterial, Antifungal, Anti-Gram+, Anti-Gram-, Antimicrobial |
| 42 | 111 | DRAMP01190 | Andersonin-W2 (Frogs, amphibians, animals) | Antibacterial, Antifungal, Anti-Gram+, Anti-Gram-, Antimicrobial |
| 43 | 112 | DRAMP01191 | Andersonin-X1 (Frogs, amphibians, animals) | Antibacterial, Antifungal, Anti-Gram+, Anti-Gram-, Antimicrobial |
| 44 | 113 | DRAMP01192 | Andersonin-Y1 (Frogs, amphibians, animals) | Antibacterial, Antifungal, Anti-Gram+, Anti-Gram-, Antimicrobial |
| 45 | 114 | DRAMP01194 | Andersonin-C1 (Frogs, amphibians, animals) | Antibacterial, Antifungal, Anti-Gram+, Anti-Gram-, Antimicrobial |
| 46 | 115 | DRAMP01195 | Andersonin-D1 (Frogs, amphibians, animals) | Antibacterial, Antifungal, Anti-Gram+, Anti-Gram-, Antimicrobial |
| 47 | 116 | DRAMP01199 | Hejiangin-A1 (Frogs, amphibians, animals) | Antibacterial, Antifungal, Anti-Gram+, Anti-Gram-, Antimicrobial |
| 48 | 117 | DRAMP01200 | Hejiangin-F1 (frog, amphibians, animals) | Antibacterial, Antifungal, Anti-Gram+, Anti-Gram-, Antimicrobial |
| 49 | 118 | DRAMP01201 | Schmackerin-C1 (Frogs, amphibians, animals) | Antibacterial, Antifungal, Anti-Gram+, Anti-Gram-, Antimicrobial |
| 50 | 122 | DRAMP01208 | Pleurain-A1 (Pleurain A1; Frogs, amphibians, animals) | Antibacterial, Antifungal, Anti-Gram+, Anti-Gram-, Antimicrobial |
| 51 | 123 | DRAMP01209 | Pleurain-A2 (Pleurain A2; Frogs, amphibians, animals) | Antibacterial, Antifungal, Anti-Gram+, Anti-Gram-, Antimicrobial |
| 52 | 124 | DRAMP01214 | Kassinatuerin-2Ma (Frogs, amphibians, animals) | Antibacterial, Anti-Gram+, Anti-Gram-, Antimicrobial |
| 53 | 125 | DRAMP01218 | Kassinatuerin-1 (Frogs, amphibians, animals) | Antibacterial, Antifungal, Anti-Gram+, Anti-Gram-, Antimicrobial |

| S.no. | PepID | DRAMP_ID | Name of the AMP | Activity |
|---|---|---|---|---|
| 54 | 127 | DRAMP01222 | Palustrin-2AJ1 (PL2AJ1; Frogs, amphibians, animals) | Antibacterial, Anti-Gram+, Anti-Gram-, Antimicrobial |
| 55 | 132 | DRAMP01232 | Palustrin-2ISc (Frogs, amphibians, animals) | Antibacterial, Anti-Gram+, Anti-Gram-, Antimicrobial |
| 56 | 136 | DRAMP01237 | Palustrin-2ISa (Frogs, amphibians, animals) | Antibacterial, Antifungal, Anti-Gram+, Anti-Gram-, Antimicrobial |
| 57 | 137 | DRAMP01238 | Palustrin-2SIb (Frogs, amphibians, animals) | Antibacterial, Anti-Gram+, Anti-Gram-, Antimicrobial |
| 58 | 138 | DRAMP01244 | Japonicin-1 (Frogs, amphibians, animals) | Antibacterial, Anti-Gram+, Anti-Gram-, Antimicrobial |
| 59 | 139 | DRAMP01245 | Japonicin-1CDYa (Frogs, amphibians, animals) | Antibacterial, Anti-Gram+, Anti-Gram-, Antimicrobial |
| 60 | 140 | DRAMP01246 | Japonicin-2 (Frogs, amphibians, animals) | Antibacterial, Anti-Gram+, Anti-Gram-, Antimicrobial |
| 61 | 142 | DRAMP01249 | Dybowskin-2 (Frogs, amphibians, animals) | Antibacterial, Anti-Gram+, Anti-Gram-, Antimicrobial |
| 62 | 143 | DRAMP01250 | Dybowskin-3 (Frogs, amphibians, animals) | Antibacterial, Anti-Gram+, Anti-Gram-, Antimicrobial |
| 63 | 144 | DRAMP01251 | Dybowskin-4 (Frogs, amphibians, animals) | Antibacterial, Antifungal, Anti-Gram+, Anti-Gram-, Antimicrobial |
| 64 | 145 | DRAMP01252 | Dybowskin-5 (Frogs, amphibians, animals) | Antibacterial, Antifungal, Anti-Gram+, Anti-Gram-, Antimicrobial |
| 65 | 147 | DRAMP01254 | Dybowskin-1CDYa (Frogs, amphibians, animals) | Antibacterial, Anti-Gram+, Anti-Gram-, Antimicrobial |
| 66 | 148 | DRAMP01255 | Dybowskin-2CDYa (Chensinin-1; Frogs, amphibians, animals) | Antibacterial, Anti-Gram+, Anti-Gram-, Antimicrobial |
| 67 | 149 | DRAMP01257 | Dermadistinctin-K (DD K; Frogs, amphibians, animals) | Antibacterial, Antifungal, Antiprotozoal, Anti-Gram+, Anti-Gram-, Antimicrobial |
| 68 | 150 | DRAMP01258 | Dermadistinctin-L (DD L; Frogs, amphibians, animals) | Antibacterial, Antifungal, Antiprotozoal, Anti-Gram+, Anti-Gram-, Antimicrobial |
| 69 | 151 | DRAMP01259 | Dermadistinctin-M (DD M; Frogs, amphibians, animals) | Antibacterial, Anti-Gram+, Anti-Gram-, Antimicrobial |
| 70 | 152 | DRAMP01260 | Dermadistinctin-Q1 (DD Q1; Frogs, amphibians, animals) | Antibacterial, Anti-Gram+, Anti-Gram-, Antimicrobial |
| 71 | 153 | DRAMP01261 | Dermadistinctin-Q2 (DD Q2; Frogs, amphibians, animals) | Antibacterial, Anti-Gram+, Anti-Gram-, Antimicrobial |
| 72 | 155 | DRAMP01301 | Phylloseptin-1 (PS-1; Frogs, amphibians, animals) | Antibacterial, Antifungal, Antiprotozoal, Anti-Gram+, Anti-Gram-, Antimicrobial |
| 73 | 156 | DRAMP01302 | Phylloseptin-2 (PS-2; Frogs, amphibians, animals) | Antibacterial, Antifungal, Anti-Gram+, Anti-Gram-, Antimicrobial |
| 74 | 157 | DRAMP01303 | Phylloseptin-3 (PS-3; Frogs, amphibians, animals) | Antibacterial, Antifungal, Anti-Gram+, Anti-Gram-, Antimicrobial |
| 75 | 158 | DRAMP01305 | Phylloseptin-7 (PS-7; Frogs, amphibians, animals) | Antibacterial, Anti-Gram+, Anti-Gram-, Antimicrobial |
| 76 | 159 | DRAMP01306 | Phylloseptin-7 (PS-7; Frogs, amphibians, animals) | Antibacterial, Anti-Gram+, Anti-Gram-, Antimicrobial |
| 77 | 161 | DRAMP01319 | Cathelicidin-AL (Gly-rich; Frogs, amphibians, animals) | Antibacterial, Antifungal, Anti-Gram+, Anti-Gram-, Antimicrobial |
| 78 | 163 | DRAMP01339 | Amolopin-2a (Frogs, amphibians, animals) | Antibacterial, Antifungal, Anti-Gram+, Anti-Gram-, Antimicrobial |
| 79 | 164 | DRAMP01341 | Amolopin-1b (Frogs, amphibians, animals) | Antibacterial, Antifungal, Anti-Gram+, Anti-Gram-, Antimicrobial |
| 80 | 165 | DRAMP01346 | Prepromelittin-related peptide (Frogs, amphibians, animals) | Antibacterial, Antifungal, Anti-Gram+, Anti-Gram-, Antimicrobial |
| 81 | 166 | DRAMP01347 | Prepromelittin-related peptide (Frogs, amphibians, animals) | Antibacterial, Antifungal, Anti-Gram+, Anti-Gram-, Antimicrobial |
| 82 | 167 | DRAMP01350 | Tigerinin-1 (Frogs, amphibians, animals) | Antibacterial, Antifungal, Anti-Gram+, Anti-Gram-, Antimicrobial |
| 83 | 168 | DRAMP01351 | Tigerinin-2 (Frogs, amphibians, animals) | Antibacterial, Antifungal, Anti-Gram+, Anti-Gram-, Antimicrobial |
| 84 | 169 | DRAMP01352 | Tigerinin-3 (Frogs, amphibians, animals) | Antibacterial, Antifungal, Anti-Gram+, Anti-Gram-, Antimicrobial |
| 85 | 170 | DRAMP01353 | Tigerinin-4 (Frogs, amphibians, animals) | Antibacterial, Antifungal, Anti-Gram+, Anti-Gram-, Antimicrobial |
| 86 | 172 | DRAMP01355 | Ranalexin (Frogs, amphibians, animals) | Antibacterial, Antifungal, Anti-Gram+, Anti-Gram-, Antimicrobial |
| 87 | 173 | DRAMP01393 | Odorranain-W1 (OdW1; Frogs, amphibians, animals) | Antimicrobial, Antibacterial, Antifungal, Anti-Gram+, Anti-Gram-, |
| 88 | 174 | DRAMP01358 | Ranalexin-Vb (Frogs, amphibians, animals) | Antibacterial, Anti-Gram+, Anti-Gram-, Antimicrobial |
| 89 | 175 | DRAMP01359 | Ranalexin-1G (Frogs, amphibians, animals) | Antibacterial, Anti-Gram+, Anti-Gram-, Antimicrobial |
| 90 | 178 | DRAMP01362 | Frenatin-3 (Frogs, amphibians, animals) | Antibacterial, Anti-Gram+, Anti-Gram-, Antimicrobial |
| 91 | 179 | DRAMP01364 | Maculatin-1.1 (Frogs, amphibians, animals) | Antibacterial, Antifungal, Antiviral, Anti-Gram+, Anti-Gram-, Antimicrobial |
| 92 | 182 | DRAMP01370 | Oh-defensin (O. hainana defensin; spiders, animals) | Antibacterial, Antifungal, Anti-Gram+, Anti-Gram-, Antimicrobial |
| 93 | 183 | DRAMP01371 | Odorranain-NR (Frogs, amphibians, animals) | Antibacterial, Antifungal, Anti-Gram+, Anti-Gram-, Antimicrobial |
| 94 | 184 | DRAMP00931 | Antimicrobial peptide 3 (Cn-AMP3; Plant defensin) | Antibacterial, Anti-Gram+, Anti-Gram-, Antimicrobial |
| 95 | 185 | DRAMP00930 | Antimicrobial peptide 2 (Cn-AMP2; Plant defensin) | Antibacterial, Anti-Gram+, Anti-Gram-, Antimicrobial |
| 96 | 186 | DRAMP00929 | Antimicrobial peptide 1 (Cn-AMP1; Plant defensin) | Antibacterial, Anti-Gram+, Anti-Gram-, Antimicrobial |
| 97 | 187 | DRAMP03542 | Neurokinin A (NKA; chicken, animals) | Neuropeptide,Antibacterial, Anti-Gram+, Anti-Gram-, Antimicrobial |
| 98 | 188 | DRAMP04532 | Myxinidin (Hagfish, animals) | Antimicrobial, Antibacterial, Antifungal, Anti-Gram+, Anti-Gram-, |
| 99 | 190 | DRAMP02993 | Abaecin (Pro-rich; insects, arthropods, invertebrates, animals) | Antibacterial, Anti-Gram+, Anti-Gram-, Antimicrobial |
| 100 | 191 | DRAMP02997 | Apidaecin-1B (Apidaecin IB; Insects, animals) | Antibacterial, Anti-Gram+, Anti-Gram-, Antimicrobial |
| 101 | 193 | DRAMP02840 | Lactoferricin B (Lfcin B; mammals, animals) | Antibacterial, Anti-Gram+, Anti-Gram-, Antimicrobial |
| 102 | 198 | DRAMP02246 | Ranatuerin-1C (Ranatuerin 1C; Frogs, amphibians, animals) | Antifungal, Anti-Gram+, Anti-Gram-, |
| 103 | 199 | DRAMP01394 | Odorranain-W2 (Frogs, amphibians, animals) | Antibacterial, Antifungal, Anti-Gram+, Anti-Gram-, Antimicrobial |
| 104 | 200 | DRAMP01395 | Odorranain-A-OA1 (Frogs, amphibians, animals) | Antibacterial, Antifungal, Anti-Gram+, Anti-Gram-, Antimicrobial |
| 105 | 201 | DRAMP01396 | Odorranain-F-OA1 (Frogs, amphibians, animals) | Antibacterial, Antifungal , Anti-Gram+, Anti-Gram-, Antimicrobial |
| 106 | 202 | DRAMP01397 | Odorranain-F-OA2 (Frogs, amphibians, animals) | Antibacterial, Antifungal , Anti-Gram+, Anti-Gram-, Antimicrobial |

| S.no. | PepID | DRAMP_ID | Name of the AMP | Activity |
|---|---|---|---|---|
| 107 | 203 | DRAMP01398 | Odorranain-F-OA3 (Frogs, amphibians, animals) | Antibacterial, Antifungal , Anti-Gram+, Anti-Gram-, Antimicrobial |
| 108 | 204 | DRAMP01399 | Odorranain-F-OA4 (Frogs, amphibians, animals) | Antibacterial, Antifungal , Anti-Gram+, Anti-Gram-, Antimicrobial |
| 109 | 206 | DRAMP01401 | Odorranain-F-OW1 (Frogs, amphibians, animals) | Antibacterial, Antifungal , Anti-Gram+, Anti-Gram-, Antimicrobial |
| 110 | 207 | DRAMP01402 | Odorranain-J-OA1 (Frogs, amphibians, animals) | Antibacterial, Antifungal , Anti-Gram+, Anti-Gram-, Antimicrobial |
| 111 | 208 | DRAMP01403 | Odorranain-J-OA2 (Frogs, amphibians, animals) | Antibacterial, Antifungal , Anti-Gram+, Anti-Gram-, Antimicrobial |
| 112 | 209 | DRAMP01409 | Nigrocin-OR1 (Frogs, amphibians, animals) | Antibacterial, Antifungal, Anti-Gram+, Anti-Gram-, Antimicrobial |
| 113 | 210 | DRAMP01410 | Nigrocin-OR2 (Frogs, amphibians, animals) | Antibacterial, Antifungal, Anti-Gram+, Anti-Gram-, Antimicrobial |
| 114 | 211 | DRAMP01411 | Nigrocin-OR3 (Frogs, amphibians, animals) | Antibacterial, Antifungal, Anti-Gram+, Anti-Gram-, Antimicrobial |
| 115 | 212 | DRAMP01412 | Nigrocin-2HSa (Frogs, amphibians, animals) | Antibacterial, Anti-Gram+, Anti-Gram-, Antimicrobial |
| 116 | 213 | DRAMP01413 | Nigrocin-2HSb (Frogs, amphibians, animals) | Antibacterial, Anti-Gram+, Anti-Gram-, Antimicrobial |
| 117 | 214 | DRAMP01414 | Nigrocin-2ISa (Frogs, amphibians, animals) | Antibacterial, Antifungal, Anti-Gram+, Anti-Gram-, Antimicrobial |
| 118 | 215 | DRAMP01415 | Nigrocin-2ISb (Frogs, amphibians, animals) | Antibacterial, Antifungal, Anti-Gram+, Anti-Gram-, Antimicrobial |
| 119 | 216 | DRAMP01416 | Nigrocin-2ISc (Frogs, amphibians, animals) | Antibacterial, Antifungal, Anti-Gram+, Anti-Gram-, Antimicrobial |
| 120 | 218 | DRAMP01418 | Nigrocin-2GRb (Frogs, amphibians, animals) | Antibacterial, Antifungal, Anti-Gram+, Anti-Gram-, Antimicrobial |
| 121 | 220 | DRAMP01420 | Nigrocin-OG4 (Frogs, amphibians, animals) | Antibacterial, Antifungal , Anti-Gram+, Anti-Gram-, Antimicrobial |
| 122 | 221 | DRAMP01421 | Nigrocin-OG5 (Frogs, amphibians, animals) | Antibacterial, Antifungal , Anti-Gram+, Anti-Gram-, Antimicrobial |
| 123 | 222 | DRAMP01422 | Nigrosin-OG21 (Frogs, amphibians, animals) | Antibacterial, Antifungal , Anti-Gram+, Anti-Gram-, Antimicrobial |
| 124 | 223 | DRAMP01423 | Nigrosin-OG13 (Frogs, amphibians, animals) | Antibacterial, Antifungal , Anti-Gram+, Anti-Gram-, Antimicrobial |
| 125 | 224 | DRAMP01426 | Nigrocin-1-OA1 (Frogs, amphibians, animals) | Antibacterial, Antifungal, Anti-Gram+, Anti-Gram-, Antimicrobial |
| 126 | 225 | DRAMP01427 | Nigrocin-1-OA2 (Frogs, amphibians, animals) | Antibacterial, Antifungal, Anti-Gram+, Anti-Gram-, Antimicrobial |
| 127 | 226 | DRAMP01428 | Nigrocin-1-OA3 (Frogs, amphibians, animals) | Antibacterial, Antifungal, Anti-Gram+, Anti-Gram-, Antimicrobial |
| 128 | 227 | DRAMP01429 | Nigrocin-1-OR1 (Frogs, amphibians, animals) | Antibacterial, Antifungal, Anti-Gram+, Anti-Gram-, Antimicrobial |
| 129 | 228 | DRAMP01430 | Nigrocin-1-OR2 (Frogs, amphibians, animals) | Antibacterial, Antifungal, Anti-Gram+, Anti-Gram-, Antimicrobial |
| 130 | 229 | DRAMP01431 | Nigrocin-1-OR3 (Frogs, amphibians, animals) | Antibacterial, Antifungal, Anti-Gram+, Anti-Gram-, Antimicrobial |
| 131 | 230 | DRAMP01432 | Nigrocin-1-OW2 (Frogs, amphibians, animals) | Antibacterial, Antifungal, Anti-Gram+, Anti-Gram-, Antimicrobial |
| 132 | 231 | DRAMP01433 | Nigrocin-1-OW3 (Frogs, amphibians, animals) | Antibacterial, Antifungal, Anti-Gram+, Anti-Gram-, Antimicrobial |
| 133 | 232 | DRAMP01434 | Nigrocin-1-OW4 (Frogs, amphibians, animals) | Antibacterial, Antifungal, Anti-Gram+, Anti-Gram-, Antimicrobial |
| 134 | 233 | DRAMP01435 | Nigrocin-1-OW5 (Frogs, amphibians, animals) | Antibacterial, Antifungal, Anti-Gram+, Anti-Gram-, Antimicrobial |
| 135 | 234 | DRAMP01436 | Nigrocin-1-OW1 (Frogs, amphibians, animals) | Antibacterial, Antifungal, Anti-Gram+, Anti-Gram-, Antimicrobial |
| 136 | 236 | DRAMP01438 | Nigrocin-2JDa (Frogs, amphibians, animals) | Antibacterial, Antifungal , Anti-Gram+, Anti-Gram-, Antimicrobial |
| 137 | 237 | DRAMP01439 | Nigrocin-2JDb (Odorranain-H2; Frogs, amphibians, animals) | Antibacterial, Antifungal , Anti-Gram+, Anti-Gram-, Antimicrobial |
| 138 | 238 | DRAMP01440 | Nigrocin-2LVb (Frogs, amphibians, animals) | Antibacterial, Anti-Gram+, Anti-Gram-, Antimicrobial |
| 139 | 242 | DRAMP01447 | Esculentin-2CHa (Frogs, amphibians, animals) | Antibacterial, Antifungal, Anti-Gram+, Anti-Gram-, Antimicrobial |
| 140 | 243 | DRAMP01452 | Esculentin-1LTa (Frogs, amphibians, animals) | Antibacterial, Antifungal, Anti-Gram+, Anti-Gram-, Antimicrobial |
| 141 | 244 | DRAMP01453 | Esculentin-2LTa (Frogs, amphibians, animals) | Antibacterial, Anti-Gram+, Anti-Gram-, Antimicrobial |
| 142 | 245 | DRAMP01454 | Esculentin-2JDa (Frogs, amphibians, animals) | Antibacterial, Anti-Gram+, Anti-Gram-, Antimicrobial |
| 143 | 246 | DRAMP01456 | Esculentin-2PLa (Frogs, amphibians, animals) | Antibacterial, Antifungal, Anti-Gram+, Anti-Gram-, Antimicrobial |
| 144 | 247 | DRAMP01457 | Esculentin-1V (Frogs, amphibians, animals) | Antibacterial, Anti-Gram+, Anti-Gram-, Antimicrobial |
| 145 | 248 | DRAMP01458 | Esculentin-2V (Frogs, amphibians, animals) | Antibacterial, Anti-Gram+, Anti-Gram-, Antimicrobial |
| 146 | 249 | DRAMP01461 | Esculentin-1S (Frogs, amphibians, animals) | Antibacterial, Anti-Gram+, Anti-Gram-, Antimicrobial |
| 147 | 250 | DRAMP01462 | Esculentin-2S (Frogs, amphibians, animals) | Antibacterial , Anti-Gram+, Anti-Gram-, Antimicrobial |
| 148 | 251 | DRAMP01469 | Esculentin-2-Ala (Frogs, amphibians, animals) | Antibacterial, Anti-Gram+, Anti-Gram-, Antimicrobial |
| 149 | 252 | DRAMP01470 | Esculentin-2-ALb (Frogs, amphibians, animals) | Antibacterial, Anti-Gram+, Anti-Gram-, Antimicrobial |
| 150 | 253 | DRAMP01471 | Esculentin-1PLa (Frogs, amphibians, animals) | Antibacterial, Anti-Gram+, Anti-Gram-, Antimicrobial |
| 151 | 254 | DRAMP01472 | Esculentin-1PLb (Frogs, amphibians, animals) | Antibacterial, Anti-Gram+, Anti-Gram-, Antimicrobial |
| 152 | 256 | DRAMP01474 | Esculentin-1ARa (Frogs, amphibians, animals) | Antibacterial, Anti-Gram+, Anti-Gram-, Antimicrobial |
| 153 | 257 | DRAMP01475 | Esculentin-1ARb (Frogs, amphibians, animals) | Antibacterial, Anti-Gram+, Anti-Gram-, Antimicrobial |
| 154 | 258 | DRAMP01476 | Esculentin-2HSa (Frogs, amphibians, animals) | Antibacterial, Anti-Gram+, Anti-Gram-, Antimicrobial |
| 155 | 259 | DRAMP01477 | Esculentin-1HSa (Frogs, amphibians, animals) | Antibacterial, Anti-Gram+, Anti-Gram-, Antimicrobial |
| 156 | 260 | DRAMP01479 | Esculentin-1CPa (Frogs, amphibians, animals) | Antibacterial, Antifungal, Anti-Gram+, Anti-Gram-, Antimicrobial |
| 157 | 261 | DRAMP01480 | Esculentin-2CPa (Frogs, amphibians, animals) | Antibacterial, Antifungal, Anti-Gram+, Anti-Gram-, Antimicrobial |
| 158 | 262 | DRAMP01482 | Esculentin-1ISa (Frogs, amphibians, animals) | Antibacterial, Anti-Gram+, Anti-Gram-, Antimicrobial |
| 159 | 263 | DRAMP01483 | Esculentin-1ISb (Frogs, amphibians, animals) | Antibacterial, Antifungal, Anti-Gram+, Anti-Gram-, Antimicrobial |

| S.no. | PepID | DRAMP_ID | Name of the AMP | Activity |
|---|---|---|---|---|
| 160 | 264 | DRAMP01484 | Esculentin-2ISa (Frogs, amphibians, animals) | Antibacterial, Antifungal, Anti-Gram+, Anti-Gram-, Antimicrobial |
| 161 | 265 | DRAMP01486 | Esculentin-1GRa (Frogs, amphibians, animals) | Antibacterial, Antifungal, Anti-Gram+, Anti-Gram-, Antimicrobial |
| 162 | 266 | DRAMP01490 | Esculentin-2A (Frogs, amphibians, animals) | Antibacterial, Anti-Gram+, Anti-Gram-, Antimicrobial |
| 163 | 267 | DRAMP01491 | Esculentin-1B (Frogs, amphibians, animals) | Antibacterial, Anti-Gram+, Anti-Gram-, Antimicrobial |
| 164 | 268 | DRAMP01493 | Esculentin-1-OA1 (Frogs, amphibians, animals) | Antibacterial, Antifungal, Anti-Gram+, Anti-Gram-, Antimicrobial |
| 165 | 269 | DRAMP01494 | Esculentin-1-OA2 (Frogs, amphibians, animals) | Antibacterial, Antifungal, Anti-Gram+, Anti-Gram-, Antimicrobial |
| 166 | 270 | DRAMP01495 | Esculentin-1-OA3 (Frogs, amphibians, animals) | Antibacterial, Antifungal, Anti-Gram+, Anti-Gram-, Antimicrobial |
| 167 | 271 | DRAMP01496 | Esculentin-1-OA4 (Frogs, amphibians, animals) | Antibacterial, Antifungal, Anti-Gram+, Anti-Gram-, Antimicrobial |
| 168 | 272 | DRAMP01497 | Esculentin-1-OA5 (Frogs, amphibians, animals) | Antibacterial, Antifungal, Anti-Gram+, Anti-Gram-, Antimicrobial |
| 169 | 273 | DRAMP01499 | Esculentin-1-OR1 (Frogs, amphibians, animals) | Antibacterial, Antifungal, Anti-Gram+, Anti-Gram-, Antimicrobial |
| 170 | 274 | DRAMP01501 | Esculentin-1-OR3 (Frogs, amphibians, animals) | Antibacterial, Antifungal, Anti-Gram+, Anti-Gram-, Antimicrobial |
| 171 | 275 | DRAMP01502 | Esculentin-1-OR4 (Frogs, amphibians, animals) | Antibacterial, Antifungal, Anti-Gram+, Anti-Gram-, Antimicrobial |
| 172 | 276 | DRAMP01503 | Esculentin-1-OR5 (Frogs, amphibians, animals) | Antibacterial, Antifungal, Anti-Gram+, Anti-Gram-, Antimicrobial |
| 173 | 277 | DRAMP01504 | Esculentin-2-OA1 (Frogs, amphibians, animals) | Antibacterial, Antifungal, Anti-Gram+, Anti-Gram-, Antimicrobial |
| 174 | 278 | DRAMP01505 | Esculentin-2-OA2 (Frogs, amphibians, animals) | Antibacterial, Antifungal, Anti-Gram+, Anti-Gram-, Antimicrobial |
| 175 | 280 | DRAMP01507 | Esculentin-2-OR1 (Frogs, amphibians, animals) | Antibacterial, Antifungal, Anti-Gram+, Anti-Gram-, Antimicrobial |
| 176 | 281 | DRAMP01508 | Esculentin-2-OR2 (Frogs, amphibians, animals) | Antibacterial, Antifungal, Anti-Gram+, Anti-Gram-, Antimicrobial |
| 177 | 282 | DRAMP01509 | Esculentin-2-OR3 (Frogs, amphibians, animals) | Antibacterial, Antifungal, Anti-Gram+, Anti-Gram-, Antimicrobial |
| 178 | 283 | DRAMP01510 | Esculentin-2-OR4 (Frogs, amphibians, animals) | Antibacterial, Antifungal, Anti-Gram+, Anti-Gram-, Antimicrobial |
| 179 | 284 | DRAMP01511 | Esculentin-2-OR5 (Frogs, amphibians, animals) | Antibacterial, Antifungal, Anti-Gram+, Anti-Gram-, Antimicrobial |
| 180 | 285 | DRAMP01513 | Esculentin-1 (Frogs, amphibians, animals) | Antibacterial, Antifungal, Anti-Gram+, Anti-Gram-, Antimicrobial |
| 181 | 286 | DRAMP01520 | Rugosin-A (Frogs, amphibians, animals) | Antibacterial, Anti-Gram+, Anti-Gram-, Antimicrobial |
| 182 | 287 | DRAMP01521 | Rugosin-B (Frogs, amphibians, animals) | Antibacterial, Anti-Gram+, Anti-Gram-, Antimicrobial |
| 183 | 288 | DRAMP01524 | Rugosin-RN1 (Frogs, amphibians, animals) | Antibacterial, Antifungal, Anti-Gram+, Anti-Gram-, Antimicrobial |
| 184 | 289 | DRAMP01525 | Rugosin-RN3 (Frogs, amphibians, animals) | Antibacterial, Antifungal, Anti-Gram+, Anti-Gram-, Antimicrobial |
| 185 | 290 | DRAMP01526 | Rugosin-RN5 (Frogs, amphibians, animals) | Antibacterial, Antifungal, Anti-Gram+, Anti-Gram-, Antimicrobial |
| 186 | 291 | DRAMP01533 | Nigroain-B1 (Frogs, amphibians, animals) | Antibacterial, Anti-Gram+, Anti-Gram-, Antimicrobial |
| 187 | 292 | DRAMP01539 | Nigroain-C2 (Frogs, amphibians, animals) | Antibacterial, Antifungal, Anti-Gram+, Anti-Gram-, Antimicrobial |
| 188 | 295 | DRAMP01546 | Nigroain-K1 (Frogs, amphibians, animals) | Antibacterial, Antifungal, Anti-Gram+, Anti-Gram-, Antimicrobial |
| 189 | 297 | DRAMP01549 | Caerin-1.1 (Frogs, amphibians, animals) | Antibacterial, Antiviral, Anti-Gram+, Anti-Gram-, Antimicrobial |
| 190 | 298 | DRAMP01550 | Caerin-1.11 (Frogs, amphibians, animals) | Antibacterial, Anti-Gram+, Anti-Gram-, Antimicrobial |
| 191 | 299 | DRAMP01552 | Caerin-1.3 (Frogs, amphibians, animals) | Antibacterial, Anti-Gram+, Anti-Gram-, Antimicrobial |
| 192 | 300 | DRAMP01553 | Caerin-1.4 (Frogs, amphibians, animals) | Antibacterial, Anti-Gram+, Anti-Gram-, Antimicrobial |
| 193 | 301 | DRAMP01555 | Caerin-1.5 (Frogs, amphibians, animals) | Antibacterial, Anti-Gram+, Anti-Gram-, Antimicrobial |
| 194 | 302 | DRAMP01560 | Caerin-1.9 (Frogs, amphibians, animals) | Antibacterial, Antifungal, Antiviral, Anti-Gram+, Anti-Gram-, Antimicrobial |
| 195 | 304 | DRAMP01563 | Caerin-2.2 (Frogs, amphibians, animals) | Antibacterial , Anti-Gram+, Anti-Gram-, Antimicrobial |
| 196 | 309 | DRAMP01574 | Caerin-4.1 (Frogs, amphibians, animals) | Antibacterial, Antiviral, Anti-Gram+, Anti-Gram-, Antimicrobial |
| 197 | 310 | DRAMP01576 | Caerin-4.3 (Frogs, amphibians, animals) | Antibacterial, Anti-Gram+, Anti-Gram-, Antimicrobial |
| 198 | 311 | DRAMP01577 | Caerin-1.10 (Frogs, amphibians, animals) | Antibacterial, Anti-Gram+, Anti-Gram-, Antimicrobial |
| 199 | 316 | DRAMP01585 | Caerin-1.18 (Frogs, amphibians, animals) | Antibacterial, Anti-Gram+, Anti-Gram-, Antimicrobial |
| 200 | 317 | DRAMP01586 | Caerin-1.19 (Frogs, amphibians, animals) | Antibacterial, Anti-Gram+, Anti-Gram-, Antimicrobial |
| 201 | 321 | DRAMP01590 | Citropin 1.1 M14 (Frogs, amphibians, animals) | Antibacterial, Anti-Gram+, Anti-Gram-, Antimicrobial |
| 202 | 322 | DRAMP01591 | Citropin 1.1 M15 (Frogs, amphibians, animals) | Antibacterial, Anti-Gram+, Anti-Gram-, Antimicrobial |
| 203 | 329 | DRAMP01607 | Aurein-1.2 (Frogs, amphibians, animals) | Antibacterial, Anticancer, Anti-Gram+, Anti-Gram-, Antimicrobial |
| 204 | 335 | DRAMP01618 | Aurein-3.3 (Frogs, amphibians, animals) | Antibacterial, Anticancer, Anti-Gram+, Anti-Gram-, Antimicrobial |
| 205 | 337 | DRAMP01621 | Bombinin-H1 (Frogs, amphibians, animals) | Antibacterial, Anti-Gram+, Anti-Gram-, Antimicrobial |
| 206 | 338 | DRAMP01623 | Bombinin-H4 (bombinin H isomers; Frogs, amphibians, animals) | Antibacterial, Anti-Gram+, Anti-Gram-, Antimicrobial |
| 207 | 339 | DRAMP01626 | Bombinin-H5 (Frogs, amphibians, animals) | Antibacterial, Anti-Gram+, Anti-Gram-, Antimicrobial |
| 208 | 340 | DRAMP01627 | Skin peptide tyrosine-tyrosine (Skin-PYY; SPYY; Frogs, amphibians, | Antibacterial, Antifungal, Anti-Gram+, Anti-Gram-, Antimicrobial |
| 209 | 341 | DRAMP01628 | Phylloxin (Frogs, amphibians, animals) | Antibacterial, Anti-Gram+, Anti-Gram-, Antimicrobial |
| 210 | 343 | DRAMP01639 | Dermaseptin-1 (DShypo01, DPh-1; Frogs, amphibians, animals) | Antibacterial, Antiprotozoal, Anti-Gram+, Anti-Gram-, Antimicrobial |
| 211 | 344 | DRAMP01643 | Dermaseptin-5 (DShypo05, DS 01; Frogs, amphibians, animals) | Antibacterial, Antiprotozoal, Anti-Gram+, Anti-Gram-, Antimicrobial |
| 212 | 345 | DRAMP01646 | Adenoregulin (Dermaseptin BII; Dermaseptin B2; Frogs, amphibians, | Antibacterial, Antifungal, Anti-Gram+, Anti-Gram-, Antimicrobial |

| S.no. | PepID | DRAMP_ID | Name of the AMP | Activity |
|---|---|---|---|---|
| 213 | 347 | DRAMP01648 | Dermaseptin-like PBN2 (DRP-PBN2; Plasticin-B1a; Frogs, | Antibacterial, Antifungal, Anti-Gram+, Anti-Gram-, Antimicrobial |
| 214 | 348 | DRAMP01649 | Dermaseptin-BI (Dermaseptin B1; Frogs, amphibians, animals) | Antibacterial, Antifungal, Anti-Gram+, Anti-Gram-, Antimicrobial |
| 215 | 349 | DRAMP01650 | Dermaseptin-B3 (Dermaseptin BIII; Frogs, amphibians, animals) | Antibacterial, Anti-Gram+, Anti-Gram-, Antimicrobial |
| 216 | 350 | DRAMP01651 | Dermaseptin-B4 (Dermaseptin BIV; Frogs, amphibians, animals) | Antibacterial, Anti-Gram+, Anti-Gram-, Antimicrobial |
| 217 | 352 | DRAMP01668 | Dermaseptin-1 (DS I; Dermaseptin-S1, DS1; Frogs, amphibians, | Antibacterial, Antifungal, Antiprotozoal, Anti-Gram+, Anti-Gram-, Antimicrobial |
| 218 | 357 | DRAMP01702 | Dermaseptin-H5 (Dermaseptin-like peptide 5, DMS5; Frogs, | Antibacterial , Anti-Gram+, Anti-Gram-, Antimicrobial |
| 219 | 359 | DRAMP01730 | Temporin-A (Frogs, amphibians, animals) | Antibacterial, Antifungal, Anti-Gram+, Anti-Gram-, Antimicrobial |
| 220 | 360 | DRAMP01731 | Temporin-ALd (Frogs, amphibians, animals) | Antibacterial, Anti-Gram+, Anti-Gram-, Antimicrobial |
| 221 | 361 | DRAMP01732 | Temporin-ALe (Frogs, amphibians, animals) | Antibacterial, Anti-Gram+, Anti-Gram-, Antimicrobial |
| 222 | 362 | DRAMP01733 | Temporin-ALf (Frogs, amphibians, animals) | Antibacterial, Anti-Gram+, Anti-Gram-, Antimicrobial |
| 223 | 363 | DRAMP01734 | Temporin-ALg (Frogs, amphibians, animals) | Antibacterial, Anti-Gram+, Anti-Gram-, Antimicrobial |
| 224 | 364 | DRAMP01735 | Temporin-ALh (Frogs, amphibians, animals) | Antibacterial, Anti-Gram+, Anti-Gram-, Antimicrobial |
| 225 | 365 | DRAMP01736 | Temporin-ALi (Frogs, amphibians, animals) | Antibacterial, Anti-Gram+, Anti-Gram-, Antimicrobial |
| 226 | 366 | DRAMP01737 | Temporin-ALj (Frogs, amphibians, animals) | Antibacterial, Anti-Gram+, Anti-Gram-, Antimicrobial |
| 227 | 367 | DRAMP01738 | Temporin-ALk (Frogs, amphibians, animals) | Antibacterial, Anti-Gram+, Anti-Gram-, Antimicrobial |
| 228 | 368 | DRAMP01739 | Temporin-B (Frogs, amphibians, animals) | Antibacterial, Antifungal, Anti-Gram+, Anti-Gram-, Antimicrobial |
| 229 | 372 | DRAMP01753 | Temporin-1CEa (Frogs, amphibians, animals) | Antibacterial, Anti-Gram+, Anti-Gram-, Antimicrobial |
| 230 | 373 | DRAMP01754 | Temporin-1CEb (Frogs, amphibians, animals) | Antibacterial, Anti-Gram+, Anti-Gram-, Antimicrobial |
| 231 | 374 | DRAMP01755 | Temporin-1TSa (Frogs, amphibians, animals) | Antibacterial, Anti-Gram+, Anti-Gram-, Antimicrobial |
| 232 | 376 | DRAMP01764 | Temporin-1TGa (Frogs, amphibians, animals) | Antibacterial, Antifungal, Anti-Gram+, Anti-Gram-, Antimicrobial |
| 233 | 377 | DRAMP01765 | Temporin-1TGb (Frogs, amphibians, animals) | Antibacterial, Anti-Gram+, Anti-Gram-, Antimicrobial |
| 234 | 378 | DRAMP01766 | Temporin-1TGc (Frogs, amphibians, animals) | Antibacterial, Anti-Gram+, Anti-Gram-, Antimicrobial |
| 235 | 381 | DRAMP01771 | Temporin-1Oa (Frogs, amphibians, animals) | Antibacterial, Anti-Gram+, Anti-Gram-, Antimicrobial |
| 236 | 383 | DRAMP01775 | Temporin-1Sa (Frogs, amphibians, animals) | Antibacterial, Anti-Gram+, Anti-Gram-, Antimicrobial |
| 237 | 384 | DRAMP01776 | Temporin-1Sb (Temporin-SHb; Frogs, amphibians, animals) | Antibacterial, Antifungal, Anti-Gram+, Anti-Gram-, Antimicrobial |
| 238 | 385 | DRAMP01777 | Temporin-1Sc (Temporin-SHc; Frogs, amphibians, animals) | Antibacterial, Antifungal, Anti-Gram+, Anti-Gram-, Antimicrobial |
| 239 | 386 | DRAMP01779 | Temporin-SHf (Frogs, amphibians, animals) | Antibacterial, Anti-Gram+, Anti-Gram-, Antimicrobial |
| 240 | 387 | DRAMP01780 | Temporin-SHa (Temporin-1Sa; Frogs, amphibians, animals) | Antibacterial, Antifungal, Anti-Gram+, Anti-Gram-, Antimicrobial |
| 241 | 390 | DRAMP01784 | Temporin-LTc (Frogs, amphibians, animals) | Antibacterial, Antiviral, Anti-Gram+, Anti-Gram-, Antimicrobial |
| 242 | 391 | DRAMP01785 | Temporin-CPa (Frogs, amphibians, animals) | Antibacterial, Anti-Gram+, Anti-Gram-, Antimicrobial |
| 243 | 392 | DRAMP01787 | Temporin-HN1 (Frogs, amphibians, animals) | Antibacterial, Antifungal, Anti-Gram+, Anti-Gram-, Antimicrobial |
| 244 | 393 | DRAMP01788 | Temporin-HN2 (Frogs, amphibians, animals) | Antibacterial, Antifungal, Anti-Gram+, Anti-Gram-, Antimicrobial |
| 245 | 394 | DRAMP01789 | Temporin-1Va (Temporin 1Va; Frogs, amphibians, animals) | Antibacterial, Antifungal, Anti-Gram+, Anti-Gram-, Antimicrobial |
| 246 | 396 | DRAMP01791 | Temporin-1Vc (Temporin 1Vc; Frogs, amphibians, animals) | Antibacterial, Anti-Gram+, Anti-Gram-, Antimicrobial |
| 247 | 397 | DRAMP01807 | Temporin-RN1 (Frogs, amphibians, animals) | Antibacterial, Antifungal, Anti-Gram+, Anti-Gram-, Antimicrobial |
| 248 | 398 | DRAMP01808 | Temporin-RN3 (Frogs, amphibians, animals) | Antibacterial, Antifungal, Anti-Gram+, Anti-Gram-, Antimicrobial |
| 249 | 399 | DRAMP01811 | Temporin-Ra (Frogs, amphibians, animals) | Antibacterial, Anti-Gram+, Anti-Gram-, Antimicrobial |
| 250 | 400 | DRAMP01812 | Temporin-Rb (Frogs, amphibians, animals) | Antibacterial, Anti-Gram+, Anti-Gram-, Antimicrobial |
| 251 | 402 | DRAMP01816 | Temporin-1CSb (Frogs, amphibians, animals) | Antibacterial, Anti-Gram+, Anti-Gram-, Antimicrobial |
| 252 | 403 | DRAMP01817 | Temporin-1CSc (Frogs, amphibians, animals) | Antibacterial, Anti-Gram+, Anti-Gram-, Antimicrobial |
| 253 | 404 | DRAMP01818 | Temporin-1CSd (Temporin-1DRb; Frogs, amphibians, animals) | Antibacterial, Antifungal, Anti-Gram+, Anti-Gram-, Antimicrobial |
| 254 | 405 | DRAMP01392 | Odorranain-V1 (OdV1; Frogs, amphibians, animals) | Antimicrobial, Antibacterial, Antifungal, Anti-Gram+, Anti-Gram-, |
| 255 | 406 | DRAMP01391 | Odorranain-U1 (OdU1; Frogs, amphibians, animals) | Antimicrobial, Antibacterial, Antifungal, Anti-Gram+, Anti- Gram-, |
| 256 | 407 | DRAMP01832 | Temporin-Eca (Frogs, amphibians, animals) | Antibacterial, Anti-Gram+, Anti-Gram-, Antimicrobial |
| 257 | 408 | DRAMP01833 | Buforin-EC (Frogs, amphibians, animals) | Antibacterial, Anti-Gram+, Anti-Gram-, Antimicrobial |
| 258 | 409 | DRAMP01834 | Cyanophlyctin (Frogs, amphibians, animals) | Antibacterial, Anti-Gram+, Anti-Gram-, Antimicrobial |
| 259 | 410 | DRAMP01840 | Ascaphin-1 (Frogs, amphibians, animals) | Antibacterial, Anti-Gram+, Anti-Gram-, Antimicrobial |
| 260 | 411 | DRAMP01842 | Ascaphin-3 (Frogs, amphibians, animals) | Antibacterial, Anti-Gram+, Anti-Gram-, Antimicrobial |
| 261 | 412 | DRAMP01844 | Ascaphin-5 (Frogs, amphibians, animals) | Antibacterial, Antifungal, Anti-Gram+, Anti-Gram-, Antimicrobial |
| 262 | 413 | DRAMP01846 | Ascaphin-7 (Frogs, amphibians, animals) | Antibacterial, Anti-Gram+, Anti-Gram-, Antimicrobial |
| 263 | 414 | DRAMP01847 | Ascaphin-8 (Frogs, amphibians, animals) | Antibacterial, Antiviral, Anti-Gram+, Anti-Gram-, Antimicrobial |
| 264 | 415 | DRAMP01849 | Jindongenin-1a (JD1a; Frogs, amphibians, animals) | Antibacterial, Antifungal, Anti-Gram+, Anti-Gram-, Antimicrobial |
| 265 | 416 | DRAMP01869 | Brevinin-1SPa (Frogs, amphibians, animals) | Antibacterial, Antifungal, Anti-Gram+, Anti-Gram-, Antimicrobial |

| S.no. | PepID | DRAMP_ID | Name of the AMP | Activity |
|---|---|---|---|---|
| 266 | 417 | DRAMP01870 | Brevinin-1SPb (Frogs, amphibians, animals) | Antibacterial, Antifungal, Anti-Gram+, Anti-Gram-, Antimicrobial |
| 267 | 418 | DRAMP01872 | Brevinin-1SPd (Frogs, amphibians, animals) | Antibacterial, Antifungal, Anti-Gram+, Anti-Gram-, Antimicrobial |
| 268 | 419 | DRAMP01873 | Brevinin-2-related peptide (Frogs, amphibians, animals) | Antibacterial, Antifungal, Anti-Gram+, Anti-Gram-, Antimicrobial |
| 269 | 420 | DRAMP01875 | Brevinin-2PRa (Frogs, amphibians, animals) | Antibacterial, Anti-Gram+, Anti-Gram-, Antimicrobial |
| 270 | 421 | DRAMP01876 | Brevinin-2PRb (Frogs, amphibians, animals) | Antibacterial, Anti-Gram+, Anti-Gram-, Antimicrobial |
| 271 | 422 | DRAMP01877 | Brevinin-2PRd (Frogs, amphibians, animals) | Antibacterial, Anti-Gram+, Anti-Gram-, Antimicrobial |
| 272 | 423 | DRAMP01878 | Brevinin-2PRe (Frogs, amphibians, animals) | Antibacterial, Anti-Gram+, Anti-Gram-, Antimicrobial |
| 273 | 424 | DRAMP01879 | Brevinin-2LTa (Frogs, amphibians, animals) | Antibacterial, Anti-Gram+, Anti-Gram-, Antimicrobial |
| 274 | 425 | DRAMP01880 | Brevinin-2LTb (Frogs, amphibians, animals) | Antibacterial, Anti-Gram+, Anti-Gram-, Antimicrobial |
| 275 | 426 | DRAMP01881 | Brevinin-2LTc (Frogs, amphibians, animals) | Antibacterial, Anti-Gram+, Anti-Gram-, Antimicrobial |
| 276 | 427 | DRAMP01885 | Brevinin-1TEa (Frogs, amphibians, animals) | Antibacterial, Anti-Gram+, Anti-Gram-, Antimicrobial |
| 277 | 428 | DRAMP01886 | Brevinin-2TEa (Frogs, amphibians, animals) | Antibacterial, Anti-Gram+, Anti-Gram-, Antimicrobial |
| 278 | 429 | DRAMP01887 | Brevinin-2TEb (Frogs, amphibians, animals) | Antibacterial, Anti-Gram+, Anti-Gram-, Antimicrobial |
| 279 | 430 | DRAMP01888 | Brevinin-1CHc (Frogs, amphibians, animals) | Antibacterial, Antifungal, Anti-Gram+, Anti-Gram-, Antimicrobial |
| 280 | 431 | DRAMP01889 | Brevinin-1TOa (Frogs, amphibians, animals) | Antibacterial, Antifungal, Anti-Gram+, Anti-Gram-, Antimicrobial |
| 281 | 432 | DRAMP01890 | Brevinin-1VLa (Frogs, amphibians, animals) | Antibacterial, Antifungal, Anti-Gram+, Anti-Gram-, Antimicrobial |
| 282 | 433 | DRAMP01891 | Brevinin-1VLc (Frogs, amphibians, animals) | Antibacterial, Antifungal, Anti-Gram+, Anti-Gram-, Antimicrobial |
| 283 | 434 | DRAMP01892 | Brevinin-1VLd (Frogs, amphibians, animals) | Antibacterial, Antifungal, Anti-Gram+, Anti-Gram-, Antimicrobial |
| 284 | 435 | DRAMP01893 | Brevinin-1VLe (Frogs, amphibians, animals) | Antibacterial, Antifungal, Anti-Gram+, Anti-Gram-, Antimicrobial |
| 285 | 436 | DRAMP01896 | Brevinin-1CG1 (Frogs, amphibians, animals) | Antibacterial, Antifungal, Anti-Gram+, Anti-Gram-, Antimicrobial |
| 286 | 437 | DRAMP01897 | Brevinin-1CG2 (Frogs, amphibians, animals) | Antibacterial, Antifungal, Anti-Gram+, Anti-Gram-, Antimicrobial |
| 287 | 438 | DRAMP01898 | Brevinin-1CG3 (Frogs, amphibians, animals) | Antibacterial, Antifungal, Anti-Gram+, Anti-Gram-, Antimicrobial |
| 288 | 439 | DRAMP01899 | Brevinin-1CG4 (Frogs, amphibians, animals) | Antibacterial, Antifungal, Anti-Gram+, Anti-Gram-, Antimicrobial |
| 289 | 440 | DRAMP01900 | Brevinin-1CG5 (Frogs, amphibians, animals) | Antibacterial, Antifungal, Anti-Gram+, Anti-Gram-, Antimicrobial |
| 290 | 442 | DRAMP01910 | Brevinin-2GHb (AMP-2; Frogs, amphibians, animals) | Antibacterial, Anti-Gram+, Anti-Gram-, Antimicrobial |
| 291 | 443 | DRAMP01911 | Brevinin-2GHc (AMP-4; Frogs, amphibians, animals) | Antibacterial, Anti-Gram+, Anti-Gram-, Antimicrobial |
| 292 | 444 | DRAMP01913 | Brevinin-1GRa (Frogs, amphibians, animals) | Antibacterial, Anti-Gram+, Anti-Gram-, Antimicrobial |
| 293 | 445 | DRAMP01914 | Brevinin-2GRa (Frogs, amphibians, animals) | Antibacterial, Antifungal, Anti-Gram+, Anti-Gram-, Antimicrobial |
| 294 | 446 | DRAMP01918 | Brevinin-1PLb (Frogs, amphibians, animals) | Antibacterial, Antifungal, Anti-Gram+, Anti-Gram-, Antimicrobial |
| 295 | 447 | DRAMP01919 | Brevinin-1PLc (Frogs, amphibians, animals) | Antibacterial, Antifungal, Anti-Gram+, Anti-Gram-, Antimicrobial |
| 296 | 448 | DRAMP01920 | Brevinin-1CSa (Frogs, amphibians, animals) | Antibacterial, Anti-Gram+, Anti-Gram-, Antimicrobial |
| 297 | 450 | DRAMP01922 | Brevinin-2SKb (Frogs, amphibians, animals) | Antibacterial, Anti-Gram+, Anti-Gram-, Antimicrobial |
| 298 | 451 | DRAMP01933 | Brevinin-2Ef (Frogs, amphibians, animals) | Antibacterial, Anti-Gram+, Anti-Gram-, Antimicrobial |
| 299 | 457 | DRAMP01940 | Brevinin-1CHa (Frogs, amphibians, animals) | Antibacterial, Antifungal, Anti-Gram+, Anti-Gram-, Antimicrobial |
| 300 | 458 | DRAMP01941 | Brevinin-1CHb (Frogs, amphibians, animals) | Antibacterial, Antifungal, Anti-Gram+, Anti-Gram-, Antimicrobial |
| 301 | 462 | DRAMP01949 | Brevinin-1HSa (Frogs, amphibians, animals) | Antibacterial, Anti-Gram+, Anti-Gram-, Antimicrobial |
| 302 | 463 | DRAMP01950 | Brevinin-1HSb (Brevinin-1JDb; Frogs, amphibians, animals) | Antibacterial, Anti-Gram+, Anti-Gram-, Antimicrobial |
| 303 | 464 | DRAMP01951 | Brevinin-1PTa (Frogs, amphibians, animals) | Antibacterial, Anti-Gram+, Anti-Gram-, Antimicrobial |
| 304 | 465 | DRAMP01953 | Brevinin-2HSa (Frogs, amphibians, animals) | Antibacterial, Anti-Gram+, Anti-Gram-, Antimicrobial |
| 305 | 466 | DRAMP01955 | Brevinin-2PTa (Frogs, amphibians, animals) | Antibacterial, Anti-Gram+, Anti-Gram-, Antimicrobial |
| 306 | 467 | DRAMP01956 | Brevinin-2PTb (Frogs, amphibians, animals) | Antibacterial, Anti-Gram+, Anti-Gram-, Antimicrobial |
| 307 | 468 | DRAMP01957 | Brevinin-2PTc (Frogs, amphibians, animals) | Antibacterial, Anti-Gram+, Anti-Gram-, Antimicrobial |
| 308 | 469 | DRAMP01959 | Brevinin-2PTe (Frogs, amphibians, animals) | Antibacterial, Anti-Gram+, Anti-Gram-, Antimicrobial |
| 309 | 470 | DRAMP01963 | Brevinin-1BLa (Frogs, amphibians, animals) | Antibacterial, Antifungal, Anti-Gram+, Anti-Gram-, Antimicrobial |
| 310 | 471 | DRAMP01965 | Brevinin-1BLc (Frogs, amphibians, animals) | Antibacterial, Antifungal, Anti-Gram+, Anti-Gram-, Antimicrobial |
| 311 | 472 | DRAMP01968 | Brevinin-1Yc (Frogs, amphibians, animals) | Antibacterial, Antifungal, Anti-Gram+, Anti-Gram-, Antimicrobial |
| 312 | 473 | DRAMP01969 | Brevinin-1Ja (Frogs, amphibians, animals) | Antibacterial, Anti-Gram+, Anti-Gram-, Antimicrobial |
| 313 | 474 | DRAMP01970 | Brevinin-1ZHa (Frogs, amphibians, animals) | Antibacterial, Antifungal, Anti-Gram+, Anti-Gram-, Antimicrobial |
| 314 | 475 | DRAMP01971 | Brevinin-1ZHb (Frogs, amphibians, animals) | Antibacterial, Antifungal, Anti-Gram+, Anti-Gram-, Antimicrobial |
| 315 | 476 | DRAMP01974 | Brevinin-2ZHa (Frogs, amphibians, animals) | Antibacterial, Antifungal, Anti-Gram+, Anti-Gram-, Antimicrobial |
| 316 | 477 | DRAMP01986 | Brevinin-2HS2 (Frogs, amphibians, animals) | Antibacterial, Antifungal, Anti-Gram+, Anti-Gram-, Antimicrobial |
| 317 | 478 | DRAMP01990 | Brevinin-1LT1 (Frogs, amphibians, animals) | Antibacterial, Anti-Gram+, Anti-Gram-, Antimicrobial |
| 318 | 479 | DRAMP01994 | Brevinin-2ISa (Frogs, amphibians, animals) | Antibacterial, Anti-Gram+, Anti-Gram-, Antimicrobial |

| S.no. | PepID | DRAMP_ID | Name of the AMP | Activity |
|---|---|---|---|---|
| 319 | 480 | DRAMP01995 | Brevinin-2ISb (Frogs, amphibians, animals) | Antibacterial, Antifungal, Anti-Gram+, Anti-Gram-, Antimicrobial |
| 320 | 481 | DRAMP01996 | Brevinin-2ISc (Frogs, amphibians, animals) | Antibacterial, Anti-Gram+, Anti-Gram-, Antimicrobial |
| 321 | 482 | DRAMP02001 | Brevinin-1HN1 (Frogs, amphibians, animals) | Antibacterial, Antifungal, Anti-Gram+, Anti-Gram-, Antimicrobial |
| 322 | 483 | DRAMP02004 | Brevinin-1V (Frogs, amphibians, animals) | Antibacterial, Antifungal, Anti-Gram+, Anti-Gram-, Antimicrobial |
| 323 | 485 | DRAMP02009 | Brevinin-1 (Frogs, amphibians, animals) | Antibacterial, Anti-Gram+, Anti-Gram-, Antimicrobial |
| 324 | 486 | DRAMP02010 | Brevinin-2 (Frogs, amphibians, animals) | Antibacterial, Anti-Gram+, Anti-Gram-, Antimicrobial |
| 325 | 487 | DRAMP02019 | Brevinin-2DYb (Frogs, amphibians, animals) | Antibacterial, Anti-Gram+, Anti-Gram-, Antimicrobial |
| 326 | 490 | DRAMP02023 | Brevinin-2DYd (Frogs, amphibians, animals) | Antibacterial, Anti-Gram+, Anti-Gram-, Antimicrobial |
| 327 | 492 | DRAMP02025 | Brevinin-2DYe (Brevinin-2CDYa; Frogs, amphibians, animals) | Antibacterial, Anti-Gram+, Anti-Gram-, Antimicrobial |
| 328 | 493 | DRAMP02026 | Brevinin-1CDYa (Frogs, amphibians, animals) | Antibacterial, Anti-Gram+, Anti-Gram-, Antimicrobial |
| 329 | 495 | DRAMP02031 | Brevinin-1Da (Frogs, amphibians, animals) | Antibacterial, Anti-Gram+, Anti-Gram-, Antimicrobial |
| 330 | 496 | DRAMP02032 | Brevinin-1TSa (Frogs, amphibians, animals) | Antibacterial, Antifungal, Anti-Gram+, Anti-Gram-, Antimicrobial |
| 331 | 497 | DRAMP02033 | Brevinin-2TSa (Frogs, amphibians, animals) | Antibacterial, Antifungal, Anti-Gram+, Anti-Gram-, Antimicrobial |
| 332 | 498 | DRAMP02034 | Brevinin-1AUa (Frogs, amphibians, animals) | Antibacterial, Antifungal, Anti-Gram+, Anti-Gram-, Antimicrobial |
| 333 | 499 | DRAMP02035 | Brevinin-1AUb (Frogs, amphibians, animals) | Antibacterial, Antifungal, Anti-Gram+, Anti-Gram-, Antimicrobial |
| 334 | 500 | DRAMP02036 | Brevinin-2-RN1 (Frogs, amphibians, animals) | Antibacterial, Antifungal, Anti-Gram+, Anti-Gram-, Antimicrobial |
| 335 | 501 | DRAMP02037 | Brevinin-2-RN2 (Frogs, amphibians, animals) | Antibacterial, Antifungal, Anti-Gram+, Anti-Gram-, Antimicrobial |
| 336 | 502 | DRAMP02038 | Brevinin-1-OA1 (Frogs, amphibians, animals) | Antibacterial, Antifungal, Anti-Gram+, Anti-Gram-, Antimicrobial |
| 337 | 504 | DRAMP02040 | Brevinin-1-OA12 (Frogs, amphibians, animals) | Antibacterial, Antifungal, Anti-Gram+, Anti-Gram-, Antimicrobial |
| 338 | 505 | DRAMP02041 | Brevinin-1-OR1 (Frogs, amphibians, animals) | Antibacterial, Antifungal, Anti-Gram+, Anti-Gram-, Antimicrobial |
| 339 | 506 | DRAMP02042 | Brevinin-1-OR3 (Frogs, amphibians, animals) | Antibacterial, Antifungal, Anti-Gram+, Anti-Gram-, Antimicrobial |
| 340 | 507 | DRAMP02043 | Brevinin-1-OR4 (Frogs, amphibians, animals) | Antibacterial, Antifungal, Anti-Gram+, Anti-Gram-, Antimicrobial |
| 341 | 508 | DRAMP02044 | Brevinin-1-OR5 (Frogs, amphibians, animals) | Antibacterial, Antifungal, Anti-Gram+, Anti-Gram-, Antimicrobial |
| 342 | 509 | DRAMP02045 | Brevinin-1-OR6 (Frogs, amphibians, animals) | Antibacterial, Antifungal, Anti-Gram+, Anti-Gram-, Antimicrobial |
| 343 | 510 | DRAMP02046 | Brevinin-1-OR7 (Frogs, amphibians, animals) | Antibacterial, Antifungal, Anti-Gram+, Anti-Gram-, Antimicrobial |
| 344 | 511 | DRAMP02047 | Brevinin-1-OR8 (Frogs, amphibians, animals) | Antibacterial, Antifungal, Anti-Gram+, Anti-Gram-, Antimicrobial |
| 345 | 512 | DRAMP02048 | Brevinin-1-OR9 (Frogs, amphibians, animals) | Antibacterial, Antifungal, Anti-Gram+, Anti-Gram-, Antimicrobial |
| 346 | 513 | DRAMP02049 | Brevinin-1-OR10 (Frogs, amphibians, animals) | Antibacterial, Antifungal, Anti-Gram+, Anti-Gram-, Antimicrobial |
| 347 | 514 | DRAMP02050 | Brevinin-1-OR11 (Frogs, amphibians, animals) | Antibacterial, Antifungal, Anti-Gram+, Anti-Gram-, Antimicrobial |
| 348 | 515 | DRAMP02051 | Lividin-1 (Brevinin-1-OR2; Frogs, amphibians, animals) | Antibacterial, Antifungal, Anti-Gram+, Anti-Gram-, Antimicrobial |
| 349 | 516 | DRAMP02052 | Lividin-2 (Brevinin-2-OR8; Frogs, amphibians, animals) | Antibacterial, Antifungal, Anti-Gram+, Anti-Gram-, Antimicrobial |
| 350 | 517 | DRAMP02053 | Lividin-3 (Brevinin-2-OR1; Frogs, amphibians, animals) | Antibacterial, Antifungal, Anti-Gram+, Anti-Gram-, Antimicrobial |
| 351 | 518 | DRAMP02054 | Brevinin-2-OA1 (Frogs, amphibians, animals) | Antibacterial, Antifungal, Anti-Gram+, Anti-Gram-, Antimicrobial |
| 352 | 519 | DRAMP02055 | Brevinin-2-OA2 (Brevinin-2E-OG1; Frogs, amphibians, animals) | Antibacterial, Antifungal, Anti-Gram+, Anti-Gram-, Antimicrobial |
| 353 | 520 | DRAMP02056 | Brevinin-2-OA3 (Frogs, amphibians, animals) | Antibacterial, Antifungal, Anti-Gram+, Anti-Gram-, Antimicrobial |
| 354 | 521 | DRAMP02057 | Brevinin-2-OA4 (Frogs, amphibians, animals) | Antibacterial, Antifungal, Anti-Gram+, Anti-Gram-, Antimicrobial |
| 355 | 522 | DRAMP02058 | Brevinin-2-OA5 (Frogs, amphibians, animals) | Antibacterial, Antifungal, Anti-Gram+, Anti-Gram-, Antimicrobial |
| 356 | 523 | DRAMP02059 | Brevinin-2-OA6 (Frogs, amphibians, animals) | Antibacterial, Antifungal, Anti-Gram+, Anti-Gram-, Antimicrobial |
| 357 | 524 | DRAMP02060 | Brevinin-2-OA7 (Frogs, amphibians, animals) | Antibacterial, Antifungal, Anti-Gram+, Anti-Gram-, Antimicrobial |
| 358 | 525 | DRAMP02061 | Brevinin-2-OA8 (Frogs, amphibians, animals) | Antibacterial, Antifungal, Anti-Gram+, Anti-Gram-, Antimicrobial |
| 359 | 526 | DRAMP02062 | Brevinin-2-OR2 (Frogs, amphibians, animals) | Antibacterial, Antifungal, Anti-Gram+, Anti-Gram-, Antimicrobial |
| 360 | 527 | DRAMP02063 | Brevinin-2-OR3 (Frogs, amphibians, animals) | Antibacterial, Antifungal, Anti-Gram+, Anti-Gram-, Antimicrobial |
| 361 | 528 | DRAMP02064 | Brevinin-2-OR4 (Frogs, amphibians, animals) | Antibacterial, Antifungal, Anti-Gram+, Anti-Gram-, Antimicrobial |
| 362 | 529 | DRAMP02065 | Brevinin-2-OR5 (Frogs, amphibians, animals) | Antibacterial, Antifungal, Anti-Gram+, Anti-Gram-, Antimicrobial |
| 363 | 530 | DRAMP02066 | Brevinin-2-OR6 (Frogs, amphibians, animals) | Antibacterial, Antifungal, Anti-Gram+, Anti-Gram-, Antimicrobial |
| 364 | 531 | DRAMP02067 | Brevinin-2-OR7 (Frogs, amphibians, animals) | Antibacterial, Antifungal, Anti-Gram+, Anti-Gram-, Antimicrobial |
| 365 | 532 | DRAMP02068 | Brevinin-2-OR9 (Frogs, amphibians, animals) | Antibacterial, Antifungal, Anti-Gram+, Anti-Gram-, Antimicrobial |
| 366 | 533 | DRAMP02069 | Brevinin-2-OR10 (Frogs, amphibians, animals) | Antibacterial, Antifungal, Anti-Gram+, Anti-Gram-, Antimicrobial |
| 367 | 534 | DRAMP02070 | Brevinin-2-OW1 (Frogs, amphibians, animals) | Antibacterial, Antifungal, Anti-Gram+, Anti-Gram-, Antimicrobial |
| 368 | 535 | DRAMP02071 | Brevinin-2-OW2 (Frogs, amphibians, animals) | Antibacterial, Antifungal, Anti-Gram+, Anti-Gram-, Antimicrobial |
| 369 | 536 | DRAMP02072 | Brevinin-2-OW3 (Frogs, amphibians, animals) | Antibacterial, Antifungal, Anti-Gram+, Anti-Gram-, Antimicrobial |
| 370 | 537 | DRAMP02073 | Brevinin-1JDa (Frogs, amphibians, animals) | Antibacterial, Antifungal, Anti-Gram+, Anti-Gram-, Antimicrobial |
| 371 | 538 | DRAMP02075 | Brevinin-1JDc (Frogs, amphibians, animals) | Antibacterial, Anti-Gram+, Anti-Gram-, Antimicrobial |

| S.no. | PepID | DRAMP_ID | Name of the AMP | Activity |
|---|---|---|---|---|
| 372 | 543 | DRAMP02078 | Brevinin-1SY (Frogs, amphibians, animals) | Antibacterial, Anti-Gram+, Anti-Gram-, Antimicrobial |
| 373 | 544 | DRAMP02081 | Brevinin-1E (Frogs, amphibians, animals) | Antibacterial, Anti-Gram+, Anti-Gram-, Antimicrobial |
| 374 | 545 | DRAMP02084 | Brevinin-2E (Frogs, amphibians, animals) | Antibacterial, Antifungal, Anti-Gram+, Anti-Gram-, Antimicrobial |
| 375 | 546 | DRAMP02101 | Brevinin-1RTa (Frogs, amphibians, animals) | Antibacterial, Anti-Gram+, Anti-Gram-, Antimicrobial |
| 376 | 547 | DRAMP02102 | Brevinin-1RTb (Frogs, amphibians, animals) | Antibacterial, Anti-Gram+, Anti-Gram-, Antimicrobial |
| 377 | 548 | DRAMP02104 | Brevinin-2RTa (Frogs, amphibians, animals) | Antibacterial, Antifungal, Anti-Gram+, Anti-Gram-, Antimicrobial |
| 378 | 549 | DRAMP02105 | Brevinin-2RTb (Frogs, amphibians, animals) | Antibacterial, Anti-Gram+, Anti-Gram-, Antimicrobial |
| 379 | 550 | DRAMP02114 | Raniseptin-1 (Rsp-1; Frogs, amphibians, animals) | Antibacterial, Anti-Gram+, Anti-Gram-, Antimicrobial |
| 380 | 551 | DRAMP02125 | Hylin-a1 (Hy-a1; Frogs, amphibians, animals) | Antibacterial, Antifungal, Anti-Gram+, Anti-Gram-, Antimicrobial |
| 381 | 554 | DRAMP02129 | Kasstasin (Frogs, amphibians, animals) | Antibacterial, Anti-Gram+, Anti-Gram-, Antimicrobial |
| 382 | 555 | DRAMP02130 | Antimicrobial peptide 1 (XT-1; Frogs, amphibians, animals) | Antibacterial, Antifungal, Anti-Gram+, Anti-Gram-, Antimicrobial |
| 383 | 556 | DRAMP02131 | Antimicrobial peptide 2 (XT-2; Frogs, amphibians, animals) | Antibacterial, Anti-Gram+, Anti-Gram-, Antimicrobial |
| 384 | 557 | DRAMP02133 | Antimicrobial peptide 4 (XT-4; Frogs, amphibians, animals) | Antibacterial, Antifungal, Anti-Gram+, Anti-Gram-, Antimicrobial |
| 385 | 558 | DRAMP02135 | Antimicrobial peptide 6 (XT-6; Frogs, amphibians, animals) | Antibacterial, Antifungal, Anti-Gram+, Anti-Gram-, Antimicrobial |
| 386 | 559 | DRAMP02136 | Antimicrobial peptide 7 (XT-7; Frogs, amphibians, animals) | Antibacterial, Antifungal, Anti-Gram+, Anti-Gram-, Antimicrobial |
| 387 | 561 | DRAMP02219 | Ranatuerin-2AUa (Frogs, amphibians, animals) | Antimicrobial, Antibacterial, Antifungal, Anti-Gram+, Anti- Gram-, |
| 388 | 568 | DRAMP02228 | Ranatuerin-1 (Frogs, amphibians, animals) | Antibacterial, Antifungal, Anti-Gram+, Anti-Gram-, Antimicrobial |
| 389 | 576 | DRAMP02237 | Ranatuerin-2Ya (Frogs, amphibians, animals) | Cytolytic,Antibacterial, Anti-Gram+, Anti-Gram-, Antimicrobial |
| 390 | 577 | DRAMP02238 | Ranatuerin-2ZHa (Frogs, amphibians, animals) | Antibacterial, Antifungal, Anti-Gram+, Anti-Gram-, Antimicrobial |
| 391 | 578 | DRAMP02239 | Ranatuerin-1Ga (Frogs, amphibians, animals) | Antibacterial, Antifungal, Anti-Gram+, Anti-Gram-, Antimicrobial |
| 392 | 579 | DRAMP02241 | Ranatuerin-2G (Frogs, amphibians, animals) | Antibacterial, Antifungal, Anti-Gram+, Anti-Gram-, Antimicrobial |
| 393 | 580 | DRAMP01390 | Odorranain-T1 (OdT1; Frogs, amphibians, animals) | Antimicrobial, Antibacterial, Antifungal, Anti-Gram+, Anti- Gram-, |
| 394 | 581 | DRAMP01389 | Odorranain-S1 (OdS1; Frogs, amphibians, animals) | Antimicrobial, Antibacterial, Antifungal, Anti-Gram+, Anti- Gram-, |
| 395 | 582 | DRAMP02251 | Ranatuerin-2CSa (Frogs, amphibians, animals) | Antibacterial, Anti-Gram+, Anti-Gram-, Antimicrobial |
| 396 | 583 | DRAMP02252 | Ranatuerin 2SKa (Frogs, amphibians, animals) | Antibacterial, Antifungal, Anti-Gram+, Anti-Gram-, Antimicrobial |
| 397 | 584 | DRAMP01108 | Maximin-2 (Toads, amphibians, animals) | Antimicrobial, Antibacterial, Antifungal, Anti-Gram+, Anti-Gram-, |
| 398 | 586 | DRAMP01107 | Maximin-1 (Toads, amphibians, animals) | Antimicrobial, Antibacterial, Antifungal, Antiviral, Anticancer, Anti-Gram+, Anti-Gram-, |
| 399 | 587 | DRAMP02268 | Xenopsin precursor fragment (XPF; Frogs, amphibians, animals) | Antibacterial, Antifungal, Anti-Gram+, Anti-Gram-, Antimicrobial |
| 400 | 588 | DRAMP02269 | Antimicrobial peptide PGQ (PGQ; Frogs, amphibians, animals) | Antibacterial, Antifungal, Anti-Gram+, Anti-Gram-, Antimicrobial |
| 401 | 589 | DRAMP02271 | Magainin-2 (Magainin II; chain of Magainins; Frogs, amphibians, | Antibacterial, Antifungal, Antiprotozoal, Anti-Gram+, Anti- Gram-, Antimicrobial |
| 402 | 590 | DRAMP02272 | PGLa (chain of PYLa/PGLa A; Frogs, amphibians, animals) | Antibacterial, Antifungal, Anti-Gram+, Anti-Gram-, Antimicrobial |
| 403 | 591 | DRAMP02273 | PGLa-H (chain of PYLa/PGLa A; Frogs, amphibians, animals) | Antibacterial, Anti-Gram+, Anti-Gram-, Antimicrobial |
| 404 | 592 | DRAMP02274 | Ranacyclin-E (Frogs, amphibians, animals) | Antibacterial, Antifungal, Anti-Gram+, Anti-Gram-, Antimicrobial |
| 405 | 593 | DRAMP02275 | Ranacyclin-T (Frogs, amphibians, animals) | Antibacterial, Antifungal, Anti-Gram+, Anti-Gram-, Antimicrobial |
| 406 | **596** | DRAMP02278 | Ranacyclin-B-RL1 (Frogs, amphibians, animals) | Antibacterial, Antifungal, Anti-Gram+, Anti-Gram-, Antimicrobial |
| 407 | 602 | DRAMP02288 | Gaegurin-RN1 (Frogs, amphibians, animals) | Antibacterial, Antifungal, Anti-Gram+, Anti-Gram-, Antimicrobial |
| 408 | 604 | DRAMP02290 | Gaegurin-RN5 (Frogs, amphibians, animals) | Antibacterial, Antifungal, Anti-Gram+, Anti-Gram-, Antimicrobial |
| 409 | 605 | DRAMP02291 | Gaegurin-1 (Gaegurin 1; GGN1; Frogs, amphibians, animals) | Antibacterial, Antifungal, Anti-Gram+, Anti-Gram-, Antimicrobial |
| 410 | 606 | DRAMP02292 | Gaegurin-2 (Gaegurin 2; GGN2; Frogs, amphibians, animals) | Antibacterial, Antifungal, Anti-Gram+, Anti-Gram-, Antimicrobial |
| 411 | 607 | DRAMP02293 | Gaegurin-3 (Gaegurin 3; GGN3; Frogs, amphibians, animals) | Antibacterial, Antifungal, Anti-Gram+, Anti-Gram-, Antimicrobial |
| 412 | 608 | DRAMP02294 | Gaegurin-4 (Gaegurin 4; GGN4; Frogs, amphibians, animals) | Antibacterial, Antifungal, Antiprotozoal, Anti-Gram+, Anti-Gram-, Antimicrobial |
| 413 | 609 | DRAMP02295 | Gaegurin-5 (Gaegurin 5; GGN5; Brevinin-1EMa; Frogs, amphibians, | Antibacterial, Antifungal, Antiprotozoal, Anti-Gram+, Anti-Gram-, Antimicrobial |
| 414 | 610 | DRAMP02296 | Gaegurin-6 (Gaegurin 6; GGN6; Frogs, amphibians, animals) | Antibacterial, Antifungal, Anti-Gram+, Anti-Gram-, Antimicrobial |
| 415 | 616 | DRAMP02314 | Hepcidin (fish, chordates, animals) | Antibacterial, Antifungal, Anti-Gram+, Anti-Gram-, Antimicrobial |
| 416 | 617 | DRAMP02315 | Chrysophsin-1 (fish, chordates, animals) | Antibacterial, Anti-Gram+, Anti-Gram-, Antimicrobial |
| 417 | 618 | DRAMP02316 | Chrysophsin-2 (fish, chordates, animals) | Antibacterial, Anti-Gram+, Anti-Gram-, Antimicrobial |
| 418 | 619 | DRAMP02317 | Chrysophsin-3 (fish, chordates, animals) | Antibacterial, Anti-Gram+, Anti-Gram-, Antimicrobial |
| 419 | 620 | DRAMP02318 | Grammistin Pp1 (Group II grammistin; fish, chordates, animals) | Antibacterial, Anti-Gram+, Anti-Gram-, Antimicrobial |
| 420 | 621 | DRAMP02320 | Grammistin PpIIb (Group II grammistin; fish, chordates, animals) | Antibacterial, Anti-Gram+, Anti-Gram-, Antimicrobial |
| 421 | 622 | DRAMP02321 | Grammistin Pp3 (Group III grammistin; fish, chordates, animals) | Antibacterial, Anti-Gram+, Anti-Gram-, Antimicrobial |
| 422 | 623 | DRAMP02324 | SAMP H1 (fish, chordates, animals) | Antibacterial, Anti-Gram+, Anti-Gram-, Antimicrobial |
| 423 | 624 | DRAMP02330 | Piscidin-1 (Pis-1; Piscidin 1; fish, chordates, animals) | Antibacterial, Antifungal, Anti-Gram+, Anti-Gram-, Antimicrobial |
| 424 | 625 | DRAMP02331 | Piscidin-2 (Pis-2; fish, chordates, animals) | Antibacterial, Antifungal, Anti-Gram+, Anti-Gram-, Antimicrobial |

| S.no. | PepID | DRAMP_ID | Name of the AMP | Activity |
|---|---|---|---|---|
| 425 | 626 | DRAMP02336 | Oncorhyncin II (Oncorhyncin 2; fish, chordates, animals) | Antibacterial, Anti-Gram+, Anti-Gram-, Antimicrobial |
| 426 | 627 | DRAMP02337 | Oncorhyncin III (Oncorhyncin 3; fish, chordates, animals) | Antibacterial, Anti-Gram+, Anti-Gram-, Antimicrobial |
| 427 | 628 | DRAMP02347 | NRC-1 (fish, chordates, animals) | Antibacterial, Antifungal, Anti-Gram+, Anti-Gram-, Antimicrobial |
| 428 | 629 | DRAMP02348 | NRC-2 (fish, chordates, animals) | Antibacterial, Antifungal, Anti-Gram+, Anti-Gram-, Antimicrobial |
| 429 | 630 | DRAMP02349 | NRC-3 (fish, chordates, animals) | Antibacterial, Antifungal, Anti-Gram+, Anti-Gram-, Antimicrobial |
| 430 | 631 | DRAMP02350 | Pleurocidin (NRC-4; fish, chordates, animals) | Antibacterial, Antifungal, Anti-Gram+, Anti-Gram-, Antimicrobial |
| 431 | 632 | DRAMP02351 | NRC-10 (fish, chordates, animals) | Antibacterial, Antifungal, Anti-Gram+, Anti-Gram-, Antimicrobial |
| 432 | 633 | DRAMP02352 | NRC-16 (fish, chordates, animals) | Antibacterial, Antifungal, Anti-Gram+, Anti-Gram-, Antimicrobial |
| 433 | 634 | DRAMP02354 | Pleurocidin-like peptide WFY (fish, chordates, animals) | Antibacterial, Antifungal, Anti-Gram+, Anti-Gram-, Antimicrobial |
| 434 | 635 | DRAMP02357 | Pleurocidin-like peptide WF3 (NRC-5; fish, chordates, animals) | Antibacterial, Antifungal, Anti-Gram+, Anti-Gram-, Antimicrobial |
| 435 | 636 | DRAMP02358 | Pleurocidin-like peptide WF4 (NRC-6; fish, chordates, animals) | Antibacterial, Antifungal, Anti-Gram+, Anti-Gram-, Antimicrobial |
| 436 | 637 | DRAMP02359 | Pleurocidin-like peptide YT2 (NRC-7; fish, chordates, animals; | Antibacterial, Antifungal, Anti-Gram+, Anti-Gram-, Antimicrobial |
| 437 | 638 | DRAMP02360 | Pleurocidin-like peptide AP1 (NRC-11; fish, chordates, animals; | Antibacterial, Antifungal, Anti-Gram+, Anti-Gram-, Antimicrobial |
| 438 | 639 | DRAMP02361 | Pleurocidin-like peptide AP2 (NRC-12; fish, chordates, animals; | Antibacterial, Antifungal, Anti-Gram+, Anti-Gram-, Antimicrobial |
| 439 | 640 | DRAMP02362 | Pleurocidin-like peptide AP3 (NRC-13; fish, chordates, animals; | Antibacterial, Antifungal, Anti-Gram+, Anti-Gram-, Antimicrobial |
| 440 | 641 | DRAMP02363 | Pleurocidin-like peptide GcSc4C5 (NRC-14; fish, chordates, animals) | Antibacterial, Antifungal, Anti-Gram+, Anti-Gram-, Antimicrobial |
| 441 | 642 | DRAMP02364 | Pleurocidin-like peptide GcSc4B7 (NRC-15; fish, chordates, animals; | Antibacterial, Antifungal, Anti-Gram+, Anti-Gram-, Antimicrobial |
| 442 | 643 | DRAMP02365 | Pleurocidin-like peptide GC3.8 (NRC-17; fish, chordates, animals; | Antibacterial, Antifungal, Anti-Gram+, Anti-Gram-, Antimicrobial |
| 443 | 644 | DRAMP02366 | Pleurocidin-like peptide GC3.2 (NRC-18; fish, chordates, animals; | Antibacterial, Antifungal, Anti-Gram+, Anti-Gram-, Antimicrobial |
| 444 | 645 | DRAMP02367 | Pleurocidin-like peptide Hb26 (NRC-19; fish, chordates, animals; | Antibacterial, Antifungal, Anti-Gram+, Anti-Gram-, Antimicrobial |
| 445 | 646 | DRAMP02368 | Pleurocidin-like peptide Hb18 (NRC-20; fish, chordates, animals; | Antibacterial, Antifungal, Anti-Gram+, Anti-Gram-, Antimicrobial |
| 446 | 648 | DRAMP02376 | Grammistin Gs 1 (Grammistin Gs F; Group I grammistin; soapfish, | Antibacterial, Anti-Gram+, Anti-Gram-, Antimicrobial |
| 447 | 649 | DRAMP02377 | Grammistin Gs 2 (Grammistin Gs G; Group I grammistin; soapfish, | Antibacterial, Anti-Gram+, Anti-Gram-, Antimicrobial |
| 448 | 650 | DRAMP02378 | Grammistin Gs A (Group III grammistin; soapfish, chordates, | Antibacterial, Anti-Gram+, Anti-Gram-, Antimicrobial |
| 449 | 651 | DRAMP02379 | Grammistin Gs B (Group II grammistin; soapfish, chordates, animals) | Antibacterial, Anti-Gram+, Anti-Gram-, Antimicrobial |
| 450 | 652 | DRAMP02380 | Grammistin Gs C (Group III grammistin; soapfish, chordates, | Antibacterial, Anti-Gram+, Anti-Gram-, Antimicrobial |
| 451 | 657 | DRAMP02390 | Astacidin 2 (crayfish, Arthropods, animals) | Antibacterial, Anti-Gram+, Anti-Gram-, Antimicrobial |
| 452 | 658 | DRAMP02391 | Hematopoietic antimicrobial peptide-37 (MgCath37; hagfishes, | Antibacterial, Antifungal, Anti-Gram+, Anti-Gram-, Antimicrobial |
| 453 | 659 | DRAMP02393 | HFIAP-1 (HFIAP-2; hagfishes, chordates, animals) | Antibacterial, Antifungal, Anti-Gram+, Anti-Gram-, Antimicrobial |
| 454 | 660 | DRAMP02394 | HFIAP-3 (hagfishes, chordates, animals) | Antibacterial, Anti-Gram+, Anti-Gram-, Antimicrobial |
| 455 | 661 | DRAMP02395 | Aurelin (jellyfish, chordates, animals) | Antibacterial, Anti-Gram+, Anti-Gram-, Antimicrobial |
| 456 | 662 | DRAMP02397 | Big defensin (RPD-1) | Anticancer, Antibacterial, Anti-Gram+, Anti-Gram-, Antimicrobial |
| 457 | 663 | DRAMP02402 | Antimicrobial peptide scolopin-1 | Antibacterial, Antifungal, Anti-Gram+, Anti-Gram-, Antimicrobial |
| 458 | 664 | DRAMP02403 | Antimicrobial peptide scolopin-2 | Antibacterial, Antifungal, Anti-Gram+, Anti-Gram-, Antimicrobial |
| 459 | 665 | DRAMP02409 | M-theraphotoxin-Gr1a (M-TRTX-Gr1a; GsMTx-4) | Antibacterial, Anti-Gram+, Anti-Gram-, Antimicrobial |
| 460 | 666 | DRAMP02410 | Antimicrobial peptide lumbricin-1 | Antibacterial, Antifungal, Anti-Gram+, Anti-Gram-, Antimicrobial |
| 461 | 667 | DRAMP02411 | Armadillidin (Glyc-rich) | Antibacterial, Anti-Gram+, Anti-Gram-, Antimicrobial |
| 462 | 668 | DRAMP02412 | Panusin (Defensin-like peptide 7, PaD7) | Antibacterial, Anti-Gram+, Anti-Gram-, Antimicrobial |
| 463 | 670 | DRAMP02421 | Arthropods, animals) | Antibacterial, Anti-Gram+, Anti-Gram-, Antimicrobial |
| 464 | 671 | DRAMP02422 | Hlsal-defensin (H. longicornis salivary gland defensin; Ticks, | Antibacterial, Anti-Gram+, Anti-Gram-, Antimicrobial |
| 465 | 672 | DRAMP02423 | HlMS-defensin (Ticks, Arthropods, animals) | Antibacterial, Antifungal, Anti-Gram+, Anti-Gram-, Antimicrobial |
| 466 | 673 | DRAMP02425 | Ixosin-B (Ticks, Arthropods, animals) | Antibacterial, Antifungal, Anti-Gram+, Anti-Gram-, Antimicrobial |
| 467 | 678 | DRAMP02432 | Antimicrobial peptide ISAMP (Ticks, Arthropods, animals) | Antibacterial, Anti-Gram+, Anti-Gram-, Antimicrobial |
| 468 | 680 | DRAMP02434 | Antimicrobial peptide lumbricin-PG (Lumbricin-PG) | Antibacterial, Anti-Gram+, Anti-Gram-, Antimicrobial |
| 469 | 683 | DRAMP02445 | Antimicrobial protein BL-A60 | Antibacterial, Anti-Gram+, Anti-Gram-, Antimicrobial |
| 470 | 684 | DRAMP02446 | Antimicrobial protein 1 (Antimicrobial protein AN5-1) | Antibacterial, Anti-Gram+, Anti-Gram-, Antimicrobial |
| 471 | 687 | DRAMP02470 | Nosiheptide (NOS; Antibiotic 9671-RP) | Antibacterial, Anti-Gram+, Anti-Gram-, Antimicrobial |
| 472 | 688 | DRAMP02473 | Cathelicidin-BF (Cathelicidin-related protein; Snakes, reptiles, | Antibacterial, Antifungal, Anti-Gram+, Anti-Gram-, Antimicrobial |
| 473 | 689 | DRAMP02474 | cathelicidin-BF15 (Snakes, reptiles, animals) | Antibacterial, Antifungal, Anti-Gram+, Anti-Gram-, Antimicrobial |
| 474 | 690 | DRAMP02478 | L-amino-acid oxidase (Bm-LAO; LAAO; LAO; Snakes, reptiles, | Antibacterial, Antiparasitic, Anti-Gram+, Anti-Gram-, Antimicrobial |
| 475 | 692 | DRAMP02522 | L-amino-acid oxidase (LAAO, LAO, Oh-LAAO; Snakes, reptiles, | Antibacterial, Anti-Gram+, Anti-Gram-, Antimicrobial |
| 476 | 693 | DRAMP02573 | Penaeidin-3a (Pen-3a; shrimps, Arthropods, animals) | Antibacterial, Antifungal, Anti-Gram+, Anti-Gram-, Antimicrobial |
| 477 | 694 | DRAMP02574 | [T8A]-Penaeidin-3a ([T8A]-Pen-3a; shrimps, Arthropods, animals) | Antibacterial, Antifungal, Anti-Gram+, Anti-Gram-, Antimicrobial |

| S.no. | PepID | DRAMP_ID | Name of the AMP | Activity |
|---|---|---|---|---|
| 478 | 697 | DRAMP02584 | Penaeidin-4a (Pen-4a; shrimps, Arthropods, animals) | Antibacterial, Antifungal, Anti-Gram+, Anti-Gram-, Antimicrobial |
| 479 | 698 | DRAMP02586 | Penaeidin-2d (Pen-2d; shrimps, Arthropods, animals) | Antibacterial, Antifungal, Anti-Gram+, Anti-Gram-, Antimicrobial |
| 480 | 700 | DRAMP02603 | Putative antimicrobial peptide A Northern Europe Heligoland | Antibacterial, Antifungal, Anti-Gram+, Anti-Gram-, Antimicrobial |
| 481 | 705 | DRAMP02740 | TBD-1 (Turtle beta-defensin 1; Reptiles, animals) | Antibacterial, Antifungal, Anti-Gram+, Anti-Gram-, Antimicrobial |
| 482 | 706 | DRAMP02768 | Pilosulin-1 (Myr b I; ants, insects, animals) | Antibacterial, Antifungal, Anti-Gram+, Anti-Gram-, Antimicrobial |
| 483 | 709 | DRAMP02777 | Rhinocerosin (Insects, animals) | Antibacterial, Anti-Gram+, Anti-Gram-, Antimicrobial |
| 484 | 710 | DRAMP02778 | Defensin (Insects, animals) | Antibacterial, Anti-Gram+, Anti-Gram-, Antimicrobial |
| 485 | 711 | DRAMP02779 | Defensin-A (Defensin A; Insects, animals) | Antibacterial , Anti-Gram+, Anti-Gram-, Antimicrobial |
| 486 | 712 | DRAMP02780 | Defensin-B (Defensin B; Insects, animals) | Antibacterial, Anti-Gram+, Anti-Gram-, Antimicrobial |
| 487 | 713 | DRAMP02802 | Paneth cell-specific alpha-defensin 1 (DEFA1; horse defensin; | Antibacterial, Antifungal, Anti-Gram+, Anti-Gram-, Antimicrobial |
| 488 | 714 | DRAMP02809 | Myticin-B (Myt B; Cys-rich; molluscas, animals) | Antibacterial, Antifungal, Anti-Gram+, Anti-Gram-, Antimicrobial |
| 489 | 715 | DRAMP02811 | Defensin MGD-1 (molluscas, animals) | Antibacterial, Anti-Gram+, Anti-Gram-, Antimicrobial |
| 490 | 717 | DRAMP02817 | Pyrrhocoricin | Antibacterial, Anti-Gram+, Anti-Gram-, Antimicrobial |
| 491 | 719 | DRAMP01381 | Odorranain-K1 (OdK1; Frogs, amphibians, animals) | Antimicrobial, Antibacterial, Antifungal, Anti-Gram+, Anti-Gram-, |
| 492 | 720 | DRAMP01383 | Odorranain-M1 (OdM1; Frogs, amphibians, animals) | Antimicrobial, Antibacterial, Antifungal, Anti-Gram+, Anti- Gram-, |
| 493 | 721 | DRAMP02841 | Lumbricin I(6-34) | Antibacterial, Antifungal, Anti-Gram+, Anti-Gram-, Antimicrobial |
| 494 | 722 | DRAMP02843 | chain a, Structure Of An Indolicidin Peptide Derivative | Antibacterial, Anti-Gram+, Anti-Gram-, Antimicrobial |
| 495 | 724 | DRAMP02845 | CP-11 (cathelicidin; mammals, animals) | Antibacterial, Anti-Gram+, Anti-Gram-, Antimicrobial |
| 496 | 726 | DRAMP02851 | Cathelicidin-1 (Bactenecin-1, Bac1; Cyclic dodecapeptide; mammals, | Antibacterial, Anti-Gram+, Anti-Gram-, Antimicrobial |
| 497 | 727 | DRAMP02854 | Cathelicidin-5 (Antibacterial peptide BMAP-28)) | Antibacterial, Antifungal , Anti-Gram+, Anti-Gram-, Antimicrobial |
| 498 | 728 | DRAMP02855 | Cathelicidin-6 (Antibacterial peptide BMAP-27) | Antibacterial, Antifungal, Anti-Gram+, Anti-Gram-, Antimicrobial |
| 499 | 730 | DRAMP02859 | Bovine Beta-defensin 2 (bBD-2; BNBD-2; BNDB-2; mammals, | Antibacterial, Anti-Gram+, Anti-Gram-, Antimicrobial |
| 500 | 731 | DRAMP02860 | Bovine Beta-defensin 3 (bBD-3; BNBD-3; BNDB-3; mammals, | Antibacterial, Anti-Gram+, Anti-Gram-, Antimicrobial |
| 501 | 732 | DRAMP02861 | Bovine Beta-defensin 4 (bBD-4; BNBD-4; BNDB-4; mammals, | Antibacterial, Anti-Gram+, Anti-Gram-, Antimicrobial |
| 502 | 734 | DRAMP02863 | Bovine Beta-defensin 6 (bBD-6; BNBD-6; BNDB-6; mammals, | Antibacterial, Anti-Gram+, Anti-Gram-, Antimicrobial |
| 503 | 735 | DRAMP02865 | Bovine Beta-defensin 8 (bBD-8; BNBD-8; BNDB-8; mammals, | Antibacterial, Anti-Gram+, Anti-Gram-, Antimicrobial |
| 504 | 736 | DRAMP02866 | Bovine Beta-defensin 9 (bBD-9; BNBD-9; BNDB-9; mammals, | Antibacterial, Anti-Gram+, Anti-Gram-, Antimicrobial |
| 505 | 737 | DRAMP02867 | Bovine Beta-defensin 10 (bBD-10; BNBD-10; BNDB-10; mammals, | Antibacterial, Anti-Gram+, Anti-Gram-, Antimicrobial |
| 506 | 738 | DRAMP02868 | Bovine Beta-defensin 11 (bBD-11; BNBD-11; BNDB-11; mammals, | Antibacterial, Anti-Gram+, Anti-Gram-, Antimicrobial |
| 507 | 739 | DRAMP02869 | Bovine Beta-defensin 12 (bBD-12; BNBD-12; BNDB-12; mammals, | Antibacterial, Anti-Gram+, Anti-Gram-, Antimicrobial |
| 508 | 740 | DRAMP02870 | Bovine Beta-defensin 13 (bBD-13; BNBD-13; BNDB-13; mammals, | Antibacterial, Anti-Gram+, Anti-Gram-, Antimicrobial |
| 509 | 741 | DRAMP02872 | Myeloid antimicrobial peptide BMAP-27 (1-18) (mammals, animals) | Antibacterial, Antifungal, Anti-Gram+, Anti-Gram-, Antimicrobial |
| 510 | 742 | DRAMP02873 | Myeloid antimicrobial peptide BMAP-28 (1-18) (mammals, animals) | Antibacterial, Antifungal, Anti-Gram+, Anti-Gram-, Antimicrobial |
| 511 | 744 | DRAMP02877 | mBMAP28 (mammals, animals) | Antibacterial, Anti-Gram+, Anti-Gram-, Antimicrobial |
| 512 | 745 | DRAMP02878 | Tracheal antimicrobial peptide (TAP; mammals, animals) | Antibacterial, Antifungal, Anti-Gram+, Anti-Gram-, Antimicrobial |
| 513 | 746 | DRAMP02903 | Bombin H7 | Antibacterial, Anti-Gram+, Anti-Gram-, Antimicrobial |
| 514 | 751 | DRAMP02922 | Canine beta-defensin (dogs, mammals, animals) | Antibacterial, Antifungal, Anti-Gram+, Anti-Gram-, Antimicrobial |
| 515 | 752 | DRAMP02923 | cBD-1 (Canine beta-defensin 1; dogs, mammals, animals) | Antibacterial, Antifungal, Anti-Gram+, Anti-Gram-, Antimicrobial |
| 516 | 755 | DRAMP02925 | Cathelicidin (dogs, mammals, animals) | Antibacterial, Antifungal, Anti-Gram+, Anti-Gram-, Antimicrobial |
| 517 | 756 | DRAMP02931 | Arasin-likeSp (crabs, Arthropods, animals) | Antibacterial, Anti-Gram+, Anti-Gram-, Antimicrobial |
| 518 | 758 | DRAMP02933 | Polyphemusin-1 (PM1; crabs, Arthropods, animals) | Antibacterial, Antifungal, Anti-Gram+, Anti-Gram-, Antimicrobial |
| 519 | 759 | DRAMP02934 | PM1-S (linear derivative of PM1) | Antibacterial, Antifungal, Anti-Gram+, Anti-Gram-, Antimicrobial |
| 520 | 761 | DRAMP02951 | PtALF6 (Portunus trituberculatus anti-lipopolysaccharide factor | Antibacterial, Antifungal, Anti-Gram+, Anti-Gram-, Antimicrobial |
| 521 | 762 | DRAMP02952 | PtALF7 (Portunus trituberculatus anti-lipopolysaccharide factor | Antibacterial, Anti-Gram+, Anti-Gram-, Antimicrobial |
| 522 | 763 | DRAMP02953 | Arasin-1 (Pro-rich, Arg-rich; crabs, Arthropods, animals) | Antibacterial, Anti-Gram+, Anti-Gram-, Antimicrobial |
| 523 | 764 | DRAMP02956 | Dolabellanin B2 | Antibacterial, Antifungal, Anti-Gram+, Anti-Gram-, Antimicrobial |
| 524 | 765 | DRAMP02959 | Antibacterial protein PR-39 (pigs, mammals, animals) | Antibacterial, Anti-Gram+, Anti-Gram-, Antimicrobial |
| 525 | 766 | DRAMP02960 | Antibacterial peptide PMAP-23 (Myeloid antibacterial peptide 23; | Antibacterial, Anti-Gram+, Anti-Gram-, Antimicrobial |
| 526 | 767 | DRAMP02961 | Antibacterial peptide PMAP-37 (Myeloid antibacterial peptide 37; | Antibacterial, Anti-Gram+, Anti-Gram-, Antimicrobial |
| 527 | 768 | DRAMP02962 | Antibacterial peptide PMAP-36 (Myeloid antibacterial peptide 36; | Antibacterial, Anti-Gram+, Anti-Gram-, Antimicrobial |
| 528 | 769 | DRAMP02963 | PMAP-36(1-20) | Antibacterial, Antifungal, Anti-Gram+, Anti-Gram-, Antimicrobial |
| 529 | 770 | DRAMP02964 | PMAP-36(1-34) | Antibacterial, Antifungal, Anti-Gram+, Anti-Gram-, Antimicrobial |
| 530 | 771 | DRAMP02965 | PMAP-36(1-35)2 | Antibacterial, Antifungal, Anti-Gram+, Anti-Gram-, Antimicrobial |

| S.no. | PepID | DRAMP_ID | Name of the AMP | Activity |
|---|---|---|---|---|
| 531 | 772 | DRAMP02966 | DBI(32-86) (pigs, mammals, animals) | Antibacterial, Anti-Gram+, Anti-Gram-, Antimicrobial |
| 532 | 773 | DRAMP02970 | Protegrin-1 (Protegrin 1; PG-1; pigs, mammals, animals) | Antibacterial, Anti-Gram+, Anti-Gram-, Antimicrobial |
| 533 | 774 | DRAMP02975 | Tritrpticin (Trp-rich; pigs, mammals, animals) | Antibacterial, Antifungal, Anti-Gram+, Anti-Gram-, Antimicrobial |
| 534 | 776 | DRAMP01376 | Odorranain-F1 (OdF1; Frogs, amphibians, animals) | Antimicrobial, Antibacterial, Antifungal, Anti-Gram+, Anti-Gram-, |
| 535 | 777 | DRAMP01377 | Odorranain-G1 (OdG1; Frogs, amphibians, animals) | Antimicrobial, Antibacterial, Antifungal, Anti-Gram+, Anti-Gram-, |
| 536 | 778 | DRAMP02995 | Hymenoptaecin (Insects, animals) | Antibacterial, Anti-Gram+, Anti-Gram-, Antimicrobial |
| 537 | 779 | DRAMP02996 | Apidaecin-2 (Apidaecin II; Insects, animals) | Antibacterial, Anti-Gram+, Anti-Gram-, Antimicrobial |
| 538 | 780 | DRAMP01378 | Odorranain-H1 (OdH1; Frogs, amphibians, animals) | Antimicrobial, Antibacterial, Antifungal, Anti-Gram+, Anti- Gram-, |
| 539 | 781 | DRAMP02998 | Apidaecin-1A (Apidaecin IA; Insects, animals) | Antibacterial, Anti-Gram+, Anti-Gram-, Antimicrobial |
| 540 | 782 | DRAMP02999 | Jellein-1 (Jelleine-I; chain of Major royal jelly protein 1; Insects, | Antibacterial, Antifungal, Anti-Gram+, Anti-Gram-, Antimicrobial |
| 541 | 783 | DRAMP03000 | Jellein-2 (Jelleine-II; chain of Major royal jelly protein 1; Insects, | Antibacterial, Antifungal, Anti-Gram+, Anti-Gram-, Antimicrobial |
| 542 | 784 | DRAMP03001 | Jellein-3 (Jelleine-III; Insects, animals) | Antibacterial, Antifungal, Anti-Gram+, Anti-Gram-, Antimicrobial |
| 543 | 785 | DRAMP03002 | Melittin (Allergen Api m 3; Allergen Api m III; Insects, animals) | Antibacterial, Antifungal, Anti-Gram+, Anti-Gram-, Antimicrobial |
| 544 | 786 | DRAMP03003 | Melectin (MEP; Insects, animals) | Antibacterial, Anti-Gram+, Anti-Gram-, Antimicrobial |
| 545 | 787 | DRAMP03007 | Osmin (Insects, animals) | Antibacterial, Antifungal, Anti-Gram+, Anti-Gram-, Antimicrobial |
| 546 | 788 | DRAMP03019 | Mastoparan PDD-B | Antibacterial, Anti-Gram+, Anti-Gram-, Antimicrobial |
| 547 | 789 | DRAMP03020 | Mastoparan PDD-A | Antibacterial, Anti-Gram+, Anti-Gram-, Antimicrobial |
| 548 | 790 | DRAMP03021 | Mastoparan PMM | Antibacterial, Anti-Gram+, Anti-Gram-, Antimicrobial |
| 549 | 791 | DRAMP03022 | Mastoparan MP | Antibacterial, Anti-Gram+, Anti-Gram-, Antimicrobial |
| 550 | 792 | DRAMP03028 | Mastoparan-1 (MP-1; Venom protein MP-1; Insects, animals) | Antibacterial, Anti-Gram+, Anti-Gram-, Antimicrobial |
| 551 | 793 | DRAMP03033 | Mastoparan-like peptide 12a (Insects, animals) | Antibacterial, Antifungal, Anti-Gram+, Anti-Gram-, Antimicrobial |
| 552 | 794 | DRAMP03034 | Mastoparan-like peptide 12b (Insects, animals) | Antibacterial, Antifungal, Anti-Gram+, Anti-Gram-, Antimicrobial |
| 553 | 795 | DRAMP03035 | Mastoparan-like peptide 12c (Insects, animals) | Antibacterial, Antifungal, Anti-Gram+, Anti-Gram-, Antimicrobial |
| 554 | 796 | DRAMP03036 | Mastoparan-like peptide 12d (Insects, animals) | Antibacterial, Antifungal, Anti-Gram+, Anti-Gram-, Antimicrobial |
| 555 | 797 | DRAMP03037 | Eumenitin (Er-12; Insects, animals) | Antibacterial, Anti-Gram+, Anti-Gram-, Antimicrobial |
| 556 | 798 | DRAMP03038 | Eumenitin-R (Insects, animals) | Antibacterial, Antifungal, Anti-Gram+, Anti-Gram-, Antimicrobial |
| 557 | 799 | DRAMP03039 | Eumenitin-F (Insects, animals) | Antibacterial, Antifungal, Anti-Gram+, Anti-Gram-, Antimicrobial |
| 558 | 800 | DRAMP03040 | Eumenine mastoparan-EF (EMP-EF; Insects, animals) | Antibacterial, Antifungal, Anti-Gram+, Anti-Gram-, Antimicrobial |
| 559 | 801 | DRAMP03041 | Eumenine mastoparan-ER (EMP-ER; Insects, animals) | Antibacterial, Antifungal, Anti-Gram+, Anti-Gram-, Antimicrobial |
| 560 | 802 | DRAMP03042 | Eumenine mastoparan-AF (EMP-AF; Af-113; Insects, animals) | Antibacterial, Anti-Gram+, Anti-Gram-, Antimicrobial |
| 561 | 803 | DRAMP03043 | Agelaia-mastoparan (Agelaia-MP; Insects, animals) | Antibacterial, Anti-Gram+, Anti-Gram-, Antimicrobial |
| 562 | 804 | DRAMP03044 | Protonectin (Agelaia-chemotactic peptide, Agelaia-CP; Insects, | Antibacterial, Anti-Gram+, Anti-Gram-, Antimicrobial |
| 563 | 805 | DRAMP03045 | Defensin-NV (Insects, animals) | Antibacterial, Antifungal, Anti-Gram+, Anti-Gram-, Antimicrobial |
| 564 | 807 | DRAMP03047 | Venom peptide 2-long (OdVP2L; analog of OdVP2; Insects, animals) | Antibacterial, Antifungal, Anti-Gram+, Anti-Gram-, Antimicrobial |
| 565 | 808 | DRAMP03050 | Dominulin-A (Insects, animals) | Antibacterial, Anti-Gram+, Anti-Gram-, Antimicrobial |
| 566 | 809 | DRAMP03051 | Dominulin-B (Insects, animals) | Antibacterial, Anti-Gram+, Anti-Gram-, Antimicrobial |
| 567 | 813 | DRAMP03055 | PP30 (Pro-rich; abaecin-like; Insects, animals) | Antibacterial, Anti-Gram+, Anti-Gram-, Antimicrobial |
| 568 | 814 | DRAMP03056 | Decoralin (Insects, animals) | Antibacterial, Antifungal, Anti-Gram+, Anti-Gram-, Antimicrobial |
| 569 | 815 | DRAMP03057 | Thanatin (Insects, animals) | Antibacterial, Antifungal, Anti-Gram+, Anti-Gram-, Antimicrobial |
| 570 | 818 | DRAMP03075 | Cecropin-D | Antibacterial, Anti-Gram+, Anti-Gram-, Antimicrobial |
| 571 | 819 | DRAMP03089 | Drosophila cecropin-A1/A2 (Insects, animals) | Antibacterial, Anti-Gram+, Anti-Gram-, Antimicrobial |
| 572 | 820 | DRAMP03090 | Drosophila cecropin B (CecB; Insects, animals) | Antibacterial, Antifungal, Anti-Gram+, Anti-Gram-, Antimicrobial |
| 573 | 821 | DRAMP03095 | Andropin (Insects, animals) | Antibacterial, Anti-Gram+, Anti-Gram-, Antimicrobial |
| 574 | 823 | DRAMP18495 | Gomesin (Gm; Spiders, arachnids, Chelicerata, arthropods, | Antimicrobial, Antibacterial, Antifungal, Antiparasitic, Antimalarial, Antic, Anti-Gram+, Anti-Gram-, |
| 575 | 825 | DRAMP03104 | Sapecin (defensins; Insects, animals) | Antibacterial, Anti-Gram+, Anti-Gram-, Antimicrobial |
| 576 | 827 | DRAMP03116 | Ceratotoxin-C (Insects, animals) | Antibacterial, Anti-Gram+, Anti-Gram-, Antimicrobial |
| 577 | 828 | DRAMP03117 | Drosophila cecropin-A1 (Insects, animals) | Antibacterial, Anti-Gram+, Anti-Gram-, Antimicrobial |
| 578 | 830 | DRAMP03138 | Cecropin-A (Insects, animals) | Antibacterial, Antifungal, Anti-Gram+, Anti-Gram-, Antimicrobial |
| 579 | 831 | DRAMP03140 | Anopheles cecropin-A amidated isoform (Insects, animals) | Antibacterial, Antifungal, Anti-Gram+, Anti-Gram-, Antimicrobial |
| 580 | 832 | DRAMP03150 | Gambicin (Insects, animals) | Antibacterial, Antifungal, Antiparasitic, Anti-Gram+, Anti- Gram-, Antimicrobial |
| 581 | 833 | DRAMP03153 | 27 kDa antibacterial protein | Antibacterial, Anti-Gram+, Anti-Gram-, Antimicrobial |
| 582 | 835 | DRAMP03166 | P15 (deer beta-defensin; ruminant, animals) | Antibacterial, Antifungal, Anti-Gram+, Anti-Gram-, Antimicrobial |
| 583 | 837 | DRAMP03173 | Arenicin-1 (Ar-1; marine polychaeta, animals) | Antibacterial, Antifungal, Cytotoxicity, Anti-Gram+, Anti- Gram-, Antimicrobial |

| S.no. | PepID | DRAMP_ID | Name of the AMP | Activity |
|---|---|---|---|---|
| 584 | 839 | DRAMP03181 | Spinigerin (Insects, animals) | Antibacterial, Antifungal, Anti-Gram+, Anti-Gram-, Antimicrobial |
| 585 | 840 | DRAMP03186 | Spheniscin-2 (Sphe-2; penguin avian beta-defensin 103b; birds, | Antibacterial, Antifungal, Anti-Gram+, Anti-Gram-, Antimicrobial |
| 586 | 841 | DRAMP03187 | Beta defensin 1(BD-1; mammals, animals) | Antibacterial, Antifungal, Anti-Gram+, Anti-Gram-, Antimicrobial |
| 587 | 845 | DRAMP03198 | Alpha-defensin PhD-4 (primates, mammals, animals) | Antibacterial, Antifungal, Anti-Gram+, Anti-Gram-, Antimicrobial |
| 588 | 846 | DRAMP03215 | Gomesin (Gm; spiders, Arthropods, animals) | Antibacterial, Antifungal, Anti-Gram+, Anti-Gram-, Antimicrobial |
| 589 | 847 | DRAMP03216 | Oxyopinin-4a (Oxt-4a; spiders, Arthropods, animals) | Antibacterial, Anti-Gram+, Anti-Gram-, Antimicrobial |
| 590 | 848 | DRAMP03217 | M-oxotoxin-Ot1a (Oxyopinin-1, Oxki1; spiders, Arthropods, animals) | Antibacterial, Insecticidal, Anti-Gram+, Anti-Gram-, Antimicrobial |
| 591 | 849 | DRAMP03222 | M-ctenitoxin-Cs1a (M-CNTX-Cs1a; Cupiennin-1a; spiders, | Antibacterial, Insecticidal, Anti-Gram+, Anti-Gram-, Antimicrobial |
| 592 | 850 | DRAMP03225 | M-ctenitoxin-Cs1d (M-CNTX-Cs1d; Cupiennin-1d; spiders, | Antibacterial, Anti-Gram+, Anti-Gram-, Antimicrobial |
| 593 | 851 | DRAMP03226 | M-zodatoxin-Lt1a (M-ZDTX-Lt1a; Latarcin-1, Ltc-1, Ltc1; spiders, | Antibacterial, Antifungal, Anti-Gram+, Anti-Gram-, Antimicrobial |
| 594 | 852 | DRAMP03227 | M-zodatoxin-Lt2a (M-ZDTX-Lt2a; Latarcin-2a, Ltc-2a, Ltc2a; spiders, | Antibacterial, Antifungal, Anti-Gram+, Anti-Gram-, Antimicrobial |
| 595 | 853 | DRAMP03229 | M-zodatoxin-Lt3a (M-ZDTX-Lt3a; Latarcin-3a, Ltc-3a; spiders, | Antibacterial, Antifungal, Anti-Gram+, Anti-Gram-, Antimicrobial |
| 596 | 854 | DRAMP03230 | M-zodatoxin-Lt3b (M-ZDTX-Lt3b; Latarcin-3b, Ltc-3b; spiders, | Antibacterial, Antifungal, Anti-Gram+, Anti-Gram-, Antimicrobial |
| 597 | 855 | DRAMP03231 | M-zodatoxin-Lt4a (M-ZDTX-Lt4a; Latarcin-4a, Ltc-4a; spiders, | Antibacterial, Antifungal, Anti-Gram+, Anti-Gram-, Antimicrobial |
| 598 | 856 | DRAMP03232 | M-zodatoxin-Lt4b (M-ZDTX-Lt4b; Latarcin-4b, Ltc-4b; spiders, | Antibacterial, Antifungal, Anti-Gram+, Anti-Gram-, Antimicrobial |
| 599 | 857 | DRAMP03233 | M-zodatoxin-Lt5a (M-ZDTX-Lt5a; Latarcin-5, Ltc-5; spiders, | Antibacterial, Antifungal, Anti-Gram+, Anti-Gram-, Antimicrobial |
| 600 | 858 | DRAMP03236 | M-zodatoxin-Lt8a (M-ZDTX-Lt8a; Cytoinsectotoxin-1a, CIT- 1a; | Antibacterial, Insecticidal, Anti-Gram+, Anti-Gram-, Antimicrobial |
| 601 | 859 | DRAMP03253 | M-lycotoxin-Ls3a (M-LCTX-Ls3a; Lycocitin-1; spiders, Arthropods, | Antibacterial, Antifungal, Anti-Gram+, Anti-Gram-, Antimicrobial |
| 602 | 860 | DRAMP03254 | M-lycotoxin-Ls3b (M-LCTX-Ls3b; Lycocitin-2; spiders, Arthropods, | Antibacterial, Antifungal, Anti-Gram+, Anti-Gram-, Antimicrobial |
| 603 | 861 | DRAMP03278 | M-lycotoxin-Hc1a (M-LCTX-Hc1a; Lycotoxin I; spiders, Arthropods, | Antibacterial, Antifungal, Anti-Gram+, Anti-Gram-, Antimicrobial |
| 604 | 862 | DRAMP03279 | M-lycotoxin-Hc2a (M-LCTX-Hc2a; Lycotoxin-2; Lycotoxin II; spiders, | Antibacterial, Antifungal, Anti-Gram+, Anti-Gram-, Antimicrobial |
| 605 | 863 | DRAMP03280 | AcAMP (A. clavatus antimicrobial peptide) | Antibacterial, Antifungal, Antiviral, Anti-Gram+, Anti-Gram-, Antimicrobial |
| 606 | 864 | DRAMP03285 | Ostricacin-1 (Beta-defensin 2; Birds, animals) | Antibacterial, Anti-Gram+, Anti-Gram-, Antimicrobial |
| 607 | 865 | DRAMP03286 | Ostricacin-2 (Beta-defensin 1; Birds, animals) | Antibacterial, Antifungal, Anti-Gram+, Anti-Gram-, Antimicrobial |
| 608 | 866 | DRAMP03287 | Ostricacin-3 (Beta-defensin 7; Birds, animals) | Antibacterial, Anti-Gram+, Anti-Gram-, Antimicrobial |
| 609 | 867 | DRAMP03288 | Ostricacin-4 (Beta-defensin 8; Birds, animals) | Antibacterial, Anti-Gram+, Anti-Gram-, Antimicrobial |
| 610 | 868 | DRAMP03311 | Stomoxyn (Insects, animals) | Antibacterial, Anti-Gram+, Anti-Gram-, Antimicrobial |
| 611 | 873 | DRAMP03405 | mCRAMP-1 (mouse cathelin-related antimicrobial peptide 1; | Antibacterial, Antifungal, Anti-Gram+, Anti-Gram-, Antimicrobial |
| 612 | 874 | DRAMP03406 | mCRAMP-2 (mouse cathelin-related antimicrobial peptide 2; | Antibacterial, Antifungal, Anti-Gram+, Anti-Gram-, Antimicrobial |
| 613 | 875 | DRAMP03419 | Neutrophil antibiotic peptide NP-1 (RatNP-1; Rodents, mammals, | Antibacterial, Antifungal, Anti-Gram+, Anti-Gram-, Antimicrobial |
| 614 | 876 | DRAMP03422 | Neutrophil antibiotic peptide NP-4 (RatNP-4; Rodents, mammals, | Antibacterial, Antifungal, Antiviral, Anti-Gram+, Anti-Gram-, Antimicrobial |
| 615 | 878 | DRAMP03464 | Cryptonin (Insects, animals) | Antibacterial, Antifungal, Anti-Gram+, Anti-Gram-, Antimicrobial |
| 616 | 880 | DRAMP03467 | Antibacterial napin (Plants) | Antibacterial, Anti-Gram+, Anti-Gram-, Antimicrobial |
| 617 | 881 | DRAMP03471 | Recombinant Crassostrea Gigas Defensin (Cg-Def; molluscs, | Antibacterial, Antifungal, Anti-Gram+, Anti-Gram-, Antimicrobial |
| 618 | 882 | DRAMP03472 | cgUbiquitin | Antibacterial, Antifungal, Anti-Gram+, Anti-Gram-, Antimicrobial |
| 619 | 885 | DRAMP03486 | Manduca Sexta Moricin (MS moricin; Insects, animals) | Antibacterial, Anti-Gram+, Anti-Gram-, Antimicrobial |
| 620 | 887 | DRAMP03507 | Cecropin-B (Insects, animals) | Antibacterial, Antifungal, Anti-Gram+, Anti-Gram-, Antimicrobial |
| 621 | 888 | DRAMP03513 | G. mellonella moricin-like peptide A (Gm-mlpA; Insects, animals; | Antibacterial, Antifungal, Anti-Gram+, Anti-Gram-, Antimicrobial |
| 622 | 889 | DRAMP03514 | G. mellonella moricin-like peptide B (Gm-mlpB; Insects, animals; | Antibacterial, Antifungal, Anti-Gram+, Anti-Gram-, Antimicrobial |
| 623 | 890 | DRAMP03515 | Moricin-like peptide C1 (Gm-mlpC1; Insects, animals; Predicted) | Antibacterial, Antifungal, Anti-Gram+, Anti-Gram-, Antimicrobial |
| 624 | 891 | DRAMP03516 | Moricin-like peptide C2 (Gm-mlpC2; Insects, animals; Predicted) | Antibacterial, Antifungal, Anti-Gram+, Anti-Gram-, Antimicrobial |
| 625 | 892 | DRAMP03517 | Moricin-like peptide C3 (Gm-mlpC3; Insects, animals; Predicted) | Antibacterial, Antifungal, Anti-Gram+, Anti-Gram-, Antimicrobial |
| 626 | 903 | DRAMP03532 | Moricin-1 (Insects, animals) | Antibacterial, Anti-Gram+, Anti-Gram-, Antimicrobial |
| 627 | 908 | DRAMP03567 | KR-20 (Derived from LL-37) | Antibacterial, Antifungal, Anti-Gram+, Anti-Gram-, Antimicrobial |
| 628 | 909 | DRAMP03568 | RK-31 (Derived from LL-37) | Antibacterial, Antifungal, Anti-Gram+, Anti-Gram-, Antimicrobial |
| 629 | 910 | DRAMP03569 | KS-30 (Derived from LL-37) | Antibacterial, Antifungal, Anti-Gram+, Anti-Gram-, Antimicrobial |
| 630 | 911 | DRAMP03570 | LL-23 (Derived from LL-37) | Antibacterial, Antifungal, Anti-Gram+, Anti-Gram-, Antimicrobial |
| 631 | 912 | DRAMP03571 | Antibacterial protein LL-37 (one chain of hCAP-18; Human, | Antibacterial, Anticancer, Anti-Gram+, Anti-Gram-, Antimicrobial |
| 632 | 915 | DRAMP03598 | Human beta-defensin 2 (hBD-2; Defensin, beta 2; Beta-defensin 4A; | Antibacterial, Antiviral, Anti-Gram+, Anti-Gram-, Antimicrobial |
| 633 | 916 | DRAMP03599 | Human beta-defensin 3 (BD-3, hBD-3; Hbd3; Beta-defensin 103; | Antibacterial, Antifungal, Anti-Gram+, Anti-Gram-, Antimicrobial |
| 634 | 917 | DRAMP03600 | Human beta-defensin 4 (hBD-4, BD-4; Beta-defensin 104; Human, | Antibacterial, Anti-Gram+, Anti-Gram-, Antimicrobial |
| 635 | 918 | DRAMP03603 | Human beta-defensin 28 (hBD-28; hBD28; Human, mammals, | Antibacterial, Anti-Gram+, Anti-Gram-, Antimicrobial |
| 636 | 919 | DRAMP03638 | VpBD (V.philippinarum beta defensin; big defensin) | Antibacterial, Anti-Gram+, Anti-Gram-, Antimicrobial |

| S.no. | PepID | DRAMP_ID | Name of the AMP | Activity |
|---|---|---|---|---|
| 637 | 920 | DRAMP03642 | Chicken heterophil peptides 1 (Antimicrobial peptide CHP1; Birds, | Antibacterial, Antifungal, Anti-Gram+, Anti-Gram-, Antimicrobial |
| 638 | 921 | DRAMP03645 | Cathelicidin-2 (CATH-2; Fowlicidin-2; Birds, animals) | Antibacterial, Anti-Gram+, Anti-Gram-, Antimicrobial |
| 639 | 922 | DRAMP03646 | Cathelicidin-3 (CATH-3; Fowlicidin-3; Birds, animals) | Antibacterial, Anti-Gram+, Anti-Gram-, Antimicrobial |
| 640 | 923 | DRAMP03647 | Cathelicidin-B1 (CATH-B1; cathelicidin; Birds, animals) | Antibacterial, Anti-Gram+, Anti-Gram-, Antimicrobial |
| 641 | 925 | DRAMP03676 | GLFcin (Lactoferrin fragment) | Antibacterial, Anti-Gram+, Anti-Gram-, Antimicrobial |
| 642 | 926 | DRAMP03677 | GLFcin II (Lactoferrin fragment) | Antibacterial, Anti-Gram+, Anti-Gram-, Antimicrobial |
| 643 | 927 | DRAMP03679 | Cathelicidin-2 (Bactenecin-5, Bac5; ChBac5; ruminant, animals) | Antibacterial, Anti-Gram+, Anti-Gram-, Antimicrobial |
| 644 | 928 | DRAMP03682 | Vespid chemotactic peptide 5e (VCP 5e; Insects, animals) | Antibacterial, Antifungal, Anti-Gram+, Anti-Gram-, Antimicrobial |
| 645 | 929 | DRAMP03683 | Vespid chemotactic peptide 5g (VCP 5g; Insects, animals) | Antibacterial, Antifungal, Anti-Gram+, Anti-Gram-, Antimicrobial |
| 646 | 930 | DRAMP03684 | Vespid chemotactic peptide 5f (VCP 5f; Insects, animals) | Antibacterial, Antifungal, Anti-Gram+, Anti-Gram-, Antimicrobial |
| 647 | 931 | DRAMP03687 | TsAP-1 (T. serrulatus antimicrobial peptide 1; scorpions, arachnids, | Antibacterial, Anticancer, Anti-Gram+, Anti-Gram-, Antimicrobial |
| 648 | 933 | DRAMP03691 | Im-1 (Arthropods, animals) | Antibacterial, Anti-Gram+, Anti-Gram-, Antimicrobial |
| 649 | 934 | DRAMP03693 | Bactridin-1 (Bact1; Bactridine 1; Arthropods, animals) | Antibacterial, Anti-Gram+, Anti-Gram-, Antimicrobial |
| 650 | 935 | DRAMP03694 | Bactridin-2 (Bact2, Bactridine 2; P-Mice-Antm-beta* NaTx14.8; | Antibacterial, Anti-Gram+, Anti-Gram-, Antimicrobial |
| 651 | 937 | DRAMP03702 | Mucroporin (Antimicrobial peptide 36.21; Arthropods, animals) | Antibacterial, Anti-Gram+, Anti-Gram-, Antimicrobial |
| 652 | 938 | DRAMP03706 | Antimicrobial peptide 1 (AamAP1; Arthropods, animals) | Antibacterial, Antifungal, Anti-Gram+, Anti-Gram-, Antimicrobial |
| 653 | 939 | DRAMP03707 | Antimicrobial peptide 2 (AamAP2; Arthropods, animals) | Antibacterial, Antifungal, Anti-Gram+, Anti-Gram-, Antimicrobial |
| 654 | 940 | DRAMP03714 | Amphiphatic peptide 5.13, NDBP-5.13; Arthropods, animals) | Antibacterial, Anti-Gram+, Anti-Gram-, Antimicrobial |
| 655 | 941 | DRAMP03715 | Amphiphatic peptide 5.14, NDBP-5.14; Arthropods, animals) | Antibacterial, Anti-Gram+, Anti-Gram-, Antimicrobial |
| 656 | 942 | DRAMP03721 | Cytotoxic linear peptide IsCT (IsCT; NDBP-5.2; Arthropods, animals) | Antibacterial, Anti-Gram+, Anti-Gram-, Antimicrobial |
| 657 | 943 | DRAMP03723 | Pandinin-1 (Pin1; Arthropods, animals) | Antibacterial, Anti-Gram+, Anti-Gram-, Antimicrobial |
| 658 | 944 | DRAMP03724 | Pandinin-2 (Pin2; Arthropods, animals) | Antibacterial, Hemolytic activity, Anti-Gram+, Anti-Gram-, Antimicrobial |
| 659 | 946 | DRAMP03734 | Parabutoporin (PP; Non-disulfide-bridged peptide 3.2, NDBP- 3.2; | Antibacterial, Antifungal, Anti-Gram+, Anti-Gram-, Antimicrobial |
| 660 | 947 | DRAMP03735 | Opistoporin-1 (OP1; Non-disulfide-bridged peptide 3.5; Opistoporin- | Antibacterial, Antifungal, Anti-Gram+, Anti-Gram-, Antimicrobial |
| 661 | 948 | DRAMP03738 | Scorpine (defensins; Arthropods, animals) | Antibacterial, Anti-Gram+, Anti-Gram-, Antimicrobial |
| 662 | 949 | DRAMP02828 | BMAP-34 (BMAP 34, bovine cathelicidin, cattle, ruminant, mammals, | Antimicrobial, Antibacterial, Antifungal, Anti-Gram+, Anti-Gram-, |
| 663 | 951 | DRAMP03746 | Peptide BmKn2 (Biologically active peptide 4; NDBP-5.1; Arthropods, | Antibacterial, Anti-Gram+, Anti-Gram-, Antimicrobial |
| 664 | 952 | DRAMP03748 | Bradykinin-potentiating peptide BmK3 (Bpp BmK3; NDBP-3.3; | Antibacterial, Antifungal, Anti-Gram+, Anti-Gram-, Antimicrobial |
| 665 | 953 | DRAMP03750 | Venom antimicrobial peptide-6 (Meucin-13; NDBP-5; Arthropods, | Antibacterial, Antifungal, Anti-Gram+, Anti-Gram-, Antimicrobial |
| 666 | 954 | DRAMP03751 | Venom antimicrobial peptide-9 (Meucin-18; NDBP-5; Arthropods, | Antibacterial, Antifungal, Anti-Gram+, Anti-Gram-, Antimicrobial |
| 667 | 955 | DRAMP03752 | Peptide BmKb1 (Non-disulfide-bridged peptide 4.2, NDBP-4.2; | Antibacterial, Anti-Gram+, Anti-Gram-, Antimicrobial |
| 668 | 956 | DRAMP03753 | Amphipathic peptide CT1 (StCT1; Non-disulfide-bridged peptide 5, | Antibacterial, Anti-Gram+, Anti-Gram-, Antimicrobial |
| 669 | 957 | DRAMP03754 | Amphipathic peptide CT2 (StCT2; Non-disulfide-bridged peptide 5, | Antibacterial, Anti-Gram+, Anti-Gram-, Antimicrobial |
| 670 | 958 | DRAMP03774 | UyCT1 (Arthropods, animals) | Antibacterial, Anti-Gram+, Anti-Gram-, Antimicrobial |
| 671 | 960 | DRAMP03776 | UyCT3 (Arthropods, animals) | Antibacterial, Anti-Gram+, Anti-Gram-, Antimicrobial |
| 672 | 961 | DRAMP03777 | UyCT5 (Arthropods, animals) | Antibacterial, Anti-Gram+, Anti-Gram-, Antimicrobial |
| 673 | 963 | DRAMP03814 | D16W (GGN4 analogue peptide with single substitution) | Antibacterial, Anti-Gram+, Anti-Gram-, Antimicrobial |
| 674 | 964 | DRAMP03815 | D16W-N23 (single amino acid substitution) | Antibacterial, Anti-Gram+, Anti-Gram-, Antimicrobial |
| 675 | 965 | DRAMP03816 | D16F-N23 (single amino acid substitution) | Antibacterial, Anti-Gram+, Anti-Gram-, Antimicrobial |
| 676 | 966 | DRAMP03823 | Dermaseptin derivative K4-S4-(1-13) | Antibacterial, Anti-Gram+, Anti-Gram-, Antimicrobial |
| 677 | 967 | DRAMP03824 | CNBr-cleaved lactoferricin Subfragment 1 | Antibacterial, Anti-Gram+, Anti-Gram-, Antimicrobial |
| 678 | 968 | DRAMP03825 | CNBr-cleaved lactoferricin Subfragment 2 | Antibacterial, Anti-Gram+, Anti-Gram-, Antimicrobial |
| 679 | 969 | DRAMP03826 | Ovispirin-1 (OV-1; N-terminal 18 amino acids of SMAP-29) | Antibacterial, Cytotoxicity, Anti-Gram+, Anti-Gram-, Antimicrobial |
| 680 | 970 | DRAMP03827 | Novispirin G-10 (mutation of Ovispirin-1) | Antibacterial, Cytotoxicity, Anti-Gram+, Anti-Gram-, Antimicrobial |
| 681 | 971 | DRAMP03828 | Novispirin T-7 (mutation of Ovispirin-1) | Antibacterial, Cytotoxicity, Anti-Gram+, Anti-Gram-, Antimicrobial |
| 682 | 973 | DRAMP03830 | Palustrin-2ISb + 3aa | Antibacterial, Anti-Gram+, Anti-Gram-, Antimicrobial |
| 683 | 974 | DRAMP03831 | Palustrin-2ISb-des-C7 | Antibacterial, Antifungal, Anti-Gram+, Anti-Gram-, Antimicrobial |
| 684 | 975 | DRAMP03832 | Palustrin-2ISb-des-C7-4D | Antibacterial, Antifungal, Anti-Gram+, Anti-Gram-, Antimicrobial |
| 685 | 976 | DRAMP03833 | Palustrin-2ISb-des-C7-12N | Antibacterial, Antifungal, Anti-Gram+, Anti-Gram-, Antimicrobial |
| 686 | 977 | DRAMP03834 | Palustrin-2ISb-des-C7-23,29S | Antibacterial, Antifungal, Anti-Gram+, Anti-Gram-, Antimicrobial |
| 687 | 979 | DRAMP03852 | G1 (Bac2A variant through single amino acid substitution) | Antibacterial, Antifungal, Anti-Gram+, Anti-Gram-, Antimicrobial |
| 688 | 980 | DRAMP03853 | G2 (Bac2A variant through single amino acid substitution) | Antibacterial, Antifungal, Anti-Gram+, Anti-Gram-, Antimicrobial |
| 689 | 981 | DRAMP03854 | R2 (Bac2A variant through single amino acid substitution) | Antibacterial, Antifungal, Anti-Gram+, Anti-Gram-, Antimicrobial |

| S.no. | PepID | DRAMP_ID | Name of the AMP | Activity |
|---|---|---|---|---|
| 690 | 982 | DRAMP03855 | R3 (Bac2A variant through single amino acid substitution) | Antibacterial, Antifungal, Anti-Gram+, Anti-Gram-, Antimicrobial |
| 691 | 983 | DRAMP03856 | W3 (Bac2A variant through single amino acid substitution) | Antibacterial, Antifungal, Anti-Gram+, Anti-Gram-, Antimicrobial |
| 692 | 984 | DRAMP03857 | R5 (Bac2A variant through single amino acid substitution) | Antibacterial, Antifungal, Anti-Gram+, Anti-Gram-, Antimicrobial |
| 693 | 985 | DRAMP03858 | K7 (Bac2A variant through single amino acid substitution) | Antibacterial, Antifungal, Anti-Gram+, Anti-Gram-, Antimicrobial |
| 694 | 986 | DRAMP03859 | W10 (Bac2A variant through single amino acid substitution) | Antibacterial, Antifungal, Anti-Gram+, Anti-Gram-, Antimicrobial |
| 695 | 987 | DRAMP03860 | R11 (Bac2A variant through single amino acid substitution) | Antibacterial, Antifungal, Anti-Gram+, Anti-Gram-, Antimicrobial |
| 696 | 988 | DRAMP03861 | G12 (Bac2A variant through single amino acid substitution) | Antibacterial, Antifungal, Anti-Gram+, Anti-Gram-, Antimicrobial |
| 697 | 989 | DRAMP03862 | Sub2 (Bac2A variant through two amino acids substitution) | Antibacterial, Antifungal, Anti-Gram+, Anti-Gram-, Antimicrobial |
| 698 | 990 | DRAMP03863 | Sub3 (Bac2A variant through three amino acids substitution) | Antibacterial, Antifungal, Anti-Gram+, Anti-Gram-, Antimicrobial |
| 699 | 991 | DRAMP03864 | Sub5 (Bac2A variant through five amino acids substitution) | Antibacterial, Antifungal, Anti-Gram+, Anti-Gram-, Antimicrobial |
| 700 | 992 | DRAMP03865 | Sub6 (Bac2A variant through six amino acids substitution) | Antibacterial, Antifungal, Anti-Gram+, Anti-Gram-, Antimicrobial |
| 701 | 993 | DRAMP03866 | Bac8a (Bac2A variant) | Antibacterial, Antifungal, Anti-Gram+, Anti-Gram-, Antimicrobial |
| 702 | 994 | DRAMP03867 | Bac8b (Bac2A variant) | Antibacterial, Antifungal, Anti-Gram+, Anti-Gram-, Antimicrobial |
| 703 | 995 | DRAMP03868 | Bac8c (Bac2A variant) | Antibacterial, Antifungal, Anti-Gram+, Anti-Gram-, Antimicrobial |
| 704 | 996 | DRAMP03869 | Bac8d (Bac2A variant) | Antibacterial, Antifungal, Anti-Gram+, Anti-Gram-, Antimicrobial |
| 705 | 997 | DRAMP03870 | Bac2A (a linear variant of bovine dodecapeptide) | Antibacterial, Antifungal, Anti-Gram+, Anti-Gram-, Antimicrobial |
| 706 | 998 | DRAMP03871 | cLf 20-29 (fragment of caprine lalctoferricin, residues 20-29) | Antibacterial, Anti-Gram+, Anti-Gram-, Antimicrobial |
| 707 | 999 | DRAMP03875 | bLf 20-29 (fragment of bovine lactoferricin, residues 20-29) | Antibacterial, Anti-Gram+, Anti-Gram-, Antimicrobial |
| 708 | 1000 | DRAMP03876 | LFB-RW (derivative of bovine lactoferrin with residues substitution) | Antibacterial, Anti-Gram+, Anti-Gram-, Antimicrobial |
| 709 | 1001 | DRAMP03877 | LFB-KW (derivative of bovine lactoferrin with residues substitution) | Antibacterial, Anti-Gram+, Anti-Gram-, Antimicrobial |
| 710 | 1002 | DRAMP03878 | LFB-Rwa (derivative of bovine lactoferrin with residues substitution) | Antibacterial, Anti-Gram+, Anti-Gram-, Antimicrobial |
| 711 | 1003 | DRAMP03879 | LFB-RF (derivative of bovine lactoferrin with residues substitution) | Antibacterial, Anti-Gram+, Anti-Gram-, Antimicrobial |
| 712 | 1004 | DRAMP03880 | LFB-RI (derivative of bovine lactoferrin with residues substitution) | Antibacterial, Anti-Gram+, Anti-Gram-, Antimicrobial |
| 713 | 1005 | DRAMP03881 | LFB-6RW (derivative of bovine lactoferrin with residues substitution) | Antibacterial, Anti-Gram+, Anti-Gram-, Antimicrobial |
| 714 | 1006 | DRAMP03882 | LFC (fragment of mature caprine lactoferrin, residues 17 to 31) | Antibacterial, Anti-Gram+, Anti-Gram-, Antimicrobial |
| 715 | 1007 | DRAMP03883 | LFH W8 (tryptophan-modified human lactoferricin derivative) | Antibacterial, Anti-Gram+, Anti-Gram-, Antimicrobial |
| 716 | 1008 | DRAMP03884 | LFC W8 (tryptophan-modified caprine lactoferricin derivative) | Antibacterial, Anti-Gram+, Anti-Gram-, Antimicrobial |
| 717 | 1009 | DRAMP03885 | LFP W8 (tryptophan-modified porcine lactoferricin derivative) | Antibacterial, Anti-Gram+, Anti-Gram-, Antimicrobial |
| 718 | 1010 | DRAMP03886 | LFB (fragment of bovine lalctoferricin, residues 17 to 31) | Antibacterial, Anti-Gram+, Anti-Gram-, Antimicrobial |
| 719 | 1011 | DRAMP03887 | LFB A1 (derivative of LFB, residue substitution with alanine at | Antibacterial, Anti-Gram+, Anti-Gram-, Antimicrobial |
| 720 | 1012 | DRAMP03888 | LFB A2 (derivative of LFB, residue substitution with alanine at | Antibacterial, Anti-Gram+, Anti-Gram-, Antimicrobial |
| 721 | 1013 | DRAMP03889 | LFB A3 (derivative of LFB, residue substitution with alanine at | Antibacterial, Anti-Gram+, Anti-Gram-, Antimicrobial |
| 722 | 1014 | DRAMP03890 | LFB A4 (derivative of LFB, residue substitution with alanine at | Antibacterial, Anti-Gram+, Anti-Gram-, Antimicrobial |
| 723 | 1015 | DRAMP03891 | LFB A5 (derivative of LFB, residue substitution with alanine at | Antibacterial, Anti-Gram+, Anti-Gram-, Antimicrobial |
| 724 | 1016 | DRAMP03892 | LFB A7 (derivative of LFB, residue substitution with alanine at | Antibacterial, Anti-Gram+, Anti-Gram-, Antimicrobial |
| 725 | 1017 | DRAMP03893 | LFB A9 (derivative of LFB, residue substitution with alanine at | Antibacterial, Anti-Gram+, Anti-Gram-, Antimicrobial |
| 726 | 1018 | DRAMP03894 | LFB A10 (derivative of LFB, residue substitution with alanine at | Antibacterial, Anti-Gram+, Anti-Gram-, Antimicrobial |
| 727 | 1019 | DRAMP03895 | LFB A11 (derivative of LFB, residue substitution with alanine at | Antibacterial, Anti-Gram+, Anti-Gram-, Antimicrobial |
| 728 | 1020 | DRAMP03896 | LFB A12 (derivative of LFB, residue substitution with alanine at | Antibacterial, Anti-Gram+, Anti-Gram-, Antimicrobial |
| 729 | 1021 | DRAMP03897 | LFB A13 (derivative of LFB, residue substitution with alanine at | Antibacterial, Anti-Gram+, Anti-Gram-, Antimicrobial |
| 730 | 1022 | DRAMP03898 | LFB A14 (derivative of LFB, residue substitution with alanine at | Antibacterial, Anti-Gram+, Anti-Gram-, Antimicrobial |
| 731 | 1025 | DRAMP03901 | LFM R1 W8 (LFM W8 derivative with residues substitution) | Antibacterial, Anti-Gram+, Anti-Gram-, Antimicrobial |
| 732 | 1027 | DRAMP03903 | LFM A1 R9 W8 (LFM W8 derivative with residues substitution) | Antibacterial, Anti-Gram+, Anti-Gram-, Antimicrobial |
| 733 | 1028 | DRAMP03904 | LFM A9 R1 W8 (LFM W8 derivative with residues substitution) | Antibacterial, Anti-Gram+, Anti-Gram-, Antimicrobial |
| 734 | 1029 | DRAMP03905 | LFM R1,9 W8 (LFM W8 derivative with residues substitution) | Antibacterial, Anti-Gram+, Anti-Gram-, Antimicrobial |
| 735 | 1032 | DRAMP03908 | LFM R1 W8 Y13 (LFM W8 derivative with residues substitution) | Antibacterial, Anti-Gram+, Anti-Gram-, Antimicrobial |
| 736 | 1034 | DRAMP03910 | LFM A1 R9 W8 Y13 (LFM W8 derivative with residues substitution) | Antibacterial, Anti-Gram+, Anti-Gram-, Antimicrobial |
| 737 | 1035 | DRAMP03911 | LFM A9 R1 W8 Y13 (LFM W8 derivative with residues substitution) | Antibacterial, Anti-Gram+, Anti-Gram-, Antimicrobial |
| 738 | 1036 | DRAMP03912 | LFM R1,9 W8 Y13 (LFM W8 derivative with residues substitution) | Antibacterial, Anti-Gram+, Anti-Gram-, Antimicrobial |
| 739 | 1037 | DRAMP03920 | Cecropin A (1-8)-melittin (1-13)hybrid peptide | Antibacterial, Anti-Gram+, Anti-Gram-, Antimicrobial |
| 740 | 1038 | DRAMP03921 | Cecropin A (1-8)-melittin (1-18)hybrid peptide | Antibacterial, Anti-Gram+, Anti-Gram-, Antimicrobial |
| 741 | 1039 | DRAMP03922 | Cecropin A (1-8)-melittin (1-12)hybrid peptide | Antibacterial, Anti-Gram+, Anti-Gram-, Antimicrobial |
| 742 | 1040 | DRAMP03923 | Cecropin A (1-8)-melittin (1-10)hybrid peptide | Antibacterial, Anti-Gram+, Anti-Gram-, Antimicrobial |

| S.no. | PepID | DRAMP_ID | Name of the AMP | Activity |
|---|---|---|---|---|
| 743 | 1041 | DRAMP03924 | Cecropin A (1-7)-melittin (1-8)hybrid peptide | Antibacterial, Anti-Gram+, Anti-Gram-, Antimicrobial |
| 744 | 1042 | DRAMP03925 | Cecropin A (1-7)-melittin (3-10)hybrid peptide | Antibacterial, Anti-Gram+, Anti-Gram-, Antimicrobial |
| 745 | 1043 | DRAMP03927 | Cecropin A (1-7)-melittin (2-9)hybrid peptide | Antibacterial, Anti-Gram+, Anti-Gram-, Antimicrobial |
| 746 | 1044 | DRAMP03928 | Cecropin A (1-7)-melittin (4-11)hybrid peptide (CAM) | Antibacterial, Anti-Gram+, Anti-Gram-, Antimicrobial |
| 747 | 1045 | DRAMP03929 | Cecropin A (1-7)-melittin (5-12)hybrid peptide | Antibacterial, Anti-Gram+, Anti-Gram-, Antimicrobial |
| 748 | 1046 | DRAMP03930 | Cecropin A (1-7)-melittin (6-13)hybrid peptide | Antibacterial, Antifungal, Anti-Gram+, Anti-Gram-, Antimicrobial |
| 749 | 1048 | DRAMP03933 | I14M (truncated isoform of thanatin, residue 8-21) | Antibacterial, Antifungal, Anti-Gram+, Anti-Gram-, Antimicrobial |
| 750 | 1050 | DRAMP03935 | V16M (truncated isoform of thanatin, residue 6-21) | Antibacterial, Antifungal, Anti-Gram+, Anti-Gram-, Antimicrobial |
| 751 | 1051 | DRAMP03936 | K18M (truncated isoform of thanatin, residue 4-21) | Antibacterial, Antifungal, Anti-Gram+, Anti-Gram-, Antimicrobial |
| 752 | 1055 | DRAMP03945 | Del 1-4 (Ranalexin analog) | Antibacterial, Anti-Gram+, Anti-Gram-, Antimicrobial |
| 753 | 1056 | DRAMP03947 | Del 1-2 (Ranalexin analog) | Antibacterial, Anti-Gram+, Anti-Gram-, Antimicrobial |
| 754 | 1057 | DRAMP03948 | Del 1 (Ranalexin analog) | Antibacterial, Anti-Gram+, Anti-Gram-, Antimicrobial |
| 755 | 1058 | DRAMP03949 | Del 20 (Ranalexin analog) | Antibacterial, Anti-Gram+, Anti-Gram-, Antimicrobial |
| 756 | 1065 | DRAMP03967 | P18 (Cecropin A(1-8)-Magainin 2(1-12) hybrid peptide analogue) | Antibacterial, Antitumour, Anti-Gram+, Anti-Gram-, Antimicrobial |
| 757 | 1066 | DRAMP03968 | [L9]-P18 (analog of P18) | Antibacterial, Antitumour, Anti-Gram+, Anti-Gram-, Antimicrobial |
| 758 | 1067 | DRAMP03969 | [S9]-P18 (analog of P18) | Antibacterial, Antitumour, Anti-Gram+, Anti-Gram-, Antimicrobial |
| 759 | 1068 | DRAMP03970 | N-1 (analog of P18) | Antibacterial, Antitumour, Anti-Gram+, Anti-Gram-, Antimicrobial |
| 760 | 1069 | DRAMP03971 | N-2 (analog of P18) | Antibacterial, Antitumour, Anti-Gram+, Anti-Gram-, Antimicrobial |
| 761 | 1070 | DRAMP03972 | N-3 (analog of P18) | Antibacterial, Antitumour, Anti-Gram+, Anti-Gram-, Antimicrobial |
| 762 | 1071 | DRAMP03973 | N-4 (analog of P18) | Antibacterial, Antitumour, Anti-Gram+, Anti-Gram-, Antimicrobial |
| 763 | 1072 | DRAMP03974 | N-5 (analog of P18) | Antibacterial, Antitumour, Anti-Gram+, Anti-Gram-, Antimicrobial |
| 764 | 1073 | DRAMP03975 | N-3L (analog of P18) | Antibacterial, Antitumour, Anti-Gram+, Anti-Gram-, Antimicrobial |
| 765 | 1074 | DRAMP03976 | N-4L (analog of P18) | Antibacterial, Antitumour, Anti-Gram+, Anti-Gram-, Antimicrobial |
| 766 | 1075 | DRAMP03977 | N-5L (analog of P18) | Antibacterial, Antitumour, Anti-Gram+, Anti-Gram-, Antimicrobial |
| 767 | 1076 | DRAMP03978 | C-1 (analog of P18) | Antibacterial, Antitumour, Anti-Gram+, Anti-Gram-, Antimicrobial |
| 768 | 1077 | DRAMP03979 | C-2 (analog of P18) | Antibacterial, Antitumour, Anti-Gram+, Anti-Gram-, Antimicrobial |
| 769 | 1078 | DRAMP03980 | C-3 (analog of P18) | Antibacterial, Antitumour, Anti-Gram+, Anti-Gram-, Antimicrobial |
| 770 | 1079 | DRAMP03981 | C-4 (analog of P18) | Antibacterial, Antitumour, Anti-Gram+, Anti-Gram-, Antimicrobial |
| 771 | 1080 | DRAMP03982 | C-5 (analog of P18) | Antibacterial, Antitumour, Anti-Gram+, Anti-Gram-, Antimicrobial |
| 772 | 1081 | DRAMP03983 | C-6 (analog of P18) | Antibacterial, Antitumour, Anti-Gram+, Anti-Gram-, Antimicrobial |
| 773 | 1082 | DRAMP03984 | C-7 (analog of P18) | Antibacterial, Antitumour, Anti-Gram+, Anti-Gram-, Antimicrobial |
| 774 | 1083 | DRAMP03985 | C-8 (analog of P18) | Antibacterial, Antitumour, Anti-Gram+, Anti-Gram-, Antimicrobial |
| 775 | 1084 | DRAMP03986 | C-9 (analog of P18) | Antibacterial, Antitumour, Anti-Gram+, Anti-Gram-, Antimicrobial |
| 776 | 1085 | DRAMP03987 | C-10 (analog of P18) | Antibacterial, Antitumour, Anti-Gram+, Anti-Gram-, Antimicrobial |
| 777 | 1088 | DRAMP03990 | L4K3W4 (LlKmWn model peptide) | Antibacterial, Anti-Gram+, Anti-Gram-, Antimicrobial |
| 778 | 1090 | DRAMP03992 | L5K3W5 (LlKmWn model peptide) | Antibacterial, Anti-Gram+, Anti-Gram-, Antimicrobial |
| 779 | 1091 | DRAMP03993 | L5K5W6 (LlKmWn model peptide) | Antibacterial, Anti-Gram+, Anti-Gram-, Antimicrobial |
| 780 | 1092 | DRAMP03994 | L6K4W6 (LlKmWn model peptide) | Antibacterial, Anti-Gram+, Anti-Gram-, Antimicrobial |
| 781 | 1093 | DRAMP03995 | L7K3W6 (LlKmWn model peptide) | Antibacterial, Anti-Gram+, Anti-Gram-, Antimicrobial |
| 782 | 1096 | DRAMP03999 | [A6]-IsCT (Mutant: W6A; IsCT analog) | Antibacterial, Anti-Gram+, Anti-Gram-, Antimicrobial |
| 783 | 1097 | DRAMP04000 | [L6]-IsCT (Mutant: W6L; IsCT analog) | Antibacterial, Anti-Gram+, Anti-Gram-, Antimicrobial |
| 784 | 1098 | DRAMP04001 | [K7]-IsCT (Mutant: E7K; IsCT analog) | Antibacterial, Anti-Gram+, Anti-Gram-, Antimicrobial |
| 785 | 1099 | DRAMP04002 | [L6, K11]-IsCT (IsCT analog through amino acids substitution) | Antibacterial, Anti-Gram+, Anti-Gram-, Antimicrobial |
| 786 | 1100 | DRAMP04003 | [K7, P8, K11]-IsCT (IsCT analog through amino acids substitution) | Antibacterial, Anti-Gram+, Anti-Gram-, Antimicrobial |
| 787 | 1101 | DRAMP04004 | Gramicidin analogue ([Scr2]-GS) | Antibacterial, Anti-Gram+, Anti-Gram-, Antimicrobial |
| 788 | 1102 | DRAMP04005 | Gramicidin analogue ([Ser2,2']-GS) | Antibacterial, Anti-Gram+, Anti-Gram-, Antimicrobial |
| 789 | 1103 | DRAMP04011 | Plasticin PD36 KF (analog of PD36) | Antibacterial, Anti-Gram+, Anti-Gram-, Antimicrobial |
| 790 | 1104 | DRAMP04012 | Plasticin PD36 K (analog of PD36) | Antibacterial, Anti-Gram+, Anti-Gram-, Antimicrobial |
| 791 | 1105 | DRAMP04013 | Plasticin ANC KF (analog of natural peptide ANC) | Antibacterial, Anti-Gram+, Anti-Gram-, Antimicrobial |
| 792 | 1109 | DRAMP04017 | LL-23V9 (LL-23 variants) | Antibacterial, Anti-Gram+, Anti-Gram-, Antimicrobial |
| 793 | 1110 | DRAMP04019 | Bac014 (Scrambled Variants of Bac2A) | Antibacterial, Antifungal, Anti-Gram+, Anti-Gram-, Antimicrobial |
| 794 | 1111 | DRAMP04020 | Bac020 (Scrambled Variants of Bac2A) | Antibacterial, Antifungal, Anti-Gram+, Anti-Gram-, Antimicrobial |
| 795 | 1112 | DRAMP04021 | Bac034 (Scrambled Variants of Bac2A) | Antibacterial, Antifungal, Anti-Gram+, Anti-Gram-, Antimicrobial |

| S.no. | PepID | DRAMP_ID | Name of the AMP | Activity |
|---|---|---|---|---|
| 796 | 1113 | DRAMP04022 | F3 (single amino acid substitution of Bac034, which is a scrambled | Antibacterial, Antifungal, Anti-Gram+, Anti-Gram-, Antimicrobial |
| 797 | 1114 | DRAMP04023 | W3 (single amino acid substitution of Bac034, which is a scrambled | Antibacterial, Antifungal, Anti-Gram+, Anti-Gram-, Antimicrobial |
| 798 | 1115 | DRAMP04024 | W4 (single amino acid substitution of Bac034, which is a scrambled | Antibacterial, Antifungal, Anti-Gram+, Anti-Gram-, Antimicrobial |
| 799 | 1116 | DRAMP04025 | R10 (single amino acid substitution of Bac034, which is a scrambled | Antibacterial, Antifungal, Anti-Gram+, Anti-Gram-, Antimicrobial |
| 800 | 1117 | DRAMP04026 | K12 (single amino acid substitution of Bac034, which is a scrambled | Antibacterial, Antifungal, Anti-Gram+, Anti-Gram-, Antimicrobial |
| 801 | 1118 | DRAMP04027 | opt1 (multiple amino acid substitution of Bac034, which is a | Antibacterial, Antifungal, Anti-Gram+, Anti-Gram-, Antimicrobial |
| 802 | 1119 | DRAMP04028 | opt2 (multiple amino acid substitution of Bac034, which is a | Antibacterial, Antifungal, Anti-Gram+, Anti-Gram-, Antimicrobial |
| 803 | 1120 | DRAMP04029 | opt3 (multiple amino acid substitution of Bac034, which is a | Antibacterial, Antifungal, Anti-Gram+, Anti-Gram-, Antimicrobial |
| 804 | 1121 | DRAMP04030 | opt4 (multiple amino acid substitution of Bac034, which is a | Antibacterial, Antifungal, Anti-Gram+, Anti-Gram-, Antimicrobial |
| 805 | 1122 | DRAMP04031 | opt5 (multiple amino acid substitution of Bac034, which is a | Antibacterial, Antifungal, Anti-Gram+, Anti-Gram-, Antimicrobial |
| 806 | 1123 | DRAMP04032 | Modified defensin | Antibacterial, Anti-Gram+, Anti-Gram-, Antimicrobial |
| 807 | 1124 | DRAMP04033 | Modified defensin | Antibacterial, Anti-Gram+, Anti-Gram-, Antimicrobial |
| 808 | 1125 | DRAMP04034 | Modified defensin | Antibacterial, Anti-Gram+, Anti-Gram-, Antimicrobial |
| 809 | 1126 | DRAMP04035 | Modified defensin | Antibacterial, Anti-Gram+, Anti-Gram-, Antimicrobial |
| 810 | 1127 | DRAMP04036 | Modified defensin | Antibacterial, Anti-Gram+, Anti-Gram-, Antimicrobial |
| 811 | 1128 | DRAMP04048 | BacR (cyclic derivative of bactenecin) | Antibacterial, Anti-Gram+, Anti-Gram-, Antimicrobial |
| 812 | 1129 | DRAMP04049 | BacP3R (cyclic derivative of bactenecin) | Antibacterial, Anti-Gram+, Anti-Gram-, Antimicrobial |
| 813 | 1130 | DRAMP04050 | BacP3R-V (cyclic derivative of bactenecin) | Antibacterial, Anti-Gram+, Anti-Gram-, Antimicrobial |
| 814 | 1131 | DRAMP04051 | Bac2I-NH2 (cyclic derivative of bactenecin) | Antibacterial, Anti-Gram+, Anti-Gram-, Antimicrobial |
| 815 | 1132 | DRAMP04052 | BacP2R-NH2 (cyclic derivative of bactenecin) | Antibacterial, Anti-Gram+, Anti-Gram-, Antimicrobial |
| 816 | 1133 | DRAMP04053 | BacP1 (cyclic derivative of bactenecin) | Antibacterial, Anti-Gram+, Anti-Gram-, Antimicrobial |
| 817 | 1134 | DRAMP04054 | BacW (cyclic derivative of bactenecin) | Antibacterial, Anti-Gram+, Anti-Gram-, Antimicrobial |
| 818 | 1135 | DRAMP04055 | BacW2R (cyclic derivative of bactenecin) | Antibacterial, Anti-Gram+, Anti-Gram-, Antimicrobial |
| 819 | 1136 | DRAMP04056 | Lin Bac2S-NH2 (linear derivative of bactenecin) | Antibacterial, Anti-Gram+, Anti-Gram-, Antimicrobial |
| 820 | 1137 | DRAMP04057 | Lin BacS-NH2 (linear derivative of bactenecin) | Antibacterial, Anti-Gram+, Anti-Gram-, Antimicrobial |
| 821 | 1153 | DRAMP04075 | Antimicrobial peptide HP (2-20) | Antibacterial, Antifungal, Anti-Gram+, Anti-Gram-, Antimicrobial |
| 822 | 1154 | DRAMP04076 | Anal 1 (antimicrobial peptide HP (2-20)analogue) | Antibacterial, Antifungal, Anti-Gram+, Anti-Gram-, Antimicrobial |
| 823 | 1155 | DRAMP04077 | Anal 2 (antimicrobial peptide HP (2-20)analogue) | Antibacterial, Antifungal, Anti-Gram+, Anti-Gram-, Antimicrobial |
| 824 | 1156 | DRAMP04078 | Anal 3 (antimicrobial peptide HP (2-20)analogue) | Antibacterial, Antifungal, Anti-Gram+, Anti-Gram-, Antimicrobial |
| 825 | 1157 | DRAMP04079 | Anal 4 (antimicrobial peptide HP (2-20)analogue) | Antibacterial, Antifungal, Anti-Gram+, Anti-Gram-, Antimicrobial |
| 826 | 1158 | DRAMP04080 | Anal 5 (antimicrobial peptide HP (2-20)analogue) | Antibacterial, Antifungal, Anti-Gram+, Anti-Gram-, Antimicrobial |
| 827 | 1159 | DRAMP04081 | Anal 6 (antimicrobial peptide HP (2-20)analogue) | Antibacterial, Antifungal, Anti-Gram+, Anti-Gram-, Antimicrobial |
| 828 | 1160 | DRAMP04082 | Anal 7 (antimicrobial peptide HP (2-20)analogue) | Antibacterial, Antifungal, Anti-Gram+, Anti-Gram-, Antimicrobial |
| 829 | 1161 | DRAMP04083 | D-amino-acid pexiganan (MSI-214) | Antibacterial, Anti-Gram+, Anti-Gram-, Antimicrobial |
| 830 | 1162 | DRAMP04095 | Cupiennin-1D (spiders, Arthropods, animals) | Antibacterial, Anti-Gram+, Anti-Gram-, Antimicrobial |
| 831 | 1163 | DRAMP04096 | 2IQ2 | Antibacterial, Antifungal, Anti-Gram+, Anti-Gram-, Antimicrobial |
| 832 | 1164 | DRAMP04097 | 2IQ3 | Antibacterial, Antifungal, Anti-Gram+, Anti-Gram-, Antimicrobial |
| 833 | 1165 | DRAMP04098 | 3IQ1 | Antibacterial, Antifungal, Anti-Gram+, Anti-Gram-, Antimicrobial |
| 834 | 1166 | DRAMP04099 | 3IQ2 | Antibacterial, Antifungal, Anti-Gram+, Anti-Gram-, Antimicrobial |
| 835 | 1167 | DRAMP04100 | 3IQ3 | Antibacterial, Antifungal, Anti-Gram+, Anti-Gram-, Antimicrobial |
| 836 | 1168 | DRAMP04101 | 3IQ4 | Antibacterial, Antifungal, Anti-Gram+, Anti-Gram-, Antimicrobial |
| 837 | 1180 | DRAMP04115 | K11 (derivative of CP-P) | Antibacterial, Anti-Gram+, Anti-Gram-, Antimicrobial |
| 838 | 1186 | DRAMP04123 | D0-NH2 | Antibacterial, Anti-Gram+, Anti-Gram-, Antimicrobial |
| 839 | 1187 | DRAMP04124 | D1-NH2 | Antibacterial, Anti-Gram+, Anti-Gram-, Antimicrobial |
| 840 | 1188 | DRAMP04125 | D2-NH2 | Antibacterial, Anti-Gram+, Anti-Gram-, Antimicrobial |
| 841 | 1189 | DRAMP04126 | D3-NH2 | Antibacterial, Anti-Gram+, Anti-Gram-, Antimicrobial |
| 842 | 1190 | DRAMP04127 | D4-NH2 | Antibacterial, Anti-Gram+, Anti-Gram-, Antimicrobial |
| 843 | 1191 | DRAMP04128 | D5-NH2 | Antibacterial, Anti-Gram+, Anti-Gram-, Antimicrobial |
| 844 | 1192 | DRAMP04129 | D6-NH2 | Antibacterial, Anti-Gram+, Anti-Gram-, Antimicrobial |
| 845 | 1193 | DRAMP04136 | LRR-1 | Antibacterial, Anti-Gram+, Anti-Gram-, Antimicrobial |
| 846 | 1194 | DRAMP04137 | LRR-2 | Antibacterial, Anti-Gram+, Anti-Gram-, Antimicrobial |
| 847 | 1205 | DRAMP04159 | LR2 (homologue of Pc-CATH1) | Antibacterial, Antifungal, Anti-Gram+, Anti-Gram-, Antimicrobial |
| 848 | 1206 | DRAMP04160 | LR3 (homologue of Pc-CATH1) | Antibacterial, Antifungal, Anti-Gram+, Anti-Gram-, Antimicrobial |

| S.no. | PepID | DRAMP_ID | Name of the AMP | Activity |
|---|---|---|---|---|
| 849 | 1207 | DRAMP04161 | LR4 (homologue of Pc-CATH1) | Antibacterial, Antifungal, Anti-Gram+, Anti-Gram-, Antimicrobial |
| 850 | 1208 | DRAMP04162 | LR5 (homologue of Pc-CATH1) | Antibacterial, Antifungal, Anti-Gram+, Anti-Gram-, Antimicrobial |
| 851 | 1209 | DRAMP04163 | LR6 (homologue of Pc-CATH1) | Antibacterial, Antifungal, Anti-Gram+, Anti-Gram-, Antimicrobial |
| 852 | 1210 | DRAMP04164 | LR7 (homologue of Pc-CATH1) | Antibacterial, Antifungal, Anti-Gram+, Anti-Gram-, Antimicrobial |
| 853 | 1211 | DRAMP04165 | LR8 (homologue of Pc-CATH1) | Antibacterial, Antifungal, Anti-Gram+, Anti-Gram-, Antimicrobial |
| 854 | 1212 | DRAMP04166 | LR9 (homologue of Pc-CATH1) | Antibacterial, Antifungal, Anti-Gram+, Anti-Gram-, Antimicrobial |
| 855 | 1213 | DRAMP04167 | LR10 (homologue of Pc-CATH1) | Antifungal, Anti-Gram+, Anti-Gram-, Antimicrobial |
| 856 | 1214 | DRAMP04168 | LR11 (homologue of Pc-CATH1) | Antifungal, Anti-Gram+, Anti-Gram-, Antimicrobial |
| 857 | 1215 | DRAMP04169 | LR13 (homologue of Pc-CATH1) | Antifungal, Anti-Gram+, Anti-Gram-, Antimicrobial |
| 858 | 1216 | DRAMP04170 | LR15 (homologue of Pc-CATH1) | Antifungal, Anti-Gram+, Anti-Gram-, Antimicrobial |
| 859 | 1217 | DRAMP04171 | LR16 (homologue of Pc-CATH1) | Antifungal, Anti-Gram+, Anti-Gram-, Antimicrobial |
| 860 | 1218 | DRAMP04174 | L2K3W2 (LlKmW2 model peptides) | Antibacterial, Anti-Gram+, Anti-Gram-, Antimicrobial |
| 861 | 1219 | DRAMP04175 | L3K2W2 (LlKmW2 model peptides) | Antibacterial, Anti-Gram+, Anti-Gram-, Antimicrobial |
| 862 | 1220 | DRAMP04176 | L2K5W2 (LlKmW2 model peptides) | Antibacterial, Anti-Gram+, Anti-Gram-, Antimicrobial |
| 863 | 1221 | DRAMP04177 | L3K4W2 (LlKmW2 model peptides) | Antibacterial, Anti-Gram+, Anti-Gram-, Antimicrobial |
| 864 | 1222 | DRAMP04178 | L4K3W2 (LlKmW2 model peptides) | Antibacterial, Anti-Gram+, Anti-Gram-, Antimicrobial |
| 865 | 1223 | DRAMP04179 | L5K2W2 (LlKmW2 model peptides) | Antibacterial, Anti-Gram+, Anti-Gram-, Antimicrobial |
| 866 | 1224 | DRAMP04180 | L3K6W2 (LlKmW2 model peptides) | Antibacterial, Anti-Gram+, Anti-Gram-, Antimicrobial |
| 867 | 1225 | DRAMP04181 | L4K5W2 (LlKmW2 model peptides) | Antibacterial, Anti-Gram+, Anti-Gram-, Antimicrobial |
| 868 | 1226 | DRAMP04182 | L5K4W2 (LlKmW2 model peptides) | Antibacterial, Anti-Gram+, Anti-Gram-, Antimicrobial |
| 869 | 1227 | DRAMP04183 | L6K3W2 (LlKmW2 model peptides) | Antibacterial, Anti-Gram+, Anti-Gram-, Antimicrobial |
| 870 | 1229 | DRAMP04185 | DFTamP1-p | Antibacterial, Anti-Gram+, Anti-Gram-, Antimicrobial |
| 871 | 1230 | DRAMP04186 | L5K5W1 (L5K5Wn model peptide) | Antibacterial, Anti-Gram+, Anti-Gram-, Antimicrobial |
| 872 | 1231 | DRAMP04187 | L5K5W2 (L5K5Wn model peptide) | Antibacterial, Anti-Gram+, Anti-Gram-, Antimicrobial |
| 873 | 1232 | DRAMP04188 | L5K5W3 (L5K5Wn model peptide) | Antibacterial, Anti-Gram+, Anti-Gram-, Antimicrobial |
| 874 | 1233 | DRAMP04189 | L5K5W4 (L5K5Wn model peptide) | Antibacterial, Anti-Gram+, Anti-Gram-, Antimicrobial |
| 875 | 1234 | DRAMP04190 | L5K5W5 (L5K5Wn model peptide) | Antibacterial, Anti-Gram+, Anti-Gram-, Antimicrobial |
| 876 | 1236 | DRAMP04192 | L5K5W7 (L5K5Wn model peptide) | Antibacterial, Anti-Gram+, Anti-Gram-, Antimicrobial |
| 877 | 1237 | DRAMP04193 | L5K5W8 (L5K5Wn model peptide) | Antibacterial, Anti-Gram+, Anti-Gram-, Antimicrobial |
| 878 | 1238 | DRAMP04194 | L5K5W9 (L5K5Wn model peptide) | Antibacterial, Anti-Gram+, Anti-Gram-, Antimicrobial |
| 879 | 1239 | DRAMP04195 | L5K5W10 (L5K5Wn model peptide) | Antibacterial, Anti-Gram+, Anti-Gram-, Antimicrobial |
| 880 | 1240 | DRAMP04196 | L5K5W11 (L5K5Wn model peptide) | Antibacterial, Anti-Gram+, Anti-Gram-, Antimicrobial |
| 881 | 1245 | DRAMP04240 | Synthetic 1 | Antibacterial, Anti-Gram+, Anti-Gram-, Antimicrobial |
| 882 | 1246 | DRAMP04241 | Synthetic 2 | Antibacterial, Anti-Gram+, Anti-Gram-, Antimicrobial |
| 883 | 1247 | DRAMP04242 | Synthetic 3 | Antibacterial, Anti-Gram+, Anti-Gram-, Antimicrobial |
| 884 | 1248 | DRAMP04243 | Synthetic 4 | Antibacterial, Anti-Gram+, Anti-Gram-, Antimicrobial |
| 885 | 1249 | DRAMP04244 | Synthetic 5 | Antibacterial, Anti-Gram+, Anti-Gram-, Antimicrobial |
| 886 | 1253 | DRAMP04359 | PDD-A-1 (PDD-A analog) | Antibacterial, Anti-Gram+, Anti-Gram-, Antimicrobial |
| 887 | 1254 | DRAMP04360 | PDD-A-2 (PDD-A analog) | Antibacterial, Anti-Gram+, Anti-Gram-, Antimicrobial |
| 888 | 1255 | DRAMP04361 | PDD-A-3 (PDD-A analog) | Antibacterial, Anti-Gram+, Anti-Gram-, Antimicrobial |
| 889 | 1256 | DRAMP04362 | PDD-A-4 (PDD-A analog) | Antibacterial, Anti-Gram+, Anti-Gram-, Antimicrobial |
| 890 | 1257 | DRAMP04363 | PDD-A-5 (PDD-A analog) | Antibacterial, Anti-Gram+, Anti-Gram-, Antimicrobial |
| 891 | 1258 | DRAMP04364 | PDD-A-6 (PDD-A analog) | Antibacterial, Anti-Gram+, Anti-Gram-, Antimicrobial |
| 892 | 1259 | DRAMP04365 | PDD-A-7 (PDD-A analog) | Antibacterial, Anti-Gram+, Anti-Gram-, Antimicrobial |
| 893 | 1260 | DRAMP04367 | PDD-A-9 (PDD-A analog) | Antibacterial, Anti-Gram+, Anti-Gram-, Antimicrobial |
| 894 | 1261 | DRAMP04368 | PDD-A-10 (PDD-A analog) | Antibacterial, Anti-Gram+, Anti-Gram-, Antimicrobial |
| 895 | 1262 | DRAMP04369 | PDD-A-11 (PDD-A analog) | Antibacterial, Anti-Gram+, Anti-Gram-, Antimicrobial |
| 896 | 1263 | DRAMP04370 | PDD-A-12 (PDD-A analog) | Antibacterial, Anti-Gram+, Anti-Gram-, Antimicrobial |
| 897 | 1264 | DRAMP04371 | PDD-B-1 (PDD-B analog) | Antibacterial, Anti-Gram+, Anti-Gram-, Antimicrobial |
| 898 | 1265 | DRAMP04372 | PDD-B-2 (PDD-B analog) | Antibacterial, Anti-Gram+, Anti-Gram-, Antimicrobial |
| 899 | 1266 | DRAMP04373 | PDD-B-3 (PDD-B analog) | Antibacterial, Anti-Gram+, Anti-Gram-, Antimicrobial |
| 900 | 1267 | DRAMP04374 | PDD-B-4 (PDD-B analog) | Antibacterial, Anti-Gram+, Anti-Gram-, Antimicrobial |
| 901 | 1268 | DRAMP04376 | MP-1 (MP analog) | Antibacterial, Anti-Gram+, Anti-Gram-, Antimicrobial |

| S.no. | PepID | DRAMP_ID | Name of the AMP | Activity |
|---|---|---|---|---|
| 902 | 1269 | DRAMP04377 | MP-2 (MP analog) | Antibacterial, Anti-Gram+, Anti-Gram-, Antimicrobial |
| 903 | 1270 | DRAMP04378 | MP-5 (MP analog) | Antibacterial, Anti-Gram+, Anti-Gram-, Antimicrobial |
| 904 | 1271 | DRAMP04379 | MP-6 (MP analog) | Antibacterial, Anti-Gram+, Anti-Gram-, Antimicrobial |
| 905 | 1272 | DRAMP04380 | PMM-1 (PMM analog) | Antibacterial, Anti-Gram+, Anti-Gram-, Antimicrobial |
| 906 | 1273 | DRAMP04381 | PMM-2 (PMM analog) | Antibacterial, Anti-Gram+, Anti-Gram-, Antimicrobial |
| 907 | 1274 | DRAMP04382 | PMM-3 (PMM analog) | Antibacterial, Anti-Gram+, Anti-Gram-, Antimicrobial |
| 908 | 1275 | DRAMP04383 | PMM-4 (PMM analog) | Antibacterial, Anti-Gram+, Anti-Gram-, Antimicrobial |
| 909 | 1276 | DRAMP04385 | PMM-6 (PMM analog) | Antibacterial, Anti-Gram+, Anti-Gram-, Antimicrobial |
| 910 | 1277 | DRAMP04386 | PMM-7 (PMM analog) | Antibacterial, Anti-Gram+, Anti-Gram-, Antimicrobial |
| 911 | 1278 | DRAMP04387 | PMM-8 (PMM analog) | Antibacterial, Anti-Gram+, Anti-Gram-, Antimicrobial |
| 912 | 1279 | DRAMP04389 | PMM-10 (PMM analog) | Antibacterial, Anti-Gram+, Anti-Gram-, Antimicrobial |
| 913 | 1280 | DRAMP04390 | PMM-11 (PMM analog) | Antibacterial, Anti-Gram+, Anti-Gram-, Antimicrobial |
| 914 | 1281 | DRAMP04391 | PMM-12 (PMM analog) | Antibacterial, Anti-Gram+, Anti-Gram-, Antimicrobial |
| 915 | 1282 | DRAMP04392 | PMM-13 (PMM analog) | Antibacterial, Anti-Gram+, Anti-Gram-, Antimicrobial |
| 916 | 1283 | DRAMP04393 | PMM-14 (PMM analog) | Antibacterial, Anti-Gram+, Anti-Gram-, Antimicrobial |
| 917 | 1285 | DRAMP04542 | Polybia-MP-I (insects, vertebrates, animals) | Antibacterial, Anti-Gram+, Anti-Gram-, Antimicrobial |
| 918 | 1286 | DRAMP04543 | Polybia-MP-II (insects, vertebrates, animals) | Antibacterial, Cytotoxicity, Anti-Gram+, Anti-Gram-, Antimicrobial |
| 919 | 1287 | DRAMP04544 | Polybia-MP-III (insects, vertebrates, animals) | Antibacterial, Cytotoxicity, Anti-Gram+, Anti-Gram-, Antimicrobial |
| 920 | 1291 | DRAMP04640 | PGLa-AN2 | Antibacterial, Anti-Gram+, Anti-Gram-, Antimicrobial |
| 921 | 1292 | DRAMP04665 | Px-cec1 | Antibacterial, Antifungal, Anti-Gram+, Anti-Gram-, Antimicrobial |
| 922 | 1293 | DRAMP04670 | PBD1-42 | Antibacterial, Anti-Gram+, Anti-Gram-, Antimicrobial |
| 923 | 1294 | DRAMP04671 | Myticusin-1 | Antibacterial, Antifungal, Anti-Gram+, Anti-Gram-, Antimicrobial |
| 924 | 1295 | DRAMP04676 | Brevinin-2HS2A | Antibacterial, Antifungal, Anti-Gram+, Anti-Gram-, Antimicrobial |
| 925 | 1296 | DRAMP04677 | Brevinin-2HS2B | Antibacterial, Antifungal, Anti-Gram+, Anti-Gram-, Antimicrobial |
| 926 | 1346 | DRAMP00052 | Mutacin-2 (Mutacin II mutacin H-29B; Bacteriocin) | Antibacterial, Anti-Gram+, Anti-Gram-, Antimicrobial |
| 927 | 1354 | DRAMP00064 | Enterocin 96 (Bacteriocin) | Antibacterial, Anti-Gram+, Anti-Gram-, Antimicrobial |
| 928 | 1358 | DRAMP00070 | Laterosporulin (Bacteriocin) | Antibacterial, Anti-Gram+, Anti-Gram-, Antimicrobial |
| 929 | 1372 | DRAMP00085 | Bacteriocin | Antibacterial, Anti-Gram+, Anti-Gram-, Antimicrobial |
| 930 | 1434 | DRAMP00169 | Enterocin AS-48 (AS-48; Bacteriocin) | Antibacterial, Anti-Gram+, Anti-Gram-, Antimicrobial |
| 931 | 1438 | DRAMP18338 | Thiocillin GE37468 (Bacteriocin) | Antibacterial, Anti-Gram+, Anti-Gram-, Antimicrobial |
| 932 | 1444 | DRAMP00182 | Thuricin-S (Bacteriocin) | Antibacterial, Anti-Gram+, Anti-Gram-, Antimicrobial |
| 933 | 1574 | DRAMP00341 | Antifungal protein ginkbilobin-1 (Ginkbilobin, GNL; Plants) | Antibacterial, Antifungal, Antiviral, Anti-Gram+, Anti-Gram-, Antimicrobial |
| 934 | 1609 | DRAMP00393 | Hedyotide B2 (hB2; Uncyclotides; Plants) | Antifungal, Anti-Gram+, Anti-Gram-, Antimicrobial |
| 935 | 1612 | DRAMP00397 | Defensin D1 (Ns-D1; Plant defensin) | Antifungal, Anti-Gram+, Anti-Gram-, Antimicrobial |
| 936 | 1613 | DRAMP00398 | Defensin D2 (Ns-D2; Plant defensin) | Antifungal, Anti-Gram+, Anti-Gram-, Antimicrobial |
| 937 | 1617 | DRAMP00402 | Defensin D1 (So-D1; Antimicrobial peptide D1; Plant defensin) | Antibacterial, Anti-Gram+, Anti-Gram-, Antimicrobial |
| 938 | 1618 | DRAMP00403 | Defensin D2 (So-D2; Antimicrobial peptide D2; Plant defensin) | Antibacterial, Antifungal, Anti-Gram+, Anti-Gram-, Antimicrobial |
| 939 | 1621 | DRAMP00406 | Defensin D5 (So-D5; Antimicrobial peptide D5; Plant defensin) | Antibacterial, Antifungal, Anti-Gram+, Anti-Gram-, Antimicrobial |
| 940 | 1622 | DRAMP00407 | Defensin D6 (So-D6; Antimicrobial peptide D6; Plant defensin) | Antibacterial, Antifungal, Anti-Gram+, Anti-Gram-, Antimicrobial |
| 941 | 1624 | DRAMP00409 | Defensin-like protein (Sesquin; Plant defensin) | Antibacterial, Antifungal, Antiviral, Anti-Gram+, Anti-Gram-, Antimicrobial |
| 942 | 1661 | DRAMP00455 | Defensin-like protein 2 (Fabatin-2; Plant defensin) | Antibacterial, Anti-Gram+, Anti-Gram-, Antimicrobial |
| 943 | 1662 | DRAMP00456 | Defensin-like protein 1 (Fabatin-1; Plant defensin) | Antibacterial, Anti-Gram+, Anti-Gram-, Antimicrobial |
| 944 | 1952 | DRAMP00746 | Flower-specific defensin (NaD1; Plant defensin) | Antifungal, Anti-Gram+, Anti-Gram-, Antimicrobial |
| 945 | 1969 | DRAMP00767 | ChaC1 (Chassatide C1; Plant defensin) | Antibacterial, Anticancer, Anti-Gram+, Anti-Gram-, Antimicrobial |
| 946 | 1970 | DRAMP00768 | ChaC2 (Chassatide C2; Plant defensin) | Antibacterial, Anticancer, Anti-Gram+, Anti-Gram-, Antimicrobial |
| 947 | 1971 | DRAMP00769 | ChaC4 (Chassatide C4; Plant defensin) | Antibacterial, Anticancer, Anti-Gram+, Anti-Gram-, Antimicrobial |
| 948 | 1972 | DRAMP00770 | ChaC10 (Chassatide C10; Plant defensin) | Antibacterial, Anticancer, Anti-Gram+, Anti-Gram-, Antimicrobial |
| 949 | 1976 | DRAMP18325 | delta-lysin I (Bacteriocin) | Antibacterial, Anti-Gram+, Anti-Gram-, Antimicrobial |
| 950 | 1977 | DRAMP00796 | Cliotide T2 (cT2; Plant defensin) | Antibacterial, Anticancer, Anti-Gram+, Anti-Gram-, Antimicrobial |
| 951 | 1978 | DRAMP00797 | Cliotide T3 (cT3; Plant defensin) | Antibacterial, Anticancer, Anti-Gram+, Anti-Gram-, Antimicrobial |
| 952 | 2071 | DRAMP00937 | Tu-AMP1 (Plant defensin) | Antibacterial, Antifungal, Anti-Gram+, Anti-Gram-, Antimicrobial |
| 953 | 2072 | DRAMP00938 | Tu-AMP2 (Plant defensin) | Antibacterial, Antifungal, Anti-Gram+, Anti-Gram-, Antimicrobial |
| 954 | 2077 | DRAMP00957 | Pp-AMP1 (P. pubescens AMP1; Plant defensin) | Antibacterial, Antifungal, Anti-Gram+, Anti-Gram-, Antimicrobial |

| S.no. | PepID | DRAMP_ID | Name of the AMP | Activity |
|---|---|---|---|---|
| 955 | 2078 | DRAMP00958 | Pp-AMP2 (P. pubescens AMP2; Plant defensin) | Antibacterial, Antifungal, Anti-Gram+, Anti-Gram-, Antimicrobial |
| 956 | 2081 | DRAMP01380 | Odorranain-J1 (OdJ1; Frogs, amphibians, animals) | Antimicrobial, Antibacterial, Antifungal, Anti-Gram+, Anti- Gram-, |
| 957 | 2090 | DRAMP00980 | Antimicrobial peptide 1a (WAMP-1a; Plant defensin) | Antibacterial, Antifungal, Anti-Gram+, Anti-Gram-, Antimicrobial |
| 958 | 2091 | DRAMP00981 | Antimicrobial peptide 1b (WAMP-1b; Plant defensin) | Antibacterial, Antifungal, Anti-Gram+, Anti-Gram-, Antimicrobial |
| 959 | 2092 | DRAMP00982 | Fa-AMP1 (Fagopyrum antimicrobial peptide 1; hevein-type; Plant | Antibacterial, Antifungal, Anti-Gram+, Anti-Gram-, Antimicrobial |
| 960 | 2093 | DRAMP00983 | Fa-AMP2 (Fagopyrum antimicrobial peptide 2; hevein-type; Plant | Antibacterial, Antifungal, Anti-Gram+, Anti-Gram-, Antimicrobial |
| 961 | 2107 | DRAMP00997 | IB-AMP4 (IBAMP4; Basic peptide AMP4; Plants) | Antibacterial, Antifungal, Anti-Gram+, Anti-Gram-, Antimicrobial |
| 962 | 2108 | DRAMP00998 | Antimicrobial peptide MBP-1 (Maize Basic Peptide 1; Plant defensin) | Antibacterial, Antifungal, Anti-Gram+, Anti-Gram-, Antimicrobial |
| 963 | 2119 | DRAMP01010 | Lunatusin (Plants) | Antibacterial, Antifungal, Antiviral, Anti-Gram+, Anti-Gram-, Antimicrobial |
| 964 | 2123 | DRAMP01015 | VaD1 (Plant defensin) | Antibacterial, Antifungal, Anti-Gram+, Anti-Gram-, Antimicrobial |
| 965 | 2126 | DRAMP01022 | Cy-AMP1 (Plant defensin) | Antibacterial, Antifungal, Anti-Gram+, Anti-Gram-, Antimicrobial |
| 966 | 2127 | DRAMP01023 | Cy-AMP2 (Plant defensin) | Antibacterial, Antifungal, Anti-Gram+, Anti-Gram-, Antimicrobial |
| 967 | 2128 | DRAMP01024 | Cy-AMP3 (Plant defensin) | Antibacterial, Antifungal, Anti-Gram+, Anti-Gram-, Antimicrobial |
| 968 | 2187 | DRAMP01100 | Bombinin-like peptide 1 (Contains: Bombinin H; toads, amphibians, | Antibacterial, Anti-Gram+, Anti-Gram-, Antimicrobial |
| 969 | 2217 | DRAMP01140 | Uperin-2.2 (toads, amphibians, animals) | Antibacterial, Anti-Gram+, Anti-Gram-, Antimicrobial |
| 970 | 2250 | DRAMP01196 | Andersonin-G1 (Frogs, amphibians, animals) | Antibacterial, Antifungal, Anti-Gram+, Anti-Gram-, Antimicrobial |
| 971 | 2251 | DRAMP01197 | Andersonin-N1 (Frogs, amphibians, animals) | Antibacterial, Antifungal, Anti-Gram+, Anti-Gram-, Antimicrobial |
| 972 | 2252 | DRAMP01198 | Andersonin-Q1 (Frogs, amphibians, animals) | Antibacterial, Antifungal, Anti-Gram+, Anti-Gram-, Antimicrobial |
| 973 | 2255 | DRAMP01207 | Galensin (Frogs, amphibians, animals) | Antibacterial, Anti-Gram+, Anti-Gram-, Antimicrobial |
| 974 | 2258 | DRAMP01212 | Pleurain-A3 (Pleurain A3; Frogs, amphibians, animals) | Antibacterial, Antifungal, Anti-Gram+, Anti-Gram-, Antimicrobial |
| 975 | 2259 | DRAMP01213 | Pleurain-A4 (Pleurain A4; Frogs, amphibians, animals) | Antibacterial, Antifungal, Anti-Gram+, Anti-Gram-, Antimicrobial |
| 976 | 2264 | DRAMP01223 | Palustrin-2AJ2 (PL2AJ12; Frogs, amphibians, animals) | Antibacterial, Anti-Gram+, Anti-Gram-, Antimicrobial |
| 977 | 2265 | DRAMP01224 | Palustrin-2AR (Palustrin-2ARa; Ranatuerin-2SEa; Frogs, | Antibacterial, Anti-Gram+, Anti-Gram-, Antimicrobial |
| 978 | 2285 | DRAMP18404 | Polybia-MPII (mastoparan; insects, arthropods, invertebrates, | Antibacterial, antifungal, Anti-Gram+, Anti-Gram-, Antimicrobial |
| 979 | 2343 | DRAMP01338 | Amolopin-1a (Frogs, amphibians, animals) | Antibacterial, Antifungal, Anti-Gram+, Anti-Gram-, Antimicrobial |
| 980 | 2345 | DRAMP01342 | Amolopin-2b (Frogs, amphibians, animals) | Antibacterial, Antifungal, Anti-Gram+, Anti-Gram-, Antimicrobial |
| 981 | 2346 | DRAMP01343 | Amolopin-1c (Frogs, amphibians, animals) | Antibacterial, Antifungal, Anti-Gram+, Anti-Gram-, Antimicrobial |
| 982 | 2347 | DRAMP01344 | Amolopin-2c (Frogs, amphibians, animals) | Antibacterial, Antifungal, Anti-Gram+, Anti-Gram-, Antimicrobial |
| 983 | 2348 | DRAMP01345 | Amolopin-1d (Frogs, amphibians, animals) | Antibacterial, Antifungal, Anti-Gram+, Anti-Gram-, Antimicrobial |
| 984 | 2363 | DRAMP01444 | Nigrocin-1 (Frogs, amphibians, animals) | Antibacterial, Antifungal, Anti-Gram+, Anti-Gram-, Antimicrobial |
| 985 | 2364 | DRAMP01445 | Nigrocin-2 (Nigrocin-2LVa; Frogs, amphibians, animals) | Antibacterial, Antifungal, Anti-Gram+, Anti-Gram-, Antimicrobial |
| 986 | 2372 | DRAMP01463 | Esculentin-1SEa (Frogs, amphibians, animals) | Antibacterial, Anti-Gram+, Anti-Gram-, Antimicrobial |
| 987 | 2373 | DRAMP01464 | Esculentin-1SEb (Frogs, amphibians, animals) | Antibacterial, Anti-Gram+, Anti-Gram-, Antimicrobial |
| 988 | 2374 | DRAMP01465 | Esculentin-1R (Frogs, amphibians, animals) | Antibacterial, Anti-Gram+, Anti-Gram-, Antimicrobial |
| 989 | 2382 | DRAMP01489 | Esculentin-1A (Frogs, amphibians, animals) | Antibacterial, Anti-Gram+, Anti-Gram-, Antimicrobial |
| 990 | 2383 | DRAMP01492 | Esculentin-IIb (Frogs, amphibians, animals) | Antibacterial, Anti-Gram+, Anti-Gram-, Antimicrobial |
| 991 | 2388 | DRAMP18312 | Propionicin PLG-1(Bacteriocin) | Antibacterial, Antifungal, Anti-Gram+, Anti-Gram-, Antimicrobial |
| 992 | 2392 | DRAMP01519 | Esculentin-2PRa (Frogs, amphibians, animals) | Antibacterial, Antifungal, Anti-Gram+, Anti-Gram-, Antimicrobial |
| 993 | 2475 | DRAMP01671 | Dermaseptin-4 (DS IV; Dermaseptin-S4, DS4; Frogs, amphibians, | Antibacterial, Antifungal, Antiviral, Anti-Gram+, Anti-Gram-, Antimicrobial |
| 994 | 2503 | DRAMP01700 | Dermaseptin-H3 (Dermaseptin-like peptide 3, DMS3; Frogs, | Antibacterial, Anti-Gram+, Anti-Gram-, Antimicrobial |
| 995 | 2518 | DRAMP01717 | Dermatoxin (Frogs, amphibians, animals) | Antibacterial, Anti-Gram+, Anti-Gram-, Antimicrobial |
| 996 | 2530 | DRAMP02857 | Indolicidin (Cathelicidin-4; mammals, animals) | Antibacterial, Anti-Gram+, Anti-Gram-, Antimicrobial |
| 997 | 2532 | DRAMP02819 | Anoplin (Insects, arthropods, invertebrates, animals) | Antimicrobial, Antibacterial, Antifungal, Anti-Gram+, Anti-Gram-, |
| 998 | 2533 | DRAMP04395 | EP3 (Earthworm,animals) | Antibacterial, Anti-Gram+, Anti-Gram-, Antimicrobial |
| 999 | 2534 | DRAMP04394 | EP2 (Earthworm,animals) | Antibacterial, Anti-Gram+, Anti-Gram-, Antimicrobial |
| 1000 | 2548 | DRAMP01772 | Temporin-1Ob (Frogs, amphibians, animals) | Antibacterial, Antifungal, Anti-Gram+, Anti-Gram-, Antimicrobial |
| 1001 | 2560 | DRAMP01799 | Temporin-1PRa (Temporin 1PRa; Frogs, amphibians, animals) | Antibacterial, Anti-Gram+, Anti-Gram-, Antimicrobial |
| 1002 | 2561 | DRAMP01800 | Temporin-1PRb (Temporin 1PRb; Frogs, amphibians, animals) | Antibacterial, Antifungal, Anti-Gram+, Anti-Gram-, Antimicrobial |
| 1003 | 2562 | DRAMP01801 | Temporin-1DYa (Frogs, amphibians, animals) | Antibacterial, Anti-Gram+, Anti-Gram-, Antimicrobial |
| 1004 | 2563 | DRAMP01802 | Temporin-PTa (Frogs, amphibians, animals) | Antibacterial, Anti-Gram+, Anti-Gram-, Antimicrobial |
| 1005 | 2565 | DRAMP01804 | Temporin-CDYb (Brevinin-1CDYb; Frogs, amphibians, animals) | Antibacterial, Anti-Gram+, Anti-Gram-, Antimicrobial |
| 1006 | 2571 | DRAMP01387 | Odorranain-P2a (OdP2a; Frogs, amphibians, animals) | Antimicrobial, Antibacterial, Antifungal, Anti-Gram+, Anti- Gram-, |
| 1007 | 2572 | DRAMP01386 | Odorranain-P1a (OdP1a; Brevinin-1HS1; Brevinin-1-OA2; Frogs, | Antimicrobial, Antibacterial, Antifungal, Anti-Gram+, Anti- Gram-, |

| S.no. | PepID | DRAMP_ID | Name of the AMP | Activity |
|---|---|---|---|---|
| 1008 | 2573 | DRAMP01824 | Temporin-1Ja (Frogs, amphibians, animals) | Antibacterial, Anti-Gram+, Anti-Gram-, Antimicrobial |
| 1009 | 2576 | DRAMP01126 | Maximin-H4 (Toads, amphibians, animals) | Antimicrobial, Antibacterial, Antifungal, Anti-Gram+, Anti- Gram-, |
| 1010 | 2577 | DRAMP01125 | Maximin-H3 (Toads, amphibians, animals) | Antimicrobial, Antibacterial, Antifungal, Anti-Gram+, Anti- Gram-, |
| 1011 | 2621 | DRAMP01924 | Brevinin-1Ea (Frogs, amphibians, animals) | Antibacterial, Anti-Gram+, Anti-Gram-, Antimicrobial |
| 1012 | 2622 | DRAMP01925 | Brevinin-1Eb (Frogs, amphibians, animals) | Antibacterial, Anti-Gram+, Anti-Gram-, Antimicrobial |
| 1013 | 2623 | DRAMP01926 | Brevinin-1Ec (Frogs, amphibians, animals) | Antibacterial, Anti-Gram+, Anti-Gram-, Antimicrobial |
| 1014 | 2624 | DRAMP01927 | Brevinin-2Ea (Frogs, amphibians, animals) | Antibacterial, Anti-Gram+, Anti-Gram-, Antimicrobial |
| 1015 | 2625 | DRAMP01928 | Brevinin-2Eb (Frogs, amphibians, animals) | Antibacterial, Anti-Gram+, Anti-Gram-, Antimicrobial |
| 1016 | 2626 | DRAMP01929 | Brevinin-2Ec (Frogs, amphibians, animals) | Antibacterial, Anti-Gram+, Anti-Gram-, Antimicrobial |
| 1017 | 2628 | DRAMP01931 | Brevinin-2Ed (Frogs, amphibians, animals) | Antibacterial, Anti-Gram+, Anti-Gram-, Antimicrobial |
| 1018 | 2629 | DRAMP01932 | Brevinin-2Ee (Frogs, amphibians, animals) | Antibacterial, Anti-Gram+, Anti-Gram-, Antimicrobial |
| 1019 | 2631 | DRAMP01945 | Brevinin-1SE (Frogs, amphibians, animals) | Antibacterial, Anti-Gram+, Anti-Gram-, Antimicrobial |
| 1020 | 2632 | DRAMP01946 | Brevinin-20a (Frogs, amphibians, animals) | Antibacterial, Antifungal, Anti-Gram+, Anti-Gram-, Antimicrobial |
| 1021 | 2633 | DRAMP01947 | Brevinin-20b (Frogs, amphibians, animals) | Antibacterial, Antifungal, Anti-Gram+, Anti-Gram-, Antimicrobial |
| 1022 | 2634 | DRAMP18294 | lactococcin Z(Bacteriocin) | Antibacterial, Anti-Gram+, Anti-Gram-, Antimicrobial |
| 1023 | 2635 | DRAMP01952 | Brevinin-1PTb (Frogs, amphibians, animals) | Antibacterial, Anti-Gram+, Anti-Gram-, Antimicrobial |
| 1024 | 2636 | DRAMP01954 | Brevinin-2HSb (Frogs, amphibians, animals) | Antibacterial, Anti-Gram+, Anti-Gram-, Antimicrobial |
| 1025 | 2637 | DRAMP01958 | Brevinin-2PTd (Frogs, amphibians, animals) | Antibacterial, Anti-Gram+, Anti-Gram-, Antimicrobial |
| 1026 | 2638 | DRAMP01960 | Brevinin-1BYa (Frogs, amphibians, animals) | Antibacterial, Antifungal, Anti-Gram+, Anti-Gram-, Antimicrobial |
| 1027 | 2639 | DRAMP01961 | Brevinin-1BYb (Frogs, amphibians, animals) | Antibacterial, Antifungal, Anti-Gram+, Anti-Gram-, Antimicrobial |
| 1028 | 2642 | DRAMP01966 | Brevinin-1Ya (Frogs, amphibians, animals) | Antibacterial, Antifungal, Anti-Gram+, Anti-Gram-, Antimicrobial |
| 1029 | 2643 | DRAMP01967 | Brevinin-1Yb (Frogs, amphibians, animals) | Antibacterial, Antifungal, Anti-Gram+, Anti-Gram-, Antimicrobial |
| 1030 | 2647 | DRAMP01976 | Brevinin-2Eg (Frogs, amphibians, animals) | Antibacterial, Anti-Gram+, Anti-Gram-, Antimicrobial |
| 1031 | 2654 | DRAMP18291 | Garviecin LG34(Bacteriocin) | Antibacterial, Anti-Gram+, Anti-Gram-, Antimicrobial |
| 1032 | 2674 | DRAMP02012 | Brevinin-1T (Brevinin-2T; Frogs, amphibians, animals) | Antibacterial, Anti-Gram+, Anti-Gram-, Antimicrobial |
| 1033 | 2675 | DRAMP02013 | Brevinin-1Ta (Frogs, amphibians, animals) | Antibacterial, Anti-Gram+, Anti-Gram-, Antimicrobial |
| 1034 | 2678 | DRAMP02016 | Brevinin-1DYa (Frogs, amphibians, animals) | Antibacterial, Anti-Gram+, Anti-Gram-, Antimicrobial |
| 1035 | 2679 | DRAMP02017 | Brevinin-2DYa (Frogs, amphibians, animals) | Antibacterial, Anti-Gram+, Anti-Gram-, Antimicrobial |
| 1036 | 2680 | DRAMP02018 | Brevinin-1DYb (Brevinin-1CDYb; Frogs, amphibians, animals) | Antibacterial, Anti-Gram+, Anti-Gram-, Antimicrobial |
| 1037 | 2681 | DRAMP02020 | Brevinin-1DYc (Frogs, amphibians, animals) | Antibacterial, Anti-Gram+, Anti-Gram-, Antimicrobial |
| 1038 | 2684 | DRAMP18281 | Plantaricin KL-1Y (Bacteriocin) | Antibacterial, Anti-Gram+, Anti-Gram-, Antimicrobial |
| 1039 | 2694 | DRAMP02089 | Brevinin-1La (Brevinin-1PRd; Frogs, amphibians, animals) | Antibacterial, Anti-Gram+, Anti-Gram-, Antimicrobial |
| 1040 | 2695 | DRAMP01124 | Maximin-H2 (Toads, amphibians, animals) | Antimicrobial, Antibacterial, Antifungal, Anti-Gram+, Anti-Gram-, |
| 1041 | 2696 | DRAMP01123 | Maximin-H1 (Toads, amphibians, animals) | Antimicrobial, Antibacterial, Antifungal, Anti-Gram+, Anti-Gram-, |
| 1042 | 2697 | DRAMP01111 | Maximin-5 (Toads, amphibians, animals) | Antimicrobial, Antibacterial, Antifungal, Antiviral, Anticancer, Anti-Gram+, Anti-Gram-, |
| 1043 | 2698 | DRAMP01110 | Maximin-4 (Toads, amphibians, animals) | Antimicrobial, Antibacterial, Antifungal, Antiviral, Anticancer, Anti-Gram+, Anti-Gram-, |
| 1044 | 2699 | DRAMP01109 | Maximin-3 (Toads, amphibians, animals) | Antimicrobial, Antibacterial, Antifungal, Antiviral, Anticancer, Anti-Gram+, Anti-Gram-, |
| 1045 | 2700 | DRAMP02100 | Brevinin-1Pe (Frogs, amphibians, animals) | Antibacterial, Antifungal, Anti-Gram+, Anti-Gram-, Antimicrobial |
| 1046 | 2706 | DRAMP18272 | Enterocin AS-48RJ (Bacteriocin) | Antibacterial, Anti-Gram+, Anti-Gram-, Antimicrobial |
| 1047 | 2718 | DRAMP02132 | Antimicrobial peptide 3 (XT-3; Levitide-like peptide; Frogs, | Antibacterial, Anti-Gram+, Anti-Gram-, Antimicrobial |
| 1048 | 2719 | DRAMP02134 | Antimicrobial peptide 5 (XT-5; PGLa-like peptide; Frogs, amphibians, | Antibacterial, Antifungal, Anti-Gram+, Anti-Gram-, Antimicrobial |
| 1049 | 2806 | DRAMP02243 | Ranatuerin-1T (Brevinin-2T; Frogs, amphibians, animals) | Antibacterial, Anti-Gram+, Anti-Gram-, Antimicrobial |
| 1050 | 2810 | DRAMP02249 | Ranatuerin-2SEB (Frogs, amphibians, animals) | Antibacterial, Anti-Gram+, Anti-Gram-, Antimicrobial |
| 1051 | 2811 | DRAMP02250 | Ranatuerin-2SEC (Frogs, amphibians, animals) | Antibacterial, Anti-Gram+, Anti-Gram-, Antimicrobial |
| 1052 | 2813 | DRAMP02258 | Ranatuerin-IIbYa (Ranatuerin-2bYa; Frogs, amphibians, animals) | Antibacterial, Anti-Gram+, Anti-Gram-, Antimicrobial |
| 1053 | 2818 | DRAMP02284 | Pseudin-1 (Pseudin 1; Frogs, amphibians, animals) | Antibacterial, Antifungal, Anti-Gram+, Anti-Gram-, Antimicrobial |
| 1054 | 2819 | DRAMP02285 | Pseudin-2 (Pseudin 2; Frogs, amphibians, animals) | Antibacterial, Antifungal, Anti-Gram+, Anti-Gram-, Antimicrobial |
| 1055 | 2820 | DRAMP02286 | Pseudin-3 (Pseudin 3; Frogs, amphibians, animals) | Antibacterial, Antifungal, Anti-Gram+, Anti-Gram-, Antimicrobial |
| 1056 | 2821 | DRAMP02287 | Pseudin-4 (Pseudin 4; Frogs, amphibians, animals) | Antibacterial, Antifungal, Anti-Gram+, Anti-Gram-, Antimicrobial |
| 1057 | 2841 | DRAMP02332 | Piscidin-3 (Pis-3; fish, chordates, animals) | Antibacterial, Antiviral, Anti-Gram+, Anti-Gram-, Antimicrobial |
| 1058 | 2851 | DRAMP01375 | Odorranain-E1 (OdE1; Frogs, amphibians, animals) | Antimicrobial, Antibacterial, Antifungal, Anti-Gram+, Anti- Gram-, |
| 1059 | 2854 | DRAMP02353 | Pleurocidin-like peptide WFX (fish, chordates, animals; Predicted) | Antibacterial, Antifungal, Anti-Gram+, Anti-Gram-, Antimicrobial |
| 1060 | 2868 | DRAMP02245 | Ranatuerin-2Cb (Ranatuerin 2Cb; Frogs, amphibians, animals) | Antimicrobial, Antibacterial, Antifungal, Anti-Gram+, Anti-Gram-, |

| S.no. | PepID | DRAMP_ID | Name of the AMP | Activity |
|---|---|---|---|---|
| 1061 | 2869 | DRAMP02398 | Antimicrobial peptide GP-19 (GP-19) | Antibacterial, Antifungal, Anti-Gram+, Anti-Gram-, Antimicrobial |
| 1062 | 2876 | DRAMP02407 | Napin-like polypeptide (Contains: Napin-like polypeptide small chain | Antibacterial, Anti-Gram+, Anti-Gram-, Antimicrobial |
| 1063 | 2890 | DRAMP02437 | Papillosin | Antibacterial, Anti-Gram+, Anti-Gram-, Antimicrobial |
| 1064 | 2891 | DRAMP02438 | Halocyntin | Antibacterial, Anti-Gram+, Anti-Gram-, Antimicrobial |
| 1065 | 2896 | DRAMP02447 | Antimicrobial protein 2 (Antimicrobial protein AN5-2) | Antibacterial, Anti-Gram+, Anti-Gram-, Antimicrobial |
| 1066 | 2901 | DRAMP02453 | S. litura moricin (Sl moricin; Insects, animals) | Antibacterial, Anti-Gram+, Anti-Gram-, Antimicrobial |
| 1067 | 2902 | DRAMP02454 | Theromacin (Arthropods, animals) | Antibacterial, Anti-Gram+, Anti-Gram-, Antimicrobial |
| 1068 | 2904 | DRAMP02457 | L-amino-acid oxidase (Balt-LAAO-I; LAAO; LAO; snakes, reptils, | Antibacterial, Anti-Gram+, Anti-Gram-, Antimicrobial |
| 1069 | 2909 | DRAMP02462 | L-amino-acid oxidase (LAAO; LAO; snakes, reptils, animals) | Antibacterial, Antiparasitic, Anti-Gram+, Anti-Gram-, Antimicrobial |
| 1070 | 2951 | DRAMP02511 | Crotamine (defensin-like toxin; Snakes, reptiles, animals) | Antibacterial, Antifungal, Cytotoxicity, Anti-Gram+, Anti-Gram-, Antimicrobial |
| 1071 | 2999 | DRAMP18253 | Piscicocin CS526(Bacteriocin) | Antibacterial, Anti-Gram+, Anti-Gram-, Antimicrobial |
| 1072 | 3003 | DRAMP02570 | Penaeidin-1 (Pen-1; shrimps, Arthropods, animals) | Antibacterial, Antifungal, Anti-Gram+, Anti-Gram-, Antimicrobial |
| 1073 | 3004 | DRAMP02571 | Penaeidin-2a (Pen-2a; shrimps, Arthropods, animals) | Antibacterial, Antifungal, Anti-Gram+, Anti-Gram-, Antimicrobial |
| 1074 | 3006 | DRAMP02575 | Penaeidin-3b (Pen-3b; shrimps, Arthropods, animals) | Antibacterial, Antifungal, Anti-Gram+, Anti-Gram-, Antimicrobial |
| 1075 | 3007 | DRAMP02576 | Penaeidin-3c (Pen-3c; shrimps, Arthropods, animals) | Antibacterial, Antifungal, Anti-Gram+, Anti-Gram-, Antimicrobial |
| 1076 | 3028 | DRAMP02602 | Clavaspirin (chordates, animals) | Antibacterial, Anti-Gram+, Anti-Gram-, Antimicrobial |
| 1077 | 3050 | DRAMP18250 | Laterosporulin (Bacteriocin) | Antibacterial, Anti-Gram+, Anti-Gram-, Antimicrobial |
| 1078 | 3052 | DRAMP18248 | Bifidin I(Bacteriocin) | Antibacterial, Anti-Gram+, Anti-Gram-, Antimicrobial |
| 1079 | 3053 | DRAMP18249 | Bac-GM100 (Bacteriocin) | Antibacterial, Antifungal, Anti-Gram+, Anti-Gram-, Antimicrobial |
| 1080 | 3059 | DRAMP18247 | Bacthuricin F4(Bacteriocin) | Antibacterial, Anti-Gram+, Anti-Gram-, Antimicrobial |
| 1081 | 3065 | DRAMP02642 | Rhesus theta-defensin 1 (RTD-1; primates, mammals, animals) | Antibacterial, Antifungal, Antiviral, Anti-Gram+, Anti-Gram-, Antimicrobial |
| 1082 | 3066 | DRAMP02643 | Rhesus theta-defensin 2 (RTD-2; primates, mammals, animals) | Antibacterial, Antifungal, Anti-Gram+, Anti-Gram-, Antimicrobial |
| 1083 | 3067 | DRAMP02644 | Rhesus theta-defensin 3 (RTD-3; primates, mammals, animals) | Antibacterial, Antiviral, Anti-Gram+, Anti-Gram-, Antimicrobial |
| 1084 | 3075 | DRAMP02653 | Neutrophil defensin 1 (RMAD-1; primates, mammals, animals) | Antibacterial, Antifungal, Anti-Gram+, Anti-Gram-, Antimicrobial |
| 1085 | 3076 | DRAMP02654 | Neutrophil defensin 2 (RMAD-2; primates, mammals, animals) | Antibacterial, Antifungal, Anti-Gram+, Anti-Gram-, Antimicrobial |
| 1086 | 3081 | DRAMP02659 | Neutrophil defensin 3 (RMAD-3; primates, mammals, animals) | Antibacterial, Antifungal, Anti-Gram+, Anti-Gram-, Antimicrobial |
| 1087 | 3082 | DRAMP02660 | Neutrophil defensin 4 (RMAD-4; primates, mammals, animals) | Antibacterial, Antifungal, Anti-Gram+, Anti-Gram-, Antimicrobial |
| 1088 | 3083 | DRAMP02661 | Neutrophil defensin 5 (RMAD-5; primates, mammals, animals) | Antibacterial, Antifungal, Anti-Gram+, Anti-Gram-, Antimicrobial |
| 1089 | 3084 | DRAMP02662 | Neutrophil defensin 6 (RMAD-6; primates, mammals, animals) | Antibacterial, Antifungal , Anti-Gram+, Anti-Gram-, Antimicrobial |
| 1090 | 3085 | DRAMP02663 | Neutrophil defensin 7 (RMAD-7; primates, mammals, animals) | Antibacterial, Antifungal, Antimicrobial, Anti-Gram+, Anti-Gram-, |
| 1091 | 3119 | DRAMP02698 | Rhesus macaque oral alpha-defensins (ROADs; primates, mammals, | Antibacterial, Antifungal, Anti-Gram+, Anti-Gram-, Antimicrobial |
| 1092 | 3128 | DRAMP18242 | Fengycin B2 (Bacteriocin) | Antibacterial, Antifungal, Anti-Gram+, Anti-Gram-, Antimicrobial |
| 1093 | 3130 | DRAMP18240 | Fengycin C(Bacteriocin) | Antibacterial, Antifungal, Anti-Gram+, Anti-Gram-, Antimicrobial |
| 1094 | 3131 | DRAMP18241 | Subtilomycin(Bacteriocin) | Antibacterial, Anti-Gram+, Anti-Gram-, Antimicrobial |
| 1095 | 3132 | DRAMP18239 | Fengycin A2(Bacteriocin) | Antibacterial, Antifungal, Anti-Gram+, Anti-Gram-, Antimicrobial |
| 1096 | 3133 | DRAMP18238 | Fengycin B(Bacteriocin) | Antibacterial, Antifungal, Anti-Gram+, Anti-Gram-, Antimicrobial |
| 1097 | 3144 | DRAMP18237 | Fengycin A(Bacteriocin) | Antibacterial, Antifungal, Anti-Gram+, Anti-Gram-, Antimicrobial |
| 1098 | 3157 | DRAMP02739 | TEWP (turtle egg-white protein; Reptiles, animals) | Antibacterial, Antiviral, Anti-Gram+, Anti-Gram-, Antimicrobial |
| 1099 | 3158 | DRAMP02741 | Pelovaterin (defensin-like AMP; Gly-rich; Reptiles, animals) | Antibacterial, Anti-Gram+, Anti-Gram-, Antimicrobial |
| 1100 | 3159 | DRAMP02742 | Defensin-like turtle egg white protein TEWP (TEWP; Reptiles, | Antibacterial, Antiviral, Anti-Gram+, Anti-Gram-, Antimicrobial |
| 1101 | 3168 | DRAMP03743 | Androctonin (Arthropods, animals) | Antimicrobial, Antibacterial, Antifungal, Anti-Gram+, Anti- Gram-, |
| 1102 | 3171 | DRAMP02755 | Ponericin G3 (ants, insects, animals) | Antibacterial, Antifungal, Anti-Gram+, Anti-Gram-, Antimicrobial |
| 1103 | 3172 | DRAMP02756 | Ponericin G4 (ants, insects, animals) | Antibacterial, Antifungal, Anti-Gram+, Anti-Gram-, Antimicrobial |
| 1104 | 3174 | DRAMP02758 | Ponericin G6 (ants, insects, animals) | Antibacterial, Antifungal, Anti-Gram+, Anti-Gram-, Antimicrobial |
| 1105 | 3176 | DRAMP02760 | Ponericin-L1 (ants, insects, animals) | Antibacterial, Anti-Gram+, Anti-Gram-, Antimicrobial |
| 1106 | 3177 | DRAMP02761 | Ponericin-L2 (ants, insects, animals) | Antibacterial, Antiviral, Anti-Gram+, Anti-Gram-, Antimicrobial |
| 1107 | 3178 | DRAMP02762 | Ponericin-W1 (ants, insects, animals) | Antibacterial, Antifungal, Insecticidal, Anti-Gram+, Anti-Gram-, Antimicrobial |
| 1108 | 3180 | DRAMP02764 | Ponericin-W3 (ants, insects, animals) | Antibacterial, Antifungal, Insecticidal, Anti-Gram+, Anti- Gram-, Antimicrobial |
| 1109 | 3181 | DRAMP02765 | Ponericin-W4 (ants, insects, animals) | Antibacterial, Antifungal, Insecticidal, Anti-Gram+, Anti- Gram-, Antimicrobial |
| 1110 | 3182 | DRAMP02766 | Ponericin-W5 (ants, insects, animals) | Antibacterial, Antifungal, Insecticidal, Anti-Gram+, Anti-Gram-, Antimicrobial |
| 1111 | 3183 | DRAMP02767 | Ponericin-W6 (ants, insects, animals) | Antibacterial, Anti-Gram+, Anti-Gram-, Antimicrobial |
| 1112 | 3184 | DRAMP02770 | Pilosulin 3 (ants, insects, animals) | Antibacterial, Anti-Gram+, Anti-Gram-, Antimicrobial |
| 1113 | 3185 | DRAMP02771 | Pilosulin 4 (ants, insects, animals) | Antibacterial, Anti-Gram+, Anti-Gram-, Antimicrobial |

| S.no. | PepID | DRAMP_ID | Name of the AMP | Activity |
|---|---|---|---|---|
| 1114 | 3189 | DRAMP02781 | Coleoptericin (Insects, animals) | Antibacterial, Anti-Gram+, Anti-Gram-, Antimicrobial |
| 1115 | 3191 | DRAMP02783 | Peptide C (Insects, animals) | Antibacterial, Anti-Gram+, Anti-Gram-, Antimicrobial |
| 1116 | 3210 | DRAMP02803 | Mytilin-A (molluscas, animals) | Antibacterial, Anti-Gram+, Anti-Gram-, Antimicrobial |
| 1117 | 3211 | DRAMP02804 | Mytilus defensin-B (molluscas, animals) | Antibacterial, Anti-Gram+, Anti-Gram-, Antimicrobial |
| 1118 | 3219 | DRAMP18235 | Gageopeptide D(Bacteriocin) | Antifungal, Antibacterial, Anti-Gram+, Anti-Gram-, Antimicrobial |
| 1119 | 3220 | DRAMP02816 | Lumbrican | Antibacterial, Antifungal, Anti-Gram+, Anti-Gram-, Antimicrobial |
| 1120 | 3221 | DRAMP02818 | Dicynthaurin | Antibacterial, Anti-Gram+, Anti-Gram-, Antimicrobial |
| 1121 | 3234 | DRAMP02832 | Reactive oxygen species modulator 1 (ROS modulator 1; mammals, | Antibacterial, Anti-Gram+, Anti-Gram-, Antimicrobial |
| 1122 | 3239 | DRAMP18233 | Gageopeptide B(Bacteriocin) | Antifungal, Antibacterial, Anti-Gram+, Anti-Gram-, Antimicrobial |
| 1123 | 3240 | DRAMP18234 | Gageopeptide C(Bacteriocin) | Antifungal, Antibacterial, Anti-Gram+, Anti-Gram-, Antimicrobial |
| 1124 | 3249 | DRAMP01356 | Ranalexin-1Ca (Ranatuerin 1Ca; Frogs, amphibians, animals) | Antimicrobial, Antibacterial, Antifungal, Anti-Gram+, Anti-Gram-, |
| 1125 | 3250 | DRAMP02864 | Bovine Beta-defensin 7 (bBD-7; BNBD-7; BNDB-7; mammals, | Antibacterial, Anti-Gram+, Anti-Gram-, Antimicrobial |
| 1126 | 3251 | DRAMP02871 | Beta-defensin 119 (Defensin, beta 119; mammals, animals) | Antibacterial, Anti-Gram+, Anti-Gram-, Antimicrobial |
| 1127 | 3281 | DRAMP02914 | Cathelicidin-1 (Bactenecin-1, Bac1; Cyclic dodecapeptide; mammals, | Antibacterial, Anti-Gram+, Anti-Gram-, Antimicrobial |
| 1128 | 3291 | DRAMP02929 | Antimicrobial protein 1 | Antibacterial, Anti-Gram+, Anti-Gram-, Antimicrobial |
| 1129 | 3292 | DRAMP02930 | Antimicrobial protein 2 (crabs, Arthropods, animals) | Antibacterial, Anti-Gram+, Anti-Gram-, Antimicrobial |
| 1130 | 3294 | DRAMP02936 | Big defensin (crabs, Arthropods, animals) | Antibacterial, Antifungal, Anti-Gram+, Anti-Gram-, Antimicrobial |
| 1131 | 3295 | DRAMP02937 | Tachycitin (crabs, Arthropods, animals) | Antibacterial, Antifungal, Anti-Gram+, Anti-Gram-, Antimicrobial |
| 1132 | 3296 | DRAMP18230 | Gageotetrin B (Bacteriocin) | Antifungal, Antibacterial, Anti-Gram+, Anti-Gram-, Antimicrobial |
| 1133 | 3297 | DRAMP18231 | Gageotetrin C (Bacteriocin) | Antifungal, Antibacterial, Anti-Gram+, Anti-Gram-, Antimicrobial |
| 1134 | 3298 | DRAMP18232 | Gageopeptide A(Bacteriocin) | Antifungal, Antibacterial, Anti-Gram+, Anti-Gram-, Antimicrobial |
| 1135 | 3301 | DRAMP02941 | Tachystatin-A1 (crabs, Arthropods, animals) | Antibacterial, Antifungal, Anti-Gram+, Anti-Gram-, Antimicrobial |
| 1136 | 3302 | DRAMP02942 | Tachystatin-A2 (crabs, Arthropods, animals) | Antibacterial, Antifungal, Anti-Gram+, Anti-Gram-, Antimicrobial |
| 1137 | 3305 | DRAMP02945 | Tachystatin-C (crabs, Arthropods, animals) | Antibacterial, Antifungal, Anti-Gram+, Anti-Gram-, Antimicrobial |
| 1138 | 3308 | DRAMP18228 | Gageostatin C (Bacteriocin) | Antibacterial, antifungal, Anti-Gram+, Anti-Gram-, Antimicrobial |
| 1139 | 3309 | DRAMP18229 | Gageotetrin A (Bacteriocin) | Antifungal, Antibacterial, Anti-Gram+, Anti-Gram-, Antimicrobial |
| 1140 | 3312 | DRAMP02955 | Hedistin (marine annelid, Metazoa) | Antibacterial, Anti-Gram+, Anti-Gram-, Antimicrobial |
| 1141 | 3316 | DRAMP02968 | Prophenin-1 (C6, PF-1; Pro-rich; pigs, mammals, animals) | Antibacterial, Anti-Gram+, Anti-Gram-, Antimicrobial |
| 1142 | 3317 | DRAMP02969 | Prophenin-2 (C12, PF-2, PR-2; Pro-rich; pigs, mammals, animals) | Antibacterial, Anti-Gram+, Anti-Gram-, Antimicrobial |
| 1143 | 3318 | DRAMP02971 | Protegrin-2 (Protegrin 2; PG-2; pigs, mammals, animals) | Antibacterial, Antifungal, Anti-Gram+, Anti-Gram-, Antimicrobial |
| 1144 | 3319 | DRAMP02972 | Protegrin-3 (Protegrin 3; PG-3; pigs, mammals, animals) | Antibacterial, Antifungal, Anti-Gram+, Anti-Gram-, Antimicrobial |
| 1145 | 3322 | DRAMP02976 | Beta-defensin 1 (BD-1; Defensin, beta 1; pigs, mammals, animals) | Antibacterial, Anti-Gram+, Anti-Gram-, Antimicrobial |
| 1146 | 3327 | DRAMP02982 | Reactive oxygen species modulator 1 (ROS modulator 1; pigs, | Antibacterial, Anti-Gram+, Anti-Gram-, Antimicrobial |
| 1147 | 3329 | DRAMP02984 | Neutrophil cationic antibacterial polypeptide of 11 kDa (CAP11; pigs, | Antibacterial, Anti-Gram+, Anti-Gram-, Antimicrobial |
| 1148 | 3330 | DRAMP02985 | Neutrophil cationic peptide 2 (CP-2; GNCP-2; pigs, mammals, | Antibacterial, Antifungal, Antiviral, Anti-Gram+, Anti-Gram-, Antimicrobial |
| 1149 | 3331 | DRAMP02986 | Neutrophil cationic peptide 1 (GPNP; Antiviral defensin; pigs, | Antibacterial, Antifungal, Antiviral, Anti-Gram+, Anti-Gram-, Antimicrobial |
| 1150 | 3334 | DRAMP02989 | Lasioglossin LL-I (Insects, animals) | Antibacterial, Anticancer, Anti-Gram+, Anti-Gram-, Antimicrobial |
| 1151 | 3335 | DRAMP02990 | Lasioglossin LL-II (Insects, animals) | Antibacterial, Anticancer, Anti-Gram+, Anti-Gram-, Antimicrobial |
| 1152 | 3336 | DRAMP02991 | Lasioglossin LL-III (Insects, animals) | Antibacterial, Anticancer, Anti-Gram+, Anti-Gram-, Antimicrobial |
| 1153 | 3348 | DRAMP03023 | Mastoparan (Protonectarina-MP; Insects, animals) | Antibacterial, Antifungal, Anti-Gram+, Anti-Gram-, Antimicrobial |
| 1154 | 3349 | DRAMP18227 | Gageostatin B (Bacteriocin) | Antibacterial, antifungal, Anti-Gram+, Anti-Gram-, Antimicrobial |
| 1155 | 3355 | DRAMP03049 | Eumenine mastoparan-OD (EMP-OD; Venom peptide 1, OdVP1; | Antibacterial, Antifungal, Anti-Gram+, Anti-Gram-, Antimicrobial |
| 1156 | 3366 | DRAMP18226 | Gageostatin A (Bacteriocin) | Antibacterial, antifungal, Anti-Gram+, Anti-Gram-, Antimicrobial |
| 1157 | 3369 | DRAMP03073 | Sapecin-C (Sapecin C; defensins; Insects, animals) | Antibacterial, Anti-Gram+, Anti-Gram-, Antimicrobial |
| 1158 | 3378 | DRAMP03083 | Sapecin-B (defensins; Insects, animals) | Antibacterial, Anti-Gram+, Anti-Gram-, Antimicrobial |
| 1159 | 3379 | DRAMP03084 | Ceratotoxin-B (Insects, animals) | Antibacterial, Anti-Gram+, Anti-Gram-, Antimicrobial |
| 1160 | 3380 | DRAMP03085 | Ceratotoxin-A (Insects, animals) | Antibacterial, Anti-Gram+, Anti-Gram-, Antimicrobial |
| 1161 | 3381 | DRAMP03086 | Ceratotoxin-D (Insects, animals) | Antibacterial, Anti-Gram+, Anti-Gram-, Antimicrobial |
| 1162 | 3400 | DRAMP18223 | Sonorensin(Bacteriocin) | Antibacterial, Anti-Gram+, Anti-Gram-, Antimicrobial |
| 1163 | 3418 | DRAMP03134 | Defensin-D (AaeDefD; Insects, animals) | Antibacterial, Anti-Gram+, Anti-Gram-, Antimicrobial |
| 1164 | 3436 | DRAMP03157 | Halocidin subunit B | Antibacterial, Anti-Gram+, Anti-Gram-, Antimicrobial |
| 1165 | 3437 | DRAMP03158 | Halocidin subunit A (invertebrates, animals; Preclinical) | Antibacterial, Anti-Gram+, Anti-Gram-, Antimicrobial |
| 1166 | 3441 | DRAMP03163 | R. prolixus defensin A (RprDefA; insect defensin; Insects, animals) | Antibacterial, Insecticidal, Anti-Gram+, Anti-Gram-, Antimicrobial |

| S.no. | PepID | DRAMP_ID | Name of the AMP | Activity |
|---|---|---|---|---|
| 1167 | 3442 | DRAMP03164 | R. prolixus defensin B (RprDefB; insect defensin; Insects, animals) | Antibacterial, Insecticidal, Anti-Gram+, Anti-Gram-, Antimicrobial |
| 1168 | 3443 | DRAMP03165 | R. prolixus defensin C (RprDefC; insect defensin; Insects, animals) | Antibacterial, Insecticidal, Anti-Gram+, Anti-Gram-, Antimicrobial |
| 1169 | 3449 | DRAMP03174 | Arenicin-2 (Ar-2; marine polychaeta, animals) | Antibacterial, Antifungal, Anti-Gram+, Anti-Gram-, Antimicrobial |
| 1170 | 3450 | DRAMP03175 | Perinerin | Antibacterial, Antifungal, Anti-Gram+, Anti-Gram-, Antimicrobial |
| 1171 | 3454 | DRAMP03180 | ASABF-alpha (ASABF; nematodes, animals) | Antibacterial, Antifungal, Anti-Gram+, Anti-Gram-, Antimicrobial |
| 1172 | 3456 | DRAMP03183 | Naegleriapore A | Antibacterial, Antiprotozoal, Cytotoxicity, Anti-Gram+, Anti-Gram-, Antimicrobial |
| 1173 | 3457 | DRAMP03184 | Naegleriapore B | Antibacterial, Antiprotozoal, Cytotoxicity, Anti-Gram+, Anti-Gram-, Antimicrobial |
| 1174 | 3465 | DRAMP03199 | PhD1 (PhD-1; Defensin-1; primates, mammals, animals) | Antibacterial, Antifungal, Anti-Gram+, Anti-Gram-, Antimicrobial |
| 1175 | 3466 | DRAMP03200 | PhD2 (PhD-2; Defensin-2; primates, mammals, animals) | Antibacterial, Antifungal, Anti-Gram+, Anti-Gram-, Antimicrobial |
| 1176 | 3467 | DRAMP03201 | PhD3 (PhD-3; Defensin-3; primates, mammals, animals) | Antibacterial, Antifungal, Anti-Gram+, Anti-Gram-, Antimicrobial |
| 1177 | 3473 | DRAMP03208 | BTD-1 (theta-defensin; primates, mammals, animals) | Antibacterial, Antifungal, Anti-Gram+, Anti-Gram-, Antimicrobial |
| 1178 | 3474 | DRAMP03209 | BTD-2 (theta-defensin; primates, mammals, animals) | Antibacterial, Antifungal, Anti-Gram+, Anti-Gram-,Antimicrobial |
| 1179 | 3476 | DRAMP03211 | BTD-4 (theta-defensin; primates, mammals, animals) | Antibacterial, Antifungal, Anti-Gram+, Anti-Gram-, Antimicrobial |
| 1180 | 3477 | DRAMP03212 | BTD-7 (theta-defensin; primates, mammals, animals) | Antibacterial, Antifungal, Anti-Gram+, Anti-Gram-, Antimicrobial |
| 1181 | 3485 | DRAMP03224 | M-ctenitoxin-Cs1c (M-CNTX-Cs1c; Cupiennin-1c; spiders, | Antibacterial, Anti-Gram+, Anti-Gram-, Antimicrobial |
| 1182 | 3489 | DRAMP03238 | M-zodatoxin-Lt8c (M-ZDTX-Lt8c; Cytoinsectotoxin-1c, CIT-1c; | Antibacterial, Insecticidal, Anti-Gram+, Anti-Gram-, Antimicrobial |
| 1183 | 3514 | DRAMP18213 | Gramicidin S(Bacteriocin) | Antibacterial, Antifungal, Anti-Gram+, Anti-Gram-, Antimicrobial |
| 1184 | 3525 | DRAMP03282 | Turkey Heterophil Peptide 1 (Antimicrobial peptide THP1; Birds, | Antibacterial, Anti-Gram+, Anti-Gram-, Antimicrobial |
| 1185 | 3543 | DRAMP03304 | Hydramacin-1 (Hm-1; annelida, animals) | Antibacterial, Anti-Gram+, Anti-Gram-, Antimicrobial |
| 1186 | 3554 | DRAMP03317 | Cathelicidin-related antimicrobial peptide (AMPs) | Antibacterial, Anti-Gram+, Anti-Gram-, Antimicrobial |
| 1187 | 3565 | DRAMP03329 | Alpha-defensin cryptdin-1 (Crp1; Rodents, mammals, animals) | Antibacterial, Anti-Gram+, Anti-Gram-, Antimicrobial |
| 1188 | 3566 | DRAMP03330 | Alpha-defensin cryptdin-2 (Defensin-related cryptdin-2; Rodents, | Antibacterial, Anti-Gram+, Anti-Gram-, Antimicrobial |
| 1189 | 3567 | DRAMP03331 | Alpha-defensin cryptdin-3 (Defensin-related cryptdin-3; Rodents, | Antibacterial, Anti-Gram+, Anti-Gram-, Antimicrobial |
| 1190 | 3568 | DRAMP03332 | Alpha-defensin cryptdin-4 (Defensin-related cryptdin4; Rodents, | Antibacterial, Anti-Gram+, Anti-Gram-, Antimicrobial |
| 1191 | 3569 | DRAMP03333 | Alpha-defensin cryptdin-5 (Defensin-related cryptdin5; Rodents, | Antibacterial, Anti-Gram+, Anti-Gram-, Antimicrobial |
| 1192 | 3570 | DRAMP03334 | Rodents, mammals, animals) | Antibacterial, Anti-Gram+, Anti-Gram-, Antimicrobial |
| 1193 | 3593 | DRAMP03357 | Cryptdin related sequence peptide (CRS4C-1a; Rodents, mammals, | Antibacterial, Anti-Gram+, Anti-Gram-, Antimicrobial |
| 1194 | 3594 | DRAMP03358 | Cryptdin related sequence peptide (CRS4C-1d; Rodents, mammals, | Antibacterial, Anti-Gram+, Anti-Gram-, Antimicrobial |
| 1195 | 3595 | DRAMP03359 | Cryptdin related sequence peptide (CRS4C-2; Rodents, mammals, | Antibacterial, Anti-Gram+, Anti-Gram-, Antimicrobial |
| 1196 | 3596 | DRAMP03360 | Cryptdin related sequence peptide (CRS4C-2b; Rodents, mammals, | Antibacterial, Anti-Gram+, Anti-Gram-, Antimicrobial |
| 1197 | 3598 | DRAMP03362 | CRS4C-3c (Cryptdin related sequence peptide; Rodents, mammals, | Antimicrobial , Anti-Gram+, Anti-Gram-, |
| 1198 | 3602 | DRAMP03366 | Beta-defensin 1 (BD-1; mBD-1; Defensin, beta 1; Rodents, | Antibacterial, Antifungal, Anti-Gram+, Anti-Gram-, Antimicrobial |
| 1199 | 3605 | DRAMP03369 | Beta-defensin 4 (BD-4, mBD-4; Defensin, beta 4; Rodents, | Antibacterial, Anti-Gram+, Anti-Gram-, Antimicrobial |
| 1200 | 3608 | DRAMP03373 | Beta-defensin 8 (BD-8, mBD-8; Defensin, beta 8; Rodents, | Antibacterial, Anti-Gram+, Anti-Gram-, Antimicrobial |
| 1201 | 3635 | DRAMP03400 | WAP four-disulfide core domain protein 12 (Rodents, mammals, | Antibacterial, Anti-Gram+, Anti-Gram-, Antimicrobial |
| 1202 | 3637 | DRAMP03402 | WAP four-disulfide core domain protein 15B (Elafin-like protein I; | Antibacterial, Anti-Gram+, Anti-Gram-, Antimicrobial |
| 1203 | 3639 | DRAMP03407 | Defr1 (Murine beta-defensin related peptide; Rodents, mammals, | Antibacterial, Anti-Gram+, Anti-Gram-, Antimicrobial |
| 1204 | 3640 | DRAMP03408 | Neutrophil defensin 1 (HANP-1; alpha-defensin; Rodents, mammals, | Antibacterial, Antifungal, Anti-Gram+, Anti-Gram-, Antimicrobial |
| 1205 | 3641 | DRAMP03409 | Neutrophil defensin 2 (HANP-2; alpha-defensin; Rodents, mammals, | Antibacterial, Antifungal, Anti-Gram+, Anti-Gram-, Antimicrobial |
| 1206 | 3642 | DRAMP03410 | Neutrophil defensin 3 (HANP-3; alpha-defensin; Rodents, mammals, | Antibacterial, Antifungal, Anti-Gram+, Anti-Gram-, Antimicrobial |
| 1207 | 3643 | DRAMP03411 | Neutrophil defensin 4 (HANP-4; alpha-defensin; Rodents, mammals, | Antibacterial, Antifungal, Anti-Gram+, Anti-Gram-, Antimicrobial |
| 1208 | 3651 | DRAMP03420 | Neutrophil antibiotic peptide NP-2 (RatNP-2; Rodents, mammals, | Antibacterial, Antifungal, Anti-Gram+, Anti-Gram-, Antimicrobial |
| 1209 | 3652 | DRAMP03421 | Neutrophil antibiotic peptide NP-3 (RatNP-3a, RatNP-3b; Rodents, | Antibacterial, Antifungal, Antiviral, Anti-Gram+, Anti-Gram-, Antimicrobial |
| 1210 | 3696 | DRAMP03470 | Defensin-1 (American oyster defensin, AOD; molluscs, animals) | Antibacterial, Anti-Gram+, Anti-Gram-, Antimicrobial |
| 1211 | 3721 | DRAMP03501 | La-LTP (LJAFP; Insects, animals) | Antibacterial, Antifungal, Anti-Gram+, Anti-Gram-, Antimicrobial |
| 1212 | 3728 | DRAMP03510 | Cecropin-A (Insects, animals) | Antibacterial, Antiviral, Anti-Gram+, Anti-Gram-, Antimicrobial |
| 1213 | 3729 | DRAMP03511 | Cecropin-B (Immune protein P9; Insects, animals) | Antibacterial, Anti-Gram+, Anti-Gram-, Antimicrobial |
| 1214 | 3730 | DRAMP03512 | Cecropin-D (Cecropin D; Insects, animals) | Antibacterial, Anti-Gram+, Anti-Gram-, Antimicrobial |
| 1215 | 3737 | DRAMP03534 | Antibacterial peptide enbocin (Moricin; Insects, animals) | Antibacterial, Anti-Gram+, Anti-Gram-, Antimicrobial |
| 1216 | 3738 | DRAMP03535 | Lebocin-1/2 (Pro-rich; Insects, animals) | Antibacterial, Anti-Gram+, Anti-Gram-, Antimicrobial |
| 1217 | 3739 | DRAMP03536 | Lebocin-3 (LEB 3; Insects, animals) | Antibacterial, Anti-Gram+, Anti-Gram-, Antimicrobial |
| 1218 | 3742 | DRAMP03554 | CCL20(1-67) (Human, mammals, animals) | Antibacterial, Anti-Gram+, Anti-Gram-, Antimicrobial |
| 1219 | 3743 | DRAMP03555 | CCL20(2-70) (Human, mammals, animals) | Antibacterial, Anti-Gram+, Anti-Gram-, Antimicrobial |

| S.no. | PepID | DRAMP_ID | Name of the AMP | Activity |
|---|---|---|---|---|
| 1220 | 3744 | DRAMP03556 | C-C motif chemokine 20 (Human, mammals, animals) | Antibacterial, Anti-Gram+, Anti-Gram-, Antimicrobial |
| 1221 | 3749 | DRAMP03564 | Human hepcidin-20 (Hepc20; one chain of Hepcidin; Human, | Antibacterial, Antifungal, Anti-Gram+, Anti-Gram-, Antimicrobial |
| 1222 | 3750 | DRAMP03565 | Human hepcidin-25 (Hepc25; one chain of Hepcidin; Human, | Antibacterial, Antifungal, Anti-Gram+, Anti-Gram-, Antimicrobial |
| 1223 | 3751 | DRAMP03566 | Salvic (Human, mammals, animals) | Antibacterial, Anti-Gram+, Anti-Gram-, Antimicrobial |
| 1224 | 3762 | DRAMP03585 | Human TC-1 (Chain of Platelet basic protein; Human, mammals, | Antibacterial, Antifungal, Anti-Gram+, Anti-Gram-, Antimicrobial |
| 1225 | 3763 | DRAMP03586 | Human TC-2 (Chain of Platelet basic protein; Human, mammals, | Antibacterial, Antifungal, Anti-Gram+, Anti-Gram-, Antimicrobial |
| 1226 | 3764 | DRAMP03587 | DCD-1 (chain of Dermcidin; Human, mammals, animals) | Antibacterial, Antifungal, Proteolytic, Anti-Gram+, Anti- Gram-, Antimicrobial |
| 1227 | 3765 | DRAMP03588 | Human MUC7 20-Mer (Human, mammals, animals) | Antibacterial, Antifungal, Anti-Gram+, Anti-Gram-, Antimicrobial |
| 1228 | 3768 | DRAMP03591 | Neutrophil defensin 1 (Defensin, alpha 1; HNP-1, HP-1; Human, | Antimicrobial, Antibacterial, Antifungal, Anti-Gram+, Anti- Gram-, |
| 1229 | 3769 | DRAMP03592 | Neutrophil defensin 2 (HNP-2, HP-2, HP2; Human, mammals, | Antifungal, Antiviral, Anti-Gram+, Anti-Gram-, Antimicrobial |
| 1230 | 3770 | DRAMP03593 | Neutrophil defensin 3 (Defensin, alpha 3; HNP-3, HP-3, HP3; | Antibacterial, Antifungal, Antiviral, Anti-Gram+, Anti-Gram-, Antimicrobial |
| 1231 | 3771 | DRAMP03594 | Neutrophil defensin 4 (Defensin, alpha 4; HNP-4, HP-4; Human, | Antibacterial, Antifungal, Antiviral, Anti-Gram+, Anti-Gram-, Antimicrobial |
| 1232 | 3772 | DRAMP03595 | Human defensin-5 (HD-5; Defensin, alpha 5; Human, mammals, | Antibacterial, Antifungal, Antiviral, Anti-Gram+, Anti-Gram-, Antimicrobial |
| 1233 | 3773 | DRAMP03596 | Human defensin-6 (HD-6; Defensin, alpha 6; Human, mammals, | Antifungal, Antiviral, Anti-Gram+, Anti-Gram-, Antimicrobial |
| 1234 | 3777 | DRAMP18203 | Panusin (beta defensins; crustaceans, arthropods, invertebrates, | Antibacterial, Anti-Gram+, Anti-Gram-, Antifungal, Antimicrobial |
| 1235 | 3789 | DRAMP18120 | BnPRP1 (Plant defensin) | Antibacterial, Antifungal, Anti-Gram+, Anti-Gram-, Antimicrobial |
| 1236 | 3799 | DRAMP18140 | VG16KRKP | Antibacterial, Antifungal, Anti-Gram+, Anti-Gram-, Antimicrobial |
| 1237 | 3802 | DRAMP18194 | AAEL000598-PA | Antibacterial, Anti-Gram+, Anti-Gram-, Antimicrobial |
| 1238 | 3808 | DRAMP03641 | Longicornsin (defensin-like; Arthropods, invertebrates, animals) | Antibacterial, Antifungal, Anti-Gram+, Anti-Gram-, Antimicrobial |
| 1239 | 3810 | DRAMP03644 | Cathelicidin-1 (CATH-1; Fowlicidin-1; Birds, animals) | Antibacterial, Cytolytic, Anti-Gram+, Anti-Gram-, Antimicrobial |
| 1240 | 3811 | DRAMP03648 | Gallinacin-1 (Gal-1; Beta-defensin 1; Birds, animals) | Antibacterial, Antifungal, Anti-Gram+, Anti-Gram-, Antimicrobial |
| 1241 | 3812 | DRAMP03649 | Gallinacin-1 alpha (Gal-1 alpha; Antimicrobial peptide CHP2; Birds, | Antibacterial, Antifungal, Anti-Gram+, Anti-Gram-, Antimicrobial |
| 1242 | 3813 | DRAMP03650 | Gallinacin-2 (Gal-2; Beta-defensin 2; Birds, animals) | Antibacterial, Anti-Gram+, Anti-Gram-, Antimicrobial |
| 1243 | 3823 | DRAMP03661 | Gallinacin-13 (Gal-13; Beta-defensin 13; Birds, animals) | Antibacterial, Anti-Gram+, Anti-Gram-, Antimicrobial |
| 1244 | 3836 | DRAMP03674 | Cystatin-1 (Cystatin-I) | Antibacterial, Anti-Gram+, Anti-Gram-, Antimicrobial |
| 1245 | 3839 | DRAMP03680 | Cathelicidin-3.4 (Bactenecin-3.4, Bac3.4; ChBac3.4; ruminant, | Antibacterial, Anti-Gram+, Anti-Gram-, Antimicrobial |
| 1246 | 3843 | DRAMP03692 | Defensin-1 (Cll-dlp; Arthropods, animals) | Antibacterial, Anti-Gram+, Anti-Gram-, Antimicrobial |
| 1247 | 3871 | DRAMP03736 | Opistoporin-2 (OP2; Non-disulfide-bridged peptide 3.6, NDBP-3.6; | Antibacterial, Antifungal, Anti-Gram+, Anti-Gram-, Antimicrobial |
| 1248 | 3872 | DRAMP03737 | Opistoporin-4 (Non-disulfide-bridged peptides 3.7, NDBP-3.7; | Antibacterial, Antifungal, Anti-Gram+, Anti-Gram-, Antimicrobial |
| 1249 | 3873 | DRAMP03739 | Buthinin (Sahara scorpion; Arthropods, animals) | Antibacterial, Anti-Gram+, Anti-Gram-, Antimicrobial |
| 1250 | 3874 | DRAMP03740 | Androctonus defensin (4 kDa defensin; Arthropods, animals) | Antibacterial, Anti-Gram+, Anti-Gram-, Antimicrobial |
| 1251 | 3875 | DRAMP03741 | Ponericin-W-like 32.1 (Arthropods, animals) | Antibacterial, Antifungal, Insecticidal, Anti-Gram+, Anti-Gram-, Antimicrobial |
| 1252 | 3876 | DRAMP03742 | Ponericin-W-like 32.2 (Arthropods, animals) | Antibacterial, Antifungal, Insecticidal, Anti-Gram+, Anti-Gram-, Antimicrobial |
| 1253 | 3880 | DRAMP03755 | Potassium channel toxin alpha-KTx 1.1 (ChTX-Lq1; charybdotoxin; | Antibacterial, Antifungal, Antiviral, Anti-Gram+, Anti-Gram-, Antimicrobial |
| 1254 | 3891 | DRAMP03766 | Heteroscorpine-1 (HS-1; defensins; Arthropods, animals) | Antibacterial, Anti-Gram+, Anti-Gram-, Antimicrobial |
| 1255 | 3907 | DRAMP03787 | Neuropeptide-like protein 31 (NLP-31; nematodes, animals) | Antifungal, Antibacterial, Anti-Gram+, Anti-Gram-, Antimicrobial |
| 1256 | 3910 | DRAMP03790 | ABF-2 (nematodes, animals) | Antibacterial, Antifungal, Anti-Gram+, Anti-Gram-, Antimicrobial |
| 1257 | 3944 | DRAMP18190 | Pantinin-3 (Non-disulfide-bridged peptide 4.22, NDBP-4.22, Non- | Antibacterial, Antifungal, Anti-Gram+, Anti-Gram-, Antimicrobial |
| 1258 | 3948 | DRAMP03872 | hLf 21-30 (fragment of human lalctoferricin, residues 21-30) | Antibacterial, Anti-Gram+, Anti-Gram-, Antimicrobial |
| 1259 | 3949 | DRAMP03873 | mLf 20-29 (fragment of murine lalctoferricin, residues 20-29) | Antibacterial, Anti-Gram+, Anti-Gram-, Antimicrobial |
| 1260 | 3950 | DRAMP03874 | pLf20-29 (fragment of porcine lactoferricin, residues 20-29) | Antibacterial, Anti-Gram+, Anti-Gram-, Antimicrobial |
| 1261 | 3961 | DRAMP03941 | Peptide 3 (Trp- and Arg-rich; derivative of Tritrpticin) | Antibacterial, Antifungal, Anti-Gram+, Anti-Gram-, Antimicrobial |
| 1262 | 3962 | DRAMP03942 | Peptide 2 (Trp- and Arg-rich; derivative of Tritrpticin) | Antibacterial, Antifungal, Anti-Gram+, Anti-Gram-, Antimicrobial |
| 1263 | 3965 | DRAMP03946 | Del 1-3 (Ranalexin analog) | Antibacterial, Anti-Gram+, Anti-Gram-, Antimicrobial |
| 1264 | 3970 | DRAMP03961 | KR-12 | Antibacterial, Anti-Gram+, Anti-Gram-, Antimicrobial |
| 1265 | 3976 | DRAMP02092 | Brevinin-1Bb (Frogs, amphibians, animals) | Antimicrobial, Antibacterial, Antifungal, Anti-Gram+, Anti-Gram-, |
| 1266 | 3978 | DRAMP18189 | Pantinin-2 (Non-disulfide-bridged peptide 4.21, NDBP-4.21, Non- | Antibacterial, Antifungal, Anti-Gram+, Anti-Gram-, Antimicrobial |
| 1267 | 3982 | DRAMP04018 | Rp-1 | Antibacterial, Antifungal, Anti-Gram+, Anti-Gram-, Antimicrobial |
| 1268 | 3983 | DRAMP04037 | Immobilized peptide E07LKK | Antibacterial, Antifungal, Anti-Gram+, Anti-Gram-, Antimicrobial |
| 1269 | 3984 | DRAMP04038 | Immobilized peptide E14LKK/H14LKK | Antibacterial, Antifungal, Anti-Gram+, Anti-Gram-, Antimicrobial |
| 1270 | 3985 | DRAMP04039 | Immobilized peptige E16KGL/H16KGL | Antibacterial, Antifungal, Anti-Gram+, Anti-Gram-, Antimicrobial |
| 1271 | 3986 | DRAMP04040 | Immobilized peptide E17KGG | Antibacterial, Antifungal, Anti-Gram+, Anti-Gram-, Antimicrobial |
| 1272 | 3987 | DRAMP04041 | Immobilized peptide E18KGG | Antibacterial, Antifungal, Anti-Gram+, Anti-Gram-, Antimicrobial |

| S.no. | PepID | DRAMP_ID | Name of the AMP | Activity |
|---|---|---|---|---|
| 1273 | 3988 | DRAMP04042 | Immobilized peptide E16LKL | Antibacterial, Antifungal, Anti-Gram+, Anti-Gram-, Antimicrobial |
| 1274 | 3989 | DRAMP04043 | Immobilized peptid E10KKL | Antibacterial, Antifungal, Anti-Gram+, Anti-Gram-, Antimicrobial |
| 1275 | 3990 | DRAMP04044 | Immobilized peptid E12LLK | Antibacterial, Antifungal, Anti-Gram+, Anti-Gram-, Antimicrobial |
| 1276 | 3991 | DRAMP04045 | Immobilized peptid E14KKL | Antibacterial, Antifungal, Anti-Gram+, Anti-Gram-, Antimicrobial |
| 1277 | 3992 | DRAMP04046 | Immobilized peptide E23GIG magainin2 | Antibacterial, Antifungal, Anti-Gram+, Anti-Gram-, Antimicrobial |
| 1278 | 3993 | DRAMP04047 | Immobilized peptide E17HSA magainin 2 deletion | Antibacterial, Antifungal, Anti-Gram+, Anti-Gram-, Antimicrobial |
| 1279 | 4020 | DRAMP04172 | LK2W2 (LlKmW2 model peptides) | Antibacterial, Anti-Gram+, Anti-Gram-, Antimicrobial |
| 1280 | 4021 | DRAMP04173 | L2KW2 (LlKmW2 model peptides) | Antibacterial, Anti-Gram+, Anti-Gram-, Antimicrobial |
| 1281 | 4169 | DRAMP04366 | PDD-A-8 (PDD-A analog) | Antibacterial, Anti-Gram+, Anti-Gram-, Antimicrobial |
| 1282 | 4170 | DRAMP04375 | PDD-B-5 (PDD-B analog) | Antibacterial, Anti-Gram+, Anti-Gram-, Antimicrobial |
| 1283 | 4171 | DRAMP04384 | PMM-5 (PMM analog) | Antibacterial, Anti-Gram+, Anti-Gram-, Antimicrobial |
| 1284 | 4172 | DRAMP04388 | PMM-9 (PMM analog) | Antibacterial, Anti-Gram+, Anti-Gram-, Antimicrobial |
| 1285 | 4424 | DRAMP04658 | Hymenochirin-5B | Antimicrobial, Anti-Gram+, Anti-Gram-, |
| 1286 | 4435 | DRAMP18188 | Pantinin-1 (Non-disulfide-bridged peptide 4.20, NDBP-4.20, Non- | Antibacterial, Antifungal, Anti-Gram+, Anti-Gram-, Antimicrobial |
| 1287 | 4443 | DRAMP04684 | Bacteriocin BAC-IB17 | Antibacterial, Anti-Gram+, Anti-Gram-, Antimicrobial |
| 1288 | 4451 | DRAMP04700 | Basic phospholipase A2 BnpTX-1 (BnPTx-I, svPLA2; | Antibacterial, Antiparasitic, Anti-Gram+, Anti-Gram-, Antimicrobial |
| 1289 | 4472 | DRAMP18187 | Toxin LyeTx 1 | Antibacterial, Anti-Gram+, Anti-Gram-, Antimicrobial |
| 1290 | 4473 | DRAMP18128 | Antimicrobial peptide HsAp4; | Antimicrobial, Antifungal, Anti-Gram+, Anti-Gram-, |
| 1291 | 4474 | DRAMP18129 | Antimicrobial peptide HsAp3; | Antimicrobial, Antifungal, Anti-Gram+, Anti-Gram-, |
| 1292 | 4475 | DRAMP18130 | Antimicrobial peptide HsAp2 | Antimicrobial, Antifungal, Anti-Gram+, Anti-Gram-, |
| 1293 | 4476 | DRAMP18131 | Antimicrobial peptide HsAp1 (HsAp) | Antimicrobial, Antifungal, Anti-Gram+, Anti-Gram-, |
| 1294 | 4485 | DRAMP18185 | Jingdongin-1 | Antibacterial, Antifungal, Anti-Gram+, Anti-Gram-, Antimicrobial |
| 1295 | 4531 | DRAMP18454 | Tepmporin-1Ee (frog, amphibians, animals) | Antibacterial, Anti-Gram+, Anti-Gram-, Antimicrobial |
| 1296 | 4533 | DRAMP18456 | Pepcon (peptide consensus sequence, synthetic) | Antibacterial, Anti-Gram+, Anti-Gram-, Antimicrobial |
| 1297 | 4555 | DRAMP18488 | Css54 (Css from the species name below; scorpions, arachnids, | Antibacterial, Cytolysis, Hemolysis, Anti-Gram+, Anti-Gram-, Antimicrobial |
| 1298 | 4564 | DRAMP03575 | LL-37(17-29) (C-terminal fragment of LL-37, LL; Human, mammals, | Antibacterial, Anticancer, Anti-Gram+, Anti-Gram-, Antimicrobial |
| 1299 | 4566 | DRAMP02094 | Brevinin-1Bd (Frogs, amphibians, animals) | Antimicrobial, Antibacterial, Antifungal, Anti-Gram+, Anti- Gram-, |
| 1300 | 4567 | DRAMP02095 | Brevinin-1Be (Frogs, amphibians, animals) | Antibacterial, Anti-Gram+, Anti-Gram-, Antimicrobial |
| 1301 | 4568 | DRAMP02096 | Brevinin-1Bf (Frogs, amphibians, animals) | Antibacterial, Anti-Gram+, Anti-Gram-, Antimicrobial |
| 1302 | 4569 | DRAMP02097 | Brevinin-1Pa (Frogs, amphibians, animals) | Antimicrobial, Antibacterial, Antifungal, Anti-Gram+, Anti-Gram-, |
| 1303 | 4570 | DRAMP02098 | Brevinin-1Pc (Frogs, amphibians, animals) | Antimicrobial, Antibacterial, Antifungal, Anti-Gram+, Anti-Gram-, |
| 1304 | 4571 | DRAMP02099 | Brevinin-1Pd (Frogs, amphibians, animals) | Antimicrobial, Antibacterial, Antifungal, Anti-Gram+, Anti-Gram-, |
| 1305 | 4572 | DRAMP02255 | Ranatuerin-2Lb (Ranatuerin 2Lb; Ranaturin-2PRd; Frogs, | Antimicronial, Antibacterial, Antifungal, Anti-Gram+, Anti- Gram-, Antimicrobial |
| 1306 | 4573 | DRAMP02254 | Ranatuerin-2La (Ranatuerin 2La; Ranatuerin-2PRa; Frogs, | Antibacterial, Anti-Gram+, Anti-Gram-, Antimicrobial |
| 1307 | 4574 | DRAMP02256 | Ranatuerin-2B (Ranatuerin 2B, Frog, amphibians, animals) | Antimicronial, Antibacterial, Antifungal, Anti-Gram+, Anti- Gram-, Antimicrobial |
| 1308 | 4575 | DRAMP02257 | Ranatuerin-2P (Ranatuerin 2P; Frogs, amphibians, animals) | Antimicronial, Antibacterial, Antifungal, Anti-Gram+, Anti- Gram-, Antimicrobial |
| 1309 | 4577 | DRAMP18497 | TSG-6 (Ixosin-B peptide derivative) | Antibacterial, Anti-Gram+, Anti-Gram-, Antimicrobial |
| 1310 | 4578 | DRAMP18498 | TSG-7 (Ixosin-B peptide derivative) | Antibacterial, Anti-Gram+, Anti-Gram-, Antimicrobial |
| 1311 | 4579 | DRAMP18499 | TSG-8 (Ixosin-B peptide derivative) | Antibacterial, Anti-Gram+, Anti-Gram-, Antimicrobial |
| 1312 | 4580 | DRAMP18500 | TSG-8-1 (Ixosin-B peptide derivative) | Antibacterial, Anti-Gram+, Anti-Gram-, Antimicrobial |
| 1313 | 4581 | DRAMP18501 | TSG-9 (Ixosin-B peptide derivative) | Antibacterial, Anti-Gram+, Anti-Gram-, Antimicrobial |
| 1314 | 4582 | DRAMP18502 | TSG-10 (Ixosin-B peptide derivative) | Antibacterial, Anti-Gram+, Anti-Gram-, Antimicrobial |
| 1315 | 4583 | DRAMP18503 | TSG-11 (Ixosin-B peptide derivative) | Antibacterial, Anti-Gram+, Anti-Gram-, Antimicrobial |
| 1316 | 4586 | DRAMP18506 | OG2 (Palustrin-OG1 peptide derivative) | Antibacterial, Anti-Gram+, Anti-Gram-, Antimicrobial |
| 1317 | 4587 | DRAMP18508 | gp41w-FKA (gp41 peptide derivative) | Antibacterial, Anti-Gram+, Anti-Gram-, Antimicrobial |
| 1318 | 4588 | DRAMP18509 | Px-cec1 (cecropin1 peptide derivative) | Antimicrobial, Antibacterial, Anitifungal, Anti-Gram+, Anti- Gram-, |
| 1319 | 4595 | DRAMP18533 | V13KL (V681 peptide derivative) | Antibacterial, Anti-Gram+, Anti-Gram-, Antimicrobial |
| 1320 | 4600 | DRAMP18538 | V681 | Antibacterial, Anti-Gram+, Anti-Gram-, Antimicrobial |
| 1321 | 4601 | DRAMP18539 | V13LL (V681 peptide derivative) | Antibacterial, Anti-Gram+, Anti-Gram-, Antimicrobial |
| 1322 | 4602 | DRAMP18540 | V13AL (V681 peptide derivative) | Antibacterial, Anti-Gram+, Anti-Gram-, Antimicrobial |
| 1323 | 4603 | DRAMP18541 | V13G (V681 peptide derivative) | Antibacterial, Anti-Gram+, Anti-Gram-, Antimicrobial |
| 1324 | 4604 | DRAMP18542 | V13SL (V681 peptide derivative) | Antibacterial, Anti-Gram+, Anti-Gram-, Antimicrobial |
| 1325 | 4605 | DRAMP18543 | V13LD (V681 peptide derivative) | Antibacterial, Anti-Gram+, Anti-Gram-, Antimicrobial |

| S.no. | PepID | DRAMP_ID | Name of the AMP | Activity |
|---|---|---|---|---|
| 1326 | 4606 | DRAMP18544 | V13VD (V681 peptide derivative) | Antibacterial, Anti-Gram+, Anti-Gram-, Antimicrobial |
| 1327 | 4607 | DRAMP18545 | V13AD (V681 peptide derivative) | Antibacterial, Anti-Gram+, Anti-Gram-, Antimicrobial |
| 1328 | 4608 | DRAMP18546 | V13SD (V681 peptide derivative) | Antibacterial, Anti-Gram+, Anti-Gram-, Antimicrobial |
| 1329 | 4609 | DRAMP18547 | V13KD (V681 peptide derivative) | Antibacterial, Anti-Gram+, Anti-Gram-, Antimicrobial |
| 1330 | 4610 | DRAMP18548 | S11LL (V681 peptide derivative) | Antibacterial, Anti-Gram+, Anti-Gram-, Antimicrobial |
| 1331 | 4611 | DRAMP18549 | S11VL (V681 peptide derivative) | Antibacterial, Anti-Gram+, Anti-Gram-, Antimicrobial |
| 1332 | 4612 | DRAMP18550 | S11AL (V681 peptide derivative) | Antibacterial, Anti-Gram+, Anti-Gram-, Antimicrobial |
| 1333 | 4613 | DRAMP18551 | S11G (V681 peptide derivative) | Antibacterial, Anti-Gram+, Anti-Gram-, Antimicrobial |
| 1334 | 4614 | DRAMP18552 | S11KL (V681 peptide derivative) | Antibacterial, Anti-Gram+, Anti-Gram-, Antimicrobial |
| 1335 | 4615 | DRAMP18553 | S11LD (V681 peptide derivative) | Antibacterial, Anti-Gram+, Anti-Gram-, Antimicrobial |
| 1336 | 4616 | DRAMP18554 | S11VD (V681 peptide derivative) | Antibacterial, Anti-Gram+, Anti-Gram-, Antimicrobial |
| 1337 | 4617 | DRAMP18555 | S11AD (V681 peptide derivative) | Antibacterial, Anti-Gram+, Anti-Gram-, Antimicrobial |
| 1338 | 4618 | DRAMP18556 | S11SD (V681 peptide derivative) | Antibacterial, Anti-Gram+, Anti-Gram-, Antimicrobial |
| 1339 | 4619 | DRAMP18557 | S11KD (V681 peptide derivative) | Antibacterial, Anti-Gram+, Anti-Gram-, Antimicrobial |
| 1340 | 4620 | DRAMP18558 | Kn2-7 (BmKn2 peptide derivative) | Antibacterial, Anti-Gram+, Anti-Gram-, Antimicrobial |
| 1341 | 4621 | DRAMP18559 | HFU3 | Antibacterial, Anti-Gram+, Anti-Gram-, Antimicrobial |
| 1342 | 4622 | DRAMP18560 | HFU4 | Antibacterial, Anti-Gram+, Anti-Gram-, Antimicrobial |
| 1343 | 4623 | DRAMP18561 | HFU5 | Antibacterial, Anti-Gram+, Anti-Gram-, Antimicrobial |
| 1344 | 4624 | DRAMP18562 | MAP-04-01 (Ixosin-B peptide derivative) | Antibacterial, Anti-Gram+, Anti-Gram-, Antimicrobial |
| 1345 | 4625 | DRAMP18563 | MAP-04-02 (Ixosin-B peptide derivative) | Antibacterial, Anti-Gram+, Anti-Gram-, Antimicrobial |
| 1346 | 4626 | DRAMP18564 | MAP-04-03 (Ixosin-B peptide derivative) | Antibacterial, Anti-Gram+, Anti-Gram-, Antimicrobial |
| 1347 | 4627 | DRAMP18565 | MAP-04-04 (Ixosin-B peptide derivative) | Antibacterial, Anti-Gram+, Anti-Gram-, Antimicrobial |
| 1348 | 4628 | DRAMP18566 | LL-IIIs-1 (lasioglossin III peptide derivative) | Antimicrobial, Antibacterial, Antifungal, Anti-Gram+, Anti-Gram-, |
| 1349 | 4629 | DRAMP18567 | LL-IIIs-2 (lasioglossin III peptide derivative) | Antimicrobial, Antibacterial, Antifungal, Anti-Gram+, Anti-Gram-, |
| 1350 | 4630 | DRAMP18568 | LL-IIIs-3 (lasioglossin III peptide derivative) | Antimicrobial, Antibacterial, Antifungal, Anti-Gram+, Anti-Gram-, |
| 1351 | 4631 | DRAMP18569 | LL-IIIs-4 (lasioglossin III peptide derivative) | Antibacterial, Anti-Gram+, Anti-Gram-, Antimicrobial |
| 1352 | 4632 | DRAMP18570 | LL-IIIs-5 cis (lasioglossin III peptide derivative) | Antibacterial, Anti-Gram+, Anti-Gram-, Antimicrobial |
| 1353 | 4633 | DRAMP18571 | LL-IIIs-5 trans (lasioglossin III peptide derivative) | Antibacterial, Anti-Gram+, Anti-Gram-, Antimicrobial |
| 1354 | 4634 | DRAMP18572 | LL-IIIs-6a (lasioglossin III peptide derivative) | Antibacterial, Anti-Gram+, Anti-Gram-, Antimicrobial |
| 1355 | 4635 | DRAMP18573 | LL-IIIs-6b (lasioglossin III peptide derivative) | Antibacterial, Anti-Gram+, Anti-Gram-, Antimicrobial |
| 1356 | 4636 | DRAMP18574 | MEP-N (melectin peptide derivative) | Antimicrobial, Antibacterial, Antifungal, Anti-Gram+, Anti-Gram-, |
| 1357 | 4637 | DRAMP18575 | MEP-Ns-1 (melectin peptide derivative) | Antimicrobial, Antibacterial, Antifungal, Anti-Gram+, Anti-Gram-, |
| 1358 | 4638 | DRAMP18576 | MEP-Ns-2 (melectin peptide derivative) | Antibacterial, Anti-Gram+, Anti-Gram-, Antimicrobial |
| 1359 | 4639 | DRAMP18577 | MEP-Ns-3 (melectin peptide derivative) | Antibacterial, Anti-Gram+, Anti-Gram-, Antimicrobial |
| 1360 | 4640 | DRAMP18578 | MEP-Ns-4 cis (melectin peptide derivative) | Antibacterial, Anti-Gram+, Anti-Gram-, Antimicrobial |
| 1361 | 4641 | DRAMP18579 | MEP-Ns-4 trans (melectin peptide derivative) | Antibacterial, Anti-Gram+, Anti-Gram-, Antimicrobial |
| 1362 | 4642 | DRAMP18580 | MEP-Ns-5 (melectin peptide derivative) | Antibacterial, Anti-Gram+, Anti-Gram-, Antimicrobial |
| 1363 | 4643 | DRAMP18581 | MEP-Ns-6 (melectin peptide derivative) | Antibacterial, Anti-Gram+, Anti-Gram-, Antimicrobial |
| 1364 | 4645 | DRAMP18583 | Tricystine cyclic cystine TP (ccTP, Tachyplesin-1 peptide derivative) | Antimicrobial, Antibacterial, Antifungal, Anti-Gram+, Anti-Gram-, |
| 1365 | 4646 | DRAMP18584 | [Arg13]ccTP (ccTP peptide derivative) | Antimicrobial, Antibacterial, Antifungal, Anti-Gram+, Anti-Gram-, |
| 1366 | 4647 | DRAMP18585 | [Arg4,8]ccTP (ccTP peptide derivative) | Antimicrobial, Antibacterial, Antifungal, Anti-Gram+, Anti- Gram-, |
| 1367 | 4648 | DRAMP18586 | [Arg4,8,13]ccTP (ccTP peptide derivative) | Antimicrobial, Antibacterial, Antifungal, Anti-Gram+, Anti- Gram-, |
| 1368 | 4649 | DRAMP18587 | [Arg4,8,13][Lys18]ccTP (ccTP peptide derivative) | Antimicrobial, Antibacterial, Antifungal, Anti-Gram+, Anti- Gram-, |
| 1369 | 4650 | DRAMP18588 | RTD | Antimicrobial, Antibacterial, Antifungal, Anti-Gram+, Anti-Gram-, |
| 1370 | 4651 | DRAMP18589 | DSE (Ctx-Ha peptide derivative) | Antimicrobial, Antibacterial, Antifungal, Anti-Gram+, Anti-Gram-, |
| 1371 | 4652 | DRAMP18590 | DEP (Ctx-Ha peptide derivative) | Antimicrobial, Antibacterial, Antifungal, Anti-Gram+, Anti- Gram-, |
| 1372 | 4653 | DRAMP18591 | DEA (Ctx-Ha peptide derivative) | Antimicrobial, Antibacterial, Antifungal, Anti-Gram+, Anti- Gram-, |
| 1373 | 4654 | DRAMP18592 | Ctx(Ile21)-Ha (Ctx-Ha peptide derivative) | Antimicrobial, Antibacterial, Antifungal, Anti-Gram+, Anti- Gram-, |
| 1374 | 4655 | DRAMP18593 | Ctx(Ile21)-Ha-VD16 (Ctx-Ha peptide derivative) | Antimicrobial, Antibacterial, Antifungal, Anti-Gram+, Anti- Gram-, |
| 1375 | 4656 | DRAMP18594 | Ctx(Ile21)-Ha-VD5,16 (Ctx-Ha peptide derivative) | Antimicrobial, Antibacterial, Antifungal, Anti-Gram+, Anti- Gram-, |
| 1376 | 4658 | DRAMP18596 | LL-I/1 (Lasioglossin LL-I peptide derivative) | Antibacterial, Anti-Gram+, Anti-Gram-, Antimicrobial |
| 1377 | 4659 | DRAMP18597 | LL-I/2 (Lasioglossin LL-I peptide derivative) | Antibacterial, Anti-Gram+, Anti-Gram-, Antimicrobial |
| 1378 | 4660 | DRAMP18598 | LL-I/3 (Lasioglossin LL-I peptide derivative) | Antibacterial, Anti-Gram+, Anti-Gram-, Antimicrobial |

| S.no. | PepID | DRAMP_ID | Name of the AMP | Activity |
|---|---|---|---|---|
| 1379 | 4661 | DRAMP18599 | LL-I/4 (Lasioglossin LL-I peptide derivative) | Antibacterial, Anti-Gram+, Anti-Gram-, Antimicrobial |
| 1380 | 4662 | DRAMP18600 | LL-II/1 (Lasioglossin LL-II peptide derivative) | Antibacterial, Anti-Gram+, Anti-Gram-, Antimicrobial |
| 1381 | 4663 | DRAMP18601 | LL-II/2 (Lasioglossin LL-II peptide derivative) | Antibacterial, Anti-Gram+, Anti-Gram-, Antimicrobial |
| 1382 | 4664 | DRAMP18602 | LL-II/3 (Lasioglossin LL-II peptide derivative) | Antibacterial, Anti-Gram+, Anti-Gram-, Antimicrobial |
| 1383 | 4665 | DRAMP18603 | LL-II/4 (Lasioglossin LL-II peptide derivative) | Antibacterial, Anti-Gram+, Anti-Gram-, Antimicrobial |
| 1384 | 4666 | DRAMP18604 | LL-III/1 (Lasioglossin LL-III peptide derivative) | Antibacterial, Anti-Gram+, Anti-Gram-, Antimicrobial |
| 1385 | 4667 | DRAMP18605 | LL-III/2 (Lasioglossin LL-III peptide derivative) | Antibacterial, Anti-Gram+, Anti-Gram-, Antimicrobial |
| 1386 | 4668 | DRAMP18606 | LL-III/3 (Lasioglossin LL-III peptide derivative) | Antibacterial, Anti-Gram+, Anti-Gram-, Antimicrobial |
| 1387 | 4669 | DRAMP18607 | LL-III/4 (Lasioglossin LL-III peptide derivative) | Antibacterial, Anti-Gram+, Anti-Gram-, Antimicrobial |
| 1388 | 4670 | DRAMP18608 | LL-III/5 (Lasioglossin LL-III peptide derivative) | Antibacterial, Anti-Gram+, Anti-Gram-, Antimicrobial |
| 1389 | 4671 | DRAMP18609 | LL-III/6 (Lasioglossin LL-III peptide derivative) | Antibacterial, Anti-Gram+, Anti-Gram-, Antimicrobial |
| 1390 | 4672 | DRAMP18610 | LL-III/7 (Lasioglossin LL-III peptide derivative) | Antibacterial, Anti-Gram+, Anti-Gram-, Antimicrobial |
| 1391 | 4675 | DRAMP18613 | TPG (Tritrpticin peptide derivative) | Antimicrobial, Antibacterial, Antifungal, Anti-Gram+, Anti-Gram-, |
| 1392 | 4676 | DRAMP18627 | D4-K9L8W (D-amino acid substitution of K9L8W) | Antibacterial, Anti-Gram+, Anti-Gram-, Antimicrobial |
| 1393 | 4677 | DRAMP18507 | SolyC (Plant defensin; tomato, plants) | Antibacterial, Anti-Gram+, Anti-Gram-, Antimicrobial |
| 1394 | 4681 | DRAMP18614 | TPA (Tritrpticin peptide derivative) | Antimicrobial, Antibacterial, Antifungal, Anti-Gram+, Anti- Gram-, |
| 1395 | 4682 | DRAMP18615 | TWF (Tritrpticin peptide derivative) | Antimicrobial, Antibacterial, Antifungal, Anti-Gram+, Anti- Gram-, |
| 1396 | 4683 | DRAMP18616 | [K22,25,27]-SMAP-29 (SMAP-29 peptide derivative) | Antimicrobial, Antibacterial, Antifungal, Anti-Gram+, Anti-Gram-, |
| 1397 | 4684 | DRAMP18617 | [A19]-SMAP-29 (SMAP-29 peptide derivative) | Antimicrobial, Antibacterial, Antifungal, Anti-Gram+, Anti-Gram-, |
| 1398 | 4685 | DRAMP18618 | SMAP-29(1-17) (SMAP-29 peptide derivative) | Antimicrobial, Antibacterial, Antifungal, Anti-Gram+, Anti- Gram-, |
| 1399 | 4686 | DRAMP18619 | [K2,7,13]-SMAP-29(1-17) (SMAP-29 peptide derivative) | Antimicrobial, Antibacterial, Antifungal, Anti-Gram+, Anti- Gram-, |
| 1400 | 4687 | DRAMP18620 | Pep-1-K (Pep-1 peptide derivative) | Antibacterial, Anti-Gram+, Anti-Gram-, Antimicrobial |
| 1401 | 4689 | DRAMP18622 | Temporin-PEb (Temporin-PE peptide derivative) | Antimicrobial, Antibacterial, Antifungal, Anti-Gram+, Anti- Gram-, |
| 1402 | 4690 | DRAMP18623 | [I5,R8] Mastoparan-L ([I5,R8] MP-L;Mastoparan-L peptide derivative) | Antimicrobial, Antibacterial, Antifungal, Anti-Gram+, Anti- Gram-, |
| 1403 | 4691 | DRAMP18624 | K9L8W | Antibacterial, Anti-Gram+, Anti-Gram-, Antimicrobial |
| 1404 | 4692 | DRAMP18625 | D3-K9L8W-1 (D-amino acid substitution of K9L8W) | Antibacterial, Anti-Gram+, Anti-Gram-, Antimicrobial |
| 1405 | 4693 | DRAMP18626 | D3-K9L8W-2 (D-amino acid substitution of K9L8W) | Antibacterial, Anti-Gram+, Anti-Gram-, Antimicrobial |
| 1406 | 4694 | DRAMP18628 | D6-K9L8W (D-amino acid substitution of K9L8W) | Antibacterial, Anti-Gram+, Anti-Gram-, Antimicrobial |
| 1407 | 4695 | DRAMP18629 | D9-K9L8W-1 (D-amino acid substitution of K9L8W) | Antibacterial, Anti-Gram+, Anti-Gram-, Antimicrobial |
| 1408 | 4696 | DRAMP18630 | D9-K9L8W-2 (D-amino acid substitution of K9L8W) | Antibacterial, Anti-Gram+, Anti-Gram-, Antimicrobial |
| 1409 | 4697 | DRAMP18631 | H5(61-90) V1 (Histone H5 peptide derivative) | Antibacterial, Anti-Gram+, Anti-Gram-, Antimicrobial |
| 1410 | 4700 | DRAMP18634 | H5(61-90) V3 (Histone H5 peptide derivative) | Antibacterial, Anti-Gram+, Anti-Gram-, Antimicrobial |
| 1411 | 4702 | DRAMP18636 | NCP-3a (CTX-1 peptide derivative) | Antimicrobial, Antibacterial, Antifungal, Anti-Gram+, Anti-Gram-, |
| 1412 | 4703 | DRAMP18637 | NCP-3b (CTX-1 peptide derivative) | Antimicrobial, Antibacterial, Antifungal, Anti-Gram+, Anti-Gram-, |
| 1413 | 4707 | DRAMP18641 | KCM11 | Antibacterial, Anti-Gram+, Anti-Gram-, Antimicrobial |
| 1414 | 4708 | DRAMP18642 | KCM12 | Antimicrobial, Antibacterial, Antifungal, Anti-Gram+, Anti-Gram-, |
| 1415 | 4709 | DRAMP18643 | KCM21 | Antimicrobial, Antibacterial, Antifungal, Anti-Gram+, Anti-Gram-, |
| 1416 | 4710 | DRAMP18644 | KRS22 | Antibacterial, Anti-Gram+, Anti-Gram-, Antimicrobial |
| 1417 | 4714 | DRAMP18648 | [Pro3,DLeu9]TL(3) (Temporin L peptide derivative) | Antimicrobial, Antibacterial, Antifungal, Anti-Gram+, Anti- Gram-, |
| 1418 | 4724 | DRAMP18658 | Temporin-PE (Edible frogs, amphibians, animals) | Antimicrobial, Antibacterial, Antifungal, Anticancer, Anti- Gram+, Anti-Gram-, |
| 1419 | 4725 | DRAMP18659 | YFGAP-OH(Yellowfin tuna GAPDH-related antimicrobial peptide; | Antibacterial, Anti-Gram+, Anti-Gram-, Antimicrobial |
| 1420 | 4726 | DRAMP18660 | YFGAP-NH2(Yellowfin tuna GAPDH-related antimicrobial peptide; | Antimicrobial, Antibacterial, Antifungal, Anti-Gram+, Anti-Gram-, |
| 1421 | 4727 | DRAMP18661 | Ctx-Ha (Frogs, amphibians, animals) | Antimicrobial, Antibacterial, Antifungal, Anti-Gram+, Anti-Gram-, |
| 1422 | 4728 | DRAMP18662 | Brevinin 21 (Brevinin-1E truncated peptide 21; Frogs, amphibians, | Antimicrobial, Antibacterial, Antifungal, Anti-Gram+, Anti-Gram-, |
| 1423 | 4729 | DRAMP18663 | Brevinin 18 (Brevinin-1E truncated peptide 18; Frogs, amphibians, | Antimicrobial, Antibacterial, Antifungal, Anti-Gram+, Anti-Gram-, |
| 1424 | 4730 | DRAMP18664 | Brevinin 15 (Brevinin-1E truncated peptide 15; Frogs, amphibians, | Antimicrobial, Antibacterial, Antifungal, Anti-Gram+, Anti- Gram-, |
| 1425 | 4731 | DRAMP18665 | Mastoparan-L (MP-L; insects, arthropods, invertebrates, animals) | Antibacterial, Anti-Gram+, Anti-Gram-, Antimicrobial |
| 1426 | 4732 | DRAMP18666 | P1 (Pilosulin-1 1-20; Ant, insects, arthropods, invertebrates, animals) | Antimicrobial, Antibacterial, Anitifungal, Anti-Gram+, Anti- Gram-, |
| 1427 | 4733 | DRAMP18667 | Pep-1 | Antibacterial, Anti-Gram+, Anti-Gram-, Antimicrobial |
| 1428 | 4737 | DRAMP18671 | TO17 (TFPI-1 C-terminal peptide) | Antibacterial, Anti-Gram+, Anti-Gram-, Antimicrobial |
| 1429 | 4738 | DRAMP18672 | Peptide 7 (Mollusca/molluscs/mollusks, invertebrates, animals) | Antibacterial, Anti-Gram+, Anti-Gram-, Antimicrobial |
| 1430 | 4739 | DRAMP18673 | Peptide 3 (Mollusca/molluscs/mollusks, invertebrates, animals) | Antibacterial, Anti-Gram+, Anti-Gram-, Antimicrobial |
| 1431 | 4740 | DRAMP18674 | Peptide 2 (Mollusca/molluscs/mollusks, invertebrates, animals) | Antibacterial, Anti-Gram+, Anti-Gram-, Antimicrobial |

| S.no. | PepID | DRAMP_ID | Name of the AMP | Activity |
|---|---|---|---|---|
| 1432 | 4741 | DRAMP18675 | Peptide 4 (Mollusca/molluscs/mollusks, invertebrates, animals) | Antibacterial, Anti-Gram+, Anti-Gram-, Antimicrobial |
| 1433 | 4742 | DRAMP18676 | Peptide 5 (Mollusca/molluscs/mollusks, invertebrates, animals) | Antibacterial, Anti-Gram+, Anti-Gram-, Antimicrobial |
| 1434 | 4743 | DRAMP18677 | Peptide 6 (Mollusca/molluscs/mollusks, invertebrates, animals) | Antibacterial, Anti-Gram+, Anti-Gram-, Antimicrobial |
| 1435 | 4744 | DRAMP18678 | Peptide 8 (Mollusca/molluscs/mollusks, invertebrates, animals) | Antibacterial, Anti-Gram+, Anti-Gram-, Antimicrobial |
| 1436 | 4745 | DRAMP18679 | Peptide 9 (Mollusca/molluscs/mollusks, invertebrates, animals) | Antibacterial, Anti-Gram+, Anti-Gram-, Antimicrobial |
| 1437 | 4759 | DRAMP18693 | Substance P (Mammals, animals) | Antimicrobial, Antibacterial, Antifungal, Anti-Gram+, Anti-Gram-, |
| 1438 | 4760 | DRAMP18694 | substance P antagonist (Mammals, animals) | Antimicrobial, Antibacterial, Antifungal, Anti-Gram+, Anti-Gram-, |
| 1439 | 4762 | DRAMP18696 | HLP1 (Lactotransferrin truncated peptide) | Antibacterial, Anti-Gram+, Anti-Gram-, Antimicrobial |
| 1440 | 4763 | DRAMP18697 | HLP2 (Lactotransferrin truncated peptide) | Antibacterial, Anti-Gram+, Anti-Gram-, Antimicrobial |
| 1441 | 4765 | DRAMP18699 | Pleurain-B1 (Frogs, amphibians, animals) | Antimicrobial, Antibacterial, Antifungal, Anti-Gram+, Anti- Gram-, |
| 1442 | 4766 | DRAMP18700 | Pleurain-C1 (Frogs, amphibians, animals) | Antimicrobial, Antibacterial, Antifungal, Anti-Gram+, Anti-Gram-, |
| 1443 | 4767 | DRAMP18701 | Pleurain-D4 (Frogs, amphibians, animals) | Antimicrobial, Antibacterial, Antifungal, Anti-Gram+, Anti-Gram-, |
| 1444 | 4768 | DRAMP18702 | Pleurain-E1 (Frogs, amphibians, animals) | Antimicrobial, Antibacterial, Antifungal, Anti-Gram+, Anti-Gram-, |
| 1445 | 4769 | DRAMP18703 | Pleurain-G1 (Frogs, amphibians, animals) | Antimicrobial, Antibacterial, Antifungal, Anti-Gram+, Anti-Gram-, |
| 1446 | 4770 | DRAMP18704 | Pleurain-J1 (Frogs, amphibians, animals) | Antimicrobial, Antibacterial, Antifungal, Anti-Gram+, Anti-Gram-, |
| 1447 | 4771 | DRAMP18705 | Pleurain-N1 (Frogs, amphibians, animals) | Antimicrobial, Antibacterial, Antifungal, Anti-Gram+, Anti- Gram-, |
| 1448 | 4772 | DRAMP18706 | Pleurain-R1 (Frogs, amphibians, animals) | Antimicrobial, Antibacterial, Antifungal, Anti-Gram+, Anti- Gram-, |
| 1449 | 4773 | DRAMP18707 | BACTENECIN 7 (bac 7, Pro-rich; bovine cathelicidin, cattle, | Antibacterial, Anti-Gram+, Anti-Gram-, Antimicrobial |
| 1450 | 4774 | DRAMP18708 | Dermaseptin-S4 (DRS-S4, DS4; frog, amphibians, animals) | Antimicrobial, Antibacterial, Antifungal, Anti-Gram+, Anti-Gram-, |
| 1451 | 4778 | DRAMP18712 | Styelin A (Tunicate, invertebrates, animals) | Antibacterial, Anti-Gram+, Anti-Gram-, Antimicrobial |
| 1452 | 4779 | DRAMP18713 | Styelin B (Tunicate, invertebrates, animals) | Antibacterial, Anti-Gram+, Anti-Gram-, Antimicrobial |
| 1453 | 4783 | DRAMP18717 | Charybdotoxin (Yellow scorpions, arachnids, Chelicerata,arthropods, | Antimicrobial, Antibacterial, Antifungal, Anti-Gram+, Anti-Gram-, |
| 1454 | 4785 | DRAMP18719 | SAAP fraction 3 (Surfactant-associated anionic peptides; Asp- rich; | Antibacterial, Anti-Gram+, Anti-Gram-, Antimicrobial |
| 1455 | 4787 | DRAMP18721 | Hinnavin I (Hin I; insects, arthropods, invertebrates, animals) | Antimicrobial, Antibacterial, Antifungal, Anti-Gram+, Anti-Gram-, |
| 1456 | 4790 | DRAMP18724 | Oncorhyncin III (Oncorhyncin-3, histone-derived; fish, animals) | Antibacterial, Anti-Gram+, Anti-Gram-, Antimicrobial |
| 1457 | 4791 | DRAMP03024 | Mastoparan B (MP-B; insects, arthropods, invertebrates, animals) | Antimicrobial, Antibacterial, Anti-Gram+, Anti-Gram-, Anti Mammalian cells, Anti-cancer |
| 1458 | 4818 | DRAMP01826 | RV-23 (Frogs, amphibians, animals) | Antimicrobial, Antibacterial, Anti-Gram+, Anti-Gram- |
| 1459 | 4820 | DRAMP03025 | Mastoparan M | Antimicrobial, Antibacterial, Anti-gram+, Anti-gram- |
| 1460 | 4825 | DRAMP03812 | Pardaxin P-4 (Pardaxin P1a; Pardaxin Pa4) | Antimicrobial, Antibacterial, Anti-Gram+, Anti-Gram-, Cytotoxic |
| 1461 | 4839 | DRAMP20771 | Acanthaporin (parasite, amoebozoa, protozoa, protists) | Antimicrobial, Antibacterial, Anti-Gram+, Anti-Gram- |
| 1462 | 4840 | DRAMP20772 | cPcAMP1/26 (ciliate, Protists) | Antimicrobial, Antibacterial, Anti-Gram+, Anti-Gram- |
| 1463 | 4842 | DRAMP20776 | HaA4 (beetles,insects,animals) | Antimicrobial, Antibacterial, Anti-Gram+, Anti-Gram- |
| 1464 | 4843 | DRAMP20777 | Cath-BF | Antimicrobial, Antibacterial, Anti-Gram+, Anti-Gram- |
| 1465 | 4844 | DRAMP20778 | Temporin-SHf (frogs,amphibians,animals) | Antimicrobial, Antibacterial, Anti-Gram+, Anti-Gram-, Antifungal |
| 1466 | 4845 | DRAMP20779 | Halictine 1 (bees,insects,animals) | Antimicrobial, Antibacterial, Anti-Gram+, Anti-Gram- |
| 1467 | 4846 | DRAMP20780 | Halictine 2 (bees,insects,animals) | Antimicrobial, Antibacterial, Anti-Gram+, Anti-Gram- |
| 1468 | 4847 | DRAMP20781 | Panurgine 1(bees,insects,animals) | Antimicrobial, Antibacterial, Anti-Gram+, Anti-Gram- |
| 1469 | 4848 | DRAMP20782 | Pleurain-D1 (Frogs,amphibians,animals) | Antimicrobial, Antibacterial, Anti-Gram+, Anti-Gram- |
| 1470 | 4849 | DRAMP20783 | Pleurain-M1 (Frogs,amphibians,animals) | Antimicrobial, Antibacterial, Anti-Gram+, Anti-Gram- |
| 1471 | 4850 | DRAMP20784 | Megin 1 | Antimicrobial, Antibacterial, Anti-Gram+, Anti-Gram-, Antifungal |
| 1472 | 4851 | DRAMP20785 | Megin 2 | Antimicrobial, Antibacterial, Anti-Gram+, Anti-Gram-, Antifungal |
| 1473 | 4852 | DRAMP20786 | mini-ChBac7.5N alpha | Antimicrobial, Antibacterial, Anti-Gram+, Anti-Gram-, Antifungal |
| 1474 | 4853 | DRAMP20787 | mini-ChBac7.5N beta | Antimicrobial, Antibacterial, Anti-Gram+, Anti-Gram-, Antifungal |
| 1475 | 4856 | DRAMP20790 | Cecropin B (Insects, arthropods, invertebrates, animals) | Antimicrobial, Antibacterial, Anti-Gram+, Anti-Gram- |
| 1476 | 4858 | DRAMP20792 | Maculatin 1.1 (Frog, amphibians, animals) | Antimicrobial, Antibacterial, Anti-Gram+, Anti-Gram- |
| 1477 | 4863 | DRAMP20797 | Uperin 3.6 (Toad, amphibians, animals) | Antimicrobial, Antibacterial, Anti-Gram+, Anti-Gram- |
| 1478 | 4864 | DRAMP20798 | Lingual antimicrobial peptide (LAP, beta defensin, cattle, ruminant, | Antimicrobial, Antibacterial, Anti-Gram+, Anti-Gram-, Antifungal |
| 1479 | 4867 | DRAMP20801 | Mussel Defensin MGD-1 (Mediterranean mussel defensin 1; | Antimicrobial, Antibacterial, Anti-Gram+, Anti-Gram- |
| 1480 | 4868 | DRAMP20802 | CPF-AM1 (caerulein precursor fragment-AM1, frogs, amphibians, | Antimicrobial, Antibacterial, Anti-Gram+, Anti-Gram- |
| 1481 | 4869 | DRAMP20803 | moronecidin-like peptide | Antimicrobial, Antibacterial, Anti-Gram+, Anti-Gram-, Antifungal |
| 1482 | 4870 | DRAMP20804 | Al-hemocidins 2 (Hb-1 truncated peptide) | Antimicrobial, Antibacterial, Anti-Gram+, Anti-Gram- |
| 1483 | 4871 | DRAMP20805 | Apo5 APOC164-88 | Antimicrobial, Antibacterial, Anti-Gram+, Anti-Gram- |
| 1484 | 4872 | DRAMP20806 | Apo6 APOC167-88 | Antimicrobial, Antibacterial, Anti-Gram+, Anti-Gram- |

| S.no. | PepID | DRAMP_ID | Name of the AMP | Activity |
|---|---|---|---|---|
| 1485 | 4873 | DRAMP20807 | A1P394-428 | Antimicrobial, Antibacterial, Anti-Gram+, Anti-Gram- |
| 1486 | 4874 | DRAMP20808 | RI21 (PMAP-36 peptide derivative) | Antimicrobial, Antibacterial, Anti-Gram+, Anti-Gram-, Antifungal |
| 1487 | 4875 | DRAMP20809 | RI18 (PMAP-36 peptide derivative) | Antimicrobial, Antibacterial, Anti-Gram+, Anti-Gram-, Antifungal |
| 1488 | 4876 | DRAMP20810 | TI15 (PMAP-36 peptide derivative) | Antimicrobial, Antibacterial, Anti-Gram+, Anti-Gram-, Antifungal |
| 1489 | 4877 | DRAMP20811 | RI12 (PMAP-36 peptide derivative) | Antimicrobial, Antibacterial, Anti-Gram+, Anti-Gram-, Antifungal |
| 1490 | 4878 | DRAMP20812 | K8 | Antimicrobial, Antibacterial, Anti-Gram+, Anti-Gram- |
| 1491 | 4879 | DRAMP20813 | L1K8 | Antimicrobial, Antibacterial, Anti-Gram+, Anti-Gram- |
| 1492 | 4880 | DRAMP20814 | S1K8 | Antimicrobial, Antibacterial, Anti-Gram+, Anti-Gram- |
| 1493 | 4881 | DRAMP20815 | F1K8 | Antimicrobial, Antibacterial, Anti-Gram+, Anti-Gram- |
| 1494 | 4882 | DRAMP20816 | K1K8 | Antimicrobial, Antibacterial, Anti-Gram+, Anti-Gram- |
| 1495 | 4883 | DRAMP20817 | RR12 | Antimicrobial, Antibacterial, Anti-Gram+, Anti-Gram- |
| 1496 | 4884 | DRAMP20818 | RR12Wpolar | Antimicrobial, Antibacterial, Anti-Gram+, Anti-Gram- |
| 1497 | 4885 | DRAMP20819 | RR12Whydro | Antimicrobial, Antibacterial, Anti-Gram+, Anti-Gram- |
| 1498 | 4886 | DRAMP20820 | FV7 | Antimicrobial, Antibacterial, Anti-Gram+, Anti-Gram- |
| 1499 | 4887 | DRAMP20821 | FV-LL (FV7 and LL(LL-37,(17-29)) hybrid peptide) | Antimicrobial, Antibacterial, Anti-Gram+, Anti-Gram- |
| 1500 | 4888 | DRAMP20822 | FV-MA (FV7 and MA(Magainin 2 (9-21)) hybrid peptide) | Antimicrobial, Antibacterial, Anti-Gram+, Anti-Gram- |
| 1501 | 4889 | DRAMP20823 | FV-CE (FV7 and CE(Cecropin A (1 | Antimicrobial, Antibacterial, Anti-Gram+, Anti-Gram- |
| 1502 | 4890 | DRAMP20824 | AM-CATH36 | Antimicrobial, Antibacterial, Anti-Gram+, Anti-Gram- |
| 1503 | 4892 | DRAMP20826 | AM-CATH21 | Antimicrobial, Antibacterial, Anti-Gram+, Anti-Gram- |
| 1504 | 4893 | DRAMP20827 | TB_L1FK | Antimicrobial, Antibacterial, Anti-Gram+, Anti-Gram- |
| 1505 | 4894 | DRAMP20828 | TB_KKG6A | Antimicrobial, Antibacterial, Anti-Gram+, Anti-Gram- |
| 1506 | 4895 | DRAMP20831 | IsCT1L1 | Antimicrobial, Antibacterial, Anti-Gram+, Anti-Gram- |
| 1507 | 4896 | DRAMP20832 | Polybia-MP1S-D8N | Antimicrobial, Antibacterial, Anti-Gram+, Anti-Gram- |
| 1508 | 4897 | DRAMP20833 | [Pro3,DLeu9]TL(1) (Temporin L peptide derivative) | Antimicrobial, Antibacterial, Anti-Gram+, Anti-Gram- |
| 1509 | 4898 | DRAMP20834 | PLS | Antimicrobial, Antibacterial, Anti-Gram+, Anti-Gram- |
| 1510 | 4899 | DRAMP20837 | Pb-CATH1 Python bivittatus antimicrobial peptides peptide derivative | Antimicrobial, Antibacterial, Anti-Gram+, Anti-Gram- |
| 1511 | 4900 | DRAMP20838 | Pb-CATH4 bivittatus antimicrobial peptides peptide derivative | Antimicrobial, Antibacterial, Anti-Gram+, Anti-Gram- |
| 1512 | 4901 | DRAMP20839 | Xylopin | Antimicrobial, Antibacterial, Anti-Gram+, Anti-Gram- |
| 1513 | 4902 | DRAMP20841 | C1b | Antimicrobial, Antibacterial, Anti-Gram+, Anti-Gram- |
| 1514 | 4903 | DRAMP20842 | C1b(1-11) | Antimicrobial, Antibacterial, Anti-Gram+, Anti-Gram- |
| 1515 | 4904 | DRAMP20843 | C1b(1-13) | Antimicrobial, Antibacterial, Anti-Gram+, Anti-Gram- |
| 1516 | 4905 | DRAMP20844 | C1b(3-13) | Antimicrobial, Antibacterial, Anti-Gram+, Anti-Gram- |
| 1517 | 4906 | DRAMP20845 | C1b(3-11) | Antimicrobial, Antibacterial, Anti-Gram+, Anti-Gram- |
| 1518 | 4907 | DRAMP20846 | C1b(3-12) | Antimicrobial, Antibacterial, Anti-Gram+, Anti-Gram- |
| 1519 | 4908 | DRAMP20847 | C1b(4-13) | Antimicrobial, Antibacterial, Anti-Gram+, Anti-Gram- |
| 1520 | 4909 | DRAMP20848 | [K4]C1b(3-11) | Antimicrobial, Antibacterial, Anti-Gram+, Anti-Gram- |
| 1521 | 4910 | DRAMP20849 | [R4]C1b(3-11) | Antimicrobial, Antibacterial, Anti-Gram+, Anti-Gram- |
| 1522 | 4911 | DRAMP20850 | [K4,K10]C1b(3-13) | Antimicrobial, Antibacterial, Anti-Gram+, Anti-Gram- |
| 1523 | 4912 | DRAMP20851 | [R4,R10]C1b(3-13) | Antimicrobial, Antibacterial, Anti-Gram+, Anti-Gram- |
| 1524 | 4920 | DRAMP20859 | TT(1-24) | Antimicrobial, Antibacterial, Anti-Gram+, Anti-Gram- |
| 1525 | 4921 | DRAMP20860 | TT(1-35) | Antimicrobial, Antibacterial, Anti-Gram+, Anti-Gram- |
| 1526 | 4923 | DRAMP20862 | rtCATH2(5-40) | Antimicrobial, Antibacterial, Anti-Gram+, Anti-Gram- |
| 1527 | 4924 | DRAMP20863 | rtCATH2(1-40) | Antimicrobial, Antibacterial, Anti-Gram+, Anti-Gram- |
| 1528 | 4925 | DRAMP20864 | SF(18-45) | Antimicrobial, Antibacterial, Anti-Gram+, Anti-Gram- |
| 1529 | 4926 | DRAMP20865 | the dimeric RRWQWR motif peptide molecule | Antimicrobial, Antibacterial, Anti-Gram+, Anti-Gram- |
| 1530 | 4927 | DRAMP20866 | the tetrameric RRWQWR motif peptide molecule | Antimicrobial, Antibacterial, Anti-Gram+, Anti-Gram- |
| 1531 | 4928 | DRAMP20867 | the palindromic RRWQWR motif peptide molecule | Antimicrobial, Antibacterial, Anti-Gram+, Anti-Gram- |
| 1532 | 4929 | DRAMP20868 | H4 | Antimicrobial, Antibacterial, Anti-Gram+, Anti-Gram- |
| 1533 | 4930 | DRAMP20869 | Pal-ano-9 (Pal-anoplin peptide derivative) | Antimicrobial, Antibacterial, Anti-Gram+, Anti-Gram-, Antifungal |
| 1534 | 4931 | DRAMP20870 | Pal-ano-8 (Pal-anoplin peptide derivative) | Antimicrobial, Antibacterial, Anti-Gram+, Anti-Gram-, Antifungal |
| 1535 | 4932 | DRAMP20871 | Pal-ano-7 (Pal-anoplin peptide derivative) | Antimicrobial, Antibacterial, Anti-Gram+, Anti-Gram-, Antifungal |
| 1536 | 4933 | DRAMP20872 | Pal-ano-6 (Pal-anoplin peptide derivative) | Antimicrobial, Antibacterial, Anti-Gram+, Anti-Gram-, Antifungal |
| 1537 | 4934 | DRAMP20873 | Pal-ano-5 (Pal-anoplin peptide derivative) | Antimicrobial, Antibacterial, Anti-Gram+, Anti-Gram-, Antifungal |

| S.no. | PepID | DRAMP_ID | Name of the AMP | Activity |
|---|---|---|---|---|
| 1538 | 4935 | DRAMP20874 | Chensinin-1b | Antimicrobial, Antibacterial, Anti-Gram+, Anti-Gram- |
| 1539 | 4936 | DRAMP20875 | OA-C1b | Antimicrobial, Antibacterial, Anti-Gram+, Anti-Gram- |
| 1540 | 4937 | DRAMP20876 | LA-C1b | Antimicrobial, Antibacterial, Anti-Gram+, Anti-Gram- |
| 1541 | 4938 | DRAMP20877 | PA-C1b | Antimicrobial, Antibacterial, Anti-Gram+, Anti-Gram- |
| 1542 | 4939 | DRAMP20878 | rVpDef | Antimicrobial, Antibacterial, Anti-Gram+, Anti-Gram- |
| 1543 | 4940 | DRAMP20879 | DAN1 | Antimicrobial, Antibacterial, Anti-Gram+, Anti-Gram- |
| 1544 | 4941 | DRAMP20880 | DAN2 | Antimicrobial, Antibacterial, Anti-Gram+, Anti-Gram-, Antifungal |
| 1545 | 4944 | DRAMP20883 | Cath-A | Antimicrobial, Antibacterial, Anti-Gram+, Anti-Gram- |
| 1546 | 4945 | DRAMP20884 | Cath-B | Antimicrobial, Antibacterial, Anti-Gram+, Anti-Gram- |
| 1547 | 4948 | DRAMP20887 | NCP-2 (CTX-1 peptide derivative) | Antimicrobial, Antibacterial, Anti-Gram+, Anti-Gram-, Antifungal |
| 1548 | 4949 | DRAMP20888 | NCP-3 (CTX-1 peptide derivative) | Antimicrobial, Antibacterial, Anti-Gram+, Anti-Gram-, Antifungal |
| 1549 | 4954 | DRAMP20893 | I16A | Antimicrobial, Antibacterial, Anti-Gram+, Anti-Gram-, Antifungal |
| 1550 | 4955 | DRAMP20894 | L19H/I20H | Antimicrobial, Antibacterial, Anti-Gram+, Anti-Gram-, Antifungal |
| 1551 | 4956 | DRAMP20895 | F1A/I2A | Antimicrobial, Antibacterial, Anti-Gram+, Anti-Gram-, Antifungal |
| 1552 | 4961 | DRAMP20900 | A12I/A15I | Antimicrobial, Antibacterial, Anti-Gram+, Anti-Gram-, Antifungal |
| 1553 | 4962 | DRAMP20901 | A12V/A15H | Antimicrobial, Antibacterial, Anti-Gram+, Anti-Gram-, Antifungal |
| 1554 | 4964 | DRAMP20903 | dC2 | Antimicrobial, Antibacterial, Anti-Gram+, Anti-Gram-, Antifungal |
| 1555 | 4965 | DRAMP20904 | R18S/R21H | Antimicrobial, Antibacterial, Anti-Gram+, Anti-Gram-, Antifungal |
| 1556 | 4967 | DRAMP20906 | dN2 | Antimicrobial, Antibacterial, Anti-Gram+, Anti-Gram-, Antifungal |
| 1557 | 4968 | DRAMP20907 | dN4 | Antimicrobial, Antibacterial, Anti-Gram+, Anti-Gram-, Antifungal |
| 1558 | 4971 | DRAMP20910 | RN7-IN7(designed based on indolicidin and ranalexin) | Antimicrobial, Antibacterial, Anti-Gram+, Anti-Gram- |
| 1559 | 4973 | DRAMP20912 | RN7-IN9(designed based on indolicidin and ranalexin) | Antimicrobial, Antibacterial, Anti-Gram+, Anti-Gram- |
| 1560 | 4974 | DRAMP20913 | Myxinidin (G1) | Antimicrobial, Antibacterial, Anti-Gram+, Anti-Gram- |
| 1561 | 4975 | DRAMP20914 | Myxinidin (I2) | Antimicrobial, Antibacterial, Anti-Gram+, Anti-Gram- |
| 1562 | 4976 | DRAMP20915 | Myxinidin (H3) | Antimicrobial, Antibacterial, Anti-Gram+, Anti-Gram- |
| 1563 | 4977 | DRAMP20916 | Myxinidin (D4) | Antimicrobial, Antibacterial, Anti-Gram+, Anti-Gram- |
| 1564 | 4978 | DRAMP20917 | Myxinidin (I5) | Antimicrobial, Antibacterial, Anti-Gram+, Anti-Gram- |
| 1565 | 4979 | DRAMP20918 | Myxinidin (L6) | Antimicrobial, Antibacterial, Anti-Gram+, Anti-Gram- |
| 1566 | 4980 | DRAMP20919 | Myxinidin (K7) | Antimicrobial, Antibacterial, Anti-Gram+, Anti-Gram- |
| 1567 | 4981 | DRAMP20920 | Myxinidin (Y8) | Antimicrobial, Antibacterial, Anti-Gram+, Anti-Gram- |
| 1568 | 4982 | DRAMP20921 | Myxinidin (G9) | Antimicrobial, Antibacterial, Anti-Gram+, Anti-Gram- |
| 1569 | 4983 | DRAMP20922 | Myxinidin (K10) | Antimicrobial, Antibacterial, Anti-Gram+, Anti-Gram- |
| 1570 | 4984 | DRAMP20923 | Myxinidin (P11) | Antimicrobial, Antibacterial, Anti-Gram+, Anti-Gram- |
| 1571 | 4985 | DRAMP20924 | Myxinidin (S12) | Antimicrobial, Antibacterial, Anti-Gram+, Anti-Gram- |
| 1572 | 4986 | DRAMP20925 | MH3R | Antimicrobial, Antibacterial, Anti-Gram+, Anti-Gram- |
| 1573 | 4987 | DRAMP20926 | IN1(designed based on indolicidin and ranalexin) | Antimicrobial, Antibacterial, Anti-Gram+, Anti-Gram- |
| 1574 | 4988 | DRAMP20927 | IN2(designed based on indolicidin and ranalexin) | Antimicrobial, Antibacterial, Anti-Gram+, Anti-Gram- |
| 1575 | 4989 | DRAMP20928 | IN3(designed based on indolicidin and ranalexin) | Antimicrobial, Antibacterial, Anti-Gram+, Anti-Gram- |
| 1576 | 4990 | DRAMP20929 | RN7-IN6(designed based on indolicidin and ranalexin) | Antimicrobial, Antibacterial, Anti-Gram+, Anti-Gram- |
| 1577 | 4991 | DRAMP20930 | BP100-Ala-NH-C16H33 | Antimicrobial, Antibacterial, Anti-Gram+, Anti-Gram- |
| 1578 | 4995 | DRAMP20935 | Macropin 1(solitary bee, insects, animals) | Antimicrobial, Antibacterial, Anti-Gram+, Anti-Gram-, Antifungal |
| 1579 | 4996 | DRAMP20936 | ΔPb-CATH1 | Antimicrobial, Antibacterial, Anti-Gram+, Anti-Gram- |
| 1580 | 4997 | DRAMP20937 | Pb-CATH3 | Antimicrobial, Antibacterial, Anti-Gram+, Anti-Gram- |
| 1581 | 4998 | DRAMP20938 | Cbf-14 | Antimicrobial, Antibacterial, Anti-Gram+, Anti-Gram- |
| 1582 | 4999 | DRAMP20939 | D-Cbf-14 | Antimicrobial, Antibacterial, Anti-Gram+, Anti-Gram- |
| 1583 | 5001 | DRAMP20941 | [Pro3,DLeu9]TL(8) (Temporin L peptide derivative) | Antimicrobial, Antibacterial, Anti-Gram+, Anti-Gram-, Antifungal |
| 1584 | 5002 | DRAMP20942 | [Pro3,DLeu9]TL(9) (Temporin L peptide derivative) | Antimicrobial, Antibacterial, Anti-Gram+, Anti-Gram-, Antifungal |
| 1585 | 5003 | DRAMP20943 | [Pro3,DLeu9]TL(10) (Temporin L peptide derivative) | Antimicrobial, Antibacterial, Anti-Gram+, Anti-Gram-, Antifungal |
| 1586 | 5004 | DRAMP20944 | [Pro3,DLeu9]TL(11) (Temporin L peptide derivative) | Antimicrobial, Antibacterial, Anti-Gram+, Anti-Gram-, Antifungal |
| 1587 | 5005 | DRAMP20945 | Recombinant Cecropin A (1–8)–LL37 (17–30) (C–L) | Antimicrobial, Antibacterial, Anti-Gram+, Anti-Gram- |
| 1588 | 5014 | DRAMP20955 | L31-P113 | Antimicrobial, Antibacterial, Anti-Gram+, Anti-Gram-, Antifungal |
| 1589 | 5015 | DRAMP20956 | AL32-P113 | Antimicrobial, Antibacterial, Anti-Gram+, Anti-Gram-, Antifungal |
| 1590 | 5016 | DRAMP20957 | StigA6 | Antimicrobial, Antibacterial, Anti-Gram+, Anti-Gram-, Antifungal, Antiparasitic, Antiproliferative |

| S.no. | PepID | DRAMP_ID | Name of the AMP | Activity |
|---|---|---|---|---|
| 1591 | 5017 | DRAMP20958 | StigA16 | Antimicrobial, Antibacterial, Anti-Gram+, Anti-Gram-, Antifungal, Antiparasitic, Antiproliferative |
| 1592 | 5021 | DRAMP20963 | Cp1 alpha s1-casein peptide derivative | Antimicrobial, Antibacterial, Anti-Gram+, Anti-Gram-, low hemolytic and toxic effects |
| 1593 | 5022 | DRAMP20964 | Synthesized Cecropin A (1–8)–LL37 (17–30) (C–L) | Antimicrobial, Antibacterial, Anti-Gram+, Anti-Gram- |
| 1594 | 5023 | DRAMP20965 | LPcin-YK3 (bovine cathelicidin, cattle, ruminant, mammals, animals) | Antimicrobial, Antibacterial, Anti-Gram+, Anti-Gram-, Antifungal |
| 1595 | 5024 | DRAMP20966 | andricin B (Andrias davidianus, Amphibians, Animals) | Antimicrobial, Antibacterial, Anti-Gram+, Anti-Gram-, Antifungal |
| 1596 | 5025 | DRAMP20967 | andricin 01 (Andrias davidianus, Amphibians, Animals) | Antimicrobial, Antibacterial, Anti-Gram+, Anti-Gram- |
| 1597 | 5026 | DRAMP20968 | Catesbeianin-1 (Ranidae, Anura, Amphibia, Animals) | Antimicrobial, Antibacterial, Anti-Gram+, Anti-Gram- |
| 1598 | 5027 | DRAMP20969 | HJH-1 (bovine cathelicidin, cattle, ruminant, mammals, animals) | Antimicrobial, Antibacterial, Anti-Gram+, Anti-Gram-, Antifungal |
| 1599 | 5028 | DRAMP20970 | P3 (bovine cathelicidin, cattle, ruminant, mammals, animals) | Antimicrobial, Antibacterial, Anti-Gram+, Anti-Gram-, Antifungal |
| 1600 | 5029 | DRAMP20971 | JH-0 (Derived from P3) | Antimicrobial, Antibacterial, Anti-Gram+, Anti-Gram- |
| 1601 | 5030 | DRAMP20972 | JH-1 (Derived from P3) | Antimicrobial, Antibacterial, Anti-Gram+, Anti-Gram- |
| 1602 | 5031 | DRAMP20973 | JH-2 (Derived from P3) | Antimicrobial, Antibacterial, Anti-Gram+, Anti-Gram-, Antifungal |
| 1603 | 5032 | DRAMP20974 | JH-3 (Derived from P3) | Antimicrobial, Antibacterial, Anti-Gram+, Anti-Gram-, Antifungal |
| 1604 | 5033 | DRAMP20975 | OH-CM6 (Derived from OH-CATH30) | Antimicrobial, Antibacterial, Anti-Gram+, Anti-Gram- |
| 1605 | 5034 | DRAMP20976 | adevonin (Derived from Adenanthera pavonina trypsin inhibitor | Antimicrobial, Antibacterial, Anti-Gram+, Anti-Gram- |
| 1606 | 5035 | DRAMP20977 | Anoplin-1 (Derived from Anoplin) | Antimicrobial, Antibacterial, Anti-Gram+, Anti-Gram- |
| 1607 | 5036 | DRAMP20978 | Anoplin-2 (Derived from Anoplin) | Antimicrobial, Antibacterial, Anti-Gram+, Anti-Gram- |
| 1608 | 5037 | DRAMP20979 | Anoplin-3 (Derived from Anoplin) | Antimicrobial, Antibacterial, Anti-Gram+, Anti-Gram- |
| 1609 | 5038 | DRAMP20980 | Anoplin-4 (Derived from Anoplin) | Antimicrobial, Antibacterial, Anti-Gram+, Anti-Gram- |
| 1610 | 5039 | DRAMP20981 | CPF-C1 (Frogs, Amphibians, Animals) | Antimicrobial, Antibacterial, Anti-Gram+, Anti-Gram- |
| 1611 | 5040 | DRAMP20982 | CPF-1 (Derived from CPF-C1) | Antimicrobial, Antibacterial, Anti-Gram+, Anti-Gram- |
| 1612 | 5041 | DRAMP20983 | CPF-2 (Derived from CPF-C1) | Antimicrobial, Antibacterial, Anti-Gram+, Anti-Gram- |
| 1613 | 5042 | DRAMP20984 | CPF-3 (Derived from CPF-C1) | Antimicrobial, Antibacterial, Anti-Gram+, Anti-Gram- |
| 1614 | 5043 | DRAMP20985 | CPF-4 (Derived from CPF-C1) | Antimicrobial, Antibacterial, Anti-Gram+, Anti-Gram- |
| 1615 | 5044 | DRAMP20986 | CPF-5 (Derived from CPF-C1) | Antimicrobial, Antibacterial, Anti-Gram+, Anti-Gram- |
| 1616 | 5045 | DRAMP20987 | CPF-6 (Derived from CPF-C1) | Antimicrobial, Antibacterial, Anti-Gram+, Anti-Gram- |
| 1617 | 5046 | DRAMP20988 | CPF-7 (Derived from CPF-C1) | Antimicrobial, Antibacterial, Anti-Gram+, Anti-Gram- |
| 1618 | 5047 | DRAMP20989 | CPF-8 (Derived from CPF-C1) | Antimicrobial, Antibacterial, Anti-Gram+, Anti-Gram- |
| 1619 | 5048 | DRAMP20990 | CPF-9 (Derived from CPF-C1) | Antimicrobial, Antibacterial, Anti-Gram+, Anti-Gram- |
| 1620 | 5049 | DRAMP20991 | CPF-10 (Derived from CPF-C1) | Antimicrobial, Antibacterial, Anti-Gram+, Anti-Gram- |
| 1621 | 5050 | DRAMP20992 | CPF-11 (Derived from CPF-C1) | Antimicrobial, Antibacterial, Anti-Gram+, Anti-Gram- |
| 1622 | 5051 | DRAMP20993 | CPF-12 (Derived from CPF-C1) | Antimicrobial, Antibacterial, Anti-Gram+, Anti-Gram- |
| 1623 | 5052 | DRAMP20994 | anoplin analog 4 | Antimicrobial, Antibacterial, Anti-Gram+, Anti-Gram-, Antifungal |
| 1624 | 5053 | DRAMP20995 | anoplin analog 5 | Antimicrobial, Antibacterial, Anti-Gram+, Anti-Gram-, Antifungal |
| 1625 | 5054 | DRAMP20996 | anoplin analog 6 | Antimicrobial, Antibacterial, Anti-Gram+, Anti-Gram-, Antifungal |
| 1626 | 5055 | DRAMP20997 | anoplin analog 7 | Antimicrobial, Antibacterial, Anti-Gram+, Anti-Gram- |
| 1627 | 5056 | DRAMP20998 | anoplin analog 8 | Antimicrobial, Antibacterial, Anti-Gram+, Anti-Gram- |
| 1628 | 5057 | DRAMP20999 | anoplin analog 9 | Antimicrobial, Antibacterial, Anti-Gram+, Anti-Gram- |
| 1629 | 5058 | DRAMP21000 | cGm (Derived from Gm) | Antimicrobial, Antibacterial, Anti-Gram+, Anti-Gram-, Antifungal, Antitumor |
| 1630 | 5059 | DRAMP21001 | [Y7W]cGm (Derived from Gm) | Antimicrobial, Antibacterial, Anti-Gram+, Anti-Gram-, Antitumor |
| 1631 | 5060 | DRAMP21002 | [Y14W]cGm (Derived from Gm) | Antimicrobial, Antibacterial, Anti-Gram+, Anti-Gram-, Antitumor |
| 1632 | 5061 | DRAMP21003 | [K8R]cGm (Derived from Gm) | Antimicrobial, Antibacterial, Anti-Gram+, Anti-Gram-, Antitumor |
| 1633 | 5062 | DRAMP21004 | [Y7W, K8R, Y14W]cGm (Derived from Gm) | Antimicrobial, Antibacterial, Anti-Gram+, Anti-Gram-, Antifungal, Antitumor |
| 1634 | 5063 | DRAMP21005 | [R4A, R18A]cGm (Derived from Gm) | Antimicrobial, Antibacterial, Anti-Gram+, Anti-Gram-, Antifungal, Antitumor |
| 1635 | 5064 | DRAMP21006 | [G1K, K8R]cGm (Derived from Gm) | Antimicrobial, Antibacterial, Anti-Gram+, Anti-Gram-, Antifungal, Antitumor |
| 1636 | 5065 | DRAMP21007 | [C/U]cGm (Derived from Gm) | Antimicrobial, Antibacterial, Anti-Gram+, Anti-Gram-, Antifungal, Antitumor |
| 1637 | 5066 | DRAMP21008 | [L5W]cGm (Derived from Gm) | Antimicrobial, Antibacterial, Anti-Gram+, Anti-Gram-, Antitumor |
| 1638 | 5067 | DRAMP21009 | [D-P L-P]cGm (Derived from Gm) | Antimicrobial, Antibacterial, Anti-Gram+, Anti-Gram-, Antifungal, Antitumor |
| 1639 | 5068 | DRAMP21010 | [G1K, L5Y, K8R]cGm (Derived from Gm) | Antimicrobial, Antibacterial, Anti-Gram+, Anti-Gram-, Antifungal, Antitumor |
| 1640 | 5069 | DRAMP21011 | [C/U, G1K, L5Y, K8R]cGm (Derived from Gm) | Antimicrobial, Antibacterial, Anti-Gram+, Anti-Gram-, Antifungal, Antitumor |
| 1641 | 5070 | DRAMP21012 | NK-2 (Mammals, Animals) | Antimicrobial, Antibacterial, Anti-Gram+, Anti-Gram-, Antifungal, Antitumor |
| 1642 | 5071 | DRAMP21013 | NK-pro (Derived from NK-2) | Antimicrobial, Antibacterial, Anti-Gram+, Anti-Gram-, Antifungal, Antitumor |
| 1643 | 5072 | DRAMP21014 | NK-dpro (Derived from NK-2) | Antimicrobial, Antibacterial, Anti-Gram+, Anti-Gram-, Antifungal, Antitumor |

| S.no. | PepID | DRAMP_ID | Name of the AMP | Activity |
|---|---|---|---|---|
| 1644 | 5073 | DRAMP21015 | A (A1R) (Derived from AR-23) | Antimicrobial, Antibacterial, Anti-Gram+, Anti-Gram- |
| 1645 | 5074 | DRAMP21016 | A (A8R) (Derived from AR-23) | Antimicrobial, Antibacterial, Anti-Gram+, Anti-Gram- |
| 1646 | 5075 | DRAMP21017 | A (I17K) (Derived from AR-23) | Antimicrobial, Antibacterial, Anti-Gram+, Anti-Gram- |
| 1647 | 5076 | DRAMP21018 | A (I17R) (Derived from AR-23) | Antimicrobial, Antibacterial, Anti-Gram+, Anti-Gram- |
| 1648 | 5077 | DRAMP21019 | A (A1R, A8R) (Derived from AR-23) | Antimicrobial, Antibacterial, Anti-Gram+, Anti-Gram- |
| 1649 | 5078 | DRAMP21020 | A (A1R, I17K) (Derived from AR-23) | Antimicrobial, Antibacterial, Anti-Gram+, Anti-Gram- |
| 1650 | 5079 | DRAMP21021 | A (A8R, I17K) (Derived from AR-23) | Antimicrobial, Antibacterial, Anti-Gram+, Anti-Gram- |
| 1651 | 5080 | DRAMP21022 | A (A1R, A8R, I17K) (Derived from AR-23) | Antimicrobial, Antibacterial, Anti-Gram+, Anti-Gram- |
| 1652 | 5081 | DRAMP21023 | A (A1R, A8R, I17R) (Derived from AR-23) | Antimicrobial, Antibacterial, Anti-Gram+, Anti-Gram- |
| 1653 | 5082 | DRAMP21024 | Stigmurin (Tityus, Scorpionida, Arachnida) | Antimicrobial, Antibacterial, Anti-Gram+, Anti-Gram-, Antifungal |
| 1654 | 5083 | DRAMP21025 | StigA25 (Derived from Stigmurin) | Antimicrobial, Antibacterial, Anti-Gram+, Anti-Gram-, Antifungal |
| 1655 | 5084 | DRAMP21026 | StigA31 (Derived from Stigmurin) | Antimicrobial, Antibacterial, Anti-Gram+, Anti-Gram-, Antifungal |
| 1656 | 5085 | DRAMP21027 | K5, 17-DPS3 (Derived from dermaseptin-PS3 (DPS3)) | Antimicrobial, Antibacterial, Anti-Gram+, Anti-Gram-, Antifungal |
| 1657 | 5086 | DRAMP21028 | L10, 11-DPS3 (Derived from dermaseptin-PS3 (DPS3)) | Antimicrobial, Antibacterial, Anti-Gram+, Anti-Gram-, Antifungal |
| 1658 | 5087 | DRAMP21029 | D5R (Derived from HD5) | Antimicrobial, Antibacterial, Anti-Gram+, Anti-Gram-, Antifungal |
| 1659 | 5088 | DRAMP21030 | D5r (Derived from HD5) | Antimicrobial, Antibacterial, Anti-Gram+, Anti-Gram-, Antifungal |
| 1660 | 5089 | DRAMP21031 | MyD5R (Derived from HD5) | Antimicrobial, Antibacterial, Anti-Gram+, Anti-Gram-, Antifungal |
| 1661 | 5090 | DRAMP21032 | MyD5r (Derived from HD5) | Antimicrobial, Antibacterial, Anti-Gram+, Anti-Gram-, Antifungal |
| 1662 | 5091 | DRAMP21033 | LaD5R (Derived from HD5) | Antimicrobial, Antibacterial, Anti-Gram+, Anti-Gram-, Antifungal |
| 1663 | 5092 | DRAMP21034 | LaD5r (Derived from HD5) | Antimicrobial, Antibacterial, Anti-Gram+, Anti-Gram-, Antifungal |
| 1664 | 5093 | DRAMP21035 | AC-UM-14W (De novo synthesis) | Antimicrobial, Antibacterial, Anti-Gram+, Anti-Gram- |
| 1665 | 5094 | DRAMP21036 | PapMA (Derived from Papiliocin and Magainin 2) | Antimicrobial, Antibacterial, Anti-Gram+, Anti-Gram- |
| 1666 | 5095 | DRAMP21037 | PapMA-k (Derived from Papiliocin and Magainin 2) | Antimicrobial, Antibacterial, Anti-Gram+, Anti-Gram- |
| 1667 | 5096 | DRAMP21038 | analog 1 (Derived from Ib-AMP1) | Antimicrobial, Antibacterial, Anti-Gram+, Anti-Gram- |
| 1668 | 5097 | DRAMP21039 | analog 2 (Derived from Ib-AMP2) | Antimicrobial, Antibacterial, Anti-Gram+, Anti-Gram- |
| 1669 | 5098 | DRAMP21040 | analog 3 (Derived from Ib-AMP2) | Antimicrobial, Antibacterial, Anti-Gram+, Anti-Gram- |
| 1670 | 5099 | DRAMP21041 | analog 4 (Derived from Ib-AMP2) | Antimicrobial, Antibacterial, Anti-Gram+, Anti-Gram- |
| 1671 | 5100 | DRAMP21042 | A2 (Derived from Indolicidin (IN)) | Antimicrobial, Antibacterial, Anti-Gram+, Anti-Gram- |
| 1672 | 5101 | DRAMP21043 | A3 (Derived from Indolicidin (IN)) | Antimicrobial, Antibacterial, Anti-Gram+, Anti-Gram- |
| 1673 | 5102 | DRAMP21044 | A4 (Derived from Indolicidin (IN)) | Antimicrobial, Antibacterial, Anti-Gram+, Anti-Gram- |
| 1674 | 5103 | DRAMP21045 | A5 (Derived from Indolicidin (IN)) | Antimicrobial, Antibacterial, Anti-Gram+, Anti-Gram- |
| 1675 | 5104 | DRAMP21046 | A6 (Derived from Indolicidin (IN)) | Antimicrobial, Antibacterial, Anti-Gram+, Anti-Gram- |
| 1676 | 5105 | DRAMP21047 | A7 (Derived from Indolicidin (IN)) | Antimicrobial, Antibacterial, Anti-Gram+, Anti-Gram- |
| 1677 | 5106 | DRAMP21048 | peptide 6 (Derived from seq2) | Antimicrobial, Antibacterial, Anti-Gram+, Anti-Gram- |
| 1678 | 5107 | DRAMP21049 | peptide 6.2 (Derived from seq2) | Antimicrobial, Antibacterial, Anti-Gram+, Anti-Gram- |
| 1679 | 5108 | DRAMP21050 | TP1[K1A] (Derived from TP1) | Antimicrobial, Antibacterial, Anti-Gram+, Anti-Gram-, Antifungal |
| 1680 | 5109 | DRAMP21051 | TP1[W2A] (Derived from TP1) | Antimicrobial, Antibacterial, Anti-Gram+, Anti-Gram-, Antifungal |
| 1681 | 5110 | DRAMP21052 | TP1[C3A, C16S] (Derived from TP1) | Antimicrobial, Antibacterial, Anti-Gram+, Anti-Gram-, Antifungal |
| 1682 | 5111 | DRAMP21053 | TP1[F4A] (Derived from TP1) | Antimicrobial, Antibacterial, Anti-Gram+, Anti-Gram-, Antifungal |
| 1683 | 5112 | DRAMP21054 | TP1[R5A] (Derived from TP1) | Antimicrobial, Antibacterial, Anti-Gram+, Anti-Gram-, Antifungal |
| 1684 | 5113 | DRAMP21055 | TP1[V6A] (Derived from TP1) | Antimicrobial, Antibacterial, Anti-Gram+, Anti-Gram-, Antifungal |
| 1685 | 5114 | DRAMP21056 | TP1[C7A, C12S] (Derived from TP1) | Antimicrobial, Antibacterial, Anti-Gram+, Anti-Gram-, Antifungal |
| 1686 | 5115 | DRAMP21057 | TP1[Y8A] (Derived from TP1) | Antimicrobial, Antibacterial, Anti-Gram+, Anti-Gram-, Antifungal |
| 1687 | 5116 | DRAMP21058 | TP1[R9A] (Derived from TP1) | Antimicrobial, Antibacterial, Anti-Gram+, Anti-Gram-, Antifungal |
| 1688 | 5117 | DRAMP21059 | TP1[G10A] (Derived from TP1) | Antimicrobial, Antibacterial, Anti-Gram+, Anti-Gram-, Antifungal |
| 1689 | 5118 | DRAMP21060 | TP1[I11A] (Derived from TP1) | Antimicrobial, Antibacterial, Anti-Gram+, Anti-Gram-, Antifungal |
| 1690 | 5119 | DRAMP21061 | TP1[C7S, C12A] (Derived from TP1) | Antimicrobial, Antibacterial, Anti-Gram+, Anti-Gram-, Antifungal |
| 1691 | 5120 | DRAMP21062 | TP1[Y13A] (Derived from TP1) | Antimicrobial, Antibacterial, Anti-Gram+, Anti-Gram-, Antifungal |
| 1692 | 5121 | DRAMP21063 | TP1[R14A] (Derived from TP1) | Antimicrobial, Antibacterial, Anti-Gram+, Anti-Gram-, Antifungal |
| 1693 | 5122 | DRAMP21064 | TP1[R15A] (Derived from TP1) | Antimicrobial, Antibacterial, Anti-Gram+, Anti-Gram-, Antifungal |
| 1694 | 5123 | DRAMP21065 | TP1[C3S, C16A] (Derived from TP1) | Antimicrobial, Antibacterial, Anti-Gram+, Anti-Gram-, Antifungal |
| 1695 | 5124 | DRAMP21066 | TP1[R17A] (Derived from TP1) | Antimicrobial, Antibacterial, Anti-Gram+, Anti-Gram-, Antifungal |
| 1696 | 5125 | DRAMP21067 | TP1[C3A, C16A] (Derived from TP1) | Antimicrobial, Antibacterial, Anti-Gram+, Anti-Gram-, Antifungal |

| S.no. | PepID | DRAMP_ID | Name of the AMP | Activity |
|---|---|---|---|---|
| 1697 | 5126 | DRAMP21068 | TP1[C7A, C12A] (Derived from TP1) | Antimicrobial, Antibacterial, Anti-Gram+, Anti-Gram-, Antifungal |
| 1698 | 5127 | DRAMP21069 | TP1[C3A, C7A, C12A, C16A] (Derived from TP1) | Antimicrobial, Antibacterial, Anti-Gram+, Anti-Gram-, Antifungal |
| 1699 | 5128 | DRAMP21070 | TP1[V6R, R9A] (Derived from TP1) | Antimicrobial, Antibacterial, Anti-Gram+, Anti-Gram-, Antifungal |
| 1700 | 5129 | DRAMP21071 | TP1[K1R] (Derived from TP1) | Antimicrobial, Antibacterial, Anti-Gram+, Anti-Gram-, Antifungal |
| 1701 | 5130 | DRAMP21072 | TP1[F4G] (Derived from TP1) | Antimicrobial, Antibacterial, Anti-Gram+, Anti-Gram-, Antifungal |
| 1702 | 5131 | DRAMP21073 | TP1[F4S] (Derived from TP1) | Antimicrobial, Antibacterial, Anti-Gram+, Anti-Gram-, Antifungal |
| 1703 | 5132 | DRAMP21074 | TP1[Y8G] (Derived from TP1) | Antimicrobial, Antibacterial, Anti-Gram+, Anti-Gram-, Antifungal |
| 1704 | 5133 | DRAMP21075 | TP1[I11G] (Derived from TP1) | Antimicrobial, Antibacterial, Anti-Gram+, Anti-Gram-, Antifungal |
| 1705 | 5134 | DRAMP21076 | TP1[F4A, Y8A, I11A] (Derived from TP1) | Antimicrobial, Antibacterial, Anti-Gram+, Anti-Gram-, Antifungal |
| 1706 | 5135 | DRAMP21077 | TP1[-R5, R17G] (Derived from TP1) | Antimicrobial, Antibacterial, Anti-Gram+, Anti-Gram-, Antifungal |
| 1707 | 5136 | DRAMP21078 | TP1[K1A, F4A] (Derived from TP1) | Antimicrobial, Antibacterial, Anti-Gram+, Anti-Gram-, Antifungal |
| 1708 | 5137 | DRAMP21079 | TP1[K1A, Y8A] (Derived from TP1) | Antimicrobial, Antibacterial, Anti-Gram+, Anti-Gram-, Antifungal |
| 1709 | 5138 | DRAMP21080 | TP1[K1A, I11A] (Derived from TP1) | Antimicrobial, Antibacterial, Anti-Gram+, Anti-Gram-, Antifungal |
| 1710 | 5139 | DRAMP21081 | TP1[R9A, R17A] (Derived from TP1) | Antimicrobial, Antibacterial, Anti-Gram+, Anti-Gram-, Antifungal |
| 1711 | 5140 | DRAMP21082 | ccTP 3 (Derived from TP2) | Antimicrobial, Antibacterial, Anti-Gram+, Anti-Gram-, Antifungal |
| 1712 | 5141 | DRAMP21083 | ccTP 5 (Derived from TP2) | Antimicrobial, Antibacterial, Anti-Gram+, Anti-Gram-, Antifungal |
| 1713 | 5142 | DRAMP21084 | ccTP 6 (Derived from TP2) | Antimicrobial, Antibacterial, Anti-Gram+, Anti-Gram-, Antifungal |
| 1714 | 5143 | DRAMP21085 | PRW4 (PR) (Derived from PMAP-36) | Antimicrobial, Antibacterial, Anti-Gram+, Anti-Gram- |
| 1715 | 5144 | DRAMP21086 | PR-FO (Derived from PRW4) | Antimicrobial, Antibacterial, Anti-Gram+, Anti-Gram- |
| 1716 | 5145 | DRAMP21087 | PR-PG (Derived from PRW4) | Antimicrobial, Antibacterial, Anti-Gram+, Anti-Gram- |
| 1717 | 5146 | DRAMP21088 | PR-TR (Derived from PRW4) | Antimicrobial, Antibacterial, Anti-Gram+, Anti-Gram- |
| 1718 | 5147 | DRAMP21089 | C4 (Derived from PRW4) | Antimicrobial, Antibacterial, Anti-Gram+, Anti-Gram- |
| 1719 | 5148 | DRAMP21090 | D4 (Derived from PRW4) | Antimicrobial, Antibacterial, Anti-Gram+, Anti-Gram- |
| 1720 | 5149 | DRAMP21091 | I4 (Derived from PRW4) | Antimicrobial, Antibacterial, Anti-Gram+, Anti-Gram- |
| 1721 | 5150 | DRAMP21092 | P4 (Derived from PRW4) | Antimicrobial, Antibacterial, Anti-Gram+, Anti-Gram- |
| 1722 | 5151 | DRAMP21093 | PRW4-d (Derived from PRW4) | Antimicrobial, Antibacterial, Anti-Gram+, Anti-Gram- |
| 1723 | 5152 | DRAMP21094 | PRW4-R (Derived from PRW4) | Antimicrobial, Antibacterial, Anti-Gram+, Anti-Gram- |
| 1724 | 5153 | DRAMP21095 | IR1 (Derived from PG-1) | Antimicrobial, Antibacterial, Anti-Gram+, Anti-Gram- |
| 1725 | 5154 | DRAMP21096 | IR2 (Derived from PG-1) | Antimicrobial, Antibacterial, Anti-Gram+, Anti-Gram- |
| 1726 | 5155 | DRAMP21097 | FR1 (Derived from PG-1) | Antimicrobial, Antibacterial, Anti-Gram+, Anti-Gram- |
| 1727 | 5156 | DRAMP21098 | FR2 (Derived from PG-1) | Antimicrobial, Antibacterial, Anti-Gram+, Anti-Gram- |
| 1728 | 5157 | DRAMP21099 | WR1 (Derived from PG-1) | Antimicrobial, Antibacterial, Anti-Gram+, Anti-Gram- |
| 1729 | 5158 | DRAMP21100 | WR2 (Derived from PG-1) | Antimicrobial, Antibacterial, Anti-Gram+, Anti-Gram- |
| 1730 | 5159 | DRAMP21101 | PR1 (Derived from PG-1) | Antimicrobial, Antibacterial, Anti-Gram+, Anti-Gram- |
| 1731 | 5160 | DRAMP21102 | PR2 (Derived from PG-1) | Antimicrobial, Antibacterial, Anti-Gram+, Anti-Gram- |
| 1732 | 5161 | DRAMP21165 | HYL-11 (Derived from HYL) | Antimicrobial, Antibacterial, Anti-Gram+, Anti-Gram-, Antifungal |
| 1733 | 5162 | DRAMP21166 | HYL-12 (Derived from HYL) | Antimicrobial, Antibacterial, Anti-Gram+, Anti-Gram-, Antifungal |
| 1734 | 5163 | DRAMP21164 | HYL-10 (Derived from HYL) | Antimicrobial, Antibacterial, Anti-Gram+, Anti-Gram-, Antifungal |
| 1735 | 5164 | DRAMP21158 | HYL-4 (Derived from HYL) | Antimicrobial, Antibacterial, Anti-Gram+, Anti-Gram-, Antifungal |
| 1736 | 5165 | DRAMP21159 | HYL-5 (Derived from HYL) | Antimicrobial, Antibacterial, Anti-Gram+, Anti-Gram-, Antifungal |
| 1737 | 5166 | DRAMP21160 | HYL-6 (Derived from HYL) | Antimicrobial, Antibacterial, Anti-Gram+, Anti-Gram-, Antifungal |
| 1738 | 5167 | DRAMP21161 | HYL-7 (Derived from HYL) | Antimicrobial, Antibacterial, Anti-Gram+, Anti-Gram-, Antifungal |
| 1739 | 5168 | DRAMP21162 | HYL-8 (Derived from HYL) | Antimicrobial, Antibacterial, Anti-Gram+, Anti-Gram-, Antifungal |
| 1740 | 5169 | DRAMP21163 | HYL-9 (Derived from HYL) | Antimicrobial, Antibacterial, Anti-Gram+, Anti-Gram-, Antifungal |
| 1741 | 5170 | DRAMP21157 | HYL-3 (Derived from HYL) | Antimicrobial, Antibacterial, Anti-Gram+, Anti-Gram-, Antifungal |
| 1742 | 5171 | DRAMP21156 | HYL-2 (Derived from HYL) | Antimicrobial, Antibacterial, Anti-Gram+, Anti-Gram-, Antifungal |
| 1743 | 5172 | DRAMP21155 | HYL-1 (Derived from HYL) | Antimicrobial, Antibacterial, Anti-Gram+, Anti-Gram-, Antifungal |
| 1744 | 5173 | DRAMP21154 | HYL (Bee, Insecta, Animals) | Antimicrobial, Antibacterial, Anti-Gram+, Anti-Gram-, Antifungal |
| 1745 | 5174 | DRAMP21153 | KR-12-a8 (Derived from KR-12) | Antimicrobial, Antibacterial, Anti-Gram+, Anti-Gram- |
| 1746 | 5175 | DRAMP21151 | KR-12-a6 (Derived from KR-12) | Antimicrobial, Antibacterial, Anti-Gram+, Anti-Gram- |
| 1747 | 5176 | DRAMP21150 | KR-12-a5 (Derived from KR-12) | Antimicrobial, Antibacterial, Anti-Gram+, Anti-Gram- |
| 1748 | 5177 | DRAMP21152 | KR-12-a7 (Derived from KR-12) | Antimicrobial, Antibacterial, Anti-Gram+, Anti-Gram- |
| 1749 | 5178 | DRAMP21149 | KR-12-a4 (Derived from KR-12) | Antimicrobial, Antibacterial, Anti-Gram+, Anti-Gram- |

| S.no. | PepID | DRAMP_ID | Name of the AMP | Activity |
|---|---|---|---|---|
| 1750 | 5179 | DRAMP21146 | KR-12-a1 (Derived from KR-12) | Antimicrobial, Antibacterial, Anti-Gram+, Anti-Gram- |
| 1751 | 5180 | DRAMP21148 | KR-12-a3 (Derived from KR-12) | Antimicrobial, Antibacterial, Anti-Gram+, Anti-Gram- |
| 1752 | 5181 | DRAMP21147 | KR-12-a2 (Derived from KR-12) | Antimicrobial, Antibacterial, Anti-Gram+, Anti-Gram- |
| 1753 | 5182 | DRAMP21145 | Myxinidin3 (Derived from Myxinidin) | Antimicrobial, Antibacterial, Anti-Gram+, Anti-Gram- |
| 1754 | 5183 | DRAMP21142 | AMP2041 (De novo synthesis) | Antimicrobial, Antibacterial, Anti-Gram+, Anti-Gram- |
| 1755 | 5184 | DRAMP21141 | AMP126 (De novo synthesis) | Antimicrobial, Antibacterial, Anti-Gram+, Anti-Gram- |
| 1756 | 5185 | DRAMP21144 | Myxinidin2 (Derived from Myxinidin) | Antimicrobial, Antibacterial, Anti-Gram+, Anti-Gram- |
| 1757 | 5186 | DRAMP21143 | Myxinidin1 (Derived from Myxinidin) | Antimicrobial, Antibacterial, Anti-Gram+, Anti-Gram- |
| 1758 | 5187 | DRAMP21140 | AMP72 (De novo synthesis) | Antimicrobial, Antibacterial, Anti-Gram+, Anti-Gram- |
| 1759 | 5188 | DRAMP21139 | GNU7 (De novo synthesis) | Antimicrobial, Antibacterial, Anti-Gram+, Anti-Gram-, Antifungal |
| 1760 | 5189 | DRAMP21138 | GNU6 (De novo synthesis) | Antimicrobial, Antibacterial, Anti-Gram+, Anti-Gram-, Antifungal |
| 1761 | 5190 | DRAMP21137 | GNU5 (De novo synthesis) | Antimicrobial, Antibacterial, Anti-Gram+, Anti-Gram-, Antifungal |
| 1762 | 5191 | DRAMP21135 | P7 (Derived from P5) | Antimicrobial, Antibacterial, Anti-Gram+, Anti-Gram- |
| 1763 | 5192 | DRAMP21136 | P8 (Derived from P5) | Antimicrobial, Antibacterial, Anti-Gram+, Anti-Gram- |
| 1764 | 5193 | DRAMP21134 | P6 (Derived from P5) | Antimicrobial, Antibacterial, Anti-Gram+, Anti-Gram- |
| 1765 | 5194 | DRAMP21133 | P5 (Derived from Octa 2) | Antimicrobial, Antibacterial, Anti-Gram+, Anti-Gram- |
| 1766 | 5195 | DRAMP21132 | P4 (Derived from P5) | Antimicrobial, Antibacterial, Anti-Gram+, Anti-Gram- |
| 1767 | 5196 | DRAMP21131 | P3 (Derived from P5) | Antimicrobial, Antibacterial, Anti-Gram+, Anti-Gram- |
| 1768 | 5197 | DRAMP21130 | P2 (Derived from P5) | Antimicrobial, Antibacterial, Anti-Gram+, Anti-Gram- |
| 1769 | 5198 | DRAMP21129 | P1 (Derived from P5) | Antimicrobial, Antibacterial, Anti-Gram+, Anti-Gram- |
| 1770 | 5199 | DRAMP21128 | T9F (Derived from RI16) | Antimicrobial, Antibacterial, Anti-Gram+, Anti-Gram- |
| 1771 | 5200 | DRAMP21127 | T9K (Derived from RI16) | Antimicrobial, Antibacterial, Anti-Gram+, Anti-Gram- |
| 1772 | 5201 | DRAMP21126 | T9I (Derived from RI16) | Antimicrobial, Antibacterial, Anti-Gram+, Anti-Gram- |
| 1773 | 5202 | DRAMP21125 | T9W (Derived from RI16) | Antimicrobial, Antibacterial, Anti-Gram+, Anti-Gram- |
| 1774 | 5203 | DRAMP21124 | RI16 (Derived from PMAP-36) | Antimicrobial, Antibacterial, Anti-Gram+, Anti-Gram- |
| 1775 | 5204 | DRAMP21123 | KR-12-a5 (7-(D)L) (Derived from LL-37) | Antimicrobial, Antibacterial, Anti-Gram+, Anti-Gram- |
| 1776 | 5205 | DRAMP21122 | KR-12-a5 (6-(D)L) (Derived from LL-37) | Antimicrobial, Antibacterial, Anti-Gram+, Anti-Gram- |
| 1777 | 5206 | DRAMP21121 | KR-12-a5 (5-(D)K) (Derived from LL-37) | Antimicrobial, Antibacterial, Anti-Gram+, Anti-Gram- |
| 1778 | 5207 | DRAMP21119 | I11R (Derived from tachyplesin I) | Antimicrobial, Antibacterial, Anti-Gram+, Anti-Gram- |
| 1779 | 5208 | DRAMP21120 | KR-12-a5 (Derived from LL-37) | Antimicrobial, Antibacterial, Anti-Gram+, Anti-Gram- |
| 1780 | 5209 | DRAMP21118 | I11S (Derived from tachyplesin I) | Antimicrobial, Antibacterial, Anti-Gram+, Anti-Gram- |
| 1781 | 5210 | DRAMP21117 | Y8R (Derived from tachyplesin I) | Antimicrobial, Antibacterial, Anti-Gram+, Anti-Gram- |
| 1782 | 5211 | DRAMP21116 | Y8S (Derived from tachyplesin I) | Antimicrobial, Antibacterial, Anti-Gram+, Anti-Gram- |
| 1783 | 5212 | DRAMP21115 | V6R (Derived from tachyplesin I) | Antimicrobial, Antibacterial, Anti-Gram+, Anti-Gram- |
| 1784 | 5213 | DRAMP21114 | V6S (Derived from tachyplesin I) | Antimicrobial, Antibacterial, Anti-Gram+, Anti-Gram- |
| 1785 | 5214 | DRAMP21111 | ASA (Derived from SLZP) | Antimicrobial, Antibacterial, Anti-Gram+, Anti-Gram-, Antifungal |
| 1786 | 5215 | DRAMP21112 | DLSA (Derived from SLZP) | Antimicrobial, Antibacterial, Anti-Gram+, Anti-Gram-, Antifungal |
| 1787 | 5216 | DRAMP21113 | PSA (Derived from SLZP) | Antimicrobial, Antibacterial, Anti-Gram+, Anti-Gram-, Antifungal |
| 1788 | 5217 | DRAMP21103 | L-RW (De novo synthesis) | Antimicrobial, Antibacterial, Anti-Gram+, Anti-Gram- |
| 1789 | 5218 | DRAMP21110 | SLZP (De novo synthesis) | Antimicrobial, Antibacterial, Anti-Gram+, Anti-Gram-, Antifungal |
| 1790 | 5219 | DRAMP21109 | FPA-Bombinin-BO (toads, amphibians, animals) | Antimicrobial, Antibacterial, Anti-Gram+, Anti-Gram-, Antifungal |
| 1791 | 5220 | DRAMP21108 | Feleucin-K3 (Derived from Feleucin-BO1) | Antimicrobial, Antibacterial, Anti-Gram+, Anti-Gram-, Antifungal |
| 1792 | 5221 | DRAMP21104 | Feleucin-2 (toads, amphibians, animals) | Antimicrobial, Antibacterial, Anti-Gram+, Anti-Gram-, Antifungal |
| 1793 | 5224 | DRAMP21107 | Feleucin-BO1 (toads, amphibians, animals) | Antimicrobial, Antibacterial, Anti-Gram+, Anti-Gram-, Antifungal |
| 1794 | 5225 | DRAMP21232 | Ranatuerin-2PLx (R2PLx; Frogs, Amphibians, Animals) | Antimicrobial, Antibacterial, Anti-Gram+, Anti-Gram- |
| 1795 | 5229 | DRAMP21227 | IsCT-P (Derived from IsCT) | Antimicrobial, Antibacterial, Anti-Gram+, Anti-Gram- |
| 1796 | 5230 | DRAMP21228 | IsCT-a (Derived from IsCT-P) | Antimicrobial, Antibacterial, Anti-Gram+, Anti-Gram- |
| 1797 | 5231 | DRAMP21225 | STPk (Derived from STP) | Antimicrobial, Antibacterial, Anti-Gram+, Anti-Gram- |
| 1798 | 5232 | DRAMP21226 | Ink (Derived from IN) | Antimicrobial, Antibacterial, Anti-Gram+, Anti-Gram- |
| 1799 | 5233 | DRAMP21223 | IsCT-p (Derived from IsCT-P) | Antimicrobial, Antibacterial, Anti-Gram+, Anti-Gram- |
| 1800 | 5234 | DRAMP21224 | TPk (Derived from TP) | Antimicrobial, Antibacterial, Anti-Gram+, Anti-Gram- |
| 1801 | 5235 | DRAMP21222 | Control-4D (Derived from IK12-all L) | Antimicrobial, Antibacterial, Anti-Gram+, Anti-Gram-, Antifungal |
| 1802 | 5236 | DRAMP21221 | Control-all D (Derived from IK12-all L) | Antimicrobial, Antibacterial, Anti-Gram+, Anti-Gram-, Antifungal |

| S.no. | PepID | DRAMP_ID | Name of the AMP | Activity |
|---|---|---|---|---|
| 1803 | 5237 | DRAMP21219 | IK12-all D (Derived from IK12-all L) | Antimicrobial, Antibacterial, Anti-Gram+, Anti-Gram-, Antifungal |
| 1804 | 5238 | DRAMP21220 | Control-all L (Derived from IK12-all L) | Antimicrobial, Antibacterial, Anti-Gram+, Anti-Gram-, Antifungal |
| 1805 | 5239 | DRAMP21218 | IK12-all L (De novo synthesis) | Antimicrobial, Antibacterial, Anti-Gram+, Anti-Gram-, Antifungal |
| 1806 | 5240 | DRAMP21217 | IK8-2D (Derived from IK8-all L) | Antimicrobial, Antibacterial, Anti-Gram+, Anti-Gram-, Antifungal |
| 1807 | 5241 | DRAMP21215 | IK4-all D (Derived from IK8-all L) | Antimicrobial, Antibacterial, Anti-Gram+, Anti-Gram-, Antifungal |
| 1808 | 5242 | DRAMP21216 | IK8-4D (Derived from IK8-all L) | Antimicrobial, Antibacterial, Anti-Gram+, Anti-Gram-, Antifungal |
| 1809 | 5243 | DRAMP21213 | IK8-all D (Derived from IK8-all L) | Antimicrobial, Antibacterial, Anti-Gram+, Anti-Gram-, Antifungal |
| 1810 | 5244 | DRAMP21214 | IK6-all D (Derived from IK8-all L) | Antimicrobial, Antibacterial, Anti-Gram+, Anti-Gram-, Antifungal |
| 1811 | 5245 | DRAMP21212 | IK8-all L (De novo synthesis) | Antimicrobial, Antibacterial, Anti-Gram+, Anti-Gram-, Antifungal |
| 1812 | 5255 | DRAMP21202 | HPA3NT3-analog (Derived from HPA3NT3) | Antimicrobial, Antibacterial, Anti-Gram+, Anti-Gram-, Antifungal |
| 1813 | 5256 | DRAMP21201 | Magainin 2a (M2a; Frogs, Amphibians, Animals) | Antimicrobial, Antibacterial, Anti-Gram+, Anti-Gram- |
| 1814 | 5257 | DRAMP21200 | GW-M4 (De novo synthesis) | Antimicrobial, Antibacterial, Anti-Gram+, Anti-Gram- |
| 1815 | 5258 | DRAMP21199 | GW-M3 (De novo synthesis) | Antimicrobial, Antibacterial, Anti-Gram+, Anti-Gram- |
| 1816 | 5259 | DRAMP21198 | GW-M1 (De novo synthesis) | Antimicrobial, Antibacterial, Anti-Gram+, Anti-Gram- |
| 1817 | 5260 | DRAMP21197 | GW-H3 (De novo synthesis) | Antimicrobial, Antibacterial, Anti-Gram+, Anti-Gram- |
| 1818 | 5261 | DRAMP21196 | GW-H1 (De novo synthesis) | Antimicrobial, Antibacterial, Anti-Gram+, Anti-Gram- |
| 1819 | 5262 | DRAMP21195 | GW-A5 (De novo synthesis) | Antimicrobial, Antibacterial, Anti-Gram+, Anti-Gram- |
| 1820 | 5263 | DRAMP21194 | GW-A4 (De novo synthesis) | Antimicrobial, Antibacterial, Anti-Gram+, Anti-Gram- |
| 1821 | 5264 | DRAMP21193 | GW-A2 (De novo synthesis) | Antimicrobial, Antibacterial, Anti-Gram+, Anti-Gram- |
| 1822 | 5265 | DRAMP21192 | GW-A1 (De novo synthesis) | Antimicrobial, Antibacterial, Anti-Gram+, Anti-Gram- |
| 1823 | 5266 | DRAMP21191 | GW-Q6 (De novo synthesis) | Antimicrobial, Antibacterial, Anti-Gram+, Anti-Gram- |
| 1824 | 5267 | DRAMP21190 | GW-Q5 (De novo synthesis) | Antimicrobial, Antibacterial, Anti-Gram+, Anti-Gram- |
| 1825 | 5268 | DRAMP21189 | GW-Q4 (De novo synthesis) | Antimicrobial, Antibacterial, Anti-Gram+, Anti-Gram- |
| 1826 | 5269 | DRAMP21188 | GW-Q3 (De novo synthesis) | Antimicrobial, Antibacterial, Anti-Gram+, Anti-Gram- |
| 1827 | 5270 | DRAMP21187 | WRL4 (Derived from leucocin A) | Antimicrobial, Antibacterial, Anti-Gram+, Anti-Gram-, Antifungal |
| 1828 | 5271 | DRAMP21186 | WRL3 (Derived from leucocin A) | Antimicrobial, Antibacterial, Anti-Gram+, Anti-Gram-, Antifungal |
| 1829 | 5272 | DRAMP21185 | WRL2 (Derived from leucocin A) | Antimicrobial, Antibacterial, Anti-Gram+, Anti-Gram-, Antifungal |
| 1830 | 5273 | DRAMP21184 | WR7 (Derived from leucocin A) | Antimicrobial, Antibacterial, Anti-Gram+, Anti-Gram-, Antifungal |
| 1831 | 5274 | DRAMP21183 | WR5 (Derived from leucocin A) | Antimicrobial, Antibacterial, Anti-Gram+, Anti-Gram-, Antifungal |
| 1832 | 5275 | DRAMP21182 | WR3 (Derived from leucocin A) | Antimicrobial, Antibacterial, Anti-Gram+, Anti-Gram-, Antifungal |
| 1833 | 5276 | DRAMP21181 | WR1 (Derived from leucocin A) | Antimicrobial, Antibacterial, Anti-Gram+, Anti-Gram-, Antifungal |
| 1834 | 5277 | DRAMP21180 | WG18 (Derived from leucocin A) | Antimicrobial, Antibacterial, Anti-Gram+, Anti-Gram-, Antifungal |
| 1835 | 5278 | DRAMP21179 | HYL-26 (Derived from HYL) | Antimicrobial, Antibacterial, Anti-Gram+, Anti-Gram-, Antifungal |
| 1836 | 5279 | DRAMP21178 | HYL-25 (Derived from HYL) | Antimicrobial, Antibacterial, Anti-Gram+, Anti-Gram-, Antifungal |
| 1837 | 5280 | DRAMP21177 | HYL-24 (Derived from HYL) | Antimicrobial, Antibacterial, Anti-Gram+, Anti-Gram-, Antifungal |
| 1838 | 5281 | DRAMP21176 | HYL-23 (Derived from HYL) | Antimicrobial, Antibacterial, Anti-Gram+, Anti-Gram-, Antifungal |
| 1839 | 5282 | DRAMP21175 | HYL-22 (Derived from HYL) | Antimicrobial, Antibacterial, Anti-Gram+, Anti-Gram-, Antifungal |
| 1840 | 5283 | DRAMP21174 | HYL-21 (Derived from HYL) | Antimicrobial, Antibacterial, Anti-Gram+, Anti-Gram-, Antifungal |
| 1841 | 5284 | DRAMP21173 | HYL-20 (Derived from HYL) | Antimicrobial, Antibacterial, Anti-Gram+, Anti-Gram-, Antifungal |
| 1842 | 5285 | DRAMP21172 | HYL-19 (Derived from HYL) | Antimicrobial, Antibacterial, Anti-Gram+, Anti-Gram-, Antifungal |
| 1843 | 5286 | DRAMP21168 | HYL-15 (Derived from HYL) | Antimicrobial, Antibacterial, Anti-Gram+, Anti-Gram-, Antifungal |
| 1844 | 5287 | DRAMP21169 | HYL-16 (Derived from HYL) | Antimicrobial, Antibacterial, Anti-Gram+, Anti-Gram-, Antifungal |
| 1845 | 5288 | DRAMP21170 | HYL-17 (Derived from HYL) | Antimicrobial, Antibacterial, Anti-Gram+, Anti-Gram-, Antifungal |
| 1846 | 5289 | DRAMP21171 | HYL-18 (Derived from HYL) | Antimicrobial, Antibacterial, Anti-Gram+, Anti-Gram-, Antifungal |
| 1847 | 5290 | DRAMP21243 | pardaxin-6 (GE-6) (Derived from pardaxin) | Antimicrobial, Antibacterial, Anti-Gram+, Anti-Gram- |
| 1848 | 5291 | DRAMP21242 | Epinecidin-8 (Derived from Epinecidin) | Antimicrobial, Antibacterial, Anti-Gram+, Anti-Gram- |
| 1849 | 5292 | DRAMP21241 | Epinecidin-1 (Derived from Epinecidin) | Antimicrobial, Antibacterial, Anti-Gram+, Anti-Gram- |
| 1850 | 5293 | DRAMP21240 | FK13-a7 (Derived from FK13) | Antimicrobial, Antibacterial, Anti-Gram+, Anti-Gram- |
| 1851 | 5294 | DRAMP21239 | FK13-a6 (Derived from FK13) | Antimicrobial, Antibacterial, Anti-Gram+, Anti-Gram- |
| 1852 | 5295 | DRAMP21238 | FK13-a5 (Derived from FK13) | Antimicrobial, Antibacterial, Anti-Gram+, Anti-Gram- |
| 1853 | 5296 | DRAMP21237 | FK13-a4 (Derived from FK13) | Antimicrobial, Antibacterial, Anti-Gram+, Anti-Gram- |
| 1854 | 5297 | DRAMP21236 | FK13-a3 (Derived from FK13) | Antimicrobial, Antibacterial, Anti-Gram+, Anti-Gram- |
| 1855 | 5298 | DRAMP21235 | FK13-a2 (Derived from FK13) | Antimicrobial, Antibacterial, Anti-Gram+, Anti-Gram- |

| S.no. | PepID | DRAMP_ID | Name of the AMP | Activity |
|---|---|---|---|---|
| 1856 | 5299 | DRAMP21167 | HYL-13 (Derived from HYL) | Antimicrobial, Antibacterial, Anti-Gram+, Anti-Gram-, Antifungal |
| 1857 | 5300 | DRAMP21234 | FK13-a1 (Derived from FK13) | Antimicrobial, Antibacterial, Anti-Gram+, Anti-Gram- |
| 1858 | 5301 | DRAMP21233 | R2PLx-22 (Derived from R2PLx) | Antimicrobial, Antibacterial, Anti-Gram+, Anti-Gram- |
| 1859 | 5302 | DRAMP21244 | TsAP-S1 (Derived from TsAP-1) | Antimicrobial, Antibacterial, Anti-Gram+, Anti-Gram-, Antifungal |
| 1860 | 5303 | DRAMP21245 | TsAP-S2 (Derived from TsAP-2) | Antimicrobial, Antibacterial, Anti-Gram+, Anti-Gram-, Antifungal |
| 1861 | 5304 | DRAMP21246 | pEM-2 (Derived from the venom of the snake Bothrops asper) | Antimicrobial, Antibacterial, Anti-Gram+, Anti-Gram- |
| 1862 | 5305 | DRAMP21247 | PV (Derived from pEM-2 and MP-VT1) | Antimicrobial, Antibacterial, Anti-Gram+, Anti-Gram- |
| 1863 | 5306 | DRAMP21248 | BVP (Derived from pEM-2 and MP-VT1 and MP-B) | Antimicrobial, Antibacterial, Anti-Gram+, Anti-Gram- |
| 1864 | 5307 | DRAMP21249 | PVP (Derived from MP-B and MP-VT1) | Antimicrobial, Antibacterial, Anti-Gram+, Anti-Gram- |
| 1865 | 5308 | DRAMP21250 | PV3 (Derived from pEM-2 and MP-VT1) | Antimicrobial, Antibacterial, Anti-Gram+, Anti-Gram- |
| 1866 | 5311 | DRAMP21253 | AaeAP1a (Derived from AaeAP1) | Antimicrobial, Antibacterial, Anti-Gram+, Anti-Gram-, Antifungal |
| 1867 | 5312 | DRAMP21254 | AaeAP2a (Derived from AaeAP2) | Antimicrobial, Antibacterial, Anti-Gram+, Anti-Gram-, Antifungal |
| 1868 | 5313 | DRAMP21255 | WL1 (Derived from CP-1) | Antimicrobial, Antibacterial, Anti-Gram+, Anti-Gram- |
| 1869 | 5314 | DRAMP21256 | WL2 (Derived from CP-1) | Antimicrobial, Antibacterial, Anti-Gram+, Anti-Gram- |
| 1870 | 5315 | DRAMP21257 | WL3 (Derived from CP-1) | Antimicrobial, Antibacterial, Anti-Gram+, Anti-Gram- |
| 1871 | 5316 | DRAMP21258 | Cecropin P1 (CP-1) (nematodes, animals) | Antimicrobial, Antibacterial, Anti-Gram+, Anti-Gram- |
| 1872 | 5317 | DRAMP21259 | Scolopendin 1 (Centipedes, Arthropoda, Animals) | Antimicrobial, Antibacterial, Anti-Gram+, Anti-Gram-, Antifungal |
| 1873 | 5318 | DRAMP21260 | KL0A10 (De novo synthesis) | Antimicrobial, Antibacterial, Anti-Gram+, Anti-Gram- |
| 1874 | 5319 | DRAMP21261 | KL4A6 (De novo synthesis) | Antimicrobial, Antibacterial, Anti-Gram+, Anti-Gram- |
| 1875 | 5320 | DRAMP21262 | KL6A4 (De novo synthesis) | Antimicrobial, Antibacterial, Anti-Gram+, Anti-Gram- |
| 1876 | 5321 | DRAMP21263 | KL10A0 (De novo synthesis) | Antimicrobial, Antibacterial, Anti-Gram+, Anti-Gram- |
| 1877 | 5322 | DRAMP21264 | LK (De novo synthesis) | Antimicrobial, Antibacterial, Anti-Gram+, Anti-Gram- |
| 1878 | 5323 | DRAMP21265 | LK-L1A (Derived from LK) | Antimicrobial, Antibacterial, Anti-Gram+, Anti-Gram- |
| 1879 | 5324 | DRAMP21266 | LK-L4A (Derived from LK) | Antimicrobial, Antibacterial, Anti-Gram+, Anti-Gram- |
| 1880 | 5325 | DRAMP21267 | LK-L5A (Derived from LK) | Antimicrobial, Antibacterial, Anti-Gram+, Anti-Gram- |
| 1881 | 5326 | DRAMP21268 | LK-L7A (Derived from LK) | Antimicrobial, Antibacterial, Anti-Gram+, Anti-Gram- |
| 1882 | 5327 | DRAMP21269 | LK-L8A (Derived from LK) | Antimicrobial, Antibacterial, Anti-Gram+, Anti-Gram- |
| 1883 | 5328 | DRAMP21270 | LK-L11A (Derived from LK) | Antimicrobial, Antibacterial, Anti-Gram+, Anti-Gram- |
| 1884 | 5329 | DRAMP21271 | LK-L12A (Derived from LK) | Antimicrobial, Antibacterial, Anti-Gram+, Anti-Gram- |
| 1885 | 5330 | DRAMP21272 | LK-L14A (Derived from LK) | Antimicrobial, Antibacterial, Anti-Gram+, Anti-Gram- |
| 1886 | 5331 | DRAMP21273 | LK-L8G (Derived from LK) | Antimicrobial, Antibacterial, Anti-Gram+, Anti-Gram- |
| 1887 | 5332 | DRAMP21274 | LK-L8S (Derived from LK) | Antimicrobial, Antibacterial, Anti-Gram+, Anti-Gram- |
| 1888 | 5333 | DRAMP21275 | LK-L8P (Derived from LK) | Antimicrobial, Antibacterial, Anti-Gram+, Anti-Gram- |
| 1889 | 5334 | DRAMP21276 | LK-L8N (Derived from LK) | Antimicrobial, Antibacterial, Anti-Gram+, Anti-Gram- |
| 1890 | 5335 | DRAMP21277 | LK-L8Q (Derived from LK) | Antimicrobial, Antibacterial, Anti-Gram+, Anti-Gram- |
| 1891 | 5336 | DRAMP21278 | LK-L8D (Derived from LK) | Antimicrobial, Antibacterial, Anti-Gram+, Anti-Gram- |
| 1892 | 5337 | DRAMP21279 | LK-L8E (Derived from LK) | Antimicrobial, Antibacterial, Anti-Gram+, Anti-Gram- |
| 1893 | 5338 | DRAMP21280 | LK-L8K (Derived from LK) | Antimicrobial, Antibacterial, Anti-Gram+, Anti-Gram- |
| 1894 | 5339 | DRAMP21281 | LK-L8H (Derived from LK) | Antimicrobial, Antibacterial, Anti-Gram+, Anti-Gram- |
| 1895 | 5340 | DRAMP21282 | Lt-F1A (Derived from Lt) | Antimicrobial, Antibacterial, Anti-Gram+, Anti-Gram- |
| 1896 | 5341 | DRAMP21283 | Lt-I4A (Derived from Lt) | Antimicrobial, Antibacterial, Anti-Gram+, Anti-Gram- |
| 1897 | 5342 | DRAMP21284 | Lt-V5A (Derived from Lt) | Antimicrobial, Antibacterial, Anti-Gram+, Anti-Gram- |
| 1898 | 5343 | DRAMP21285 | Lt-I8A (Derived from Lt) | Antimicrobial, Antibacterial, Anti-Gram+, Anti-Gram- |
| 1899 | 5344 | DRAMP21286 | Lt-F11A (Derived from Lt) | Antimicrobial, Antibacterial, Anti-Gram+, Anti-Gram- |
| 1900 | 5345 | DRAMP21287 | Lt-F12A (Derived from Lt) | Antimicrobial, Antibacterial, Anti-Gram+, Anti-Gram- |
| 1901 | 5346 | DRAMP21288 | Lt-I4G (Derived from Lt) | Antimicrobial, Antibacterial, Anti-Gram+, Anti-Gram- |
| 1902 | 5347 | DRAMP21289 | Lt-I4S (Derived from Lt) | Antimicrobial, Antibacterial, Anti-Gram+, Anti-Gram- |
| 1903 | 5348 | DRAMP21290 | Lt-I4N (Derived from Lt) | Antimicrobial, Antibacterial, Anti-Gram+, Anti-Gram- |
| 1904 | 5349 | DRAMP21291 | Lt-I4Q (Derived from Lt) | Antimicrobial, Antibacterial, Anti-Gram+, Anti-Gram- |
| 1905 | 5350 | DRAMP21292 | Lt-I4H (Derived from Lt) | Antimicrobial, Antibacterial, Anti-Gram+, Anti-Gram- |
| 1906 | 5351 | DRAMP21293 | Lt-V5G (Derived from Lt) | Antimicrobial, Antibacterial, Anti-Gram+, Anti-Gram- |
| 1907 | 5352 | DRAMP21294 | Lt-V5S (Derived from Lt) | Antimicrobial, Antibacterial, Anti-Gram+, Anti-Gram- |
| 1908 | 5353 | DRAMP21295 | Lt-V5N (Derived from Lt) | Antimicrobial, Antibacterial, Anti-Gram+, Anti-Gram- |

| S.no. | PepID | DRAMP_ID | Name of the AMP | Activity |
|---|---|---|---|---|
| 1909 | 5354 | DRAMP21296 | Lt-V5Q (Derived from Lt) | Antimicrobial, Antibacterial, Anti-Gram+, Anti-Gram- |
| 1910 | 5355 | DRAMP21297 | Lt-V5H (Derived from Lt) | Antimicrobial, Antibacterial, Anti-Gram+, Anti-Gram- |
| 1911 | 5356 | DRAMP21298 | Lt-F11G (Derived from Lt) | Antimicrobial, Antibacterial, Anti-Gram+, Anti-Gram- |
| 1912 | 5357 | DRAMP21299 | Lt-F11S (Derived from Lt) | Antimicrobial, Antibacterial, Anti-Gram+, Anti-Gram- |
| 1913 | 5358 | DRAMP21300 | Lt-F11N (Derived from Lt) | Antimicrobial, Antibacterial, Anti-Gram+, Anti-Gram- |
| 1914 | 5359 | DRAMP21301 | Lt-F11Q (Derived from Lt) | Antimicrobial, Antibacterial, Anti-Gram+, Anti-Gram- |
| 1915 | 5360 | DRAMP21302 | Lt-F11H (Derived from Lt) | Antimicrobial, Antibacterial, Anti-Gram+, Anti-Gram- |
| 1916 | 5361 | DRAMP21303 | A7-PMAP-23 (Derived from PMAP-23) | Antimicrobial, Antibacterial, Anti-Gram+, Anti-Gram- |
| 1917 | 5362 | DRAMP21304 | A21-PMAP-23 (Derived from PMAP-23) | Antimicrobial, Antibacterial, Anti-Gram+, Anti-Gram- |
| 1918 | 5363 | DRAMP21305 | R8 (De novo synthesis) | Antimicrobial, Antibacterial, Anti-Gram+, Anti-Gram- |
| 1919 | 5364 | DRAMP21306 | TL-1 (Derived from Temporin-1Tl (TL)) | Antimicrobial, Antibacterial, Anti-Gram+, Anti-Gram- |
| 1920 | 5365 | DRAMP21307 | TL-2 (Derived from Temporin-2Tl (TL)) | Antimicrobial, Antibacterial, Anti-Gram+, Anti-Gram- |
| 1921 | 5366 | DRAMP21308 | TL-3 (Derived from Temporin-3Tl (TL)) | Antimicrobial, Antibacterial, Anti-Gram+, Anti-Gram- |
| 1922 | 5367 | DRAMP21309 | TL-4 (Derived from Temporin-4Tl (TL)) | Antimicrobial, Antibacterial, Anti-Gram+, Anti-Gram- |
| 1923 | 5368 | DRAMP21311 | 2W-1 (Derived from PMAP-36) | Antimicrobial, Antibacterial, Anti-Gram+, Anti-Gram- |
| 1924 | 5369 | DRAMP21312 | 2W-2 (Derived from PMAP-36) | Antimicrobial, Antibacterial, Anti-Gram+, Anti-Gram- |
| 1925 | 5370 | DRAMP21313 | 2W-3 (Derived from PMAP-36) | Antimicrobial, Antibacterial, Anti-Gram+, Anti-Gram- |
| 1926 | 5371 | DRAMP21314 | 3W-1 (Derived from PMAP-36) | Antimicrobial, Antibacterial, Anti-Gram+, Anti-Gram- |
| 1927 | 5372 | DRAMP21315 | 3W-2 (Derived from PMAP-36) | Antimicrobial, Antibacterial, Anti-Gram+, Anti-Gram- |
| 1928 | 5373 | DRAMP21316 | 3W-3 (Derived from PMAP-36) | Antimicrobial, Antibacterial, Anti-Gram+, Anti-Gram- |
| 1929 | 5374 | DRAMP21317 | 3W-4 (Derived from PMAP-36) | Antimicrobial, Antibacterial, Anti-Gram+, Anti-Gram- |
| 1930 | 5375 | DRAMP21318 | 3W-5 (Derived from PMAP-36) | Antimicrobial, Antibacterial, Anti-Gram+, Anti-Gram- |
| 1931 | 5376 | DRAMP21319 | 3V (Derived from PMAP-36) | Antimicrobial, Antibacterial, Anti-Gram+, Anti-Gram- |
| 1932 | 5377 | DRAMP21320 | 3L (Derived from PMAP-36) | Antimicrobial, Antibacterial, Anti-Gram+, Anti-Gram- |
| 1933 | 5378 | DRAMP21321 | 4W (Derived from PMAP-36) | Antimicrobial, Antibacterial, Anti-Gram+, Anti-Gram- |
| 1934 | 5379 | DRAMP21322 | RTV (Derived from PMAP-36) | Antimicrobial, Antibacterial, Anti-Gram+, Anti-Gram- |
| 1935 | 5380 | DRAMP21323 | RTI (Derived from PMAP-36) | Antimicrobial, Antibacterial, Anti-Gram+, Anti-Gram- |
| 1936 | 5381 | DRAMP21324 | RTF (Derived from PMAP-36) | Antimicrobial, Antibacterial, Anti-Gram+, Anti-Gram- |
| 1937 | 5382 | DRAMP21325 | RTL (Derived from PMAP-36) | Antimicrobial, Antibacterial, Anti-Gram+, Anti-Gram- |
| 1938 | 5383 | DRAMP21326 | RLR (Derived from PMAP-36) | Antimicrobial, Antibacterial, Anti-Gram+, Anti-Gram- |
| 1939 | 5384 | DRAMP21327 | RVR (Derived from PMAP-36) | Antimicrobial, Antibacterial, Anti-Gram+, Anti-Gram- |
| 1940 | 5385 | DRAMP21328 | RTR (Derived from PMAP-36) | Antimicrobial, Antibacterial, Anti-Gram+, Anti-Gram- |
| 1941 | 5386 | DRAMP21329 | RFR (Derived from PMAP-36) | Antimicrobial, Antibacterial, Anti-Gram+, Anti-Gram- |
| 1942 | 5387 | DRAMP21330 | KVK (Derived from PMAP-36) | Antimicrobial, Antibacterial, Anti-Gram+, Anti-Gram- |
| 1943 | 5388 | DRAMP21331 | KLK (Derived from PMAP-36) | Antimicrobial, Antibacterial, Anti-Gram+, Anti-Gram- |
| 1944 | 5389 | DRAMP21332 | KIK (Derived from PMAP-36) | Antimicrobial, Antibacterial, Anti-Gram+, Anti-Gram- |
| 1945 | 5390 | DRAMP21333 | RVK (Derived from PMAP-36) | Antimicrobial, Antibacterial, Anti-Gram+, Anti-Gram- |
| 1946 | 5391 | DRAMP21334 | Ranatuerin-2Pb (Frogs, amphibians, animals) | Antimicrobial, Antibacterial, Anti-Gram+, Anti-Gram-, Antifungal |
| 1947 | 5392 | DRAMP21335 | RPa (Frogs, amphibians, animals) | Antimicrobial, Antibacterial, Anti-Gram+, Anti-Gram-, Antifungal |
| 1948 | 5393 | DRAMP21336 | RPb (Frogs, amphibians, animals) | Antimicrobial, Antibacterial, Anti-Gram+, Anti-Gram-, Antifungal |
| 1949 | 5394 | DRAMP21337 | BMAP-27 (Bovine, mammals, animals) | Antimicrobial, Antibacterial, Anti-Gram+, Anti-Gram- |
| 1950 | 5395 | DRAMP21338 | [Arg]3-VmCT1-NH2 (Derived from VmCT1) | Antimicrobial, Antibacterial, Anti-Gram+, Anti-Gram-, Antifungal |
| 1951 | 5396 | DRAMP21339 | [Arg]7-VmCT1-NH2 (Derived from VmCT1) | Antimicrobial, Antibacterial, Anti-Gram+, Anti-Gram-, Antifungal |
| 1952 | 5397 | DRAMP21340 | [Arg]11-VmCT1-NH2 (Derived from VmCT1) | Antimicrobial, Antibacterial, Anti-Gram+, Anti-Gram-, Antifungal |
| 1953 | 5398 | DRAMP21341 | [Gly]1-VmCT1-NH2 (Derived from VmCT1) | Antimicrobial, Antibacterial, Anti-Gram+, Anti-Gram-, Antifungal |
| 1954 | 5399 | DRAMP21342 | [Pro]8-VmCT1-NH2 (Derived from VmCT1) | Antimicrobial, Antibacterial, Anti-Gram+, Anti-Gram-, Antifungal |
| 1955 | 5400 | DRAMP21343 | [Leu]9-VmCT1-NH2 (Derived from VmCT1) | Antimicrobial, Antibacterial, Anti-Gram+, Anti-Gram-, Antifungal |
| 1956 | 5401 | DRAMP21344 | [Phe]9-VmCT1-NH2 (Derived from VmCT1) | Antimicrobial, Antibacterial, Anti-Gram+, Anti-Gram-, Antifungal |
| 1957 | 5402 | DRAMP21345 | [Leu]12-VmCT1-NH2 (Derived from VmCT1) | Antimicrobial, Antibacterial, Anti-Gram+, Anti-Gram-, Antifungal |
| 1958 | 5403 | DRAMP21346 | [Tyr]12-VmCT1-NH2 (Derived from VmCT1) | Antimicrobial, Antibacterial, Anti-Gram+, Anti-Gram-, Antifungal |
| 1959 | 5423 | DRAMP21366 | [Lys]1-VmCT1-NH2 (Derived from VmCT1) | Antimicrobial, Antibacterial, Anti-Gram+, Anti-Gram-, Antifungal |
| 1960 | 5425 | DRAMP21368 | [Lys]1[Lys]12-VmCT1-NH2 (Derived from VmCT1) | Antimicrobial, Antibacterial, Anti-Gram+, Anti-Gram-, Antifungal |
| 1961 | 5426 | DRAMP21369 | [Lys]3[Lys]7-VmCT1-NH2 (Derived from VmCT1) | Antimicrobial, Antibacterial, Anti-Gram+, Anti-Gram-, Antifungal |

| S.no. | PepID | DRAMP_ID | Name of the AMP | Activity |
|---|---|---|---|---|
| 1962 | 5427 | DRAMP21370 | [Lys]3[Lys]11-VmCT1-NH2 (Derived from VmCT1) | Antimicrobial, Antibacterial, Anti-Gram+, Anti-Gram-, Antifungal |
| 1963 | 5428 | DRAMP21371 | [Lys]7[Lys]11-VmCT1-NH2 (Derived from VmCT1) | Antimicrobial, Antibacterial, Anti-Gram+, Anti-Gram-, Antifungal |
| 1964 | 5429 | DRAMP21372 | [Lys]3[Lys]7[Lys]11-VmCT1-NH2 (Derived from VmCT1) | Antimicrobial, Antibacterial, Anti-Gram+, Anti-Gram-, Antifungal |
| 1965 | 5447 | DRAMP21390 | K17 (Derived from ATG16) | Antimicrobial, Antibacterial, Anti-Gram+, Anti-Gram-, Antifungal |
| 1966 | 5448 | DRAMP21391 | K18 (Derived from ATG16) | Antimicrobial, Antibacterial, Anti-Gram+, Anti-Gram-, Antifungal |
| 1967 | 5449 | DRAMP21392 | K22 (Derived from ATG16) | Antimicrobial, Antibacterial, Anti-Gram+, Anti-Gram-, Antifungal |
| 1968 | 5450 | DRAMP21393 | K22.2 (Derived from ATG16) | Antimicrobial, Antibacterial, Anti-Gram+, Anti-Gram-, Antifungal |
| 1969 | 5451 | DRAMP21394 | K30 (Derived from ATG16) | Antimicrobial, Antibacterial, Anti-Gram+, Anti-Gram-, Antifungal |
| 1970 | 5452 | DRAMP21395 | K31 (Derived from ATG16) | Antimicrobial, Antibacterial, Anti-Gram+, Anti-Gram-, Antifungal |
| 1971 | 5453 | DRAMP21396 | K33 (Derived from ATG16) | Antimicrobial, Antibacterial, Anti-Gram+, Anti-Gram-, Antifungal |
| 1972 | 5454 | DRAMP21397 | K36 (Derived from ATG16) | Antimicrobial, Antibacterial, Anti-Gram+, Anti-Gram-, Antifungal |
| 1973 | 5455 | DRAMP21398 | K46 (Derived from ATG16) | Antimicrobial, Antibacterial, Anti-Gram+, Anti-Gram-, Antifungal |
| 1974 | 5457 | DRAMP21400 | NBC2253 (De Novo Synthesis) | Antimicrobial, Antibacterial, Anti-Gram+, Anti-Gram- |
| 1975 | 5458 | DRAMP21401 | NBC2254 (De Novo Synthesis) | Antimicrobial, Antibacterial, Anti-Gram+, Anti-Gram- |
| 1976 | 5459 | DRAMP21402 | B1 (Derived from LL-37 and BMAP-27) | Antimicrobial, Antibacterial, Anti-Gram+, Anti-Gram- |
| 1977 | 5460 | DRAMP21403 | peptide 1 (De Novo Synthesis) | Antimicrobial, Antibacterial, Anti-Gram+, Anti-Gram- |
| 1978 | 5461 | DRAMP21404 | peptide 2 (De Novo Synthesis) | Antimicrobial, Antibacterial, Anti-Gram+, Anti-Gram- |
| 1979 | 5462 | DRAMP21405 | LGL13K (De Novo Synthesis) | Antimicrobial, Antibacterial, Anti-Gram+, Anti-Gram- |
| 1980 | 5463 | DRAMP21406 | DGL13K (De Novo Synthesis) | Antimicrobial, Antibacterial, Anti-Gram+, Anti-Gram- |
| 1981 | 5464 | DRAMP21407 | Bac4K (Derived from CAMPs) | Antimicrobial, Antibacterial, Anti-Gram+, Anti-Gram- |
| 1982 | 5465 | DRAMP21408 | Bac3W (Derived from CAMPs) | Antimicrobial, Antibacterial, Anti-Gram+, Anti-Gram- |
| 1983 | 5466 | DRAMP21409 | dBac (Derived from CAMPs) | Antimicrobial, Antibacterial, Anti-Gram+, Anti-Gram- |
| 1984 | 5467 | DRAMP21410 | dBac4K (Derived from CAMPs) | Antimicrobial, Antibacterial, Anti-Gram+, Anti-Gram- |
| 1985 | 5468 | DRAMP21411 | dBac3W (Derived from CAMPs) | Antimicrobial, Antibacterial, Anti-Gram+, Anti-Gram- |
| 1986 | 5469 | DRAMP21412 | dBacK (Derived from CAMPs) | Antimicrobial, Antibacterial, Anti-Gram+, Anti-Gram- |
| 1987 | 5470 | DRAMP21413 | dBacK- (cap) (Derived from CAMPs) | Antimicrobial, Antibacterial, Anti-Gram+, Anti-Gram- |
| 1988 | 5471 | DRAMP21414 | CecB Q53 (Derived from CecB E53) | Antimicrobial, Antibacterial, Anti-Gram+, Anti-Gram-, Antifungal |
| 1989 | 5472 | DRAMP21415 | α4-short (Derived from α4) | Antimicrobial, Antibacterial, Anti-Gram+, Anti-Gram- |
| 1990 | 5473 | DRAMP21416 | WV (De Novo Synthesis) | Antimicrobial, Antibacterial, Anti-Gram+, Anti-Gram- |
| 1991 | 5474 | DRAMP21417 | WI (De Novo Synthesis) | Antimicrobial, Antibacterial, Anti-Gram+, Anti-Gram- |
| 1992 | 5475 | DRAMP21418 | WF (De Novo Synthesis) | Antimicrobial, Antibacterial, Anti-Gram+, Anti-Gram- |
| 1993 | 5476 | DRAMP21419 | WW (De Novo Synthesis) | Antimicrobial, Antibacterial, Anti-Gram+, Anti-Gram- |
| 1994 | 5481 | DRAMP21424 | B1 (De Novo Synthesis) | Antimicrobial, Antibacterial, Anti-Gram+, Anti-Gram- |
| 1995 | 5482 | DRAMP21425 | peptide 2 (Derived from B1) | Antimicrobial, Antibacterial, Anti-Gram+, Anti-Gram- |
| 1996 | 5483 | DRAMP21426 | peptide 3 (Derived from B1) | Antimicrobial, Antibacterial, Anti-Gram+, Anti-Gram- |
| 1997 | 5484 | DRAMP21427 | peptide 4 (Derived from B1) | Antimicrobial, Antibacterial, Anti-Gram+, Anti-Gram- |
| 1998 | 5485 | DRAMP21428 | peptide 5 (Derived from B1) | Antimicrobial, Antibacterial, Anti-Gram+, Anti-Gram- |
| 1999 | 5486 | DRAMP21429 | peptide 6 (Derived from B1) | Antimicrobial, Antibacterial, Anti-Gram+, Anti-Gram- |
| 2000 | 5487 | DRAMP21430 | peptide 7 (Derived from B1) | Antimicrobial, Antibacterial, Anti-Gram+, Anti-Gram- |
| 2001 | 5488 | DRAMP21431 | peptide 8 (Derived from B1) | Antimicrobial, Antibacterial, Anti-Gram+, Anti-Gram- |
| 2002 | 5489 | DRAMP21432 | peptide 9 (Derived from B1) | Antimicrobial, Antibacterial, Anti-Gram+, Anti-Gram- |
| 2003 | 5490 | DRAMP21433 | peptide 10 (Derived from B1) | Antimicrobial, Antibacterial, Anti-Gram+, Anti-Gram- |
| 2004 | 5491 | DRAMP21434 | peptide 11 (Derived from B1) | Antimicrobial, Antibacterial, Anti-Gram+, Anti-Gram- |
| 2005 | 5494 | DRAMP21437 | peptide 14 (Derived from B1) | Antimicrobial, Antibacterial, Anti-Gram+, Anti-Gram- |
| 2006 | 5495 | DRAMP21438 | peptide 15 (Derived from B1) | Antimicrobial, Antibacterial, Anti-Gram+, Anti-Gram- |
| 2007 | 5496 | DRAMP21439 | peptide 16 (Derived from B1) | Antimicrobial, Antibacterial, Anti-Gram+, Anti-Gram- |
| 2008 | 5497 | DRAMP21440 | peptide 17 (Derived from B1) | Antimicrobial, Antibacterial, Anti-Gram+, Anti-Gram- |
| 2009 | 5501 | DRAMP21444 | peptide 21 (Derived from B1) | Antimicrobial, Antibacterial, Anti-Gram+, Anti-Gram- |
| 2010 | 5505 | DRAMP21448 | peptide 25 (Derived from B1) | Antimicrobial, Antibacterial, Anti-Gram+, Anti-Gram- |
| 2011 | 5508 | DRAMP21451 | peptide 28 (Derived from B1) | Antimicrobial, Antibacterial, Anti-Gram+, Anti-Gram- |
| 2012 | 5509 | DRAMP21452 | peptide 29 (Derived from B1) | Antimicrobial, Antibacterial, Anti-Gram+, Anti-Gram- |
| 2013 | 5510 | DRAMP21453 | Hybrid (Derived from Melittin and thanatin) | Antimicrobial, Antibacterial, Anti-Gram+, Anti-Gram- |
| 2014 | 5511 | DRAMP21454 | PLP1 (Insects, animals) | Antimicrobial, Antibacterial, Anti-Gram+, Anti-Gram-, Antifungal |

| S.no. | PepID | DRAMP_ID | Name of the AMP | Activity |
| --- | --- | --- | --- | --- |
| 2015 | 5512 | DRAMP21455 | PLP2 (Insects, animals) | Antimicrobial, Antibacterial, Anti-Gram+, Anti-Gram-, Antifungal |
| 2016 | 5513 | DRAMP21456 | PLP3 (Insects, animals) | Antimicrobial, Antibacterial, Anti-Gram+, Anti-Gram-, Antifungal |
| 2017 | 5514 | DRAMP21457 | PLP4 (Insects, animals) | Antimicrobial, Antibacterial, Anti-Gram+, Anti-Gram-, Antifungal |
| 2018 | 5515 | DRAMP21458 | PLP5 (Insects, animals) | Antimicrobial, Antibacterial, Anti-Gram+, Anti-Gram-, Antifungal |
| 2019 | 5516 | DRAMP21459 | PLP6 (Insects, animals) | Antimicrobial, Antibacterial, Anti-Gram+, Anti-Gram-, Antifungal |
| 2020 | 5517 | DRAMP21460 | PQ (De Novo Synthesis) | Antimicrobial, Antibacterial, Anti-Gram+, Anti-Gram- |
| 2021 | 5518 | DRAMP21461 | PP (De Novo Synthesis) | Antimicrobial, Antibacterial, Anti-Gram+, Anti-Gram- |
| 2022 | 5519 | DRAMP21462 | GG (De Novo Synthesis) | Antimicrobial, Antibacterial, Anti-Gram+, Anti-Gram- |
| 2023 | 5520 | DRAMP21463 | Qa (De Novo Synthesis) | Antimicrobial, Antibacterial, Anti-Gram+, Anti-Gram- |
| 2024 | 5521 | DRAMP21464 | Qna (De Novo Synthesis) | Antimicrobial, Antibacterial, Anti-Gram+, Anti-Gram- |
| 2025 | 5522 | DRAMP21465 | P1-LI-1577 (De Novo Synthesis) | Antimicrobial, Antibacterial, Anti-Gram+, Anti-Gram- |
| 2026 | 5523 | DRAMP21466 | P2-LI-1298 (De Novo Synthesis) | Antimicrobial, Antibacterial, Anti-Gram+, Anti-Gram- |
| 2027 | 5524 | DRAMP21467 | P3-LI-2085 (De Novo Synthesis) | Antimicrobial, Antibacterial, Anti-Gram+, Anti-Gram- |
| 2028 | 5525 | DRAMP21310 | RK12 (Derived from PMAP-36) | Antimicrobial, Antibacterial, Anti-Gram+, Anti-Gram- |
| 2029 | 5528 | DRAMP21494 | MEP-N | Antimicrobial, Antibacterial, Anti-Gram+, Anti-Gram-, Antifungal |
| 2030 | 5529 | DRAMP21579 | Val-nHSLP | Antimicrobial, Antibacterial, Anti-Gram+, Anti-Gram- |
| 2031 | 5538 | DRAMP21616 | DRIM | Antimicrobial, Antibacterial, Anti-Gram+, Anti-Gram- |
| 2032 | 5539 | DRAMP21618 | WWSP | Antimicrobial, Antibacterial, Anti-Gram+, Anti-Gram- |
| 2033 | 5540 | DRAMP21620 | KFGF | Antimicrobial, Antibacterial, Anti-Gram+, Anti-Gram- |
| 2034 | 5541 | DRAMP21622 | MAP-1 | Antimicrobial, Antibacterial, Anti-Gram+, Anti-Gram- |
| 2035 | 5543 | DRAMP21627 | E2EM15W | Antimicrobial, Antibacterial, Anti-Gram+, Anti-Gram- |
