## Supplementary material for "‘Targeting’ the Search: An Upgraded Structural and Functional Repository of Antimicrobial Peptides for Biofilm Studies (B-AMP v2.0) with a Focus on Biofilm Protein Targets": Suppl File 2

**Supplementary Table 2: Distribution of biofilm targets in B-AMP v2.0 based on bacterial class, family, genera or description as per NCBI Taxonomy**

| Serial Number | Bacterial Class, Family, Genus or Description as per NCBI Taxonomy | Count |
|---|---|---|
| 1 | *Pseudomonas* | 315 |
| 2 | *Escherichia* | 259 |
| 3 | *Salmonella* | 216 |
| 4 | *Acinetobacter* | 160 |
| 5 | *Staphylococcus* | 123 |
| 6 | *Stenotrophomonas* | 79 |
| 7 | *Klebsiella* | 70 |
| 8 | *Yersinia* | 69 |
| 9 | *Xanthomonas* | 49 |
| 10 | *Enterobacter* | 48 |
| 11 | *Bacillus* | 46 |
| 12 | *Vibrio* | 31 |
| 13 | *Deltaproteobacteria* | 27 |
| 14 | *Xenorhabdus* | 27 |
| 15 | *Bordetella* | 26 |
| 16 | *Citrobacter* | 26 |
| 17 | *Burkholderia* | 25 |
| 18 | *Shewanella* | 25 |
| 19 | *Candidatus* | 23 |
| 20 | *Gammaproteobacteria* | 22 |
| 21 | *Serratia* | 22 |
| 22 | *Ralstonia* | 21 |
| 23 | *Lysobacter* | 18 |
| 24 | *Pectobacterium* | 18 |
| 25 | *Lentisphaerae* | 17 |
| 26 | *Porphyromona* | 17 |
| 27 | *Achromobacter* | 15 |
| 28 | *Enterobacteriaceae* | 15 |
| 29 | *Streptococcus* | 15 |
| 30 | *Geobacter* | 14 |
| 31 | *Leptospira* | 14 |
| 32 | *Paraburkholderia* | 12 |
| 33 | *Aeromonas* | 11 |
| 34 | *Bradyrhizobium* | 11 |
| 35 | *Porphyromonas* | 11 |
| 36 | *Shigella* | 11 |
| 37 | *Actinobacillus* | 10 |
| 38 | *Chromobacterium* | 10 |
| 39 | *Cronobacter* | 10 |
| 40 | *Cupriavidus* | 10 |
| 41 | *Xanthomonadaceae* | 10 |

| | | |
|---|---|---|
| 42 | *Clostridium* | 9 |
| 43 | *Comamonae* | 9 |
| 44 | *Moraxellaceae* | 9 |
| 45 | *Sulfurimona* | 9 |
| 46 | *Aggregatibacter* | 8 |
| 47 | *Chloroflexus* | 8 |
| 48 | *Deinococcus* | 8 |
| 49 | *Edwardsiella* | 8 |
| 50 | *Lautropia* | 8 |
| 51 | *Synechocystis* | 8 |
| 52 | *Gallionellales* | 7 |
| 53 | *Methylococcaceae* | 7 |
| 54 | *Proteobacteria* | 7 |
| 55 | *Rhodopirellula* | 7 |
| 56 | *Burkholderiales* | 6 |
| 57 | *Comamonadaceae* | 6 |
| 58 | *Desulfobulbaceae* | 6 |
| 59 | *Halomona* | 6 |
| 60 | *Lentisphaeria* | 6 |
| 61 | *Macrococcus* | 6 |
| 62 | *Methylomona* | 6 |
| 63 | *Methylotenera* | 6 |
| 64 | *Nitrospiraceae* | 6 |
| 65 | *Photorhabdus* | 6 |
| 66 | *Priestia* | 6 |
| 67 | *Variovorax* | 6 |
| 68 | *Alcaligenaceae* | 5 |
| 69 | *Carnobacterium* | 5 |
| 70 | *Desulfobacteraceae* | 5 |
| 71 | *Dictyoglomus* | 5 |
| 72 | *Firmicutes* | 5 |
| 73 | *Herbaspirillum* | 5 |
| 74 | *Neisseria* | 5 |
| 75 | *Pelosinus* | 5 |
| 76 | *Pusillimona* | 5 |
| 77 | *Thermotoga* | 5 |
| 78 | *Acidithiobacillales* | 4 |
| 79 | *Alicyclobacillus* | 4 |
| 80 | *Anaeromyxobacter* | 4 |
| 81 | *Companilactobacillus* | 4 |
| 82 | *Hafnia* | 4 |
| 83 | *Janthinobacterium* | 4 |
| 84 | *Listeria* | 4 |
| 85 | *Metakosakonia* | 4 |
| 86 | *Methylocucumis* | 4 |
| 86 | *Oxalobacteraceae* | 4 |
| 87 | *Pseudogulbenkiania* | 4 |

| | | |
|---|---|---|
| 88 | *Pseudomonadales* | 4 |
| 89 | *Sulfuricurvum* | 4 |
| 90 | *Thiotrichales* | 4 |
| 91 | *Trabulsiella* | 4 |
| 92 | *uncultured* | 4 |
| 93 | *Xylella* | 4 |
| 94 | *Alcaligenes* | 3 |
| 95 | *Aquifex* | 3 |
| 96 | *Aquitalea* | 3 |
| 97 | *Betaproteobacteria* | 3 |
| 98 | *blood* | 3 |
| 99 | *Caldanaerobacter* | 3 |
| 100 | *Castellaniella* | 3 |
| 101 | *Chromatiales* | 3 |
| 102 | *Desulfovibrio* | 3 |
| 103 | *Ensifer* | 3 |
| 104 | *Gamma* | 3 |
| 105 | *Geoalkalibacter* | 3 |
| 106 | *Hyphomonaceae* | 3 |
| 107 | *Kangiella* | 3 |
| 108 | *Lachnospiraceae* | 3 |
| 109 | *Lacticaseibacillus* | 3 |
| 110 | *Lactobacillus* | 3 |
| 111 | *Leclercia* | 3 |
| 112 | *Luteibacter* | 3 |
| 113 | *Luteimona* | 3 |
| 114 | *Mammaliicoccus* | 3 |
| 115 | *Novosphingobium* | 3 |
| 116 | *Oceanisphaera* | 3 |
| 117 | *Piscirickettsiaceae* | 3 |
| 118 | *Rhizobium* | 3 |
| 119 | *Rhodocyclales* | 3 |
| 120 | *Sedimenticola* | 3 |
| 121 | *Streptomyces* | 3 |
| 122 | *Syntrophotalea* | 3 |
| 123 | *Tatumella* | 3 |
| 124 | *Thermodesulfovibrio* | 3 |
| 125 | *Thioalkalivibrio* | 3 |
| 126 | *Xanthomonadales* | 3 |
| 127 | *Acidiferrobacteraceae* | 2 |
| 128 | *Acidovorax* | 2 |
| 129 | *Actinobacteria* | 2 |
| 130 | *Agrobacterium* | 2 |
| 131 | *Alphaproteobacteria* | 2 |
| 132 | *Azospirillum* | 2 |
| 133 | *Bacteroides* | 2 |
| 134 | *Bibersteinia* | 2 |

| | | |
|---|---|---|
| 135 | *Collinsella* | 2 |
| 136 | *Dehalococcoidia* | 2 |
| 137 | *Desulfuromonadales* | 2 |
| 138 | *Elusimicrobia* | 2 |
| 139 | *Gordonibacter* | 2 |
| 140 | *Hahella* | 2 |
| 141 | *Helicobacter* | 2 |
| 142 | *Lentilactobacillus* | 2 |
| 143 | *Mannheimia* | 2 |
| 144 | *Marine* | 2 |
| 145 | *Nitrospira* | 2 |
| 146 | *Nitrospirae* | 2 |
| 147 | *Oceanospirillales* | 2 |
| 148 | *Parabacteroides* | 2 |
| 149 | *Pasteurellaceae* | 2 |
| 150 | *Pediococcus* | 2 |
| 151 | *Photobacterium* | 2 |
| 152 | *Planctomycetaceae* | 2 |
| 153 | *Pluralibacter* | 2 |
| 154 | *Polaromona* | 2 |
| 155 | *Rhodocyclaceae* | 2 |
| 156 | *Romboutsia* | 2 |
| 157 | *Rubrivivax* | 2 |
| 158 | *Sphingobium* | 2 |
| 159 | *Sporolactobacillus* | 2 |
| 160 | *Syntrophobacter* | 2 |
| 161 | *Tetragenococcus* | 2 |
| 162 | *Thauera* | 2 |
| 163 | *Tistrella* | 2 |
| 164 | *Acidithiobacillus* | 1 |
| 165 | *Acidobacteria* | 1 |
| 166 | *Advenella* | 1 |
| 167 | *Aerococcus* | 1 |
| 168 | *Arenimona* | 1 |
| 169 | *Bacteroidetes* | 1 |
| 170 | *Brucella* | 1 |
| 171 | *Campylobacter* | 1 |
| 172 | *candidate* | 1 |
| 173 | *Capnocytophaga* | 1 |
| 174 | *Clostridiaceae* | 1 |
| 175 | *Coxiella* | 1 |
| 176 | *Derxia* | 1 |
| 177 | *Desulfitobacterium* | 1 |
| 178 | *Desulfotalea* | 1 |
| 179 | *Desulfuromona* | 1 |
| 180 | *Eggerthella* | 1 |
| 181 | *Eikenella* | 1 |

| | | |
|---|---|---|
| 182 | *Exiguobacterium* | 1 |
| 183 | *Furfurilactobacillus* | 1 |
| 184 | *Gallionella* | 1 |
| 185 | *Gemmataceae* | 1 |
| 186 | *Geobacteraceae* | 1 |
| 187 | *Guyparkeria* | 1 |
| 188 | *Halomonas* | 1 |
| 189 | *Helicobacteraceae* | 1 |
| 190 | *Hydrogenophaga* | 1 |
| 191 | *Koribacter* | 1 |
| 192 | *Kosakonia* | 1 |
| 193 | *Lactococcus* | 1 |
| 194 | *Legionella* | 1 |
| 195 | *Levilactobacillus* | 1 |
| 196 | *Ligilactobacillus* | 1 |
| 197 | *Limosilactobacillus* | 1 |
| 198 | *Magnetospirillum* | 1 |
| 199 | *Marinobacter* | 1 |
| 200 | *Mesorhizobium* | 1 |
| 201 | *Methylomicrobium* | 1 |
| 202 | *Methylophaga* | 1 |
| 203 | *Methylosinus* | 1 |
| 204 | *Mycetohabitans* | 1 |
| 205 | *Mycolicibacterium* | 1 |
| 206 | *Nitrosomonadales* | 1 |
| 207 | *Nitrosospira* | 1 |
| 208 | *Pelagibacterium* | 1 |
| 209 | *Phocaeicola* | 1 |
| 210 | *Proteus* | 1 |
| 211 | *Pseudoalteromona* | 1 |
| 212 | *Rahnella* | 1 |
| 213 | *Rhodanobacter* | 1 |
| 214 | *Rhodopila* | 1 |
| 215 | *Rhodospirillales* | 1 |
| 216 | *Schinkia* | 1 |
| 217 | *Serpentinomona* | 1 |
| 218 | *Serpentinomonas* | 1 |
| 219 | *Shimwellia* | 1 |
| 220 | *Sinorhizobium* | 1 |
| 221 | *Solimona* | 1 |
| 222 | *Spirochaetae* | 1 |
| 218 | *Strain* | 1 |
| 219 | *Syntrophomona* | 1 |
| 220 | *Syntrophus* | 1 |
| 221 | *Thermoanaerobacter* | 1 |
| 222 | *Thermodesulfatator* | 1 |
| 223 | *Tissierellia* | 1 |

| | | |
|---|---|---|
| 224 | *Treponema* | 1 |
| 225 | *Verrucomicrobiales* | 1 |
| 226 | *Weissella* | 1 |
| 227 | *Zetaproteobacteria* | 1 |
| | **Total** | **2502** |
