## Supplementary material for "‘Targeting’ the Search: An Upgraded Structural and Functional Repository of Antimicrobial Peptides for Biofilm Studies (B-AMP v2.0) with a Focus on Biofilm Protein Targets": Suppl File 3

**Supplementary Table 3: Distribution of biofilm targets in B-AMP v2.0 based on bacterial species or description as per NCBI taxonomy**

| Serial Number | Bacterial Species | Count |
|---|---|---|
| 1 | *Escherichia coli* | 249 |
| 2 | *Salmonella enterica* | 118 |
| 3 | *Pseudomonas spp.* | 99 |
| 4 | *Staphylococcus aureus* | 81 |
| 5 | *Acinetobacter baumannii* | 72 |
| 6 | *Klebsiella pneumoniae* | 64 |
| 7 | *Salmonella typhimurium* | 58 |
| 8 | *Pseudomonas fluorescens* | 51 |
| 9 | *Pseudomonas aeruginosa* | 45 |
| 10 | *Stenotrophomonas spp.* | 41 |
| 11 | *Yersinia enterocolitica* | 38 |
| 12 | *Stenotrophomonas maltophilia* | 31 |
| 13 | *Acinetobacter spp.* | 28 |
| 14 | *Deltaproteobacteria bacterium* | 27 |
| 15 | *Bacillus subtilis* | 26 |
| 16 | *Vibrio cholerae* | 25 |
| 17 | *Shewanella oneidensis* | 24 |
| 18 | *Gammaproteobacteria bacterium* | 22 |
| 19 | *Xanthomonas campestris* | 20 |
| 20 | *Citrobacter freundii* | 19 |
| 21 | *Yersinia pestis* | 19 |
| 22 | *Pseudomonas putida* | 18 |
| 23 | *Lentisphaerae bacterium* | 17 |
| 24 | *Porphyromonas gingivalis* | 17 |
| 25 | *Enterobacteriaceae bacterium* | 15 |
| 26 | *Acinetobacter oleivorans* | 14 |
| 27 | *Leptospira interrogans* | 13 |
| 28 | *Ralstonia solanacearum* | 12 |
| 29 | *Serratia marcescens* | 12 |
| 30 | *Bradyrhizobium diazoefficiens* | 11 |
| 31 | *Enterobacter spp.* | 11 |
| 32 | *Porphyromonas gingivalis* | 11 |
| 33 | *Bordetella pertussis* | 10 |
| 34 | *Geobacter sulfurreducens* | 10 |
| 35 | *Staphylococcus epidermidis* | 10 |
| 36 | *Xanthomonadaceae bacterium* | 10 |
| 37 | *Acinetobacter ursingii* | 9 |
| 38 | *Bacillus cereus* | 9 |
| 39 | *Burkholderia multivorans* | 9 |
| 40 | *Comamonae bacterium* | 9 |
| 41 | *Enterobacter hormaechei* | 9 |
| 42 | *Moraxellaceae bacterium* | 9 |

| | | |
|---|---|---|
| 43 | *Xanthomonas oryzae* | 9 |
| 44 | *Yersinia pseudotuberculosis* | 9 |
| 45 | *Bordetella bronchiseptica* | 8 |
| 46 | *Bordetella parapertussis* | 8 |
| 47 | *Chloroflexus aurantiacus* | 8 |
| 48 | *Deinococcus radiodurans* | 8 |
| 49 | *Lautropia spp.* | 8 |
| 50 | *Pseudomonas veronii* | 8 |
| 51 | *Synechocystis spp.* | 8 |
| 52 | *Burkholderia pyrrocinia* | 7 |
| 53 | *Enterobacter cloacae* | 7 |
| 54 | *Enterobacter ludwigii* | 7 |
| 55 | *Gallionellales bacterium* | 7 |
| 56 | *Methylococcaceae bacterium* | 7 |
| 57 | *Proteobacteria bacterium* | 7 |
| 58 | *Pseudomonas brassicacearum* | 7 |
| 59 | *Pseudomonas chlororaphis* | 7 |
| 60 | *Pseudomonas lactis* | 7 |
| 61 | *Rhodopirellula baltica* | 7 |
| 62 | *Achromobacter marplatensis* | 6 |
| 63 | *Acinetobacter calcoaceticus* | 6 |
| 64 | *Acinetobacter pittii* | 6 |
| 65 | *Acinetobacter seifertii* | 6 |
| 66 | *Actinobacillus pleuropneumoniae* | 6 |
| 67 | *Aggregatibacter actinomycetemcomitans* | 6 |
| 68 | *Burkholderiales bacterium* | 6 |
| 69 | *Clostridium botulinum* | 6 |
| 70 | *Comamonadaceae bacterium* | 6 |
| 71 | *Desulfobulbaceae bacterium* | 6 |
| 72 | *Escherichia spp.* | 6 |
| 73 | *Halomona spp.* | 6 |
| 74 | *Lentisphaeria bacterium* | 6 |
| 75 | *Lysobacter capsici* | 6 |
| 76 | *Methylotenera versatilis* | 6 |
| 77 | *Nitrospiraceae bacterium* | 6 |
| 78 | *Priestia megaterium* | 6 |
| 79 | *Staphylococcus capitis* | 6 |
| 80 | *Variovorax spp.* | 6 |
| 81 | *Xenorhabdus nematophila* | 6 |
| 82 | *Xenorhabdus spp.* | 6 |
| 83 | *Acinetobacter venetianus* | 5 |
| 84 | *Alcaligenaceae bacterium* | 5 |
| 85 | *Burkholderia spp.* | 5 |
| 86 | *Carnobacterium divergens* | 5 |
| 87 | *Desulfobacteraceae bacterium* | 5 |
| 88 | *Dictyoglomus turgidum* | 5 |
| 89 | *Edwardsiella tarda* | 5 |

| | | |
|---|---|---|
| 90 | *Firmicutes bacterium* | 5 |
| 91 | *Neisseria meningitidis* | 5 |
| 92 | *Pelosinus spp.* | 5 |
| 93 | *Pseudomonas proteolytica* | 5 |
| 94 | *Pusillimona spp.* | 5 |
| 95 | *Salmonella enteritidis* | 5 |
| 96 | *Salmonella paratyphi* | 5 |
| 97 | *Serratia plymuthica* | 5 |
| 98 | *Shigella flexneri* | 5 |
| 99 | *Staphylococcus haemolyticus* | 5 |
| 100 | *Streptococcus gordonii* | 5 |
| 101 | *Thermotoga maritima* | 5 |
| 102 | *Acidithiobacillales bacterium* | 4 |
| 103 | *Acinetobacter schindleri* | 4 |
| 104 | *Aeromona salmonicida* | 4 |
| 105 | *Alicyclobacillus acidocaldarius* | 4 |
| 106 | *Anaeromyxobacter spp.* | 4 |
| 107 | *Chromobacterium violaceum* | 4 |
| 108 | *Citrobacter pasteurii* | 4 |
| 109 | *Enterobacter agglomerans* | 4 |
| 110 | *Escherichia fergusonii* | 4 |
| 111 | *Geobacter spp.* | 4 |
| 112 | *Hafnia alvei* | 4 |
| 113 | *Janthinobacterium spp.* | 4 |
| 114 | *Klebsiella aerogenes* | 4 |
| 115 | *Listeria monocytogenes* | 4 |
| 116 | *Metakosakonia spp.* | 4 |
| 117 | *Methylocucumis oryzae* | 4 |
| 118 | *Oxalobacteraceae bacterium* | 4 |
| 119 | *Pectobacterium atrosepticum* | 4 |
| 120 | *Pectobacterium parmentieri* | 4 |
| 121 | *Pectobacterium peruviense* | 4 |
| 122 | *Pectobacterium polonicum* | 4 |
| 123 | *Pseudogulbenkiania spp.* | 4 |
| 124 | *Pseudomonadales bacterium* | 4 |
| 125 | *Pseudomonas baetica* | 4 |
| 126 | *Pseudomonas fuscovaginae* | 4 |
| 127 | *Pseudomonas gessardii* | 4 |
| 128 | *Pseudomonas kairouanensis* | 4 |
| 129 | *Pseudomonas moraviensis* | 4 |
| 130 | *Pseudomonas ogarae* | 4 |
| 131 | *Pseudomonas panacis* | 4 |
| 132 | *Pseudomonas protegens* | 4 |
| 133 | *Pseudomonas silesiensis* | 4 |
| 134 | *Pseudomonas synxantha* | 4 |
| 135 | *Pseudomonas trivialis* | 4 |
| 136 | *Pseudomonas viciae* | 4 |

| | | |
|---|---|---|
| 137 | *Shigella spp.* | 4 |
| 138 | *Staphylococcus caprae* | 4 |
| 139 | *Staphylococcus lugdunensis* | 4 |
| 140 | *Stenotrophomonas rhizophila* | 4 |
| 141 | *Streptococcus pneumoniae* | 4 |
| 142 | *Thiotrichales bacterium* | 4 |
| 143 | *Trabulsiella odontotermitis* | 4 |
| 144 | *Xylella fastidiosa* | 4 |
| 145 | *Achromobacter arsenitoxydans* | 3 |
| 146 | *Achromobacter spp.* | 3 |
| 147 | *Achromobacter xylosoxidans* | 3 |
| 148 | *Acinetobacter baylyi* | 3 |
| 149 | *Acinetobacter cumulans* | 3 |
| 150 | *Acinetobacter wuhouensis* | 3 |
| 151 | *Aeromona hydrophila* | 3 |
| 152 | *Alcaligenes spp.* | 3 |
| 153 | *Aquifex aeolicus* | 3 |
| 154 | *Aquitalea magnusonii* | 3 |
| 155 | *Betaproteobacteria bacterium* | 3 |
| 156 | *blood disease* | 3 |
| 157 | *Caldanaerobacter subterraneus* | 3 |
| 158 | *Candidatus Competibacteraceae* | 3 |
| 159 | *Candidatus Desulfofervidus* | 3 |
| 160 | *Candidatus Manganitrophus* | 3 |
| 161 | *Candidatus Muproteobacteria* | 3 |
| 162 | *Candidatus thiomargarita* | 3 |
| 163 | *Castellaniella defragrans* | 3 |
| 164 | *Chromatiales bacterium* | 3 |
| 165 | *Chromobacterium haemolyticum* | 3 |
| 166 | *Chromobacterium spp.* | 3 |
| 167 | *Clostridium spp.* | 3 |
| 168 | *Cronobacter dublinensis* | 3 |
| 169 | *Cronobacter sakazakii* | 3 |
| 170 | *Cupriavidus gilardii* | 3 |
| 171 | *Ensifer adhaerens* | 3 |
| 172 | *Enterobacter asburiae* | 3 |
| 173 | *Enterobacter chengduensis* | 3 |
| 174 | *Enterobacter kobei* | 3 |
| 175 | *Gamma proteobacterium* | 3 |
| 176 | *Geoalkalibacter subterraneus* | 3 |
| 177 | *Herbaspirillum spp.* | 3 |
| 178 | *Hyphomonaceae bacterium* | 3 |
| 179 | *Kangiella koreensis* | 3 |
| 180 | *Lachnospiraceae bacterium* | 3 |
| 181 | *Leclercia spp.* | 3 |
| 182 | *Luteibacter yeojuensis* | 3 |
| 183 | *Luteimona cucumeris* | 3 |

| | | |
|---|---|---|
| 184 | *Lysobacter antibioticus* | 3 |
| 185 | *Lysobacter enzymogenes* | 3 |
| 186 | *Lysobacter gummosus* | 3 |
| 187 | *Methylomona denitrificans* | 3 |
| 188 | *Methylomona methanica* | 3 |
| 189 | *Oceanisphaera profunda* | 3 |
| 190 | *Paraburkholderia phytofirmans* | 3 |
| 191 | *Photorhabdus spp.* | 3 |
| 192 | *Piscirickettsiaceae bacterium* | 3 |
| 193 | *Pseudomonas aestus* | 3 |
| 194 | *Pseudomonas nabeulensis* | 3 |
| 195 | *Pseudomonas orientalis* | 3 |
| 196 | *Pseudomonas prosekii* | 3 |
| 197 | *Pseudomonas yamanorum* | 3 |
| 198 | *Ralstonia pickettii* | 3 |
| 199 | *Ralstonia solanaarum* | 3 |
| 200 | *Ralstonia syzygii* | 3 |
| 201 | *Rhodocyclales bacterium* | 3 |
| 202 | *Salmonella newport* | 3 |
| 203 | *Sedimenticola spp.* | 3 |
| 204 | *Serratia spp.* | 3 |
| 205 | *Staphylococcus saprophyticus* | 3 |
| 206 | *Stenotrophomonas geniculata* | 3 |
| 207 | *Streptomyces coelicolor* | 3 |
| 208 | *Sulfuricurvum kujiense* | 3 |
| 209 | *Sulfurimona denitrificans* | 3 |
| 210 | *Sulfurimona gotlandica* | 3 |
| 211 | *Sulfurimona spp.* | 3 |
| 212 | *Syntrophotalea carbinolica* | 3 |
| 213 | *Tatumella spp.* | 3 |
| 214 | *Thermodesulfovibrio yellowstonii* | 3 |
| 215 | *Thioalkalivibrio sulfidiphilus* | 3 |
| 216 | *Xanthomonadales bacterium* | 3 |
| 217 | *Xanthomonas axonopodis* | 3 |
| 218 | *Xanthomonas citri* | 3 |
| 219 | *Xanthomonas gardneri* | 3 |
| 220 | *Xanthomonas hortorum* | 3 |
| 221 | *Xanthomonas vasicola* | 3 |
| 222 | *Xanthomonas vesicatoria* | 3 |
| 223 | *Xenorhabdus budapestensis* | 3 |
| 224 | *Xenorhabdus ehlersii* | 3 |
| 225 | *Xenorhabdus ishibashii* | 3 |
| 226 | *Xenorhabdus kozodoii* | 3 |
| 227 | *Xenorhabdus stockiae* | 3 |
| 228 | *Yersinia entomophaga* | 3 |
| 229 | *Acidiferrobacteraceae bacterium* | 2 |
| 230 | *Acidovorax citrulli* | 2 |

| | | |
|---|---|---|
| 231 | *Actinobacillus succinogenes* | 2 |
| 232 | *Actinobacillus suis* | 2 |
| 233 | *Actinobacteria bacterium* | 2 |
| 234 | *Aeromona sobria* | 2 |
| 235 | *Aeromona veronii* | 2 |
| 236 | *Aggregatibacter aphrophilus* | 2 |
| 237 | *Alphaproteobacteria bacterium* | 2 |
| 238 | *Azospirillum lipoferum* | 2 |
| 239 | *Bacillus licheniformis* | 2 |
| 240 | *Bibersteinia trehalosi* | 2 |
| 241 | *Burkholderia ambifaria* | 2 |
| 242 | *Burkholderia cenocepacia* | 2 |
| 243 | *Candidatus Accumulibacter* | 2 |
| 244 | *Candidatus Rokubacteria* | 2 |
| 245 | *Collinsella aerofaciens* | 2 |
| 246 | *Companilactobacillus farciminis* | 2 |
| 247 | *Cronobacter malonaticus* | 2 |
| 248 | *Cupriavidus basilensis* | 2 |
| 249 | *Cupriavidus necator* | 2 |
| 250 | *Dehalococcoidia bacterium* | 2 |
| 251 | *Desulfovibrio spp.* | 2 |
| 252 | *Desulfuromonadales bacterium* | 2 |
| 253 | *Elusimicrobia bacterium* | 2 |
| 254 | *Hahella chejuensis* | 2 |
| 255 | *Helicobacter pylori* | 2 |
| 256 | *Klebsiella oxytoca* | 2 |
| 257 | *Lacticaseibacillus fabifermentans* | 2 |
| 258 | *Lysobacter arseniciresistens* | 2 |
| 259 | *Macrococcus caseolyticus* | 2 |
| 260 | *Macrococcus goetzii* | 2 |
| 261 | *Mannheimia succiniciproducens* | 2 |
| 262 | *Marine gamma* | 2 |
| 263 | *Nitrospira bacterium* | 2 |
| 264 | *Nitrospirae bacterium* | 2 |
| 265 | *Novosphingobium nitrogenifigens* | 2 |
| 266 | *Oceanospirillales bacterium* | 2 |
| 267 | *Parabacteroides distasonis* | 2 |
| 268 | *Paraburkholderia azotifigens* | 2 |
| 269 | *Paraburkholderia caribensis* | 2 |
| 270 | *Paraburkholderia hospita* | 2 |
| 271 | *Paraburkholderia spp.* | 2 |
| 272 | *Pasteurellaceae bacterium* | 2 |
| 273 | *Pectobacterium brasiliense* | 2 |
| 274 | *Pediococcus damnosus* | 2 |
| 275 | *Photobacterium profundum* | 2 |
| 276 | *Planctomycetaceae bacterium* | 2 |
| 277 | *Pluralibacter gergoviae* | 2 |

| | | |
|---|---|---|
| 278 | *Pseudomonas furukawaii* | 2 |
| 279 | *Rhodocyclaceae bacterium* | 2 |
| 280 | *Salmonella dublin* | 2 |
| 281 | *Salmonella gallinarum* | 2 |
| 282 | *Salmonella typhi* | 2 |
| 283 | *Sporolactobacillus laevolacticus* | 2 |
| 284 | *Staphylococcus cohnii* | 2 |
| 285 | *Staphylococcus condimenti* | 2 |
| 286 | *Staphylococcus delphini* | 2 |
| 287 | *Streptococcus mutans* | 2 |
| 288 | *Syntrophobacter spp.* | 2 |
| 289 | *Thauera spp.* | 2 |
| 290 | *Tistrella mobilis* | 2 |
| 291 | *uncultured bacterium* | 2 |
| 292 | *uncultured Desulfobacterium* | 2 |
| 293 | *Vibrio scophthalmi* | 2 |
| 294 | *Xanthomonas cucurbitae* | 2 |
| 295 | *Acidithiobacillus ferrivorans* | 1 |
| 296 | *Acidobacteria bacterium* | 1 |
| 297 | *Acinetobacter nosocomialis* | 1 |
| 298 | *Advenella kashmirensis* | 1 |
| 299 | *Aerococcus urinaehominis* | 1 |
| 300 | *Agrobacterium rubi* | 1 |
| 301 | *Agrobacterium salinitolerans* | 1 |
| 302 | *Arenimona spp.* | 1 |
| 303 | *Bacillus atrophaeus* | 1 |
| 304 | *Bacillus glycinifermentans* | 1 |
| 305 | *Bacillus megaterium* | 1 |
| 306 | *Bacillus methanolicus* | 1 |
| 307 | *Bacillus pumilus* | 1 |
| 308 | *Bacillus sonorensis* | 1 |
| 309 | *Bacillus spizizenii* | 1 |
| 310 | *Bacillus spp.* | 1 |
| 311 | *Bacillus stratosphericus* | 1 |
| 312 | *Bacteroides ovatus* | 1 |
| 313 | *Bacteroides uniformis* | 1 |
| 314 | *Bacteroidetes bacterium* | 1 |
| 315 | *Brucella melitensis* | 1 |
| 316 | *Campylobacter jejuni* | 1 |
| 317 | *candidate division* | 1 |
| 318 | *Candidatus Eisenbacteria* | 1 |
| 319 | *Candidatus Fermentibacter* | 1 |
| 320 | *Candidatus Lambdaproteobacteria* | 1 |
| 321 | *Candidatus melainabacteria* | 1 |
| 322 | *Capnocytophaga ochracea* | 1 |
| 323 | *Citrobacter amalonaticus* | 1 |
| 324 | *Citrobacter rodentium* | 1 |

| | | |
|---|---|---|
| 325 | *Citrobacter spp.* | 1 |
| 326 | *Clostridiaceae bacterium* | 1 |
| 327 | *Companilactobacillus mindensis* | 1 |
| 328 | *Companilactobacillus salsicarnum* | 1 |
| 329 | *Coxiella burnetii* | 1 |
| 330 | *Cronobacter condimenti* | 1 |
| 331 | *Cronobacter turicensis* | 1 |
| 332 | *Cupriavidus oxalaticus* | 1 |
| 333 | *Cupriavidus spp.* | 1 |
| 334 | *Cupriavidus taiwanensis* | 1 |
| 335 | *Derxia gummosa* | 1 |
| 336 | *Desulfitobacterium hafniense* | 1 |
| 337 | *Desulfotalea psychrophila* | 1 |
| 338 | *Desulfovibrio magneticus* | 1 |
| 339 | *Desulfuromona spp.* | 1 |
| 340 | *Edwardsiella anguillarum* | 1 |
| 341 | *Edwardsiella ictaluri* | 1 |
| 342 | *Edwardsiella piscicida* | 1 |
| 343 | *Eggerthella spp.* | 1 |
| 344 | *Eikenella corrodens* | 1 |
| 345 | *Enterobacter lignolyticus* | 1 |
| 346 | *Exiguobacterium chiriqhucha* | 1 |
| 347 | *Furfurilactobacillus rossiae* | 1 |
| 348 | *Gallionella spp.* | 1 |
| 349 | *Gemmataceae bacterium* | 1 |
| 350 | *Geobacteraceae bacterium* | 1 |
| 351 | *Gordonibacter pamelaeae* | 1 |
| 352 | *Gordonibacter urolithinfaciens* | 1 |
| 353 | *Guyparkeria spp.* | 1 |
| 354 | *Halomonas spp.* | 1 |
| 355 | *Helicobacteraceae bacterium* | 1 |
| 356 | *Herbaspirillum aquaticum* | 1 |
| 357 | *Herbaspirillum frisingense* | 1 |
| 358 | *Hydrogenophaga spp.* | 1 |
| 359 | *Koribacter versatilis* | 1 |
| 360 | *Kosakonia radicincitans* | 1 |
| 361 | *Lacticaseibacillus rhamnosus* | 1 |
| 362 | *Lactobacillus acidophilus* | 1 |
| 363 | *Lactobacillus johnsonii* | 1 |
| 364 | *Lactobacillus spp.* | 1 |
| 365 | *Lactococcus lactis* | 1 |
| 366 | *Legionella drancourtii* | 1 |
| 367 | *Lentilactobacillus diolivorans* | 1 |
| 368 | *Lentilactobacillus kisonensis* | 1 |
| 369 | *Leptospira licerasiae* | 1 |
| 370 | *Levilactobacillus koreensis* | 1 |
| 371 | *Ligilactobacillus apodemi* | 1 |

| | | |
|---|---|---|
| 372 | *Limosilactobacillus mucosae* | 1 |
| 373 | *Lysobacter ruishenii* | 1 |
| 374 | *Macrococcus bohemicus* | 1 |
| 375 | *Macrococcus epidermidis* | 1 |
| 376 | *Magnetospirillum gryphiswaldense* | 1 |
| 377 | *Mammaliicoccus fleurettii* | 1 |
| 378 | *Mammaliicoccus lentus* | 1 |
| 379 | *Mammaliicoccus sciuri* | 1 |
| 380 | *Marinobacter nauticus* | 1 |
| 381 | *Mesorhizobium huakuii* | 1 |
| 382 | *Methylomicrobium album* | 1 |
| 383 | *Methylophaga spp.* | 1 |
| 384 | *Methylosinus sporium* | 1 |
| 385 | *Mycetohabitans rhizoxinica* | 1 |
| 386 | *Mycolicibacterium smegmatis* | 1 |
| 387 | *Nitrosomonadales bacterium* | 1 |
| 388 | *Nitrosospira multiformis* | 1 |
| 389 | *Novosphingobium barchaimii* | 1 |
| 390 | *Paraburkholderia madseniana* | 1 |
| 391 | *Pelagibacterium spp.* | 1 |
| 392 | *Phocaeicola vulgatus* | 1 |
| 393 | *Photorhabdus khanii* | 1 |
| 394 | *Photorhabdus temperata* | 1 |
| 395 | *Photorhabdus thracensis* | 1 |
| 396 | *Polaromona naphthalenivorans* | 1 |
| 397 | *Polaromona spp.* | 1 |
| 398 | *Proteus mirabilis* | 1 |
| 399 | *Pseudoalteromona translucida* | 1 |
| 400 | *Pseudomonas caspiana* | 1 |
| 401 | *Pseudomonas chengduensis* | 1 |
| 402 | *Pseudomonas mandelii* | 1 |
| 403 | *Rahnella spp.* | 1 |
| 404 | *Rhizobium freirei* | 1 |
| 405 | *Rhizobium radiobacter* | 1 |
| 406 | *Rhizobium spp.* | 1 |
| 407 | *Rhodanobacter spp.* | 1 |
| 408 | *Rhodopila globiformis* | 1 |
| 409 | *Rhodospirillales bacterium* | 1 |
| 410 | *Romboutsia maritimum* | 1 |
| 411 | *Romboutsia weinsteinii* | 1 |
| 412 | *Rubrivivax benzoatilyticus* | 1 |
| 413 | *Rubrivivax gelatinosus* | 1 |
| 414 | *Salmonella abortus-equi* | 1 |
| 415 | *Salmonella agona* | 1 |
| 416 | *Salmonella anatum* | 1 |
| 417 | *Salmonella bongori* | 1 |
| 418 | *Salmonella choleraesuis* | 1 |

| | | |
|---|---|---|
| 419 | *Salmonella derby* | 1 |
| 420 | *Salmonella diarizonae* | 1 |
| 421 | *Salmonella hadar* | 1 |
| 422 | *Salmonella heidelberg* | 1 |
| 423 | *Salmonella houtenae* | 1 |
| 424 | *Salmonella infantis* | 1 |
| 425 | *Salmonella montevideo* | 1 |
| 426 | *Salmonella moscow* | 1 |
| 427 | *Salmonella muenchen* | 1 |
| 428 | *Salmonella oranienberg* | 1 |
| 429 | *Salmonella potsdam* | 1 |
| 430 | *Salmonella schwarzengrund* | 1 |
| 431 | *Salmonella senftenberg* | 1 |
| 432 | *Salmonella thompson* | 1 |
| 433 | *Salmonella typhisuis* | 1 |
| 434 | *Salmonella virchow* | 1 |
| 435 | *Schinkia azotoformans* | 1 |
| 436 | *Serpentinomona mccroryi* | 1 |
| 437 | *Serpentinomonas raichei* | 1 |
| 438 | *Serratia fonticola* | 1 |
| 439 | *Serratia inhibens* | 1 |
| 440 | *Shewanella spp.* | 1 |
| 441 | *Shigella dysenteriae* | 1 |
| 442 | *Shigella sonnei* | 1 |
| 443 | *Shimwellia blattae* | 1 |
| 444 | *Sinorhizobium terangae* | 1 |
| 445 | *Solimona terrae* | 1 |
| 446 | *Sphingobium baderi* | 1 |
| 447 | *Sphingobium spp.* | 1 |
| 448 | *Spirochaetae bacterium* | 1 |
| 449 | *Staphylococcus pseudintermedius* | 1 |
| 450 | *Staphylococcus simiae* | 1 |
| 451 | *Staphylococcus simulans* | 1 |
| 452 | *Staphylococcus spp.* | 1 |
| 453 | *Strain Marseille* | 1 |
| 454 | *Streptococcus anginosus* | 1 |
| 455 | *Streptococcus parasanguinis* | 1 |
| 456 | *Streptococcus pyogenes* | 1 |
| 457 | *Streptococcus salivarius* | 1 |
| 458 | *Sulfuricurvum spp.* | 1 |
| 459 | *Syntrophomona wolfei* | 1 |
| 460 | *Syntrophus aciditrophicus* | 1 |
| 461 | *Tetragenococcus halophilus* | 1 |
| 462 | *Tetragenococcus koreensis* | 1 |
| 463 | *Thermoanaerobacter spp.* | 1 |
| 464 | *Thermodesulfatator atlanticus* | 1 |
| 465 | *Tissierellia bacterium* | 1 |

| | | |
|---|---|---|
| 466 | *Treponema denticola* | 1 |
| 467 | *Verrucomicrobiales bacterium* | 1 |
| 468 | *Vibrio alginolyticus* | 1 |
| 469 | *Vibrio parahaemolyticus* | 1 |
| 470 | *Vibrio spp.* | 1 |
| 471 | *Vibrio vulnificus* | 1 |
| 472 | *Weissella cibaria* | 1 |
| 473 | *Zetaproteobacteria bacterium* | 1 |
| | **Total** | **2502** |
