## Supplementary material for "‘Targeting’ the Search: An Upgraded Structural and Functional Repository of Antimicrobial Peptides for Biofilm Studies (B-AMP v2.0) with a Focus on Biofilm Protein Targets": Suppl File 4

**Supplementary Table 4: PDB functions of the biofilm targets in B-AMP v2.0**

| Serial Number | Classification as per PDB | Number of biofilm targets |
|---|---|---|
| 1. | Cell adhesion | 23 |
| 2. | Transferase | 10 |
| 3. | Hydrolase / Hydrolase inhibitor | 9 |
| 4. | Signaling protein / inhibitor | 9 |
| 5. | Structural protein | 8 |
| 6. | Transcription | 6 |
| 7. | Immune system / RNA | 4 |
| 8. | Protein fibril | 3 |
| 9. | Sugar binding protein/ inhibitor | 3 |
| 10. | Gene regulation | 2 |
| 11. | Protein binding | 2 |
| 12. | Membrane protein | 2 |
| 13. | RNA binding protein / chaperone | 2 |
| 14. | DNA binding protein / toxin | 2 |
| 15. | Transcriptional regulator | 2 |
| 16. | Transport protein | 1 |
| 17. | Fimbrial protein | 1 |
| 18. | Protein transport | 1 |
| 19. | Metal binding protein | 1 |
| 20. | Signaling protein activator | 1 |
| 21. | Peptide binding protein | 1 |
| 22. | Toxin inhibitor | 1 |
| 23. | Calcium binding protein | 1 |
| 24. | Ribosome | 1 |
| 25. | Chaperone / protein transport | 1 |
| 26. | Structural genomics / biosynthetic protein | 1 |
