## Supplementary material for "‘Targeting’ the Search: An Upgraded Structural and Functional Repository of Antimicrobial Peptides for Biofilm Studies (B-AMP v2.0) with a Focus on Biofilm Protein Targets": Suppl File 5

### Supplementary Table 5: List of UniProt functions of biofilm targets in B-AMP v2.0

| Serial Number | Classification as per UniProt | Number of biofilm targets |
|---|---|---|
| 1. | Acetyl glucosaminyl transferase activity | 474 |
| 2. | Hydrolase activity | 442 |
| 3. | Acting on carbon-nitrogen (but not peptide) bonds | 415 |
| 4. | Diguanylate cyclase activity | 188 |
| 5. | DNA binding | 69 |
| 6. | ATP binding | 34 |
| 7. | Phosphorelay sensor kinase activity | 33 |
| 8. | Metal ion binding | 22 |
| 9. | DNA-directed DNA polymerase activity | 18 |
| 10. | GTP binding | 15 |
| 11. | Identical protein binding | 14 |
| 12. | Transcription cis-regulatory region binding | 10 |
| 13. | Acting on glycosyl bonds | 10 |
| 14. | Structural proteins | 9 |
| 15. | DNA-binding Transcription activator activity | 9 |
| 16. | N-acyl homoserine lactone synthase activity | 9 |
| 17. | S-ribosylhomocysteine lyase activity | 9 |
| 18. | Iron ion binding | 9 |
| 19. | RNA binding | 8 |
| 20. | Protein dimerization activity | 7 |
| 21. | AcylTransferase activity | 7 |
| 22. | Endoribonuclease activity | 6 |
| 23. | Phosphorelay response regulator activity | 6 |
| 24. | Zinc ion binding | 6 |
| 25. | Host cell extracellular matrix binding | 6 |
| 26. | Toxin activity | 6 |
| 27. | Sigma factor activity | 5 |
| 28. | Transmembrane transporter activity | 5 |
| 29. | Glycosyl transferase activity | 5 |
| 30. | DNA-binding Transcription repressor activity | 4 |
| 31. | DNA-binding Transcription factor activity | 4 |
| 32. | Catalytic activity | 4 |
| 33. | Sequence-specific DNA binding | 4 |
| 34. | Transferring groups other than amino-acyl groups | 4 |

| | | |
|---|---|---|
| 35. | Phosphopantetheine binding | 3 |
| 36. | Kinase activity | 3 |
| 37. | Calcium ion binding | 3 |
| 38. | tRNA binding | 3 |
| 39. | Oxidoreductase activity | 5 |
| 40. | Phosphoprotein phosphatase activity | 3 |
| 41. | DNA-directed 5'-3' RNA polymerase activity | 3 |
| 42. | Protein histidine kinase activity | 2 |
| 43. | Transferase activity | 2 |
| 44. | Nucleotide binding | 2 |
| 45. | PhosphoTransferase activity | 2 |
| 46. | DNA binding function | 2 |
| 47. | 3-oxoacyl-[acyl-carrier-protein] synthase activity | 2 |
| 48. | Efflux transmembrane transporter activity | 2 |
| 49. | Exodeoxyribonuclease III activity | 2 |
| 50. | Protein-N(PI)-phosphohistidine-sugar phosphotransferase activity | 2 |
| 51. | Cyclic-di-GMP binding | 2 |
| 52. | Phosphoric diester hydrolase activity | 2 |
| 53. | Transcription antitermination factor activity | 2 |
| 54. | Helicase activity | 2 |
| 55. | Endonuclease activity | 2 |
| 56. | Mannose binding | 2 |
| 57. | Hydrolyzing O-glycosyl compounds | 2 |
| 58. | 5'-3' exoribonuclease activity | 2 |
| 59. | mRNA 5'-UTR binding | 2 |
| 60. | Ribosome binding | 2 |
| 61. | Acting on the aldehyde or oxo group of donors | 2 |
| 62. | HexosylTransferase activity | 1 |
| 63. | GTPase activity | 1 |
| 64. | RNA strand annealing activity | 1 |
| 65. | Oxygen binding | 1 |
| 66. | Ribosomal large subunit binding | 1 |
| 67. | RNA helicase activity | 1 |
| 68. | Isomerase activity | 1 |
| 69. | Pheromone activity | 1 |
| 70. | Enzyme binding | 1 |
| 71. | Serine-type peptidase activity | 1 |
| 72. | Protein Kinase activator activity | 1 |

| | | |
|---|---|---|
| 73. | ABC-type bacteriocin transporter activity | 1 |
| 74. | Deacetylase activity | 1 |
| 75. | Pyrophosphatase activity | 1 |
| 76. | Polyphosphate kinase activity | 1 |
| 77. | 5'-bis(diphosphate) 3'-diphosphatase activity | 1 |
| 78. | Toxic substance binding | 1 |
| 79. | Ligase activity | 1 |
| 80. | Producing 3'-phosphomonoesters | 1 |
| 81. | For other substituted phosphate groups | 1 |
| 82. | Double-stranded DNA 3'-5' exodeoxyribonuclease activity | 1 |
| 83. | DNA strand exchange activity | 1 |
| 84. | Protein homodimerization activity | 1 |
| 85. | Transaminase activity | 1 |
| 86. | Cellulose synthase (UDP-forming) activity | 1 |
| 87. | Serine-type endopeptidase activity | 1 |
| 88. | Cholesterol binding | 1 |
| 89. | Cis-regulatory region Sequence-specific DNA binding | 1 |
| 90. | Ribosomal small subunit binding | 1 |
| 91. | tRNA nucleotidylTransferase activity | 1 |
| 92. | 3-hydroxyisobutyryl-CoA Hydrolase activity | 1 |
| 93. | Endoribonuclease inhibitor activity | 1 |
| 94. | ATP hydrolysis activity | 1 |
| 95. | Fimbrial usher porin activity | 1 |
| 96. | Phosphate ion binding | 1 |
| 97. | Transmembrane signaling receptor activity | 1 |
| 98. | dCMP deaminase activity | 1 |
| 99. | Carbohydrate binding | 1 |
| 100. | Exopolyphosphatase activity | 1 |
| 101. | Bacterial-type RNA polymerase core enzyme binding | 1 |
| 102. | Cysteine-type peptidase activity | 1 |
| 103. | Nickel cation binding | 1 |
| 104. | DNA-(apurinic or apyrimidinic site) endonuclease activity | 1 |
| 105. | Flavin adenine dinucleotide binding | 1 |
| 106. | Alpha-L-fucosidase activity | 1 |
| 107. | Porin activity | 1 |
| 108. | Adenosylmethionine decarboxylase activity | 1 |
| 109. | Guanosine-3' | 1 |
| 110. | Beta-ketoacyl-acyl-carrier-protein synthase III activity | 1 |

| | | |
|---|---|---|
| 111. | O-acetyl-ADP-ribose deacetylase activity | 1 |
| 112. | Acting on the CH-CH group of donors | 1 |
| 113. | Alcohol group as acceptor | 1 |
| 114. | Adenosylhomocysteinase activity | 1 |
| 115. | Complement component C3b binding | 1 |
| 116. | Cellulase activity | 1 |
| 117. | Pyridoxal phosphate binding | 1 |
| 118. | ADP-dependent short-chain-acyl-CoA hydrolase activity | 1 |
| 119. | Purine nucleoside binding | 1 |
| 120. | Translation activator activity | 1 |
| 121. | Amino acid binding | 1 |
| 122. | NucleotidylTransferase activity | 1 |
| 123. | rRNA binding | 1 |
| 124. | Carbon monoxide binding | 1 |
| 125. | MethylTransferase activity | 1 |
| 126. | Protein-containing complex binding | 1 |
| 127. | Heme binding | 1 |
| 128. | Autoinducer-2 kinase activity | 1 |
| 129. | Beta-lactamase activity | 1 |
| 130. | Protein-exporting ATPase activity | 1 |
| 131. | 3'-5'-exoribonuclease activity | 1 |
| 132. | Protein kinase activity | 1 |
