## Supplementary material for "‘Targeting’ the Search: An Upgraded Structural and Functional Repository of Antimicrobial Peptides for Biofilm Studies (B-AMP v2.0) with a Focus on Biofilm Protein Targets": Suppl File 7

**Supplementary File 7: List of bacterial species with targets with PDB structures in B-AMP v2.0**

| Serial Number | Bacterial species with PDB structures available | Number of PDB structures |
|---|---|---|
| 1. | *Acinetobacter spp.* | 1 |
| 2. | *Aggregatibacter actinomycetemcomitans* | 1 |
| 3. | *Bacillus subtilis* | 9 |
| 4. | *Bacteroides ovatus* | 1 |
| 5. | *Bacteroides uniformis* | 1 |
| 6. | *Bordetella bronchiseptica* | 1 |
| 7. | *Escherichia coli* | 54 |
| 8. | *Geobacter sulfurreducens* | 1 |
| 9. | *Marinobacter nauticus* | 1 |
| 10. | *Neisseria meningitidis* serogroup B | 1 |
| 11. | *Parabacteroides distasonis* | 2 |
| 12. | *Phocaeicola vulgatus* | 1 |
| 13. | *Porphyromonas gingivalis* | 12 |
| 14. | *Proteus mirabilis* | 1 |
| 15. | *Pseudomonas aeruginosa* | 67 |
| 16. | *Salmonella typhimurium* | 12 |
| 17. | *Shewanella oneidensis* | 1 |
| 18. | *Staphylococcus aureus* | 13 |
| 19. | *Staphylococcus epidermidis* | 1 |
| 20. | *Streptococcus gordonii* | 9 |
| 21. | *Streptococcus mutans* | 2 |
| 22. | *Streptococcus parasanguinis* | 1 |
| 23. | *Streptococcus pneumoniae* serotype 4 | 1 |
| 24. | *Streptococcus pyogenes* serotype M1 | 3 |
| 25. | *Xanthomonas campestris* pv. *campestris* | 1 |
