## Supplementary material for "‘Targeting’ the Search: An Upgraded Structural and Functional Repository of Antimicrobial Peptides for Biofilm Studies (B-AMP v2.0) with a Focus on Biofilm Protein Targets": Suppl File 8

**Supplementary Table 8: *Pseudomonas aeruginosa* biofilm protein targets modeled using RoseTTAfold**

| Serial Number | UniProt Entry | Target IDs | Protein names | Length of protein |
|---|---|---|---|---|
| 1. | Q9HU67 | 533 | 50S ribosomal subunit assembly factor BipA, EC 3.6.5.- (GTP-binding protein BipA) | 605 |
| 2. | K7QUT8 | 843 | Adenosylhomocysteinase, EC 3.3.1.1 (S-adenosyl-L-homocysteine hydrolase, AdoHcyase) | 469 |
| 3. | Q9HT84 | 2248 | Diguanylate cyclase DgcP, EC 2.7.7.65 | 671 |
| 4. | G3XDB0 | 528 | Exodeoxyribonuclease III, EC 3.1.11.2 | 259 |
| 5. | Q9ZN70 | 106 | Exopolyphosphatase, ExopolyPase, EC 3.6.1.11 (Polyphosphate:ADP phosphotransferase, PolyP:ADP phosphotransferase, EC 2.7.4.1) | 506 |
| 6. | Q9HTS0 | 529 | Fimbrial domain-containing protein | 304 |
| 7. | Q9I1Y7 | 520 | Fimbrial subunit CupA1 | 183 |
| 8. | Q9I4X7 | 531 | Fimbrial subunit CupC1 | 205 |
| 9. | Q9HZX6 | 709 | GGDEF domain-containing protein | 525 |
| 10. | Q9I2P4 | 699 | GGDEF domain-containing protein | 401 |
| 11. | Q9I4M8 | 527 | GGDEF domain-containing protein | 398 |
| 12. | Q9HYQ2 | 526 | GGDEF domain-containing protein | 389 |
| 13. | Q9HZ57 | 532 | GGDEF domain-containing protein | 307 |
| 14. | Q9I6W3 | 525 | GGDEF domain-containing protein | 235 |
| 15. | Q9HWU2 | 530 | Probable fimbrial subunit CupB1 | 189 |
| 16. | Q9HUW7 | 711 | Probable two-component response regulator | 542 |
| 17. | G3XD78 | 535 | Regulatory protein RsaL | 80 |
| 18. | Q51373 | 2268 | Response regulator GacA (Global activator) | 214 |

| | | | | |
|---|---|---|---|---|
| 19. | Q9I4N3 | 2272 | Response regulator protein FleR | 473 |
| 20. | A0A0H2ZGR9 | 2279 | RNA polymerase sigma-54 factor | 497 |
| 21. | P49988 | 2212 | RNA polymerase sigma-54 factor | 497 |
| 22. | Q9HWA4 | 107 | Two-component response regulator PprB | 275 |
| 23. | Q9I6K0 | 534 | Uncharacterized protein | 323 |
