## Supplementary material for "‘Targeting’ the Search: An Upgraded Structural and Functional Repository of Antimicrobial Peptides for Biofilm Studies (B-AMP v2.0) with a Focus on Biofilm Protein Targets": Suppl File 9

**Supplementary Table 9: *Staphylococcus aureus* biofilm protein targets modeled using RoseTTAfold**

| Serial Number | UniProt Entry | Target IDs | Protein names | Length of protein |
|---|---|---|---|---|
| 1. | Q9RQQ0 | 2362 | Biofilm operon icaADBC HTH-type negative transcriptional regulator IcaR (Intercellular adhesion protein R) | 186 |
| 2. | Q2FWH5 | 2261 | DEAD-box ATP-dependent RNA helicase CshA, EC 3.6.4.13 | 506 |
| 3. | A0A0H2XKG3 | 2259 | Fibronectin-binding protein B | 940 |
| 4. | Q2G1U6 | 2285 | Global transcriptional regulator Spx | 131 |
| 5. | Q9RQP5 | 404 | Glycerol ester hydrolase | 108 |
| 6. | Q83ZH9 | 402 | IcaA | 124 |
| 7. | Q9RQP7 | 2355 | Poly-beta-1,6-N-acetyl-D-glucosamine N-deacetylase, PNAG N-deacetylase, Poly-beta-1,6-GlcNAc N-deacetylase, EC 3.5.1.- (Biofilm polysaccharide intercellular adhesin deacetylase, Biofilm PIA deacetylase) (Intercellular adhesion protein B) | 290 |
| 8. | Q5HCN1 | 2347 | Poly-beta-1,6-N-acetyl-D-glucosamine synthase, PNAG synthase, Poly-beta-1,6-GlcNAc synthase, EC 2.4.1.- (Biofilm polysaccharide intercellular adhesin synthesis protein IcaA, Biofilm PIA synthesis protein IcaA) (Intercellular adhesion protein A) (N-acetylglucosaminyltransferase IcaA) | 412 |
| 9. | Q6G608 | 2351 | Poly-beta-1,6-N-acetyl-D-glucosamine synthase, PNAG synthase, Poly-beta-1,6-GlcNAc synthase, EC 2.4.1.- (Biofilm polysaccharide intercellular adhesin synthesis protein IcaA, Biofilm PIA synthesis protein IcaA) (Intercellular adhesion protein A) (N-acetylglucosaminyltransferase IcaA) | 412 |

| | | | | |
|---|---|---|---|---|
| 10. | Q6GDD8 | 2350 | Poly-beta-1,6-N-acetyl-D-glucosamine synthase, PNAG synthase, Poly-beta-1,6-GlcNAc synthase, EC 2.4.1.- (Biofilm polysaccharide intercellular adhesin synthesis protein IcaA, Biofilm PIA synthesis protein IcaA) (Intercellular adhesion protein A) (N-acetylglucosaminyltransferase IcaA) | 412 |
| 11. | Q7A351 | 2349 | Poly-beta-1,6-N-acetyl-D-glucosamine synthase, PNAG synthase, Poly-beta-1,6-GlcNAc synthase, EC 2.4.1.- (Biofilm polysaccharide intercellular adhesin synthesis protein IcaA, Biofilm PIA synthesis protein IcaA) (Intercellular adhesion protein A) (N-acetylglucosaminyltransferase IcaA) | 412 |
| 12. | Q8NUI7 | 2352 | Poly-beta-1,6-N-acetyl-D-glucosamine synthase, PNAG synthase, Poly-beta-1,6-GlcNAc synthase, EC 2.4.1.- (Biofilm polysaccharide intercellular adhesin synthesis protein IcaA, Biofilm PIA synthesis protein IcaA) (Intercellular adhesion protein A) (N-acetylglucosaminyltransferase IcaA) | 412 |
| 13. | Q99QX3 | 2348 | Poly-beta-1,6-N-acetyl-D-glucosamine synthase, PNAG synthase, Poly-beta-1,6-GlcNAc synthase, EC 2.4.1.- (Biofilm polysaccharide intercellular adhesin synthesis protein IcaA, Biofilm PIA synthesis protein IcaA) (Intercellular adhesion protein A) (N-acetylglucosaminyltransferase IcaA) | 412 |
| 14. | Q9RQP9 | 2346 | Poly-beta-1,6-N-acetyl-D-glucosamine synthase, PNAG synthase, Poly-beta-1,6-GlcNAc synthase, EC 2.4.1.- (Biofilm polysaccharide intercellular adhesin synthesis protein IcaA, Biofilm PIA synthesis protein IcaA) (Intercellular adhesion protein A) (N-acetylglucosaminyltransferase IcaA) | 412 |
| 15. | A0A0H2XJC1 | 1320 | Poly-beta-1,6-N-acetyl-D-glucosamine synthase, Poly-beta-1,6-GlcNAc synthase, EC 2.4.1.- | 412 |

| | | | | |
|---|---|---|---|---|
| 16. | A0A0H3KAW6 | 462 | Poly-beta-1,6-N-acetyl-D-glucosamine synthase, Poly-beta-1,6-GlcNAc synthase, EC 2.4.1.- | 412 |
| 17. | A0A1D0C161 | 396 | Poly-beta-1,6-N-acetyl-D-glucosamine synthase, Poly-beta-1,6-GlcNAc synthase, EC 2.4.1.- | 410 |
| 18. | A0A2S6DQT8 | 823 | Poly-beta-1,6-N-acetyl-D-glucosamine synthase, Poly-beta-1,6-GlcNAc synthase, EC 2.4.1.- | 412 |
| 19. | A0A6A9GS78 | 403 | Poly-beta-1,6-N-acetyl-D-glucosamine synthase, Poly-beta-1,6-GlcNAc synthase, EC 2.4.1.- | 412 |
| 20. | W8TQA2 | 1786 | Poly-beta-1,6-N-acetyl-D-glucosamine synthase, Poly-beta-1,6-GlcNAc synthase, EC 2.4.1.- | 412 |
| 21. | Q9RQP8 | 2361 | Poly-beta-1,6-N-acetyl-D-glucosamine synthesis protein IcaD, PGA synthesis protein IcaD, Poly-beta-1,6-GlcNAc synthesis protein IcaD (Biofilm polysaccharide intercellular adhesin synthesis protein IcaD, Biofilm PIA synthesis protein IcaD) (Intercellular adhesion protein D) | 101 |
| 22. | Q9RQP6 | 2358 | Probable poly-beta-1,6-N-acetyl-D-glucosamine export protein, PGA export protein, Poly-beta-1,6-GlcNAc export protein (Biofilm polysaccharide intercellular adhesin export protein, Biofilm PIA export protein) (Intercellular adhesion protein C) | 350 |
| 23. | P60612 | 2228 | Sensor histidine kinase/phosphatase LytS, EC 2.7.13.3, EC 3.1.3.- (Autolysin sensor kinase) | 584 |
| 24. | P60613 | 2229 | Sensor histidine kinase/phosphatase LytS, EC 2.7.13.3, EC 3.1.3.- (Autolysin sensor kinase) | 584 |
| 25. | P60614 | 2232 | Sensor histidine kinase/phosphatase LytS, EC 2.7.13.3, EC 3.1.3.- (Autolysin sensor kinase) | 584 |

| | | | | |
|---|---|---|---|---|
| 26. | Q2FK10 | 2224 | Sensor histidine kinase/phosphatase LytS, EC 2.7.13.3, EC 3.1.3.- (Autolysin sensor kinase) | 584 |
| 27. | Q2YV68 | 2226 | Sensor histidine kinase/phosphatase LytS, EC 2.7.13.3, EC 3.1.3.- (Autolysin sensor kinase) | 584 |
| 28. | Q53705 | 2225 | Sensor histidine kinase/phosphatase LytS, EC 2.7.13.3, EC 3.1.3.- (Autolysin sensor kinase) | 584 |
| 29. | Q5HJB6 | 2227 | Sensor histidine kinase/phosphatase LytS, EC 2.7.13.3, EC 3.1.3.- (Autolysin sensor kinase) | 584 |
| 30. | Q6GCL2 | 2231 | Sensor histidine kinase/phosphatase LytS, EC 2.7.13.3, EC 3.1.3.- (Autolysin sensor kinase) | 584 |
| 31. | Q6GK52 | 2230 | Sensor histidine kinase/phosphatase LytS, EC 2.7.13.3, EC 3.1.3.- (Autolysin sensor kinase) | 584 |
| 32. | Q5HFT1 | 2215 | Sensor protein SrrB, EC 2.7.13.3 (Staphylococcal respiratory response protein B) | 583 |
| 33. | Q6G973 | 2301 | Sensor protein SrrB, EC 2.7.13.3 (Staphylococcal respiratory response protein B) | 583 |
| 34. | Q6GGK7 | 2296 | Sensor protein SrrB, EC 2.7.13.3 (Staphylococcal respiratory response protein B) | 583 |
| 35. | Q7A5H7 | 2295 | Sensor protein SrrB, EC 2.7.13.3 (Staphylococcal respiratory response protein B) | 583 |
| 36. | Q8NWF3 | 2294 | Sensor protein SrrB, EC 2.7.13.3 (Staphylococcal respiratory response protein B) | 583 |
| 37. | Q99TZ9 | 2197 | Sensor protein SrrB, EC 2.7.13.3 (Staphylococcal respiratory response protein B) | 583 |
| 38. | Q5HEN9 | 2319 | Sensor protein VraS, EC 2.7.13.3 | 347 |

| | | | | |
|---|---|---|---|---|
| 39. | Q6G849 | 2323 | Sensor protein VraS, EC 2.7.13.3 | 347 |
| 40. | Q6GFH2 | 2299 | Sensor protein VraS, EC 2.7.13.3 | 347 |
| 41. | Q7A0H9 | 2310 | Sensor protein VraS, EC 2.7.13.3 | 347 |
| 42. | Q99SZ7 | 2333 | Sensor protein VraS, EC 2.7.13.3 | 347 |
| 43. | Q9KWK8 | 2320 | Sensor protein VraS, EC 2.7.13.3 | 347 |
| 44. | O86487 | 2284 | Serine-aspartate repeat-containing protein C | 947 |
| 45. | Q2FFR1 | 2202 | Signal transduction protein TRAP (Target of RNAIII-activating protein) | 167 |
| 46. | Q2YU04 | 2203 | Signal transduction protein TRAP (Target of RNAIII-activating protein) | 167 |
| 47. | Q5HEU0 | 2204 | Signal transduction protein TRAP (Target of RNAIII-activating protein) | 167 |
| 48. | Q6G8A1 | 2208 | Signal transduction protein TRAP (Target of RNAIII-activating protein) | 167 |
| 49. | Q6GFM2 | 2207 | Signal transduction protein TRAP (Target of RNAIII-activating protein) | 167 |
| 50. | Q7A2Q4 | 2205 | Signal transduction protein TRAP (Target of RNAIII-activating protein) | 167 |
| 51. | Q7A4W3 | 2206 | Signal transduction protein TRAP (Target of RNAIII-activating protein) | 167 |
| 52. | Q8NVW1 | 2209 | Signal transduction protein TRAP (Target of RNAIII-activating protein) | 167 |
| 53. | P60609 | 2219 | Transcriptional regulatory protein LytR (Sensory transduction protein LytR) | 246 |
| 54. | P60610 | 2220 | Transcriptional regulatory protein LytR (Sensory transduction protein LytR) | 246 |
| 55. | Q2FK09 | 2216 | Transcriptional regulatory protein LytR (Sensory transduction protein LytR) | 246 |
| 56. | Q2YV67 | 2217 | Transcriptional regulatory protein LytR (Sensory transduction protein LytR) | 246 |

| | | | | |
|---|---|---|---|---|
| 57. | Q5HJB5 | 2218 | Transcriptional regulatory protein LytR (Sensory transduction protein LytR) | 246 |
| 58. | Q6GCL1 | 2222 | Transcriptional regulatory protein LytR (Sensory transduction protein LytR) | 246 |
| 59. | Q6GK51 | 2221 | Transcriptional regulatory protein LytR (Sensory transduction protein LytR) | 246 |
| 60. | Q8NYH3 | 2223 | Transcriptional regulatory protein LytR (Sensory transduction protein LytR) | 246 |
| 61. | Q5HFT0 | 2289 | Transcriptional regulatory protein SrrA (Staphylococcal respiratory response protein A) | 241 |
| 62. | Q6G972 | 2198 | Transcriptional regulatory protein SrrA (Staphylococcal respiratory response protein A) | 241 |
| 63. | Q6GGK6 | 2283 | Transcriptional regulatory protein SrrA (Staphylococcal respiratory response protein A) | 241 |
| 64. | Q7A0U4 | 2200 | Transcriptional regulatory protein SrrA (Staphylococcal respiratory response protein A) | 241 |
| 65. | Q7A2R6 | 2196 | Transcriptional regulatory protein SrrA (Staphylococcal respiratory response protein A) | 241 |
| 66. | Q7A5H6 | 2290 | Transcriptional regulatory protein SrrA (Staphylococcal respiratory response protein A) | 241 |
| 67. | Q9L524 | 2199 | Transcriptional regulatory protein SrrA (Staphylococcal respiratory response protein A) | 241 |
